# Limb proportions predict aquatic habits and soft-tissue flippers in extinct amniotes

**DOI:** 10.1101/2025.06.25.661573

**Authors:** Caleb M. Gordon, Lisa S. Freisem, Christopher T. Griffin, Jacques A. Gauthier, Bhart-Anjan S. Bhullar

## Abstract

Gordon et al. use a phylogenetic machine-learning approach incorporating linear and geometric morphometric analyses (including >11,000 original linear measurements) to characterize osteological correlates of interdigital webbing, soft-tissue flippers, and aquatic habits in amniotes. They find that relative hand length can reconstruct soft-tissue phenotypes and aquatic habits in extinct amniotes with >90% accuracy, enabling a more phylogenetically comprehensive investigation of limb evolution among bona fide aquatic and terrestrial species. This investigation reveals distinct morphometric landscapes for limb evolution in mammals and reptiles and clarifies the aquatic habits of multiple extinct groups with historically ambiguous ecologies. In addition, it debuts a versatile supervised machine-learning approach for reconstructing cryptic phenotypes from fossil material.

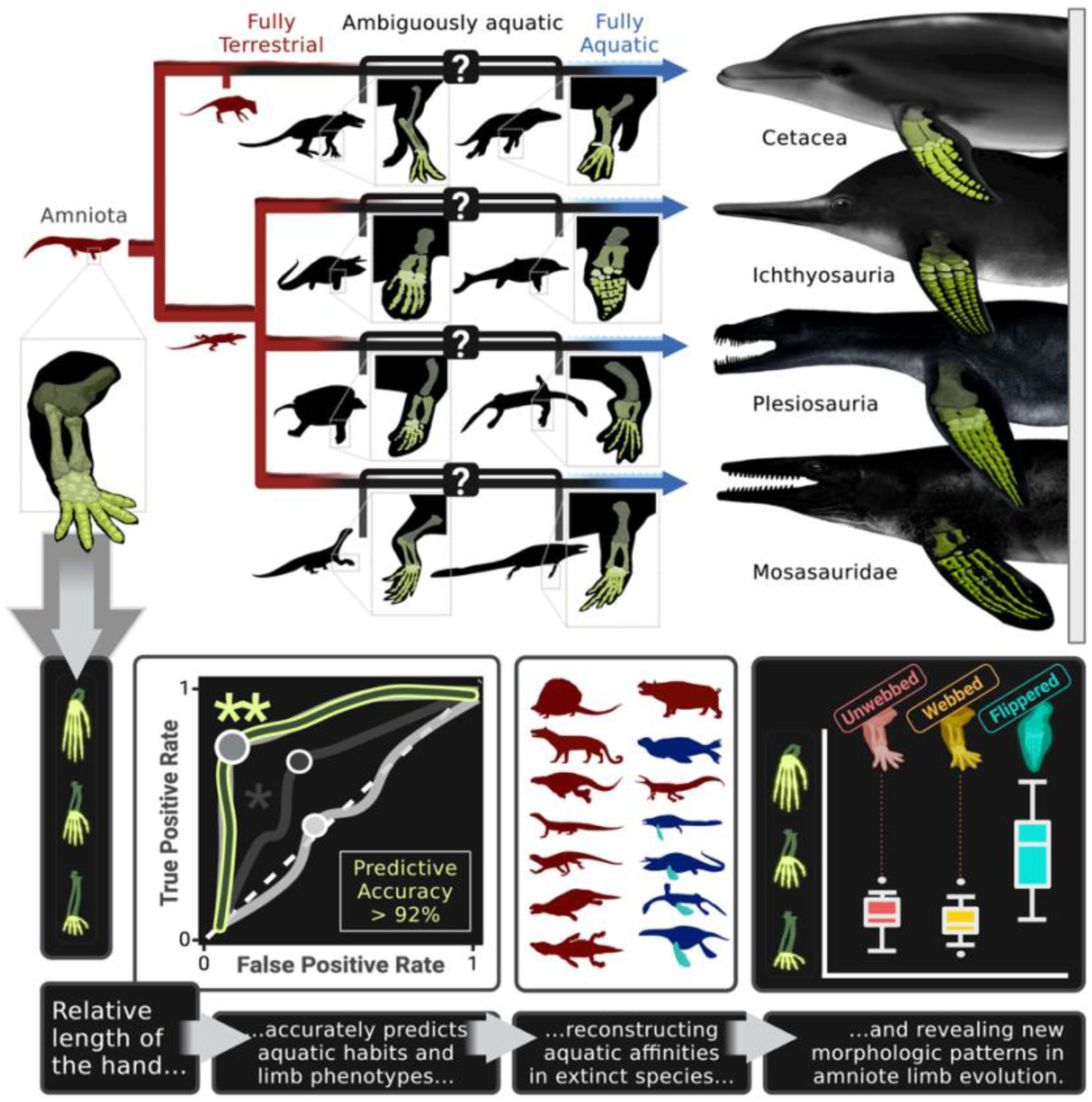

**HIGHLIGHTS:**

- Forelimb proportions reliably predict soft-tissue flippers and highly/fully aquatic habits with >90% accuracy across amniotes.
- Interdigital webbing cannot be predicted from the bones alone using previously suggested correlates.
- All Paleozoic marine reptiles known to date lived at most an amphibious lifestyle, regularly voyaging onto land.
- Phylogenetic Receiver-Operating Characteristic (ROC) analysis effectively reconstructs cryptic phenotypes in extinct species.

## INTRODUCTION

Amniotes descended from terrestrial ancestors^1^, but independently adapted to life in the water dozens of times over the last 300 million years, making them an ideal system for studying ecological and evolutionary constraints across deep time.^2,3,4^ The most aquatically specialized of these groups have limb morphologies that betray a fully marine lifestyle, but the transitional forms on each of their stem lineages have more ambiguous features, making it difficult to determine which fossil species were aquatic and resolve precisely how aquatic they were (Figs. 1, S1). These semi-aquatic taxa often retain plesiomorphic features associated with life on land and thus have some derived features that would (on their own) suggest a more aquatic lifestyle and some incompletely overprinted features that would (on their own) suggest a more terrestrial lifestyle. Conflicting lines of evidence have consequently prompted ongoing controversy about the aquatic affinities of many extinct groups—including spinosaurids,^5,6,7,8^ mesosaurs,^9,10^ and the earliest fossil ancestors of monotremes,^11,12^ cetaceans,^13,14,15^ pinnipeds,^16,17^ turtles,^18,19,20^ ichthyopterygians,^21,22^ sauropterygians,^23,24,25^ and mosasaurs (Text S1–S5).^26,27,28^

**Figure 1.**
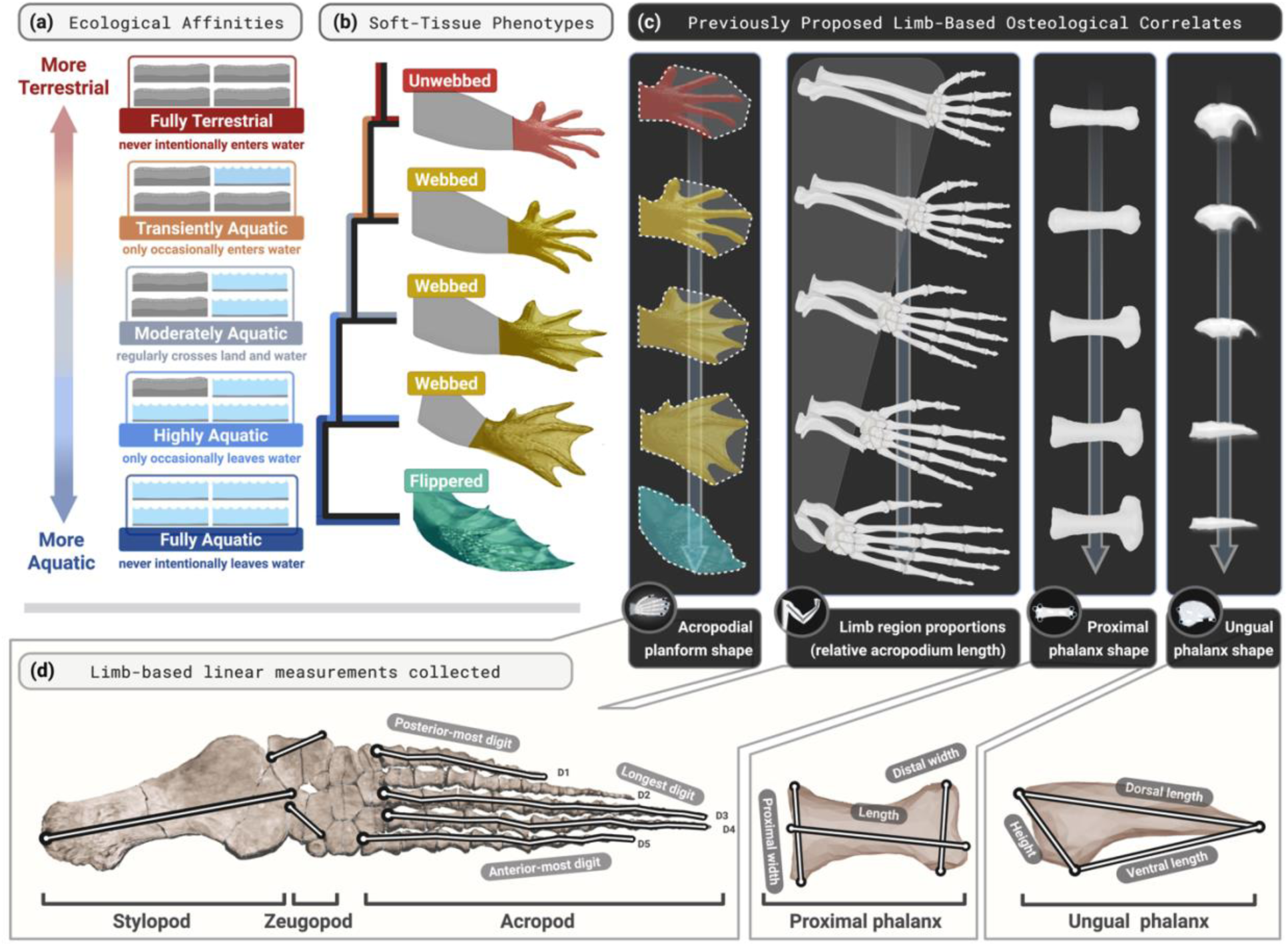
Scoring system and measurements used in this study. (A) The terrestrial-to-aquatic continuum can be captured across amniotes with five ecological bins or stages. (B) As a lineage passes through these stages, its members often evolve soft-tissue limb features (i.e., interdigital webbing and flippers) that help them swim. (C) Several osteological correlates have been proposed for these soft-tissue features or for aquatic habits directly. (D) We collected various linear measurements to test the validity of these proposed correlates. Colors for aquatic affinity bins and soft- tissue phenotypes are consistent across figures.

Predicted soft-tissue limb features, such as interdigital webbing and flippers, may help to discern the semi- or fully aquatic habits of extinct species,^5,9,14,15,29,30,31,32,33^ since these features often reflect the functional ecologies of extant amniotes^34^ (Figs. 1a–b, S2). Here we distinguish *webbed* limbs as having flimsy soft tissues between adjacent phalangeal elements in the hand/foot, and a *flipper* as a continuous soft-tissue surface that encompasses the whole hand/foot and lower arm/leg. Since the direct soft-tissue preservation of webbing and flippers is restricted to exceptional circumstances,^32,35,36,37,38^ researchers have typically identified them using osteological correlates, such as a relatively symmetrical hand or foot planform^9,31,32,39^, a lengthened anterior-most digit,^5,39^ flattened unguals,^5,14,15^ distally widened proximal phalanges,^12^ and differences in the proportions of the three main limb regions (Fig. 1c).^16,28,29,30^ Some of these correlates are corroborated by compelling modern examples,^16,34^ such as the flattened pedal unguals on web-footed birds,^40^ but none have been validated using an extant phylogenetic bracket^41^ that encompasses the extinct taxa of interest (Text S6).

Here, we use a phylogenetic machine-learning approach to assess each of these previously proposed osteological correlates for their validity across amniotes (Figs. S3–S5). Assembling the largest morphometric dataset of amniote limbs to date (11,410 linear measurements on *n* = 747 specimens and three geometric morphometric analyses on *n* = 174 specimens), we compare the proportions and shapes of the major limb regions and the proximal and ungual phalanges among webbed, unwebbed, and flippered species of differing aquatic affinities, which we classify on a five-point scale from “fully terrestrial” to “fully aquatic” (Figs. 1d, S4–S7). For each purported osteological correlate, we train phylogenetic logistic regression models on extant species with known aquatic affinities or soft-tissue phenotypes. We then assess the correlate’s validity with Receiver-Operating Characteristic (ROC) analysis^42^—a machine-learning technique that compares the predictive accuracies of competing models. Finally, we use the best of these correlates to predict the aquatic habits and soft-tissue phenotypes of contentious extinct taxa. In doing so, we shed new light on the evolutionary histories of marine adaptation in seafaring amniotes and offer a new method for reconstructing cryptic phenotypes in extinct species.

## RESULTS

### There are no unambiguous osteological correlates of interdigital webbing

Purported osteological correlates of webbing, such as limb region proportions, acropodial planform shape, proximal phalanx shape, and ungual shape (Fig. 1c), have been used to support contentious habitat predictions for several extinct species.^5,14,15,16,29,31,32^ To test the validity of these correlates, we took linear measurements or (on a subset of specimens; see STAR Methods) performed geometric morphometric analyses on webbed and unwebbed amniote skeletons. We then compared these measurements between webbed and unwebbed taxa, used each feature to iteratively train and test phylogenetic logistic regression models on extant taxa over 100 runs of 3-fold cross-validation, and assessed whether each model and associated metric could predict webbing phenotypes in extant species using ROC analysis. Ultimately, we found no consistent osteological correlates for interdigital webbing across amniotes (Figs. 2, S8–S12). Webbed taxa did not have longer or differently shaped acropodial regions (Figs. S8a–b), flatter unguals (Figs. 2c–d, S9, S10c), or more distally widened proximal phalanges (Figs. S8a–b, S9) than their unwebbed counterparts. In line with these results, limb and phalanx proportions failed to reliably predict webbing (Figs. 2h, S12a). We did identify some more nuanced, clade-specific osteological correlates of webbing (Figs. 2b, S10c, S15; Text S10–S11), but none of these features consistently predicted interdigital webbing across *Amniota*.

**Figure 2.**
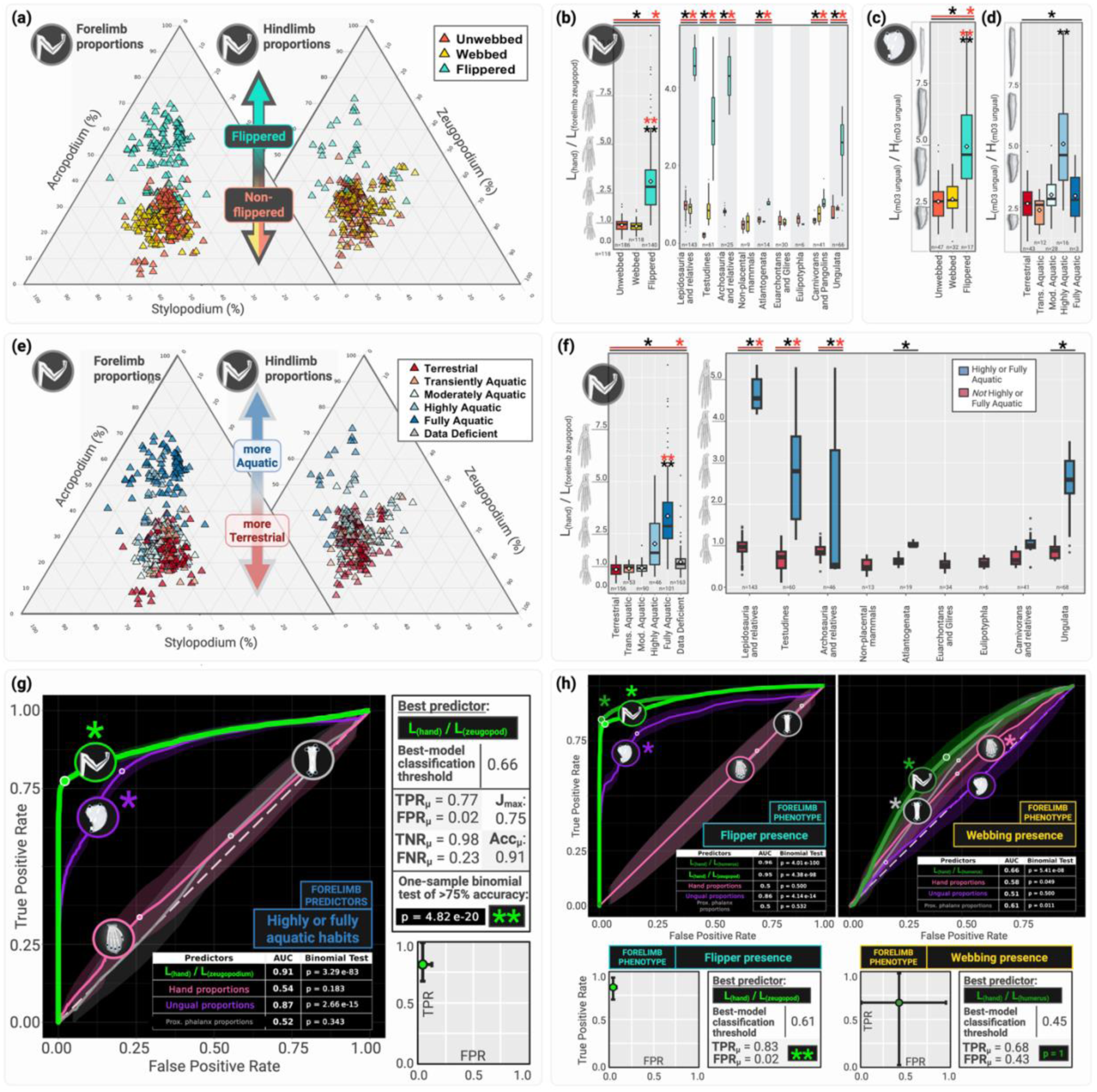
Amniote limb proportions accurately predict flipper phenotypes and aquatic habits. (A) Ternary plots show differences in fore- and hindlimb region proportions among flippered, unwebbed, and webbed species. (B) Box plots show that relative hand length is significantly higher and more variable for flippered species both across amniotes and within each individual clade sampled. (C–D) Flippered and highly aquatic species also have significantly longer, flatter unguals. (E) Ternary plots show differences in fore- and hindlimb region proportions among amniotes of differing aquatic affinities. (F) Box plots show that highly or fully aquatic species have significantly longer hand-zeugopodium length ratios than less aquatic species, both across amniotes and within each individual clade sampled. (G–F) Consensus Receiver- Operating Characteristic (ROC) curves compare the predictive accuracies of forelimb proportions (green), and the shapes of the acropodial planform (pink), ungual phalanx (purple), and proximal phalanx (gray) for (G) highly/fully aquatic habits and (H) soft-tissue limb phenotypes in amniotes for which these features were known *a priori*. Forelimb region proportions (hand-zeugopodium length ratios) are recovered as the most accurate predictors in all cases. “Best predictor” insets show the characteristics of the best predictive model for each phenotype, noting its optimal threshold probability for classifying binary phenotypes, and its mean and 95% confidence interval values for TPR and FPR. In (B–F), asterisks denote significant results (*p* < 0.05) of phylogenetic ANOVAs (black) and Levene’s tests (red)—single asterisks for omnibus tests and double asterisks for pairwise post-hoc tests (here indicating difference from the “terrestrial” group). In (G– H), double asterisks indicate that the best model has a success rate significantly higher than 75% (one- tailed binomial test, *p* < 0.05). Abbreviations: Acc., accuracy; AUC, area under the ROC curve; D3, digit 3; FNR, false negative rate; FPR, false positive rate; H, height; Jmax, maximum Youden’s J value; L, length; m, manual; n, sample size; prox., proximal; TNR, true negative rate; TPR, true positive rate. Additional morphometric and ROC analysis results are provided in the Supplementary Material (Document S1).

### Amniote limb proportions accurately predict flipper phenotypes and highly or fully aquatic habits

In contrast to webbing, flippers were reliably predicted across amniotes (Figs. 2, S8–S12). Flippered species have significantly longer hand regions and flatter unguals than non-flippered species, as well as significantly more variable limb region proportions, ungual dimensions, and hand planform shapes (Figs. 2a–c, S8a–b, S12a). These morphometric differences hold independently for each clade with flippered members (Fig. 2b).

Consistent with the observed morphometric trends, limb region and ungual proportions reliably predicted flipper phenotypes across amniotes (Figs. 2h, S12a). Relative hand length achieved the highest predictive accuracies (Fig. 2h) and remained a significant predictor in nearly all individual clades (Fig. S15). The predictive accuracy of relative hand length was maximized at a binary classification threshold of 0.61 (such that taxa with predicted probabilities significantly above 0.61 would be classified as flippered), yielding a predictive model with true and false positive rates of 0.83 and 0.02, respectively (Fig. 2h). Taken together, these results support the use of forelimb region proportions for predicting flipper presence across amniotes.

Limb proportions also scaled consistently with aquatic affinity across amniotes (Figs. 2e–f, S8c, S12–S13). Highly and fully aquatic species had consistently higher and more variable relative hand and foot lengths and distinct hand planform dimensions relative to their moderately aquatic, transiently aquatic, and terrestrial counterparts (Figs. 2e–f, S8c). To score aquatic affinity as a binary variable suitable for binomial logistic regression models, we recombined our moderately, highly, and fully aquatic species bins in various ways to create binary “more aquatic” and “less aquatic” categories (Fig. S6) and compared these to alternative aquatic affinity classification systems (Fig. S7). In each case, regardless of how we recombined these bins, relative acropodium (hand or foot) length was higher for all aquatic affinity bins that included fully aquatic taxa, both across amniotes (Fig. S6) and within each individual clade with aquatic and terrestrial members (Figs. 2f, S13a–b).

Accordingly, forelimb proportions were significant predictors of both “aquatic or semi-aquatic” habits (Fig. S13c–d) and “highly or fully aquatic” habits (Fig. 2g) both across amniotes and within most individually sampled clades with highly/fully aquatic members (Fig. S14). The predictive accuracy of relative hand length for the latter was maximized at a binary classification threshold of 0.66 (such that all taxa with predicted probabilities significantly above 0.66 would be classified as highly/fully aquatic), yielding a predictive model with a mean false positive rate of 0.02 (Fig. 4g). Hindlimb proportions were also significant predictors of highly/fully aquatic habits (Fig. S12b), but less accurate than forelimb proportions. We therefore selected relative hand (rather than foot) length as the single best-performing osteological correlate with which to predict highly/fully aquatic habits in extinct taxa.

### Reptiles show lineage-specific patterns of aquatic adaptation

By feeding relative hand lengths gathered for extinct taxa into our best-performing logistic regression models (see STAR Methods), we computed the mean probability (across 10,000 simulated iterations) that these extinct species were flippered and highly/fully aquatic. We then used a one-tailed t-test to determine whether this predicted probability was significantly higher than the optimal classification threshold identified by ROC analysis. This process yielded binary phenotype classifications (“highly/fully aquatic” or not, “flippered” or not) for all tips on the tree with formerly ambiguous phenotypes (Figs. 3, S16), which we propagated down the tree using ancestral state reconstructions.

**Figure 3.**
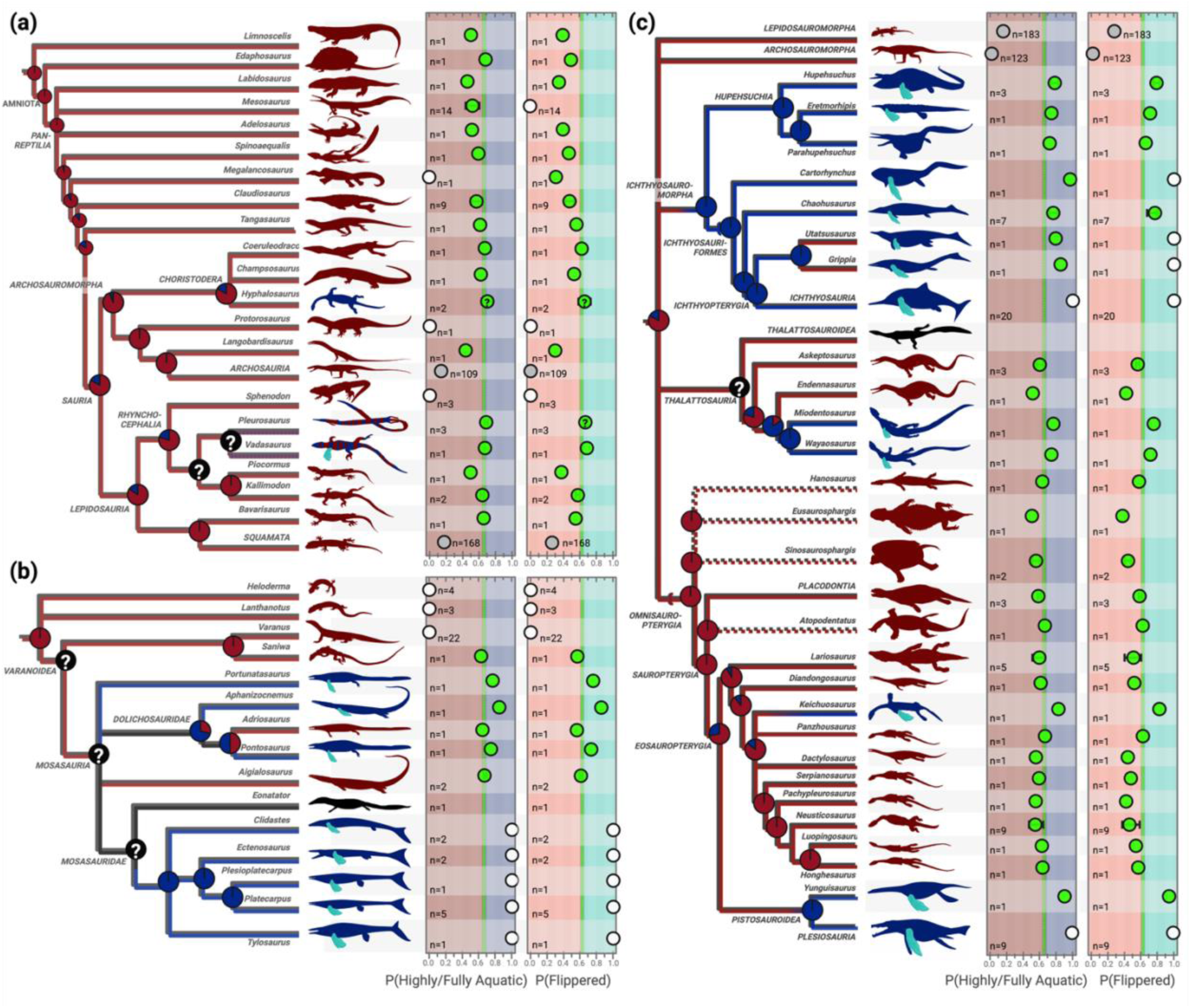
Reconstructed evolutionary history of flippers and aquatic habits in extinct pan-reptiles. (A–C) Our best-performing models (using Lhand / Lzeugopodium values) give the predicted probabilities (from 0 to 1) of highly/fully aquatic habits (dark blue) and flippers (cyan) in extinct stem reptiles (A), mosasaurians (B), and other Mesozoic marine reptiles (C). Green circles indicate mean phenotype probabilities predicted by our model, with error bars showing full mean prediction ranges for all taxon replicates. White circles indicate phenotype probabilities that were fixed *a priori* based on morphological considerations or direct observation of extant species. Gray circles indicate phenotype probabilities that were derived from a combination of fixed and predicted tree tips using ancestral-state reconstructions. Each dashed green line indicates the probability classification threshold for the associated model. Tip taxa with recovered phenotype probabilities significantly higher than the classification threshold (one-tailed t-test, *p* < 0.05) are colored accordingly, as either dark red (possibly semi-aquatic, but regularly returning to land) or dark blue (highly or fully aquatic, seldom if ever leaving the water), with cyan limb icons indicating flipper presence. Pie charts and branch colors show ancestral-state reconstructions for aquatic habits given tip phenotypes. Taxa in black are unsampled, and black nodes have ambiguous reconstructions that depend highly on contentious tree topologies. Phylopic attributions and similar predictions for mammals are provided in the Supplementary Material (Document S1).

**Figure 4.**
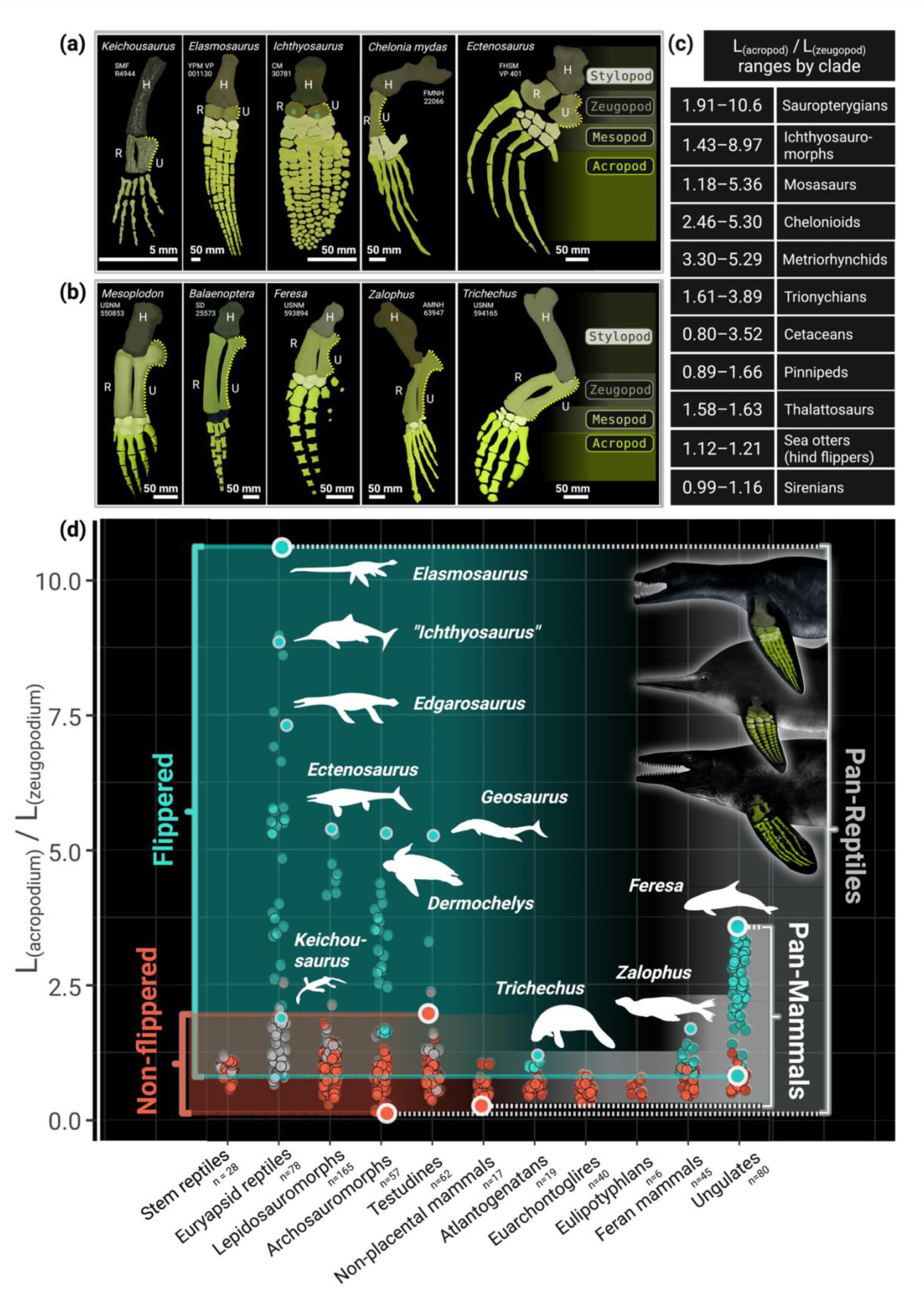
Morphospace occupancy and anatomical differences among reptile and mammal limbs. (A–B) Select forelimb skeletons of flippered pan-reptiles (A) and pan-mammals (B) in our dataset are shown color-coded by limb region. In contrast to many flippered reptiles, those flipper-bearing mammals with the highest acropodium-to-zeugopodium length ratios retain well-defined, medially sculpted posterior ulnar margins, indicating more strongly maintained long bone identity in the zeugopodials. Those flippered reptiles that retain similar sculpting (e.g., *Chelonia*, *Ectenosaurus*) elongate the individual distal phalanges noticeably more than any flippered mammals. (C) Table shows absolute acropodium-to-zeugopodium length ratios for all flippered amniote clades. (D) Strip plot compares the breadth of relative hand lengths among flippered vs. non-flippered pan-mammals and pan-reptiles, illustrating the limited morphospace occupancy of mammals. Phylopic attributions and additional discussion are provided in the Supplementary Material (Document S1).

*Amniota*, *Mammalia*, *Ungulata*, *Artiodactyla*, *Pan-Reptilia*, and *Sauria* were all recovered as ancestrally terrestrial regardless of competing hypotheses for pan-reptile interrelations (Figs 3a, S16). Among pan-reptiles, highly or fully aquatic habits were absent outside crown *Sauria*, even for the semi- aquatic taxa *Mesosaurus*, *Spinoaequalis*, and *Claudiosaurus* (Fig. 3a). Within *Lepidosauria*, mosasaurians showed a dynamic pattern of aquatic adaptation, reminiscent of previous suggestions^26,28,43^ (Fig. 3b). Highly/fully aquatic habits and flippers were recovered in some Mesozoic mosasaurians (i.e., the dolichosaurs *Portunatasaurus*, *Aphanizocnemus*, and *Pontosaurus*), but not in others (i.e., *Adriosaurus*, *Aigialosaurus dalmaticus*), suggesting multiple origins for this ecology within *Mosasauria*.

In contrast, the Triassic marine reptile clades *Ichthyosauromorpha* and *Thalattosauria* showed single origins of highly/fully aquatic habits (Fig. 3c). All sampled ichthyosauromorphs (including hupehsuchians, nasorostrans, basal ichthyosauriforms, and *Chaohusaurus* spp.) were recovered as flippered and highly/fully aquatic, pushing the origin of these phenotypes back to the earliest Triassic^44^ or Permian^45^ (Fig. 5d). In contrast, highly/fully aquatic habits arose deep within *Thalattosauria* (Fig. 3c). The basal-most askeptosauroids lacked flippers and retained ties to land, whereas the derived *Wayaosaurus* and *Miodentosaurus* were recovered as flippered and highly/fully aquatic, suggesting a loss of consistent terrestrial ties (i.e., a transition from moderately to highly/fully aquatic habits) in the common ancestor of these taxa in the Ladinian or Carnian, 242–228 Ma^46^ (Fig. S17; Table S10).

The third major clade of Triassic marine reptiles—which, for lack of an alternative, we call “*Omnisauropterygia*” (tax. nov.), defined as the largest clade comprising *Sauropterygia* but not *Ichthyosauromorpha*, *Thalattosauria*, *Archosauria*, or *Lepidosauria* (Text S2)—shows multiple independent origins of flippers and highly/fully aquatic habits (Fig. 3c). The basal-most omnisauropterygians, as well as all sampled placodonts and most eosauropterygians, lacked flippers and retained consistent ties to land, living at most an amphibious lifestyle (one-tailed t-tests, *p* > 0.05). Among *Eosauropterygia*, flippers and highly/fully aquatic habits were recovered only in *Keichousaurus hui* and pistosaurs. Ancestral-state reconstructions thus robustly recovered two separate origins of flippers and highly/fully aquatic lifestyles— one in *Keichousaurus* and one in *Pistosauria*, both of which originated during the Early Triassic^47^ (Fig. S17).

## DISCUSSION

Our model predictions lend new clarity to ongoing disputes about the semi-aquatic habits of various extinct taxa, and thus shed light on previously ambiguous temporal patterns of marine adaptation in reptiles. All semi-aquatic reptiles from the Paleozoic have ecologies that are either disputed (e.g., for mesosaurs^9,10^) or sparsely explored (e.g., for *Spinoaequalis*^48^ and *Claudiosaurus*^31^). Our best models confidently reject highly/fully aquatic habits in all these taxa, clarifying that despite their apparent morphological adaptations to life in the water,^9,31,32,33,48^ these semi-aquatic taxa maintained regular ties to land. These results dovetail with recent studies^10,33,49^ on the anatomy of the axial skeleton in mesosaurs that have suggested semi- terrestrial habits. They also make sense of the plesiomorphic limb features (well-ossified limb joints and autopodial elements) retained in most basal askeptosauroids, omnisauropterygians, and mosasaurians, which would have enabled some overland travel.^17,30,50,51,52,53^

These semi-aquatic predictions contrast with our recovery of both highly/fully aquatic habits and flippers in the eosauropterygian *Keichousaurus*; the thalattosauroids *Miodentosaurus* and *Wayaosaurus*; the basal mosasaurians *Adriosaurus*, *Aphanizocnemus*, and *Portunatasaurus*; and all known ichthyosauromorphs, which highlights lineage-specific patterns of marine adaptation in Mesozoic reptiles. In particular, we report two independent origins of highly/fully aquatic habits and flippers in sauropterygians, confirm existing suggestions for multiple independent origins of highly/fully aquatic habits and flippers among mosasaurs,^26,28,43^ and clarify that a semi-terrestrial, non-flippered ichthyosauromorph has yet to be found. These predictions clarify the timing and extent of aquatic invasions and concomitant morphological innovations by Mesozoic reptiles, with implications for interpreting their functional ecologies and eventual extinctions (Text S13–S14).

Taken more broadly, the morphometric trends recovered here shed light on general patterns of amniote limb evolution. For example, the high predictability of flippers and the low predictability of interdigital webbing likely reflects a weaker association between interdigital webbing and aquatic (especially highly/fully aquatic) habits than we tend to intuitively assume (Fig. S18; Text S15). In addition, our results quantify the textbook^2,29,30,54^ convergence of amniote flippers, long noted but sparingly quantified across *Amniota*,^18,55,56,57,58^ and demonstrate that highly/fully aquatic amniotes converge on a distinct morphometric profile beyond the bounds of terrestrial morphospace (Figs. 2, 4; Text S16). In particular, we find that distal limb elongation takes place in *all* flippered amniote lineages, despite the myriad axial, appendicular, lift- based, and drag-based locomotor modes they employ.^59,60^ This recurring limb morphometry suggests strong biomechanical incentives for acropodial elongation in groups committed to a highly/fully aquatic lifestyle. For underwater fliers (like sea turtles and plesiosaurs) who use their flippers as hydrofoils,^60^ acropodial elongation should optimize thrust efficiency by reducing tip vortex separation in the foil’s wake, increasing lift and minimizing induced drag.^61^ For rowers (like trionychids) who use their flippers as oars or paddles, longer digits should increase paddle surface area, producing more propulsive drag when oriented perpendicular to flow.^18,60^ Even in axial swimmers (like cetaceans), who use their flippers primarily for stability or maneuverability, increasing flipper surface area often enhances hydrodynamic performance, affording either more rapid or more energy-efficient maneuvers, depending on flipper design.^62^ Thus, we suggest that acropodial elongation tends to confer a biomechanical advantage to flippered species regardless of swimming mode, explaining the ubiquity of this trend in flippered mammals and reptiles.

Moreover, our results hint at clade- and habitat-specific adaptive landscapes for aquatic and terrestrial mammals and reptiles. The greater range of relative hand lengths observed among flippered species (Fig. 4c) indicates that highly/fully aquatic amniotes have more disparate limb morphometries than terrestrial ones, extending a recently discovered mammalian^56^ trend to reptiles for the first time, and hinting at distinct evolutionary landscapes for aquatic and terrestrial limbs across amniotes. Even among aquatic taxa, our results suggest clade-specific adaptive landscapes, contoured by developmental constraints that prohibit certain evolutionary outcomes (Text S17). Aquatic reptiles repeatedly achieve higher and more variable acropodium-zeugopodium ratios than any aquatic mammals (Fig. 4), regardless of their flippers’ propulsive function or degree of hyperphalangy. Reptiles may have attained these extreme morphometries by “blurring” the identities of the major limb regions, so that their zeugopodials became less like long bones and more like wrist bones.^54,63,64^ Previous studies have linked this tendency to *HoxA*/*HoxD* repatterning within the early limb bud,^63,64^ as these genes specify limb region identity along the limb’s proximodistal axis.^65^ Experimental data support the potential role of these genes in distal limb truncation,^66,67,68^ suggesting a plausible developmental mechanism that seems to have been exploited naturally in several groups of flippered reptiles, but never among flippered mammals (Fig. 4a). This may hint at a fundamental constraint against *HoxA*/*HoxD* repatterning in the mammalian limb bud, which bars mammals from entering the full breadth of morphospace accessible to flippered reptiles.

Moving beyond amniote limbs, the methods employed here offer a highly transferable new analytical pipeline for reconstructing the evolutionary histories of extinct species. Our statistical results were robust against pairwise trait correlations (Figs. S19–27), differences in measurement methods (Fig. S28), and major changes in tree topology and data transformation methods (Fig. S29; Text S8), demonstrating a strong biological signal. This signal was derived from simple input data (length measurements and ratios) that are easy to obtain from fragmentary specimens, suggesting an approach for reconstructing cryptic phenotypes that is broadly applicable to other questions in vertebrate evolution, such as the origins of flight in theropods and terrestriality in tetrapods. This pipeline highlights the power of ROC analysis^42^—largely unutilized by paleontologists and incorporated here into a large-scale phylogenetic statistical framework for the first time—to compare the predictive accuracies of competing models for a given region of tree space. Future studies can build upon the framework we establish here and employ phylogenetic ROC analysis to greater ends, answering formerly intractable questions about the evolutionary history of life on Earth with unprecedented clarity.

## RESOURCE AVAILABILITY

### Lead contact

Further information and requests for this paper should be directed to and will be fulfilled by the lead contact, Caleb Gordon.

### Data and code availability

- All the data associated with this study are provided on Dryad: https://doi.org/10.5061/dryad.08kprr5fn.
- All original code for this project is available on Dryad: https://doi.org/10.5061/dryad.08kprr5fn.
- Any additional information required to reanalyze the data reported in this paper is available from the lead contact upon request.

## Supporting information

Supplementary R Scripts

## ACKNOWLEDGMENTS

This paper required the support of many generous people and institutions. We would especially like to thank Rebecca Vitkus and Armita Manafzadeh for their helpful edits; Thomas Langford, Aya Nawano, and the Yale Center for Research Computing (YCRC) for assistance with Yale’s Grace High-Performance Computing Cluster; Brandon Mercado and Yale’s Chemical and Biophysical Instrumentation Center for access to and training on the micro-CT scanner; Raquel Jaramillo for designing the photorealistic silhouettes shown in Figures 1 and 6; and Jessica Utrup and Susan Butts for help with image stacking. This project also benefited from helpful discussion with Gregory Watkins-Colwell, Kristof Zyskowski, Zachary Morris, and Antoine Verrière on character scorings; David Polly and Savannah Olroyd on landmarking; Jonathan D. Reuning-Scherer, Carlo Meloro, Lucinda Sisk (Yale StatLab), and Kyle Stanley on statistical methods; and Kelsey M. Jenkins, Dalton Meyer, Alexander Ruebenstahl, and Luis Leonardo Hostos Olivera on amniote systematics. For collections access or specimen images we are indebted to Gregory Watkins-Colwell, Kristof Zyskowski, Vanessa Rhue, Daniel Brinkman, Ryosuke Motani, Michael Caldwell, Amy Henrici, Eleanor Hoeger, Carl Mehling, and dozens of others (listed in our extended acknowledgments: Supplementary Text S17). This research was made possible by financial support to C. M. G. via the Yale Institute of Biospheric Studies (Doctoral Pilot Grant, Doctoral Dissertation Improvement Grant), the IUCN Crocodile Specialist Group (Fritz-Huchzermeyer Student Research Assistance Scheme), and a National Science Foundation Graduate Research Fellowship under Grant No. DGE1752134.

## AUTHOR CONTRIBUTIONS

Conceptualization, C. M. G.; methodology, C.M.G., L.S.F., C.T.G., J.A.G., and B.-A.S.B.; investigation, C.M.G. and L.S.F.; writing—original draft, C.M.G.; writing—review & editing, C.M.G., L.S.F., C.T.G., J.A.G., and B.-A.S.B.; funding acquisition, C.M.G. and B.-A.S.B.; resources, C.M.G., B.-A.S.B., and J.A.G.; supervision, C.T.G., J.A.G., and B.-A.S.B.

## DECLARATION OF GENERATIVE AI AND AI-ASSISTED TECHNOLOGIES

During the preparation of this work, C.M.G. used ChatGPT v. 2 to generate <0.1% of the original code required for data analysis, in order to assist with trivial aspects of list processing, graph color-coding, and correlation matrix construction in R. After using this tool or service, C.M.G. and L.S.F. reviewed and edited the content as needed and take full responsibility for the content of the publication.

## SUPPLEMENTAL INFORMATION

**Document S1. Text S1–S17 and Figures S1–S30** (this is the main PDF).

This document contains all of the supplementary text and supplementary figures associated with this study.

Table S1. All linear morphometric, limb phenotype, and aquatic affinity data.

Additional information on the measurements taken is provided in Figure 1 and the STAR Methods. Abbreviations: Antmost, anterior-most; DistWidth, distal epiphysis width; FOD, first occurrence datum, in mya; EFZL, effective average forelimb zeugopodial length [Average (LRadius, LUlna)]; f_ASR, forelimb acropodium-to-stylopodium length ratio; f_AZR, forelimb acropodium-to-zeugopodium length ratio; f_subtype, forelimb soft-tissue phenotype (narrow classification); f_SZR, forelimb stylopodium-to- zeugopodium length ratio; fore_type, forelimb soft-tissue phenotype (broad classification); h_subtype, hindlimb soft-tissue phenotype (narrow classification); hind_type, hindlimb soft-tissue phenotype (broad classification); H, height; ID, specimen identification number; L, length; LOD, last occurrence datum, in mya; mD, manual digit; mD3, manual digit III; mm, millimeters; mya, million years ago; PBDB, the Paleobiology Database; pD, pedal digit; pD3, pedal digit III; Postmost, posterior-most; proxp, proximal phalanx; proxWidth, proximal epiphysis width; Symmetry_index1, acropodial symmetry index 1 (Lanterior-most digit / Lposterior-most digit); Symmetry_index2, acropodial symmetry index 2 [Llongest digit / Average (Lanterior-most digit, Lposterior- most digit)]; up, ungual phalanx; FI, one of three ungual phalanx flatness indices (see STAR Methods). All institutional abbreviations are given in Text S2 of Document S1. All literature sources are linked using either stable URLs or full-length citations inline within the table at each mention.

Table S2. All repeat length measurements and ratios for Bland-Altman analysis.

Additional information on the Bland-Altman analysis performed is provided in the STAR Methods. Abbreviations: fAnAR, forelimb acropodium-to-*non*-acropodium length ratio; fASR, forelimb acropodium-to- stylopodium length ratio; fAZR, forelimb acropodium-to-zeugopodium length ratio; fSZR, forelimb stylopodium-to-zeugopodium length Ratio; hAnAR, hindlimb acropodium-to-*non*-acropodium length ratio; hASR, hindlimb acropodium-to-stylopodium length ratio; hAZR, hindlimb acropodium-to-zeugopodium length ratio; hSZR, hindlimb stylopodium-to-zeugopodium length ratio; Index, arbitrary specimen index number used during data collection; Method, broad measurement method used; Submethod, narrower measurement method used; All institutional abbreviations are given in Text S2 of Document S1.

Table S3. Stitched results from all phylogenetic ANOVAs, Levene’s, and post-hoc tests.

These results were generated using the phyloBoxPlot function with 10,000 simulations and the Benjamini-Hochberg (BH) correction method for post-hoc tests. These results were generated in script_S2.R and stitched together in script S5. Abbreviations: Fstat, f-statistic; phylANOVA, phylogenetic ANOVA; phyloLev, phylogenetic Levene’s test; pval, p-value. Additional abbreviations and names for input datasets, variables, and supertrees are given in script S1.

Table S4. Stitched results from all phylogenetic correlation tests.

These results were generated using the phycorr function (REML = TRUE, optim_method = Nelder-Mead, constrain.d = TRUE, maxit.NM = 500), within script_S3.R in massively parallel fashion on 10,000 CPUs using job arrays. All outputs from script_S3 jobs were stitched together to form the CSV file corresponding to this table in script_S5.R. Abbreviations: AIC, Akaike Information Criterion; BIC, Bayesian Information Criterion; corvariable, one of two variables being correlated; logLik, log likelihood; p.val, p-value; Rsq, R-squared; Std.Err, standard error. Additional abbreviations and names for input datasets, variables, and supertrees are given in scripts S1 and S3.

Table S5. Stitched results from all phylogenetic binomial logistic regression (phybLR) and Receiver-Operating Characteristic (ROC) analyses.

These results were generated by means of the phybLR function (Kfold = 3, CV_runs = 100, bootnum = 10000, btol = 35) within script_S4.R in massively parallel fashion on 10,000 CPUs using job arrays. All outputs from script S4 jobs were stitched together to form the CSV file corresponding to this table in script S5. Abbreviations in column headings: 95ci, 95% confidence interval; µ, sample mean across 10,000 bootstrap replicates of the specified variable; τ, threshold probability for classifying binary phenotypes; Acc, Accuracy; AIC, Akaike Information Criterion; AUC, Area Under the ROC Curve; bestmod, the best phybLR model for this combination of inputs using the optimal classification threshold and fit to all the data; binomt, binomial test; ds, dataset; FPR, False Positive Rate; highb, high bound; lowb, low bound; McF, McFadden’s; n_obs, number of observations; p, p-value; Rsq, R-squared; TNR, True Negative Rate; TPR, True Positive Rate. Additional abbreviations and names for input datasets, variables, and supertrees are given in scripts S1 and S4. Additional metadata from each joint phybLR and ROC analysis are available in Data S4.

Table S6. Phylogenetic comparative test metadata for sensitivity analyses concerning tree topology and data transformation method.

These results were the input data for Figure S29, generated within script_S5.R using statistical test results stitched together from scripts S2–S4. Table S6a summarizes the correspondence among significance test verdicts for input datasets that were (prior to testing) either not transformed, Box-Cox transformed, or log10- transformed. Table S6b summarizes the correspondence among significance test verdicts for input datasets given the eight different input tree topologies considered in this study. The table summarizes the count of significance tests with identical significance verdicts (based on p-values) across all considered classes and the count of significance tests with divergent (non-identical) significance verdicts among the considered classes, and the results of an associated binomial test to test for the significance of the disagreement among test classes in each case. More details about these sensitivity analyses and the associated abbreviations and metadata are given in script S5, the STAR Methods, and Text S7 within Document S1.

Table S7. Tip phenotype predictions made by our best-performing phylogenetic binomial logistic regression models for all specimens in our dataset.

These results were generating using the tipPreds function within script S5. Abbreviations in column headings: mean, sample mean across 10,000 bootstrap replicates of the specified variable; Acc, Accuracy; bestmod, the best phybLR model for this combination of inputs using the optimal classification threshold and fit to all the data; bestmod_threshold, threshold probability for classifying binary phenotypes; binomt, binomial test; df, degrees of freedom; FPR, False Positive Rate; known Prob, known probability in cases when probability of phenotype was fixed based on observations from extant taxa or other considerations; predictor_var, predictor variable in phybLR model; predProb, predicted probability of phenotype; prob_type, denotes whether tip probability was fixed or predicted by the phybLR model; pval, p-value; response_var, response variable in phybLR model; sd, standard deviation; TPR, True Positive Rate; ttest, t-test comparing mean predicted probability to classification threshold; tval, t-value from t-test; tx, tx score. All institutional abbreviations are given in Text S2 of Document S1. Additional abbreviations and names for input datasets, variables, and supertrees are given in scripts S1 and S5.

Table S8. Internal node phenotypes inferred from the tip states in Table S7.

These results were generated using the nodePreds function within script S5. Abbreviations in column headings: asr, ancestral state reconstruction; P(), probability. All institutional abbreviations are given in Text S2 of Document S1. Additional abbreviations and names for input datasets, variables, and supertrees are given in scripts S1 and S5.

Table S9. Summary of geometric morphometric data generated for this study.

This table includes a summary of the metadata, Procrustes coordinates, and PC scores generated for the three geometric morphometric analyses associated with this study. Abbreviations: GM_Analysis, geometric morphometric analysis performed; PC N; Principal Component #N; PERMANOVA, permutational ANOVA; PCA, principal component analysis; phyLANOVAs, phylogenetic ANOVAs; n_fixed, number of fixed landmarks, n_sliding, number of sliding landmarks; Xn, Procrustes X-coordinate #N. All institutional abbreviations are given in Text S2 of Document S1. Additional abbreviations and explanations for input variable names are provided in scripts S1 and S6. Additional metadata associated with all geometric morphometric analyses are provided in Data S6.

Table S10. Temporal ranges for partial and complete aquatic invasions in extinct amniotes.

This table includes the sources and considerations employed to give the age ranges for aquatic invasions illustrated in Figure S17. Abbreviations: Ma, *Mega anna*. Additional context for this table is provided in Text S14 within Document S1.

**Script S1.** This script begins the analysis by processing the dataset, calibrating all supertrees, completing a few computationally inexpensive analyses, and defining original functions and R objects required for downstream scripts. For more information about this script’s function in the context of the study as a whole, please see the STAR Methods, as well as Text S1 and Figure S30 within Document S1.

**Script S2.** This script performs all phylogenetic ANOVAs, Levene’s tests, and post-hoc tests, and generates associated box plots and ternary plots referenced in the paper. For more information about this script’s function in the context of the study as a whole, please see the STAR Methods, as well as Text S1 and Figure S30 within Document S1.

**Script S3.** This script performs all the phylogenetic correlation tests reported in Document S1. For more information about this script’s function in the context of the study as a whole, please see the STAR Methods, as well as Text S1, Text S9, and Figure S30 within Document S1.

**Script S4.** This script fits all the phylogenetic binomial logistic regression models referenced in the paper. For more information about this script’s function in the context of the study as a whole, please see the STAR Methods, as well as Text S1 and Figure S30 within Document S1.

**Script S5.** This script stitches together the outputs from previous scripts, uses Receiver-Operating Characteristic (ROC) analysis to select the best-performing models and classification thresholds from script S4, and uses those selected model-threshold pairs and ancestral-state estimations to reconstruct the tip and internal node phenotypes for all the extinct taxa in the dataset. This script also includes a sensitivity analysis to assess the impact of differing tree topologies and data-transformation methods on our results. For more information about this script’s function in the context of the study as a whole, please see the STAR Methods, as well as Text S1, Text S8, and Figures S29–S30 within Document S1.

**Script S6.** This script contains all of the R code required to perform the geometric morphometric analyses associated with this study. For more information about this script’s function in the context of the study as a whole, please see the STAR Methods, as well as Text S1, Text S7, Figure S10, and Figure S30 within Document S1.

**Data S1.** This zip folder contains the BASH scripts required for running the R scripts above on a high-performance computing cluster; the Newick files required to generate time-calibrated trees in script S1; and all script S1 outputs, including high-resolution versions (both ultrametric and tip-dated) of the eight supertrees assembled for this project, data transformation plots, and category association plots. For more information about how this data file ties into the workflow of the study as a whole, please see Text S1 and Figure S30 within Document S1.

**Data S2.** This zip folder contains PNGs of all the box plots and associated statistical test results generated in script S2. For more information about how this data file ties into the workflow of the study as a whole, please see Text S1 and Figure S30 within Document S1.

**Data S3.** This zip folder contains high-resolution PNGs of the nine correlation matrices generated from the analyses run in job arrays with script S3. For more information about how this data file ties into the workflow of the study as a whole, please see Text S1 and Figure S30 within Document S1.

**Data S4.** This zip folder contains all script S4 outputs, including unstitched “RData” files, phylogenetic logistic regression analysis results, and associated metadata files. For more information about how this data file ties into the workflow of the study as a whole, please see Text S1 and Figure S30 within Document S1.

**Data S5.** This zip folder contains all script S5 outputs, including assembled consensus ROC curves, best-model prediction histograms, and supertrees with overlain ancestral state reconstructions. For more information about how this data file ties into the workflow of the study as a whole, please see Text S1 and Figure S30 within Document S1.

**Data S6.** This zip folder contains key script S6 inputs and outputs, including TPS files, sliding semi-landmark designations, and PAST files. For more information about how this data file ties into the workflow of the study as a whole, please see Text S1 and Figure S30 within Document S1.

## REFERENCES

[The references are currently formatted as endnotes at the end of this document for ease of revision during peer review. We will reformat them as a numbered list, placed here, to meet journal guidelines, prior to publication.]

## STAR★METHODS

### KEY RESOURCES TABLE

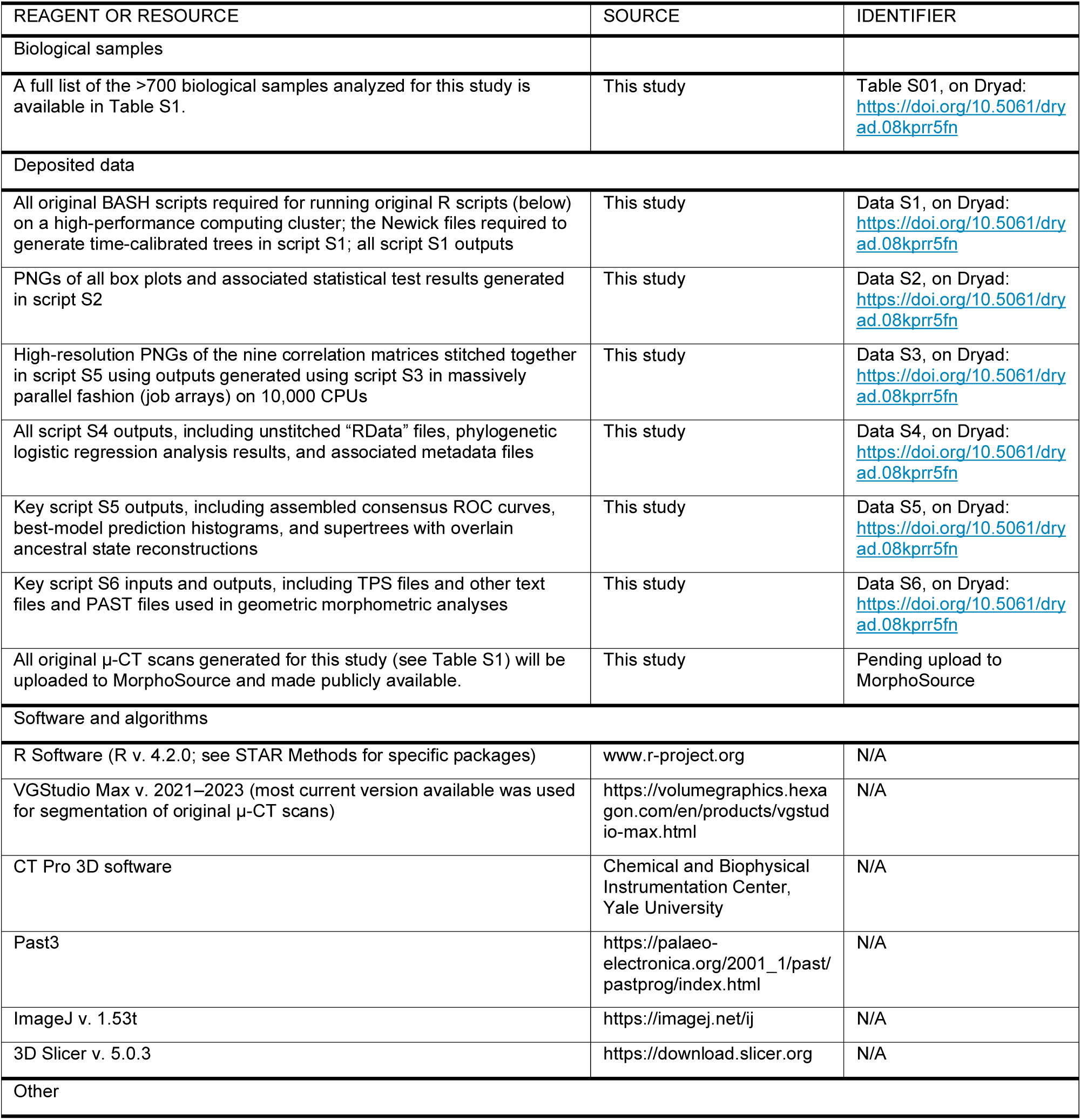
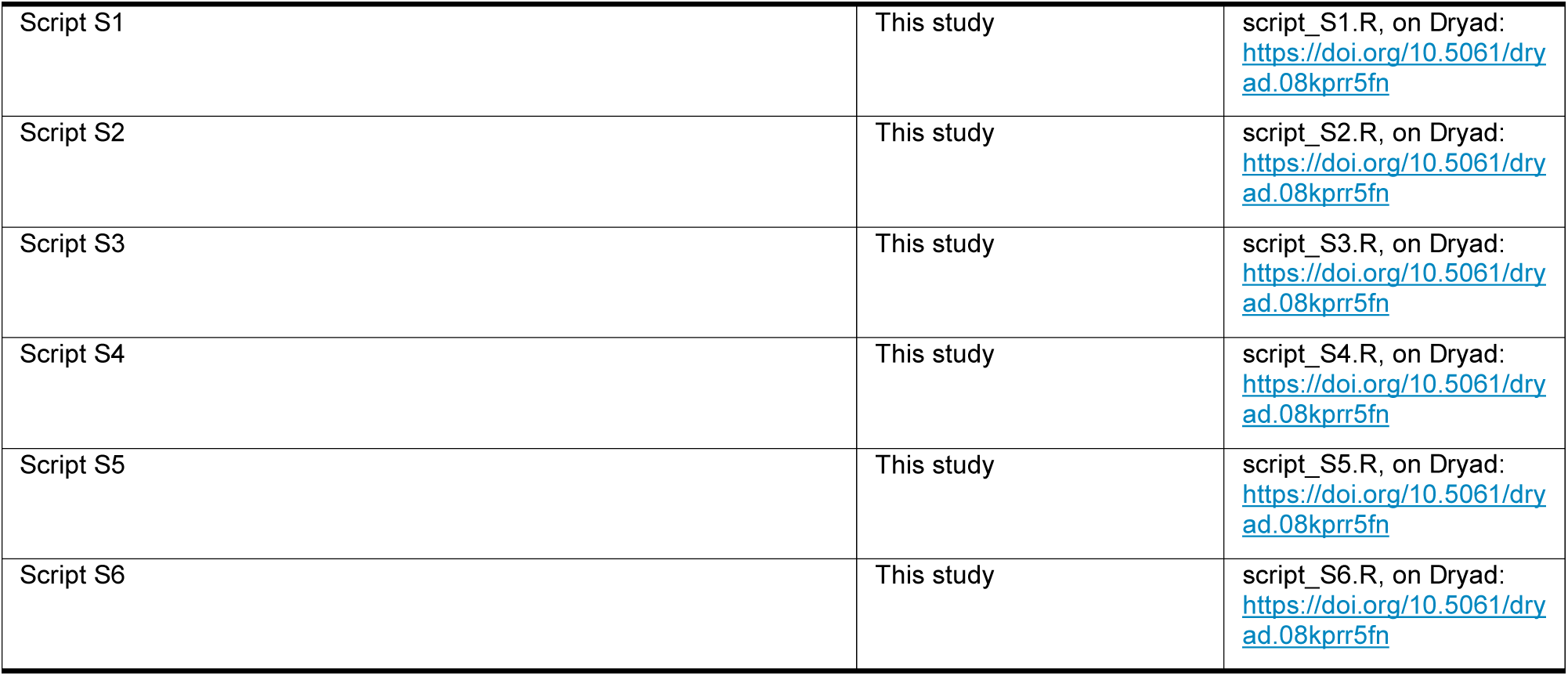

### METHOD DETAILS

To test previously posited osteological correlates of interdigital webbing, flipper phenotype, and aquatic affinity, we performed linear and geometric morphometric analyses of tetrapod limbs within a phylogenetic comparative framework and assessed the predictive accuracy of each posited correlate using phylogenetic Receiver-Operating Characteristic (ROC) analysis. Linear measurements were collected on 759 specimens: 469 extant, 290 extinct, comprising 524 specimens of pan-reptiles, 223 of pan-mammals, and 12 of tetrapods from outside of crown *Amniota* (see Table S1). Our dataset excluded limbs involved in flapping flight, so the forelimbs of avialans, and the fore- and hindlimbs of bats and pterosaurs, were omitted from our analyses. We scored webbing and flipper phenotypes using fixed or dried zoological specimens, published anatomical descriptions, or (in rare cases) exceptional soft-tissue preservation from fossil taxa, and scored aquatic habits from the natural history literature (see Table S1). Limb skeletons in our dataset were measured or landmarked and the resulting shape data analyzed among taxa with different aquatic affinities and soft-tissue phenotypes using phylogenetic comparative tests, as described below. Measurements for specimens with known phenotypes were fed into phylogenetic binomial logistic regression models, which were trained and tested with a 3-fold cross-validation procedure repeated 100 times to generate a consensus ROC curve (cROCC), as described below. We used ROC analysis to select the best of all tested models for their ability to predict the presence of webbing, flippers, or aquatic habits in species whose phenotypes are known. These best-performing models were then used to reconstruct all significantly predictable features in extinct taxa with contentious phenotypes.

#### Obtaining length measurements

Linear measurements were taken from articulated limb skeletons in one of three ways. For most specimens (n=480), measurements were obtained from either physical skeletons (n=365) or 3D mesh files (n=115). Mesh files were downloaded from MorphoSource or obtained by original µ-CT scans of Yale Peabody Museum (YPM) Vertebrate Zoology specimens. All scans were constructed using CT Pro 3D software (Nikon, Tokyo, Japan) and post-processed, examined, and segmented using VGStudio MAX v. 2021–2023 (Volume Graphics, Heidelberg, Germany). We measured physical specimens with digital calipers (VINEN model no. DCLA-1205: Valencia, CA), and 3D meshes in 3D Slicer (v. 5.0.3) using the “line” and “curve” tools (curve type: linear). For a large subset of specimens (n=279) that were splayed within a single 2D plane (e.g., fossils preserved on slabs), linear measurements were taken from scaled high-resolution photographs in planform view, which we either obtained from museum curators or collected ourselves using a DMC-GX8 camera (Panasonic Lumix, 60 mm lens) mounted to a copy stand.

Length measurements were then collected from photographs in ImageJ (v. 1.53t)^69^ using the straight- and segmented-line tools.

To test the consistency of 2D measurements taken on ImageJ and 3D measurements collected with calipers, a small subset of specimens (n = 21) were first measured with calipers and then photographed and measured with ImageJ software. Corresponding ImageJ-caliper pairs were tested for congruence with Bland-Altman analyses^70^ (Fig. S2; Script S1), which reported no significant systematic directional error for any of the linear measurements taken. For both direct length measurements and computed ratios, differences in value between measurement methods were statistically indistinguishable from zero (one-tailed t-tests, each *p* > 0.3). Mean and median absolute random errors between measurement methods were ∼0.1 mm for direct lengths measurements (Fig. S2a), and < 0.01 units for computed ratios (Fig. S2b), with a slight correlation between absolute error size and direct measurement length (Pearson coefficient = 0.3, *p* < 0.05; Fig. S2a). To reduce the observed random error, and account for the slight tendency for increased error with larger measurements, all length measurements and computed ratios were rounded (lengths ≥10 mm to the nearest 1 mm, lengths <10 mm to the nearest 0.1 mm, and computed ratios to the nearest 0.01 units).

#### Measuring limb region proportions

Limb proportions were determined by computing ratios from direct length measurements of the fore- and hindlimb stylopodium, zeugopodium, and acropodium (Fig. 2d). Humerus length was taken from the proximal-most point of the anterior process of the proximal epiphysis to the distal-most point of the distal epiphysis. Femur length was taken from the proximal-most exposed point of the femoral head to the distal-most point of the distal epiphysis. If the anterior process of the humerus was obscured (in the case of some mounted specimens), or if the processes on the proximal epiphysis were poorly defined (in the case of some cetaceans and Mesozoic marine reptiles), then the proximal-most exposed point of the humerus was used as a substitute. Zeugopodial (radius, ulna, or tibia) length was taken from the proximal-most exposed point of the element to the lateral-most point of the element’s distal epiphysis along its lateral margin. Radius and ulna lengths were averaged to provide a single estimate for forelimb zeugopodium length. For birds, hindlimb zeugopodium length was taken from the proximal- to distal-most points of the tibiotarsus. Acropodial length was proxied using the combined length of the metacarpal and the un-clawed phalanges of either digit III (for reptiles) or the longest digit (for mammals and outgroups). The ungual phalanx was excluded from the acropodial measurement of all taxa with keratinized ungual claws on the digit(s) measured. For birds, hindlimb acropodium length was taken as the combined length of the tarsometatarsus and pedal digit III, again excluding the clawed ungual phalanx. Tarsometatarsus length was taken from the proximal-most point of the *Eminentia intercotylaris* to the distal-most point of *Trochlea metatarsi III* (*sensu* Baumel & Witmer, 1993).^71^ Given that the tibiotarsus and tarsometatarsus each include portions of the mesopodium, the associated length measurements for birds have the issue of not being from strictly homologous points to those from which lengths were measured in other tetrapods (as those latter lengths did not incorporate any mesopodial elements). This concession was necessary for us to include birds within the dataset. However, we achieve the same results with or without the inclusion of birds in the dataset, highlighting that our results continue to hold when all lengths compared among species are strictly homologous.

Stylopodium, zeugopodium, and acropodium length were visually compared with ternary plots^18^ and combined in various ways to reduce multivariate data on limb proportions into one-dimensional metrics. These included the ratios of acropodium-to-zeugopodium length, acropodium-to-stylopodium length, and stylopodium-to-zeugopodium length, among others. These ratios were then compared among limb phenotypes and aquatic affinity bins visually with box plots and statistically with phylogenetic comparative tests, as described below.

#### Measuring acropodial planform shape

We quantified acropodium shape using a combination of direct length measurements and 2D geometric morphometrics (Figs. 1d, S10). Length of the anterior-most, posterior-most, and longest digits were taken as described above (but excluding the hallux in birds). From these measurements, we computed two acropodium “symmetry indices” (SIs) from these measurements: one capturing the relative symmetry of the acropodium (*i.e.*, the length ratio of the first and last digits: Lanterior-most / Lposterior-most), and one capturing the difference in length between its longest and lateral-most digits (Llongest / (Lanterior-most + Lposterior-most)/2). The usage of “anterior-most,” “posterior-most,” and “longest” digit descriptors, rather than digit identities based on homology (e.g., “digit I”, “digit V”), enables the comparison of acropodial planform shapes across all limbed amniotes regardless of digit number—so that, for example, we can include artiodactyls with four digits and ichthyosaurs with seven digits in the same analysis. Not all limbed amniotes have a “digit V”, but all do have a posterior-most digit. In the limiting case where a single full digit is present (e.g., for extant equids), the anterior- and posterior-most digits are equivalent.

These SI ratios based on linear measurements serve as effective proxies for certain aspects of hand or foot planform shape, whose relative symmetry or asymmetry has been used as evidence to suggest the presence of interdigital webbing or particular swimming styles in extinct species by previous authors.^5,9,31,32,39^ The first SI, captured by Lanterior-most / Lposterior-most, tends to reflect whether the acropodium tapers in one direction. An Lanterior-most / Lposterior-most of 1 indicates that the leading and lagging edges of the acropodium have the same length, suggesting a symmetrical acropodial planform. An Lanterior-most / Lposterior- most of >1 or <1 would imply an asymmetrical planform—that is either longer anteriorly (if this first SI >1) or longer posteriorly (if this first SI <1). The second SI ratio, (Llongest / (Lanterior-most + Lposterior-most)/2), captures another aspect of acropodial planform shape—whether the outer digits differ markedly in shape from the longest digit. This second SI ratio reflects the variation in length across all digits, and whether the longest digit is located near to or far from the center of the acropodium’s main length axis. If (Llongest / (Lanterior-most + Lposterior-most)/2) = 1, it implies that the longest digit is roughly equal in length to the outermost digits. If this SI is >1 or <1, it suggests that the longest digit deviates in length from that of the outermost digits. This deviation can result if one of the outermost digits is much longer than the other (as is the case of sea lions, where the anterior-most digit is typically the longest and the posterior-most digit typically the shortest). This deviation can also result if any of the inner digits is very long relative to the outermost one (as is the case in most terrestrial lizards, which tend to have rather large digit III and digit IV length values). Taken together, these two SI ratios [(Lanterior-most / Lposterior-most) and (Llongest / (Lanterior-most + Lposterior- most)/2)] allow us to capture two major axes of shape variation that intuitively reflect acropodial planform symmetry with only three linear measurements, enabling a large sampling of taxa that avoids difficulties associated with disputed digital homologies, differing numbers of digits, and varied *in situ* splaying positions in sampled taxa. To supplement this linear morphometric approach and obtain a more comprehensive picture of acropodial planform shape, we also performed two geometric morphometric analyses of the acropodial planform using a subset of 103 fully articulated specimens from our dataset with limbs photographed in dorsal or ventral planform view (Text S6).

#### Scoring habitat preference and webbing phenotype

We binned all samples by webbing/flipper phenotype and aquatic affinity using an original scoring scheme (Figs. 1a, S5). We defined interdigital webbing as any soft, flimsy tissue between the digits and distal to the stiff portion of the proximal acropodium (typically comprising the metacarpals but sometimes including proximal phalanges). A flipper was defined as any continuous soft-tissue structure integrating the acropodium and zeugopodium. Webbing and flipper phenotypes were further subdivided by the nature of any interdigital soft-tissue present and its anteroposterior and proximodistal extents across the acropodium (Fig. S5). Wherever possible, webbing and flipper phenotypes were determined from fluid-preserved or taxidermized zoological specimens. In some cases, we also scored webbing phenotypes from exceptionally preserved fossils (*e.g*., in the case of *Mesosaurus tenuidens*, where a few specimens preserve actual fossilized soft-tissue evidence for webbing^33^) or anatomical descriptions (*e.g.*, in the case of some carnivorans, for which webbing has been described in detail^72^). We binned samples by aquatic affinity score using a new scheme that classifies species by the relative frequencies of time they spend on land and in the water (Fig. 1a). In tandem, we also scored aquatic affinity based on saltwater vs. freshwater preferences and using a previously published scheme that classifies species based on the depth and energy level of the waters they inhabit^18^ (Fig. S7). In most cases, we scored extant taxa for aquatic affinity using field guides and primary natural history literature. However, for a select few fossil groups (*i.e.*, ichthyosaurs, plesiosaurs, mosasaurids, and metriorhynchids), morphological considerations were used to infer fully aquatic habits *a priori*. In addition, limb phenotype and aquatic affinity bins were variously re-partitioned into binary sets (*i.e.*, “Webbed” vs. “Unwebbed,” “Flippered” vs. “Non-Flippered,” “Aquatic or Semi-Aquatic” vs. “Terrestrial,” and “Terrestrial or Semi-Aquatic” vs. “Highly or Fully Aquatic”) for downstream binomial logistic regression analyses. All specimen and literature sources for soft-tissue phenotype and aquatic affinity, as well as all multinomial limb and ecotype scores, are detailed in Table S1. All binary classifications are detailed and derived within Script S1.

### QUANTIFICATION AND STATISTICAL ANALYSIS

#### Phylogenetic comparative tests

Relationships among all aforementioned length measurements, ratios, shape indices, and PC scores were compared among limb phenotype and aquatic affinity bins using statistical tests within a phylogenetic framework. All statistical tests were performed in R (v. 4.2.0-foss-2020b)^73^ on the Grace High-Performance Computing Cluster at Yale University, with varying degrees of parallelization to accelerate computational time. Our full data-analytical pipeline is summarized in Fig. S2 and detailed in scripts S1–S6.

In defining “consensus” trees for downstream statistical tests, we identified three pairs of competing phylogenetic hypotheses that might drastically influence test results given our taxonomic sample (Fig. S29; Supplementary Data S1; Text S7). We address each of these hypotheses in turn below:

• The first of these competing phylogenetic hypotheses concerns the basal-most clade of stem reptiles: Some recent analyses have recovered *Captorhinidae* (Text S2) in this position^74^, but for the preceding two decades [e.g., as captured by Müller (2003)^75^, Modesto et al. (2015)^76^, and many others] this position has typically been occupied instead by a monophyletic grouping of “parareptiles” [*sensu* Gauthier et al., 1988b^77^]. These inconsistent branching patterns might plausibly alter the results of our analyses for basally branching stem reptiles.

• The second of these competing phylogenetic hypotheses concerns the basal-most member of the largest clade that includes *Sauropterygia* but none of *Archosauria*, *Lepidosauria*, *Ichthyopterygia* or *Thalattosauria*. Unfortunately, there is no existing name for this branch-based clade, and recently coined alternatives (*Sauropterygomorpha*^25^ and *Sauropterygiformes*^47^) are explicitly node-based and sensitive to the contentious basal interrelations of *Sauropterygia* relatives. Thus, for lack of an alternative, we call this clade “*Omnisauropterygia*” and its members “omnisauropterygians” (Text S2). *Omnisauropterygia* is the clade comprising *Sauropterygia* and all taxa more closely related to *Sauropterygia* than to any of *Archosauria*, *Lepidosauria*, *Ichthyopterygia*, or *Thalattosauria*. The most basally branching member of *Omnisauropterygia* (thus defined) has often been recovered as *Saurosphargidae* in the years since this latter clade was defined.^78,79,80^ However, a recent study^47^ instead recovered *Hanosaurus hupehensis* as the most basally branching omnisauropterygian—a placement that has major implications for the ancestral character states of the clade^47^ and might, as a consequence, plausibly alter the results of our analyses for basal omnisauropterygians and any other closely related Triassic marine reptiles.

• The third of these competing phylogenetic hypotheses concerns the tree topology of *Squamata* (Text S2). Conflicting tree topologies for *Squamata* have been suggested by phylogenetic analyses employing molecular^81^ and morphological^82^ datasets. These conflicting topologies might plausibly alter the results of our analyses for extinct and potentially semi-aquatic lepidosaurs.

To address these disagreements, we created a separate maximal supertree for each combination of these hypotheses, constructing a total of 2^3^ = 8 maximal supertrees as inputs for downstream statistical tests (Fig. S29b). These different tree topologies are summarized in Fig. S29b, and their full topologies are available in Data S1. Specimens in our dataset were placed within each of these supertrees based on current phylogenetics literature (sources in Table S1). All species replicates were treated as separate tips and placed in a polytomy, as were any major taxonomic uncertainties. Thus, for example, the Mesozoic marine reptile “superclade” recovered in most recent analyses^17,45^ was placed in a polytomy with *Archosauromorpha* and *Lepidosauromorpha* within each supertree.

For *Mammalia* (Text S2), we assumed the consensus topology recovered from recent phylogenomic analyses of mammals.^83^ We accounted for uncertainty in certain portions of the tree—e.g., for the branching order near the base of *Euarchontoglires*, which has often been contested^84^—by placing the contenders for its most basally branching member in a polytomy. Additional conflicting hypotheses for the tree topology of *Mammalia*—those that required rerunning all analyses with separate, alternative supertrees—were not considered, for two reasons: First, because incorporating these additional trees would have increased the computational resources required to run all analyses exponentially; and second, because these were not taken to bear on any of the sampled extinct taxa with ambiguous phenotypes in our dataset. One notable conflicting set of hypotheses concerns the contested clade *Atlantogenata*,^85^ comprising *Afrotheria* + *Xenarthra* to the exclusion of all other placentals (Text S2).

*Atlantogenata* is recovered as monophyletic by most molecular phylogenetic analyses [see Álvarez- Carretero, et al. (2022)^83^ and references therein] as well as some recent morphological phylogenetic analyses.^86^ However, some molecular analyses^87^ and several morphological and combined-evidence analyses [e.g., O’Leary et al. (2013)^88^ and references therein] reject *Xenarthra*-*Afrotheria* monophyly, instead recovering *Xenarthra* and *Afrotheria* as separate, successive outgroups to the rest of placental mammals. Because our dataset included no extinct xenarthrans or afrotheres of dubious aquatic affinity (e.g., no potentially semi-aquatic stem sirenians or other atlantogenatans with disputed aquatic habits), these conflicting hypotheses were not considered likely to alter any of our predictions for extinct transitional species, and were thus not worth the exponential increase in computational time that would have been required to consider them both explicitly. As a result, the three sets of alternative pan-reptile tree topologies above were the only competing hypotheses considered, giving us a total of 2^3^ = 8 maximal supertrees as inputs for downstream statistical tests, as described above.

All eight supertrees were written manually in Newick format and converted to a “phylo” object with the read.tree command in the R package *ape* (v. 5.6).^89^ The resulting tree was then annotated and visualized using the R *ggtree* package (v. 3.4.4),^90^ and tip-dated with the DatePhylo command in the R *strap* package (v. 1.6),^91^ using first and last occurrence data siphoned from the Paleobiology Database (https://paleobiodb.org/) and the literature (sources in Table S1). Each maximal tree was pruned and aligned to our morphometric datasets, and then used as input to perform the downstream phylogenetic comparative tests described below, using original R code provided in Script S1.

To check for relationships among all quantitative variables, we performed phylogenetic correlation tests on all variable pairs and repeated this process once for each of our 8 supertrees (Figs. S3–S11).

Phylogenetic correlation tests were performed using corphylo command (REML=T, constrain.d=T, method="Nelder-Mead", maxit.NM=500; *ape* v. 5.6).^89^ Phylogenetically corrected correlation matrices were then constructed using the cor.test() function (*stats* v. 4.2.1)^73^ and visualized with the corrplot.mixed command (*corrplot* v. 0.92).^92^ To compare variable distributions among all scored limb type and aquatic affinity bins, we performed simulation-based phylogenetic analyses of variance (phylANOVAs)^93^ and phylogenetic Levene’s tests (phyloLevs) with pairwise *post-hoc* Tukey comparisons for all predictor-response variable pairs in our dataset, using original R functions (Script S1) incorporating the phylANOVA command (nsim=10000, posthoc=T, p.adj="BH") from the R package *phytools* (v. 1.2-0).^94^

To assess the impacts of competing phylogenetic hypotheses and different data processing methods on the results of our phylogenetic comparative tests, we performed a comprehensive sensitivity analysis (Fig. S29). All phylogenetic comparative tests (*i.e.*, phylogenetic correlation tests, phylANOVAs, phyloLevs, pairwise post-hoc tests, and all tests associated with the phylogenetic logistic regression analyses detailed below) were rerun (for every dataset-variable combination) under each of our eight competing tree topologies and following three different data-processing methods: one using the raw, untransformed data; one using log-transformed data; and one using Box-Cox–transformed data.^95^ Equivalent and inconsistent significance verdicts (*p* > 0.05 vs. *p* < 0.05) were then determined for a given dataset-variable combination as follows: If *any* of the significance verdicts were different among the eight supertrees or among the three data-transformation methods, that test result was considered “inconsistent”; otherwise, if *all* significance verdicts were the same among the eight supertrees or among the three data-transformation methods, that test result was considered “equivalent.” For each phylogenetic comparative test, frequencies of equivalent and inconsistent significance verdicts were visualized using stacked bar plots, and a one-tailed binomial test^96^ was performed to assess whether the frequency of “inconsistent” significance verdicts was significantly greater than 0 (in which case *p* < 0.05). Phylogenetic correlation test verdicts differed at high frequencies across testing groups, but the results of all other phylogenetic comparative tests were highly robust against differences in tree topology or prior data transformation (Fig. S29; Text S7). Given the reliance of phylogenetic correlation results on tree topology, we present a separate phylogenetic correlation matrix for every supertree in our analysis (Figs. S20–S27). For other comparative tests, associated main-text figures and supplementary figures present only the results from untransformed data assuming the “supertree1” topology (Fig. S29b), unless stated otherwise. However, corresponding results for all other supertrees and transformation methods are available in Tables S3–S4 and Supplementary Data S2–S5.

#### Phylogenetic binomial logistic regression and Receiver-Operating Characteristic (ROC) analyses

After performing phylogenetic comparative tests, we filtered our raw, untransformed datasets to include just those taxa with “known” phenotypes (i.e., species whose phenotypes we can directly observe or for which we have direct soft-tissue evidence). Most of these were Recent taxa, whose limb phenotypes and aquatic affinities could be verified directly by observation of zoological specimens and reference to the natural history literature. However, some of the taxa with known phenotypes in this filtered dataset were extinct: For example, we know that ichthyosaurs were fully aquatic, as they could not physically support their weight on land,^51^ and we know that mesosaurs had webbed hands and feet, because multiple specimens of this monospecific taxon preserve exceptional soft-tissue webbing between all digits^33^ (Table S1). On this filtered dataset of taxa with known phenotypes, we used a machine-learning approach to fit phylogenetic binomial logistic regression curves^97^ that we could use to predict the ambiguous phenotypes of other extinct species. For a given dataset and tree, all limb phenotype and aquatic affinity bins were converted into binary classification schemes (Fig. S14), and a phylogenetic binomial logistic regression (phybLR) model was trained and tested using a tree subsampling procedure involving 3-fold cross-validation with 100 iterations (Fig. S3b). For each iteration of this procedure, the dataset of taxa with known phenotypes was split randomly into thirds, two thirds of the data was used to fit (or “train”) a phybLR model, and the remaining third was fed into the trained model to test its performance.

We assessed model performance using Receiver-Operating Characteristic (ROC) curve analysis—a collection of methods that was originally developed by the U.S. Army Signal Corps in World War II^98^ and has since become a common technique in medical decision-making research.^42^ ROC analysis provides an effective set of tools for researchers to visualize, report, and compare the performance of multiple competing predictive models.^42,99^ It centers around the ROC curve—a plot of true positive rate (TPR) vs. false positive rate (FPR) values for a particular predictive model whose coordinates differ depending on the predicted probability threshold for classifying a binary response variable (Fig. S3). Competing predictive models can be compared in ROC space using the area under the curve (AUC)—a value between 0 and 1 that corresponds to a model’s predictive accuracy across all classification thresholds.^42^ In the context of the present study, the area under the curve (AUC) of a given ROC plot equals the probability that the associated logistic regression model will assign higher predicted probabilities to specimens with a given phenotype than specimens without it.^42^ In this way, AUC correlates with and serves as a scalar proxy for model accuracy, as models with higher AUCs will tend to predict more true positives and fewer false negatives across classification thresholds.^42^ For each of the 1,000 iterations in our phybLR training-and-testing procedure, we assessed model performance by computing model TPR and FPR for every predicted probability classification threshold (from 0.00 to 1.00, in intervals of 0.01), and used the resulting set of TPR-FPR coordinate pairs to plot an ROC curve. We repeated this process for each iteration to generate 100 ROC curves, whose coordinates we averaged to produce a “consensus ROC curve” (cROCC) that would reliably report phybLR model performance. One-tailed binomial tests were performed to check whether each phybLR model makes accurate predictions across thresholds in more than 50% and 75% of cases, respectively. A given phybLR model was considered significant if its pooled success rate across classification thresholds was significantly greater than 0.5, and highly significant if its pooled success rate was significantly greater than 0.75 (one-tailed binomial tests, each *p* < 0.05). Such significant test results would imply that the phybLR model tends, across thresholds, to assign higher predicted probabilities to specimens *with* a given phenotype in more than 50% and 75% of cases, respectively. In order to select the most reliable probability classification threshold for each phybLR model, we calculated Youden’s J-statistic (TPR – FPR) for the given model under every classification threshold and selected that threshold with the highest corresponding J-value.

As a final test of model performance, we used the TPR and FPR for each phybLR model at its most accurate classification threshold to perform a second one-tailed binomial test, to check whether that best-performing model-threshold pair made accurate predictions significantly more than 75% of the time.

To obtain another ROC-independent assessment of model performance, we also assessed the “fit” of each tested phybLR model with two R^2^-like metrics suited for binomial logistic regression. There are several R^2^ analogues for assessing the “fit” of logistic regression models, but no clear consensus as to which it preferred [reviewed in Smith et al. 2021].^100^ McFadden’s pseudo-R^2^ is calculated as 1 – logLik(full model)/logLik(null model), where the full model is fit using all model parameters, and the null model is fit using only the intercept term.^101^ McFadden’s pseudo-R^2^ is somewhat analogous to regular R^2^ in that it proxies variance explained by the regression model.^101^ Tjur’s R^2^ (his “coefficient of discrimination”)^102^ is calculated by taking the difference between the mean predicted probabilities for successes and failures.^101^ In our case, for a given phenotype, the Tjur’s R^2^ value corresponds to the difference between the mean predicted probability that specimens have the phenotype and the mean predicted probability that specimens lack the phenotype. Both of these R^2^ analogues range in value from 0 to 1.^101^ To assess model “fit” independently of ROC analysis, we computed McFadden’s pseudo-R^2^ and Tjur’s R^2^ for each trained phybLR model (one for each iteration run) and took the average across runs for each model (Table S5).

Among all significant phybLR models that passed the binomial test requirements above, we selected the most accurate predictor variable (that cROCC with the highest AUC) for each of our limb region proportion, acropodial planform dimension, ungual proportion, and proximal phalanx proportion metrics. Among those, we used AUC values and binomial exact test results to select the single most accurate predictor metric for each binary response variable in our dataset. In this way, we determined which morphometric variables most reliably predicted limb phenotype and aquatic affinity among amniotes for a given region of tree space. Finally, these “best” phybLR models were used to derive a normal distribution of n=10,000 predicted probabilities for each of the tip taxa in our dataset with ambiguous phenotypes. A one-tailed t-test was then used to determine the binary phenotype classifications for all tip taxa. Specifically, a tip taxon was determined to possess the phenotype of interest (*e.g.*, to have flippers) if and only if its mean predicted probability of having the phenotype was significantly higher than the ideal binary classification threshold recovered for the associated cROCC. Internal node phenotypes were then determined using ancestral-state reconstructions.

This phylogenetic machine-learning approach was carried out with five original R functions (phybLR, phycROCCs, filter_bLR_mods, tipPreds, and nodePreds, all detailed in Fig. S2 and Script S1), which themselves made use of previously published functions for fitting phylogenetic logistic regression models, performing statistical tests, plotting ROC curves, and reconstructing ancestral states for internal nodes. Phylogenetic binomial logistic regression curves were fitted using the phyloglm function (boot=10000, btol=35, method = "logistic_MPLE") in the R *phylolm* package (v. 2.6.2^103^). Binomial tests were performed using the binom.test function (x=c(n_successes, n_failures), p=0.5 or 0.75, alternative = "greater") in the R *stats* package (v. 4.2.1).^73^ ROC curve data was generated by either manually calculating TPR and FPR values (as described above) or using the roc function within the R *pROC* package (v. 1.18.0)^104^. AUC was calculated for a given manually generated ROC curve using trapezoidal integration via the trapz function in the R *pracma* package (v. 2.4.4),^105^ and Tjur’s R^2^ calculated using the r2_tjur function within the R *performance* package (v. 0.10.9).^106^ Predicted probabilities for contentious tip taxa were resolved by feeding those taxa into the given phybLR model and predicted probabilities for internal nodes determined by performing ancestral state reconstruction with the asr_independent_contrasts function in the R package *castor* (v. 1.8.0).^107^

#### Testing associations among categorical variables

Associations among all pairs of categorical variables (e.g., forelimb phenotype and aquatic affinity) were assessed by calculating Cramer’s V statistic^108^ and running Fisher’s exact test^109^ on the pair’s associated contingency table. Cramer’s V was calculated using the assocstats function in the *vcd* package (v. 1.4- 11),^110^ and Fisher’s exact test p-value using the chisq.test function in the R *stats* package^73^ (v. 4.2.1; Script S1). Associations among categorical variables were visualized using stacked bar plots to display percentage overlaps among all variable categories (Figs. S7, S18).

#### Data visualization

Primary outputs from statistical analyses were generated in R (v. 4.2.0)^73^ or PAST (v. 4.11)^111^ and compiled into multi-panel figures with BioRender.com under the Student Plan Promo. Morphospaces and thin-plate splines were generated with PAST, ternary plots with the R *Ternary* package (v. 2.1.0),^112^ correlation matrices with the package *corrplot* (v. 0.92),^92^ and all other graph types (box plots, histograms, normal-quantile plots, ROC plots, dot-and-whisker plots, stacked bar plots, etc.) with the R *ggplot2* package (v. 3.4.2).^113^ Taxon silhouettes for main-text and supplementary figures were either designed in Adobe Photoshop 2022 (v. 23.5.1) or downloaded from phylopic.org. For the latter, full image and license attributions are provided in the Supplementary Information (Text S17).

#### Data availability

All of the R scripts, BASH scripts, tree files, and supplementary data tables (including linear and geometric morphometric data, limb phenotype scorings, aquatic affinity scorings, and key phylogenetic comparative test outputs) are available on Dryad [https://doi.org/10.5061/dryad.08kprr5fn]. The full citation for this Dryad dataset is provided below.^114^

