## Supplementary Text and Figures for "Limb proportions predict aquatic habits and soft-tissue flippers in extinct amniotes"

### Supplementary Information
#### for *Limb proportions predict aquatic habits and soft-tissue flippers in extinct amniotes*

#### Table of Contents

| | | |
|---|---|---|
| Text S1. | Overview of supplementary text and materials | 1 |
| Text S2. | Institutional abbreviations in supplementary data tables | 3 |
| Text S3. | Phylogenetic taxonomy and cladistic terminology | 6 |
| Text S4. | Aquatic affinity scoring system and ecotype justifications | 12 |
| Text S5. | How contentious are the aquatic affinities of extinct species? | 14 |
| Text S6. | Previously considered lines of evidence for webbing and flipper form | 18 |
| Text S7. | Geometric morphometric analyses of the acropodium and zeugoautopodium | 19 |
| Text S8. | Sensitivity analysis rationale and results | 21 |
| Text S9. | Pairwise correlations among forelimb, hindlimb, and phalanx proportions | 24 |
| Text S10. | Potential clade-specific osteological correlates of interdigital webbing | 24 |
| Text S11. | Future directions for investigating osteological correlates of interdigital webbing | 25 |
| Text S12. | Additional comments on model predictions for extinct taxa | 26 |
| Text S13. | Implications of soft-tissue flippers for functional ecology in Triassic marine reptiles | 28 |
| Text S14. | Additional comments on temporal patterns of aquatic invasion by extinct amniotes | 29 |
| Text S15. | To what extent are webbing, flippers, and aquatic habits related? | 30 |
| Text S16. | Additional comments on amniote flipper morphometry | 31 |
| Text S17. | Possible developmental factors constraining flipper form in mammals | 32 |
| Text S18. | Note on usage of binomial testing in tandem with ROC analysis | 33 |
| Text S19. | Extended acknowledgments and image attributions | 34 |
| Supplementary Figures | | 41 |
| Supplementary References | | 72 |

#### Text S1. Overview of supplementary text and materials.

This document contains all supplementary text and figures for Gordon et al., "Limb proportions predict aquatic habits and soft-tissue flippers in extinct amniotes." All of the files listed below are accessible on Dryad (Gordon et al. 2025: https://doi.org/10.5061/dryad.08kprr5fn). In the present section, we summarize its contents, and that of the other supplementary tables, scripts, and data files associated with this paper. The supplementary material for this paper includes the following items:

- **Document_S1.pdf:** This is the current document, which contains supplementary text portions S1–S17, and supplementary figures S1–S30. The supplementary text portions go over additional aspects of the introduction, methods, results, and discussion that did not fit in the main text of the manuscript. The supplementary figures likewise cover analyses and results that did not fit in the main text (including outputs from Bland-Altman analyses, phylogenetic-correlation tests, geometric-morphometric analyses, and ROC analysis, among other features).
- The ten data tables referenced in the main-text and supplementary text portions:
  - **Table_S1.csv:** All linear-morphometric, limb-phenotype, and aquatic-affinity data.
  - **Table_S2.csv:** All repeat length measurements and ratios for Bland-Altman analysis.
  - **Table_S3.csv:** The stitched results from all phylogenetic ANOVAs, Levene's, and post-hoc tests (performed in script S2 as described below).
  - **Table_S4.csv:** The stitched results from all phylogenetic-correlation tests (performed in script S3 as described below).
  - **Table_S5.csv:** The stitched results from all phylogenetic binomial-logistic-regression analyses (performed in script S4 as described below).

- **Table_S6.csv:** The combined and processed phylogenetic comparative-test metadata for our sensitivity analyses (performed in script S5 as described below).
- **Table_S7.csv:** All tip-phenotype predictions made by our best-performing phylogenetic Binomial-logistic-regression models (run in script S5 as described below).
- **Table_S8.csv:** All internal-node phenotypes inferred from the tip states in Table S7 (computed in script S5 as described below).
- **Table_S9.csv:** A summary of the geometric-morphometric data generated in script S6.
- **Table_S10.xlsx:** The input data used to generate Figure S17.

- The six R scripts (described in Figure S30) used to run all of our data analyses:
  - **_script_S1.R:** This script begins the analysis by processing the dataset, calibrating all supertrees, completing a few computationally inexpensive analyses, and defining original functions and R-objects required for downstream scripts.
  - **_script_S2.R:** This script performs all phylogenetic ANOVAs, Levene's tests, and post-hoc tests, and generates associated box plots and ternary plots referenced in the paper.
  - **_script_S3.R:** This script performs all the phylogenetic-correlation tests reported here.
  - **_script_S4.7z:** This script fits all the phylogenetic binomial-logistic-regression models referenced in the paper.
  - **_script_S5.R:** This script stitches together the outputs from previous scripts, uses Receiver-Operating Characteristic (ROC) analysis to select the best-performing models and classification thresholds from script S4, and uses those selected model-threshold pairs and ancestral-state estimations to reconstruct the tip- and internal-node phenotypes for all extinct taxa in the dataset. This script also includes a sensitivity analysis to assess the impact of differing tree topologies and data-transformation methods on our results (see Figure S29).
  - **_script_S6.R:** This script contains all of the R code required to perform the geometric-morphometric analyses associated with this study.
  - Please see Figure S30 for more information about how to proceed with these scripts to replicate our data analysis.

- The six supplementary data files, which are compressed zip files, include all the shareable inputs and outputs associated with each of the six supplementary scripts described above.
  - **Data_S1.zip:** All the BASH scripts required for running the R scripts above on a high-performance computing cluster; the Newick files required to generate time-calibrated trees in script S1; and all script S1 outputs, including high-resolution versions (both ultrametric and tip-dated) of the eight supertrees assembled for this project, data-transformation plots, and category-association plots.
  - **Data_S2.zip:** PNGs of all the box plots and associated statistical-test results generated in script S2.
  - **Data_S3.zip:** High-resolution PNGs of the nine correlation matrices generated from the analyses run in job arrays with script S3.
  - **Data_S4.zip:** All script S4 outputs, including unstitched "RData" files, phylogenetic Logistic-regression analysis results, and associated metadata files.
  - **Data_S5.zip:** All script S5 outputs, including assembled consensus ROC curves, best-model prediction histograms, and supertrees with overlain ancestral-state reconstructions.
  - **Data_S6.zip:** Key script S6 inputs and outputs, including TPS files, curveslide.txt, and

PAST .dat files.

If you want to repeat the analyses performed for this paper, you should first review Figure S30, expand Data_S1.zip in a new project folder within a high-performance computing cluster, and then open and begin script S1. Alternatively, if you just want to review our results and conclusions in more detail, the supplementary text and figures below should suffice. If you have any questions about this project, please feel free to contact the lead author (Caleb Gordon) via the following two. Questions about the sensitivity analysis (Text S8) and binomial testing approach (Text S18) should also be directed to the second author, Lisa S. Freisem, at.

The supplementary-text sections below pass roughly through aspects of the main-text Introduction, STARR Methods, Results, and Discussion sections of the paper. Supplementary Text S2 is a list of the institutional abbreviations given in the supplementary tables for this manuscript. Supplementary Text S3 clarifies the meaning of taxon names, and their attendant composition, used throughout the main text and the present supplementary document. Supplementary Text S4 expands on our aquatic-affinity binning system, and supplementary Text S5 briefly summarizes some of the less contentious and more contentious scorings for extinct species in our dataset under this binning system. Supplementary Text S6 clarifies our rationale for investigating particular morphometric features associated with interdigital webbing and flipper form. Supplementary Text S7 provides additional details and comments on the benefits and limitations of our geometric morphometric analyses. Supplementary Text S8 discusses the rationale and results of our sensitivity analysis for the effects of tree topology and transformation method on our phylogenetic comparative tests. Supplementary Text S9 discusses the results of our phylogenetic-correlation matrices (Figs. S19–S27). Supplementary Text S10 and Text S11 cover nuanced morphometric results for webbed taxa that merit further investigation. Supplementary Text S12 includes additional comments on the flipper phenotypes and aquatic affinities predicted for extinct pan-mammals in our dataset, along with certain taxa (i.e., extinct rhynchocephalians) whose phenotypes remained ambiguous following our analysis. Supplementary Text S13 touches on some of the implications for the functional ecology and extinction of Triassic marine reptiles, and Supplementary Text S14 puts our predictions for aquatic habits in extinct species within a broader temporal and geological context. Supplementary Text S15 discusses the results of our category association plots. Supplementary Text S16 provides additional comments on the observed trends in amniote limb morphometry, and supplementary Text S17 provides an extended discussion of possible developmental constraints explaining the reduced morphospace occupancy of pan-mammals. Supplementary Text S18 clarifies our use of binomial testing in tandem with ROC analysis. Finally, supplementary Text S19 provides an extended list of acknowledgments for project funding, collections/specimen access, and image attributions.

### Text S2. Institutional abbreviations in supplementary data tables.

Below, in alphabetical order, are the institutional and field-number abbreviations given for specimens in the supplementary data tables associated with this manuscript.

- **AGB / AGM** – Anhui Geological Museum, Hefei, People's Republic of China
- **AM** – Australian Museum, Sydney, New South Wales, Australia
- **AMB** – field number given for some specimens at the Florida Museum of Natural History in Gainesville, Florida, United States of America (hereafter abbreviated "United States")
- **AMNH** – American Museum of Natural History, New York City, New York, United States
- **BMNH** – British Museum of Natural History, London, England, United Kingdom
- **BMNHC** – Beijing Museum of Natural History, Beijing, People's Republic of China

- **BPI** – Evolutionary Studies Institute (formerly Bernard Price Institute) for Palaeontological Research, University of the Witwatersrand, Johannesburg, South Africa
- **BSP / BSPG** – Bayerische Staatsammlung für Paläontologie und Geologie [Bavarian State Collection for Palaeontology and Geology], Munich, Bavaria, Germany
- **CM** – Carnegie Museum of Natural History, Pittsburgh, Pennsylvania, United States
- **CMN NUFV** – Canadian Museum of Nature, Palaeobiology Section (Nunavut Fossil Vertebrates), Ottawa, Ontario, Canada
- **DINO** – Dinosaur National Monument, Utah and Colorado, United States
- **DLC** – Duke Lemur Center Museum of Natural History, North Carolina, United States
- **DPC OST** – Duke Lemur Center Museum of Natural History (alternative specimen identification prefix), North Carolina, United States
- **DU EA** – Duke University Evolutionary Anthropology Dept., North Carolina, United States
- **FHSM** – Fort Hayes State University's Sternberg Museum of Natural History, Hays, Kansas, United States
- **FLMNH** – Florida Museum of Natural History, Gainesville, Florida, United States
- **FMNH** – Field Museum of Natural History, Chicago, Illinois, United States
- **FSAC** – Faculté des Sciences Aïn Chock, University Hassan II, Casablanca, Morocco
- **GPIT** – Geologisch-Paläontologisches Institut Tübingen, Germany
- **GMPKU** – Geological Museum of Peking University, Peking, People's Republic of China
- **GZDB** – Guizhou Geological Museum, Guiyang, Guizhou, People's Republic of China
- **HPM** – Hrvatski Prirodo-slovni Muzej, Zagreb, Croatia
- **HUJ** – Hebrew University of Jerusalem, Israel
- **IGM** – Instituto Nacional de Investigaciones Geológico-Mineras, Bogotá, Colombia
- **IGPS** – Institute of Geology and Paleontology, Tohoku University, Sendai, Japan
- **IVPP** – Institute of Vertebrate Paleontology and Paleoanthropology, Beijing, People's Republic of China
- **JME** – Jura Museum Eichstätt, Eichstätt, Germany
- **JMP** – Jinzhou Museum of Paleontology, Jinzhou, Liaoning, People's Republic of China
- **KUVP** – Kansas University Biodiversity Institute and Natural History Museum, Division of Vertebrate Paleontology, Lawrence, Kansas, United States
- **LACM** – Natural History Museum of Los Angeles County, Los Angeles, California, United States
- **LSUMZ** – Louisiana State University Museum of Natural History, Baton Rouge, Louisiana, United States
- **MB.R** – specimen identification prefix for some specimens from the Museum für Naturkunde Berlin in Berlin, Germany
- **MCSNB** – Museo Civico di Scienze Naturali Bergamo, Bergamo, Italy
- **MCZ** – Museum of Comparative Zoology, Cambridge, Massachusetts, United States
- **MFSN** – Museo Friulano Di Storia Naturale, Udine, Italy
- **MGP** – specimen identification prefix for sauropterygians from the Museum of Nature and Humankind of Padova (MNH), Padua, Veneto, Italy
- **MNHN** – Muséum National d'Histoire Naturelle, Paris, France
- **MNN** – Musée National du Niger, Niamey, Niger
- **MOR** – Museum of the Rockies, Bozeman, Montana, United States

- **MPC** – Institute of Paleontology and Geology, Mongolian Academy of Sciences, Ulanbatar, Mongolia
- **MSNM** – Museo di Storia Naturale di Milano Sezione di Paleontologia, Milan, Italy
- **MVZ** – Museum of Vertebrate Zoology, Berkeley, California, United States
- **MZSP** – Museum of Zoology of the University of São Paulo, Brazil
- **NCSM** – North Carolina Museum of Natural Sciences, Raleigh, North Carolina, United States
- **NHMUK** – Natural History Museum, London, England, United Kingdom
- **NMNS** – National Museum of Natural Science, Taichung, Taiwan, People's Republic of China
- **NMW** – Naturhistorisches Museum Wien, Vienna, Austria
- **NPS** – National Park Service prefix for specimens from Glacier Bay National Park, Alaska, United States
- **NYF** – Yuan'an Geological Museum, Yuan'an County, Yichang, Hubei, People's Republic of China
- **OUVC** – Ohio University Vertebrate Collection, Athens, Ohio, United States
- **OUMNH** – Oxford University Museum of Natural History, England, United Kingdom
- **PIMUZ** – Paläontologisches Institut Universitüat Zürich, Switzerland
- **PMOL** – Palaeontological Museum of Liaoning, Yizhou, People's Republic of China
- **PMU** – Paleontologiska Museet, Uppsala Universitet, Uppsala, Sweden
- **PVL** – Paleontología de Vertebrados, Instituto 'Miguel Lillo', San Miguel de Tucumán, Argentina
- **PVSJ** – Museo de Ciencias Naturales, Universidad Nacional de San Juan, San Juan, Argentina
- **PZO** – Museo Archeologico dell'Alto Adige, Bolzano (Bozen), Italy
- **RC** – Former specimen identification prefix for some specimens from the Instituto Nacional de Investigaciones Geológico-Mineras (IGM), Bogotá, Colombia
- **RVC** – Royal Veterinary College, University of London, England, United Kingdom
- **SAM** – Iziko South African Museum, Cape Town, South Africa
- **SDSNH** – San Diego Natural History Museum, San Diego, California, United States
- **SMF** – Senckenberg Museum Frankfurt, Frankfurt, Germany
- **SMC** – Save the Manatee Club, Longwood, Florida, United States
- **SMNK** – Staatliches Museum für Naturkunde Karlsruhe, Karlsruhe, Baden-Württemberg, Germany
- **SMNS** – Staatliches Museum für Naturkunde Stuttgart, Stuttgart, Germany
- **SNHM** – Shanghai Natural History Museum, Shanghai, People's Republic of China
- **SSSC** – Sitka Sound Science Center, Sitka, Alaska, United States
- **TCWC** – Texas A&M University Biodiversity Research and Teaching Collections, College Station, Texas, United States
- **TMAG** – Tasmanian Museum and Art Gallery, Hobart, Australia
- **TMP** – Royal Tyrrell Museum of Palaeontology, Drumheller, Alberta, Canada
- **UCM** – University of Colorado Museum of Natural History, Boulder, Colorado, United States
- **UCRC** – University of Chicago Research Collection, Chicago, Illinois, United States
- **UFSCar** – The São Carlos Federal University Advanced Materials and Energy Research Center, São Carlos, Brazil
- **UMZC** – University of Cambridge Museum of Zoology, Cambridge, England, United Kingdom
- **UNIMORE** – Collezioni del Polo Museale, Università di Modena e Reggio Emilia, Modena, Italy
- **UPI** – Uppsala University Paleontological Institute, Sweden
- **USNM** – National Museum of Natural History, Washington, DC, United States of America
- **WGSC** – Wuhan Centre of China Geological Survey, Wuhan, China
- **WIGM** – Wuhan Institute of Geology and Mineral Resources, Wuhan, People's Republic of China

- **XNGM** – Xingyi National Geopark Museum, Xingyi City, Guizhou Province, People's Republic of China
- **YPM** – Yale Peabody Museum, New Haven, Connecticut, United States
- **ZMB** – specimen identification prefix for some specimens from the Museum für Naturkunde Berlin, Germany
- **ZMNH** – Zhejiang Museum of Natural History, Hangzhou, Zhejiang, People's Republic of China

### Text S3. Phylogenetic taxonomy and cladistic terminology.

This dissertation employs a phylogenetic taxonomy following the rules and recommendations given in the International Code of Phylogenetic Nomenclature (PhyloCode), Version 6 (Cantino & de Queiroz 2020) and the associated companion volume, *Phylonyms* (de Queiroz et al. 2020). As per PhyloCode guidelines, all clade names are italicized. For more information on PhyloCode rules, recommendations, and conceptual definitions (e.g., of "crown" vs. "total" clades), please refer to the PhyloCode (Cantino & de Queiroz 2020).

Below, we list the clade names used in the main- and supplementary texts in this study, with brief descriptions of their composition, and the proper attributions to their phylogenetic definitions to facilitate the interpretation of our results and give due credit to previous researchers. Following PhyloCode guidelines, in-text attribution immediately following the clade name is for the author(s) who coined the clade name (nominal author[s]), and the later parenthetical attribution is for the author(s) who explicitly gave the clade name a phylogenetic definition (definitional author[s]), but we provide these names below in a more tabular format for clarity. All authors are cited in the Supplementary References section near the end of this document. Clade names below are given alphabetically (rather than systematically). Genus- and species-level clades are not formally defined but instead given in-text nominal author citations. "Definitional" and "Definition" are placed in quotations because not all the clade names below have been explicitly defined with reference to specifier species and assigned formal PhyloCode registration numbers.

| Clade Name | Nominal Author(s) | "Definitional" Author(s) | Phylogenetic "Definition" (and notes, if applicable) |
|---|---|---|---|
| ***Afrotheria*** | M. J. Stanhope et al. 1998 | Stanhope et al. 1998 | *Afrotheria* is the crown clade stemming from the most recent common ancestor of tenrecs, golden moles, elephants, sirenians, hyracoids, aardvark, and elephant shrews. |
| ***Alligatoridae*** | G. Cuvier 1807 | Norell et al. 1994; Rio & Mannion 2021 | *Alligatoridae* is the crown clade stemming from the most recent common ancestor of *Alligator mississippiensis* (originally "*Crocodilus mississipiensis*" F. M. Daudin 1802) and *Caiman crocodilus* C. Linnaeus 1758. |
| ***Alligatoroidea*** | J. E. Gray 1844 | Norell et al. 1994; Rio & Mannion 2021 | *Alligatoroidea* is the total clade including *Alligatoridae* and all crocodylians more closely related to it than to *Crocodylus niloticus* J. N. Laurenti 1768 and *Gavialis gangeticus* J. F. Gmelin 1789. |
| ***Amniota*** | E. Haeckel 1866 | Laurin & Reisz 2020a | *Amniota* is the crown clade stemming from the most recent common ancestor of mammals (e.g., *Homo sapiens* C. Linnaeus 1758) and reptiles (e.g., *Gallus gallus* C. Linnaeus 1758). |

| | | | |
|---|---|---|---|
| ***Araeoscelidia*** | S. W. Williston 1913 | Laurin 1991; Laurin & Reisz 1995 | *Araeoscelidia* is the clade stemming from the most recent common ancestor of *Araeoscelis* S. W. Williston 1910 and *Petrolacosaurus* H. H. Lane 1945. |
| ***Archosauria*** | E. D. Cope 1869b | Gauthier & Padian 2020 | *Archosauria* is the crown clade stemming from the most recent common ancestor of birds (e.g., *Gallus gallus* C. Linnaeus 1758) and crocodylians (e.g., *Alligator mississippiensis* [originally "*Crocodilus mississipiensis*" F. M. Daudin 1802]). |
| ***Archosauromorpha*** | F. von Huene 1946 | Gauthier 2020a | *Archosauromorpha* is the clade stemming from the most recent common ancestor of birds (e.g., *Gallus gallus* C. Linnaeus 1758), crocodylians (e.g., *Alligator mississippiensis* [originally "*Crocodilus mississipiensis*" F. M. Daudin 1802]), *Mesosuchus browni* D. M. S. Watson 1912, *Trilophosaurus buettneri* E. C. Case 1928, *Prolacerta broomi* F. R. Parrington 1935, and *Protorosaurus speneri* H. Von Meyer 1832. |
| ***Artiodactyla*** | R. Owen 1848 | Spaulding et al. 2009 | *Artiodactyla* is the crown clade stemming from the most recent common ancestor of extant cattle, antelope, deer, giraffes, musk deer, chevrotains, hippos, pigs, peccaries, and camels. |
| ***Askeptosauroidea*** | E. Kuhn-Schnyder 1971 | Müller 2005, with tip taxa updated to include *M. brevis* | The clade stemming from the most recent common ancestor of *Askeptosaurus italicus* F. V. Nopcsa 1925, *Anshunsaurus huangguoshuensis* J. Liu 1999, *Endennasaurus acutirostris* S. Renesto 1984, and *Miodentosaurus brevis* Y. N. Cheng et al. 2007. |
| ***Atlantogenata*** | P. J. Waddell et al. 1999 | Waddell et al. 1999 | *Atlantogenata* is the crown clade stemming from the most recent common ancestor of *Afrotheria* (defined above) and *Xenarthra* (defined below). |
| ***Captorhinidae*** | E. C. Case 1911 | Laurin & Reisz 1995 | *Captorhinidae* is the clade stemming from the most recent common ancestor of *Captorhinus angusticeps* E. D. Cope 1895, *Labidosaurus hamatus* E. D. Cope 1896, and *Romeria texana* L. I. Price 1937. |
| ***Carnivora*** | T. E. Bowdich 1821 | Flynn et al. 2020 | *Carnivora* is the crown clade stemming from the most recent common ancestor of *Panthera leo* C. Linnaeus 1758, *Canis lupus* C. Linnaeus 1758, and *Phoca vitulina* C. Linnaeus 1758. |
| ***Cetacea*** | M. Brisson 1756 | Deméré 2020 | *Cetacea* is the crown clade stemming from the most recent common ancestor of odontocetes (e.g., *Tursiops truncatus* G. Montagu 1821) and mysticetes (e.g., *Balaena mysticetus* C. Linnaeus 1758). |
| ***Crocodylia*** | J. F. Gmelin 1789 | Benton & Clark 1988; Rio & Mannion 2021 | *Crocodylia* is the crown clade stemming from the most recent common ancestor of *Alligator mississippiensis* (originally "*Crocodilus mississipiensis*" F. M. Daudin 1802), *Crocodylus niloticus* J. N. Laurenti 1768, and *Gavialis gangeticus* J. F. Gmelin 1789. |
| ***Crocodylidae*** | G. Cuvier 1807 | Brochu 1999 | *Crocodylidae* is the crown clade stemming from the most recent common ancestor of *Crocodylus niloticus* J. N. Laurenti 1768, *Osteolaemus tetraspis* E. D. Cope 1860, and *Tomistoma schlegelli* (originally "*Cr. Schlegelii*" S. Müller 1838). |

| | | | |
|---|---|---|---|
| ***Dolichosauridae*** | P. Gervais 1852 | Augusta et al. 2022 | *Dolichosauridae* is the clade stemming from the most recent common ancestor of *Dolichosaurus longicollis* R. Owen 1850, *Aphanizocnemus libanensis* C. Dal Sasso & G. Pinna 1997, *Pontosaurus lesinensis* A. Kornhuber 1873, and *Adriosaurus suessi* H. G. Seeley 1881. |
| ***Eosauropterygia*** | O. Rieppel 1994 | Rieppel 1994, with tip taxa simplified and revised to avoid referencing "Pachy-pleurosauroidea" | *Eosauropterygia* is the clade stemming from the most recent common ancestor of *Corosaurus alcovensis* E. C. Case 1936, *Neusticosaurus* (originally "*Pachypleurosaura*) *edwardsi* E. Cornalia 1854), *Lariosaurus balsami* G. Curioni 1847 (*sensu* Rieppel 1998), *Pistosaurus longaevus* H. Von Meyer 1839, and *Plesiosaurus dolichodeirus* W. D. Conybeare 1824. |
| ***Euarchontoglires*** | W. J. Murphy et al. 2001 | Benton et al. 2015 (with precise tip taxa selected here arbitrarily) | *Euarchontoglires* is the clade stemming from the most recent common ancestor of archontans (e.g., *Homo sapiens* C. Linnaeus 1758) and glires [e.g., *Rattus norvegicus* (originally "*Mus Norvegicus"*) J. Berkenhout 1769]. |
| ***Ferae*** | C. Linnaeus 1758 | Asher et al. 2009 | *Ferae* is the clade stemming from the most recent common ancestor of *Carnivora* (e.g., *Canis lupus* C. Linnaeus 1758) and pangolins (e.g., *Manis pentadactyla* C. Linnaeus 1758). |
| ***Hupehsuchia*** | R. L. Carroll & Z.-M. Dong 1991 | Inferred from Motani et al. 2015 | *Hupehsuchia* is the clade containing all taxa more closely related to *Hupehsuchus nanchangensis* C. C. Young & Z.-M. Dong 1972 than to *Ichthyosaurus communis* H. T. De la Beche & W. D. Conybeare 1821. |
| ***Ichthyosauriformes*** | R. Motani et al. 2015 | Motani et al. 2015 | *Ichthyosauriformes* is the clade containing all taxa more closely related to *Ichthyosaurus communis* H. T. De la Beche & W. D. Conybeare 1821 than to *Hupehsuchus nanchangensis* C. C. Young & Z.-M. Dong 1972. |
| ***Ichthyopterygia*** | R. Owen 1859 | Motani 1999 | *Ichthyopterygia* is the clade stemming from the most recent common ancestor of *Ichthyosaurus communis* H. T. De la Beche & W. D. Conybeare 1821, *Utatsusaurus hataii* T. Shikama and T. Kamei 1978, and *Parvinatator wapitiensis* E. L. Nicholls & D. B. Brinkman 1995. |
| ***Ichthyosauromorpha*** | R. Motani et al. 2015 | Motani et al. 2015 | *Ichthyosauromorpha* is the clade stemming from the most recent common ancestor of *Ichthyosaurus communis* H. T. De la Beche & W. D. Conybeare 1821 and *Hupehsuchus nanchangensis* C. C. Young & Z.-M. Dong 1972. |
| ***Lepidosauria*** | E. Haeckel 1866 | Gauthier et al. 1988a; De Queiroz & Gauthier 2020a | *Lepidosauria* is clade stemming from the most recent common ancestor of *Squamata* (defined below; e.g., *Lacerta agilis* C. Linnaeus 1758) and *Sphenodon* (originally "*Hatteria*") *punctatus* J. E. Gray 1842. |
| ***Lepidosauromorpha*** | J. A. Gauthier et al. 1988b | Branch-based definition: Gauthier et al. 1988b; Gauthier & de Queiroz 2020b | *Lepidosauromorpha* is used here as the clade containing all species more closely related to *Lepidosauria* (e.g., *Lacerta agilis* C. Linnaeus 1758) than to *Archosauria* (e.g., *Gallus gallus* C. Linnaeus 1758). This name is used here as a synonym for *Pan-Lepidosauria* (defined below). An alternative, node-based definition would be "the clade stemming from the most recent common ancestor of *Lacerta agilis* C. |

| | | | |
|---|---|---|---|
| | | | Linnaeus 1758, *Sphenodon* (originally "*Hatteria*") *punctatus* J. E. Gray 1842, and the extinct species *Megachirella wachtleri* S. Renesto & R. Posenato 2003, *Sophineta cracoviensis* S. E. Evans & M. Borsuk-Białynicka 2009, *Marmoretta oxoniensis* S. E. Evans 2008," which are consistently recovered as being closer to crown lepidosaurs than they are to archosaurs (Gauthier & de Queiroz 2020b). |
| ***Mammalia*** | C. Linnaeus 1758 | Rowe 2020a | *Mammalia* is the crown clade stemming from the most recent common ancestor of monotremes (e.g., *Tachyglossus aculeatus* G. Shaw 1792), marsupials (e.g., *Didelphis marsupialis* C. Linnaeus 1758), and placentals (e.g., *Homo sapiens* C. Linnaeus 1758). |
| ***Mesosauridae* (=*Mesosaurus tenuidens*?)** | G. Baur 1889 | Laurin & Reisz 1995 | *Mesosauridae* is the clade stemming from the most recent common ancestor of the putative mesosaur taxa *Mesosaurus tenuidens* P. Gervais 1865*, Brazilosaurus sanpauloensis* T. Shikama & H. Ozaki 1966, and *Stereosternum tumidum* E. D. Cope 1886. Recent morphological and morphometric work has synonymized these three species, suggesting that the clade is monospecific (Piñeiro et al. 2021; Verrière & Fröbisch 2022). Wherever possible, we refer to the constituents of this clade as "mesosaurs" to remain agnostic about this taxonomic distinction. |
| ***Mosasauria*** | T. H. Huxley 1871 | Augusta et al. 2022 | *Mosasauria* is the clade stemming from the most recent common ancestor of *Dolichosauridae* (e.g., *Dolichosaurus longicollis* R. Owen 1850), defined above, and *Mosasauridae* (e.g., *Mosasaurus hoffmanni* G. Mantell 1829). Herein, we call members of *Mosasauria* "mosasaurians," and refer to derived mosasaurians within *Mosasauridae* as "mosasaurids." |
| ***Mosasauridae*** | P. Gervais 1852 | Madzia & Conrad 2020 | *Mosasauridae* is the clade stemming from the most recent common ancestor of *Mosasaurus hoffmanni* G. Mantell 1829, *Tylosaurus* (originally "*Macrosaurus*") *proriger* E. D. Cope 1869a, and *Halisaurus platyspondylus* O. C. Marsh 1869. |
| ***Mosasauroidea*** | C. L. Camp 1923 | Madzia & Conrad 2020; Augusta et al. 2022, with reference tip taxa specified | *Mosasauroidea* is the clade stemming from the most recent common ancestor of *Aigialosaurus dalmaticus* D. Gorjanović-Kramberger 1892 and *Mosasaurus hoffmanni* G. Mantell 1829. Augusta et al. (2022) define *Mosasauroidea* as "the minimally inclusive clade including aigialosaurids and mosasaurids." We follow this definition but refer to a particular aigialosaurid specifier species given the uncertainty of aigialosaurid monophyly. |
| ***Omnisauropterygia*** | New taxon | This study | *Omnisauropterygia* is the largest clade that includes *Placodus gigas* L. Agassiz 1833 and *Plesiosaurus dolichodeirus* W. D. Conybeare 1824 but none of *Ichthyosaurus communis* H. T. De la Beche & W. D. Conybeare 1821, *Thalattosaurus alexandrae* J. C. Merriam 1904, *Gallus gallus* C. Linnaeus 1758, *Alligator* [originally "*Crocodilus*"] *mississippiensis* F. M. Daudin 1802, *Lacerta agilis* C. Linnaeus 1758, or *Sphenodon punctatus* J. E. Gray 1842. Said more briefly, *Omnisauropterygia* is the largest clade that includes *Sauropterygia* but none of *Archosauria*, *Lepidosauria*, *Ichthyopterygia,* or *Thalattosauria*. The clade *Omnisauropterygia* may be coextensive with *Sauropterygomorpha* or *Sauropterygiformes* (defined below), but differs from these latter clade names in being branch-based, and therefore obtaining regardless of whether *Prosaurosphargis* |

| | | | |
|---|---|---|---|
| | | | *yingzishanensis* A. Wolniewicz et al. 2023, *Eusaurosphargis dalsassoi* S. Nosotti & O. Rieppel 2003, *Sinosaurosphargi yunguiensis* C. Li et al. 2011, *Hanosaurus hupehensis* C. C. Young 1972, or another close relative of *Sauropterygia* is recovered as basal to the others. As far as we know, this clade is unnamed, and we name it here for lack of an alternative option that remains valid across all the tree topologies considered in this study. |
| ***Pan-Archosauria*** | J. A. Gauthier 2020b | Gauthier 1984; Gauthier 2020b | *Pan-Archosauria* is the clade containing *Archosauria* (defined above; e.g., *Gallus gallus* C. Linnaeus 1758) and all taxa more closely related to it than they are to *Lepidosauria* (defined above; e.g., *Lacerta agilis* C. Linnaeus 1758). |
| ***Pan-Lepidosauria*** | J. A. Gauthier & K. de Queiroz 2020a | Gauthier & de Queiroz 2020a | *Pan-Lepidosauria* is the clade containing *Lepidosauria* (defined above; e.g., *Lacerta agilis* C. Linnaeus 1758) and all taxa more closely related to it than they are to *Archosauria* (defined above; e.g., *Gallus gallus* C. Linnaeus 1758). This name has precedence over its synonym: *Lepidosauromorpha* (defined above). |
| ***Pan-Mammalia*** | T. B. Rowe 2004 | Rowe 2020b | *Pan-Mammalia* is the clade containing *Mammalia* (defined above; e.g., *Homo sapiens* C. Linnaeus 1758) and all taxa more closely related to it than they are to *Reptilia* (defined below; e.g., *Lacerta agilis* C. Linnaeus 1758). |
| ***Pan-Reptilia*** | G. S. Bever et al. 2015 | Gauthier in Gordon 2025 | *Pan-Reptilia* is the clade containing *Reptilia* (defined above; e.g., *Lacerta agilis* C. Linnaeus 1758) and all taxa more closely related to it than they are to *Mammalia* (defined above; e.g., *Homo sapiens* C. Linnaeus 1758). |
| ***Pistosauria*** | O. Rieppel 1998 | Rieppel 1998 | *Pistosauria* is the clade stemming from the most recent common ancestor of *Cymatosaurus fridericianus* K. Von Fritsch 1894, *Pistosaurus longaevus* H. Von Meyer 1839, and *Plesiosaurus dolichodeirus* W. D. Conybeare 1824. |
| ***Placodontiformes*** | J. M. Neenan et al. 2013 | Inferred from Neenan et al. 2013 | *Placodontiformes* is the clade containing *Placodus gigas* L. Agassiz 1833 and all taxa more closely related to it than they are to *Eosauropterygia* (defined above). |
| ***Plesiosauria*** | De Blainville 1835 | Ketchum & Benson 2010 | *Plesiosauria* is the clade containing all taxa more closely related to *Plesiosaurus dolichodeirus* W. D. Conybeare 1824 than they are to *Augustasaurus hagdorni* P. M. Sander et al. 1997. |
| ***Reptilia*** | C. Linnaeus 1758 | Laurin & Reisz 2020b | The crown clade stemming from the most recent common ancestor of *Testudo graeca* C. Linnaeus 1758, *Iguana iguana* C. Linnaeus 1758, and *Crocodylus* (originally "*Lacerta"*) *niloticus* J. N. Laurenti 1768. |
| ***Sauria*** | J. Macartney 1802 | Gauthier & de Queiroz 2020b, here revised to avoid excluding turtles | The crown clade stemming from the most recent common ancestor of *Archosauria* [defined above; e.g., *Alligator* (= "*Crocodilus*") *mississippiensis* F. M. Daudin 1802] and *Lepidosauria* [defined above; e.g., *Sphenodon* (= "*Hatteria*") *punctatus* J. E. Gray 1842]. |
| ***Sauropterygia*** | R. Owen 1859 | Rieppel 1994 | The clade stemming from the most recent common ancestor of *Placodontiformes* (defined above; e.g., *Placodus gigas* L. Agassiz 1833) and *Eosauropterygia* (which is itself defined above as the |

| | | | |
|---|---|---|---|
| | | | most recent common ancestor of *Corosaurus alcovensis* E. C. Case 1936, *Neusticosaurus* [originally "*Pachypleurosaura"*] *edwardsi* E. Cornalia 1854, *Lariosaurus balsami* G. Curioni 1847, *Pistosaurus longaevus* H. Von Meyer 1839, and *Plesiosaurus dolichodeirus* W. D. Conybeare 1824). |
| ***Sauropterygiformes*** | W. Wang et al. 2022 | Wang et al. 2022, with reference tip taxa revised for clarity | Wang et al. define *Sauropterygiformes* as "a clade including all the eosauropterygians, placodontiforms, saurosphargids, *Helveticosaurus*, and *Atopodentatus*." Given the disputed monophyly of saurosphargids and the general instability of basal sauropterygiform interrelations, we revise this definition slightly to refer to explicit tip taxa as specifiers. Thus, we define *Sauropterygiformes* as the smallest clade stemming from the most recent common ancestor of *Corosaurus alcovensis* E. C. Case 1936, *Neusticosaurus* (originally "*Pachypleurosaura*) *edwardsi* E. Cornalia 1854), *Lariosaurus balsami* G. Curioni 1847 (*sensu* Rieppel 1998), *Pistosaurus longaevus* H. Von Meyer 1839, and *Plesiosaurus dolichodeirus* W. D. Conybeare 1824, *Placodus gigas* L. Agassiz 1833, *Eusaurosphargis dalsassoi* S. Nosotti & O. Rieppel 2003, *Sinosaurosphargi yunguiensis* C. Li et al. 2011, and *Hanosaurus hupehensis* C. C. Young 1972. |
| ***Sauropterygomorpha*** | Wolniewicz et al. 2023 | Wolniewicz et al. 2023 | *Sauropterygomorpha* is the clade stemming from the most recent common ancestor of *Eusaurosphargis dalsassoi* S. Nosotti & O. Rieppel 2003 and *Eosauropterygia* (which is itself defined above as the most recent common ancestor of *Corosaurus alcovensis* E. C. Case 1936, *Neusticosaurus* [originally "*Pachypleurosaura"*] *edwardsi* E. Cornalia 1854, *Lariosaurus balsami* G. Curioni 1847, *Pistosaurus longaevus* H. Von Meyer 1839, and *Plesiosaurus dolichodeirus* W. D. Conybeare 1824). |
| ***Saurosphargidae*** | C. Li et al. 2011 | Li et al. 2014 | *Saurosphargidae* is the clade stemming from the most recent common ancestor of *Saurosphargis volzi* F. von Huene 1936, *Sinosaurosphargis yunguiensis* C. Li et al. 2011, and *Largocephalosaurus qianensis* C. Li et al. 2014. |
| ***Squamata*** | M. Oppel 1811 | De Queiroz & Gauthier 2020b | *Squamata* is the largest crown clade containing *Lacerta agilis* C. Linnaeus 1758 but not *Sphenodon* (originally "*Hatteria*") *punctatus* J. E. Gray 1842. |
| ***Tangasaurinae*** | J. Piveteau 1926 | Buffa et al. 2025 | *Tangasaurinae* is the clade stemming from the most recent common ancestor of *Tangasaurus mennelli* S. H. Haughton 1924 and *Hovasaurus boulei* J. Piveteau 1926. |
| ***Testudines*** | A. J. G. C. Batsch 1788 | Joyce et al. 2020 | *Testudines* is the smallest crown clade containing the pleurodire *Chelus* (originally "*Testudo*") *fimbriatus* J. G. Schneider 1783, the trionychian *Trionyx* (originally "*Testudo*") *triunguis* P. Forskål 1775, and the chelonioid *Chelonia* (originally "*Testudo*") *mydas* C. Linnaeus 1758. |
| ***Thalattosauria*** | J. C. Merriam 1904 | Inferred from Chai et al. 2023, with tip | *Thalattosauria* is the clade stemming from the most recent common ancestor of *Thalattosauroidea* (defined below; e.g., *Thalattosaurus* |

| | | | |
|---|---|---|---|
| | | taxa specified for clarity | *alexandrae* J. C. Merriam 1904) and *Askeptosauroidea* (defined above; e.g., *Askeptosaurus italicus* F. V. Nopcsa 1925). |
| ***Thalattosauroidea*** | F. V. Nopcsa 1928 | Inferred from Druckenmiller et al. 2020 and Chai et al. 2023, with clarified specifiers | *Thalattosauroidea* is the clade stemming from the most recent common ancestor of *Thalattosaurus alexandrae* J. C. Merriam 1904, *Clarazia schinzi* B. Peyer 1936, and *Gunakadeit joseeae* P. S. Druckenmiller et al. 2020. |
| ***Ungulata*** | C. Linnaeus 1766 | Archibald 2020 | The crown clade stemming from the most recent common ancestor of artiodactyls (see *Artiodactyla* defined above; e.g., *Bos taurus* C. Linnaeus 1758) and perissodactyls (e.g., *Equus caballus* C. Linnaeus 1758). |
| ***Xenarthra*** | E. D. Cope 1889 | Shockey 2020 | The crown clade stemming from the most recent common ancestor of armadillos (e.g., *Dasypus novemcinctus* C. Linnaeus 1758), sloths (e.g., *Choloepus didactylus* C. Linnaeus 1758), and anteaters (e.g., *Myrmecophaga tridactyla* C. Linnaeus 1758). |

**Text S4. Aquatic affinity scoring system and ecotype justifications.**

The aquatic habits of amniotes exist on a continuous and multidimensional spectrum, and any effort to discretize them is inevitably subjective. All we can hope to do is bin species consistently and transparently, in a way that captures broad differences between "more" and "less" aquatic habits, and makes these differences discrete and applicable across amniotes, despite their diverse and nuanced ecologies. To do this, we settled on an original scoring system (Figure 1a; Figure S5) that separates species into five aquatic affinity "guilds", where a "guild" is defined as a group of species that exploit the same class of environmental resources in the same way, without regard to their phylogenetic positions (Root 1967). Guild groupings are commonly used to classify marine reptiles—most notably into "feeding guilds" based on tooth morphology (Massare 1987; Foffa et al. 2018; Fischer et al. 2022). For this study, we developed a new guild system under which we classified all sampled specimens into one of five aquatic affinity guilds (fully terrestrial, transiently aquatic, moderately aquatic, highly aquatic, and fully aquatic) based on the relative amount of time they typically spend on (and the relative portion of niche space in which they use) land vs. water:

- **Fully terrestrial** species spend nearly all of their time on land, seldom intentionally crossing water, and include most desert-dwelling lizards and mammals, as well as tortoises and songbirds.

- **Transiently aquatic** species spend the vast majority of their time on land, but routinely make transient forays into the water to escape predators, find new suitable terrestrial habitats, or snag the occasional subaqueous morsel. Water anoles and marsh rabbits fall into this category, but so do more stereotypically terrestrial megafauna that make rare but extensive marine journeys, such as wild boars, which have been known to swim on occasion from England to France (see supplementary tables within Table S1).

- **Moderately aquatic** species are amphibious, spending roughly equal amounts of time on land and in the water—regularly crossing and residing in both media. Platypuses, many otters, beavers, some aquatic lizards, and most waterfowl and shorebirds fall into this category.

- **Highly aquatic** species spend the vast majority of their time in the water but routinely bask or nest for short periods on land. Extant sea turtles and sea lions fall into this category.

- Finally, **fully aquatic** species never intentionally leave the water, and include extant sirenians and cetaceans, along with some extant viviparous sea snakes and many extinct marine reptiles (such as plesiosaurs and ichthyosaurs).

These bins together form a five-rung scale that stretches the terrestrial-aquatic continuum and allows us to apply the same classification system to all amniotes. Most of these aquatic-affinity bins for extant taxa are intuitive, with short, literature-based justifications provided in the supplementary tables within Table S1. However, scoring the aquatic affinities of extant *crocodilians* was particularly challenging, and our crocodilian ecotype scores require further clarification. All extant crocodilians are amphibious predators that spend extensive time hunting and wading in the water, but also routinely bask, nest, and travel to some degree on land (Whitaker & Basu 1982; Ferguson 1985; Vitt & Caldwell 2014; Murray et al. 2020). These species nevertheless span a continuum of semi-aquatic lifestyles, which differ intraspecifically, ontogenetically, and depending on weather and other external factors (Vitt & Caldwell 2014; Isberg et al. 2019). In order to bin crocodilians by their aquatic habits in an analysis that could plausibly encompass all other amniotes, we inevitably oversimplify the complex behavioral repertoire of this clade. We try only to score them in a consistent way that reflects their ecological preferences and permits both inter- and intra-clade comparisons. To that end, we use this supplementary text to describe our reasoning for these scores.

Briefly, we call *Gavialis gangeticus* J. F. Gmelin 1789 and *Crocodylus acutus* G. Cuvier 1807 "highly aquatic," and all other crown crocodilian species "moderately aquatic." Perhaps because they descend from marine ancestors (Wilberg et al. 2019), the gavialids *G. gangeticus* and *Tomistoma schlegelli* S. Müller 1838 have longirostrine snouts often treated as specializations for hunting fish, though their diet is actually more varied (Whitaker & Basu 1982). While *T. schlegelli* occupies lentic marsh waters (Staniewicz et al. 2018), *G. gangeticus* often resides in deep, fast-moving rivers, seldom (if ever) moves far from the water's edge, often remains submerged during nest attendance, and has more limited mobility on land than other modern crocodilians (Whitaker & Basu 1982; Vitt & Caldwell 2014; Choudhary et al. 2018; Murray et al. 2020). As a result, *G. gangeticus* is frequently considered the most aquatic crown crocodilian (Vitt & Caldwell 2014). In a similar vein, the saltwater crocodile (*Crocodylus porosus* J. G. Schneider 1801) may be the most active long-distance swimmer among extant crocodilians, as it regularly makes extended (>1000 km) journeys over open ocean (Fukuda et al. 2019; Spennemann 2020). Because *G. gangeticus* and *C. acutus* spend most of their time in large or fast-moving water bodies, but still occasionally travel over land, we consider these species "highly aquatic." Given that no other extant species of crocodilian typically spends this much time active in large or fast-moving bodies of water, we consider all other crown crocodilians—including alligatorids, other crocodylids, and *Tomistoma schlegelli*—to be "moderately aquatic." This scoring reflects the fact that these species spend extensive time on both land and (typically shallower, more lentic) water. This "moderately aquatic" designation does not distinguish between time spent being "active" (e.g., hunting) vs. "passive" (e.g., basking) on land vs. water, mainly because this proved too difficult to score consistently across taxa, and this additional behavioral complexity would be valuable to incorporate into our scoring system for a more restricted set of taxa in a future study.

Given that, based on evidence from depositional environments and certain body proportions (limb length vs. body length), Wilberg et al. (2019) report no major changes in habitat preference over the evolutionary history of crown *Alligatoroidea*, we tentatively score extinct *Alligator* F. M. Daudin 1802 species in the same way as we do extant species. Of course, we cannot observe the natural history of these extinct species directly, so our ecotype scores for extinct *Alligator* species remain speculative. We used a similar approach to score articulated skeletons of extant crocodilians with traditional family-level designations, basing their aquatic affinity scores on sampled species from each group in our dataset. More specifically, we scored "*Crocodylidae*, indet." specimens as "data deficient" (given that they might represent either a highly aquatic or a moderately aquatic species), and "*Alligatoridae*, indet." specimens as "moderately aquatic," given that we treat all crown alligatorids this way. Additional details on the scorings of related extinct crocodylians are given in the supplementary tables associated with this chapter.

Within this single bin of "moderately aquatic" crocodilians, we stress the existence of a finer continuum. In particular, we acknowledge that the brevirostrine species *Osteolaemus tetraspis* E. D. Cope 1860 and *Paleosuchus trigonatus* J. G. Schneider 1801 are less aquatic, overall, than the other "moderately aquatic" crocodilian species noted above, despite the fact that we give them here the same ecotype score. *O. tetraspis* often inhabits sparsely inundated shallow inland streams and cave systems (Crocodile Specialist Group 1996; Vitt & Caldwell 2014; Shirley et al. 2017), and it regularly hunts small terrestrial prey (Pauwels et al. 2007), which may make up the entirety of its adult diet in cave-dwelling populations (Shirley et al. 2017). Though quantitative comparisons of water and land habitation are sparse, one study found that some (but not all) captive *O. tetraspis* individuals spent the overwhelming majority of their time on land (Uwakaneme et al. 2004)—a finding consistent with the highly terrestrial diet of some populations (Shirley et al. 2017). *P. trigonatus* is likewise often found about small inland streams in heavily forested areas (Medem 1958; Magnusson et al. 1987; Brochu 2001). Indeed, *P. trigonatus* adults primarily consume terrestrial prey (Jackson et al. 1974; Magnusson et al. 1987), and spend vast stretches of time on land, with one wild individual apparently residing under dry ground for 10 days (Magnusson & Lima 1991). These relatively terrestrial habits have led some authors to suggest a connection between terrestriality and brevirostry (Medem 1958 and references therein; Brochu 2001). However, the closely related *P. palpebrosus*, which is more conspicuously brevirostrine than its congener (Medem 1958), resides about larger bodies of water and consumes relatively less terrestrial prey (Magnusson et al. 1987), casting doubt on this suggestion. Regardless, these species (or some of their subpopulations) still typically spend extended periods of time in the water and have at least partly aquatic diets, with some populations making more use of aquatic resources than others. As a result, we still consider *P. trigonatus* and *O. tetraspis* moderately aquatic, with equal-to-subequal regions of niche space and habitation time devoted to land and water.

### Text S5. How contentious are the aquatic affinities of extinct species?

Now that we have provided a classification system for describing the aquatic habits of extant and extinct species, we should discuss how feasible or difficult it was to place extinct species within this system. We might phrase this question another way: Given our classification system for assigning aquatic affinities in a consistent way across amniotes, which extinct taxa were possible for us to confidently classify given current lines of evidence?

We provide a brief written overview of confident *a priori* assignments here and then go through, broadly, the many taxa for which *a priori* classification was infeasible, due either to a lack of data on the taxon or ongoing disputes about their aquatic habits by different researchers. We consider the aquatic habits for all such taxa "contentious," "ambiguous," or otherwise "unclear" as noted in the main-text Introduction and Discussion. This section (Text S6) thus serves as a short, written justification for our decision to label certain taxa as having contentious aquatic affinities. However, a more comprehensive set of justifications and reviews of existing literature are available in Table S1, which contains a row-by-row summary of previous comments on the aquatic habits of each sampled taxon in our linear morphometric dataset.

*Text S5.1. Extinct taxa in our dataset with clear and non-contentious aquatic affinities.*

The most clearly flippered marine reptiles of the Mesozoic—including ichthyopterygians, plesiosaurs, mosasaurids, metriorhynchids, and *Archelon ischyros* G. R. Wieland 1896 (described as having flippers in Huxley 1871; Wieland 1902; Romer 1956; Caldwell 2002; Maxwell 2012)—have been interpreted as unable to come onto or traverse land without great difficulty (Romer & Parsons 1949; Griebeler & Klein 2019). Their limb and skull osteologies are highly adapted for aquatic and pelagic lifestyles. Moreover, live birth in mosasauroids (Carsosaurus?) (Caldwell & Lee 2001), plesiosaurs (O'Keefe & Chiappe 2001), and ichthyopterygians (Maxwell & Caldwell 2003; Motani et al. 2014; Miedema et al. 2023) would certainly facilitate fully aquatic habits in those clades (even if 'live' birth does not imply aquatic habits). Likewise, the lack of dorsal ornamentation with thermoregulatory osteoderms in metriorhynchids suggests that they either did not engage in routine basking behavior or primarily relied on muscular contraction associated with near-constant swimming to generate heat, supporting highly or fully aquatic habits for this group (Spindler et al. 2021; De Araújo Sena & Cubo 2023). There is a general consensus about the highly or fully aquatic habits of these groups, and we found no sources disputing their highly or fully aquatic nature. Given the likelihood that metriorhynchids returned to land to lay their eggs (like all other archosaurs), we classified them as either highly or fully aquatic. We classified plesiosaurs, ichthyopterygians, and mosasauroids as fully aquatic.

Fully terrestrial habits were more difficult to assign *a priori*, as species may pursue a transiently aquatic lifestyle without any obvious gross morphological or physiological adaptations to life in the water (Dawson et al. 1977). However, we did sparingly assign fully terrestrial habits *a priori* to highly derived arboreal taxa (e.g., *Megalancosaurus preonensis* M. Calzavara et al. 1981; see Table S1). We inferred either fully terrestrial *or* transiently aquatic habits (i.e., "predominantly terrestrial" habits) *a priori* for most extinct taxa lacking any previously reported morphological adaptations to an aquatic lifestyle and appearing instead specialized for cursorial or arboreal habits or for consuming terrestrial vegetation. This included most sampled dinosaurs (but see some exceptions below); the stem avians (viz., basal dinosaurs) *Eoraptor lunensis* P. C. Sereno et al. 1993 and *Herrerasaurus ischigualastensis* O. A. Reig 1963; the stem archosaurs *Protorosaurus speneri* H. Von Meyer 1832, *Prolacerta broomi* F. R. Parrington 1935, and *Mesosuchus browni* D. M. S. Watson 1912; the basal stem crocodylians "Popobuddy", *Poposaurus gracilis* M. G. Mehl 1915, and *Neoaetosauroides engaeus* J. F. Bonaparte 1969; *Thrinaxodon liorhinus* H. G. Seeley 1894; and all sampled extinct stem artiodactyls, stem perissodactyls, and stem equids. For more details about which particular taxa were assigned predominantly terrestrial habits, please see Table S1.

*Text S5.2. Extinct taxa in our dataset with unclear or contentious aquatic affinities.*

For the majority of fossil taxa in our dataset, we could not classify their aquatic affinities *a priori*. We briefly touch on why this was, broadly, for the taxa listed below, but more details for each sampled taxon are given in Table S1.

*Text S5.2.1. Various basal stem reptiles and stem mammals*.

Many of the extinct stem reptiles and stem mammals in our dataset have been sparsely studied (e.g., *Adelosaurus huxleyi* [originally "*Proterosaurus huxleyi*" in A. Hancock & R. Howse 1870; see Evans 1988 for most recent description]; *Spinoaequalis schultzei* M. De Braga & R. R. Reisz 1995). Others have had conflicting interpretations or lines of evidence that might suggest alternatively more semi-aquatic or terrestrial habits. For example, *Cabarzia trostheidi* F. Spindler et al. 2019 has been reconstructed as a terrestrial facultatively bipedal ground-dweller, but it has a broad ulnare similar to those of some aquatic stem reptiles, and the taxon comes from a deposit with mixed terrestrial and freshwater fauna (Spindler et al. 2019). Likewise, *Edaphosaurus* E. D. Cope 1882 shows no obvious morphological adaptations to an aquatic lifestyle, and it seems intuitive to assume that it was fully terrestrial. However, *Edaphosaurus* specimens are commonly found in depositional environments representing swamps or swampy lakes (Romer 1956; Bakker 1982), and at least one fully articulated *Edaphosaurus* skeleton has been found in marine deposits (Reisz et al. 1982). As a result, we cannot rule out semi-aquatic habits for the taxon *a priori*.

Conversely, various subtle morphological features or facies evidence, along with variable levels of pachyostosis (see Houssaye 2009 and references therein), suggest that the stem reptiles *Claudiosaurus germaini* R. L. Carroll 1981, *Tangasaurus mennelli* S. H. Haughton 1924, and (to a lesser extent) *Thadeosaurus colcanapi* R. L. Carroll 1981 may have had a semi-aquatic lifestyle (Harris and Carroll 1977; Carroll 1981; Currie 1981; Sues 2019). However, these taxa all retain well-developed jointed appendages, and it remains unclear whether and to what extent they traversed on land. We have insufficient evidence to say whether their habits were—for *T. colcanapi*—fully terrestrial, transiently aquatic, or moderately aquatic, and—for the others—transiently, moderately, or highly aquatic.

The aquatic affinities of sampled "parareptiles" are particularly contentious. *Barasaurus besairiei* J. Piveteau 1955 has been varyingly described as aquatic (McMenamin 2018) and terrestrial (Ketchum &

Barrett 2004), in either case with little justification. It is found in the same stratum as *C. germaini* and *Hovasaurus boulei* J. Piveteau 1926 (Ketchum & Barrett 2004), but it has no osteological features obviously indicating adaptation to an aquatic lifestyle (see Meckert 1995). Thus, the extent of its aquatic habits remains unclear. Likewise, the aquatic affinities of mesosaurs have garnered persistent dispute. Mesosaurs were viviparous (Piñeiro et al. 2012) and clearly adapted for aquatic life in the water (Romer 1956; Demarco et al. 2018), leading many authors to assume that mesosaurs were fully aquatic, but recent morphological and morphometric considerations have led others to suggest a more semi-aquatic lifestyle for this group (reviewed in Demarco et al. 2018). Adult mesosaurs may have occupied pelagic environments (Holmes 1953; Verrière & Frobisch 2022), but we lack direct evidence concerning whether and with what frequency they may have actually traversed land, so it remains unclear whether mesosaurs were moderately, highly, or fully aquatic.

*Text S5.2.2. Basal ichthyosauromorphs.*
As noted above, we scored all ichthyopterygians as fully aquatic *a priori*. However, the aquatic affinities of non-ichthyopterygian ichthyosauromorphs were more difficult to score. Although viviparity has been demonstrated in some ichthyopterygians (Maxwell & Caldwell 2003; Motani et al. 2014; Miedema et al. 2023), there is no direct evidence for oviparity or viviparity non-ichthyopterygian ichthyosauromorphs (with the exception of *Chaohusaurus* C. C. Young & Z.-M. Dong 1972, which may or may not be an ichthyopterygian [Moon 2019]) and egg-laying cannot be ruled out for these taxa. Nasorostrans and hupehsuchians are both highly modified for an aquatic lifestyle (see Carroll & Dong 1991; Motani et al. 2015). Hupehsuchians are thought to have had a difficult time clambering onto land (see Carroll & Dong 1991), but whether and to what extent they went on land has not to our knowledge been discussed. Nasorostrans had highly derived flippers, but may have been capable of dorsoventral flexion at the carpus, enabling limited locomotion on land and potentially an amphibious lifestyle (Motani et al. 2015). Given these ambiguities, we could not confidently determine *a priori* whether nasorostrans and hupehsuchians were fully aquatic, highly aquatic, or moderately aquatic. *Chaohusaurus* C. C. Young & Z.-M. Dong 1972, with its demonstrated viviparity and highly derived aquatic specializations (Motani et al. 2014) seems likely to be fully aquatic, but its uncertain position near the base of the ichthyosauromorph phylogeny (see Moon 2019) prompted us to remain agnostic about its aquatic habits in order to avoid biasing the results of our model, since its aquatic affinities would exert a strong and variable influence on the ancestral state reconstruction for the aquatic habits of *Ichthyopterygia* depending on the position of *Chaohusaurus* within *Ichthyosauromorpha*.

*Text S5.2.3. Basal omnisauropterygians.*
Despite a long history of study, the aquatic affinities of non-plesiosaurian omnisauropterygians are seldom obvious or explicitly discussed. The basal omnisauropterygians *Eusaurosphargis dalsassoi* S. Nosotti & O. Rieppel 2003 and *Helveticosaurus zollingeri* B. Peyer 1955 have received some of the few explicit ecological interpretations for particular species: *E. dalsassoi* has been interpreted as either predominantly terrestrial (Scheyer et al. 2017) or semi-aquatic (Klein & Scheyer 2024), and the basal omnisauropterygian *H. zollingeri* has been interpreted tentatively as being capable of clambering onto land (Rieppel 1989; Houssaye 2009). One of the few more comprehensive allusions to omnisauropterygian aquatic habits is by Motani and Vermeij (2021), who interpret placodonts and saurosphargids as having minimized their terrestrial travel and eaten exclusively in the water, and they interpret other non-pistosauroid omnisauropterygians (e.g., "nothosaurs" and "pachypleurosaurs") as either doing the same or making use of both terrestrial and marine food sources. Based on the inferred flexibilities of their appendicular skeletons, all of these taxa were interpreted as retaining the ability to locomote on land (Motani & Vermeij, 2021). Another explicit consideration of basal omnisauropterygian habits comes from Thewissen and Nummela (2008), who tentatively (i.e., with a "?") interpret all sauropterygians as making use of both terrestrial and aquatic food sources and occasionally leaving the water. Another explicit allusion to basal omnisauropterygian aquatic habits (Klein & Sander 2019) notes that the earliest pachypleurosaurs may have been terrestrial or semiterrestrial, or were at least not as aquatic as previous researchers have suggested.

However, taken together, these conflicting or uncertain reconstructions made it difficult for us to assign one of moderately, highly, or fully aquatic habits to non-plesiosaurian omnisauropterygians *a priori*.

*Text S5.2.4. Thalattosaurs*.
Although clearly adapted to an aquatic lifestyle and known exclusively from marine deposits (Merriam 1904; Romer 1956; Rieppel et al. 2005), most thalattosaurs seem to retain well-developed jointed limbs, suggesting that they may have retained the ability to make limited trips on land. The degree to which they traveled onto land remains unclear, so we cannot resolve whether thalattosaurs were moderately, highly, or fully aquatic. Motani and Vermeij (2021) and Thewissen and Nummela (2008) tentatively interpret them as making use of both terrestrial and aquatic food sources and occasionally leaving the water, but this interpretation may be difficult to reconcile with other ecological interpretations for some derived askeptosauroids (Druckenmiller et al. 2020), which may have had more difficulty traversing land, and for askeptosauroids, which might have had a semi-terrestrial lifestyle characterized by regular sojourns onto land (Bastiaans 2024). Aside from these occasional allusions, the precise aquatic affinities of thalattosaurs are seldom discussed, and we were unable to confidently classify them as one of moderately, highly, or fully aquatic.

*Text S5.2.5. Basal mosasaurians* and other anguimorphs.
Although many derived mosasaurians (e.g., mosasaurids, *sensu* Augusta et al. 2022) are usually considered fully aquatic (Bell & Polcyn 2005; Motani & Vermeij 2021; Augusta et al. 2022; though see Thewissen & Nummela 2008), the basal interrelations of mosasaurians and the evolution of aquatic habits within this clade remain contested, with different authors arguing for varying degrees of facultatively aquatic or semi-aquatic habits in non-mosasaurids (see Bell & Polcyn 2005; Dutchak & Caldwell 2006; Dutchak & Caldwell 2009; Lindgren et al. 2011; Simões et al. 2017; Augusta et al. 2022). As a result, we classified all mosasaurids in our dataset as fully aquatic, and treated the aquatic habits of all non-mosasaurid mosasaurians as ambiguous.

The aquatic affinities of *Saniwa ensidens* J. Leidy 1870 are unresolved (Rieppel & Grande 2007). *Saniwa* is recovered from an aquatic paleoenvironment (Grande 1984) and displays some features (*e.g.,* reduced mesopodial ossification and tail elongation) which track aquatic habits in closely related aquatic lepidosaurs (Rieppel & Grande 2007), but these features alone do not imply a moderately or highly aquatic lifestyle (Rieppel & Grande 2007). We were consequently unable to determine the aquatic habits of this taxon *a priori*.

*Text S5.2.6. Pleurosaurs and other extinct rhynchocephalians*.
Pleurosaurs and *Vadasaurus herzogi* G. Bever & M. Norell 2017 are stem rhynchocephalians that are both thought to have been secondarily aquatic (Dupret 2004; Bever & Norell 2017; Motani & Vermeij 2021), but whether and to what extent they ventured onto land remains unclear. *V. herzogi* has some features associated with aquatic habits (Bever & Norell 2017), and pleurosaurs have more derived and likely more aquatically adaptive morphologies (Dupret 2004). However, both clades show a blend of aquatic and terrestrial features, making it difficult to confidently assign them one aquatic-affinity bin. More details on the aquatic and terrestrial features in these taxa is provided in <u>Text S12</u> and <u>Table S1</u>.

*Text S5.2.7. Potentially semi-aquatic non-avialan dinosaurs*.
As we mentioned above, we classified most extinct dinosaurs in our dataset as being predominantly terrestrial. There were a few notable exceptions—namely *Halszkaraptor escuilliei* A. Cau et al. 2017, *Juravenator starki* U. B. Göhlich & L. M. Chiappe 2006, and *Spinosaurus aegyptiacus* E. Stromer 1915. We briefly summarize existing contentions associated with the aquatic habits of these taxa below.

*Halszkaraptor escuilliei* was a basal dromaeosaurid from the Cretaceous of Mongolia (Cau et al. 2017). Noting several histological and morphological features considered adaptive for an aquatic environment, Cau et al. (2017) suggested that *H. escuilliei* was amphibious (i.e., “moderately aquatic” under our binning procedure). Brownstein (2019) challenged many of the arguments used by Cau et al. (2017) to infer aquatic habits in *H. escuilliei*, casting doubt on the aquatic lifestyle of the species. However, Cau (2020) disputed many of these counterpoints, defending the initial inference that *H. escuilliei* was moderately aquatic. This discourse suggests that the aquatic affinity of *H. escuilliei* remains contentious, and we score it accordingly as ambiguous.

*Juravenator starki* is a compsognathid recovered from a limestone deposit in the Late Jurassic Solnhofen reef archipelago of Bavaria (Göhlich & Chiappe 2006). *J. starki* has no obvious gross morphological features strongly suggesting an aqauatic lifestyle (Göhlich & Chiappe 2006; Chiappe & Göhlich 2010). However, *J. starki* was recovered from a lagoonal/shallow subtidal palaeoenvironment (Bell & Hendrickx 2021), in association with fish remains (Chiappe & Göhlich 2010), has a tooth morphology reminiscent of some other opportunistically piscivorous coelurosaurs (Bell & Hendrickx 2021), and has unique ornamental scales on its tail that have been interpreted as resembling integumentary sense organs useful for detecting prey in the water (Bell & Hendrickx 2020; Bell & Hendrickx 2021). As a result, Bell and Hendrickx (2020) proposed a "semi-aquatic opportunist" lifestyle for *J. starki*, in which it hunted by "submerging its tail (and other body regions with integumentary sense organs) to acquire sensory information" about "aquatic prey on the lagoon margins and other water bodies on the Solnhofen reef archipelago" (Bell & Hendrickx 2020). This interpretation is consistent with both transiently and moderately aquatic habits. In addition, this recent interpretation remains admittedly "speculative" (Bell & Hendrickx 2020), so we treat the precise aquatic affinities of *J. starki* as uncertain.

*Spinosaurus aegyptiacus* was a tetanuran theropod from the Cretaceous of North Africa (Arden et al. 2019) and the dispute surrounding its aquatic affinity is notoriously fraught. Spinosaurids likely consumed fish (Arden et al. 2019; Brusatte 2021, and references therein), and were thus at least slightly aquatic. Isotopic data supports a somewhat aquatic lifestyle for many spinosaurids, which have ∂18 oxygen phosphate signatures more similar to semi-aquatic crocodilians and turtles than to other (presumably terrestrial) theropods (Amiot et al. 2010). However, it remains unclear precisely how aquatic spinosaurids really were. Several morphological and histological features have supported more extreme aquatic habits in *S. aegyptiacus*, but it remains disputed whether this species was merely a wading shoreline piscivore (Sereno et al. 2022) or a highly aquatic pursuit predator (Ibrahim et al. 2020). On the one hand, tall neural spines on the caudal vertebrae supported a contentiously paddle-like tail that may acted as an efficient propulsive organ in water (Ibrahim et al. 2020), and the bone compactness of its femur falls within the range of modern diving or highly aquatic reptiles (Fabbri et al. 2022). On the other hand, most 3D computer models of *S. aegyptiacus* have found it to be a competent walker on land, and a highly buoyant, slow, and inefficient swimmer (Henderson 2018; Sereno et al. 2022). Moreover, at least two *Spinosaurus* specimens have been identified in inland fluvial deposits, suggesting a tolerance for less permanently aquatic and non-marine habitats (Sereno et al. 2022). A recent study (Smart & Sakamoto 2024) used cluster analysis on 2D linear measurements of the cranium to distinguish terrestrial from aquatic species and confidently recovered *S. aegyptiacus* as being either semi-aquatic or fully aquatic (their results could not distinguish between these groups). The results of this recent analysis are thus consistent with both the "shoreline wader" and the "subaqueous forager" hypotheses and fail to discriminate between them. Synthesizing from these conflicting accounts, we cannot resolve whether *S. aegyptiacus* was transiently, moderately, or highly aquatic. We therefore considered its aquatic habits ambiguous and refrained from scoring them *a priori*.

### Text S6. Previously considered lines of evidence for webbing and flipper form.

Interdigital webbing and flippers can be difficult to identify in the fossil record. Some exceptional fossils preserve the soft-tissue remnants of flippers (Lindgren et al. 2013; Frey et al. 2017; Vincent et al. 2017; Joyce et al. 2021; Eriksson et al. 2022), interdigital webbing (Rösler & Tatizana 1985; Fielding et al. 2005; Ji et al. 2006; MacDougall et al. 2020), or their traces, providing direct evidence for these structures.

There are also abundant fossil trackways that can provide evidence of interdigital webbing in extinct species—as considered, for example, by King & Benton (1996), Vila et al. (2014), Lee et al. (2020), Helm (2022), and many others. However, trackways can be deceptively unreliable lines of evidence for interdigital webbing. False positive “webbing” traces can result from the interaction of certain hand or foot geometries and application forces with certain substrates (Falkingham et al. 2009). Moreover, even in cases where such misleading autopod-substrate interactions can be discounted, the connection of a particular trace to a particular producer taxon can be difficult to verify and thus remains uncertain (except, of course, in the circumstance that the producer is preserved as a body fossil at one end of its own trackway). This difficulty complicates any effort to validate interdigital webbing in a given extinct species using trace-fossil data alone, as such inferences necessarily rely (much like osteological correlates) on indirect morphological comparisons between the body fossils of various producers and the shape and arrangement of trackway footprints, and very similar trackways might be realized by very different taxa—especially when the variables of gait, weight application, and the material give of the substrate remain unfixed across traces (see Falkingham et al. 2009).

As a result, in most cases, unless exceptional preservation obtains, the presence of soft-tissue structures must be inferred by validating osteological correlates within a phylogenetic framework (Witmer 1995). Previous studies have suggested various putative osteological correlates for interdigital webbing, including a relatively symmetrical or ovoid hand/foot planform (Carroll 1981; Currie 1981; Thewissen & Fish 1997; Modesto 2010), a lengthened anterior-most digit (Thewissen & Fish 1997; Ibrahim et al. 2014), lengthened and flattened unguals (Bejder & Hall 2002; Ibrahim et al. 2014), distally flared proximal phalanges (Madar 2007), and differences in the proportions of the three main limb regions (the stylopodium, zeugopodium, and autopod [Williston 1914]). Many of these correlates have compelling extant exemplars. For example, interdigital webbing has been linked to the overall shape of the foot in some carnivorans (Thewissen & Fish 1997), and to dorsoventrally flattened unguals in some wading and diving birds (Manegold 2006). In addition, the limb proportions of turtles track their preferences for different aquatic habitats (Joyce & Gauthier 2004), and the proximal phalanges of some webbed geckos (e.g., *Ptychozoon* spp.) have dramatically flared distal epiphyses (personal observation of specimens in Table S1). Even so, none of these proposed soft-tissue correlates have been evaluated within an extant phylogenetic bracket encompassing the extinct stem groups of interest (Witmer 1995). Consequently, given the great variety of webbing forms in modern amniotes (Figure S2), these correlates require strict validation within a phylogenetic framework before we can use them to reconstruct the aquatic habits of extinct species.

The identification of soft-tissue flippers poses a similar problem. Many flippers have hyper-derived osteologies that make them easy to recognize, with flattened long bones (Blob et al. 2016), immobilized joints (Cooper 2008), hyperphalangeal digits (Fedak & Hall 2004), accessory digits (Maxwell et al. 2014), reduced perichondral ossification (Caldwell 2002), or hyper-elongated hands and feet (Romer 1956; Joyce & Gauthier 2004; Joyce et al. 2021). However, many extant species (including sirenians, beaked whales, and trionychids) have flippers that lack some or all of these features (personal observation of specimens in Table S1). Instead, they have jointed and often terrestrially proportioned limbs that would make it difficult to identify them as flippers from the bones alone. Consequently, the flipper phenotypes of some extinct groups (including pleurosaurs, hupehsuchians, thalattosaurs, and basal sauropterygians) remain unclear *a priori*, with historically varied interpretations that have been central to previous discussions of their aquatic habits (e.g., Williston 1914; Romer 1956; Mekarski et al. 2019; Krahl 2021).

#### Text S7. Geometric morphometric analyses of the acropodium and zeugoautopodium.

As described in the STARR Methods, we quantified acropodial planform shape using a linear morphometric approach, by combining three length measurements ($L_{anterior\text{-}most}$ digit, $L_{posterior\text{-}most}$ digit, and $L_{longest}$ digit) in various ways to create two ratio-based “symmetry indices” that serve to capture acropodial asymmetry. This enabled us to quantify the acropodial planform shapes of a large number of amniotes with disputed digital homologies, differing numbers of digits, and varied *in situ* splaying positions in sampled taxa. However, while this approach is time-efficient (requiring only three linear measurements per specimen) and avoids many of the taphonomic and anatomical variations that might skew other analyses (as discussed below), it does not provide a comprehensive description of acropodial planform shape.

Thus, in order to supplement our linear morphometric approach and obtain a more comprehensive picture of acropodial planform shape, we performed two geometric-morphometric analyses on a subset of 103 specimens with limbs photographed in dorsal planform view (Figure S10a–b). These analyses considered the shape of fully articulated hands (autopodia) or distal forearms (zeugo- and autopodia). Landmarks were placed in ImageJ (v. 1.53t) at the proximal, distal, and/or lateral extremities of constituent bones in each articulated skeleton. Hand shape was captured with 15 fixed landmarks, and forelimb zeugoautopod shape with 11 fixed landmarks and 20 sliding semilandmarks (Figure S10a–b). Fixed acropodial landmarks were defined with reference to the anterior-most and posterior-most digits, again enabling us to compare acropodial planform shapes in taxa with differing numbers of digits. All fixed landmarks are described in Figure S10.

Downstream of landmarking, geometric morphometric analyses of the acropodium and zeugoautopodium were completed in the same manner described in the STARR Methods for our geometric-morphometric analysis of ungual phalanges. Briefly, generalized Procrustes Superimposition was performed using the gpagen command within the R geomorph package (v. 4.0.4). Resulting Procrustes coordinates were then written to a TPS file and uploaded to PAST (v. 4.11) for principal component analysis (PCA) and thin-plate spline comparisons. Principal Component (PC) scores were subsequently compared qualitatively in 2D morphospaces by observing the convex hulls for different limb phenotype and aquatic affinity guilds.

These geometric morphometric analyses for assessing acropodial symmetry were intended just to supplement our linear morphometric approach (using the "symmetry indices" described above). However, these analyses have a major shortcoming that deserves special mention: They involve landmarking articulated elements that can in most cases (depending on the taxon sampled) change position independent of one another. This can lead to significant in-species variation. For example, consider the fossil of a putatively semi-aquatic taxon preserved in a slab. If its left hand was preserved with the fingers fully splayed (abducted from the autopodial midline) and the right hand was preserved with the fingers fully appressed (adducted to the autopodial midline), these two hands would have drastically different acropodial planforms, and one would need to account and correct for this greater possible intraspecies variation in shape in order to clearly isolate any ecological or behavioral signals in shape variation across all sampled taxa. However, even bearing this caveat in mind, such analyses can still prove insightful, for two reasons:

• The first reason these analyses remain potentially insightful is that the ecological or limb-phenotype signal in acropodial planform shape could be strong enough that it manifests clearly even despite the extra noise generated by potential intraspecific variation in the relative preservation positions of landmarked elements. For example, if flippers happen to have very different shapes from non-flippered limbs, their average shape might be recovered significantly different from that of webbed and un-webbed limbs, even with the added variation resulting from the stochastically preserved differences in orientation of their landmarked elements. This high-enough signal-to-noise ratio is observed in the results of our geometric analysis of the zeugoautopodium (Figure S10a). In this analysis, the degree of digital splaying and the angle formed by main axes of the zeugopodium and autopodium (what we might call the "wrist angle") were uncorrected, and thus permitted to stochastically vary both inter- and intraspecifically. That is, some specimens happened to have splayed fingers, and others not; some specimens happened to have a hand angled clockwise away from the zeugopodials and others in counterclockwise, and so on. Even so, along principal component (PC) 1, which accounted for nearly half of all variance in shape (49.9%), the majority of flippered limbs occupied a distinct region of morphospace untapped by webbed and non-webbed specimens (Figure S10a). This uniquely 'flippered' region of morphospace (captured by more negative PC1 scores) mainly reflects the relative lengths of the zeugopodium and autopodium and the relative widths at the base and proximodistal center of the acropodium. Thus, despite the tremendous extra possible variance permitted by allowing individual digits and limb regions to move with respect to one another as a consequence of *in vivo* movements, the broader limb-phenotype signal manages to obtain and in fact remains the strongest signal observed in the dataset. Rigorously quantifying this signal would require additional phylogenetic comparative tests that correct for the greater potential variance enabled by these permitted inter-element movements. This rigorous quantification would be time-consuming and only further support the results from our linear morphometric data (i.e., by demonstrating that flippered taxa have relatively longer acropodia and relatively shorter zeugopodia) so we refrain from quantifying this signal here. Even so, this geometric morphometric analysis of the zeugoautopodial planform lends additional qualitative support to our linear morphometric results, highlighting that the observed differences in limb region proportions are robust enough to obtain in a more comprehensive multidimensional analysis of zeugoautopodial shape. This is one reason that we should refrain from fully dismissing geometric morphometric analyses that incorporate independently mobile elements: Though we should view them with caution and take extra care to correct for the extra non-informative variation they enable, they may nevertheless yield insights robust enough to shine through this noise and reveal key morphometric differences among the groups of interest.

• The second reason these analyses remain potentially informative is that the possible intraspecific variance in the relative positions of landmarked elements (i.e., the extra "un-informative" variance in shape that results from the movement of landmarked elements relative to one another) may itself actually inform our understanding of their biology, by reflecting differences in the soft-tissue anatomy or aquatic habits of these species. For example, even though hundreds of articulated plesiosaur flippers have been recovered, none have ever been observed with their fingers splayed. This suggests that the un-fossilized soft-tissue anatomy of the plesiosaur flipper restricts the movement of plesiosaur digits relative to one another, both *in vivo* and even during *post-mortem* biostratinomic processes and burial. Likewise, interdigital webbing might constrain the number of likely *in vivo* digital splaying positions, reducing the likelihood for some of these positions to be preserved during fossilization. It is thus notable that, in our geometric morphometric analysis of acropodial planforms (Figure S10b), webbed acropodial planforms occupied a much narrower band of PC1 (representing 40.8% of total shape variance and capturing variation in acropodial symmetry and direction of taper). Given that we did not rigorously quantify and correct for the potential intraspecific variation in digital splaying that we observe for webbed, unwebbed, and flippered species, we refrain from applying phylogenetic comparative tests to assess the significance of these differences. However, the qualitative difference in morphospace occupancy is striking and merits further investigation. The potential significance of this difference in PC1 variance among limb phenotypes offers a second reason for why we should refrain from fully dismissing geometric morphometric analyses that incorporate independently mobile elements: Though we should again view them with caution and take extra care to correct for the extra variation they enable, this extra variation may not be distributed equally among sampled groups, and may thus itself be informative for characterizing limb phenotypes in extinct species.

### Text S8. Sensitivity analysis rationale and results.

*Text S8.1. Examining the effects of data-transformation method on phylogenetic statistical test results.*
Transforming raw data before performing statistical analyses can have severe effects on test results. However, data transformation can in some instances be necessary to make the data suitable for the method of analysis (e.g., increasing likelihood of homoscedasticity of a dataset through Box-Cox transformation [Box & Cox 1964, 1982]). For this study, all phylogenetic comparative tests, including phylogenetic analyses of variance (phylANOVAs [Garland et al. 1993]), correlation tests (Paradis & Schliep 2019), and phylogenetic binomial logistic regression analyses (Ho & Ané 2014) were performed here using presumably non-parametric, simulation-based methods, which should, theoretically, not require the raw input data to be normally distributed or homoscedastic. However, several previous studies have tended to log-transform datasets as a precaution to minimize homoscedasticity or achieve a more proportional scale prior to performing phylogenetic comparative tests (e.g., Benson et al. 2012; Soul & Benson 2017; Simões et al. 2022). To our knowledge, the benefits and consequences of this practice have not been rigorously investigated for phylogenetic comparative methods, and the size of our dataset and range of statistical tests performed provide a unique opportunity for us to explore the influence of data-transformation practices in

more detail. For this reason, we sought to test whether data transformation was required or impactful for all downstream statistical analyses.

In order to check whether prior transformation of our data affected the significance verdicts resulting from any of the phylogenetic comparative tests reported in this paper, we directly compared p-value significance across (i) raw (un-transformed) data, (ii) $log_{10}$-transformed data, and (iii) Box-Cox transformed data (Figure S29a; script S5). For each yielded p-value, statistical significance was coded as either "significant" ($p < 0.05$) or "insignificant" ($p \geq 0.05$) and compared across all three transformation methods to quantify accordance between them. Only when all transformation methods agreed on either "significant" or "insignificant" results for a specific p-value would it be coded as "identical" across transformation method; otherwise it was coded "divergent."

The resulting stacked bar plots (Figure S29c) show a strong (>95%) consensus in significance verdicts between the raw data and both transformation methods for phylANOVA- and phyloLev-associated tests (100.00% and 95.06% consensus values for phylANOVA omnibus and post-hoc tests, respectively; 100.00% and 99.04% consensus for phyloLev omnibus and post-hoc tests, respectively), and moderate-to-high (>75%) consensus in significance verdicts for phybLR-associated tests (binomial tests for AUC [recording best model performance across thresholds] and the optimal model [recording best model performance at its optimal threshold]). These results (reported in Table S6a) confirm the suitability of the raw data for analysis under use of these methods and call into question the need for a blanket transformation prior to phylogenetic statistical analysis when performing phylogenetic ANOVAs, phylogenetic logistic regression analyses, or other tests derived from them. Because p-values are treated as binomial values ("significant" or "insignificant") in this instance, we tested for divergence of consensus among transformation methods from a random binomial distribution using a binomial exact test. All our binomial tests indicate strong divergence from random binomial distribution and therefore suggest high statistical significance of our results (all binomial exact test p-values << 0.001). We can therefore say with confidence that our raw data is suitable for processing in phylANOVA and phyloLev analyses and not statistically significantly affected by $log_{10}$- or BoxCox transformation.

In contrast, we observed a major divergence in significance verdicts for different transformation methods when testing the significance of phybLR model "fit" via McFadden's pseudo-$R^2$ (McFadden 1974), with consensus values of 54.64% and 68.50% for tests of significant McFadden pseudo-$R^2$ values averaged across 100 subsampling iterations of 3-fold cross-validation and for the whole dataset, respectively. This reduced consensus for McFadden's pseudo-$R^2$ verdicts suggest that this metric for assessing phybLR model fit is less robust against prior data transformation than a simple binomial test. We therefore suggest that, in cases where data transformation may be required for the project at hand, future phylogenetic ROC analyses should evaluate model performance using binomial tests rather than McFadden's pseudo-$R^2$ metrics for model likelihood.

We observed even greater divergence of significance verdicts for phylogenetic correlation tests in our analysis. Consensus values for tests assessing the intercept and slope of linear regression equations were 13.41% and 13.21%, respectively, indicating that the great majority of test results differed for raw and transformed datasets (Figure S29c). However, even for these parameters, consensus across data transformation methods is significantly high compared to random chance (all binomial t-test p-values << 0.001), indicating that transformation method, while affecting the phybLR testing procedure, does not necessarily significantly alter test results or choice of best-model fit. Even so, the low consensus observed among transformation methods leads us to recommend caution when transforming data prior to performing phylogenetic correlation tests, as data transformation does seem to alter correlation significance verdicts at high frequencies. However, as we suggest in Text S8.2 below, this result may be due to the weak and somewhat spurious nature of pairwise correlations among variables in our dataset, rather than the nature of the phylogenetic correlation tests themselves.

*Text S8.2. Examining the effects of tree topology on phylogenetic statistical test results.*

It has long been known that tree topology can bias and alter the results of statistical tests when the data are phylogenetically structured (Felsenstein 1985). As a result, the choice of one tree topology (one set of phylogenetic hypotheses) over another can potentially have major effects on the outcomes of phylogenetic comparative tests. Given that we were investigating limb evolution across amniotes, there were several contentious (often competing) phylogenetic hypotheses that we needed to fix prior to statistical analysis. Some of these disputes were resolved by defining polytomies in our supertrees. For example, the highly contentious placement of *Pan-Testudines* was addressed by placing it in a polytomy with *Pan-Lepidosauria*, *Pan-Archosauria*, and the frequently recovered clade of "euryapsid" Mesozoic marine reptiles (*Omnisauropterygia* + *Ichthyosauromorpha* + *Thalattosauria*) at the base of *Sauria* (see Text S3 for phylogenetic definitions of these clade names). All of our simulation-based statistical tests spontaneously collapsed polytomies over 10,000 iterations, so that *Pan-Testudines* was placed variably as the sister to each of *Pan-Lepidosauria*, *Pan-Archosauria*, the euryapsids, and different combinations thereof at roughly equal frequencies >1,000 times for each simulation-based test performed. However, in other cases, polytomies were insufficient to remove potential phylogenetic bias *a priori* from statistical tests. For example, the morphological and molecular topologies of squamates differ drastically in branching order, and reducing the entire subtree of *Squamata* to a comb-like polytomy would drastically reduce the phylogenetic precision of our analyses. For cases like these, we would need to repeat phylogenetic statistical tests given supertrees with distinct topologies. Of course, examining every combination of competing phylogenetic hypotheses for this project would have been computationally infeasible, requiring resources that far exceed Yale's High-Performance Computing clusters. Given this computational limitation, we selected three pairs of competing phylogenetic hypotheses that might have major impacts on our statistical test results, given the taxa for which we made predictions. As described in the Online Methods, these phylogenetic hypotheses concerned concerned (i) the basal-most clade of stem reptiles (*Captorhinidae* [Ford & Benson 2020] vs. a monophyletic grouping of "parareptiles" [Gauthier et al. 1988b]), (ii) the basal-most clade of *Omnisauropterygia* (*Hanosaurus hupehensis* C. C. Young 1972 [Wang et al. 2022] vs. a monophyletic clade of saurosphargids [Li et al. 2011]), and (iii) the tree topology of *Squamata* (*i.e.*, under molecular [Burbrink et al. 2020] vs. morphological [Gauthier et al. 2012; Mongiardino Koch & Gauthier 2018] topologies). To address these disagreements, we created a separate maximal supertree for each combination of these hypotheses, constructing a total of $2^3 = 8$ maximal supertrees as inputs for downstream statistical tests (Figure S29b). These different tree topologies are summarized in Figure S29b, and their full topologies are available in Data S1.

To test the effect of supertree choice on our phylogenetic comparative test results, we repeated the sensitivity analyses described above (Text S8.1) to assess consensus among all eight supertrees used in this study (Figure S29d; script S5). Each time a p-value significance verdict differed ("significant" vs. "insignificant") in at least one of the eight supertrees, the respective p-value was scored as "divergent" across supertrees. Only in the case that all eight supertrees showed consensus (all eight p-values implying significance or all eight p-values implying non-significance) was the test result scored as "identical." Overall, our phylogenetic comparative tests were highly robust against the choice of supertree. Neither phylANOVA (omnibus consensus frequency: 100.00%; post-hoc consensus frequency: 91.64%) nor phylogenetic Levene's test results (omnibus consensus frequency: 100%, post-hoc consensus frequency: 98.34%) are significantly affected by tree choice (Table S6b). Likewise, binomial tests for best-model performance had high consensus frequencies (>90%) across all supertrees, and even McFadden-based estimates of significant model fit were generally in agreement for all topologies (consensus frequencies >72%). Binomial exact tests were run to check divergence of significance verdicts from a random distribution yield p-values << 0.001 for all tested variables. Taken broadly, these results suggest a general robustness of our results against ongoing disputes on the interrelationships of basal pan-reptiles, basal sauropterygiforms, and squamates. However, we should note that this sensitivity analysis investigated the effects of tree topology on all statistical tests in aggregate and may overlook more nuanced influences of tree topology on phylogenetic test results for certain clades of interest.

It is noteworthy that, in contrast to the other phylogenetic comparative tests mentioned above, phylogenetic correlation test verdicts did show divergence among datasets using different supertrees (consensus frequencies were respectively 13.41% and 13.21% for regression intercept and slope significances). Because these tests also had the lowest consensus frequencies among transformation methods (Text S8.1), this low divergence may reflect the generally weak statistical signal (manifested here as low $R^2$ values) recovered for many of the significant phylogenetic correlations identified between variables in our dataset (Text S9, below). We therefore suspect that the poor consensus among phylogenetic correlation test verdicts, both across supertrees and across data transformation methods, highlights the weakness of many observed phylogenetic correlations, rather than the vulnerability of phylogenetic correlation tests themselves to alteration. However, additional work is required to investigate the mathematical causes of this divergence more rigorously to assess whether it is specific to our dataset. Conversely, the high consensus frequencies for phylANOVA, phyloLev, and phybLR test verdicts across supertrees and data transformation methods indicate particularly strong signals in the data—clear trends or patterns—which are robust against subtle shifts in tree topology at the base of major clades or systemic changes to data normality and homoscedasticity.

#### Text S9. Pairwise correlations among forelimb, hindlimb, and phalanx proportions.

To check for significant linear relationships among all length measurements taken, Pearson correlation coefficients (PCCs) were computed from both raw measurement data (Figure S19) and one of eight variance-covariance matrices derived from a maximal tip-dated supertree of all specimens in the dataset under a Brownian motion model of trait evolution (Figures S20–S27; script S3). Because phylogenetic correlation tests were uniquely *in*consistent among tree topologies in our analysis (see Text S8.2), we present the phylogenetic correlation matrices for all eight supertrees (Figures S20–S27) and discuss these results below. Both before and after accounting for phylogenetic autocorrelation (Figures S20–S27), most of our direct length measurements were strongly and significantly correlated (PCC > 0.70; $p < 0.05$). Correlations corresponding fore- and hindlimb measurements (e.g., humerus vs. femur, radius vs. ulna) were particularly strong across all supertrees (PCC ≥ 0.80; $p < 0.05$). This supports previous work identifying significant positive associations between humerus and femur length across amniotes (Campione & Evans 2012) and highlights that fore- and hindlimb lengths tend to scale monotonically in amniotes. We likewise observed strong, significant correlations between most corresponding fore- and hindlimb phalangeal measurements, including ungual height and length along its dorsal and ventral margins (PCC ≥ 0.89, $p < 0.05$), as well as proximal phalanx length, proximal epiphysis width, and distal epiphysis width: (PCC ≥ 0.80, $p < 0.05$ with significance verdicts that depended on tree topology). Ratios capturing forelimb proportions (e.g., $L_{(hand)}$ / $L_{(zeugopod)}$, and $L_{(humerus)}$ / $L_{(zeugopod)}$) displayed vanishingly small correlations with hand, radius, and humerus measurements (PCC ≤ 0.07; $p < 0.05$), weak and variable correlations with ulna length (PCCs ranging from 0.01–0.38; $p < 0.05$), and moderate but highly variable correlations with proximal phalanx measurements (PCCs ranging from 0.10–0.51; $p < 0.05$ with significance verdicts depending on tree topology). Corresponding ratios that captured hindlimb proportions ($L_{(foot)}$ / $L_{(tibia)}$, and $L_{(femur)}$ / $L_{(tibia)}$) showed a more consistent weak-to-trivial correlation with their constituent length measurements (PCC ≤ 0.33 but usually ≤ 0.15; $p < 0.05$).

Notably, corresponding fore- and hindlimb proportions exhibited consistent, moderate-to-strong pairwise correlations. Fore- and hindlimb stylopodium-to-zeugopodium ratios (or "SZRs": $L_{(humerus)}$ / $L_{(forelimb\ zeugopod)}$ and $L_{(femur)}$ / $L_{(tibia)}$) had a significant PCC of 0.58 across all eight supertrees, while the two corresponding acropodium-to-zeugopodium ratios (or "AZRs": $L_{hand}$ / $L_{zeugopodium}$ and $L_{foot}$ / $L_{tibia}$) had a significant PCC of 0.77–0.78 across all trees. For this study, we opted to examine forelimb and hindlimb segment proportions separately. However, the significant correlations observed among corresponding limb region proportions suggest that future studies might examine these proportions together for a broader analysis of limb evolution. That said, the distinct predictive capacities of fore- and hindlimb proportions with respect to aquatic habits and soft-tissue phenotypes (Figure 2; Figure S12) hint that such correlations may mask a more complex evolutionary interplay among dissociated fore- and hindlimb phenotypes.

### Text S10. Potential clade-specific osteological correlates of interdigital webbing.

We found no consistent differences in limb, acropodium, ungual, or proximal phalanx morphometry that could reliably distinguish webbed and unwebbed species across amniotes (Figure 2; Figures S8–S12). However, some morphometric features may reliably distinguish webbed and unwebbed species within particular clades (Figure 2b; Figures S8b, S15). In most of the clades or grades we sampled, differences in acropodium-to-zeugopodium length ratios between webbed and unwebbed species were negligible (Figure 2b). In some groups (e.g., ungulates, non-placental mammals [Figure 2b; Figure S8b]), hindlimb differences were notable, but small sample sizes ($n < 10$) precluded rigorous statistical comparison. However, within both *Testudines* (crown-group turtles) and *Ferae* (carnivorans + pangolins), we observed clear differences between webbed and unwebbed species. In both clades, webbed species were observed to have significantly longer hands and feet relative to both zeugo- and stylopodia (Figure 2b; Figure S8b; phylANOVAs and *post-hoc* Tukeys, each $p < 0.05$). Webbed turtles also tend to have relatively longer anterior digits, more distally widened proximal phalanges, and more elongate, flattened unguals than their unwebbed counterparts, although (likely due to small sample sizes) none of these differences were significant (Data S2). In line with these trends, ferans and turtles were unique among sampled clades in having predictable webbing phenotypes. For ferans, hindlimb region proportions were significant predictors of pedal webbing (one-tailed binomial tests, each $p < 0.05$; Figure S15). For turtles, proximal phalanx proportions were significant predictors of manual webbing, and limb region proportions significant predictors of pedal webbing (one-tailed binomial tests, each $p < 0.05$; Figure S15), although the low sample size considered for the former makes the predictive power of proximal phalanx shape dubious, and we encourage future studies to sample proximal phalanx shape using traditional and geometric morphometric methods with a broader cross-clade sample.

These clade-specific trends also merit broader caution. First, these trends rely on a particular definition of webbing (Figures S3, S5) that treats pangolins and tortoises as unwebbed. Tortoises, elephants, and sloths do have connective tissues between the digits (personal observation of specimens in Table S1). However, the stiff, immobilizing nature of these tissues leads us to consider their "stubbed" feet unwebbed. Second, these trends rely on a distribution of webbing that is phylogenetically biased within each of these clades. Webbing is highly concentrated in particular subclades within Testudines and Ferae. All unwebbed turtles in our dataset were testudinoids (either emydids, geoemydids, or testudinids). Likewise, most of the unwebbed ferans in our dataset were procyonids, canids, and pangolins, whose webbing phenotypes were often difficult to determine from available museum specimens (Table S1). As a result, though our preliminary results do hint at some promising correlates of webbing in turtles and ferans, these initial trends require validation with larger sample sizes and greater taxonomic sampling among the subclades of *Testudines* and *Carnivora*.

### Text S11. Future directions for investigating osteological correlates of interdigital webbing.

Although we found no unambiguous osteological correlates of interdigital webbing that held across amniotes (Figure 2; Figures S8–S12), a few more nuanced or spurious differences between webbed and unwebbed species merit further investigation, and may reveal new osteological correlates in the future. In particular, webbed taxa in our sample had notably (but non-significantly) less variable foot dimensions and flatter pedal unguals than unwebbed species (Figure S8b), with subtly different ungual curvatures that were captured by a small portion of the variance in landmark placements (Figure S10c). Perhaps as a result, we recovered foot planform shape (captured by $L_{(longest\ digit)} / L_{(marginal\ digits)}$) and ungual proportions (captured by $L_{(ungual)} / H_{(ungual)}$) as weak but significant predictors of pedal webbing in extant taxa (Figure 3h; one-tailed binomial tests: each $p < 0.05$). These trends lend suggestive (but spurious and inconclusive) support to previous claims that webbed rowers or paddlers tend to have more symmetrical foot planforms associated with rowing or paddling locomotor modes (Robinson 1975; Thewissen & Fish 1997; Modesto 2010) and longer, dorsoventrally narrower ungual phalanges (Ibrahim et al. 2014). We describe these results in more detail below. As a result, while our existing regression models for foot morphometry have a low predictive power, future studies with larger sample sizes might be able to validate this correlate more robustly. If this osteological correlate were validated, one could then use significant deviations from foot symmetry as evidence against the presence of webbing in particular fossil species.

Likewise, ungual proportions were poor cross-clade predictors of interdigital webbing in our sample (AUC = 0.62), but may hold promise to distinguish webbed and unwebbed species in future studies. Webbed amniotes in our dataset did not have significantly longer or flatter unguals than their unwebbed counterparts as captured by either traditional (Figure 2c) or geometric morphometrics (Figure 10c; Figure S8b). However, webbed amniotes in our dataset do seem to have notably (but non-significantly) more variable pedal ungual proportions than unwebbed and flippered species (Figure S8b), show notable (but again non-significant) differences in pedal ungual flatness (captured by $L_{(pD3\ ungual)}$ / $H_{(pD3\ ungual)}$ in Figure S8b and PC2 score [22.0% total variance] in Figure S10c), and exhibit significant differences in the dorsoventral trajectory of ungual curvature (captured by PC4 score [7.4% total variance] in Figure S10c). Taken together, these results may suggest that webbed species have distinct ungual shape profiles too nuanced for our models to identify. Thus, assessing the relationship between ungual shape and interdigital webbing with a larger sample size and a more complex, three-dimensional landmarking scheme may help to identify significant morphometric differences between webbed and unwebbed species, in line with previous suggestions (Bejder & Hall 2002; Ibrahim et al. 2014).

Although we tested many previously proposed osteological correlates of webbing in this study, a few other plausible (and yet untested) correlates remain. In particular, webbing may modify the force loads acting on webbed finger bones, producing small grooves, tuberosities, or osteohistological features. For example, Madar (2007) noted the presence of dorsolateral flanges on the proximal phalanges of some pakicetids, and although we recovered no significant differences in proximal phalanx proportions between webbed and unwebbed species, there may be subtler differences in phalanx shape that our measurement scheme failed to identify. Three-dimensional geometric morphometrics may help to compare the subtle difference in shape of non-ungual phalanges between webbed and unwebbed species. Histological features offer similar promise. When viewed in thin section, frayed periosteal margins and Sharpey's fibers can help to identify entheses and other osseous sites under tension at a higher resolution (Petermann & Sander 2013). A systematic comparison of phalanx thin sections in extant webbed and unwebbed species may shed light on whether webbing has any similar histological correlates across amniotes.

### Text S12. Additional comments on model predictions for extinct taxa.

*Text S12.1. Model predictions for extinct pan-mammals.*

Our dataset contained 223 pan-mammal specimens, the majority of which had measurable forelimb zeugopodia and acropodia and could thus be fed into our best-performing phybLR model. This latter set of specimens included four extinct non-mammalian synapsids, six monotremes, eight marsupials, and 186 placentals. We summarize our model predictions for extinct species within this set in Figure S16. Briefly, all sampled extinct pan-mammals—including the non-mammalian synapsid taxa *Edaphosaurus* E. D. Cope 1882, *Cabarzia trostheidi* F. Spindler et al. 2019, *Theriodesmus* H. G. Seeley 1888, and *Thrinaxodon* H. G. Seeley 1894; the caniform *Daphoenocyon* J. R. Hough 1948, the stem ungulates *Dromocyon* O. C. Marsh 1876 and *Pachyaena* E. D. Cope 1875, the stem perissodactyls *Theosodon* and *Adinotherium* (both named by F. Ameghino 1887 [cited in Ameghino 1891]), the stem equid *Palaeotherium* G. Cuvier 1804, the rhinoceratoid *Teleoceras* J. B. Hatcher 1894, and the basal artiodactyl *Diplobunops* O. A. Peterson 1919 were confidently identified as retaining terrestrial ties (*i.e.*, as *not* being highly or fully aquatic: one-tailed t-test, $p > 0.05$). Highly or fully aquatic habits have not been seriously considered for the majority of these taxa, so these results are not surprising.

However, a few of these fossil taxa (*Edaphosaurus*, *Teleoceras*, and *Diplobunops*) deserve special consideration. The basal synapsid *Edaphosaurus* shows no obvious morphological adaptations to an aquatic lifestyle, but *Edaphosaurus* specimens are commonly found in depositional environments representing swamps or swampy lakes (Romer 1956), and at least one fully articulated *Edaphosaurus* skeleton has been found in marine deposits (Reisz et al. 1982). Our results suggest that such aquatic paleoenvironments, when recovered, are misleading, and that *Edaphosaurus* likely retained consistent ties to land. *Teleoceras* has been more controversial. The hippo-like proportions of this animal (its relatively short limbs and portentous ribcage) led Osborn (1898) to consider it semi-aquatic. However, isotopic evidence from tooth enamel has suggested a terrestrial lifestyle (Clementz et al. 2008; Wang & Secord 2020). We cannot rule out a semi-aquatic lifestyle for *Teleoceras*, but our best phybLR model confidently rejects the hypothesis that it spent the majority of its time in the water. We thus find morphometric support for the previous claim (based on isotopic evidence) that *Teleoceras* retained ties to land and had at least a semi-terrestrial (if not fully terrestrial) lifestyle. Even so, we encourage future studies to repeat this analysis with a broader sample of rhinoceratoids in order to verify the ubiquity of semi- or fully terrestrial habits in this clade.

Highly or fully aquatic habits have evolved at least four times independently within Mammalia—in the ancestor of sirenians, the ancestor of pinnipeds, the ancestor of extant *Enhydra*[1] spp., and *Cetacea* (Figure S16). We were unable to sample informative stem taxa for any of these lineages. In order to reconstruct the evolutionary history of aquatic habits in these lineages, we encourage future studies to compute their acropod-zeugopodium length ratios and feed these measurements into our best phybLR model to predict their aquatic affinities.

*Text S12.2. Additional comments on model predictions for extinct rhynchocephalians.*

As described in the main-text Results, the aquatic habits and flipper phenotypes predicted by forelimb acropod-zeugopodium length ratios ($L_{hand}$ / $L_{zeugopod}$) for *Vadasaurus* G. S. Bever & M. A. Norell 2017 and *Pleurosaurus* H. Von Meyer 1831 were dependent on tree topology. The table shown inline below summarizes these predictions from our best forelimb acropodium-zeugopodium model by supertree, though all of the actual test results and associated metadata are available in Table S7.

| TREE TOPOLOGY | SQUAMATA TOPOLOGY | *VADASAURUS* HIGHLY OR FULLY AQUATIC? | *PLEUROSAURUS* HIGHLY OR FULLY AQUATIC? | *VADASAURUS* FORELIMB FLIPPERED? | *PLEUROSAURUS* FORELIMB FLIPPERED? |
|---|---|---|---|---|---|
| ***SUPERTREE1*** | **Molecular** | No | No | Yes | Varied by specimen |
| ***SUPERTREE2*** | **Molecular** | No | No | Yes | Varied by specimen |
| ***SUPERTREE3*** | **Morphological** | Yes | Yes | No | No |
| ***SUPERTREE4*** | **Morphological** | Yes | No | No | Varied by specimen |
| ***SUPERTREE5*** | **Molecular** | Yes | Varied by specimen | No | No |
| ***SUPERTREE6*** | **Molecular** | No | Varied by specimen | Yes | No |
| ***SUPERTREE7*** | **Morphological** | No | Varied by specimen | No | Varied by specimen |
| ***SUPERTREE8*** | **Morphological** | Yes | Varied by specimen | Yes | Varied by specimen |

In *Vadasaurus*, flippered forelimbs were recovered more often when assuming the molecular topology of squamates, whereas for *Pleurosaurus*, flippered forelimbs were slightly more common under the morphological topology. For both genera, highly or fully aquatic habits were recovered more often under the morphological topology. Given the generally small effects of these topological changes for phybLR predictions overall (Text S8), it is interesting that they had such a major impact on phenotype predictions for *Vadasaurus* and *Pleurosaurus*, which (as rhynchocephalians) fall outside Squamata. However, given the lack of prediction consistency for these taxa under either the molecular or morphological topologies, it seems there may be more at play affecting their predicted phenotypes. For example, it may be that *Vadasaurus* and *Pleurosaurus* had such liminal acropod-zeugopodium ratios—of 1.40–1.45 (see Table S1),

[1] See Wilson et al. (1991) and references therein for a taxonomic history of the genus *Enhydra*, including nominal authorship.

squarely within both the terrestrial and the aquatic ranges—and were in just such a phylogenetic position to be influenced heavily by the ancestral acropod-zeugopodium ratio of squamates and the ancestral acropod-zeugopodium ratio of Sauria—both of which were highly variable across topologies.

Our ambiguous predictions join a slew of ambiguous morphological and taphonomic evidence for aquatic habits or flippers in these taxa. The precise aquatic habits and flipper phenotypes of both genera have received scarce attention. In addition to its relatively long hands and recovery from a marine limestone, *Vadasaurus* has a number of subtle osteological features suggestive of aquatic habits, including elongated external nares, reduced ossification of the mesopodials and limb long bone epiphyses, and an elongated tail (Bever & Norell 2017). However, aside from these features, the limbs of *Vadasaurus* seem *prima facie* to be generally plesiopedal and capable of supporting its weight on it. *Vadasaurus* retains well defined shoulder and elbow joints, and tall, recurved ungual phalanges that would have sported keratinous claws, making its occasional recovery as "flippered" surprising. *Pleurosaurus* is more derived, and more obviously adapted for an aquatic lifestyle, but its precise aquatic affinity and flipper phenotype present similar difficulties. Highly or fully aquatic habits would be consistent with the recovery of *Pleurosaurus* from marine limestones, its dorsally retracted orbits and external nares (Dupret 2004; Jones 2008), and its highly reduced limbs, which would have complicated overland travel. Like *Vadasaurus*, sampled pleurosaurs have high hand-zeugopodium ratios, within the low range of extant flippered species, but their stylo- and zeugopodia retain cylindrical (rather than flat) shafts (Dupret 2004) and well-defined distal articular surfaces, while their unguals remain tall and recurved (Cocude-Michel 1963), more like those of non-flippered species. Taken together, these features suggest more terrestrially adapted limbs (Benson & Butler 2011) with functional elbow and wrist joints, and keratinous claws that could have manipulated terrestrial substrate. However, while these features would be surprising in a flippered or highly aquatic taxa, they do have a precedent, as some highly aquatic and flippered extant species retain these features as well: Sirenians and turtles retain mobile elbow joints (Wyneken 2001; Cooper 2008), and some flippered trionychids retain dorsoventrally tall, recurved unguals on their anterior digits (pers. obs.). Thus, although *Vadasaurus* and *Pleurosaurus* limbs lack any obvious osteological cues for soft-tissue flippers, we cannot rule out their presence given the variety of flipper osteologies observed in extant species, and our best model presents no decisive evidence for either reconstruction. To investigate this question further, we believe our model could be rerun with a greater taxonomic sample of extinct rhynchocephalians and stem lepidosaurs. More detailed osteological descriptions of the limb osteologies of these taxa would also be highly beneficial for assessing their limb function and likelihood of bearing true flippers.

#### Text S13. Implications of soft-tissue flippers for functional ecology in Triassic marine reptiles.

In the main-text Results, we note that soft-tissue fore-flippers were recovered from our best model (FPR = 0.02) in the following Triassic marine reptile taxa: the eosauropterygian *Keichousaurus hui* C. C. Young 1958; the non-plesiosaurian pistosaur *Yunguisaurus liae* Y.-N. Cheng et al. 2006; all sampled hupehsuchians and Triassic ichthyosauriforms; and the thalattosauroids *Wayaosaurus bellus* X. G. Zhou in G. Z. Yin et al. 2000 and *Miodentosaurus brevis* Y.-N. Cheng et al. 2006. Here we suggest that soft-tissue flipper phenotypes recovered for these taxa may shed light on their functional ecologies, with possible implications for the causes of their extinction.

Soft-tissue flippers integrating the autopodia and zeugopodia tend to facilitate greater or more efficient thrust generation underwater (see Fish et al. 2008; Blob et al. 2016) and have thus often been associated with the potential for a more pelagic lifestyle (Benson & Butler 2011; Gutarra & Rahman 2022), as they free taxa from shallow near-shore environments by enabling longer-distance oceanic travel. Such abundant flipper phenotypes in Triassic taxa may help to refine our understanding of why certain marine reptiles ultimately went extinct. These flippered limbs would have been powerful propulsive agents, calling into question the claim that some of these groups were functionally restricted to nearshore marine settings (Benson & Butler 2011; Kelley et al. 2014). This restriction to shallow waters has previously been implicated in the selective extinction of basal sauropterygians and thalattosaurs in the Middle or Late

Triassic, which coincided with the fall of global sea levels and consequent regression of epicontinental seaways (Figure S17). However, the prevalence of robust flippers in *Keichousaurus*, *Wayaosaurus*, *Miodentosaurus*, and the non-plesiosaurian pistosaurs—all of which are thought to have gone extinct during the latter half of the Triassic (Kelley et al. 2014)—suggests that propulsive limitations played a minimal role in their selective extinction. Instead, it seems likely that their extinction resulted from subtler ecological factors, such as a reliance on benthic food resources, as other authors have suggested (Kelley et al. 2014; Stubbs & Benton 2016).

**Text S14. Additional comments on temporal patterns of aquatic invasion by extinct amniotes.**

When considered in tandem with the most conservative predictions for when major Mesozoic clades first diverged, our predictions about the aquatic habits of extinct species help to clarify the timing and extent of aquatic invasions by extinct species (Figure S17). These results thus inform our understanding of when (and perhaps why) various marine reptiles radiated in the context of the broader Earth-system (Figure S17). For example, as noted in the main text, we confirm that amniotes were restricted from aquatic environments until the latest Carboniferous and note that all aquatic experiments preceding the latest Permian were sparse and partial, never resulting in full commitment to an aquatic lifestyle. This may be due to challenging environmental conditions associated with severe glaciation during the later Paleozoic or competitive exclusion by other marine animals.

In contrast, during the latest Permian and earliest Triassic, amniotes radiated into aquatic environments at a high frequency, with at least nine incipient invasions, and at least three additional commitments to a highly/fully aquatic lifestyle. The latter half of this radiation presumably represents an adaptive burst following the End-Permian Mass Extinction. However, at least two of these aquatic invasions (and perhaps more, as discussed below), must have preceded the End-Permian Mass Extinction's most acute phase (at ~252.3 Ma [Shen et al. 2011]). In particular, the ancestors of *Claudiosaurus germaini* and *Tangasaurinae* (Text S3) independently adapted to a semi-aquatic lifestyle, and both clades are known strictly from the Permian Lower Sakamena Formation (Carroll 1981; Buffa et al. 2025). The precise age of the Lower Sakamena is poorly constrained but has been roughly correlated to the Tatarian stage of the Russian Platform, and the vertebrate-bearing sandstone beds within the Lower Sakamena Formation are likely early Wuchiapingian (see Buffa et al. 2025; see also Henderson et al. 2020). They thus have likely a hard maximum (Wuchiapingian) age of 259.55 Ma and a soft minimum (Tatarian) age of ~257 Ma (Henderson et al. 2020). *C. germaini* and *Tangasaurinae* thus invaded aquatic environments either during or before this interval, and thus several million years before the End-Permian Mass Extinction.

Two other clades that may (or may not) have invaded aquatic environments in the Late Permian are *Omnisauropterygia* and *Ichthyosauromorpha* (see Text S3 for definitions of clade names). There are no *bona fide* omnisauropterygian or ichthyosauromorph fossils known from the Permian (Jiang et al. 2023), and thus no direct evidence that these clades originated prior to the extinction's most acute phase at ~252.3 Ma (Shen et al. 2011). The stratigraphically oldest omnisauropterygian and ichthyosauromorph fossils about whose ages we can be confident are part of the later Early Triassic (Olenekian) Chaohu Fauna of Anhui Province in southeastern China (Jiang et al. 2023), where they were excavated from the Middle and Upper Members of the Nanlinghu Formation, from a unit that likely falls within or straddles the middle–upper Spathian substage of the Olenekian. The Spathian dates from 248.1–246.7 Ma (Ogg et al. 2020), and models of astrochronological cycles detected in carbon isotopes have dated the enaliosaur-bearing unit within the Nanlinghu Formation to 248.8–247.7 Ma (Motani et al. 2017; Jiang et al. 2023). This implies only that the clades *Ichthyosauromorpha* and *Omnisauropterygia* must have originated before 248.8 Ma.

However, it remains possible that these Chaohu marine reptile clades originated earlier. The Chaohu Fauna includes all known nasorostrans, the basal ichthyosauriform or ichthyopterygian genus *Chaohusaurus*, and the eosauropterygian *Majiashanosaurus discocoracoidis* D.-Y. Jiang et al. 2014. These taxa are all highly adapted for an aquatic lifestyle (nasorostrans and *Chaohusaurus* again appear to be highly/fully aquatic), they show a variety of specialized feeding modes (Jiang et al. 2023), and they are not the basal-most members of their respective clades. Thus, there is reason to expect that *Omnisauropterygia* and *Ichthyosauromorpha* originated much earlier. In line with this suspicion, based on the existing stratigraphic ranges for *Sauropterygia* and *Ichthyosauromorpha*, at least three previous morphological clock analyses have placed their origins prior to the Permian-Triassic boundary, in either the latest Carboniferous (Simões et al. 2018) or the Late Permian (Simões et al. 2022; Wang et al. 2022).

Even so, these later Paleozoic origination times should be treated with caution. Motani et al. (2017) showed that intraclade rates of morphological evolution varied greatly across time bins, and that divergence times based on morphological clocks were highly sensitive to the time span over which the rate of morphological evolution was computed. For *Ichthyosauromorpha*, the highest rates of morphological evolution were observed in the earliest-occurring (Spathian) taxa (Motani et al. 2017). Using this highest computed rate of morphological evolution, Motani et al. (2017) predicted a conservative mean divergence time for *Ichthyosauromorpha* of 251.5 Ma (range 246.2–253.4 Ma), which would suggest (but not imply) that the clade postdates the End-Permian Mass Extinction. This timing has been supported by Jiang et al. (2023), who note that the warming seas at the beginning of the Early Triassic would be more hospitable to air-breathing ichthyosauromorph predators than the colder waters more typical of the later Permian. This timing assumes exceptionally high rates of morphological evolution immediately following the End-Permian Mass Extinction (six times higher than the average computed by Motani et al. 2017 for all of *Ichthyosauromorpha*), but such rates may well have been plausible in the post-extinction environment.

Thus, depending on when the progenitors of *Omnisauropterygia* and *Ichthyosauromorpha* first took to the water (taken together with *Tangasaurinae* and *Claudiosaurus*), 2–4 independent amniote invasions of aquatic environments took place in the Late Permian. One trigger for these invasions could have been the final melting of Permian glaciers 260–255 Ma (Montañez & Poulsen 2013; Marcilly et al. 2022), which would have prompted a major burst in shallow marine productivity and opened new niches formerly vacated by ice cover. Another set of triggers could have been the precipitous drop in global sea level during the latest Permian (Marcilly et al. 2022), and the rapid closure of the Paleotethys ocean in southeastern Asia (Scotese & Langford 1995), both of which would have split formerly continuous intercontinental seaways into isolated inland seas, creating various ecological incubators for marine experimentation. The subsequent break-up of Pangaea in the Mesozoic (Dietz & Holden 1970; Seton et al. 2012) and associated climatic changes and large-scale habitat fragmentations may have then facilitated the rapid diversifications of the major clades of Triassic marine reptiles that survived the End-Permian mass extinction. At present, such Earth-system connections to marine reptile diversification remain necessarily vague and speculative. Resolving more precisely the evolutionary patterns associated with marine colonization in pan-reptiles, particularly within the Permian and Triassic, will require additional study of the precise stratigraphic ranges of *Tangasaurinae* and *Claudiosaurus*, and the recovery of earlier Triassic (Induan) or Permian (Lopingian) fossils referable to either *Omnisauropterygia* or *Ichthyosauromorpha*.

### Text S15. To what extent are webbing, flippers, and aquatic habits related?

We have found morphometric correlates of flipper form and aquatic habits but not of interdigital webbing. This may seem counterintuitive, given that webbing is often canonically associated with aquatic adaptation (Thewissen & Taylor 2007) and may be a common transitional stage in the developmental evolution of flippers (Cooper et al. 2018). Surprisingly, however, we find that both flippers and aquatic habits have diagnostic limb morphometries, whereas webbing does not. This disconnect may result from the diversity of interdigital webbing forms and functions among amniotes (Figure S2; Thewissen & Taylor 2007), but soft-tissue flippers also display a variety of forms and functions, and our models can recognize them regardless. An alternative explanation for this disconnect may be that all flippered species live in the water, whereas many webbed species live on land. Interdigital webbing does abound in terrestrial amniotes—hyraxes, bats, colugos, various terrestrial rodents, some ursids, many felids and terrestrial mustelids, several species of gecko, and the extinct pterosaurs all have some degree of interdigital webbing (Figure S2; personal observation of specimens in Table S1). However, within our current dataset, the majority of terrestrial species lacked webbing (Figure S18a,b), and webbing had a significant positive association with aquatic habits (Figure S18c–f: Fisher's exact tests, each $p < 0.05$), suggesting that terrestrial webbed species alone fail to explain the observed dissociation.

The remainder of the explanation for this disconnect may relate to the *extent* to which species in our dataset inhabit aquatic vs. terrestrial environments, which differed greatly between webbed and flippered taxa. Most webbed species were either transiently or moderately aquatic, whereas all flippered species were either highly or fully aquatic (Figure S18c–f). Highly or fully aquatic habits accordingly had a more than five-fold stronger association (as captured by Cramer's V statistic) with flippers than they did with interdigital webbing (Figure S18c–f). This is notable, as transiently and moderately aquatic species had far less distinct limb proportions from their terrestrial counterparts than did highly/fully aquatic species (Figure 2f; Figure S8c). As a result, the observed contrast in morphometric signal between webbing and flippers probably corresponds in part to the contrast between transiently/moderately aquatic and highly/fully aquatic habits. This contrast, in turn, corresponds to the disconnect between species partially committed and species fully or near-fully committed to life in the water. Webbed species are more often "on the fence" between terrestrial and aquatic habits, whereas flippered and highly/fully aquatic species are more often "all in" (Blob et al. 2016). The conflicting demands of terrestrial and aquatic locomotion force amphibious and semi-aquatic taxa to make functional trade-offs in their behavior and morphology that allow them to traverse both media (Blob et al. 2016). These trade-offs are less pronounced in highly aquatic taxa, and altogether absent in fully aquatic species that never voyage onto land. Key among these trade-offs is the need, retained in amphibious and semi-aquatic species, for terrestrially proportioned limbs that can bear weight on land and manipulate terrestrial substrate. This requirement maintains terrestrially specialized limb joints (Coombs 1978) and elongate zeugopodials that should support efficient lever mechanics on land. As a result, amphibious and semi-aquatic species maintain the functional constraints imposed on limb morphometry by terrestrial locomotion. And because most webbed aquatic species have amphibious habits, they should retain both more terrestrial and more constrained (less variable) limb proportions. We believe this retained evolutionary trade-off may amount to a functional constraint that prevents webbed species from acquiring their own diagnostic morphometric profile.

#### Text S16. Additional comments on amniote flipper morphometry.

In the main-text Discussion, we noted that flippered taxa have relatively longer acropodial regions than their non-flippered counterparts in every amniote lineage with flippered and non-flippered representatives. It is worth clarifying here that flippered species do not of course have *universally* longer acropods than non-flippered species. Indeed, there is overlap in the limb proportions of flippered and non-flippered taxa (visible in Figure 2f and Figure 4). Even so, flippered species do have longer acropods than their closest non-flippered relatives. For example, sirenians, beaked (i.e., ziphiid) whales, fin whales (*Balaenoptera* [B. G. E. De La Cépède 1804] spp.), and trionychids have hand-zeugopodium ratios that fall squarely within the non-flippered range of most amniotes (sirenians: 0.99–1.16; *Mesoplodon* [P. Gervais 1852] and *Balaenoptera* spp.: 0.84–1.00; trionychids: 1.55–1.64). However, these groups still have higher hand-zeugopodium length ratios than most non-flippered ungulates (mean: 0.81), atlantogenatans (mean: 0.64), and turtles (mean: 0.69). As a result, these relatively short-handed flippers still conform to the broad morphometric pattern observed across clades.

Moreover, in addition to having relatively longer acropodium regions, we found that flippered and highly/fully aquatic species also have a significantly greater *range* of relative acropodium lengths (Figure 2f; Figure S6; Figure S7a–b; Figure S8c). Indeed, the hand-to-zeugopodium length ratios of flippered species ranged from 0.89–10.6, a nearly ten-fold increase from the non-flippered range of 0.11–1.93 (Figure 4c). Amniotes with flippers thus invade both the shared terrestrial-aquatic region of morphospace and new swaths of morphospace unexplored by terrestrial species. This result supports the recent finding that aquatic mammals have more disparate limb morphometries than terrestrial species (Rothier et al. 2024) and reveals the same trend in reptiles for the first time, demonstrating that aquatic habits dramatically increase limb diversity across amniotes. This disparity of limb configurations may hint at distinct evolutionary landscapes for limbs that are adapted to aquatic and terrestrial media (Text S15). In terrestrial amniotes, cursorial habits are associated with the accentuation of hinge-like limb joints, and the parallel elongation of the zeugo- and acropodia (Coombs 1978). In contrast, among highly or fully aquatic amniotes, movement speed, maneuverability, and efficiency tend to favor reduced joint movement and anti-correlated zeugo- and autopodial length changes. This should promote the evolutionary dis-integration of limb segments, allowing for a greater range of limb proportions in flippered species. Such an interpretation lends support to recent anatomical network analyses (Fernández et al. 2020), in which the limbs of flippered species explored patterns of anatomical integration unseen among terrestrial taxa. Additional work could test this prediction more directly by comparing patterns of evolutionary integration (Goswami & Finarelli 2016) among limb segments between flippered and non-flippered species, to see if relatively dis-integrated limb segments account for the great range of morphospace explored by flippered clades.

Our predictions of soft-tissue fore-flippers in many taxa whose flipper phenotypes were previously overlooked or understudied (e.g., hupehsuchians, *Keichousaurus*, derived askeptosauroids) also suggest new avenues of research into the morphological and functional variety exemplified by reptilian flippers. For example, our results lend new functional significance to the massive proximal ulnar prominence of *Keichousaurus* (Carroll & Gaskill 1985), which may have served as a spar for the soft-tissue base of the flipper and a lever for elbow extension and digital flexion in a manner similar to the olecranon process in sea lions (English 1977). Likewise, the presence of soft-tissue flippers in hupehsuchians, and some derived askeptosauroids should lead us to re-evaluate the functional osteology and myology of their limb skeletons assuming a continuous soft-tissue zeugoautopodial system facilitates thrust generation in these swimmers.

### Text S17. Possible developmental factors constraining flipper form in mammals.

In the main-text Discussion, we discussed how differences in the flexibility of *HoxA* and *HoxD* cluster patterning during early limb morphogenesis may account for the reduced morphometric breadth of mammalian limbs when compared to those of reptiles. Several extinct flippered reptiles (including plesiosaurs, ichthyosaurs, mosasaurians, and metriorhynchids) exhibit reduced perichondral ossification in the radius and ulna, which "blur[s]" the autopod-zeugopodium boundary by degrading the long-bone identity and associated posterior sculpturing of zeugopodials (Caldwell 2002; Maxwell et al. 2014). This process, called "mesopodialization" (Wagner & Chiu 2001), likely enabled many extinct marine reptiles to hyper-truncate the zeugopodium (Caldwell 2002). Previous studies have linked mesopodialization to *HoxA*/HoxD cluster patterning in the early limb bud (Wagner & Chiu 2001; Caldwell 2002; Maxwell et al. 2014). This mechanism is intuitive, because *HoxA*/*HoxD* genes specify limb segment (stylopodium, zeugopodium, mesopodium, and acropodium) identity in a collinearly restricted sequence along the limb's proximodistal axis (Wolpert et al. 2011). In addition, existing experimental data support the plausibility of this mechanism in extinct flippered species. In *hoxa-11/hoxd-11* double-mutant mice, for example, zeugopodia have stunted growth and less clearly defined shaft margins (Davis et al. 1995). Likewise, in *hoxd-13* mutant mice and people, palm bones become short and nodular, more like mesopodial elements (Muragaki et al. 1996; Horsnell et al. 2006). The presence of similarly mesopodialized metapodial elements, acropodial elements, and (especially) zeugopodial elements in various extinct flippered reptiles (e.g., metriorhynchids, mosasaurians, plesiosaurs, and ichthyosaurs), and the absence of mesopodialized zeugopodia in any living or extinct mammals (Figure 4a), make this developmental mechanism a plausible and compelling explanation for the latter's limited morphometric range. In theory, functional tests knocking down early *hoxa-11/hoxd-11* expression in the reptilian limb bud (and observing resultant limb osteologies during later developmental stages) could test this hypothesis. However, until such additional experiments are performed, we lack rigorous developmental evidence for this hypothesis within a closed extant phylogenetic bracket that encompasses the extinct flippered reptiles of interest.

A related factor that might influence the relative tendency of mammals and reptiles to mesopodialize is the clade-specific distribution of epiphyseal growth plates in limb long bones. In therian mammals, physiological signals tend to promote epiphyseal-diaphyseal fusion and cessation of growth—

i.e., they have secondary ossification centers, whereas reptiles, ancestrally, do not. The secondary ossification centers (SOCs) of long bones may have evolved to make articular surfaces less sensitive to heavy mechanical loads on land (Xie et al. 2020), though there may be more plausible alternative explanations. To that point, while SOCs are present ancestrally in therians, some (but not all) crown cetaceans have secondarily reduced their SOCs (Xie et al. 2020). In contrast, pan-crocodylians, turtles, and (presumably) ancestral saurians lack(ed) SOCs, which might help to explain the reduced posterior sculpting of the ulna in flippered ichthyosaurians, pistosaurians, *Keichousaurus*, metriorhynchids, and turtles (Figure 4a). That said, SOCs are likely ancestral for lepidosaurs (Gauthier et al. 1988b; Regnault et al. 2016; Frýdlová et al. 2020). This is notable, as many mosasaur flippers (including those of *Tylosaurus* and *Ectenosaurus* D. A. Russell 1967, shown respectively in Figure S1 and Figure 4a) have less clearly mesopodialized zeugopodia that retain sculpted posterior margins. This may help to explain why the highly diverse mosasaurians never attained the same range of limb proportions as flippered euryapsid reptiles (Figure 4a), but it fails to explain how mosasaurians manage to invade regions of limb morphospace beyond flippered mammals. Thus, while the presence of epiphyseal growth plates in mammals may serve as an additional constraint against loss of zeugopodial long bone identity, it does not suffice on its own explain the reduced range of limb region proportions observed among pan-mammals.

An additional, perhaps independent developmental factor that might constrain limb development concerns the length of individual distal phalanges. Mammals in our dataset had a notably narrower range of ungual proportions (as captured by $L_{(\text{mD3 ungual})} / H_{(\text{mD3 ungual})}$) than reptiles, although the greater part of the reptile range was taken up by chelonioids, whose D3 unguals are exceedingly long and tapered. This was noted in Figure 4a, as even reptiles that retain zeugopodial long-bone identity and the associated posterior sculpting of the ulna (e.g., *Chelonia mydas* C. Linnaeus 1758 and *Ectenosaurus*) have higher acropod-zeugopodium length ratios than all sampled mammals, in part because they drastically elongate the distal (ungual and penultimate) phalanges, to a greater extent than any observed pinnipeds or sirenians. This casts doubt on differences in the plasticity of zeugopodial long bone form (due to some combination of *HoxA*/*HoxD* patterning, the presence of epiphyseal growth plates, and other developmental constraints) as the sole explanation for the limited range of mammalian limb region proportions.

Of course, in the absence of additional experimental data in reptiles, these explanations remain untested. A combination of these factors, and myriad alternative developmental or functional anatomical factors as-yet unconceived (e.g., pertaining to the contrasting architecture of appendicular musculature in mammals and reptiles), might well constrain mammalian limb morphometry in ways we have yet to understand. Regardless, the morphometric patterns observed herein suggest that some set of constraints has limited the range of relative acropodium and zeugopodium lengths available to mammals.

#### Text S18. Note on usage of binomial testing in tandem with ROC analysis.

Binomial testing is a standard approach to testing the likelihood of an outcome assuming an evenly distributed sample with binary output (Clopper & Pearson 1934). In contrast to the AUC value resulting from an ROC analysis, binomial tests do not test for sensitivity and specificity of a model, but rather accuracy of the predictions. While AUC measures how likely a classifier is to rank positive instances higher than negative instances (Fawcett 2006), a binomial test in this context assesses the correct classification rate of a given model or set of models. We employ one-tailed binomial tests to determine whether the following conditions hold:

- We test whether the correct classification rate of a given phylogenetic binomial logistical regression (phybLR) model, with classification results pooled across classification thresholds, is significantly higher than would be expected by random chance (50%).

- We test whether the correct classification rate of a given phybLR model, with classification results pooled across classification thresholds, is significantly higher than 75%.

- We test whether the correct classification rate of a given "best" phybLR model (that with the classification threshold that maximizes Youden's J-statistic [Youden 1950]) has a correct classification rate significantly higher than 75%.

A limitation with this approach is that binomial tests typically do not account for skewness of the data, making them most useful for datasets that have an even distribution of correct category classifications. For example, if 90% of a sample are assigned to category A and only 10% to category B, a model always predicting category A would have a 90% accuracy according to binomial tests, although the model itself does not represent the real data. However, measuring model classification performance is an intuitive measure to assess overall predictability of our data set and can be, in combination with other model assessment strategies, highly informative. We therefore employ one-tailed binomial tests not as a substitute, but a complementary measure to ROC analyses to analyze performance of our phybLR models. However, a more ROC-specific statistical testing approach—using, for example, DeLong's test (DeLong et al. 1988) to compare the AUCs of competing consensus Receiver Operating Characteristic curves (cROCCs)—would provide an effective complement to our technique for model comparison across thresholds.

### Text S19. Extended acknowledgments and image attributions.

This project would not have been possible without access to hundreds of specimens and dozens of high-resolution specimen images from multiple institutions. We are therefore indebted to the following institutions and people who provided access to physical specimens or specimen photographs for linear and geometric morphometric analyses: Gregory Watkins-Colwell, Kristof Zyskowsky, Vanessa Rhue, Daniel Brinkman, Alex Ruebenstahl, and Marilyn Fox (Yale Peabody Museum of Natural History); Eleanor Hoeger, Sara Ketelsen, Marisa Surovy, Carl Mehling, Roger Benson, David Kizirian, and Amelia Zietlow (American Museum of Natural History); Amy Henrici, Matt Lamanna, Alyssa McNece, and Andrew McAfee (Carnegie Museum of Natural History); Michael Caldwell (University of Alberta); Ryosuke Motani (University of California Davis); John Ososky, Michael McGowen, Michael Brett-Surman, and George Robert King (National Museum of Natural History, Smithsonian Institution); Torsten Scheyer and Christian Klug (Paläontologisches Institut und Museum Universität Zürich); Christina Ifrim (Jura Museum Eichstätt); Sven Sachs (Naturkunde-Museum Bielefeld, Abteilung Geowissenschaften); Cecilia Apaldetti (Instituto y Museo de Ciencias Naturales, Universidad Nacional de San Juan); ReBecca Hunt-Foster (Dinosaur National Monument); Keqin Gao, Da-yong Jiang, and Jun Chai (Peking University); Wei Wang (Chinese Academy of Sciences); Zichen Fang (China University of Geosciences); Tamaki Sato (Kanagawa University); Giovanni Serafini (Università degli Studi di Modena e Reggio Emilia); Annalisa Aiello (Museo Civico di Scienze Naturali Bergamo); Xiao-Chun Wu (Canadian Museum of Nature); Jamie Pescatore (Loggerhead Marinelife Center); Jun Liu and Guang-Hui Xu (Institute of Vertebrate Paleontology and Paleoanthropology, Chinese Academy of Sciences); Eric Metz and John Scannella (Museum of the Rockies); Tom Deméré (San Diego Natural History Museum); Tyler Keillor, Matteo Fabbri, Alan Resetar, and William Simpson (Field Museum of Natural History); Chris Beard, Megan Sims, Ana Motta, and Rich Glor (Kansas University Biodiversity Institute and Natural History Museum); Marc Jones (Natural History Museum, London); Zaituna Skosan and Devonne Kortje (Iziko SAM); Peggy Vincent, Lilian Zazes, and Nour-Eddine Jalil (Muséum National D'Histoire Naturelle); Julien Kimmig and Jannik Weidtke (Staatliches Museum für Naturkunde Karlsruhe); Gunnar Riedel, Rainer Brocke, and Sven Tränkner (Senckenberg Naturmuseum Frankfurt); Yu-nan Wang (Shanghai Science and Technology Museum); Lida Xing (China University of Geosciences); Phil Bell (University of New England); Ronny Maik Leder (Naturkundemuseum Leipzig); Erin Maxwell (Staatliches Museum für Naturkunde, Stuttgart); Oliver Wings (Natural History Museum, Bamberg); Sylvia Humphrey (Great North Museum, Hancock); Luca Simonetto (Museo Friulano di Storia Naturale); Meghan Cohorst (Save the Manatee Club); Francois Therrien (Royal Tyrrell Museum of Palaeontology); Cristiano Dal Sasso (Museo di Storia Naturale di

Milano); Neal Immega (Houston Gem and Mineral Society); and Mark McMenamin (Mount Holyoke College). We are also grateful to Michael Durham (Oregon Zoo), Thomas Odermatt (Tom's Pet Supply), and Matt Shetzers (Shetzers Photography), all of whom provided access to live animal photographs, either for use in Figure S2 or to facilitate limb phenotype scoring.

In addition, we thank the following people and institutions for granting access to micro-computerized tomography (µCT) scans on MorphoSource: Stephanie Baumgart, Paul Sereno, Tyler Keillor, Lauren Conroy, Daniel Vidal, Kate Webbink, Adam Ferguson, and Janeen Jones (Field Museum of Natural History); David Blackburn, Edward Stanley, and Jaimi Gray (Florida Museum of Natural History); Stevie Kennedy-Gold and Joe Martinez (Museum of Comparative Zoology, Harvard University); Hilary Ketchum, Mark Carnall, and Duncan Murdock (Oxford University Museum of Natural History); Emily Braker (University of Colorado Boulder Museum of Natural History); Kevin Conway (Texas A&M University); Douglas Boyer (University of Nebraska State Museum); Laurel Lamb (University of Arkansas Museum); Carol Spencer, Michelle Koo, and Carla Cicero (Museum of Vertebrate Zoology, UC Berkeley); Cody Thompson and Gregory Schneider (University of Michigan); Lachlan Hart (University of New South Wales, Sydney); Emma Schachner (Louisiana State University Health Sciences Center New Orleans); the Louisiana State University & Agricultural and Mechanical College; the University of Kansas Center for Research, Inc.; the Duke Lemur Center Museum of Natural History; the University of Michigan Museum of Zoology; and the California Academy of Sciences Division of Herpetology.

The researchers would again like to thank the National Science Foundation, Yale Institute for Biospheric Studies, and the IUCN Crocodile Specialist Group for funding this research and its lead author. We also thank the Society of Vertebrate Paleontology, the Geological Society of America, the Society of Integrative and Comparative Biology, and the International Meeting on the Secondary Adaptation of Tetrapods to Life in Water (SECAD) for allowing us to present our preliminary findings at their annual or quadrennial conferences. Their enthusiasm for this project kept us going!

Digital coding and image resources require additional thanks and attributions. BioRender.com's exceptional built-in icon library made it possible to construct final figures that matched our vision for the paper. For Figure S2, all photographs were taken by the lead author with the exception of the *Bradypus* hand, which was photographed by Matt Shetzers (Shetzers Photography). ChatGPT (versions 3.0 and 3.5) was used to write or assist in writing ~100 lines of code in scripts S1, S2, and S4, as noted therein, and to brainstorm methods for performing phylogenetic correlation tests. The photorealistic silhouettes in main-text Figs. 1 and 6c were designed in Adobe Photoshop 2024 as follows: PhyloPic silhouettes for *Tylosaurus* sp. (credit: Scott Hartman, CC Attribution 3.0 Unported: https://www.phylopic.org/images/8ae20f15-d9c1-45ff-a117-77f001b6eb0a/tylosaurus), *Rhomaleosaurus cramptoni* (credit: Scott Hartman, 2019, Attribution 3.0 Unported, with adjustments made to original silhouette proportions: https://www.phylopic.org/images/a3f68690-f6b5-4265-89bd-36363e8d64ea/morturneria-seymourensis), *Ichthyosaurus communis* (credit: Jagged Fang Designs, 2021, CC0 1.0 Universal Public Domain Dedication: https://www.phylopic.org/images/447bec69-acdf-473c-8df7-4f2f79040bd0/ichthyosaurus-communis), and *Stenella coeruleoalbus* (credit: T. Michael Keesey, Attribution-ShareAlike 3.0 Unported: https://www.phylopic.org/images/f00a3f93-cd4b-48d4-bc25-1482eadd1bef/stenella-coeruleoalbus) were cropped, stunningly colored and textured by Raquel Jaramillo, and then blackened and overlain with stylized fore-flipper skeletons by Caleb M. Gordon, who adapted these skeletons from existing specimen images of FHSM VP 0003 (*Tylosaurus proriger*, photo credit: Amelia Zietlow), NHM PV R 2864 (*Muraenosaurus* sp., photo credit: Marc Jones), SMF R 4166 (*Stenopterygius quadriscissus*, photo credit: Sven Tränkner, Senckenberg Museum Frankfurt), and USNM 593987 (*Stenella coeruleoalba*, photo credit: John Ososky and the Smithsonian Whale Collection Database; EZID: http://n2t.net/ark:/65665/3fd11b878-e9c3-43b9-be2b-0eebd2bd48a9). Similarly colorized limb skeletons from Figure 4 and Figure S1 were adapted or redrawn from images of the following specimens, with image attributions given in Table S1: AMNH MAM 63947, CM 30781, FMNH 22066, FMNH PR2378, FHSM VP 401, IVPP V3232, SD 25573, SMF R4944, USNM 550853, USNM 593894, USNM 594165, Shanghai Natural History Museum specimen 20140727, AGM MT 10010 (new catalog number unknown; see Table S1 for specimen details), YPM VP 001130 (photo by

Division of Vertebrate Paleontology, Yale Peabody Museum, 2017: https://collections.peabody.yale.edu/search/Record/YPM-VP-001130), and a published but currently uncatalogued NMW specimen of *Aigialosaurus buccichi* (see Table S1). Two additional colorized limb skeletons were adapted or redrawn from reconstructions of *Pakicetus* (as shown in Muizon 2009, Fig. 2, original photograph image by H. J. G. Thewissen [article accessible via https://doi.org/10.1016/j.crpv.2008.07.002]) and the stem cetacean forelimb SNTB 2011-01 (as shown in Vautrin et al. 2020, Fig. 4 [https://doi.org/10.18563/journal.m3.92]).

Figure creation for this manuscript was also greatly enhanced by the availability of taxon silhouettes downloaded from PhyloPic.org, and we thank T. Michael Keesey for making this resource available to the public. For Figures 2–4, Figure S1, and Figures S14–S17, taxon silhouettes were downloaded from PhyloPic.org with the following attributions: *Karpinskiosaurus secundus* (by T. Michael Keesey, based on illustration by Dmitry Bogdanov, Attribution 3.0 Unported: https://www.phylopic.org/images/24c49c6a-0052-4004-b571-b0ba2cb1d073/karpinskiosaurus-secundus), *Limnoscelis paludis* (by Diego Alexander Canque Sosa, based on original illustration by Nobu Tamura, Attribution-ShareAlike 3.0 Unported: https://www.phylopic.org/images/67c1695a-19d2-42e9-b9e5-0a090ffc18a9/limnoscelis-paludis), *Edaphosaurus* (by T. Michael Keesey, based on illustration by Nobu Tamura, Attribution-ShareAlike 3.0 Unported: https://www.phylopic.org/images/2969e396-ed30-439c-8e2a-3fa94903774e/edaphosaurus), *Cabarzia trostheidei* (by Zak Lewis, based on work by Nix Illustration, Attribution-NonCommercial 3.0 Unported: https://www.phylopic.org/images/9900614e-d86c-4f8b-9c3a-1d1056a62118/cabarzia-trostheidei), *Scylacosaurus* sp. (used for *Theriodesmus*; by T. Michael Keesey, Public Domain Mark 1.0: https://www.phylopic.org/images/6ecdac12-8832-401b-b6e2-8474dbc41b6a/scylacosaurus), *Thrinaxodon liorhinus* (by Christine Axon, CC0 1.0 Universal Public Domain Dedication: https://www.phylopic.org/images/94fda119-5435-4246-ad23-20b15f3bcb8d/thrinaxodon-liorhinus), *Mesonyx obtusidens* (used for both *Pachyaena* and *Dromocyon* / *Synoplotherium*; by Mimics (Julián Bayona), Attribution-ShareAlike 3.0 Unported: https://www.phylopic.org/images/73624419-f6f1-41a7-becf-7f1c90d5b203/mesonyx-obtusidens), *Ornithorhynchus anatinus* (by Sarah Werning, Attribution 3.0 Unported: https://www.phylopic.org/images/b406c409-2735-4a3d-a7aa-8afe0b6e72dc/ornithorhynchus-anatinus), *Didelphis virginiana* (used for Marsupialia; by Gabriela Palomo-Munoz, Attribution-NonCommercial 3.0 Unported: https://www.phylopic.org/images/f9f6ae2b-ed15-4e25-976c-ab4e15e0fe41/didelphis-virginiana), *Tamandua mexicana* (used for Xenarthra; by Xavier Jenkins, CC0 1.0 Universal Public Domain Dedication: https://www.phylopic.org/images/6df900f7-0a90-47ca-88e2-99d901814c22/tamandua-mexicana), *Tenrec ecaudatus* (used for Afroinsectophilia; by Yan Wong, CC0 1.0 Universal Public Domain Dedication (https://www.phylopic.org/images/7777eeb6-87bd-48c2-ba02-15cc8e2bfac9/tenrec-ecaudatus), *Dendrohyrax* sp. (used for Procaviidae; by Steven Traver, CC0 1.0 Universal Public Domain Dedication: https://www.phylopic.org/images/bae84fd8-937a-4390-b123-0a22bfcd7df7/dendrohyrax), *Loxodonta africana* (used for Proboscidea; by Chuanxin You, CC0 1.0 Universal Public Domain Dedication: https://www.phylopic.org/images/910d853a-1a15-4953-a1d3-b81208994d35/loxodonta-africana), *Pezosiren portelli* (by T. Michael Keesey, CC0 1.0 Universal Public Domain Dedication: https://www.phylopic.org/images/ba8c3e75-fdc2-489a-b331-50024fe0ee08/pezosiren-portelli), *Trichechus manatus* (by T. Michael Keesey based on work by Vince Smith, Attribution-NonCommercial-ShareAlike 3.0 Unported: https://www.phylopic.org/images/12597d6c-9c80-4510-9826-975eabddc597/trichechus-manatus), *Rattus norvegicus* (used for Euarchontoglires; by Rebecca Groom, Attribution-ShareAlike 3.0 Unported: https://www.phylopic.org/images/00d12cef-0904-4bf8-a915-901feeecbb2e/rattus-norvegicus), *Solenodon paradoxus* (used for Eulipotyphla; by Inessa VOET, CC0 1.0 Universal Public Domain Dedication (https://www.phylopic.org/images/3717bf88-d959-4bdd-aa69-54e3dccc53b6/solenodon-paradoxus), *Manis culionensis* (by Steven Traver, CC0 1.0 Universal Public Domain Dedication: https://www.phylopic.org/images/f628f6fe-6e2d-4173-a20f-265dc7d2d04f/manis-culionensis), *Urocyon litorali santacruzae* (by T. Michael Keesey based on work by Björn S, Attribution-NonCommercial 3.0 Unported: https://www.phylopic.org/images/f1a044ce-86f4-4395-a85a-ba6dd0e5492c/urocyon-littoralis-santacruzae), *Neofelis nebulas* (used for Aeluroidea; by Margot Michaud, CC0 1.0 Universal Public

Domain Dedication: https://www.phylopic.org/images/37dc018a-f356-4a26-91bb-e5c522a89f47/neofelis-nebulosa), *Taxidea taxus* (by Gabriela Palomo-Munoz, Attribution 4.0 International: https://www.phylopic.org/images/8a3105db-4928-47ab-aaf6-10a96dc8504b/taxidea-taxus), *Galictis vittata* (used for Mustelinae; by Margot Michaud, CC0 1.0 Universal Public Domain Dedication: https://www.phylopic.org/images/3e13e4f5-fd4d-4a0f-9df1-5cfcdf4c57ed/galictis-vittata), *Aonyx cinereus* (by Margot Michaud, CC0 1.0 Universal Public Domain Dedication: https://www.phylopic.org/images/95ead4b3-34c0-461a-97fc-827c10d9dc70/aonyx-cinereus), *Enhydra lutris* (by Margot Michaud, CC0 1.0 Universal Public Domain Doedication: https://www.phylopic.org/images/d3a78afb-1b9e-45e0-b6f4-144d79f399f0/enhydra-lutris), *Puijila darwini* (by T. Michael Keesey, CC0 1.0 Universal Public Domain Dedication: https://www.phylopic.org/images/1ffc19b2-82f4-4715-9c85-6f6095bb9eb5/puijila-darwini), *Lutra lutra* (by Margot Michaud, CC0 1.0 Universal Public Domain Dedication: https://www.phylopic.org/images/3cdae098-058b-4ce2-8f46-ad3101b624db/lutra-lutra), *Lontra canadensis* (by Margot Michaud, CC0 1.0 Universal Public Domain Dedication: https://www.phylopic.org/images/51510f2d-8803-4386-89b3-b04e90a32c4d/lontra-canadensis), *Zalophus japonicus* (by Margot Michaud, CC0 1.0 Universal Public Domain Dedication: https://www.phylopic.org/images/6b7f24dc-6525-4263-b480-6f93f9a7a31d/zalophus-japonicus), *Palaeotherium magnum* (by Michael Tripoli, Attribution 4.0 International: https://www.phylopic.org/images/317ff59a-363e-4559-a3ea-c4566575ccea/palaeotherium-magnum), *Adinotherium* sp. (by Zimices (Julián Bayona), Attribution-NonCommericial 3.0 Unported: https://www.phylopic.org/images/28e0a139-de00-4a8a-ad69-47e74d8d22ea/adinotherium), *Theosodon* sp. (by Zimices (Julián Bayona), Attribution-ShareAlike 3.0 Unported: https://www.phylopic.org/images/c9c5af3a-1fed-4abf-a734-3473f89cae18/theosodon), *Camelus bactrianus* (by T. Michael Keesey based on work by Lukasiniho, Attribution-NonCommercial-ShareAlike 3.0 Unported: https://www.phylopic.org/images/918ad1f7-fc6e-43f5-9708-b2da8a71a820/camelus-bactrianus), *Tapirus augustus* (by Zimices (Julián Bayona), Attribution 3.0 Unported: https://www.phylopic.org/images/21142af0-58f1-494d-8b2c-b2d9d0eada4c/tapirus-augustus), *Phacochoerus africanus* (by T. Michael Keesey based on work by Jan A. Venter, Herbert H. T. Prins, David A. Balfour & Rob Slotow, Attribution 3.0 Unported: https://www.phylopic.org/images/d594a0c4-2708-4cde-ba41-08fde8b1184f/phacochoerus-africanus), *Bos bison* (used for Ruminantia; by T. Michael Keesey based on work by Lukasiniho, Attribution-NonCommerical-ShareAlike 3.0 Unported: https://www.phylopic.org/images/9257e657-c494-450d-b8d4-2ef6fd94f81f/bos-bison), *Mesohippus* sp. (used for Equidae; by T. Michael Keesey, based on work by Heinrich Harder, Public Domain Mark 1.0: https://www.phylopic.org/images/de4f0606-7e53-4439-8105-797546c7d1c4/mesohippus), *Hyracodon* sp. (by T. Michael Keesey, based on work by Charles R. Knight, Public Domian Mark 1.0: https://www.phylopic.org/images/a53719a5-267a-425e-8b4b-a0f46604a73e/hyracodon), *Teleoceras dossier* (by Zimices (Julián Bayona), Attribution-ShareAlike 3.0 Unported: https://www.phylopic.org/images/ebbf856d-bd68-4c3e-b177-7b471586a793/teleoceras-fossiger), *Rhinoceros unicornis* (by T. Michael Keesey based on work by H. F. O. March, Public Domain Mark 1.0: https://www.phylopic.org/images/abad209c-2ca1-423b-a66b-317ca52f4c78/rhinoceros-unicornis), *Promerycochoerus carrier* (used for Diplobunops; Zimices (Julián Bayona), Attribution 3.0 Unported: https://www.phylopic.org/images/0b68ecc3-2aa0-4a34-b97d-5179d68be658/ promerycochoerus-carrikeri), *Leptomeryx* evens (by T. Michael Keesey, based on work by Nobu Tamura, Attribution 3.0 Unported: https://www.phylopic.org/images/e69e1841-7a09-4936-bc94-19ca766cba84/ leptomeryx-evansi), *Amphicyon ingens* (used for Daphoenocyon; by T. Michael Keesey based on work by Rom-diz, Public Domain Mark 1.0: https://www.phylopic.org/images/33ada84e-026c-4af8-9de3-01eade53d730/amphicyon-ingens), *Hippopotamus amphibius* (by T. Michael Keesey, based on work by Jan A. Venter, Herbert H. T. Prins, David A. Balfour & Rob Slotow, Attribution 3.0 Unported: https://www.phylopic.org/images/6336f90c-8f02-48f5-94d1-1d85c0100473/hippopotamus-amphibius), *Choeropsis liberiensis* (by T. Michael Keesey based on work by Marek Velechovsky, Attribution 3.0 Unported: https://www.phylopic.org/images/72c02812-d2c8-4d0d-83ce-3f0677a01ace/choeropsis-liberiensis), *Ambulocetus natans* (by T. Michael Keesey, based on original illustration by Nobu Tamura, Attribution 3.0 Unported:

https://www.phylopic.org/images/70382e3e-de05-49da-a542-ba9c13ff4bfd/ ambulocetus-natans), *Pakicetus* (by T. Michael Keesey, based on illustration by Conty, Attribution 3.0 Unported: https://www. phylopic.org/images/c1cbc4b7-1d87-4461-a818-6d3a23feb7d6/pakicetus), *Dorudon atrox* (by T. Michael Keesey, CC0 1.0 Universal Public Domain Dedication: https://www. phylopic.org/images/b02ddb7c-a946-4d95-bd5d-57ce575e8b76/dorudon-atrox), *Feresa attenuata* (by T. Michael Keesey based on work by Chris huh, Attribution-ShareAlike 3.0 Unported: https://www. phylopic.org/images/b61a3c6f-66ae-4c16-8ff9-4616aba24f98/feresa-attenuata), *Mesosaurus* (by Antoine Verrière, based on illustration by Nobu Tamura, Attribution-NonCommercial-ShareAlike 3.0 Unported: https://www.phylopic.org/ images/293052d9-23df-40b4-893a-22e3cb093732/mesosaurus), *Drepanosaurus unguicaudatus*, (by Xavier Jenkins, CC0 1.0 Universal Public Domain Dedication: https://www.phylopic.org/images/ 63aefb22-c38a-475d-8a9c-6c14dced258f/drepanosaurus-unguicaudatus); *Claudiosaurus germaini* (by T. Michael Keesey, based on illustration by Nobu Tamura, CC BY-SA 3.0: https://www.phylopic.org/ images/009f32c8-cd70-4c5b-8db7-6a7991447d85/ claudiosaurus-germaini), *Araeoscelis gracilis* (by Zak Lewis, based on original illustration by Nix Illustration, Attribution-NonCommercial 3.0 Unported, CC BY-NC 3.0: https://www.phylopic.org/ images/49f89510-3749-4c6e-973e-87d3863a4957/araeoscelis-gracilis), *Spinoaequalis schultzei* (by Caleb Gordon, CC0 1.0 Universal Public Domain Dedication: https://www.phylopic.org/images/9f5ee601-a9da-4e9b-b518-918629cd55dd/spinoaequalis-schultzei), *Sinosaurosphargis yunguiensis* (by Davide Vecchio, CC0 1.0 Universal Public Domain Dedication: https://www.phylopic.org/images/73fbad1a-708c-4cae-a2e3-f88e38c86ecc/sinosaurosphargis-yunguiensis), *Atopodentatus unicus* (by Scott Reid, Attribution 3.0 Unported: https://www.phylopic.org/ images/bd7e8e3b-6c81-4e37-a10f-2b1def7dd77f/atopodentatus-unicus), *Placodus gigas* (by Neil Kelley, CC0 1.0 Universal Public Domain Dedication: https://www. phylopic.org/images/6cc9ddd8-c1bb-4f05-be2d-650492aa3de7/placodus-gigas), *Keichousaurus hui* (by Gareth Monger, CC BY 3.0: https://www. phylopic.org/images/bc166d59-31c3-49fd-8538-0d6904b784cb/keichousaurus-hui), *Nothosaurus mirabilis* (by Dan Niel, CC0 1.0 Universal Public Domain Dedication: https://www.phylopic.org/ images/8f252288-5164-4105-b322-e417bf39fe2d/ nothosaurus-mirabilis), *Pistosaurus longaevus* (by T. Michael Keesey, based on illustration by Nobu Tamura, Attribution-ShareAlike 3.0 Unported: https://www.phylopic.org/images/7999c103-932b-4647-815b-711a1967f0f4/pistosaurus-longaevus), *Kronosaurus* (by dannj, CC0 1.0 Universal Public Domain Dedication: https://www.phylopic.org/ images/cc9e2603-b8f9-414c-a09b-8b8ae3da6343/kronosaurus), *Tuarangisaurus keyesi* (by Arthur S. Brum, CC0 1.0 Universal Public Domain Dedication: https://www.phylopic.org/images/d3a6f6eb-1b95-4d5f-9c3b-99d8b271911a/tuarangisaurus-keyesi), *Hupehsuchus nanchangensis* (by Davide Vecchio, CC0 1.0 Universal Public Domain Dedication: https://www.phylopic.org/images/302dc673-14d9-45d0-8d39-09d2e6b4ffa8/hupehsuchus-nanchangensis), *Hupehsuchus* (by Scott Reid, 2021, Attribution 3.0 Unported: https://www.phylopic.org/ images/9578e572-e0f3-4913-bb17-836329fe0121/hupehsuchus), *Cartorhynchus lenticarpus* (by Davide Vecchio, CC0 1.0 Universal Public Domain Dedication: https://www.phylopic.org/images/59c50f5f-6c0f-4725-8d97-1e951cf96391/cartorhynchus-lenticarpus), *Chaohusaurus geishanensis* (by Gareth Monger, 2013, Attribution 3.0 Unported: https://www.phylopic .org/images/d940c3c4-2b59-4e68-b31b-2cf6c89130f3/chaohusaurus-geishanensis), *Utatsusaurus hataii* (by T. Michael Keesey, based on illustration by Nobu Tamura, Attribution-ShareAlike 3.0 Unported: https://www.phylopic.org/images/cf07f025-6977-430a-8ea1-a2297f3f864c/utatsusaurus-hataii), *Grippia longirostris* (by T. Michael Keesey, based on illustration by Dmitry Bogdanov, Attribution 3.0 Unported: https://www.phylopic.org/images/ab7194d3-1d74-4029-a6b1-81663a25dc13/grippia-longirostris), *Askeptosaurus brevis* (by T. Michael Keesey, based on illustration by Nobu Tamura, CC BY-SA 3.0: https://www.phylopic.org/images/3f653f44-f60e-47e6-860c-790045020a70/askeptosaurus-brevis), *Askeptosaurus italicus* (by Davide Vecchio, CC0 1.0 Universal Public Domain Dedication: https://www. phylopic.org/images/ fb96fb4f-8c21-4b6c-9472-4fce5158b8e8/askeptosaurus-italicus, used in both its original state to represent *Askeptosaurus* and a slightly modified state [with a more rounded trunk and flattened snout] to represent *Endennasaurus*), *Champsosaurus* (by Steven Traver, CC0 1.0 Universal Public Domain Dedication: https://www.phylopic.org/images/833cb3e5-9016-441a-8d57-42d3e0ae9484/ champsosaurus), *Coeruleodraco jurassicus* (by T. Michael Keesey, Attribution 4.0 International:

https://www.phylopic. org/images/199fac5d-0473-4647-ba73-338f59bac28d/coeruleodraco-jurassicus), *Protorosaurus speneri* (by T. Michael Keesey, based on illustration by SeismicShrimp, Attribution 4.0 International: https://www.phylopic.org/images/3f260f5a-229a-4bce-9647-0e5a54a4adc9/protorosaurus-speneri), *Langobardisaurus Pandolfi* (by Cy Marchant, Attribution 4.0 International: https://www. phylopic.org/images/7414551d-5914-4156-83ec-ce9831b85218/langobardisaurus-pandolfii), *Hyperodapedon huxleyi* (by T. Michael Keesey, based on illustration by Nobu Tamura, Attribution 4.0 International: https://www.phylopic.org/images/0287d1e0-eccc-460e-bdc5-b36ee0fe86b1/ hyperodapedon-huxleyi), *Poposaurus gracilis* (by Scott Hartman, Attribution 3.0 Unported: https:// www.phylopic.org/images/ 694830cf-1e9d-4795-8165-ef7fce9c088d/poposaurus-gracilis), *Steneosaurus bollensis* (by Gareth Monger, Attribution 3.0 Unported: https://www.phylopic.org/ images/b15e46eb-be9a-42b4-9a06-8b718249ab23/steneosaurus-bollensis), *Geosaurus giganteus* (by T. Michael Keesey, based on illustration by Dmitry Bogdanov, Attribution 3.0 Unported: https://www.phylopic.org/images/ 80315bbf-e7e9-4f9a-b598-6a8a68f3b169/geosaurus-giganteus), Tomistominae (by Armin Reindl, Attribution-NonCommercial 3.0 Unported: https://www.phylopic.org/images/f1a14495-1852-4a81-bafb-a58fa9cc7f76/tomistominae), *Gavialis gangeticus* (by Jagged Fang Designs, CC0 1.0 Universal Public Domain Dedication: https://www.phylopic.org/images/f963f388-8297-4ceb-91f6-493b38dccba5/gavialis-gangeticus), *Hesperornis regalis* (by T. Michael Keesey, based on illustration by Nobu Tamura, Attribution 3.0 Unported: https://www.phylopic.org/images/6a96ce0a-ef5c-4986-ac48-01e66cfb1531/ hesperornis-regalis), *Protostega gigas* (by T. Michael Keesey, based on illustration by Dmitry Bogdanov, Attribution 3.0 Unported: https://www.phylopic.org/images/d0ba41b6-78d8-415a-ae57-24df5fc501ee/ protostega-gigas), *Dermochelys coriacea* (by Ray Chatterji, CC0 1.0 Universal Public Domain Dedication: https://www.phylopic.org/images/1c65c811-4caa-4be3-9936-7a16e6905131/ dermochelys-coriacea), *Ardeosaurus brevipes* (by T. Michael Keesey based on former work by Ghedo and T. Michael Keesey, Attribution-ShareAlike 3.0 Unported: https://www.phylopic.org/images/ 83053aee-0f56-4cf3-bbfa-9207e6f13f46/ardeosaurus-brevipes), *Marmoretta oxoniensis* (by T. Michael Keesey, CC0 1.0 Universal Public Domain Dedication: https://www.phylopic.org/images/ddd87c97-827b-4c59-93e6-67d7d8493989/ marmoretta-oxoniensis), *Sphenodon punctatus* (by Steven Traver, CC0 1.0 Universal Public Domain Dedication: https://www.phylopic.org/images/f2a5ae73-c899-4e47-b0ad-b6eac3a99350/ sphenodon-punctatus), *Gekko gecko* (by Steven Traver, CC0 1.0 Universal Public Domain Dedication: https://www. phylopic.org/images/9aca34d8-4dde-418d-9fdc-2d58b6a7b267/gekko-gecko), *Hydrophis curtus* (by Christina Zdenek, Attribution 4.0 International: https://www.phylopic.org/ images/38588c8f-cd7b-4051-b5e3-6433ed1bba89/hydrophis-curtus), *Lanthanotus borneensis* (by François-Louis PELISSIER, CC0 1.0 Universal Public Domain Dedication: https://www.phylopic.org/ images/7761ca72-d764-4700-93f5-cac1bb2db0b0/lanthanotus-borneensis), *Heloderma suspectum* (by Nicolas Mongiardino Koch, CC0 1.0 Universal Public Domain Dedication: https://www.phylopic.org/ images/9cae2028-126b-416f-9094-250782c5bc22/heloderma-suspectum) *Varanus gouldii* (by Rachel T. Mason, CC0 1.0 Universal Public Domain Dedication: https://www.phylopic.org/images/de23dbe6-70b4-4eca-987f-fceed2ba5641/varanus-gouldii), *Tethysaurus nopcsai* (by Zimices, Julián Bayona, Attribution-ShareAlike 3.0 Unported: https://www.phylopic.org/ images/52044499-6262-48f0-a61a-b4f193c536f7/ tethysaurus-nopcsai), *Aigialosaurus bucchichi* (by thefunkmonk, CC0 1.0 Universal Public Domain Dedication: https://www. phylopic.org/images/4e498bf6-bc90-41d1-9c3c-00badad7a92f/ aigialosaurus-bucchichi), *Pannoniasaurus inexpectatus* (by T. Michael Keesey, based on work by László Makádi, Michael W. Caldwell, and Attila Ősi, Attribution 4.0 International: https://www.phylopic.org/images/ 3e73bbac-998a-4fbc-b00c-41f3e1c0f705/pannoniasaurus-inexpectatus), *Clidastes propython* (by Cy Marchant, Attribution 4.0 International: https://www.phylopic.org/images/05d5c2c0-78c5-48e3-b8ca-5243a6bba401/clidastes-propython), *Taniwhasaurus* (by Craig Dylke, Attribution 3.0 Unported: https://www.phylopic.org/images/b20f473b-03a1-4267-a9b6-879022456f7d/taniwhasaurus), *Platecarpus* (by Matthew Crook, Attribution-ShareAlike 3.0 Unported: https://www.phylopic.org/images/5253b3b3-a1f8-4839-a67b-10f6606ffc0b/platecarpus),

Additional original silhouettes were created in Adobe Photoshop (v. 23.5.1) for the following taxa:

*Eretmorhipis carrolldongi*, traced from Long et al. 2019 [https://doi.org/10.1038/s41598-018-37754-6], Fig. 1c reconstruction; *Eusaurosphargis hupehensis* (based on photograph of PIMUZ A III 4380, provided by Torsten Scheyer); *Hanosaurus hupehensis* (based on photograph of IVPP V 15911, taken and provided by Wei Wang); *Hyphalosaurus lingyuanensis* (based on photograph of IVPP V11075, provided by Keqin Gao); *Labidosaurus hamatus* (based on the 2008 Wikimedia Commons illustration by A. Кац: https://commons.wikimedia. org/wiki/File:Labidosaurus_ hamatus.jpg); *Lariosaurus* (based on photograph of PIMUZ T4288, provided by Christian Klug); *Langobardisaurus pandolfi* (based on photograph of MFSN 1921, provided by Luca Simonetto); *Luopingosaurus imparilis* (based on photograph of IVPP V 19049, provided by Guang Hui Xu); *Miodentosaurus brevis* (based on photographs of NMNS 004727 and ZMNH M8742, taken and provided by Xiao-Chun Wu; *Neusticosaurus peyeri* (based on photograph of MGP PD 27538, taken and provided by Giovanni Serafini); *Pachypleurosaurus edwardsi* (based on photograph of SMF R4964, provided by Rainer Brocke); *Panzhousaurus rotundirostris* (based on photograph of GMPKU P 1059, provided by Da-yong Jiang); *Pleurosaurus ginsburgi* (based on photograph of JME MOE 170-R77, belonging to the collection of the Bishops Seminar Eichstätt, taken and provided by Christina Ifrim at SNSB Jura Museum Eichstätt); *Saniwa ensidens* (based on photograph of FMNH PR2378, taken by John Weinstein and provided by William Simpson: © The Field Museum, GEO86871-03-17d); *Vadasaurus herzogi* (traced from published photograph of AMNH FARB 32768, as figured in Bever & Norell 2017 [http://dx.doi.org/10.1098/rsos.170570], Fig. 1a); and *Wayaosaurus bellus* (based on photograph of GZD-B 0005, taken and provided by Jun Chai). Phylopic and original silhouettes were recolored and positioned within larger figures with BioRender.com.

### Supplementary Figures

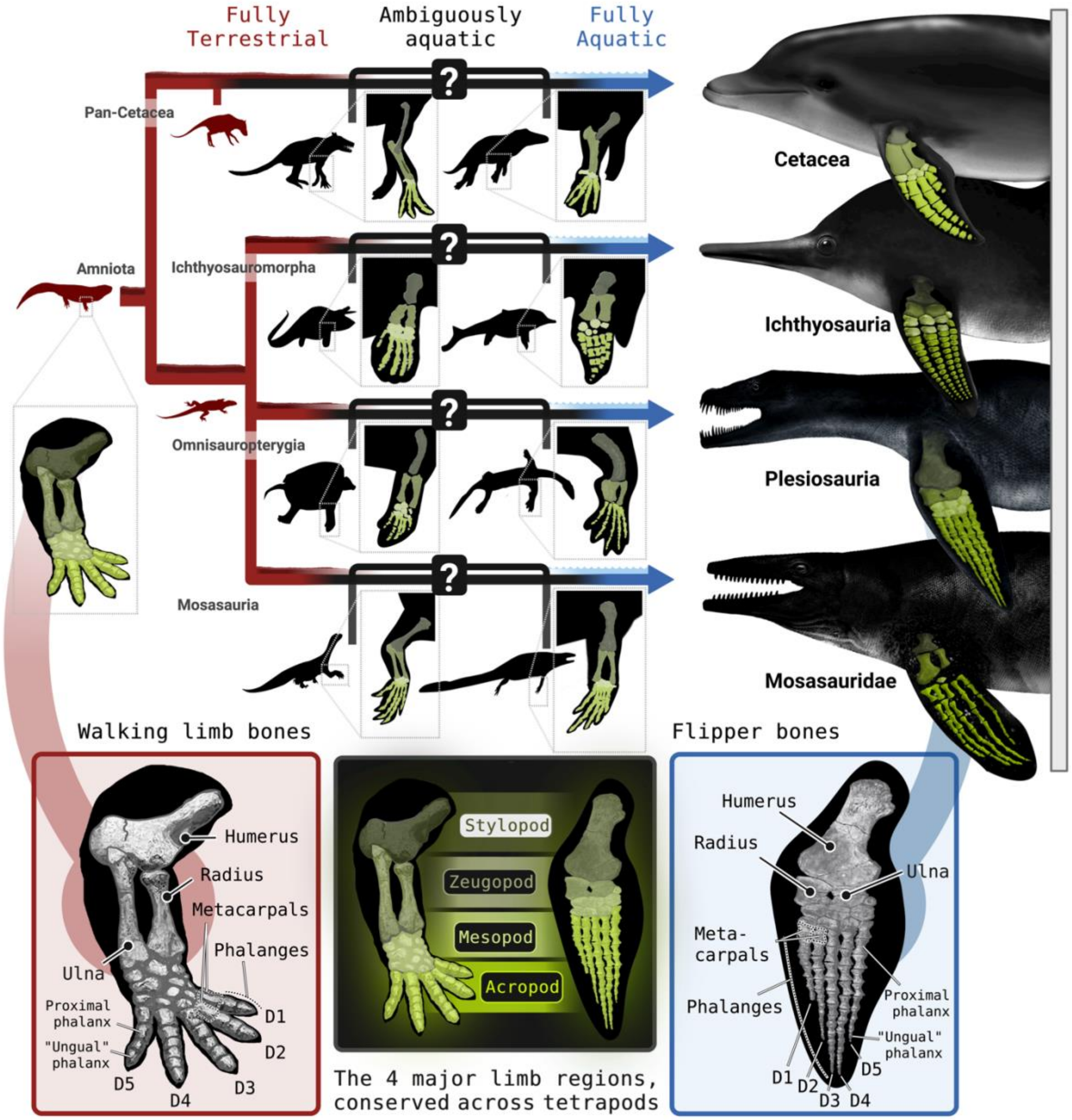


**Figure S1. Extended rationale and anatomical context.** The four clades shown on the right are highly adapted for an aquatic lifestyle, and their limbs are easily identified as flippers, with elongated and often supernumerary acropodial elements. However, the more basally branching taxa on the line leading to each of these highly aquatic clades have ambiguous limb morphologies, which make their precise aquatic habits difficult to discern. The limb interfaces directly with the locomotor media of these taxa, and thus serves as a treasure trove of potential information on their habitat preferences. Although limbs are highly modified across amniotes (particularly in highly or fully aquatic species), they all retain four major limb regions: the stylopodium (upper arm or thigh), zeugopodium (lower arm or shin), mesopodium (ankle or wrist), and acropodium (digital region, consisting of the palm and fingers or the metatarsals and toes). These conserved limb regions allow us to compare the shapes and proportions of the limb across amniotes despite their varied morphologies. D1–D5 denote digits 1–5. Figure silhouettes on the right were illustrated by Raquel Jaramillo. Phylopic attributions are provided in Text S19.

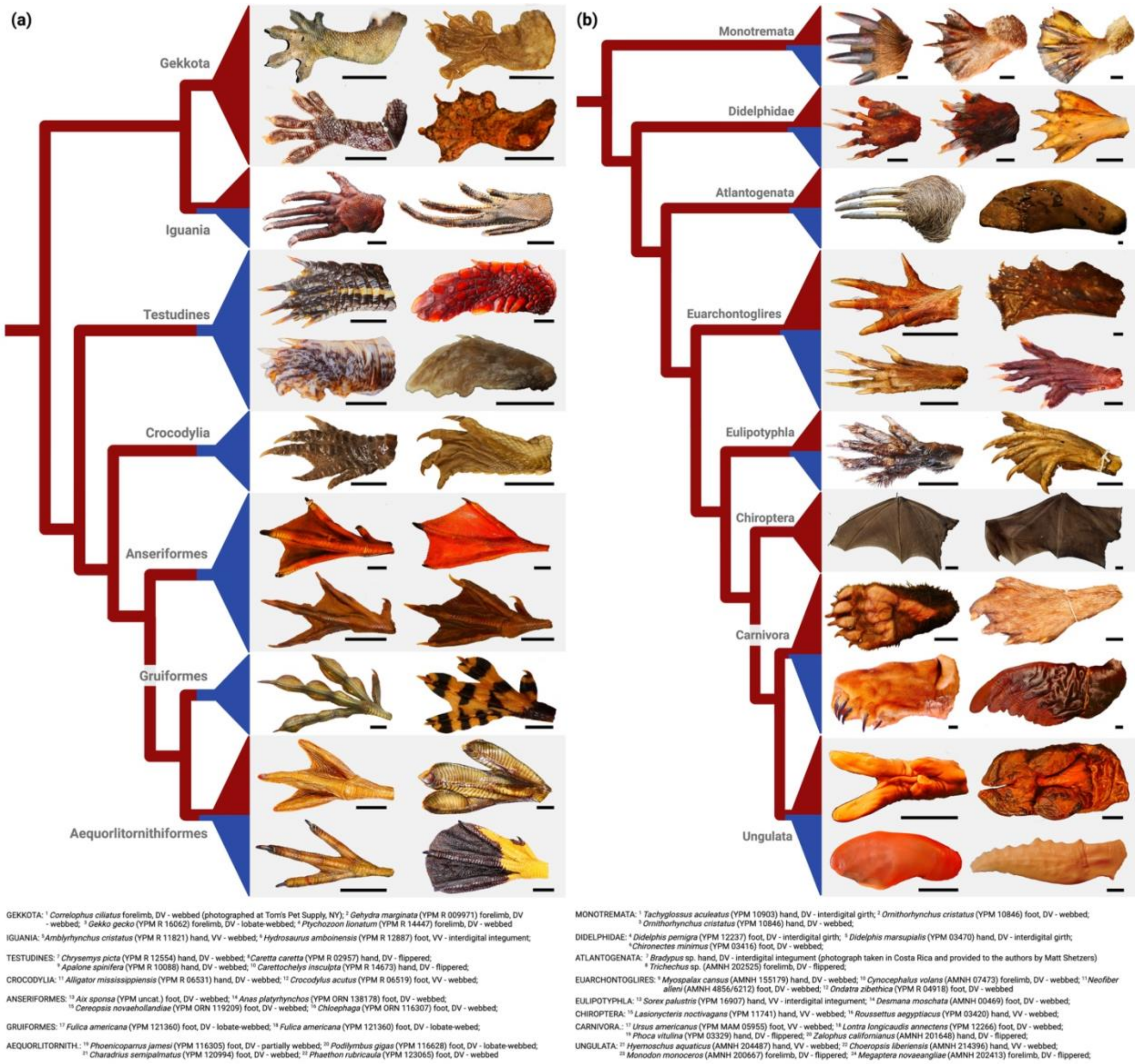


**Figure S2. Varieties of interdigital tissue in extant amniotes. (a)** Reptiles and **(b)** mammals show a striking diversity of interdigital tissue morphologies. These include stiff integumentary structures (interdigital hairs and scales); palmar girth that invades interdigital spaces; flippers of varying stiffness that encompass the whole distal limb; and flimsy, cellularized interdigital tissues (true "webbing") of various shapes and extents. Limbs are shown in either dorsal or ventral planform view, oriented with the distal end of the limb to the left. Tree branches are colored to show aquatic habits (blue: nearly all species aquatic; red: nearly all species terrestrial; mixed blue/red: many species aquatic and many terrestrial). The broad distribution of webbing and aquatic habits highlights that ecotype and interdigital webbing are often but not always associated. All specimen images are original photographs by C. M. G. unless stated otherwise. Abbreviations: AMNH, American Museum of Natural History; DV, dorsal view; VV, ventral view; YPM, Yale Peabody Museum.

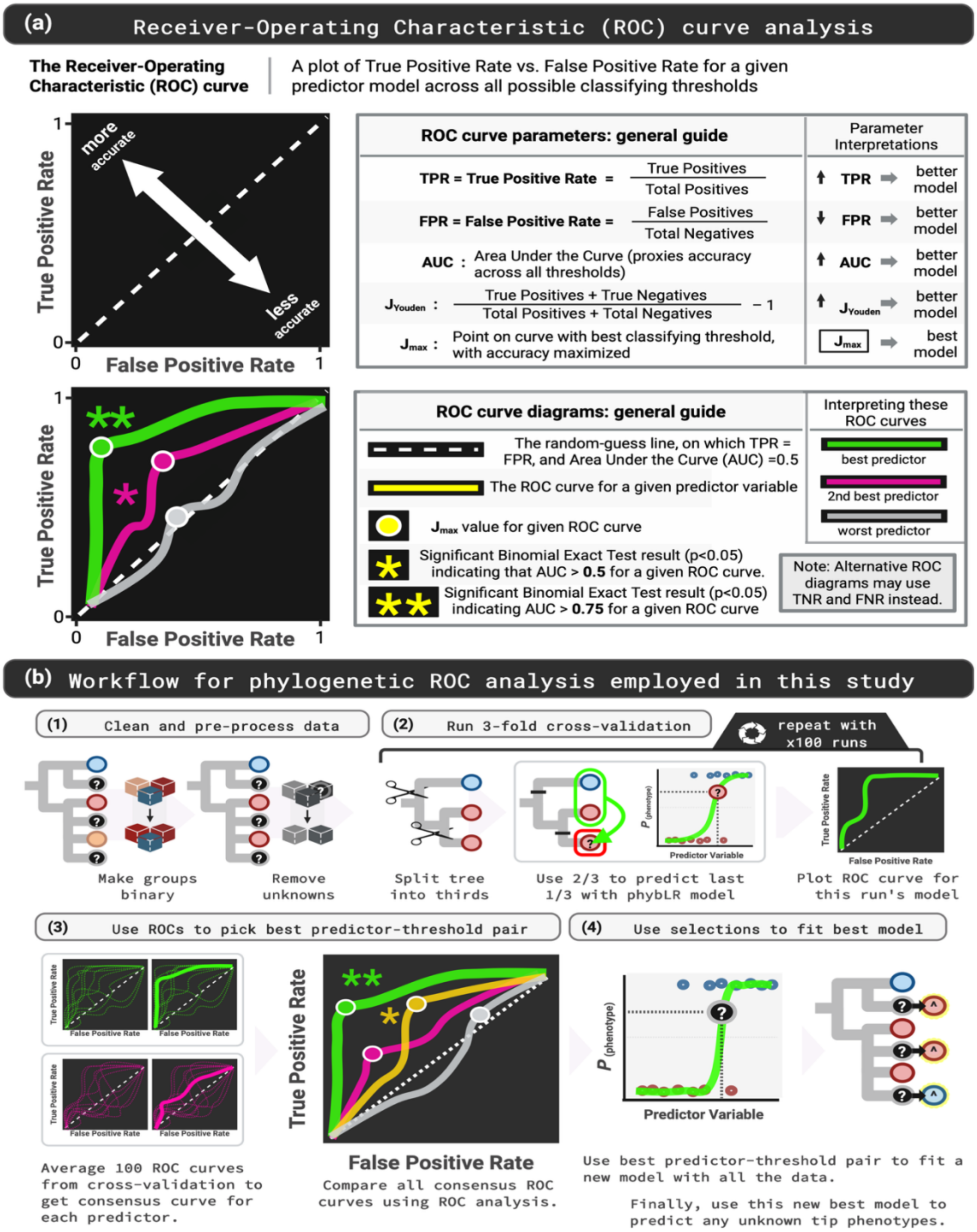


**Figure S3. Supervised machine-learning pipeline used in the study. (a)** A guide illustrates the major aspects of Receiver-Operating Characteristic (ROC) curve analysis, which was used to compare the accuracies of competing predictive models. **(b)** A workflow schematic details the major steps in our data analysis pipeline, which involved (1) processing and aligning data to one of eight supertrees, (2) splitting the aligned data into thirds 100 times to train and test phylogenetic binomial logistic regression models, (3) averaging the resulting 100 ROC curves to generate a single consensus curve for each predictor and select the best predictor-threshold pairs, and (4) using this predictor-threshold pair to fit a new logistic regression model with all the data, which we then applied to predict contentious phenotypes in extinct taxa.

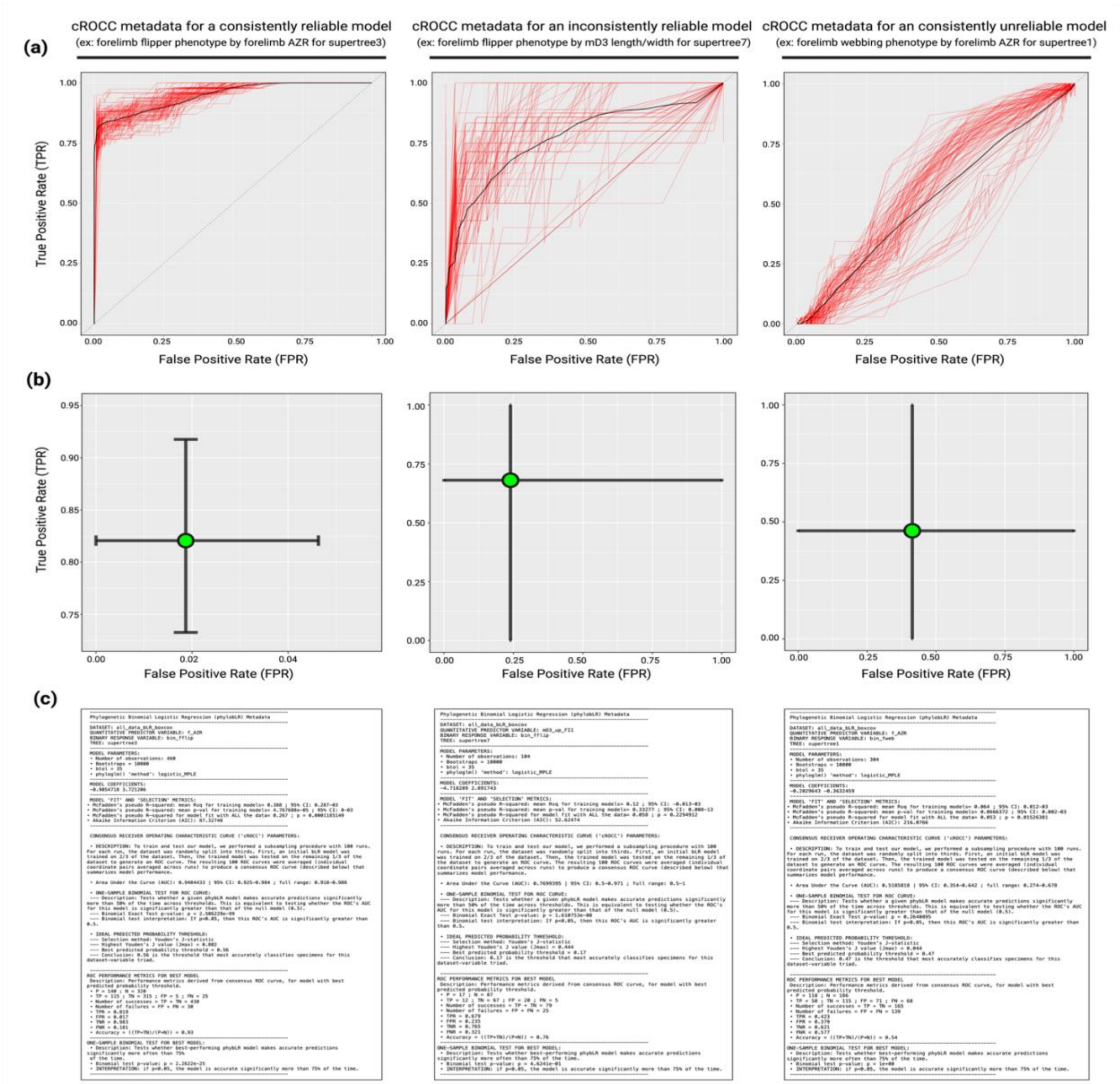


**Figure S4. Example metadata from phylogenetic ROC analysis, contrasting performance of reliable, semi-reliable, and unreliable phylogenetic binomial logistic regression (phybLR) models in ROC space. (a)** ROC curves (red lines) generated across 100 runs of 3-fold cross-validation with the whole dataset, averaged to generate the consensus ROC curve (cROCC, thick black line) for that model. Consistently reliable phybLR models had ROC curves that varied little across runs, with high TPRs and low FPRs, whereas less reliable models had more variable ROC curves, and consistently *un*reliable models had ROC curves with lower average TPRs and higher average FPRs. **(b)** 95% confidence intervals for the TPRs and FPRs of each of the above phybLR models given their optimal probability classification threshold (that threshold which maximized Youden's J-statistic [see Figure S3]). As in (a), the more reliable phybLR model has a high mean TPR and a low mean FPR with minimal variance, the less reliable phybLR model retains a high mean TPR and a low mean FPR but with greater variance, and the consistently unreliable phybLR model retains a high variance but with a relatively low mean TPR and high mean FPR. **(c)** Full set of metadata associated with the cross-validation and ROC analysis of each phybLR model, providing a richer set of information about model performance. The plots and metadata within this figure are available for every tested phybLR model from our study in Data S4.

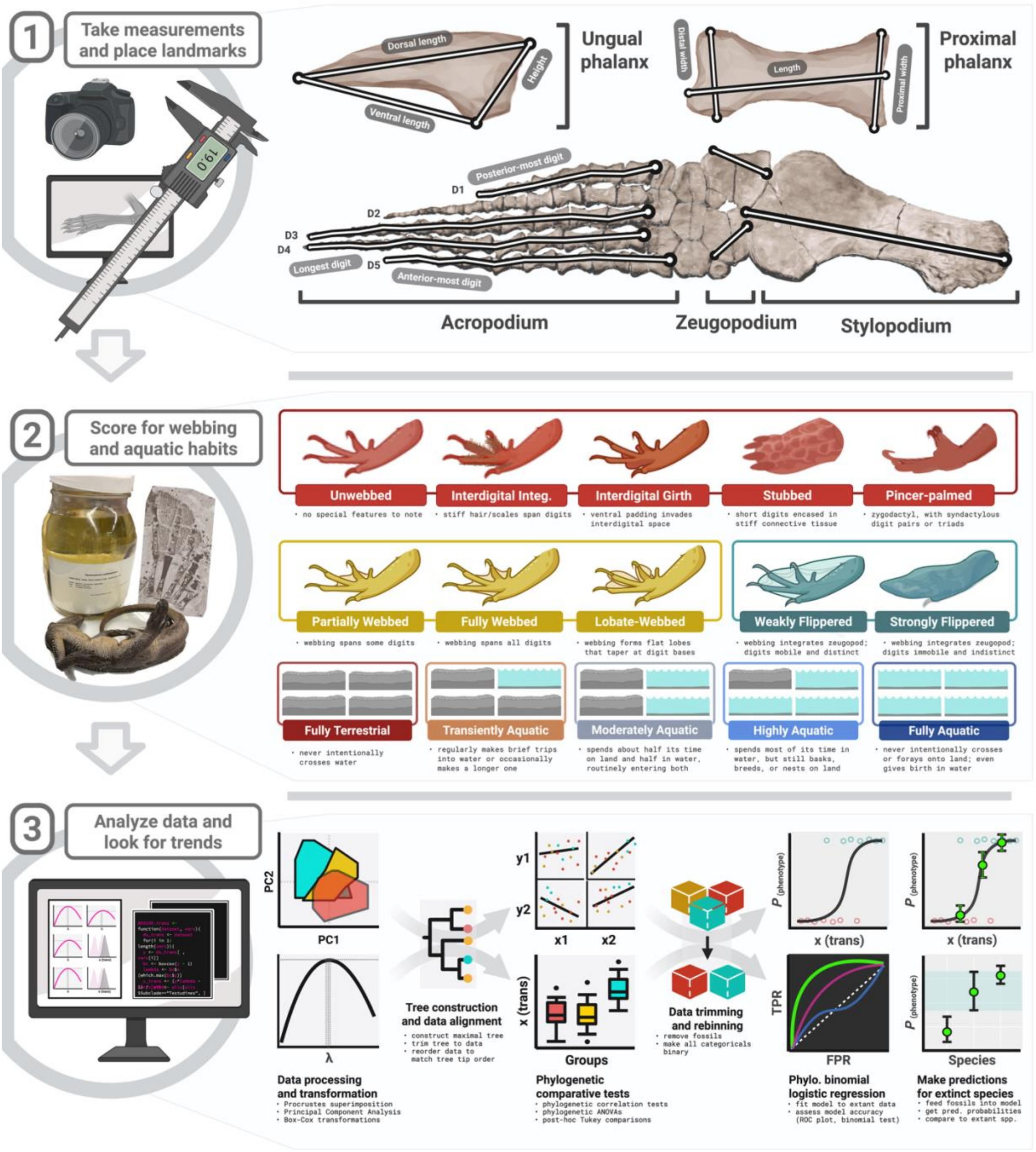


**Figure S5. Phylogenetic ROC analysis in the context of the broader workflow for this study.** We began by (1) collecting limb measurements or placing landmarks on a broad sample of extant and fossil amniotes. We then (2) scored all measured taxa in our sample for their soft-tissue limb morphologies and aquatic affinities using the classification system shown above. Finally, we (3) analyzed the data in a phylogenetic comparative framework, ultimately fitting phylogenetic binomial logistic regression (phybLR) models to the data for every morphometric-categorical variable pair, selecting the most accurate predictors of soft-tissue phenotype and aquatic affinity using ROC analysis, and using these most accurate models to predict the soft-tissue limb phenotypes and aquatic affinities of extinct species.

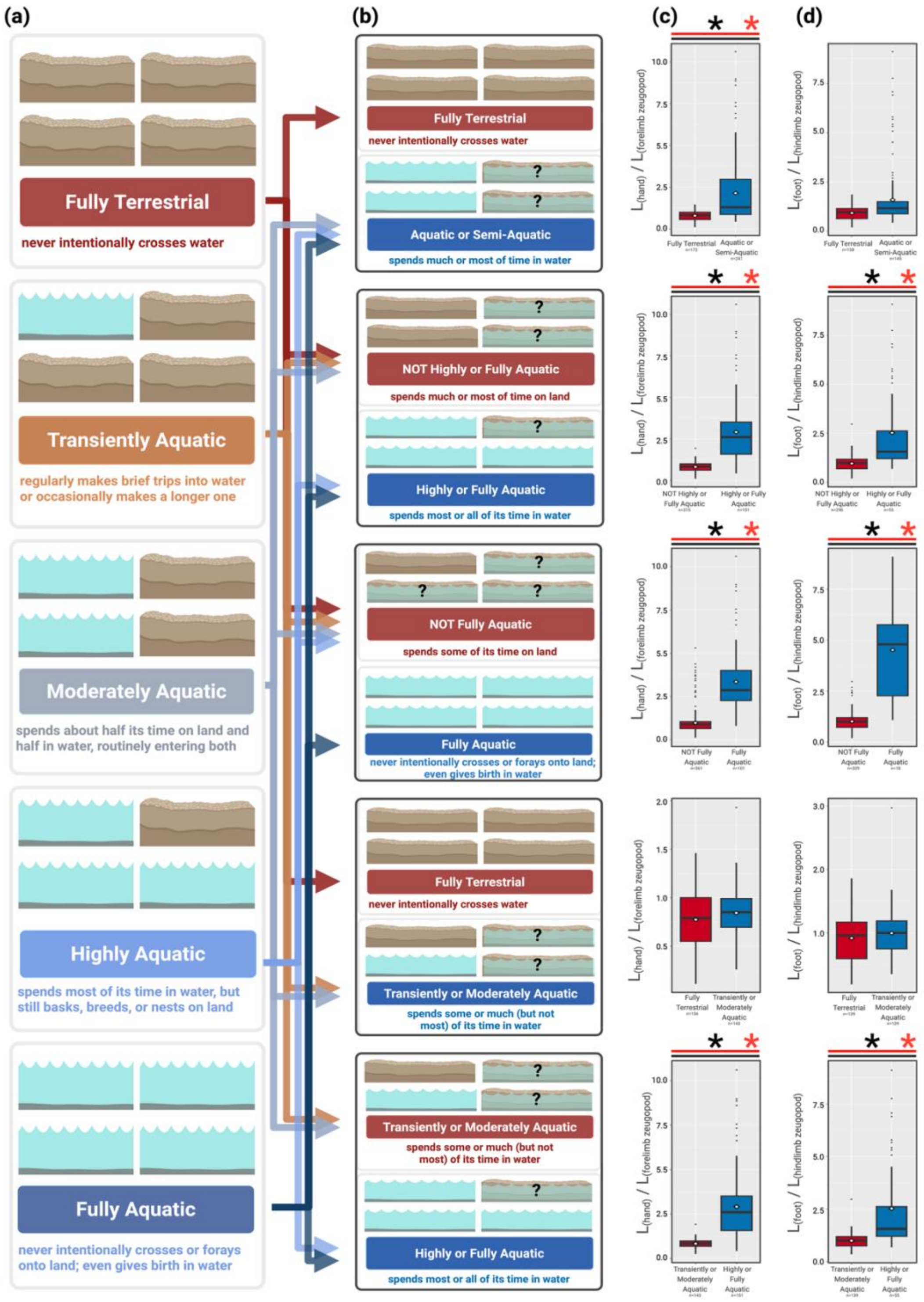


**Figure S6. Re-binning procedure and associated measurement comparisons for aquatic affinity bins.** **(a)** Original aquatic affinity guilds. **(b)** Recombined binary aquatic affinity bins used to score the response variable in downstream binomial logistic regression models. **(c–d)** Box plots show comparable trends in forelimb (c) and hindlimb (d) region proportions for all binary classification systems including highly or fully aquatic taxa. Asterisks denote significant results ($p < 0.05$) of phylogenetic ANOVAs (black) and Levene's tests (red). Omnibus and post-hoc tests are equivalent because there are only two categories.

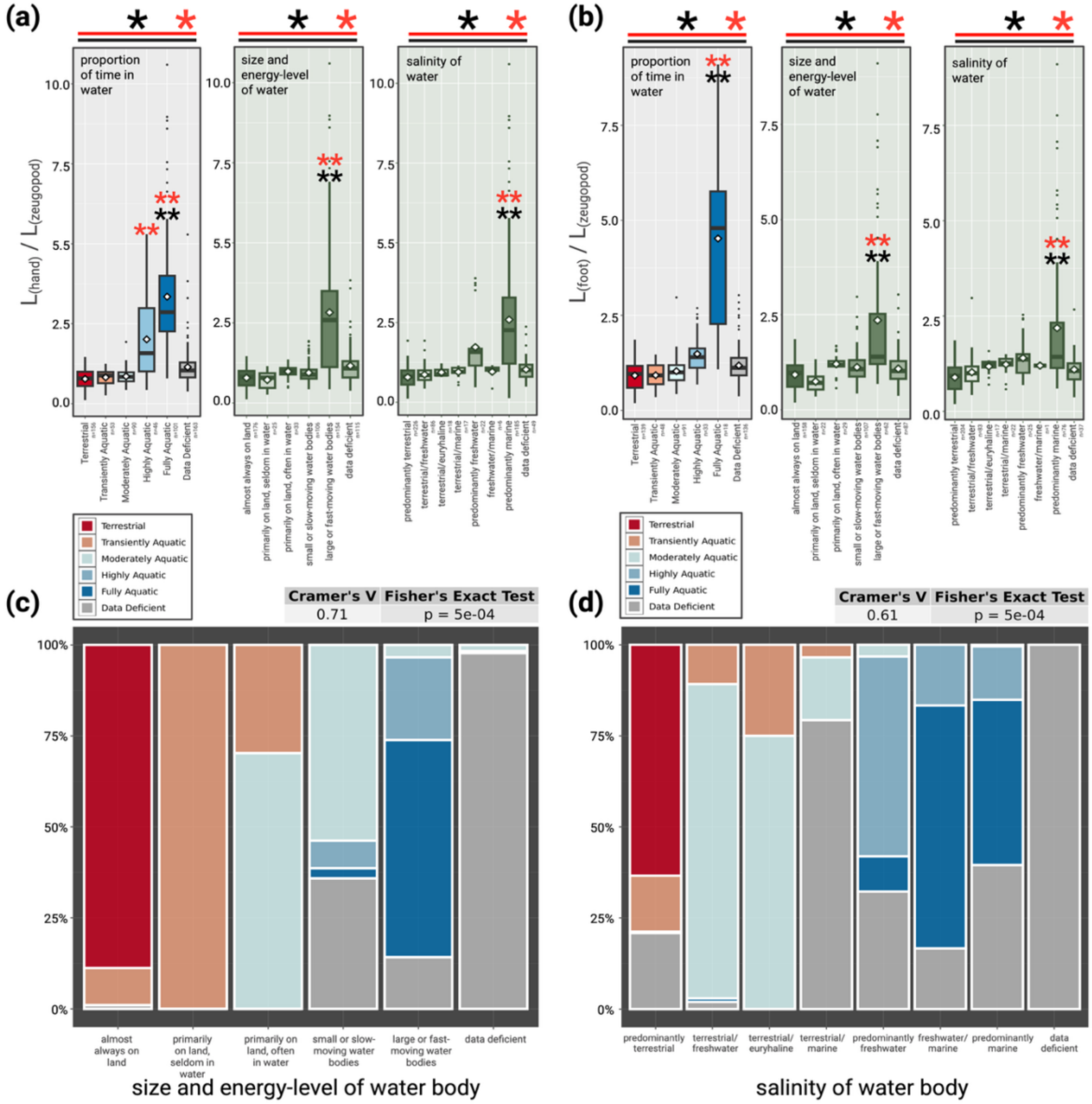


**Figure S7. Similar results for alternative ecotype classification systems.** In addition to our own ecological binning system based on the relative amounts of time spent on land vs. water, we scored aquatic affinities using two binning systems based on the nature of the bodies of water inhabited—one based on their size and energy level (Joyce & Gauthier 2004) and another based on their salinity. **(a–b)** Box plots show how relative hand (a) and foot (b) lengths differ among aquatic affinity bins for all three classification systems. Asterisks denote significant results ($p < 0.05$) of phylogenetic ANOVAs (black) and Levene's tests (red). Single asterisks indicate significant omnibus test results, and double asterisks denote post-hoc test results (showing significant difference from the "most terrestrial" bin). The latter two binning systems yielded results that were broadly similar to our own, with the most aquatic group recovered as having significantly longer hands and feet than the most terrestrial group. **(c–d)** Category association plots show the percentage overlap in classification for all specimens in our dataset, between the time- vs. size/energy-based binning systems (c), and the time- vs. salinity-based systems (d). Cramer's V reports the strength (from 0 to 1) of the association between each pair of categorical variables, and Fisher's exact tests indicate that character scores across both pairs of binning systems are significantly associated (each $p < 0.05$).

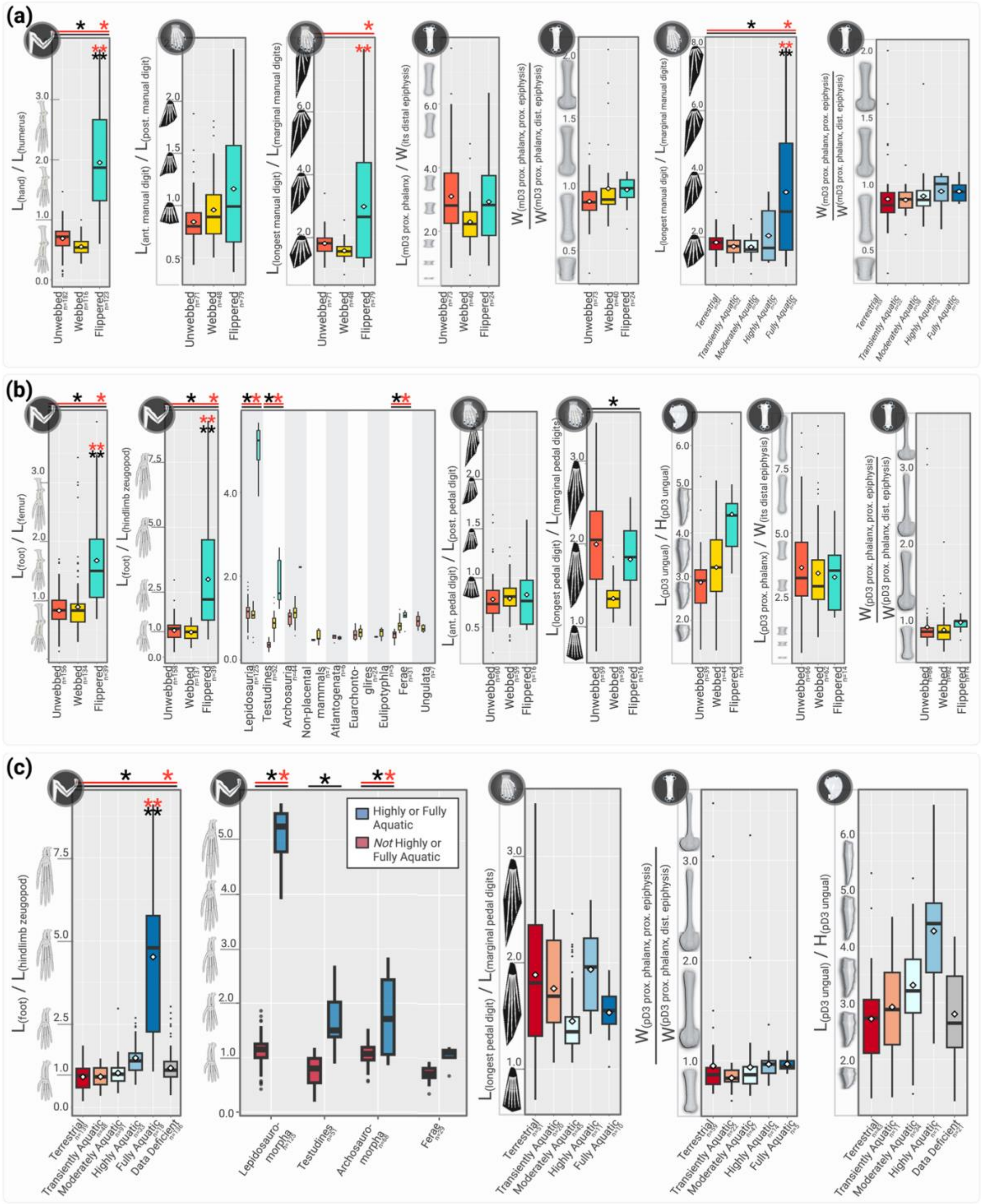

**Figure S8. Additional linear morphometric results for soft-tissue limb phenotypes and aquatic affinity guilds across amniotes. (a)** Key additional forelimb results recovered across amniotes. **(b–c)** Key hindlimb results recovered across amniotes, for **(b)** limb soft-tissue phenotypes and **(c)** aquatic affinity bins. Asterisks denote significant results ($p < 0.05$) of phylogenetic ANOVAs (black) and Levene's tests (red)—single asterisks for omnibus tests and double asterisks for pairwise post-hoc tests (here indicating difference from the "terrestrial" group). Abbreviations: ant., anterior; D3, digit 3; dist., distal; H, height; L, length; m, manual; n, sample size; post., posterior; p, pedal; prox., proximal; W, width.

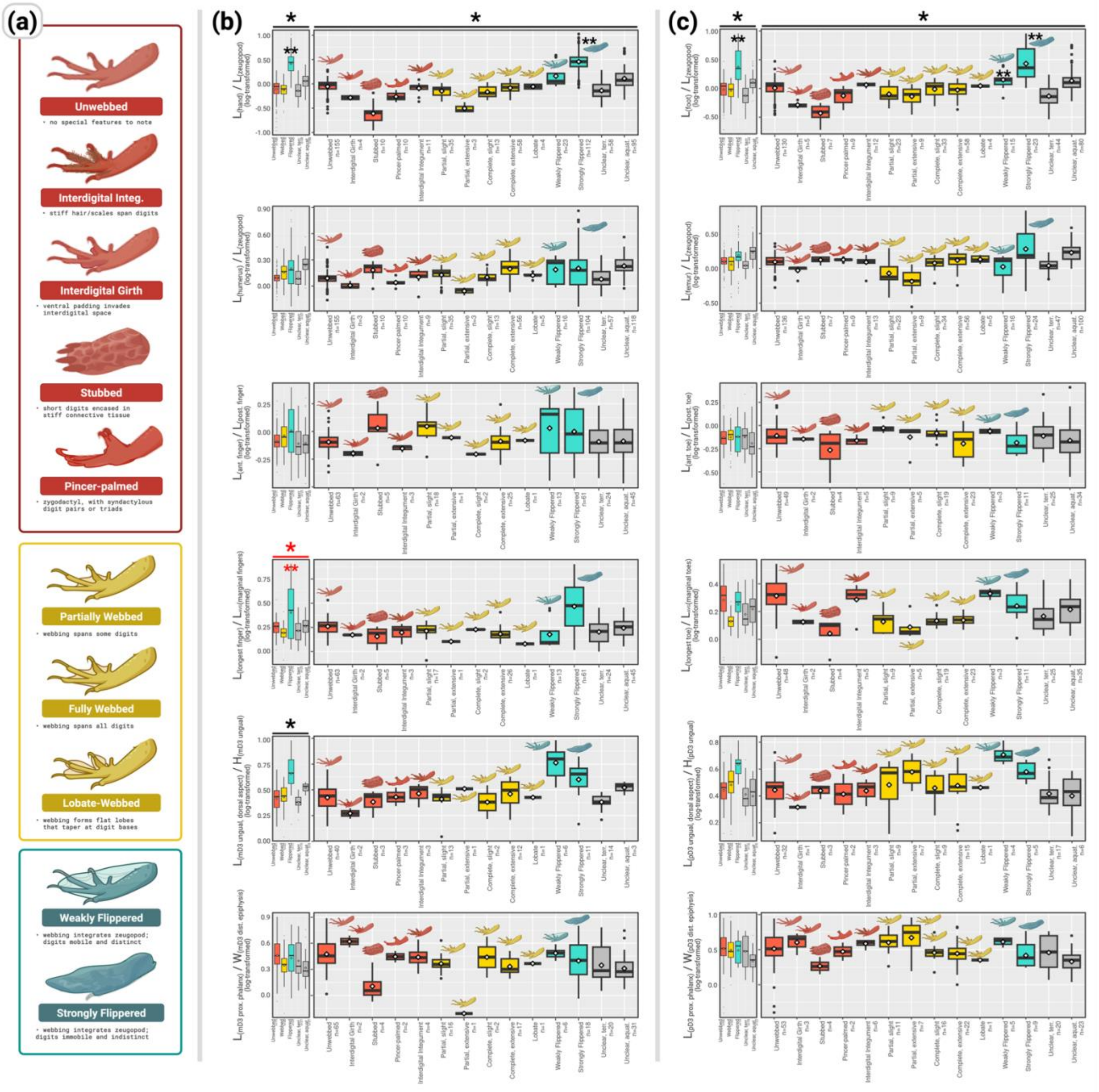


**Figure S9. Morphological differences observed among narrower limb phenotype categories. (a)** Given the diversity of extant webbing phenotypes (Figure S2), unwebbed (red), webbed (gold), and flippered (turquoise) taxa were further subdivided into narrower soft-tissue phenotype bins as shown, and classified (wherever possible) following direct observation of zoological specimens. **(b–c)** Morphometric patterns observed in the forelimb (b) and hindlimb (c) among both broad and narrow phenotype bins. All values are $log_{10}$-transformed to make differences more visible. Asterisks denote significant results ($p < 0.05$) of phylogenetic ANOVAs (black) and Levene's tests (red). Single asterisks correspond to omnibus tests and double asterisks to pairwise post-hoc tests (here indicating significant difference from the "unwebbed" group). Ultimately, taxa with interdigital webbing lacked a distinct morphometric profile regardless of how many operational webbing categories we defined. Abbreviations: aquat., aquatic; integ., integument; n, sample size; terr., terrestrial. Additional anatomical abbreviations are given in Figure S8.

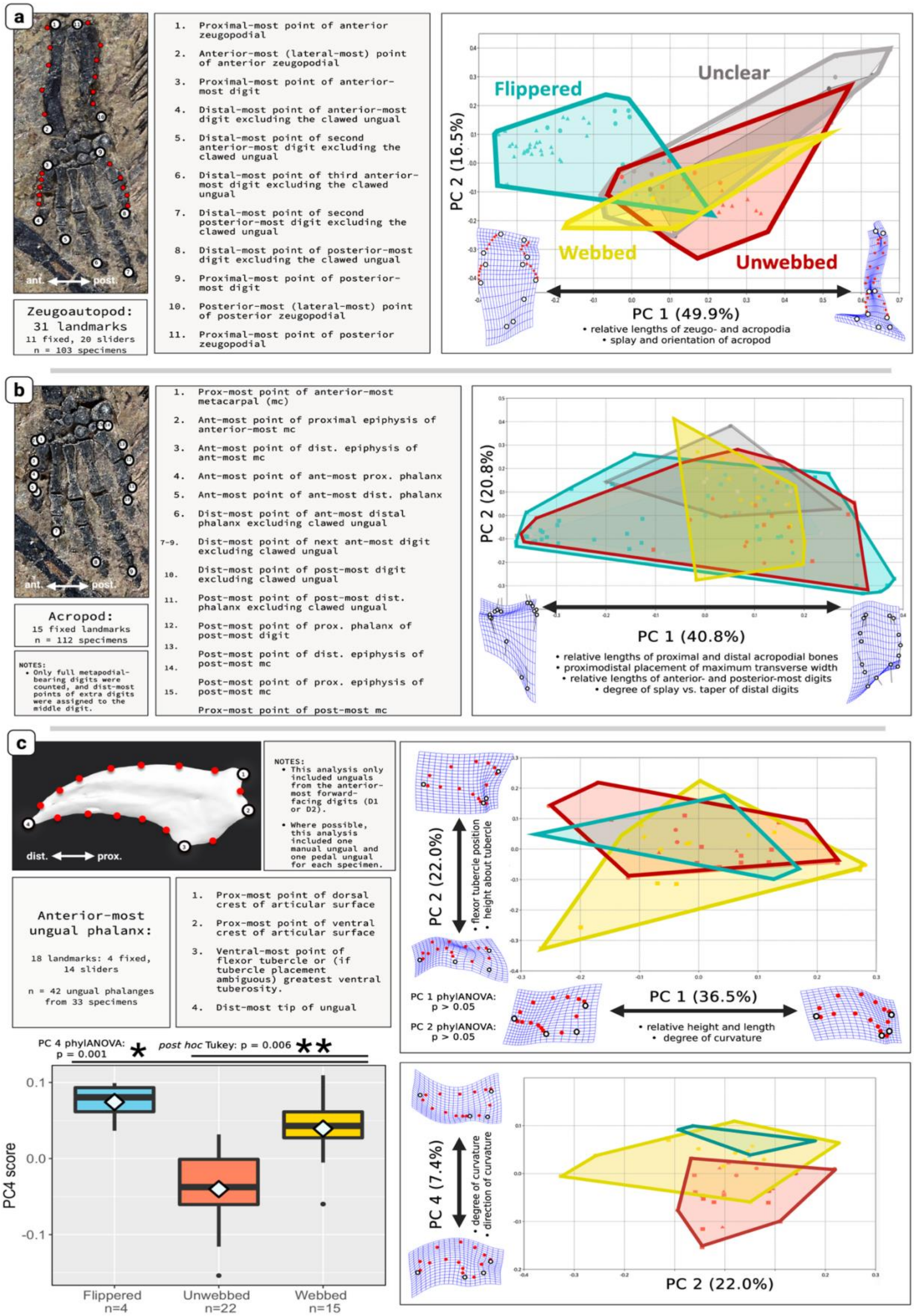
a
ant. post.
Zeugoautopod: 31 landmarks
11 fixed, 20 sliders
n = 103 specimens
1. Proximal-most point of anterior zeugopodial
2. Anterior-most (lateral-most) point of anterior zeugopodial
3. Proximal-most point of anterior-most digit
4. Distal-most point of anterior-most digit excluding the clawed ungual
5. Distal-most point of second anterior-most digit excluding the clawed ungual
6. Distal-most point of third anterior-most digit excluding the clawed ungual
7. Distal-most point of second posterior-most digit excluding the clawed ungual
8. Distal-most point of posterior-most digit excluding the clawed ungual
9. Proximal-most point of posterior-most digit
10. Posterior-most (lateral-most) point of posterior zeugopodial
11. Proximal-most point of posterior zeugopodial
Flippered
Unclear
Webbed
Unwebbed
PC 2 (16.5%)
PC 1 (49.9%)
• relative lengths of zeugo- and acropodia
• splay and orientation of acropod
b
ant. post.
Acropod:
15 fixed landmarks
n = 112 specimens
NOTES:
• Only full metapodial-bearing digits were counted, and dist-most points of extra digits were assigned to the middle digit.
1. Prox-most point of anterior-most metacarpal (mc)
2. Ant-most point of proximal epiphysis of anterior-most mc
3. Ant-most point of dist. epiphysis of ant-most mc
4. Ant-most point of ant-most prox. phalanx
5. Ant-most point of ant-most dist. phalanx
6. Dist-most point of ant-most distal phalanx excluding clawed ungual
7-9. Dist-most point of next ant-most digit excluding clawed ungual
10. Dist-most point of post-most digit excluding clawed ungual
11. Post-most point of post-most dist. phalanx excluding clawed ungual
12. Post-most point of prox. phalanx of post-most digit
13. Post-most point of dist. epiphysis of post-most mc
14. Post-most point of prox. epiphysis of post-most mc
15. Prox-most point of post-most mc
PC 2 (20.8%)
PC 1 (40.8%)
• relative lengths of proximal and distal acropodial bones
• proximodistal placement of maximum transverse width
• relative lengths of anterior- and posterior-most digits
• degree of splay vs. taper of distal digits
c
dist. prox.
NOTES:
• This analysis only included unguals from the anterior-most forward-facing digits (D1 or D2).
• Where possible, this analysis included one manual ungual and one pedal ungual for each specimen.
Anterior-most ungual phalanx:
18 landmarks: 4 fixed, 14 sliders
n = 42 ungual phalanges from 33 specimens
1. Prox-most point of dorsal crest of articular surface
2. Prox-most point of ventral crest of articular surface
3. Ventral-most point of flexor tubercle or (if tubercle placement ambiguous) greatest ventral tuberosity.
4. Dist-most tip of ungual
PC 2 (22.0%)
• flexor tubercle position
• height about tubercle
PC 1 phylANOVA: p > 0.05
PC 2 phylANOVA: p > 0.05
PC 1 (36.5%)
• relative height and length
• degree of curvature
PC 4 phylANOVA: p = 0.001
post hoc Tukey: p = 0.006
PC4 score
Flippered n=4
Unwebbed n=22
Webbed n=15
PC 4 (7.4%)
• degree of curvature
• direction of curvature
PC 2 (22.0%)

**Figure S10 (previous page). Geometric morphometric (GM) analyses comparing unwebbed, webbed, and flippered amniotes. (a)** Landmarking scheme and morphospace for 2D GM analysis of the distal arm (forelimb zeugoautopod) skeleton in dorsal planform view. This analysis supported the results of linear measurements (Figure 2a–b), as flippered forelimbs had relatively longer acropodial regions. **(b)** Landmarking scheme and morphospace for 2D GM analysis of the forelimb acropodial planform in dorsal planform view. **(c)** Landmarking scheme, morphospaces, and phylogenetic comparative tests for 2D GM analysis of the ungual phalanx in lateral view. Abbreviations: ant., anterior; dist., distal; n, sample size; PC, principal component; post., posterior; prox., proximal.

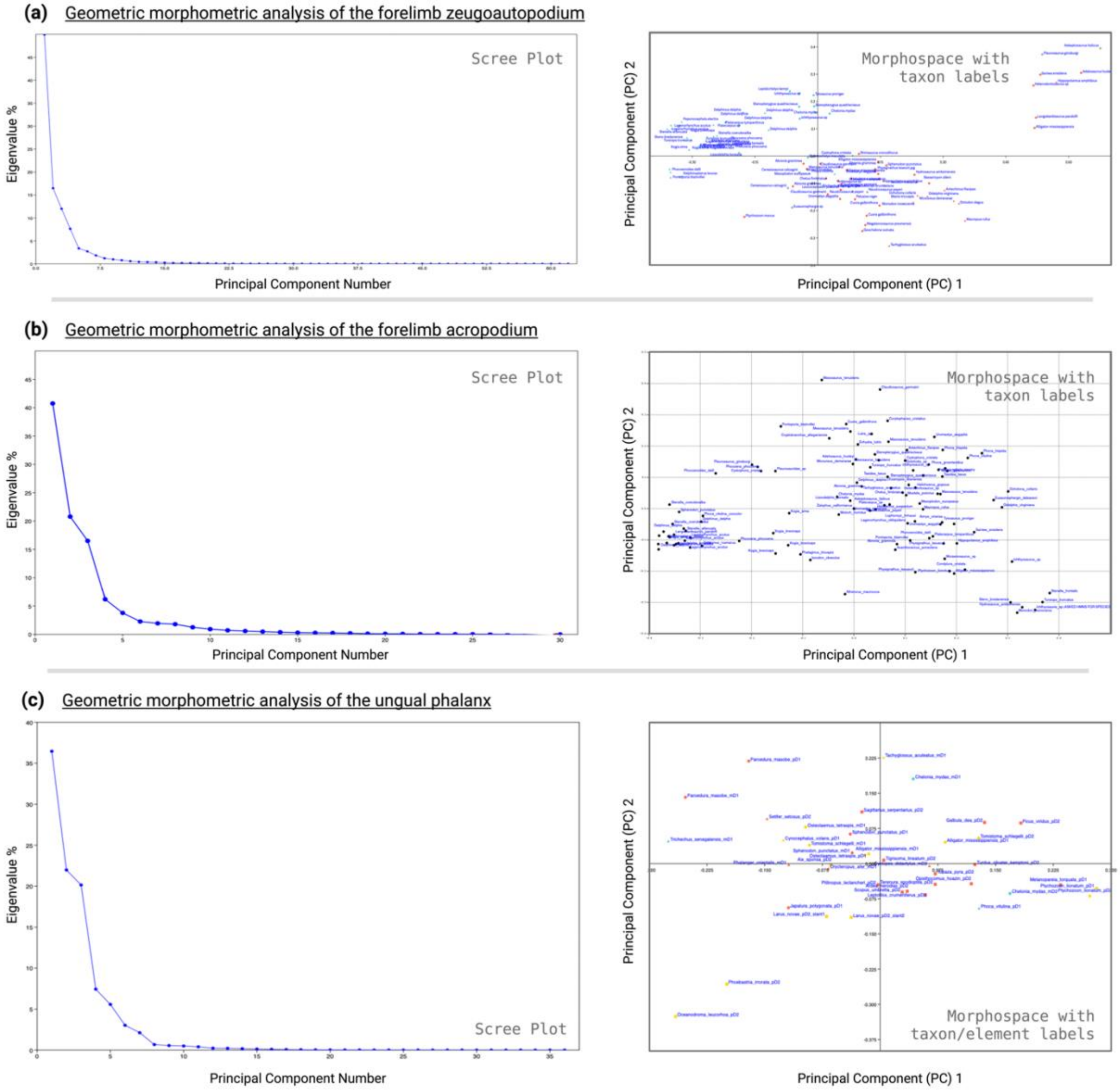


**Figure S11. Scree plots and morphospaces with taxon labels for all geometric morphometric analyses.** Scree plots show the variance captured by every principal component (PC) following principal component analysis (PCA) of the Procrustes coordinates for all specimens; morphospaces show taxon labels for PC 1 vs. PC 2 plots shown in Figure S10. These results are shown for geometric morphometric analyses of **(a)** the distal forelimb (forelimb zeugoautopod) skeleton, **(b)** the forelimb acropodial skeleton, and **(c)** the ungual phalanx. Higher-resolution plots are available in Data S6.

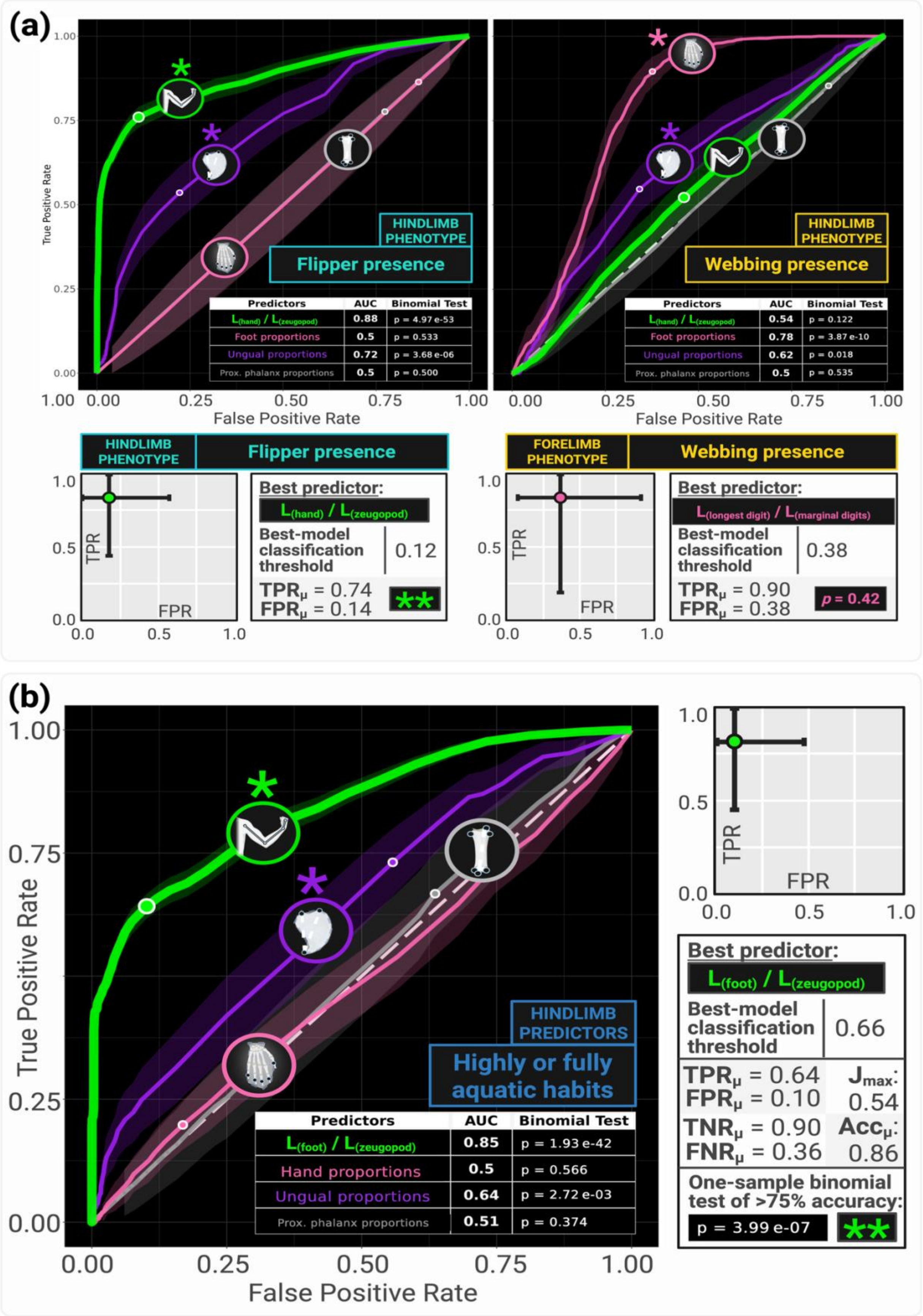
(a)
True Positive Rate
False Positive Rate
HINDLIMB PHENOTYPE
Flipper presence
Predictors | AUC | Binomial Test
L(hand) / L(zeugopod) | 0.88 | p = 4.97 e-53
Foot proportions | 0.5 | p = 0.533
Ungual proportions | 0.72 | p = 3.68 e-06
Prox. phalanx proportions | 0.5 | p = 0.500
HINDLIMB PHENOTYPE
Webbing presence
Predictors | AUC | Binomial Test
L(hand) / L(zeugopod) | 0.54 | p = 0.122
Foot proportions | 0.78 | p = 3.87 e-10
Ungual proportions | 0.62 | p = 0.018
Prox. phalanx proportions | 0.5 | p = 0.535
HINDLIMB PHENOTYPE
Flipper presence
TPR
FPR
Best predictor:
L(hand) / L(zeugopod)
Best-model classification threshold 0.12
TPRμ = 0.74
FPRμ = 0.14
**
FORELIMB PHENOTYPE
Webbing presence
TPR
FPR
Best predictor:
L(longest digit) / L(marginal digits)
Best-model classification threshold 0.38
TPRμ = 0.90
FPRμ = 0.38
p = 0.42
(b)
True Positive Rate
False Positive Rate
HINDLIMB PREDICTORS
Highly or fully aquatic habits
Predictors | AUC | Binomial Test
L(foot) / L(zeugopod) | 0.85 | p = 1.93 e-42
Hand proportions | 0.5 | p = 0.566
Ungual proportions | 0.64 | p = 2.72 e-03
Prox. phalanx proportions | 0.51 | p = 0.374
TPR
FPR
Best predictor:
L(foot) / L(zeugopod)
Best-model classification threshold 0.66
TPRμ = 0.64
FPRμ = 0.10
Jmax: 0.54
TNRμ = 0.90
FNRμ = 0.36
Accμ: 0.86
One-sample binomial test of >75% accuracy:
p = 3.99 e-07
**

**Figure S12 (previous page). Consensus Receiver-Operating Characteristic (ROC) curves and associated model performance data for hindlimb features.** ROC plots compare the accuracies of limb region proportions (green), acropodial planform shape (pink), ungual dimensions (purple), and proximal phalanx shape (gray) for **(a)** webbing and flipper phenotypes and **(b)** highly or fully aquatic habits, in amniotes with known phenotypes. A white circle indicates the best model for a given predictor, and a single asterisk indicates that the area under the curve (AUC) is significantly higher than 0.5 (one-tailed binomial test, $p < 0.05$). Characteristics of the best predictive model are shown for each phenotype, noting the model's optimal threshold probability for classifying binary phenotypes, and its mean and 95% confidence interval values for true and false positive rate. Double asterisks indicate that the best model has a success rate significantly higher than 75% (one-tailed binomial test, $p < 0.05$). Abbreviations: Acc., accuracy; AUC, area under the ROC curve; FNR, false negative rate; FPR, false positive rate; $J_{max}$, maximum Youden's J value; L, length; p, pedal; prox., proximal; TNR, true negative rate; TPR, true positive rate.

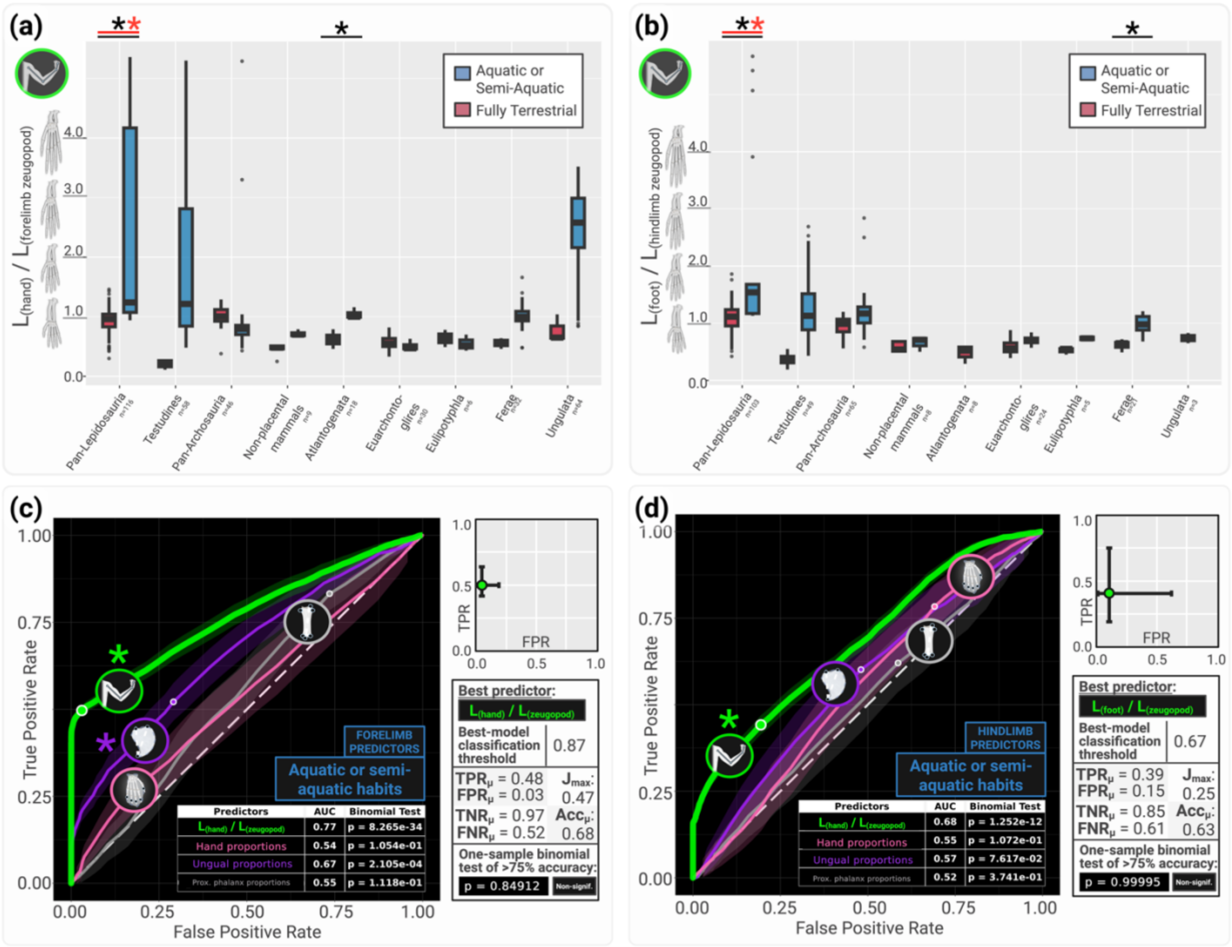


**Figure S13. Morphometric differences and predictive accuracy of acropodium-zeugopodium length for aquatic or semi-aquatic habits. (a–b)** Box plots grouped by clade, showing differences in relative hand length (a) and relative foot length (b) between fully terrestrial and (semi-)aquatic taxa. Asterisks mark significant phylogenetic ANOVAs (black) and Levene's tests (red). **(c–d)** Corresponding consensus Receiver-Operating Characteristic (ROC) curves show the best predictors of aquatic or semi-aquatic habits across amniotes. A single asterisk indicates an area under the curve (AUC) significantly higher than 0.5, and a white circle marks the best-performing model for a given predictor. Double asterisks note a best model success rate significantly > 75%. Abbreviations: Acc., accuracy; FNR, false negative rate; FPR, false positive rate; $J_{max}$, maximum Youden's J value; L, length; TNR, true negative rate; TPR, true positive rate.

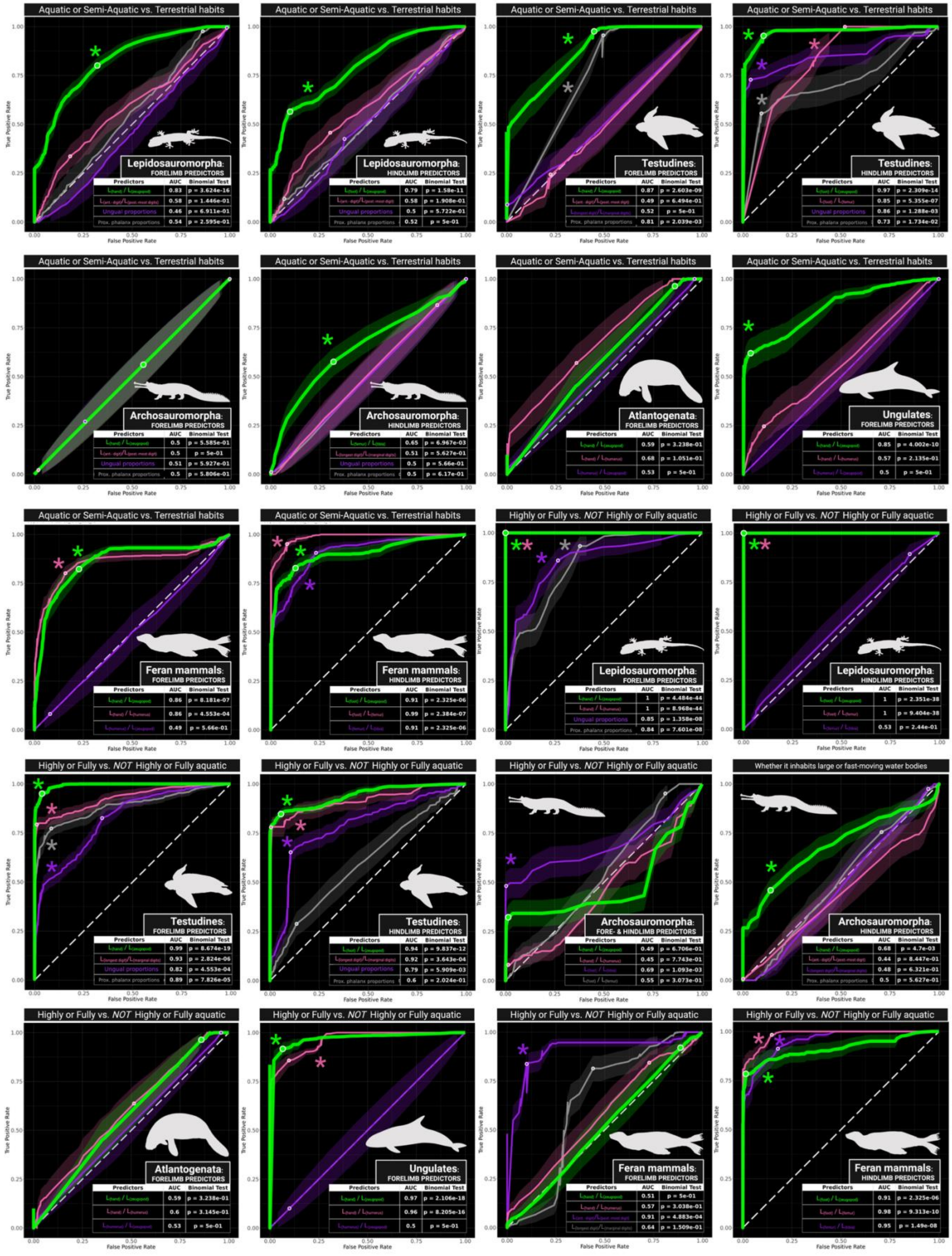
Aquatic or Semi-Aquatic vs. Terrestrial habits
Lepidosauromorpha: FORELIMB PREDICTORS
Predictors | AUC | Binomial Test
0.83 | p = 3.624e-16
0.58 | p = 1.446e-01
Ungual proportions | 0.46 | p = 6.911e-01
Prox. phalanx proportions | 0.54 | p = 2.595e-01
Aquatic or Semi-Aquatic vs. Terrestrial habits
Lepidosauromorpha: HINDLIMB PREDICTORS
Predictors | AUC | Binomial Test
0.79 | p = 1.58e-11
0.58 | p = 1.908e-01
Ungual proportions | 0.5 | p = 5.722e-01
Prox. phalanx proportions | 0.52 | p = 5e-01
Aquatic or Semi-Aquatic vs. Terrestrial habits
Testudines: FORELIMB PREDICTORS
Predictors | AUC | Binomial Test
0.87 | p = 2.603e-09
0.49 | p = 6.494e-01
0.52 | p = 5e-01
Prox. phalanx proportions | 0.81 | p = 2.039e-03
Aquatic or Semi-Aquatic vs. Terrestrial habits
Testudines: HINDLIMB PREDICTORS
Predictors | AUC | Binomial Test
0.97 | p = 2.309e-14
0.85 | p = 5.355e-07
Ungual proportions | 0.86 | p = 1.288e-03
Prox. phalanx proportions | 0.73 | p = 1.734e-02
Aquatic or Semi-Aquatic vs. Terrestrial habits
Archosauromorpha: FORELIMB PREDICTORS
Predictors | AUC | Binomial Test
0.5 | p = 5.585e-01
0.5 | p = 5e-01
Ungual proportions | 0.51 | p = 5.927e-01
Prox. phalanx proportions | 0.5 | p = 5.806e-01
Aquatic or Semi-Aquatic vs. Terrestrial habits
Archosauromorpha: HINDLIMB PREDICTORS
Predictors | AUC | Binomial Test
0.65 | p = 6.967e-03
0.51 | p = 5.627e-01
Ungual proportions | 0.5 | p = 5.66e-01
Prox. phalanx proportions | 0.5 | p = 6.17e-01
Aquatic or Semi-Aquatic vs. Terrestrial habits
Atlantogenata: FORELIMB PREDICTORS
Predictors | AUC | Binomial Test
0.59 | p = 3.238e-01
0.68 | p = 1.051e-01
0.53 | p = 5e-01
Aquatic or Semi-Aquatic vs. Terrestrial habits
Ungulates: FORELIMB PREDICTORS
Predictors | AUC | Binomial Test
0.85 | p = 4.002e-10
0.57 | p = 2.135e-01
0.5 | p = 5e-01
Aquatic or Semi-Aquatic vs. Terrestrial habits
Feran mammals: FORELIMB PREDICTORS
Predictors | AUC | Binomial Test
0.86 | p = 8.181e-07
0.86 | p = 4.553e-04
0.49 | p = 5.66e-01
Aquatic or Semi-Aquatic vs. Terrestrial habits
Feran mammals: HINDLIMB PREDICTORS
Predictors | AUC | Binomial Test
0.91 | p = 2.325e-06
0.99 | p = 2.384e-07
0.91 | p = 2.325e-06
Highly or Fully vs. NOT Highly or Fully aquatic
Lepidosauromorpha: FORELIMB PREDICTORS
Predictors | AUC | Binomial Test
1 | p = 4.484e-44
1 | p = 8.968e-44
Ungual proportions | 0.85 | p = 1.358e-08
Prox. phalanx proportions | 0.84 | p = 7.601e-08
Highly or Fully vs. NOT Highly or Fully aquatic
Lepidosauromorpha: HINDLIMB PREDICTORS
Predictors | AUC | Binomial Test
1 | p = 2.351e-38
1 | p = 9.404e-38
0.53 | p = 2.44e-01
Highly or Fully vs. NOT Highly or Fully aquatic
Testudines: FORELIMB PREDICTORS
Predictors | AUC | Binomial Test
0.99 | p = 8.674e-19
0.93 | p = 2.824e-06
Ungual proportions | 0.82 | p = 4.553e-04
Prox. phalanx proportions | 0.89 | p = 7.826e-05
Highly or Fully vs. NOT Highly or Fully aquatic
Testudines: HINDLIMB PREDICTORS
Predictors | AUC | Binomial Test
0.94 | p = 9.837e-12
0.92 | p = 3.643e-04
Ungual proportions | 0.79 | p = 5.909e-03
Prox. phalanx proportions | 0.6 | p = 2.024e-01
Highly or Fully vs. NOT Highly or Fully aquatic
Archosauromorpha: FORE- & HINDLIMB PREDICTORS
Predictors | AUC | Binomial Test
0.49 | p = 6.706e-01
0.45 | p = 7.743e-01
0.69 | p = 1.093e-03
0.55 | p = 3.073e-01
Whether it inhabits large or fast-moving water bodies
Archosauromorpha: HINDLIMB PREDICTORS
Predictors | AUC | Binomial Test
0.68 | p = 4.7e-03
0.44 | p = 8.447e-01
0.48 | p = 6.321e-01
Prox. phalanx proportions | 0.5 | p = 5.627e-01
Highly or Fully vs. NOT Highly or Fully aquatic
Atlantogenata: FORELIMB PREDICTORS
Predictors | AUC | Binomial Test
0.59 | p = 3.238e-01
0.6 | p = 3.145e-01
0.53 | p = 5e-01
Highly or Fully vs. NOT Highly or Fully aquatic
Ungulates: FORELIMB PREDICTORS
Predictors | AUC | Binomial Test
0.97 | p = 2.106e-18
0.96 | p = 8.205e-16
0.5 | p = 5e-01
Highly or Fully vs. NOT Highly or Fully aquatic
Feran mammals: FORELIMB PREDICTORS
Predictors | AUC | Binomial Test
0.51 | p = 5e-01
0.57 | p = 3.038e-01
0.91 | p = 4.883e-04
0.64 | p = 1.509e-01
Highly or Fully vs. NOT Highly or Fully aquatic
Feran mammals: HINDLIMB PREDICTORS
Predictors | AUC | Binomial Test
0.91 | p = 2.325e-06
0.98 | p = 9.313e-10
0.95 | p = 1.49e-08
True Positive Rate
False Positive Rate

**Figure S14 (previous page). Consensus Receiver-Operating Characteristic (ROC) curves showing the best predictors of different binary aquatic affinity classifications for each clade.** Different curves highlight clade-specific predictive abilities for different previously proposed correlates of aquatic affinity. Results for three different binary classification systems are shown; each is described in the STARR Methods and in Figs. S6–S7. Note that curve colors differ by ROC plot as shown. For each plot, a single asterisk indicates that the area under the curve (AUC) is significantly greater than 0.5, and each white circle marks the best-performing model for a given predictor (that model with the phenotype classification threshold for which Youden's J is maximized). The clade-specific performance of some osteological correlates could not be estimated; for example, there are no highly or fully aquatic atlantogenatans or ungulates that retain hindlimbs, so we could not assess the predictive accuracy of any hindlimb metrics for highly or fully aquatic habits in these taxa. Abbreviations: AUC, area under the ROC curve; prox., proximal. PhyloPic attributions are provided in Text S19.

**Figure S15 (next page). Consensus Receiver-Operating Characteristic (ROC) curves showing the best predictors of soft-tissue limb phenotype (webbing or flippers) for each clade.** Different curves highlight clade-specific predictive abilities for different previously proposed correlates. A single asterisk indicates that the area under the curve (AUC) is significantly greater than 0.5, and each white circle marks the best-performing model for a given predictor (that model with the phenotype classification threshold for which Youden's J is maximized). Note that curve colors differ by ROC plot as shown. The clade-specific performance of some osteological correlates could not be estimated; for example, no atlantogenatan mammals have hind-flippers, so we could not assess hind-flipper predictability for Atlantogenata in ROC space. Abbreviations: AUC, area under the ROC curve; prox., proximal. PhyloPic attributions are provided in Text S19.

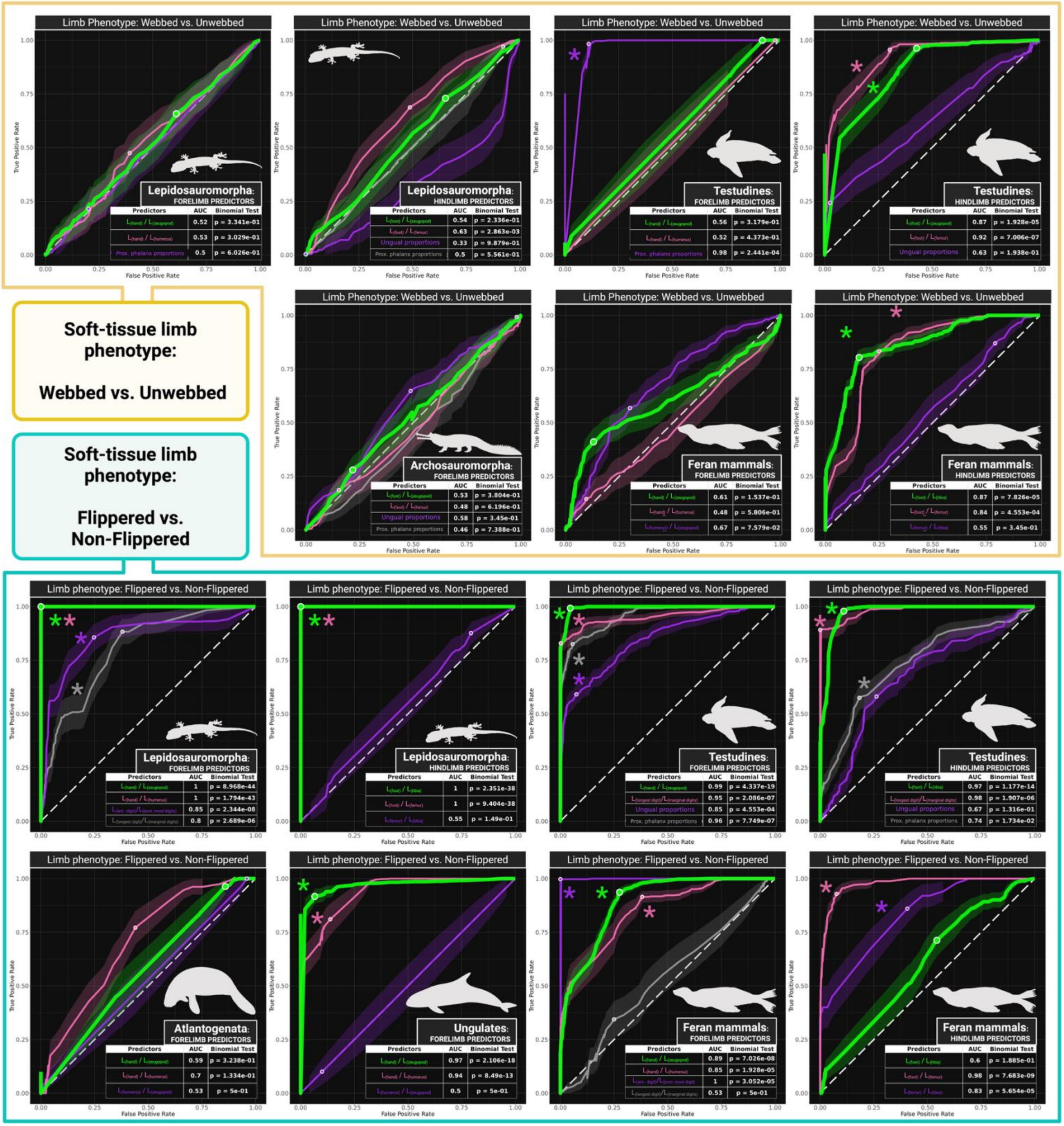
Soft-tissue limb phenotype:
Webbed vs. Unwebbed
Soft-tissue limb phenotype:
Flippered vs. Non-Flippered
Limb Phenotype: Webbed vs. Unwebbed
Limb phenotype: Flippered vs. Non-Flippered
True Positive Rate
False Positive Rate
Lepidosauromorpha: FORELIMB PREDICTORS
Lepidosauromorpha: HINDLIMB PREDICTORS
Testudines: FORELIMB PREDICTORS
Testudines: HINDLIMB PREDICTORS
Archosauromorpha: FORELIMB PREDICTORS
Feran mammals: FORELIMB PREDICTORS
Feran mammals: HINDLIMB PREDICTORS
Atlantogenata: FORELIMB PREDICTORS
Ungulates: FORELIMB PREDICTORS
Predictors
AUC
Binomial Test
Ungual proportions
Prox. phalanx proportions

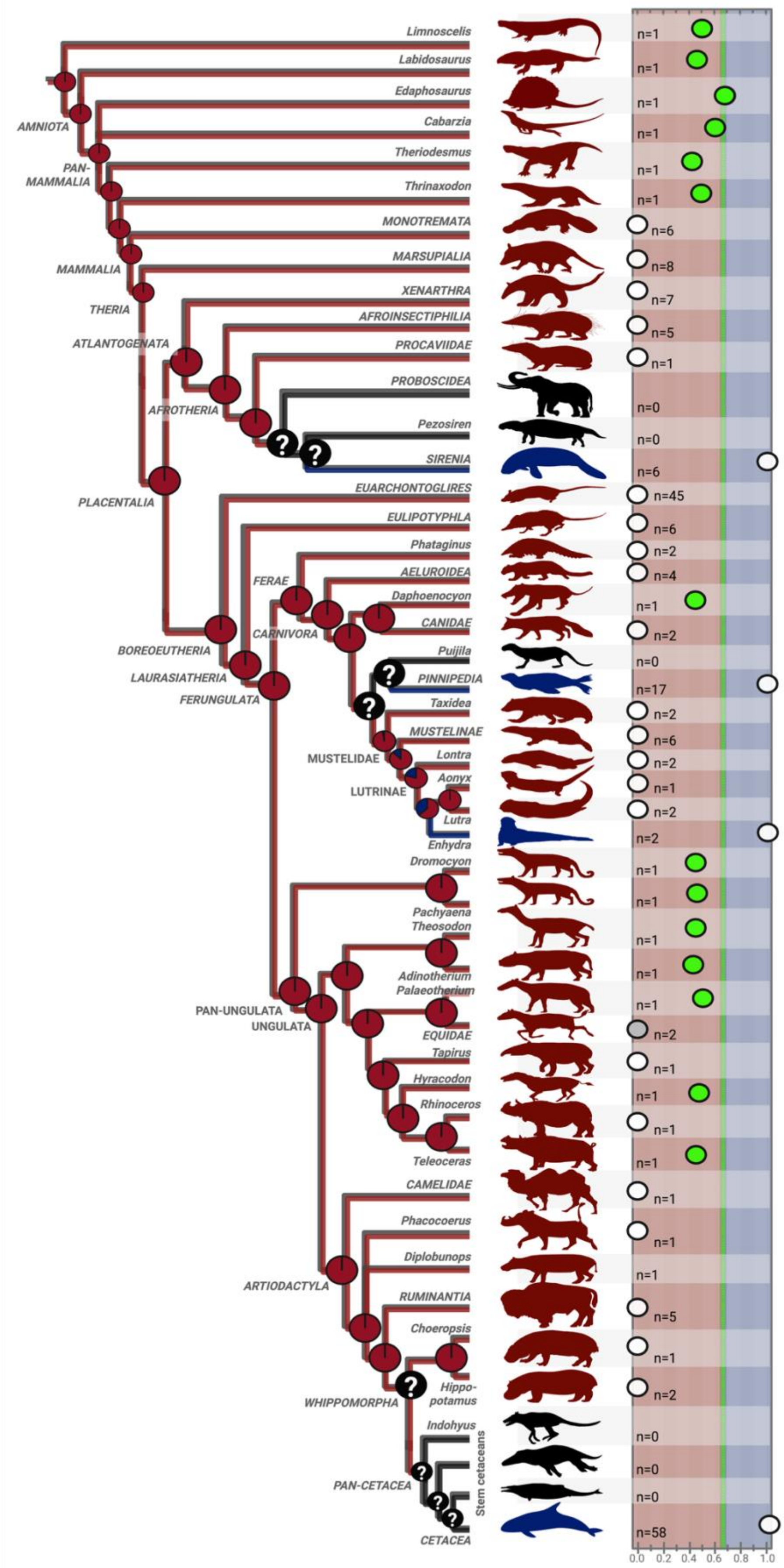
Limnoscelis
Labidosaurus
Edaphosaurus
Cabarzia
Theriodesmus
Thrinaxodon
MONOTREMATA
MARSUPIALIA
XENARTHRA
AFROINSECTIPHILIA
PROCAVIIDAE
PROBOSCIDEA
Pezosiren
SIRENIA
EUARCHONTOGLIRES
EULIPOTYPHLA
Phataginus
AELUROIDEA
Daphoenocyon
CANIDAE
Puijila
PINNIPEDIA
Taxidea
MUSTELINAE
Lontra
Aonyx
Lutra
Enhydra
Dromocyon
Pachyaena
Theosodon
Adinotherium
Palaeotherium
EQUIDAE
Tapirus
Hyracodon
Rhinoceros
Teleoceras
CAMELIDAE
Phacocoerus
Diplobunops
RUMINANTIA
Choeropsis
Hippo-
potamus
Indohyus
Stem cetaceans
CETACEA
AMNIOTA
PAN-
MAMMALIA
MAMMALIA
THERIA
ATLANTOGENATA
AFROTHERIA
PLACENTALIA
FERAE
CARNIVORA
BOREOEUTHERIA
LAURASIATHERIA
FERUNGULATA
MUSTELIDAE
LUTRINAE
PAN-UNGULATA
UNGULATA
ARTIODACTYLA
WHIPPOMORPHA
PAN-CETACEA
n=1
n=1
n=1
n=1
n=1
n=1
n=6
n=8
n=7
n=5
n=1
n=0
n=0
n=6
n=45
n=6
n=2
n=4
n=1
n=2
n=0
n=17
n=2
n=6
n=2
n=1
n=2
n=2
n=1
n=1
n=1
n=1
n=1
n=2
n=1
n=1
n=1
n=1
n=1
n=1
n=1
n=5
n=1
n=2
n=0
n=0
n=0
n=58
0.0 0.2 0.4 0.6 0.8 1.0

**Figure S16 (previous page). Reconstructed evolutionary history of highly or fully aquatic habits in pan-mammals.** Our best-performing model (using $L_{hand}$ / $L_{zeugopodium}$ values) gives the predicted probabilities (from 0 to 1) of highly/fully aquatic habits (dark blue) in extinct pan-mammals. Green circles indicate mean phenotype probabilities predicted by our model. White circles indicate phenotype probabilities that were fixed *a priori* based on morphological considerations or direct observation of extant species. Gray circles indicate phenotype probabilities that were derived from a combination of fixed and predicted tree tips using ancestral-state reconstructions. The dashed green line indicates the probability classification threshold for the associated model. Tip taxa with recovered phenotype probabilities significantly higher than the classification threshold (one-tailed t-test, $p < 0.05$) are colored accordingly, as either dark red (possibly semi-aquatic, but regularly returning to land) or dark blue (highly or fully aquatic, seldom if ever leaving the water). Pie charts and branch colors show ancestral-state reconstructions for aquatic habits given tip phenotypes. Taxa in black are unsampled, and black nodes have ambiguous reconstructions that depend on unsampled taxa. The absence of stem-group sirenians, pinnipeds, and cetaceans in our dataset highlights an opportunity for future studies to investigate the evolutionary history of highly/fully aquatic habits in mammals. PhyloPic attributions are provided in Text S19.

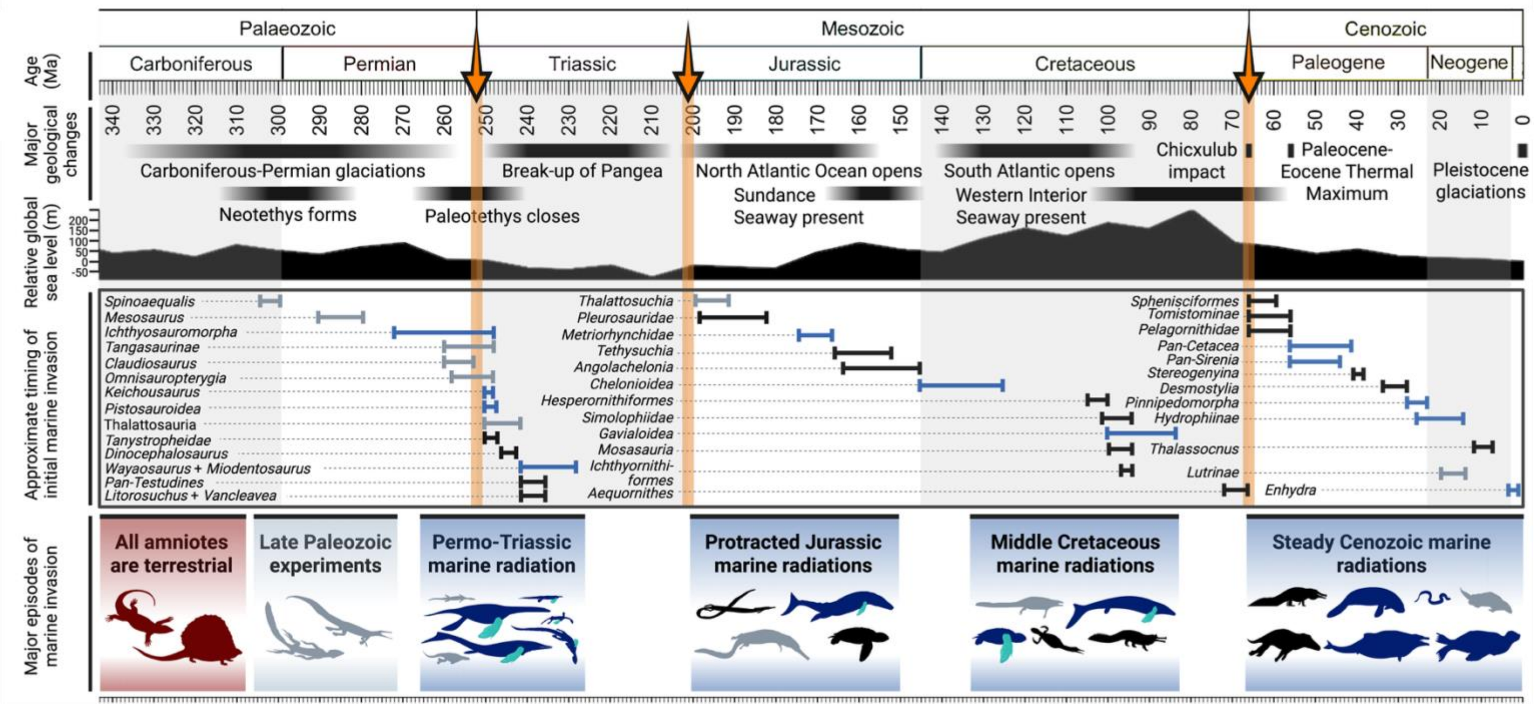


**Figure S17. Approximate timing and geological context for the invasion of aquatic environments by amniotes, with a focus on taxa analyzed in this study.** Geological context for amniote marine invasions. Orange arrows mark major mass extinction events; timings—given the trees above—of initial marine invasions (dark red) and initial commitments to a highly or fully aquatic lifestyle (dark blue) are shown for the Paleozoic, Mesozoic, and Cenozoic in chronological order from top to bottom, revealing broad-scale temporal patterns in the utilization of marine environments by amniotes. All ranges shown for clade origins and marine invasions are approximate ranges synthesized from previous studies. They are not the results of original analyses from this study, and are intended only to illustrate broad-scale trends. More information on these ranges for clade origination times are provided in Text S14 and Table S10. PhyloPic attributions are provided in Text S19.

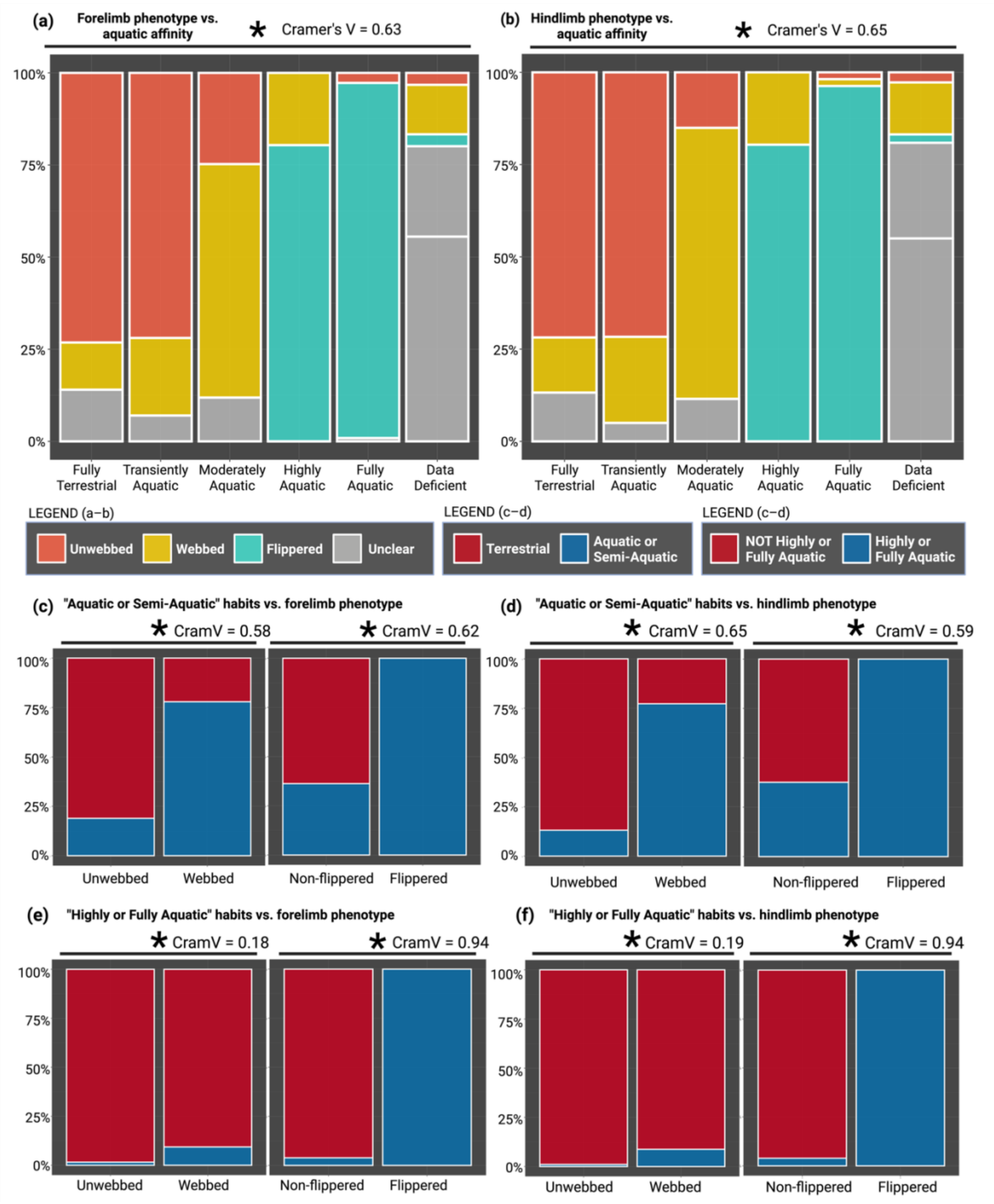


**Figure S18. Category association plots showing the percentage overlap between the major aquatic affinity and limb phenotype bins in our dataset. (a–b)** Category associations between aquatic habits and soft-tissue limb phenotypes for forelimbs (a) and hindlimbs (b) show significant overlap among groups. **(c–d)** Binary category associations between "aquatic or semi-aquatic" habits and limb phenotypes show similar association strengths for aquatic habits and flippers. **(e–f)** Binary category associations between "highly or fully aquatic" habits and limb phenotypes show greater association strengths for flippers. Cramer's V statistic (abbreviated "CramV") reports the strength of association between two categorical variables (from 0 to 1). Asterisks report that these associations are significant (Fisher's exact test: $p < 0.05$).

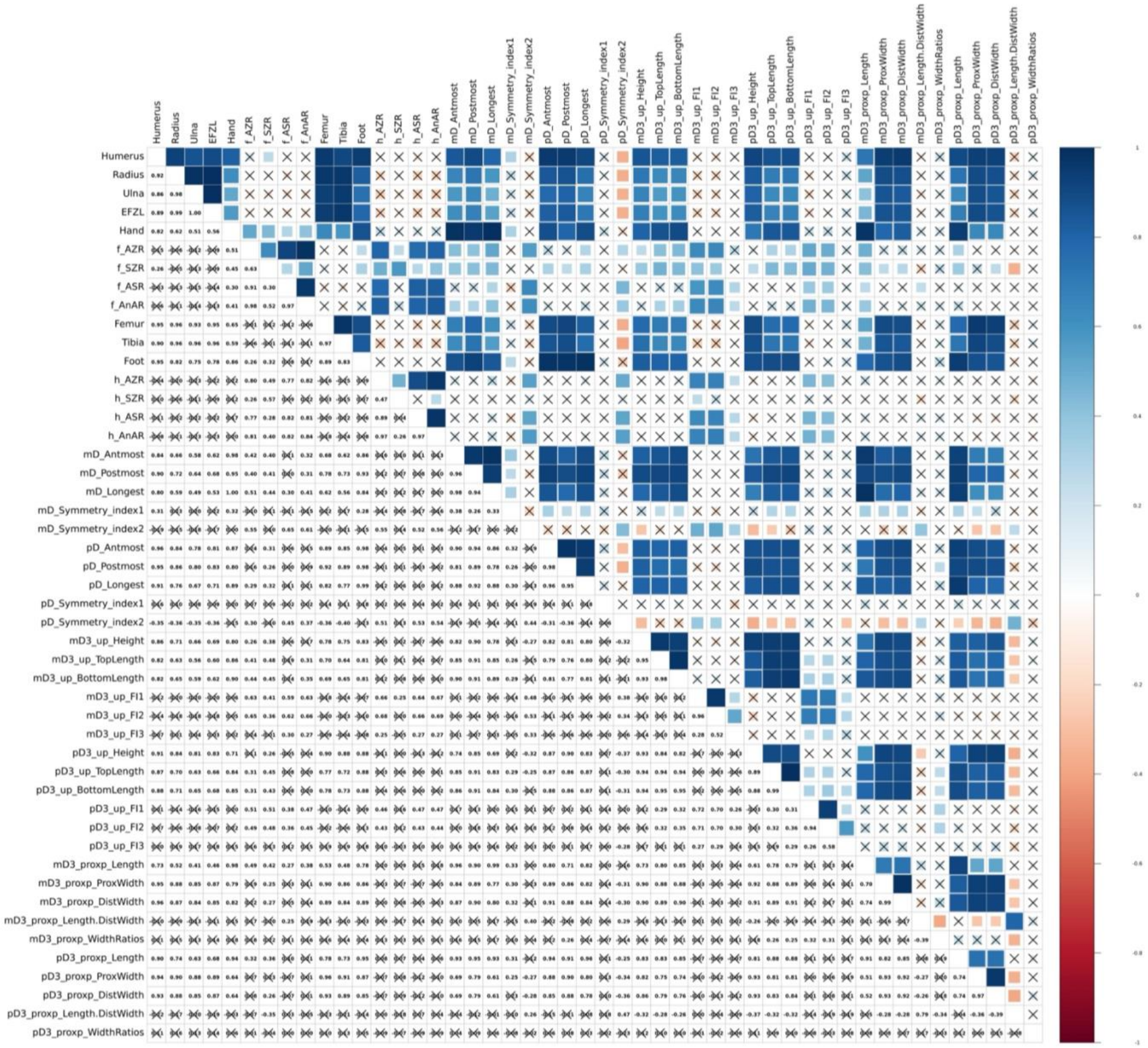


**Figure S19. Correlation matrix for all quantitative variables in our dataset, prior to removing phylogenetic autocorrelation.** Pearson correlation coefficients scale from 0 (dark red) to 1 (dark blue), and an "X" marks each non-significant correlation ($p > 0.05$). Abbreviations: AZR, acropod-zeugopodium ratio; AnAR, acropod-to-non-acropodium length ratio (acropodium length divided by the average of all stylopodial and zeugopodial lengths); Antmost, anterior-most; ASR, acropod-stylopodium ratio; BottomLength, length of the ungual about its ventral margin; DistWidth, width of the distal epiphysis; FI, flatness index; mD, manual digit; mD3, manual digit 3; pD, pedal digit; pD3, pedal digit 3; Postmost, posterior-most; proxp, proximal phalanx; ProxWidth, width of the proximal epiphysis; TopLength, length of the ungual about its dorsal margin; up, ungual phalanx; WidthRatios, ratio of the width of the distal epiphysis to the length of the proximal epiphysis. Note that a "." above replaces a "/", so "Length.DistWidth" should be read as the fraction "Length/DistWidth." All these metrics are defined and related to the main-text results within script S1. A higher-resolution version of this plot is available in the supplementary files associated with Data S3.

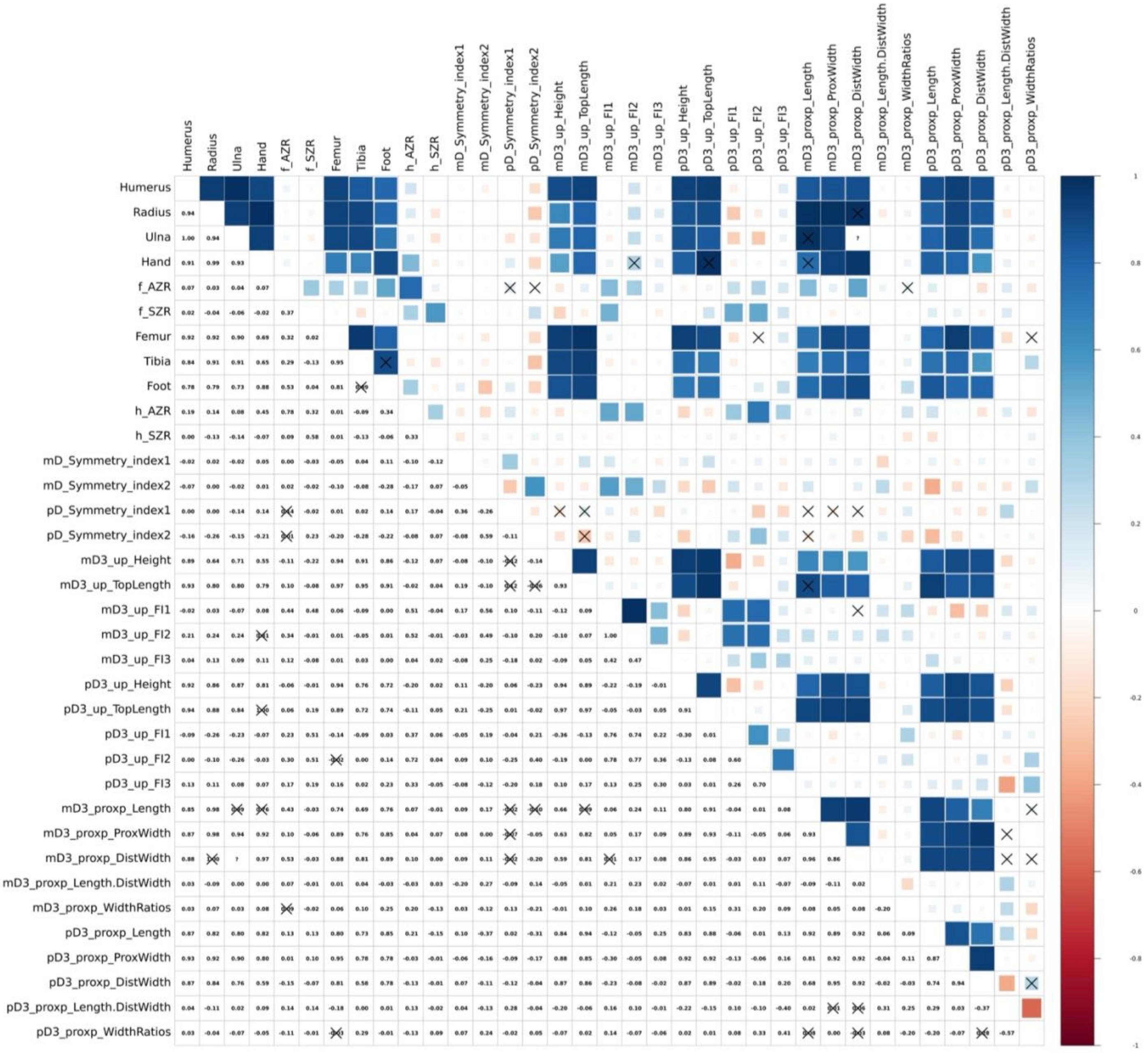


**Figure S20. Phylogenetic correlation matrix for all quantitative variables in our dataset under supertree 1.** Pearson correlation coefficients scale from 0 (dark red) to 1 (dark blue), and an "X" marks each non-significant correlation ($p > 0.05$). Abbreviations: AZR, acropod-zeugopodium ratio; AnAR, acropod-to-non-acropodium length ratio (acropodium length divided by the average of all stylopodial and zeugopodial lengths); Antmost, anterior-most; ASR, acropod-stylopodium ratio; BottomLength, length of the ungual about its ventral margin; DistWidth, width of the distal epiphysis; FI, flatness index; mD, manual digit; mD3, manual digit 3; pD, pedal digit; pD3, pedal digit 3; Postmost, posterior-most; proxp, proximal phalanx; ProxWidth, width of the proximal epiphysis; TopLength, length of the ungual about its dorsal margin; up, ungual phalanx; WidthRatios, ratio of the width of the distal epiphysis to the length of the proximal epiphysis. Note that a "." above replaces a "/", so "Length.DistWidth" should be read as the fraction "Length/DistWidth." All these metrics are defined and related to the main-text results within script S1. A higher-resolution version of this plot is available in the supplementary files associated with Data S3.

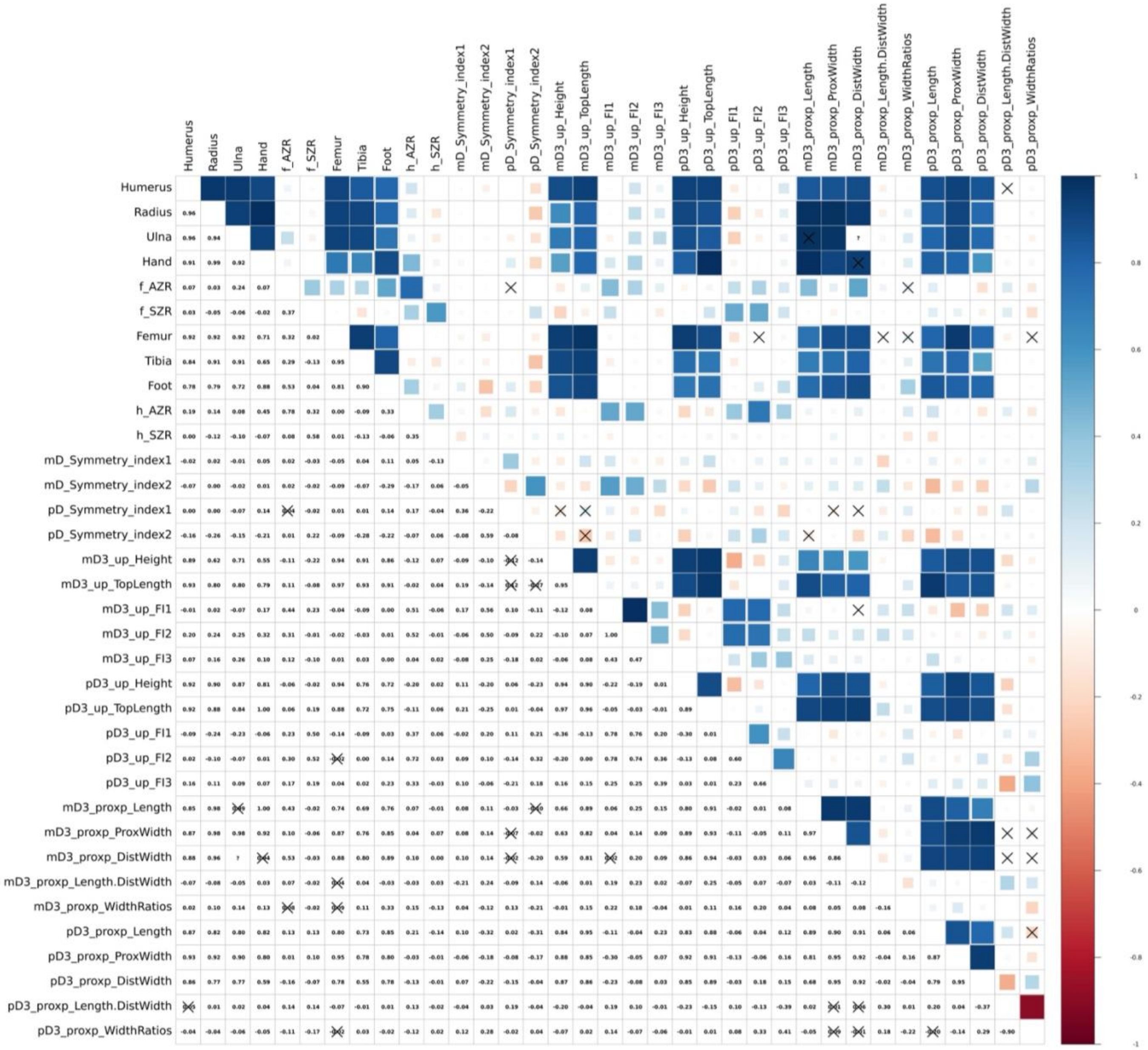

**Figure S21. Phylogenetic correlation matrix for all quantitative variables in our dataset under supertree 2.** Pearson correlation coefficients scale from 0 (dark red) to 1 (dark blue), and an "X" marks each non-significant correlation ($p > 0.05$). Abbreviations: AZR, acropod-zeugopodium ratio; AnAR, acropod-to-non-acropodium length ratio (acropodium length divided by the average of all stylopodial and zeugopodial lengths); Antmost, anterior-most; ASR, acropod-stylopodium ratio; BottomLength, length of the ungual about its ventral margin; DistWidth, width of the distal epiphysis; FI, flatness index; mD, manual digit; mD3, manual digit 3; pD, pedal digit; pD3, pedal digit 3; Postmost, posterior-most; proxp, proximal phalanx; ProxWidth, width of the proximal epiphysis; TopLength, length of the ungual about its dorsal margin; up, ungual phalanx; WidthRatios, ratio of the width of the distal epiphysis to the length of the proximal epiphysis. Note that a "." above replaces a "/", so "Length.DistWidth" should be read as the fraction "Length/DistWidth." All these metrics are defined and related to the main-text results within script S1. A higher-resolution version of this plot is available in the supplementary files associated with Data S3.

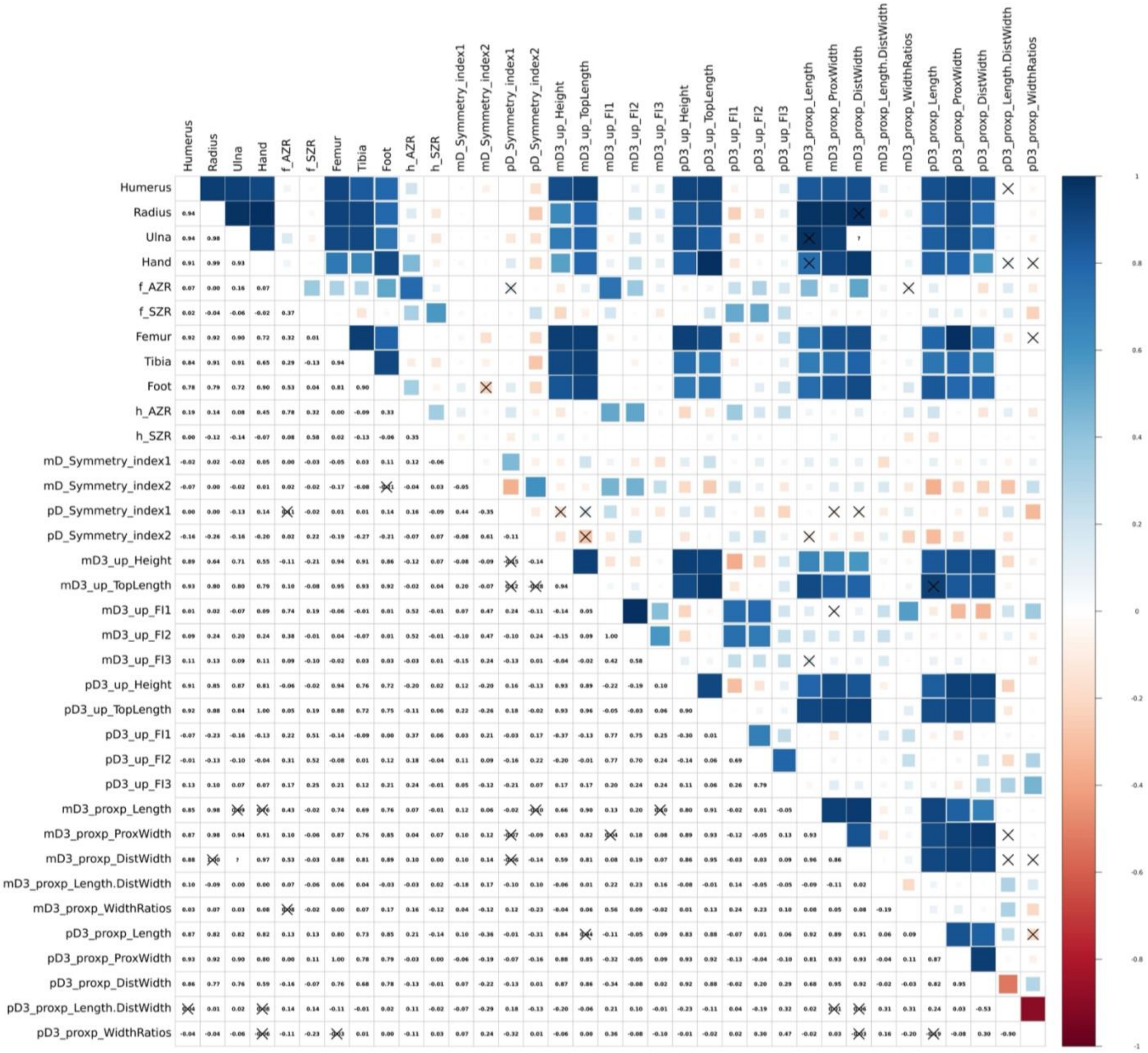


**Figure S22. Phylogenetic correlation matrix for all quantitative variables in our dataset under supertree 3.** Pearson correlation coefficients scale from 0 (dark red) to 1 (dark blue), and an "X" marks each non-significant correlation ($p > 0.05$). Abbreviations: AZR, acropod-zeugopodium ratio; AnAR, acropod-to-non-acropodium length ratio (acropodium length divided by the average of all stylopodial and zeugopodial lengths); Antmost, anterior-most; ASR, acropod-stylopodium ratio; BottomLength, length of the ungual about its ventral margin; DistWidth, width of the distal epiphysis; FI, flatness index; mD, manual digit; mD3, manual digit 3; pD, pedal digit; pD3, pedal digit 3; Postmost, posterior-most; proxp, proximal phalanx; ProxWidth, width of the proximal epiphysis; TopLength, length of the ungual about its dorsal margin; up, ungual phalanx; WidthRatios, ratio of the width of the distal epiphysis to the length of the proximal epiphysis. Note that a "." above replaces a "/", so "Length.DistWidth" should be read as the fraction "Length/DistWidth." All these metrics are defined and related to the main-text results within script S1. A higher-resolution version of this plot is available in the supplementary files associated with Data S3.

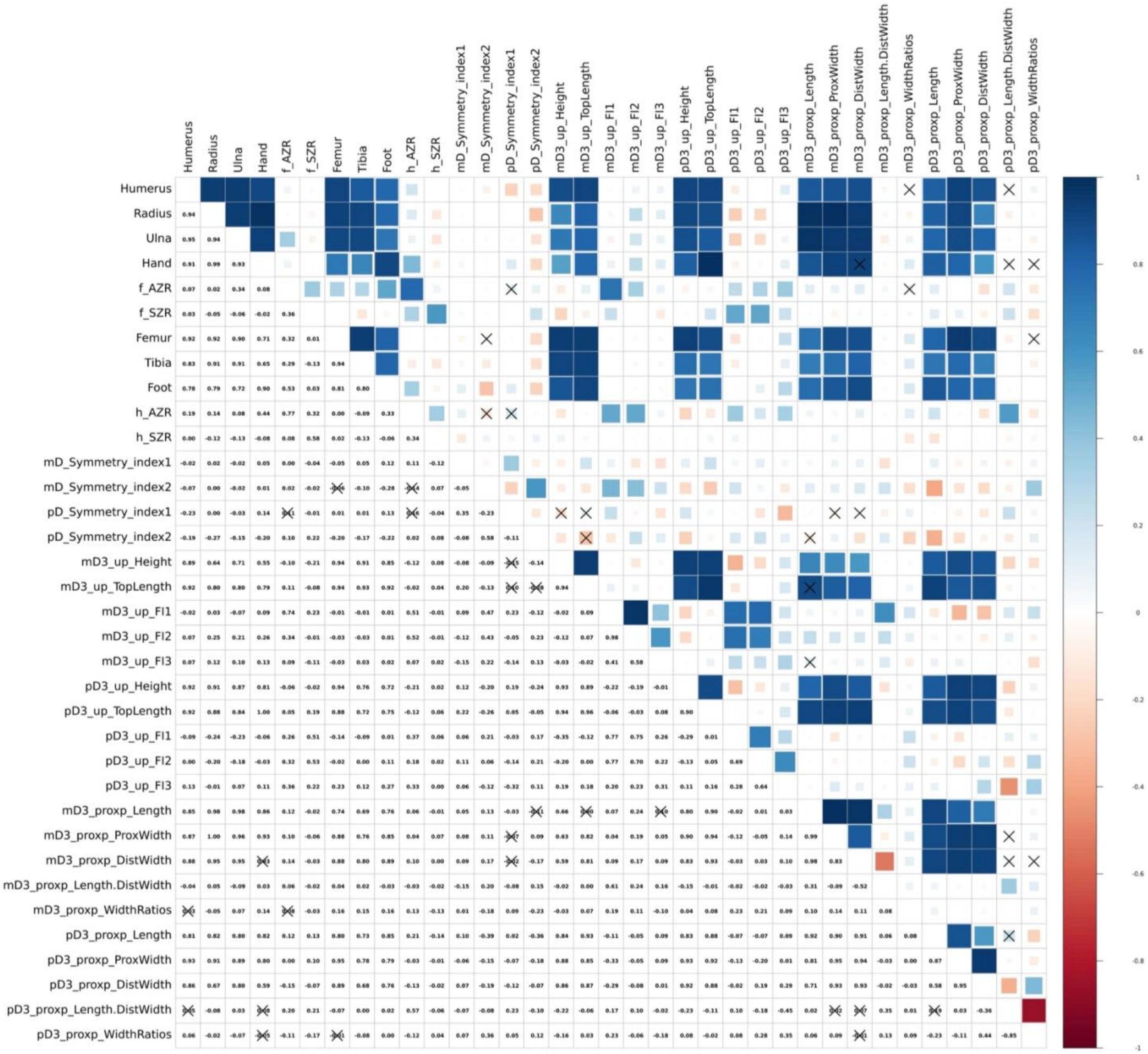


**Figure S23. Phylogenetic correlation matrix for all quantitative variables in our dataset under supertree 4.** Pearson correlation coefficients scale from 0 (dark red) to 1 (dark blue), and an "X" marks each non-significant correlation ($p > 0.05$). Abbreviations: AZR, acropod-zeugopodium ratio; AnAR, acropod-to-non-acropodium length ratio (acropodium length divided by the average of all stylopodial and zeugopodial lengths); Antmost, anterior-most; ASR, acropod-stylopodium ratio; BottomLength, length of the ungual about its ventral margin; DistWidth, width of the distal epiphysis; FI, flatness index; mD, manual digit; mD3, manual digit 3; pD, pedal digit; pD3, pedal digit 3; Postmost, posterior-most; proxp, proximal phalanx; ProxWidth, width of the proximal epiphysis; TopLength, length of the ungual about its dorsal margin; up, ungual phalanx; WidthRatios, ratio of the width of the distal epiphysis to the length of the proximal epiphysis. Note that a "." above replaces a "/", so "Length.DistWidth" should be read as the fraction "Length/DistWidth." All these metrics are defined and related to the main-text results within script S1. A higher-resolution version of this plot is available in the supplementary files associated with Data S3.

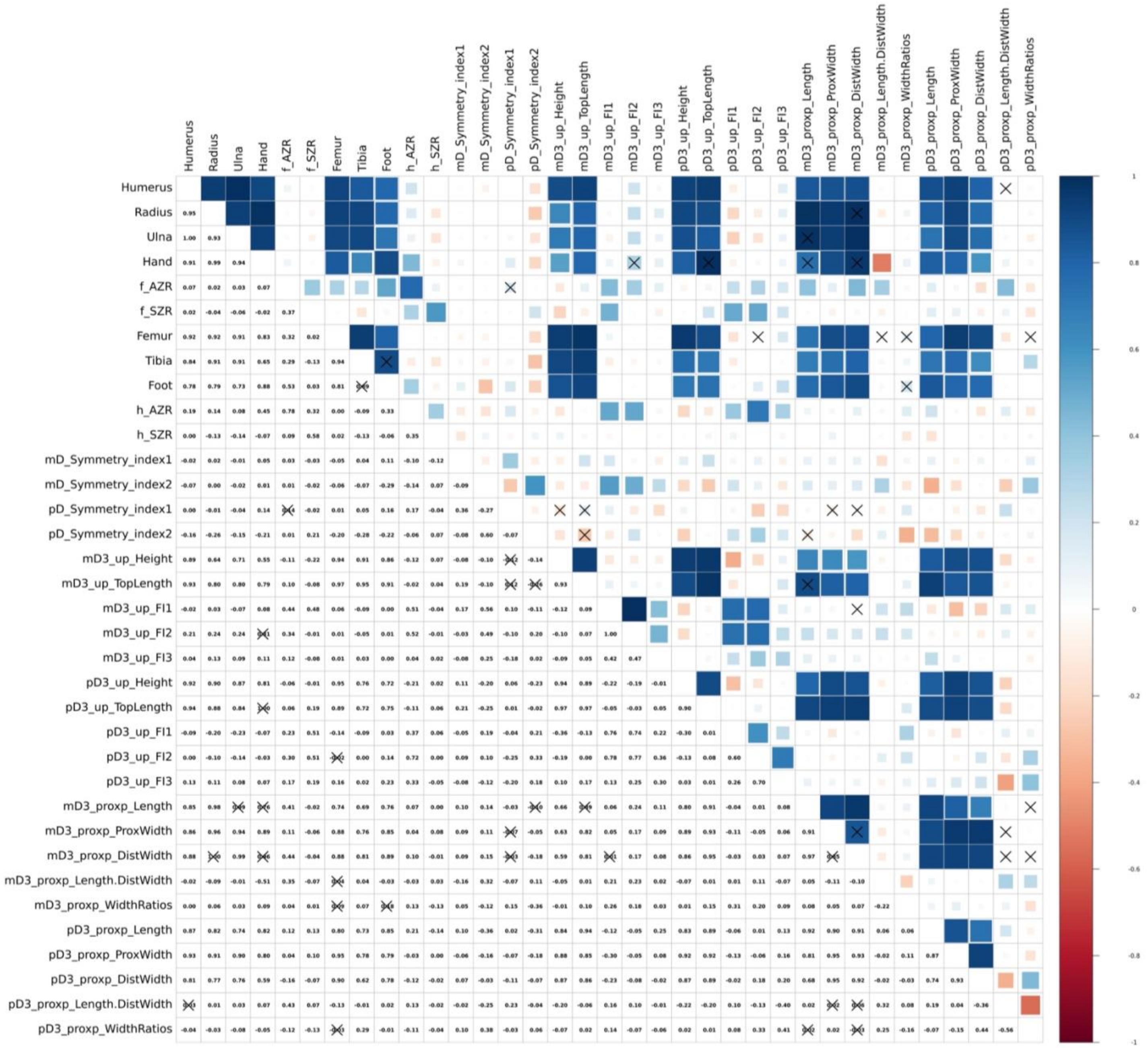


**Figure S24. Phylogenetic correlation matrix for all quantitative variables in our dataset under supertree 5.** Pearson correlation coefficients scale from 0 (dark red) to 1 (dark blue), and an "X" marks each non-significant correlation ($p > 0.05$). Abbreviations: AZR, acropod-zeugopodium ratio; AnAR, acropod-to-non-acropodium length ratio (acropodium length divided by the average of all stylopodial and zeugopodial lengths); Antmost, anterior-most; ASR, acropod-stylopodium ratio; BottomLength, length of the ungual about its ventral margin; DistWidth, width of the distal epiphysis; FI, flatness index; mD, manual digit; mD3, manual digit 3; pD, pedal digit; pD3, pedal digit 3; Postmost, posterior-most; proxp, proximal phalanx; ProxWidth, width of the proximal epiphysis; TopLength, length of the ungual about its dorsal margin; up, ungual phalanx; WidthRatios, ratio of the width of the distal epiphysis to the length of the proximal epiphysis. Note that a "." above replaces a "/", so "Length.DistWidth" should be read as the fraction "Length/DistWidth." All these metrics are defined and related to the main-text results within script S1. A higher-resolution version of this plot is available in the supplementary files associated with Data S3.

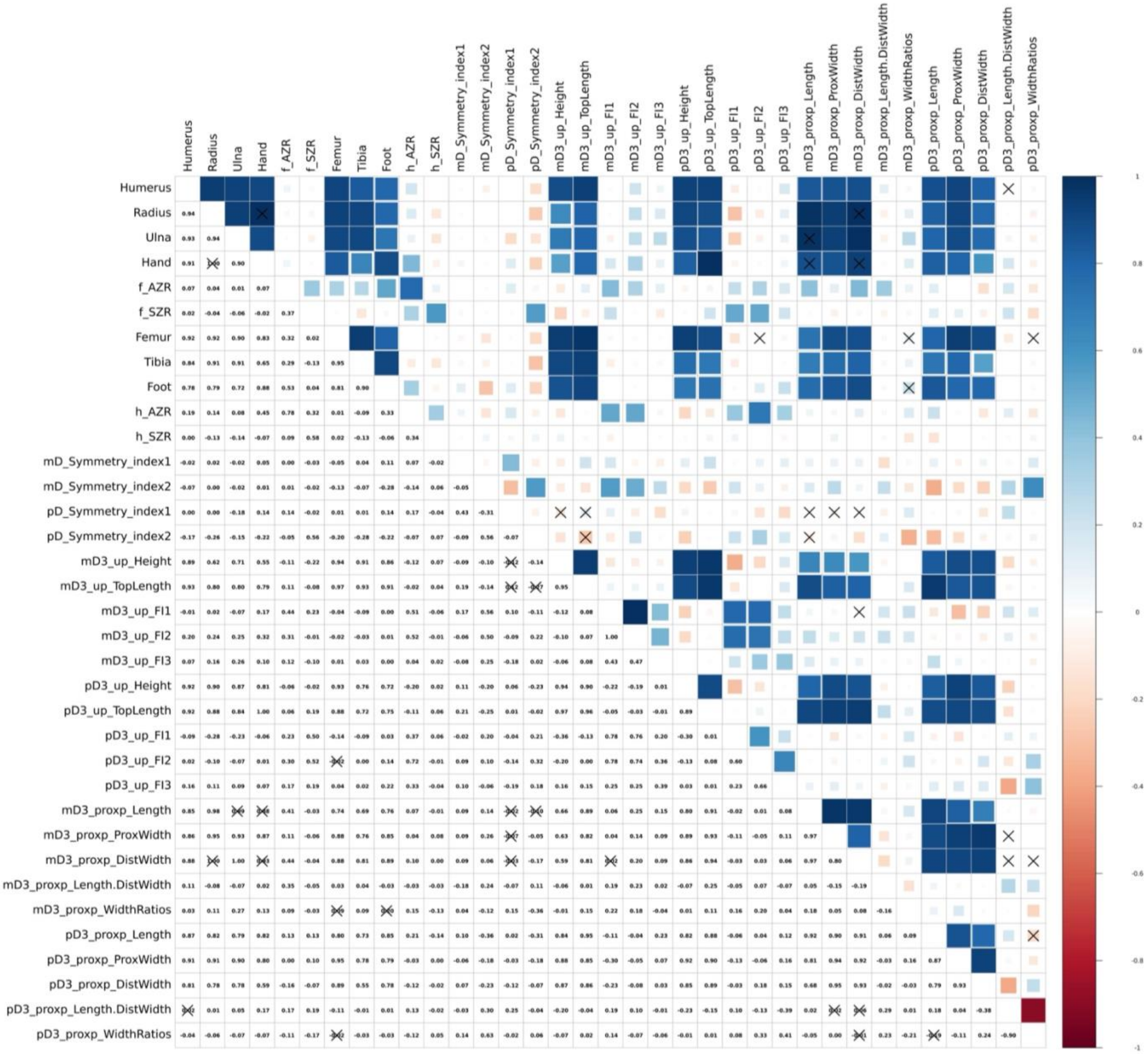


**Figure S25. Phylogenetic correlation matrix for all quantitative variables in our dataset under supertree 6.** Pearson correlation coefficients scale from 0 (dark red) to 1 (dark blue), and an "X" marks each non-significant correlation ($p > 0.05$). Abbreviations: AZR, acropod-zeugopodium ratio; AnAR, acropod-to-non-acropodium length ratio (acropodium length divided by the average of all stylopodial and zeugopodial lengths); Antmost, anterior-most; ASR, acropod-stylopodium ratio; BottomLength, length of the ungual about its ventral margin; DistWidth, width of the distal epiphysis; FI, flatness index; mD, manual digit; mD3, manual digit 3; pD, pedal digit; pD3, pedal digit 3; Postmost, posterior-most; proxp, proximal phalanx; ProxWidth, width of the proximal epiphysis; TopLength, length of the ungual about its dorsal margin; up, ungual phalanx; WidthRatios, ratio of the width of the distal epiphysis to the length of the proximal epiphysis. Note that a "." above replaces a "/", so "Length.DistWidth" should be read as the fraction "Length/DistWidth." All these metrics are defined and related to the main-text results within script S1. A higher-resolution version of this plot is available in the supplementary files associated with Data S3.

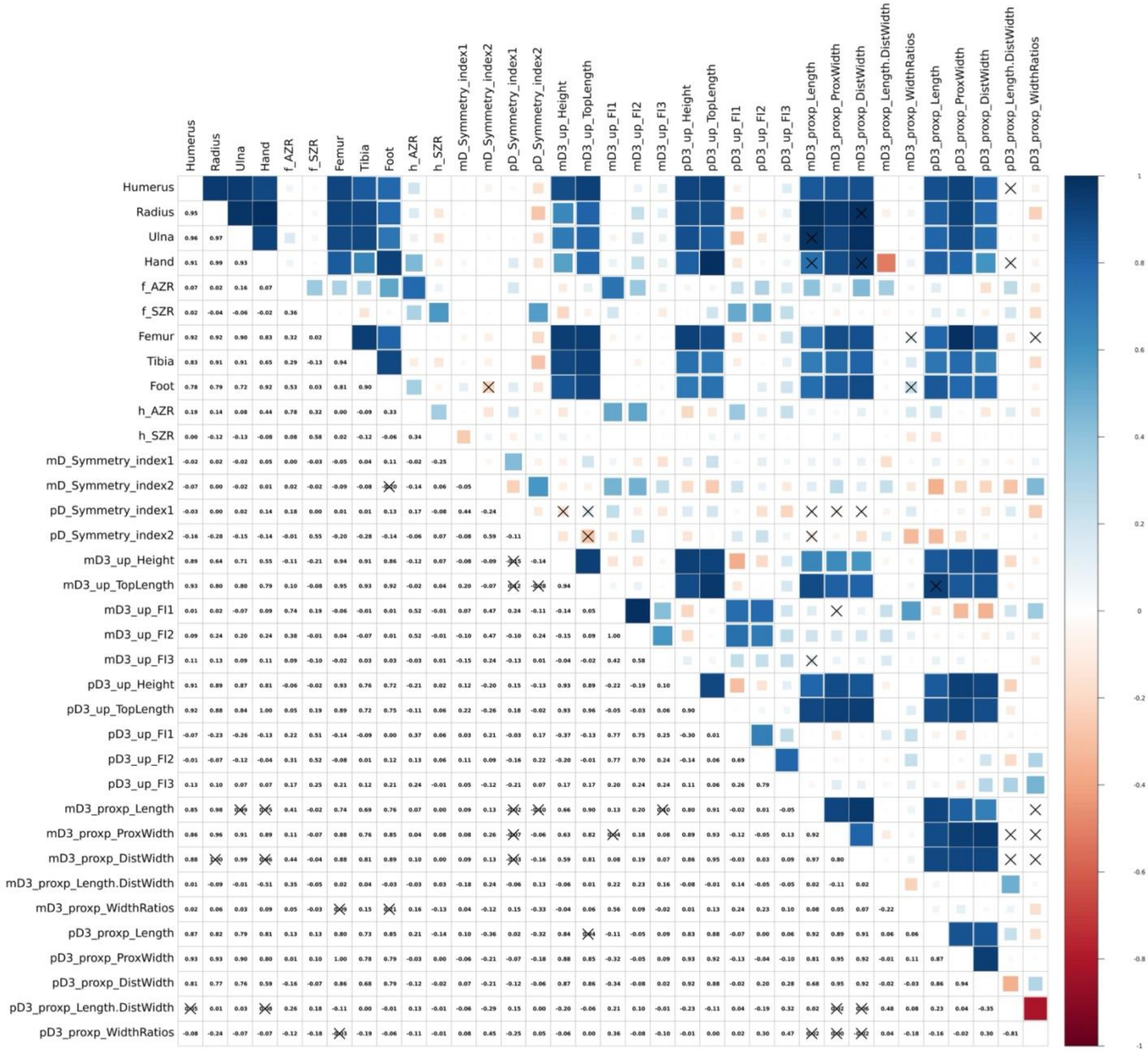

**Figure S26. Phylogenetic correlation matrix for all quantitative variables in our dataset under supertree 7.** Pearson correlation coefficients scale from 0 (dark red) to 1 (dark blue), and an "X" marks each non-significant correlation ($p > 0.05$). Abbreviations: AZR, acropod-zeugopodium ratio; AnAR, acropod-to-non-acropodium length ratio (acropodium length divided by the average of all stylopodial and zeugopodial lengths); Antmost, anterior-most; ASR, acropod-stylopodium ratio; BottomLength, length of the ungual about its ventral margin; DistWidth, width of the distal epiphysis; FI, flatness index; mD, manual digit; mD3, manual digit 3; pD, pedal digit; pD3, pedal digit 3; Postmost, posterior-most; proxp, proximal phalanx; ProxWidth, width of the proximal epiphysis; TopLength, length of the ungual about its dorsal margin; up, ungual phalanx; WidthRatios, ratio of the width of the distal epiphysis to the length of the proximal epiphysis. Note that a "." above replaces a "/", so "Length.DistWidth" should be read as the fraction "Length/DistWidth." All these metrics are defined and related to the main-text results within script S1. A higher-resolution version of this plot is available in the supplementary files associated with Data S3.

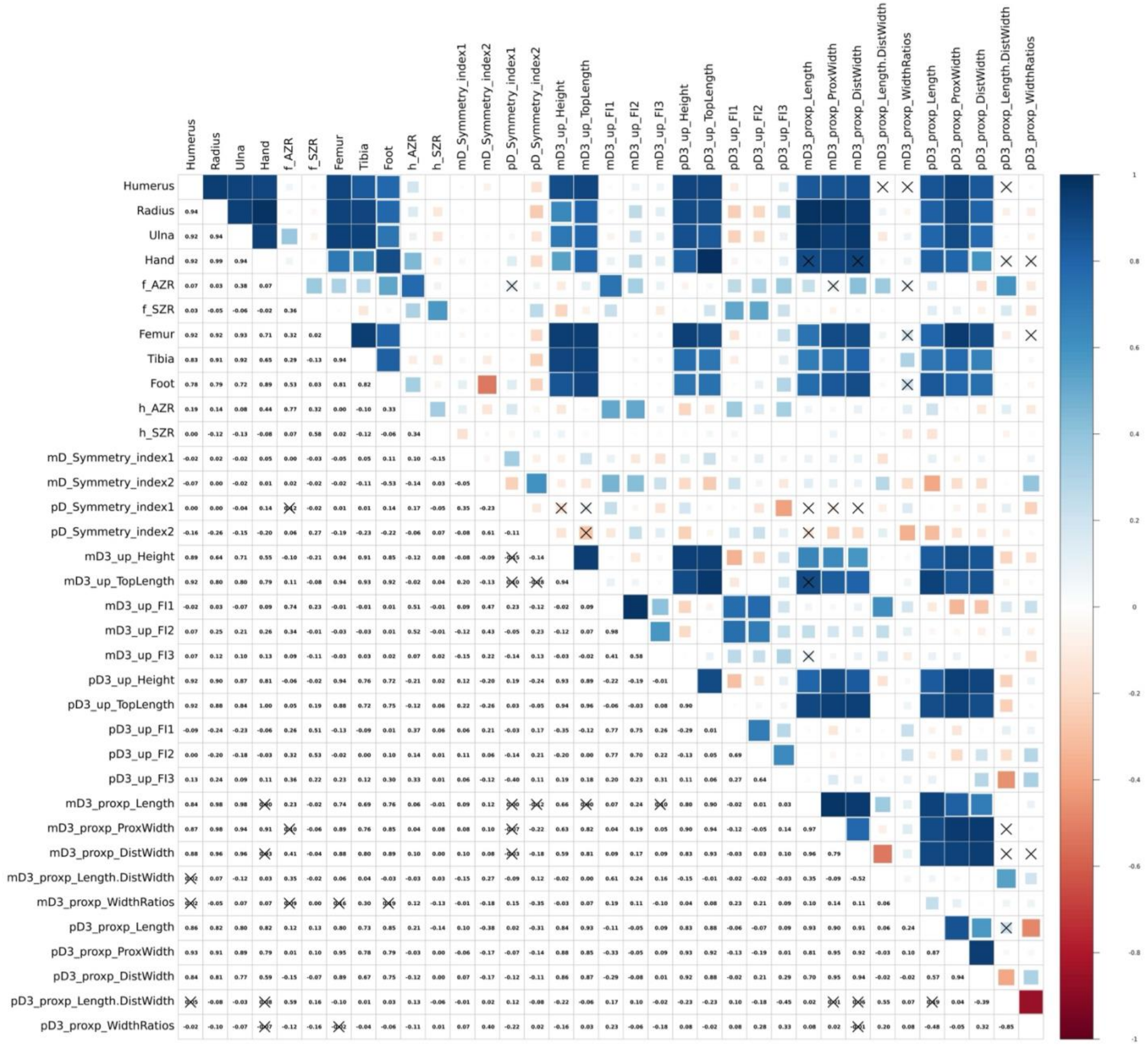

**Figure S27. Phylogenetic correlation matrix for all quantitative variables in our dataset under supertree 8.** Pearson correlation coefficients scale from 0 (dark red) to 1 (dark blue), and an "X" marks each non-significant correlation ($p > 0.05$). Abbreviations: AZR, acropod-zeugopodium ratio; AnAR, acropod-to-non-acropodium length ratio (acropodium length divided by the average of all stylopodial and zeugopodial lengths); Antmost, anterior-most; ASR, acropod-stylopodium ratio; BottomLength, length of the ungual about its ventral margin; DistWidth, width of the distal epiphysis; FI, flatness index; mD, manual digit; mD3, manual digit 3; pD, pedal digit; pD3, pedal digit 3; Postmost, posterior-most; proxp, proximal phalanx; ProxWidth, width of the proximal epiphysis; TopLength, length of the ungual about its dorsal margin; up, ungual phalanx; WidthRatios, ratio of the width of the distal epiphysis to the length of the proximal epiphysis. Note that a "." above replaces a "/", so "Length.DistWidth" should be read as the fraction "Length/DistWidth." All these metrics are defined and related to the main-text results within script S1. A higher-resolution version of this plot is available in the supplementary files associated with Data S3.

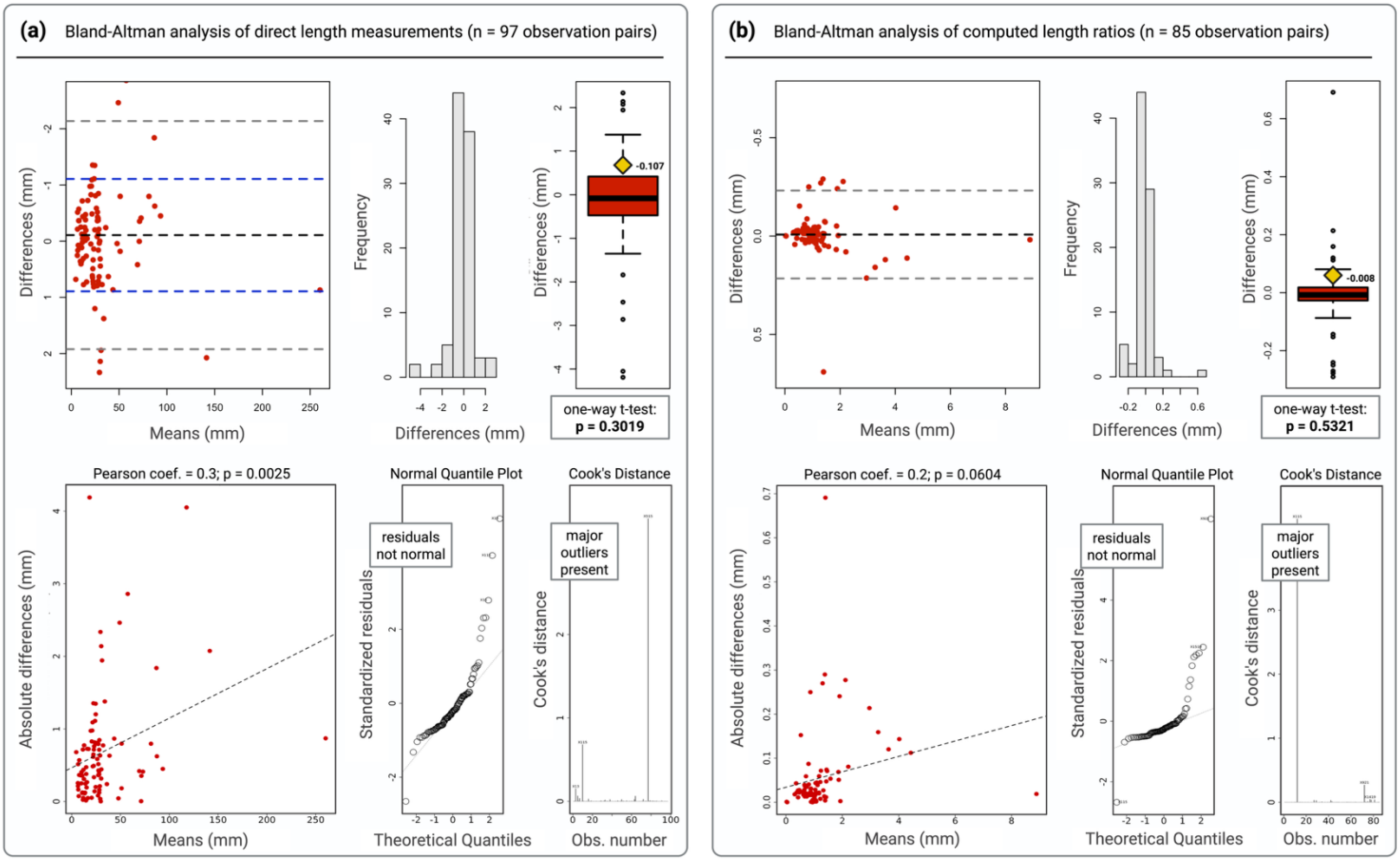


**Figure S28. Bland-Altman analyses assessing consistency between caliper and ImageJ measurements.** These analyses assess the relationship between the difference and mean of each pair of measurements collected by both methods for **(a)** direct length measurements and **(b)** computed limb measurement ratios capturing limb, acropodium, and phalanx proportions. Histograms and boxplots show the distribution of differences between caliper and ImageJ measurement pairs. One-sample t-tests show that differences are not significantly different from zero (each $p > 0.05$). Scatterplots and associated normal-quantile and Cook's distance plots for each Bland-Altman analysis show slight positive correlations (Pearson $\leq 0.3$) between absolute difference and mean absolute value. However, these correlations are either non-significant (b) or confounded by outliers and non-normally distributed residuals (a). Abbreviations: coef., coefficient; mm, millimeters; n, sample size; normal, normally distributed; Obs., observation.

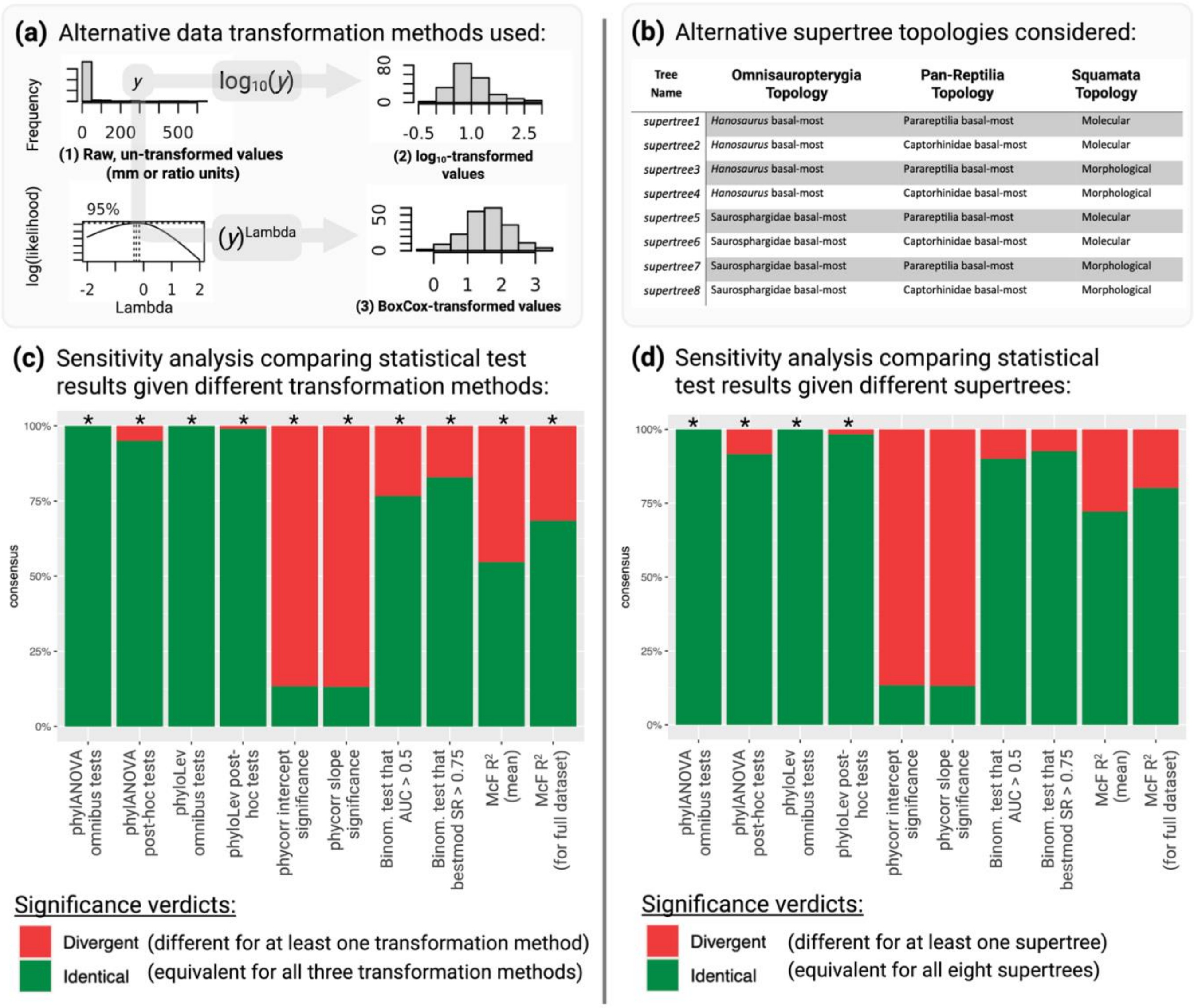


**Figure S29. Sensitivity analysis investigating the effects of data transformation method and tree topology on all phylogenetic comparative tests. (a)** The three data transformation methods applied. **(b)** The eight different combinations of phylogenetic hypotheses considered (corresponding full trees given in Data S1). **(c)** Sensitivity analysis results for phylogenetic comparative tests repeated following different data transformation methods. **(d)** Sensitivity analysis results for phylogenetic comparative tests repeated using different tree topologies. In (c–d), a test verdict is considered consistent (green) if and only if it is equivalent across either all eight supertrees (c) or all three transformation methods (d), and inconsistent (red) if it differs at least once among any of them. An asterisk indicates that the frequency of divergent test results is significantly greater than would be expected by chance alone (one-tailed binomial exact test, $p < 0.05$). Abbreviations: AUC, area under the curve; bestmod; best model (using probability classification threshold at which Youden's J-value was maximized); Binom., binomial; McF $R^2$, McFadden's Pseudo-$R^2$; mm, millimeters; phycorr, phylogenetic correlation test; phylANOVA, phylogenetic ANOVA; phyloLev, phylogenetic Levene's test; SR, success rate; *y*, arbitrary sample value of morphometric feature used to illustrate data transformation methods employed in sensitivity analysis. See Text S8 for additional details.

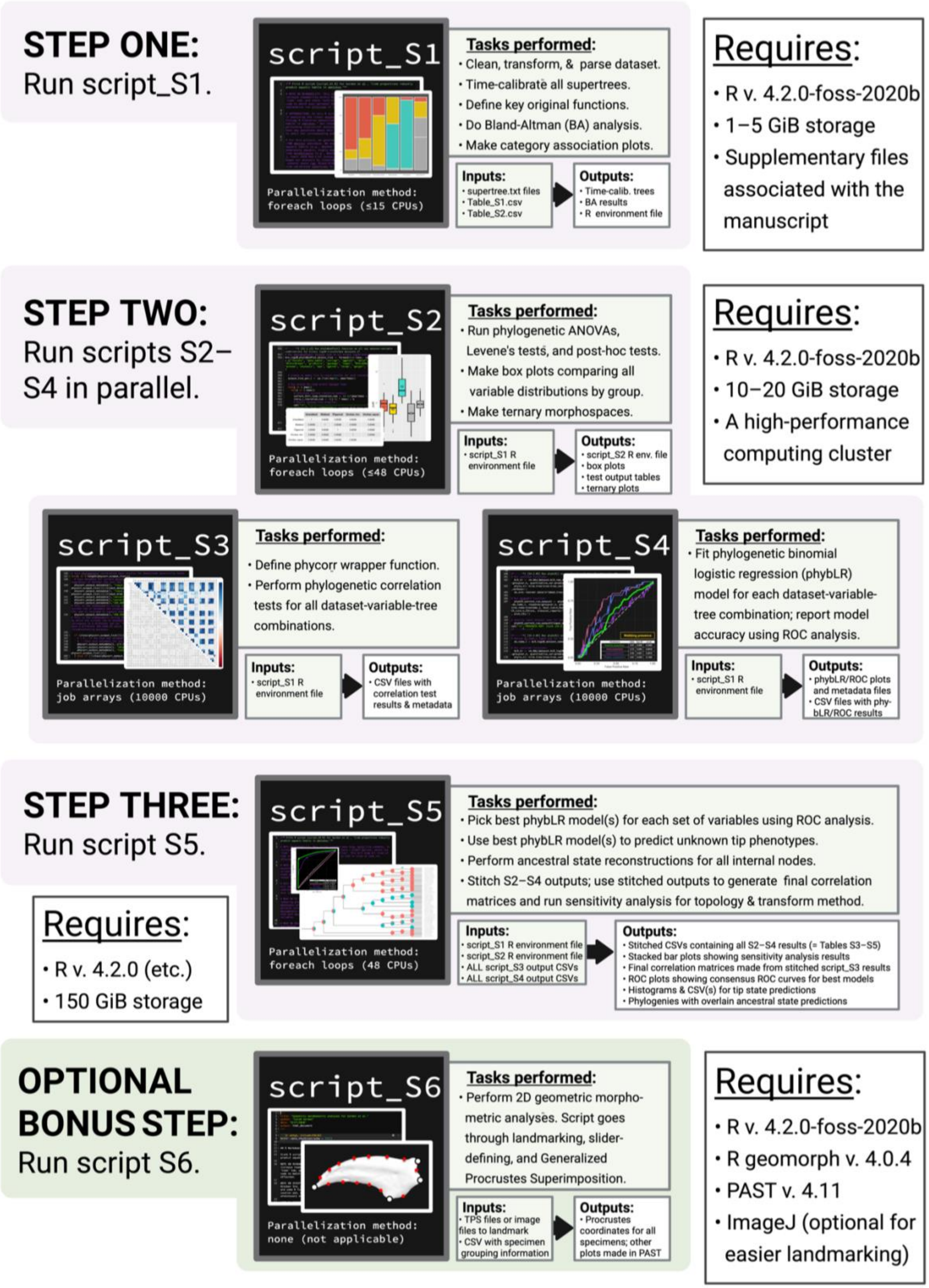


**Figure S30. Data analysis workflow.** This schematic describes the purpose, inputs, outputs, and requirements of each supplementary script (scripts S1–S6) to help anyone interested in repeating this study.

### Supplementary References


- Agassiz, L. *Recherches sur les Poissons Fossiles, Tome 1: Contenant l'introduction et toutes les questions générales, anatomiques, zoologiques et géologiques.* Imprimerie de Petitpierre: Neuchatel. 188 p. (1833).

- Ameghino, F. *Revista Argentina de Historia Natural, Tomo I.* Jacobo Peuser: Buenos Aires. 456 pp (1891).

- Amiot, R., Buffetaut, E., Lécuyer, C., Wang, X., Aoudad, L., Ding, Z., Fouler, F., Hutt, S., Martinaeu, F., Medeiros, M. A., Mo, J., Simon, L., Suteethorn, S., Tong, H., Zhang, F. & Zhou, Z. Oxygen isotope evidence for semi-aquatic habits among Spinosauridae theropods. *Geology* **38**, 139–142 (2010). https://dx.doi.org/10.1130/g30402.1.

- Archibald, J. D. *Ungulata* C. Linnaeus 1766 [J. D. Archibald], converted clade name. Registration Number 292. In: De Queiroz, K., Cantino, P. D. & Gauthier, J. A. (eds.) *Phylonyms: A Companion to the PhyloCode.* pp. 943–947 (2020).

- Arden, T. M. S., Klein, C. G., Zouhri, S., Longrich, N. R. Aquatic adaptation in the skull of carnivorous dinosaurs (Theropoda: Spinosauridae) and the evolution of aquatic habits in spinosaurids. *Cretaceous Research* **93**, 275–284 (2019). https://doi.org/10.1016/j.cretres.2018.06.013.

- Asher, R. J., Bennett, N. & Lehmann, T. The new framework for understanding placental mammal evolution. *BioEssays* **31**, 853–864 (2009). https://doi.org/10.1002/bies.200900053.

- Augusta, B. G., Zaher, H., Polcyn, M. J., Fiorillo, A. R. & Jacobs, L. L. A Review of Non-Mosasaurid (Dolichosaur and Aigialosaur) Mosasaurians and Their Relationships to Snakes. Chapter 7, in: Gower, D. J. & Zaher, H. (eds.) *The Origin and Early Evolutionary History of Snakes.* Cambridge University Press: Cambridge. pp. 157–179. (2022).

- Bakker, R. T. Juvenile-adult habitat shift in Permian fossil reptiles and amphibians. *Science* **217**, 53–55 (1982). https://doi.org/10.1126/science.217.4554.53.

- Bastiaans, D. Thalattosauria in time and space: a review of thalattosaur spatiotemporal occurrences, presumed evolutionary relationships and current ecological hypotheses. *Swiss Journal of Palaeontology* **143**, 36 (2024). https://doi.org/10.1186/s13358-024-00333-6.

- Batsch, A. J. G. C. *Versuch Einer Anleitung, zur Kenntniß und Geschichte der Tiere und Mineralien*. Akademische Buchhandlung: Jena. 528 pp. excluding plates (1788).

- Baur, G. *Palaeohatteria* Credner, and the Proganosauria. *American Journal of Science* **27**, 310–313 (1889). https://doi.org/10.2475/ajs.s3-37.220.310.

- Bejder, L. & Hall, B. K. Limbs in whales and limblessness in other vertebrates: mechanisms of evolutionary and developmental transformation and loss. *Evolution & Development* **4**, 445–458 (2002). https://doi.org/10.1046/j.1525-142x.2002.02033.x.

- Bell, G. L. & Polcyn, M. J. *Dallasaurus turneri*, a new primitive mosasauroid from the Middle Turonian of Texas and comments on the phylogeny of Mosasauridae (Squamata). *Netherlands Journal of Geosciences* **84**, 177–194 (2005). https://doi.org/10.1017/S0016774600020965.

- Bell, P. R. & Hendrickx, C. Crocodile-like sensory scales in a Late Jurassic theropod dinosaur. *Current Biology* **30**, R1063–R1070 (2020). https://doi.org/10.1016/j.cub.2020.08.066.

- Bell, P. R. & Hendrickx, C. Epidermal complexity in the theropod dinosaur *Juravenator* from the Upper Jurassic of Germany. *Palaeontology* **64**, 203–223 (2021). https://doi.org/10.1111/pala.12517.

- Benson, R. B. J. & Butler, R. J. Uncovering the diversification history of marine tetrapods: ecology influences the effect of geological sampling biases. In: McGowan, A. J. & Smith, A. B. (eds.) Comparing the Geological and Fossil Records: Implications for Biodiversity Studies. *Geological Society, London, Special Publications* **358**, 191–208 (2011). https://doi.org/10.1144/SP358.13.

- Benson, R. B. J., Evans, M. & Druckenmiller, P. S. High diversity, low disparity and small body size in plesiosaurs (Reptilia, Sauropterygia) from the Triassic–Jurassic boundary. *PLoS ONE* **7**, e31838 (2012). https://doi.org/10.1371/journal.pone.0031838.

- Benton, M. J. & Clark, J. M. Archosaur phylogeny and the relationships of the Crocodylia. In: Benton MJ, ed. The Phylogeny and Classification of the Tetrapods, Vol. 1: Amphibians, Reptiles, Birds. Oxford: Clarendon Press, 295–338 (1988).

- Benton, M. J., Donoghue, P. C. J., Asher, R. J., Friedman, M., Near, T. J. & Vinther, J. Constraints on the timescale of animal evolutionary history. *Palaeontologia Electronica* **18.1.1FC**, 1-106 (2015). https://palaeo-electronica.org/content/pdfs/424.pdf.

- Berkenhout, J. *Outlines of the Natural History of Great Britain and Ireland*, *Volume 1: Comprehending the Animal Kingdom.* P. Elmsly: London. 230 pp. (1769).

- Bever, G. S. & Norell, M. A. A new rhynchocephalian (Reptilia: Lepidosauria) from the Late Jurassic of Solnhofen (Germany) and the origin of the marine Pleurosauridae. *Royal Society Open Science* **4**, 170570 (2017). http://dx.doi.org/10.1098/rsos.170570.

- Bever, G. S., Lyson, T. R., Field, D. J. & Bhullar, B.-A. S. Evolutionary origin of the turtle skull. *Nature* **525**, 239–242 (2015). https://doi.org/10.1038/nature14900.

- Blob, R. W., Mayerl, C. J., Rivera, A. R. V., Rivera, G. & Young, V. K. H. ‘‘On the fence’’ versus ‘‘all in’’: Insights from turtles for the evolution of aquatic locomotor specializations and habitat transitions in tetrapod vertebrates. *Integrative and Comparative Biology* **56**, 1310–1322 (2016). https://doi.org/10.1093/icb/icw121.

- Bonaparte, J. F. Los tetrápodos triásicos de Argentina. *1º International Symposium of Gondwana* (Mar del Plata, 1967): 307-325 (1969).

- Bowdich, T. E. *An Analysis of the Natural Classifications of Mammalian, for the Use of Students and Travelers*. J. Smith: Paris. 115 pp. excluding plates (1821).

- Box, G. E. P. & Cox, D. R. An analysis of transformations. *Journal of the Royal Statistical Society, Series B* **26**, 211–252 (1964). http://links.jstor.org/sici?sici=0035-9246%281964%2926%3A2%3C211%3AAAOT%3E2.0.CO%3B2-6.

- Box, G. E. P. & Cox, D. R. An analysis of transformations revisited, rebutted. *Journal of the American Statistical Association* **77**, 209–210 (1982). http://dx.doi.org/10.1080/01621459.1982.10477788.

- Brisson, M. *Le Regne Animal divisé en IV Classes Ou Méthode Contenant la Division Generale des Animaux en IX Classes, & la Division Particulière des deux Premières Classes, Sçavoir de Celle des Quadrupedes & de Celle des Cetacées, en Ordres, Serions, Genres & Espéces*. Jean-Baptiste Bauche: Paris. 375 pp. (1756).

- Brochu, C. A. Phylogenetics, taxonomy, and historical biogeography of Alligatoroidea. *Society of Vertebrate Paleontology Memoirs* **6**, 9–100 (1999). https://doi.org/10.1080/02724634.1999.10011201.

- Brochu, C. A. Crocodylian snouts in space and time: Phylogenetic approaches toward adaptive radiation. *American Zoologist* **41**, 564–585 (2001). https://doi.org/10.1093/icb/41.3.564.

- Brownstein, C. D. *Halszkaraptor escuilliei* and the evolution of the paravian *bauplan*. *Scientific Reports* **9**, 16455 (2019). https://doi.org/10.1038/s41598-019-52867-2.

- Brusatte, S. L. *Spinosaurus*. *Current Biology* **31**, R1369–1371 (2021). https://doi.org/10.1016/j.cub.2021.09.002.

- Buffa, V., Jalil, N.-E., Falconnet, J. & Vincent, P. The Neodiapsida *Thadeosaurus colcanapi* from the upper Permian of Madagascar. *Papers in Palaeontology* **11**, e70008 (2025). https://doi.org/10.1002/spp2.70008.

- Burbrink, F. T., Grazziotin, F. G., Pyron, R. A., Cundall, D., Donnellan, S., Irish, F., Keogh, S., Kraus, F., Murphy, R. W., Noonan, B., Raxworthy, C. J., Ruane, S., Lemmon, A. L., Lemmon, E. M. & Zaher, H. 2020. Interrogating genomic-scale data for Squamata (lizards, snakes, and amphisbaenians) shows no support for key traditional morphological relationships. *Systematic Biology* **69**, 502–520 (2020). https://doi.org/10.1093/sysbio/syz062.

- Caldwell, M. W. From fins to limbs to fins: Limb evolution in fossil marine reptiles. *American Journal of Medical Genetics* **112**, 236–249 (2002). https://doi.org/10.1002/ajmg.10773.

- Caldwell, M. W. & Lee, M. S. Y. Live birth in Cretaceous marine lizards (mosasauroids). *Proceedings of the Royal Society of London B* **268**, 2397–2401 (2001). https://doi.org/10.1098/rspb.2001.1796.

- Calzavara, M., Muscio, G. & Wild, R. *Megalancosaurus preonensis* n. g., n. sp., a new reptile from the Norian of Friuli, Italy. Gortania - *Atti del Museo Friulano di Storia Naturale* **2**, 49-64 (1981).

- Camp, C. L. Classification of the lizards. *Bulletin of the American Museum of Natural History* **48**, 289–480 (1923).

- Campione, N. E. & Evans, D. C. A universal scaling relationship between body mass and proximal limb bone dimensions in quadrupedal terrestrial tetrapods. *BMC Biology* **10**, 60 (2012). https://doi.org/10.1186/1741-7007-10-60.

- Cantino, P. D. & de Queiroz, K. *International Code of Phylogenetic Nomenclature (PhyloCode)*, Version 6. CRC Press: Boca Raton. 149 pp (2020).

- Carroll, R. L. Plesiosaur ancestors from the Upper Permian of Madagascar. *Philosophical Transactions of the Royal Society of London B* **293**, 315–383 (1981). https://doi.org/10.1098/rstb.1981.0079.

- Carroll, R. L. & Dong, Z.-M. *Hupehsuchus,* an enigmatic aquatic reptile from the Triassic of China, and the problem of establishing relationships. *Philosophical Transactions of the Royal Society of London B* **331**, 131–153 (1991). https://doi.org/10.1098/rstb.1991.0004.

- Carroll, R. L. & Gaskill, P. The nothosaur *Pachypleurosaurus* and the origin of plesiosaurs. *Philosophical Transactions of the Royal Society of London B* **309**, 343–393 (1985). https://doi.org/10.1098/rstb.1985.0091.

- Case, E. C. *A revision of the Cotylosauria of North America.* Carnegie Institution of Washington: Washington, D. C. 121 pp. excluding plates (1911).

- Case, E. C. A cotylosaur from the Upper Triassic of western Texas. *Journal of the Washington Academy of Sciences* **18**, 177–178 (1928). https://www.jstor.org/stable/24522527.

- Case, E. C. A nothosaur from the Triassic of Wyoming. *University of Michigan Contributions from the Museum of Paleontology* **5**, 1–36 (1936).

- Cau, A. The body plan of *Halszkaraptor escuilliei* (Dinosauria, Theropoda) is not a transitional form along the evolution of dromaeosaurid hypercarnivory. *PeerJ* **8**, e8672 (2020). https://doi.org/10.7717/peerj.8672.

- Cau, A., Beyrand, V., Voeten, D. F. A. E., Fernandez, V., Tafforeau, P., Stein, K., Barsbold, R., Tsogtbaatar, K., Currie, P. J. & Godefroit, P. Synchrotron scanning reveals amphibious ecomorphology in a new clade of bird-like dinosaurs. *Nature* **552**, 395–399 (2017). https://doi.org/10.1038/nature24679.

- Chai, J., Lu, H., Jiang, D.-Y., Motani, R., Druckenmiller, P. S., Tintori, A. & Kelley, N. P. Reidentification of *Wayaosaurus bellus* and the conservative trunk and tail shape of Thalattosauria. *Historical Biology* **36**, 2537–2556 (2023). https://doi.org/10.1080/08912963.2023.2264886.

- Cheng, Y.-N. Sato, T., Wu, X.-C. & Li, C. First complete pistosauroid from the Triassic of China. *Journal of Vertebrate Paleontology* **26**, 501–504 (2006). https://www.jstor.org/stable/4524593.

- Cheng, Y.-N., Wu, X.-C. & Sato, T. A new thalattosaurian (Reptilia: Diapsida) from the Upper Triassic of Guizhou, China. *Vertebrata PalAsiatica* **45**, 246–260 (2007).

- Chiappe, L. M. & Göhlich, U. B. Anatomy of *Juravenator starki* (Theropoda: Coelurosauria) from the Late Jurassic of Germany. *Neues Jahrbuch fur Geologie und Palaontologie* **258**, 257–296 (2010).

- Choudhary, S., Choudhary, B. C. & Gopi, G. V. Spatio-temporal partitioning between two sympatric crocodilians (Gavialis gangeticus & Crocodylus palustris) in Katarniaghat Wildlife Sanctuary, India. *Aquatic Conservation: Marine and Freshwater Ecosystems* **28**, 1067–1076 (2018).

- Clementz, M. T., Holroyd, P. A. & Koch, P. L. Identifying aquatic habits of herbivorous mammals through stable isotope analysis. *PALAIOS* **23**, 574–585 (2008). http://dx.doi.org/10.2110/palo.2007.p07-054r.

- Clopper, C. J. & Pearson, E. S. The use of confidence or fiducial limits illustrated in the case of the binomial. *Biometrika* **26**, 404–413 (1934). https://doi.org/10.2307/2331986.

- Cocude-Michel, M. Les rhynchocéphales et les sauriens des calcaires lithographiques (Jurassique supérieur) d'Europe occidentale. *Nouvelles archives du Muséum d'histoire naturelle de Lyon* **7**, 3–224 (1963).

- Conybeare, W. D. XXI.—On the discovery of an almost perfect skeleton of the *Plesiosaurus*. *Transactions of the Geological Society of London* **6**, 381–389 (1824).

- Coombs, W. P. Theoretical aspects of cursorial adaptations in dinosaurs. *The Quarterly Review of Biology* **53**, 393–418 (1978). https://www.jstor.org/stable/2826581.

- Cooper, L. N. “Forelimb Anatomy.” In: Wursig, B., et al. (eds.). *Encyclopedia of Marine Mammals*. Elsevier Science & Technology, pp. 449–452 (2008). Accessed via ProQuest Ebook Central, http://ebookcentral.proquest.com/lib/yale-ebooks/detail.action?docID=405935.

- Cooper, L. N., Sears, K. E., Armfield, B. A., Kala, B., Hubler, M. & Thewissen, J. G. M. Review and experimental evaluation of the embryonic development and evolutionary history of flipper development and hyperphalangy in dolphins (Cetacea: Mammalia). *genesis* **56**, e23076 (2018). https://doi.org/10.1002/dvg.23076.

- Cope, E. D. “*Osteolaemus tetraspis*.” In: *Proceedings of the Academy of Natural Science of Philadelphia* **12**, 549–550 (1860).

- Cope, E. D. Remarks on *Holops brevispinus*, *Ornithotarsus immanis* and *Macrosaurus proriger*. *Proceedings of the Academy of Natural Sciences of Philadelphia* **11**, 1–123 (1869a).

- Cope, E. D. Synopsis of the extinct Batrachia and Reptilia of North America, Part I. *Transactions of the American Philosophical Society, New Series* **14**, 1–235 (1869b).

- Cope, E. D. *Systematic Catalogue of Vertebrata of the Eocene of New Mexico: Collected in 1874*. US Government Printing Office: Washington. 37 pp. (1875).

- Cope, E. D. Third contribution to the history of the Vertebrata of the Permian formation of Texas. *Proceedings of the American Philosophical Society* **20**, 447–461 (1882). https://www.jstor.org/stable/982692.

- Cope, E. D. A contribution to the vertebrate paleontology of Brazil. *Proceedings of the American Philosophical Society* **25**, 7–15 (1886). https://www.jstor.org/stable/982910.

- Cope, E. D. The *Edentata* of North America. *American Naturalist* **23**, 657–664 (1889). https://www.jstor.org/stable/2451326.

- Cope, E. D. The Reptilian Order Cotylosauria. *Proceedings of the American Philosophical Society* **34**, 436-457 (1895). https://www.jstor.org/stable/982900.

- Cope, E. D. Second contribution to the history of Cotylosauria. *Proceedings of the American Philosophical Society* **35**, 122–139 (1896). https://www.jstor.org/stable/983258.

- Cornalia, E. Notizie zoologiche sul *Pachypleura edwardsii* Cor. Nuovo sauro acrodonte degli strati Triasici di Lombardia. *Giornale dell'istituto Lombardo di Scienze* **6**, 1–46 (1854).

- Crocodile Specialist Group. Osteolaemus tetraspis. *The IUCN Red List of Threatened Species* 1996, e.T15635A4931429 (1996). http://dx.doi.org/10.2305/IUCN.UK.1996.RLTS.T15635A4931429.en.

- Curioni, G. Cenni sopra un nuovo saurio fossile dei monti di Perledo sul Lario e sul terreno che lo racchiude. *Giornale dell' I. R. Istituto Lombardo di Scienze, Lettre ed Arti* **16**, 159–170 (1847).

- Currie, P. J. *Hovasaurus boulei*, an aquatic eosuchian from the Upper Permian of Madagascar. *Palaeontologica Africana* **24**, 99–168 (1981). https://wiredspace.wits.ac.za/bitstreams/dfdaeb86-87d3-4099-88af-57c49882335f/download.

- Cuvier, G. Sur les espèces d'animaux dont proviennent les os fossiles répandus dans la pierre à plâtre des environs de Paris. *Annales du Muséum National d'Histoire Naturelle, Paris* **3**, 275–303 (1804).

- Cuvier, G. Sur les diffèrentes espèces de crocodiles vivans et sur leurs caractères distinctifs. *Annales du Muséum d'Histoire Naturelle* **10**, 8–66 (1807).

- Dal Sasso, C. & Pinna, G. *Aphanizocnemus libanesi* n. Gen. g. Sp., a new dolichosaur (Reptilia, Varanoidea) from the Upper Cretaceous of Lebanon. *Paleontologia Lombarda*: *Sezione di Paleontologia del Museo Civico di Storia Naturale di Milano* **7**, 3–31 (1997).

- Daudin, F. M. *Histoire Naturelle, Générale et Particulière des Reptiles*, Tome Second (Vol. 2). F. Dufart: Paris. 432 pp. (1802).

- Davis, A. P., Witte, D. P., Hsieh-Li, H. M., Potter, S. S. & Capecchi, M. R. Absence of radius and ulna in mice lacking *hoxa-11* and *hoxd-11*. *Nature* **375**, 791–795 (1995).

- Dawson, W. R., Bartholomew, G. A. & Bennett, A. F. A reappraisal of the aquatic specializations of the Galapagos Marine Iguana (*Amblyrhynchus cristatus*). *Evolution* **31**, 891–897 (1977).

- De Araújo Sena, M. V. & Cubo, J. Inferring the lifestyles of extinct Crocodyliformes using osteoderm ornamentation. *The Science of Nature* **110**, 41 (2023). https://doi.org/10.1007/s00114-023-01871-8.

- De Blainville, H. D. Description de quelques espèces de reptiles de la Californie, précédée de l'analyse d'un système general d'Erpetologie et d'Amphibiologie. — *Nouvelles Annales du Muséum (National) d'History Naturaelle, Paris* **4**, 233–296 (1835).

- De Braga, M. & Reisz, R. R. A new diapsid reptile from the uppermost Carboniferous (Stephanian) of Kansas. *Palaeontology* **38**, 199–212 (1995).

- De Braga, M. & Rieppel, O. Reptile phylogeny and the interrelationships of turtles. *Zoological Journal of the Linnean Society* **120**, 281–3554 (1997). https://doi.org/10.1111/j.1096-3642.1997.tb01280.x.

- De la Beche, H. T. & Conybeare, W. D. XXX.—Notice on the discovery of a new Fossil Animal, forming a link between the Ichthyosaurus and Crocodile, together with general remarks on the Osteology of the Ichthyosaurus. *Transactions of the Geological Society* **5**, 559–594 (1821).

- De La Cépède, B. G. E. *Histoire Naturelle Des Cétacées.* Plassan, Imprimeur-Libraire: Paris. 329 pp. (1804).

- De Queiroz, K. & Gauthier, J. A. *Lepidosauria* E. Haeckel 1866 [K. De Queiroz and J. A. Gauthier], converted clade name. Registration Number 61. In: De Queiroz, K., Cantino, P. D. & Gauthier, J. A. (eds.) *Phylonyms: A Companion to the PhyloCode.* CRC Press, Taylor & Francis: Boca Raton. pp. 1079–1085 (2020a).

- De Queiroz, K. & Gauthier, J. A. *Squamata* M. Oppel 1811 [K. de Queiroz and J. A. Gauthier], converted clade name. Registration Number 101. In: de Queiroz, K., Cantino, P. D. & Gauthier, J. A. (eds.) *Phylonyms: A Companion to the PhyloCode.* CRC Press, Taylor & Francis: Boca Raton. pp. 1093–1101 (2020b).

- De Queiroz, K., Cantino, P. D. & Gauthier, J. A. (eds.) *Phylonyms: A Companion to the PhyloCode.* CRC Press, Taylor & Francis: Boca Raton. 1324 pp. (2020).

- DeLong, E. R., DeLong,  D. M. & Clarke-Pearson, D. L. Comparing the areas under two or more correlated Receiver Operating Characteristic curves: A nonparametric approach. *Biometrics* **44**, 837–845 (1988). https://doi.org/10.2307/2531595.

- Demarco, P. N., Meneghal, M., Laurin, M. & Piñeiro, G. Was *Mesosaurus* a fully aquatic reptile? *Frontiers in Ecology & Evolution* **6**, 109 (2018). https://doi.org/10.3389/fevo.2018.00109.

- Deméré, T. A. *Cetacea* M. J. Brisson 1762 [T. A. Deméré], converted clade name. Registration Number 261. In: De Queiroz, K., Cantino, P. D. & Gauthier, J. A. (eds.) *Phylonyms: A Companion to the PhyloCode.* CRC Press, Taylor & Francis: Boca Raton. pp. 960–962 (2020).

- Dietz, R. S. & Holden, J. C. Reconstruction of Pangaea: Breakup and dispersion of continents, Permian to present. *Journal of Geophysical Research* **75**, 4939–4956 (1970). https://doi.org/10.1029/JB075i026p04939.

- Druckenmiller, P. S., Kelley, N. P., Metz, E. T. & Baichtal, J. An articulated Late Triassic (Norian) thalattosauroid from Alaska and ecomorphology and extinction of Thalattosauria. *Scientific Reports* **10**, 1746 (2020). https://doi.org/10.1038/s41598-020-57939-2.

- Dupret, V. The pleurosaurs: Anatomy and phylogeny. *Revue de Paleobiologie* **9**, 61–80 (2004). https://www.researchgate.net/publication/243971261_The_pleurosaurs_Anatomy_and_phylogeny.

- Dutchak, A. R. & Caldwell, M. W. Redescription of *Aigialosaurus dalmaticus* Kramberger, 1892, a Cenomainian Mosasauroidea lizard from Hvar Island, Croatia. *Canadian Journal of Earth Sciences* **43**, 1821–1834 (2006). https://doi.org/10.1139/e06-086.

- Dutchak, A. R. & Caldwell, M. W. A redescription of *Aigialosaurus* (= *Opetiosaurus*) *bucchichi* (Kornhuber, 1901) (Squamata: Aigialosauridae) with comments on mosasauroid systematics. *Journal of Vertebrate Paleontology* **29**, 437–452 (2009). https://doi.org/10.1671/039.029.0206.

- English, A. W. M. Structural correlates of forelimb function in fur seals and sea lions. *Journal of Morphology* **151**, 325–352 (1977). https://doi.org/10.1002/jmor.1051510303.

- Eriksson, M. E., De La Garza, R., Horn, E. & Lindgren, J. A review of ichthyosaur (Reptilia, Ichthyopterygia) soft tissues with implications for life reconstructions. *Earth-Science Reviews* **226**, 103965 (2022). https://doi.org/10.1016/j.earscirev.2022.103965.

- Evans, S. E. The Upper Permian reptile *Adelosaurus* from Durham. *Palaeontology* **31**, 957–964 (1988).

- Evans, S. E. & Borsuk-Białynicka, M. A small lepidosauromorph reptile from the Early Triassic of Poland. *Paleontologica Polonica* **65**, 179–202 (2009).

- Fabbri, M., Navalón, G., Benson, R. B. J., Pol, D., O'Connor, J., Bhullar, B.-A. S., Erickson, G. M., Norell, M. A., Orkney, A., Lamanna, M. C., Zouhri, S., Becker, J., Emke, A., Dal Sasso, C., Bindellini, G., Maganuco, S., Auditore, M. & Ibrahim, N. Subaqueous foraging among carnivorous dinosaurs. *Nature* **603**, 852–857 (2022). https://doi.org/10.1038/s41586-022-04528-0.

- Falkingham, P. L., Margetts, L., Smith, I. M. & Manning, P. L. Reinterpretation of palmate and semi-palmate (webbed) fossil tracks; insights from finite element modelling. *Palaeogeography, Palaeoclimatology, Palaeoecology* **271**, 69–76 (2009). https://doi.org/10.1016/j.palaeo.2008.09.011.

- Fawcett, T. An introduction to ROC analysis. *Pattern Recognition Letters* 27, 861–874 (2006). http://dx.doi.org/10.1016/j.patrec.2005.10.010.

- Fedak, T. J. & Hall, B. K. Perspectives on hyperphalangy: patterns and processes. *Journal of Anatomy* **204**, 151–163 (2004). https://doi.org/10.1111/j.0021-8782.2004.00278.x.

- Felsenstein, J. Phylogenies and the comparative method. *The American Naturalist* **125**, 1–15 (1985). https://doi.org/10.1086/284325.

- Ferguson, M. W. J. Reproductive Biology and Embryology of the Crocodilians. In: Gans, C., Billett, F., Maderson, P. F. A. (Eds.) *Development A. Biology of the Reptilia*. New York: John Wiley & Sons: 331–491 (1985).

- Fernández, M. S., Vlachos, E., Buono, M. R., Alzugaray, L., Campos, L., Sterli, J., Herrera, Y. & Paolucci, F. Fingers zipped up or baby mittens? Two main tetrapod strategies to return to the sea. *Biology Letters* **16**, 20200281 (2020). http://dx.doi.org/10.1098/rsbl.2020.0281.

- Fielding, S., Martill, D. M. & Naish, D. Solnhofen-style soft-tissue preservation in a new species of turtle from the Crato Formation (Early Cretaceous, Aptian) of north-east Brazil. *Palaeontology* **48**, 1301–1310 (2005). https://doi.org/10.1111/j.1475-4983.2005.00508.x.

- Fischer, V., Bennion, R. F., Foffa, D., MacLaren, J. A., McCurry, M. R., Melstrom, J. M. & Bardet, N. Ecological signal in the size and shape of marine amniote teeth. *Proceedings of the Royal Society of London B* **289**, 20221214 (2022). https://doi.org/10.1098/rspb.2022.1214.

- Fish, F. E., Howle, L. E. & Murray, M. M. Hydrodynamic flow control in marine mammals. *Integrative and Comparative Biology* **48**, 788–800 (2008). https://doi.org/10.1093/icb/icn029.

- Flynn, J. J., Wyss, A. A. & Wolsan, M. *Carnivora* T. Bowdich 1821 [J. J Flynn, A. R. Wyss, and M. Wolsan], converted clade name. Registration Number 268. In: De Queiroz, K., Cantino, P. D. & Gauthier, J. A. (eds.) *Phylonyms: A Companion to the PhyloCode.* CRC Press, Taylor & Francis: Boca Raton. pp. 983–987 (2020).

- Foffa, D., Young, M. T., Stubbs, T. L., Dexter, K. G. & Brusatte, S. L. The long-term ecology and evolution of marine reptiles in a Jurassic seaway. *Nature Ecology & Evolution* **2**, 1548–1555 (2018). https://doi.org/10.1038/s41559-018-0656-6.

- Ford, D. P. & Benson, R. B. J. The phylogeny of early amniotes and the affinities of Parareptilia and Varanopidae. *Nature Ecology & Evolution* **4**, 57–65 (2020). https://doi.org/10.1038/s41559-019-1047-3.

- Forskål, P. *Descriptiones Animalium Avium, Amphibiorum, Piscium, Insectorum, Vermium; Quae in Itinere Orientali Observavit*. Möllerus: Haunia (1775).

- Frey, E., Mulder, E. W. A., Stinnesbeck, W., Rivera-Sylva, H. E., Padilla-Gutiérrez, J. M., González-González, A. H. A new polycotylid plesiosaur with extensive soft-tissue preservation from the early Late Cretaceous of northeast Mexico. *Boletín de la Sociedad Geológica* Mexicana **69**, 87–134 (2017). http://dx.doi.org/10.18268/BSGM2017v69n1a5.

- Frýdlová, P., Mrzílková, J., Šeremeta, M., Křemen, J., Dudák, J., Žemlička, J., Minnich, B., Kverková, K., Němec, P., Zach, P. & Frynta, D. Determinate growth is predominant and likely ancestral in squamate reptiles. *Proceedings of the Royal Society of London B* **287**, 20202737 (2020). https://doi.org/10.1098/rspb.2020.2737.

- Fukuda, Y., Webb, G., Manolis, C., Lindner, G. & Banks, S. Translocation, genetic structure and homing ability confirm geographic barriers disrupt saltwater crocodile movement and dispersal. *PLoS ONE* **14**, e0205862 (2019). https://doi.org/10.1371/journal.pone.0205862.

- Garland, T., Dickerman, A. W., Janis, C. M. & Jones, J. A. Phylogenetic analysis of covariance by computer simulation. *Systematic Biology* **42**, 265–292 (1993). https://doi.org/10.1093/sysbio/42.3.265.

- Gauthier, J. A. A cladistic analysis of the higher systematic categories of the Diapsida. University Microfilms International, Ann Arbor, MI. #85-12825, vii + 564 pp. (1984).

- Gauthier, J. A. *Archosauromorpha* F. Von Huene 1946 [J. A. Gauthier], converted clade name. Registration Number 175. In: De Queiroz, K., Cantino, P. D. & Gauthier, J. A. (eds.) *Phylonyms: A Companion to the PhyloCode.* CRC Press, Taylor & Francis: Boca Raton. pp. 1179–1182 (2020a).

- Gauthier, J. A. *Pan-Archosauria* J. A. Gauthier, new clade name. Registration Number 176. In: De Queiroz, K., Cantino, P. D. & Gauthier, J. A. (eds.) *Phylonyms: A Companion to the PhyloCode.* CRC Press, Taylor & Francis: Boca Raton. pp. 1175–1177 (2020b).

- Gauthier, J. A. & De Queiroz, K. *Pan-Lepidosauria* J. A. Gauthier and K. De Queiroz, new clade name. Registration Number 118. In: de Queiroz, K., Cantino, P. D. & Gauthier, J. A. (eds.) *Phylonyms: A Companion to the PhyloCode.* CRC Press, Taylor & Francis: Boca Raton. pp. 1075–1078 (2020a).

- Gauthier, J. A. & De Queiroz, K. *Sauria* J. Macartney 1802 [J. A. Gauthier and K. de Queiroz], converted clade name. Registration Number 114. In: de Queiroz, K., Cantino, P. D. & Gauthier, J. A. (eds.) *Phylonyms: A Companion to the PhyloCode.* CRC Press, Taylor & Francis: Boca Raton. pp. 1065–1073 (2020b).

- Gauthier, J. A. & Padian, K. *Archosauria* E. D. Cope 1869 [J. A. Gauthier and K. Padian], converted clade name. Registration Number 173. In: De Queiroz, K., Cantino, P. D. & Gauthier, J. A. (eds.) *Phylonyms: A Companion to the PhyloCode.* CRC Press, Taylor & Francis: Boca Raton. pp. 1187–1193 (2020).

- Gauthier, J. A., Kluge, A. G. & Rowe, T. Amniote phylogeny and the importance of fossils. *Cladistics* **4**, 105–209 (1988a). https://doi.org/10.1111/j.1096-0031.1988.tb00514.x.

- Gauthier, J. A., Estes, R. & de Queiroz, K. A Phylogenetic Analysis of Lepidosauromorpha. In: Estes, R. & Pregill, G. (eds.) *Phylogenetic Relationships of the Lizard Families: Essays Commemorating Charles L. Camp.* Stanford University Press: Stanford. pp. 15–98 (1988b).

- Gauthier, J. A., Kearney, M., Maisano, J. A., Rieppel, O. & Behlke, A. D. B. Assembling the squamate tree of life: Perspectives from the phenotype and the fossil record. *Bulletin of the Peabody Museum of Natural History* **53**, 3–308 (2012). https://doi.org/10.3374/014.053.0101.

- Gervais, P. *Zoologie et Paléontologie Françaises (Animaux Vertébrés) ou Nouvelles Recherches sur les Animaux Vivants et Fossiles de la France*. Arthus Bertrand: Paris. 274 pp. (1852).

- Gervais, P. Description du *Mesosaurus tenuidens*, Reptile fossile de l'Afrique australe. *Académie des Sciences et Lettres de Montpellier. Mémoires de la Section des Sciences* **6**, 169–175 (1865).

- Gmelin, J. F. *Systema Naturae, per Regna Tria Natura: Secundum Classes, Ordines, Genera, Species, cum Characteribus, Differentiis, Synonymis, Locis*. Georg Emanuel Beer: Leipzig. pp. 501–1032 (1789).

- Göhlich, U. B. & Chiappe, L. M. A new carnivorous dinosaur from the Late Jurassic Solnhofen archipelago. *Nature* **440**, 329–332 (2006). https://doi.org/10.1038/nature04579.

- Gordon, C. M. *Reconstructing the Evolutionary Histories of Aquatic and Macrocarnivorous Lifestyles in Reptiles*. A Dissertation Presented to the Faculty of the Graduate School of Yale University in Candidacy for the Degree of Doctor of Philosophy. 1527 pp. (2025).

- Gordon, C. M., Freisem, L. S., Griffin, C. T., Gauthier, J. A. & Bhullar, B.-A. S. Supplementary scripts and data for "Limb proportions predict aquatic habits and soft-tissue flippers in extinct amniotes" [Dataset]. Dryad (2025). https://doi.org/10.5061/dryad.08kprr5fn.

- Gorjanović-Kramberger, D. *Aigialosaurus*, eine neue Eidechse aus den Kreideschiefern der Insel Lesina, mit Rücksicht auf die bereits beschriebenen Lacertiden von Comen und Lesina. Glasnik hrvatskoga naravoslovnoga drustva (Societas historico-naturalis croatica) u Zagrebu **7**, 74–106 (1892).

- Goswami, A. & Finarelli, J. A. EMMLi: A maximum likelihood approach to the analysis of modularity. *Evolution* **70**, 1622–1637 (2016). https://doi.org/10.1111/evo.12956.

- Grande, L. Paleontology of the Green River Formation, with a review of the fish fauna. The Geological *Survey of Wyoming Bulletin* **63**, xviii + 333 pp (1984).

- Gray, J. E. Descriptions of two hitherto unrecorded species of reptiles from New Zealand; presented to the British Museum by Dr. Diefenbach. *Zoological Miscellany* **72** (1842).

- Gray, J. E. Catalogue of the tortoises, crocodiles, and amphisbaenians, in the collection of the British Museum. London: Edward Newman, 80 (1844).

- Griebeler, E. M. & Klein, N. Life-history strategies indicate live-bearing in *Nothosaurus* (Sauropterygia). *Palaeontology* **62**, 697–713 (2019). https://doi.org/10.1111/pala.12425.

- Gutarra, S. & Rahman, I. A. The locomotion of extinct secondarily aquatic tetrapods. *Biological Reviews* **97**, 67–98 (2022). https://doi.org/10.1111/brv.12790.

- Haeckel, E. *Generelle Morphologie der Organismen*, *Band 2: Allgemeine Entwicklungsgeschichte der Organismen*. Georg Reimer: Berlin. 462 pp. excluding plates (1866).

- Hancock, A. & Howse, R. On *Protorosaurus speneri* von Meyer, and a new species, *Protorosaurus huxleyi*, from the Marl Slate of Middridge, Durham. *Quarterly Journal of the Geological Society of London* **26**, 565–572 (1870).

- Harris, J. M. & Carroll, R. L. *Kenyasaurus*, a new eosuchian reptile from the Early Triassic of Kenya. *Journal of Paleontology* **51**, 139–149 (1977). https://www.jstor.org/stable/1303471.

- Hatcher, J. B. A median horned rhinoceros from the Loup Fork Beds of Nebraska. *The American Geologist* **18**, 149 (1894).

- Haughton, S. H. On reptilian remains from the Karroo beds of East Africa. *Quarterly Journal of the Geological Society* **80**, 1–11 (1924). https://doi.org/10.1144/GSL.JGS.1924.080.01-04.03.

- Helm, C. Inferred evidence of webbing in a possible Cretaceous avian track from northeastern British Columbia. *British Columbia Birds* **33**, 14–19 (2022).

- Henderson, C. M., Shen, S. Z., Gradstein, F. M. & Agterberg, F. P. The Permian Period. In: *Geologic Time Scale 2020*, Volume 2. Elsevier B. V.: Amsterdam. pp. 875–902 (2020). https://doi.org/10.1016/B978-0-12-824360-2.00024-3.

- Henderson, D. M. A buoyancy, balance and stability challenge to the hypothesis of a semi-aquatic *Spinosaurus* Stromer, 1915 (Dinosauria: Theropoda). *PeerJ* **6**, e5409 (2018). https://doi.org/10.7717/peerj.5409.

- Ho, L. S. T. & Ané, C. A linear-time algorithm for Gaussian and Non-Gaussian trait evolution models. *Systematic Biology* **63**, 397–408 (2014). https://doi.org/10.1093/sysbio/syu005.

- Holmes, A. The South Atlantic: Land bridges or continental drift? *Nature* **171**, 669–671 (1953). https://doi.org/10.1038/171669a0.

- Hough, J. R. A systematic revision of *Daphoenus* and some allied genera. *Journal of Paleontology* **22**, 573–600 (1948). https://www.jstor.org/stable/1299593.

- Houssaye, A. "Pachyostosis" in aquatic amniotes: a review. *Integrative Zoology* **4**, 325–340 (2009). https://doi.org/10.1111/j.1749-4877.2009.00146.x.

- Horsnell, K., Ali, M., Malik, S., Wilson, L., Hall, C., Debeer, P. & Crow, Y. Clinical phenotype associated with homozygosity for a HOXD13 7-residue polyalanine tract expansion. *European Journal of Medical Genetics* **49**, 396–401 (2006). https://doi.org/10.1016/j.ejmg.2006.01.004.

- Huxley, T. H. *A Manual of the Anatomy of Vertebrated Animals*. J. A. Churchill: London. 431 pp. (1871).

- Ibrahim, N., Sereno, P. C., Dal Sasso, C., Maganuco, S., Fabbri, M., Martill, D. M., Zouhri, S., Myhrvold, N. & Iurino, D. A. Semiaquatic adaptations in a giant predatory dinosaur. *Science* **345**, 1613–1616 (2014). https://doi.org/10.1126/science.1258750.

- Ibrahim, N., Maganuco, S., Dal Sasso, C., Fabbri, M., Auditore, M., Bindellini, G., Martill, D. M., Zouhri, S., Mattarelli, D. A., Unwin, D. M., Wiemann, J., Bonadonna, D., Amane, A., Jakubczak, J., Joger, U., Lauder, G. V. & Pierce, S. E. Tail-propelled aquatic locomotion in a theropod dinosaur. *Nature* **581**, 67–70 (2020). https://doi.org/10.1038/s41586-020-2190-3.

- Isberg, S., Combrink, X., Lippai, C. & Balaguera-Reina, S. A. Crocodylus niloticus. *The IUCN Red List of Threatened Species* 2019: e.T45433088A3010181 (2019). http://dx.doi.org/10.2305/IUCN.UK.2019-1.RLTS.T45433088A3010181.en.

- Jackson, J. F., Campbell, H. W. & Campbell, K. E. The feeding habits of crocodilians: Validity of the evidence from stomach contents. *Journal of Herpetology* **8**, 378–381 (1974). https://doi.org/10.2307/1562912.

- Ji, Q., Luo, Z.-X., Yuan, C.-X. & Tabrum, A. R. A Swimming mammaliaform from the Middle Jurassic and ecomorphological diversification of early mammals. *Science* **311**, 1123–1127 (2006). https://doi.org/10.1126/science.1123026.

- Jiang, D.-Y., Motani, R., Tintori, A., Rieppel, O., Chen, G.-B., Huang, J.-D., Zhang, R., Sun, Z.-Y. & Ji, C. The Early Triassic eosauropterygian *Majiashanosaurus discocoracoidis*, gen. et sp. nov. (Reptilia, Sauropterygia), from Chaohu, Anhui Province, People's Republic of China. *Journal of Vertebrate Paleontology* **34**, 1044–1052 (2014). https://doi.org/10.1080/02724634.2014.846264.

- Jiang, D.-Y., Zhou, M., Motani, R., Tintori, A., Fraser, N. C., Huang, J.-d., Rieppel, O., Ji, C., Fun, W.-l., Sun, Z.-y. & Lu, H. Emergence and ecological transition of the Mesozoic marine reptiles: Evidence from the Early Triassic Chaohu and the Middle Triassic Xingyi Faunas. *Palaeogeography, Palaeoclimatology, Palaeoecology* **628**, 111750 (2023). https://doi.org/10.1016/j.palaeo.2023.111750.

- Jones, M. E. H. Skull shape and feeding strategy in *Sphenodon* and other Rhynchocephalia (Diapsida: Lepidosauria). *Journal of Morphology* **269**, 945–966 (2008). https://doi.org/10.1002/jmor.10634.

- Joyce, W. G. & Gauthier, J. A. Palaeoecology of Triassic stem turtles sheds new light on turtle origins. *Proceedings of the Royal Society of London B* **271**, 1–5 (2004). https://doi.org/10.1098/rspb.2003.2523.

- Joyce, W. G., Parham, J. F., Anquetin, J., Claude, J., Danilov, I. G., Iverson, J. B., Kear, B., Lyson, T. R., Rabi, M. & Sterli, J. *Testudines* A. J. G. C. Batsch 1788 [W. G. Joyce, J. F. Parham, J. Anquetin, J. Claude, I. G. Danilov, J. B. Iverson, B. Kear, T. R. Lyson, M. Rabi, and J. Sterli], converted clade name. Registration Number 274. In: de Queiroz, K., Cantino, P. D. & Gauthier, J. A. (eds.) *Phylonyms: A Companion to the PhyloCode.* CRC Press, Taylor & Francis: Boca Raton. pp. 1049–1051 (2020).

- Joyce, W. G., Mäuser, M. & Evers, S. W. Two turtles with soft tissue preservation from the platy limestones of Germany provide evidence for marine flipper adaptations in Late Jurassic thalassochelydians. *PLoS ONE* **16**, e0252355 (2021). https://doi.org/10.1371/journal.pone.0252355.

- Kelley, N. P., Motani, R., Jiang, D.-Y., Rieppel, O. & Schmitz, L. Selective extinction of Triassic marine reptiles during long-term sea-level changes illuminated by seawater strontium isotopes. *Palaeogeography, Palaeoclimatology, Palaeoecology* **400** , 9–16 (2014). http://dx.doi.org/10.1016/j.palaeo.2012.07.026.

- Ketchum, H. F. & Barrett, P. M. New reptile material from the Lower Triassic of Madagascar: implications for the Permian–Triassic extinction event. *Canadian Journal of Earth Sciences* **41**, 1–8 (2004). https://doi.org/10.1139/e03-084.

- Ketchum, H. F. & Benson, R. B. J. Global interrelationships of Plesiosaur (Reptilia, Sauropterygia) and the pivotal role of taxonomy sampling in determining the outcome of phylogenetic analyses. *Biological Reviews* **85**, 361–392 (2010). https://doi.org/10.1111/j.1469-185X.2009.00107.x.

- King, M. J. & Benton, M. J. Dinosaurs in the Early and Mid Triassic?—The footprint evidence from Britain. *Palaeogeography, Palaeoclimatology, Palaeoecology* **122**, 213–225 (1996). https://doi.org/10.1016/0031-0182(95)00095-X.

- Klein, N. & Sander, M. *Anarosaurus heterodontus*, the least aquatic pachypleurosaur. *Staringia* **16**, 226–233 (2019). https://natuurtijdschriften.nl/pub/1021544/.

- Klein, N. & Scheyer, T. M. Vertebral, rib, and osteoderm morphology and histology of Middle Triassic diapsid *Eusaurosphargis*. *Acta Palaeontologica Polonica* **69**, 633–648 (2024). https://doi.org/10.4202/app.01163.2024.

- Kornhuber, A. Uber einen neuen fossilen saurier aus Lesina. Herausgegeben von der k. k. geologischen Reichsanstalt. *Wien* **5**, 75–90 (1873).

- Krahl, A. The locomotory apparatus and paraxial swimming in fossil and living marine reptiles: comparing Nothosauroidea, Plesiosauria, and Chelonioidea. *PalZ* **95**, 483–501 (2021). https://doi.org/10.1007/s12542-021-00563-w.

- Kuhn-Schnyder, E. Über einen Schädel von *Askeptosaurus italicus* Nopsca aus der mittleren Trias des Monte San Giorgio (Kt. Tessin, Schweiz). *Abhandlungen des Hessischen Landesamts für Bodenforschung* **60**, 89–98 (1971).

- Lane, H. H. New mid-Pennsylvanian reptiles from Kansas. *Transactions of the Kansas Academy of Science* **47**, 381–390 (1945). https://doi.org/10.2307/3625454.

- Laurenti, J. N. *Specimen Medicum, Exhibens Synopsin Reptilium Emendatam cum Experimentis circa Venena et Antidota Reptilium Austracorum, quod Authoritate et Consensu*. Joan Thomae: Vienna. 217 pp. (1768).

- Laurin, M. The osteology of a Lower Permian eosuchian from Texas and a review of diapsid phylogeny. *Zoological Journal of the Linnean Society* **101**, 59–95 (1991). https://doi.org/10.1111/j.1096-3642.1991.tb00886.x.

- Laurin, M. & Reisz, R. R. A reevaluation of early amniote phylogeny. *Zoological Journal of the Linnean Society* **113**, 165–223 (1995). https://doi.org/10.1111/j.1096-3642.1995.tb00932.x.

- Laurin, M. & Reisz, R. R. *Amniota* E. Hackel 1866 [M. Laurin and R. R. Reisz], converted clade name. Registration Number 7. In: De Queiroz, K., Cantino, P. D. & Gauthier, J. A. (eds.) *Phylonyms: A Companion to the PhyloCode.* pp. 1027–1031 (2020a).

- Laurin, M. & Reisz, R. R. *Reptilia* C. Linnaeus 1758 [M. Laurin and R. R. Reisz], converted clade name. Registration Number 88. In: De Queiroz, K., Cantino, P. D. & Gauthier, J. A. (eds.) *Phylonyms: A Companion to the PhyloCode.* pp. 1027–1031 (2020b).

- Lee, Y.-N., Kong, D.-Y. & Jung, S.-H. The first possible choristoderan trackway from the Lower Cretaceous Daegu Formation of South Korea and its implications on choristoderan locomotion. *Scientific Reports* **10**, 14442 (2020). https://doi.org/10.1038/s41598-020-71384-1.

- Leidy, J. Descriptions of *Emys jaenesi*, *E. haydeni*, *Baena arenosa*, and *Saniwa ensidens*. *Proceedings of the American Philosophical Society* **1870**, 123–124 (1870).

- Li, C., Rieppel, O., Wu, X.-C., Zhao, L.-J. & Wang, L.-T. A new Triassic marine reptile from southwestern China. *Journal of Vertebrate Paleontology* **31**, 303–312 (2011). http://dx.doi.org/10.2307/25835826.

- Li, C., Jiang, D.-Y., Cheng, L., Wu, X.-C. & Rieppel, O. A new species of *Largocephalosaurus* (Diapsida: Saurosphargidae), with implications for the morphological diversity and phylogeny of the group. *Geological Magazine* **151**, 100–120 (2014). https://doi.org/10.1017/S001675681300023X.

- Lindgren, J., Polcyn, M. J. & Young, B. A. Landlubbers to leviathans: evolution of swimming in mosasaurine mosasaurs. *Paleobiology* **37**, 445–469 (2011). https://doi.org/10.1666/09023.1.

- Lindgren, J., Kaddumi, H. F. & Polcyn, M. J. Soft tissue preservation in a marine fossil lizard with a bilobed tail fin. *Nature Communications* **4**, 2423 (2013). https://doi.org/10.1038/ncomms3423.

- Linnaeus, C. *Systema Naturae Per Regna Tria Naturae, Secundum Classes, Ordines, Genera, Species, cum Characteribus, Diferentiis, Synonymis, Locis*. Tomus I, Editio X, reformata. Laurentius Salvius, Holmiae: Stockholm (1758).

- Linnaeus, C. *Systema Naturae Per Regna Tria Naturae, Secundum Classes, Ordines, Genera, Species, cum Characteribus, Diferentiis, Synonymis, Locis*. Tomus I, Editio XII. Laurentius Salvius, Holmiae: Stockholm (1766).

- Liu, J. Sauropterygian from Triassic of Guizhou, China. *Chinese Science Bulletin* **44**, 1312–1316 (1999). https://doi.org/10.1007/BF02885852.

- Macartney, J. "Tables of classifcation." In: Cuvier, G. (ed.) *Lectures on Comparative Anatomy*. Translated by William Ross under the inspection of James Macartney. T. N. Longman and O. Rees: London (1802).

- MacDougall, M. J., Verrière, A., Wintrich, T., LeBlanc, A. R. H., Fernandez, V. & Fröbisch, J. Conflicting evidence for the use of caudal autotomy in mesosaurs. *Scientific Reports* **10**, 7184 (2020). https://doi.org/10.1038/s41598-020-63625-0

- Madar, S. I. The postcranial skeleton of early Eocene pakicetid cetaceans. *Journal of Paleontology* **81**, 176–200 (2007). https://www.jstor.org/stable/4133804.

- Madzia, D. & Conrad, J. L. *Mosasauridae* P. Gervais 1853 [D. Madzia and J. L. Contrad], converted clade name. Registration Number 200. In: De Queiroz, K., Cantino, P. D. & Gauthier, J. A. (eds.) *Phylonyms: A Companion to the PhyloCode.* pp. 1103–1107 (2020).

- Magnusson, W. E. & Lima, A. P. The ecology of a cryptic predator, *Paleosuchus trigonatus*, in a tropical rainforest. *Journal of Herpetology* **25**, 41–48 (1991). https://doi.org/10.2307/1564793.

- Magnusson, W. E., da Silva, E. V. & Lima, A. P. Diets of Amazonian crocodilians. *Journal of Herpetology* **21**, 85–95 (1987). https://doi.org/10.2307/1564468.

- Magnusson, W. E., Campos, Z. & Muniz, F. *Paleosuchus palpebrosus*. *The IUCN Red List of Threatened Species* 2019: e.T46587A3009946 (2019). http://dx.doi.org/10.2305/IUCN.UK.2019-1.RLTS.T46587A3009946.en.

- Manegold, A. Two additional synapomorphies of grebes Podicipedidae and flamingos Phoenicopteridae. *Acta Ornithologica* **41**, 79–82 (2006). https://doi.org/10.3161/068.041.0113.

- Mantell, G. A tabular arrangement of the organic remains of the county of Sussex. *Transactions of the Geological Society of London* **3**, 201–216 (1829).

- Marcilly, C. M., Torsvik, T. H. & Conrad, C. P. Global Phanerozoic sea levels from paleogeographic flooding maps. *Gondwana Research* **110**, 128–142 (2022). https://doi.org/10.1016/j.gr.2022.05.011.

- Marsh, O. C. Notice of some new mosasauroid reptiles from the Greensand of New Jersey. *American Journal of Science* **48**, 392–397 (1869).

- Marsh, O. C. Notice of new Tertiary mammals. *American Journal of Science* **3**, 401–404 (1876).

- Massare, J. A. Tooth morphology and prey preference of Mesozoic marine reptiles. *Journal of Vertebrate Paleontology* **7**, 121–137 (1987). https://doi.org/10.1080/02724634.1987.10011647.

- Maxwell, E. E. Unraveling the influences of soft-tissue flipper development on skeletal variation using an extinct taxon. *Journal of Experimental Zoology Part B: Molecular and Developmental Evolution* **318B**, 545–554 (2012). https://doi.org/10.1002/jez.b.22459.

- Maxwell, E. E. & Caldwell, M. W. First record of live birth in Cretaceous ichthyosaurs: closing an 80 million year gap. *Proceedings of the Royal Society of London B* **270**, S104–S107 (2003). https://doi.org/10.1098/rsbl.2003.0029.

- Maxwell, E. E., Scheyer, T. M. & Fowler, D. A. An evolutionary and developmental perspective on the loss of regionalization in the limbs of derived ichthyosaurs. *Geological Magazine* **151**, 29–40 (2014). https://doi.org/10.1017/S0016756812001070.

- McFadden, D. Conditional logit analysis of qualitative choice behavior, in: Zarembka, P. (ed) *Frontiers in Econometrics*. Academic Press, New York, pp. 105–142 (1974).

- McMenamin, M. A. S. "Barasaurus Squamation", in: McMenamin, M. A. S. *Deep Time Analysis: A Coherent View of the History of Life*. Springer Geology. pp. 159–204 (2018).

- Meckert, D. The Procolophonid Barasaurus and the Phylogeny of Early Amniotes. PhD Thesis, Biology Department, McGill University. 149 pp (1995).

- Medem, F. J. The crocodilian genus *Paleosuchus*. *Fieldiana*: *Zoology* **39**, 227–247 (1958).

- Mehl, M. G. *Poposaurus gracilis*, a new reptile from the Triassic of Wyoming. *Journal of Geology* **23**, 516–522 (1915).

- Mekarski, M. C., Japundžić, D., Krizmanić, K. & Caldwell, M. W. Description of a new basal mosasauroid from the Late Cretaceous of Croatia, with comments on the evolution of the mosasauroid forelimb. *Journal of Vertebrate Paleontology* **39**, e1577872 (2019). https://doi.org/10.1080/02724634.2019.1577872.

- Merriam, J. C. A new marine reptile from the Triassic of California. *University of California Publications: Bulletin of the Department of Geology* **3**, 419–421 (1904).

- Miedema, F., Klein, N., Blackburn, D. G., Sander, P. M., Maxwell, E. E., Griezelen, E. M. & Scheyer, T. M. Heads or tails first? Evolution of fetal orientation in ichthyosaurs, with a scrutiny of the prevailing hypothesis. *BMC Ecology and Evolution* **23**, 12 (2023).

- Modesto, S. P. The postcranial skeleton of the aquatic parareptile *Mesosaurus tenuidens* from the Gondwanan Permian. *Journal of Vertebrate Paleontology* **30**, 1378–1395 (2010). https://doi.org/10.1080/02724634.2010.501443.

- Mongiardino-Koch, N. & Gauthier, J. A. Noise and biases in genomic data may underlie radically different hypotheses for the position of Iguania within Squamata. *PLoS ONE* **13**, e0202729 (2018). https://doi.org/10.1371/journal.pone.0202729.

- Montagu, G. Description of a species of *Delphinus*, which appears to be new. *Memoirs of the Wernerian Natural History Society* **3**, 75–82 (1821).

- Montañez, I. P. & Poulsen, C. J. The Late Paleozoic ice age: An evolving paradigm. *Annual Review of Earth and Planetary Sciences* **41**, 629–656 (2013). https://doi.org/10.1146/annurev.earth.031208.100118.

- Moon, B. C. A new phylogeny of ichthyosaurs (Reptilia: Diapsida). *Journal of Systematic Paleontology* **17**, 129–155 (2019). https://doi.org/10.1080/14772019.2017.1394922.

- Motani, R. Phylogeny of the Ichthyopterygia. *Journal of Vertebrate Paleontology* **19**, 473–496 (1999). https://www.jstor.org/stable/4524011.

- Motani, R. & Vermeij, G. J. Ecophysiological steps of marine adaptation in extant and extinct non-avian tetrapods. *Biological Reviews* **96**, 1769–1798 (2021). https://doi.org/10.1111/brv.12724.

- Motani, R., Jiang, D.-y., Tintori, A., Rieppel, O. & Chen, G.-b. Terrestrial origin of viviparity in Mesozoic marine reptiles indicated by Early Triassic embryonic fossils. *PLOS ONE* **9**, e88640 (2014). https://doi.org/10.1371/journal.pone.0088640.

- Motani, R., Jiang, D.-Y., Chen, G.-B., Tintori, A., Rieppel, O., Ji, C. & Huang, J.-D. A basal ichthyosauriform with a short snout from the Lower Triassic of China. *Nature* **517**, 485–488 (2015). https://doi.org/10.1038/nature13866.

- Motani, R., Jiang, D.-Y., Tintori, A., Ji, C. & Huang, J.-d. Pre- versus post-mass extinction divergence of Mesozoic marine reptiles dictated by time-scale dependence of evolutionary rates. *Proceedings of the Royal Society of London B* **284**, 20170241 (2017). https://doi.org/10.1098/rspb.2017.0241.

- Müller, S. Waarnemingen over de Indische Krokodillen en Beschrijving Van Enne Nieuwe Soort. *Tijdschrift voor Natuurlijke Geschiedenis en Physiologie* **5**, 61–87 (1838).

- Müller, J. The anatomy of Askeptosaurus italicus from the Middle Triassic of Monte San Giorgio and the interrelationships of thalattosaurs (Reptilia, Diapsida). *Canadian Journal of Earth Sciences* **42**, 1347–1367 (2005). https://doi.org/10.1139/e05-030.

- Muragaki, Y., Mundlos, S., Upton, J. & Olsen, B. R. Altered growth and branching patterns in synpolydactyly caused by mutations in HOXD13. *Science* **272**, 548–551 (1996). https://doi.org/10.1126/science.272.5261.548.

- Murphy, W. J., Eizirik, E., O'Brien, S. J., Madsen, O., Scally, M., Douady, C. J., Teeling, E., Ryder, O. A., Stanhope, M. J., de Jong, W. W. & Springer, M. S. Resolution of the early placental mammal radiation using Bayesian phylogenetics. *Science* **294**, 2348–2351 (2001). https://doi.org/10.1126/science.1067179.

- Murray, C. M., Crother, B. I. & Doody, J. S. The evolution of crocodilian nesting ecology and behavior. *Ecology and Evolution* **10**, 131–149 (2020). https://doi.org/10.1002/ece3.5859.

- Neenan, J. M., Klein, N. & Scheyer, T. M. European origin of placodont marine reptiles and the evolution of crushing dentition in Placodontia. *Nature Communications* **4**, 1621 (2013). https://doi.org/10.1038/ncomms2633.

- Nicholls, E. L. & Brinkman, D. B. "A new ichthyosaur from the Triassic Sulphur Mountain formation of British Columbia." In, Sarjeant, W. A. S. (ed.) *Vertebrate fossils and the evolution of scientific concepts*. Gordon and Breach: Switzerland, pp. 521–535 (1995).

- Nopcsa, F. V. *Askeptosaurus, ein neues reptil der Trias von Besano: Centralblatt für Mineralogie, Geologie und Paläontologie B*. pp. 265–267 (1925).

- Nopcsa, F. V. The Genera of Reptiles. *Palaeobiologica* **1**, 163–188 (1928).

- Norell, M. A., Clark, J. M. & Hutchison, J. H. The Late Cretaceous alligatoroid Brachychampsa montana (Crocodylia): new material and putative relationships. *American Museum Novitates* **3116**, 1–26 (1994).

- Nosotti, S. & Rieppel, O. *Eusaurosphargis dalsassoi* n. gen n. sp., a new, unusual diapsid reptile from the Middle Triassic of Besano (Lombardy, N Italy). *Memorie della Società italiana di scienze naturali Museo Civico di Storia Naturale di Milano* **31**, 3–33 (2003).

- Ogg, J. G., Chen, Z.-Q., Orchard, M. J. & Jiang, H. S. The Triassic Period. Chapter 25 in: Gradstein, F. M., Ogg, J. G., Schmitz, M. D., Ogg, G. M. *Geologic Time Scale 2020*. Elsevier BV, pp. 903–953 (2020). https://doi.org/10.1016/B978-0-12-824360-2.00025-5.

- O'Keefe, F. R. & Chiappe, L. M. Viviparity and K-selected life history in a Mesozoic marine plesiosaur (Reptilia, Sauropterygia). *Science* **333**, 870–873 (2001).

- Oppel, M. *Die Ordnungen, Familien und Gattungen der Reptilien als Prodrom einer Naturgeschichte Derselben.* Joseph Lindauer: München. 87 pp. (1811).

- Osborn, H. F. A complete skeleton of *Teleoceras fossiger*. Notes upon the growth and sexual characters of this species. *Bulletin of the American Museum of Natural History* **10**, 51–59 (1898).

- Owen, R. Description of teeth and portions of jaws of two extinct anthracotherioid quadrupeds (*Hyopotamus vectianus* and *Hyop. bovinus*) discovered by the marchioness of Hastings in the Eocene deposits on the N.W. coast of the Isle of Wight: with an attempt to develop Cuvier's idea of the classification of pachyderms by the number of their toes. *Quarterly Journal of the Geological Society of London* **4**,103–293 (1848).

- Owen, R. Description of the fossil reptiles of the chalk formation. In: *The Geology and Fossils of the Tertiary and Cretaceous Formations of Sussex.* Longman, Brown, Green and Longman*:* London, pp. 378–404 (1850).

- Owen, R. On the orders of fossil and recent Reptilia, and their distribution in time. *Report of the British Association for the Advancement of Science* **29**, 153–166 (1859).

- Paradis, E. & Schliep, K. ape 5.0: an environment for modern phylogenetics and evolutionary analyses in R. *Bioinformatics* **35**, 526–528 (2019). https://doi.org/10.1093/bioinformatics/bty633.

- Parrington, F. R. XVI.— *On* Prolacerta broomi, gen. et sp. n., and the origin of lizards. *Annals and Magazine of Natural History* **16**, 197–205 (1935).

- Pauwels, O. S. G., Barr, B., Sanchez, M. L. & Burger, M. Diet records for the Dwarf Crocodile, *Osteolaemus tetraspis tetraspis* in Rabi Oil fields and Loango National Park, southwestern Gabon. *Hamadryad* **31**, 258–264 (2007).

- Petermann, H. & Sander, M. Histological evidence for muscle insertion in extant amniote femora: implications for muscle reconstruction in fossils. *Journal of Anatomy* **222**, 419–436 (2013). https://doi.org/10.1111/joa.12028.

- Peterson, O. A. II. Report upon the material discovered in the Upper Eocene of the Uinta Basin by Earl Douglas in the years 1908–1909, and by O. A. Peterson in 1912. *Annals of the Carnegie Museum* **12**, 40–168 (1919).

- Peyer, B. *Clarazia schinzi* nov. gen. nov. spec. *Abhandlungen der Schweizerischen Paläontologischen Gesellschaft* **57**, 1–61 (1936).

- Peyer, B. *Helveticosaurus zollingeri* n. g. n. sp. *Schweizerische palaeontologische Abhandlungen [Mémoires suisses de paléontologie* **72**, 1–50 excluding plates (1955).

- Piñeiro, G., Ferigolo, J., Ramos, A. & Laurin, M. Cranial morphology of the Early Permian mesosaurid *Mesosaurus tenuidens* and the evolution of the lower temporal fenestration reassessed. *Comptes Rendus Palevol* **11**, 379–391 (2012). https://doi.org/10.1016/j.crpv.2012.02.001.

- Piñeiro, G., Ferigolo, J., Mones, A. & Demarco, P. N. Mesosaur taxonomy reappraisal: Are *Stereosternum* and *Brazilosaurus* valid taxa? *Revista Brasileira de Paleontologia* **24**, 205–235 (2021). https://doi.org/10.4072/rbp.2021.3.04.

- Piveteau, J. Paleontologie de Madagascar: XII. Amphibiens et reptiles Permiens. *Annales de Paleontologie* **15**, 55–179 (1926).

- Piveteau, J. Existence d'un Reptile du groupe des Procolophonidés à Madagascar - Conséquences stratigraphiques et paléontologiques. *Comptes Rendus Hebdomadaires des Séances de l'Academie des Sciences* **241**, 1325–1327 (1955).

- Price, L. I. Two new cotylosaurs from the Permian of Texas. *Proceedings of the New England Zoölogical Club* **16**, 97–102 (1937).

- Regnault, S., Jones, M. E. H., Pitsillides, A. A. & Hutchinson, J. R. Anatomy, morphology and evolution of the patella in squamate lizards and tuatara (*Sphenodon punctatus*). *Journal of Anatomy* **228**, 864–876 (2016). https://doi.org/10.1111/joa.12435.

- Reig, O. A. La presencia de dinosaurios saurisquios en los "Estratos de Ischigualasto" (Mesotriasico Superior) de las provincias de San Juan y La Rioja (República Argentina) [The presence of saurischian dinosaurs in the "Ischigualasto beds" (upper Middle Triassic) of San Juan and La Rioja Provinces (Argentine Republic)]. *Ameghiniana* **3**, 3–20 (1963).

- Reisz, R. R., Heaton, M. J. & Pynn, B. R. Vertebrate fauna of Late Pennsylvanian Rock Lake Shale near Garnett, Kansas: Pelycosauria. *Journal of Paleontology* **56**, 741–750 (1982). https://www.jstor.org/stable/1304403.

- Renesto, S. A new lepidosaur (Reptilia) from the Norian beds of the Bergamo Prealps (Preliminary Note). *Rivista Italiana di Paleontologia e Stratigrafia* **90**, 165–176 (1984). https://doi.org/10.54103/2039-4942/13332.

- Renesto, S. & Posenato, R. A new lepidosauromorph reptile from the Middle Triassic of the Dolomites. *Rivista Italiana di Paleontologia e Stratigrafia* **109**, 463–474 (2003). http://dx.doi.org/10.13130/2039-4942/5517.

- Rieppel, O. *Helveticosaurus zollingeri* Peyer (Reptilia, Diapsida) skeletal paedomorphosis, functional anatomy and systematic affinities. *Palaeontographica Abteilung. A* **208**, 123–152 (1989).

- Rieppel, O. Osteology of *Simosaurus gaillardoti* and the relationships of stem-group Sauropterygia. *Fieldiana Geology, New Series* **28**, 1–85 (1994). https://doi.org/10.5962/bhl.title.3399.

- Rieppel, O. *Corosaurus alcovensis* Case and the phylogenetic interrelationships of Triassic stem-group Sauropterygia. *Zoological Journal of the Linnean Society* **124**,1–41 (1998). https://doi.org/10.1111/j.1096-3642.1998.tb00568.x.

- Rieppel, O., Müller, J. & Liu, J. Rostral structure in Thalattosauria (Reptilia, Diapsida). *Canadian Journal of Earth Sciences* **42**, 2081–2086 (2005). https://doi.org/10.1139/e05-076.

- Rieppel, O. & Grande, L. The anatomy of the fossil varanid lizard *Saniwa ensidens* Leidy, 1870, based on a newly discovered complete skeleton. *Journal of Paleontology* **81**, 643–665 (2007). https://www.jstor.org/stable/4498815.

- Rio, J. P. & Mannion, P. D. Phylogenetic analysis of a new morphological dataset elucidates the evolutionary history of Crocodylia and resolves the long-standing gharial problem. *PeerJ* **9**, e12094 (2021). http://dx.doi.org/10.7717/peerj.12094.

- Robinson, J. A. The locomotion of plesiosaurs. *Neues Jahrbuch für Geologie und Paläontologie, Abhandlungen* **149**, 286–332 (1975). https://doi.org/10.1127/njgpa/149/1975/286.

- Romer, A. S. *Osteology of the Reptiles*. Krieger Publishing Company: Malabar, 772 pp. (1956).

- Romer, A. S. & Parsons, T. S. *The Vertebrate Body*, Sixth Edition (1986). Saunders College Publishing: Philadelphia. 679 pp. (1949).

- Root, R. B. The niche exploitation pattern of the blue-gray gnatcatcher. *Ecological Monographs* **37**, 317–350 (1967). https://doi.org/10.2307/1942327.

- Rösler, O. & Tatizana, C. As membranas natatorias em *Stereosternum tumidum*. *Coletânea de trabalhos paleontológicos* **27**, 129–31 (1985).

- Rothier, P. S., Fabre, A.-C., Benson, R. B. J., Martinez, Q., Fabre, P.-H., Hedrick, B. P. & Herrel, A. Of flippers and wings: The locomotor environment as a driver of the evolution of forelimb morphological diversity in mammals. *Functional Ecology* **00**, 1–16 (2024). https://doi.org/10.1111/1365-2435.14632.

- Rowe, T. B. Chordate phylogeny and development. In: Cracraft, J. & Donoghue, M. J. (eds.) *Assembling the Tree of Life*. Oxford University Press: Oxford. pp. 384–409 (2004).

- Rowe, T. B. *Mammalia* C. Linnaeus 1758 [T. B. Rowe], converted clade name. Registration Number 220. In: De Queiroz, K., Cantino, P. D. & Gauthier, J. A. (eds.) *Phylonyms: A Companion to the PhyloCode.* CRC Press, Taylor & Francis: Boca Raton. pp. 859–865 (2020a).

- Rowe, T. B. *Pan-Mammalia* T. B. Rowe 2004 [T. B. Rowe], converted clade name. Registration Number 224. In: De Queiroz, K., Cantino, P. D. & Gauthier, J. A. (eds.) *Phylonyms: A Companion to the PhyloCode.* CRC Press, Taylor & Francis: Boca Raton. pp. 801–810 (2020b).

- Russell, D. A. Systematics and morphology of American mosasaurs. *Bulletin of the Peabody Museum of Natural History* **23**, 1–240 (1967). https://doi.org/10.2307/j.ctvxkn75d.

- Sander, P. M., Rieppel, O. C. & Bucher, H. A new pistosaurid (Reptilia: Sauropterygia) from the Middle Triassic of Nevada and its implications for the origins of plesiosaurs. *Journal of Vertebrate Paleontology* **17**, 526–533 (1997). https://doi.org/10.1080/02724634.1997.10010999.

- Scheyer, T. M., Neenan, J. M., Bodogan, T., Furrer, H., Obrist, C. & Plamondon, M. A new, exceptionally preserved juvenile specimen of *Eusaurosphargis dalsassoi* (Diapsida) and implications for Mesozoic marine diapsid phylogeny. *Scientific Reports* **7**, 4406 (2017). https://doi.org/10.1038/s41598-017-04514-x.

- Schneider, J. G. *Allgemeine Naturgeschichte der Schildkröten, Nebst Einem Systematischen Verzeichnisse der Einzelnen Arten und Zwey Kupfern*. Johann Gotfried Müllersche Buchhandlung: Leipzig (1783).

- Schneider, J. G. *Historiae Amphibiorum naturalis et literariae Fasciculus Secundus continens Crocodilos, Scincos, Chamaesauras, Boas, Pseudoboas, Elapes, Angues, Amphisbaenas et Caecilias*. Jenae: Wesselhoeft. 364 pp. Excluding plates (1801).

- Scotese, C. R. & Langford, R. P. Pangea and the paleogeography of the Permian. In: Scholle, P.A., Peryt, T.M. & Ulmer-Scholle, D. S. (eds.) *The Permian of Northern Pangea.* Springer, Berlin, pp. 3–19 (1995). https://doi.org/10.1007/978-3-642-78593-1_1.

- Seeley, H. G. On remains of a small lizard from Neocomian rocks of Comen, near Trieste, preserved in the Geological Museum of the University of Vienna. *Quarterly Journal of the Geological Society of London* **37**, 52–56 (1881). https://doi.org/10.1144/GSL.JGS.1881.037.01-04.07.

- Seeley, H. G. "V. Researches on the structure, organization, and classification of the fossil reptilia.—III. On Parts of the Skeleton of a Mammal from Triassic Rocks of Klipfontein, Fraserberg, South Africa (Theriodesmus phylarchus, Seeley), illustrating the Reptilian Inheritance in the Mammalian Hand." *Philosophical Transactions of the Royal Society of London B* **179**, 141–155 (1888).

- Seeley, H. G. "XI. Researches on the Structure, Organisation, and Classification of the Fossil Reptilia.—Part IX., Section 1. On the Therosuchia." *Philosophical Transactions of the Royal Society of London B* **185**, 987–1018 (1894).

- Sereno, P. C., Forster, C. A., Rogers, R. R. & Monetta, A. M. Primitive dinosaur skeleton from Argentina and the early evolution of Dinosauria. *Nature* **361**, 64–66 (1993). https://doi.org/10.1038/361064a0.

- Sereno, P. C., Myhrvold, N., Henderson, D. M., Fish, F. E., Vidal, D., Baumgart, S. L., Keillor, T. M., Formoso, K. K. & Conroy, L. L. *Spinosaurus* is not an aquatic dinosaur. *eLife* **11**, e80092 (2022). https://doi.org/10.7554/eLife.80092.

- Seton, M., Müller, R. D., Zahirovic, S., Gaina, C., Torsvik, T., Shephard, G., Talsma, A., Gurnis, M., Turner, M., Maus, S. & Chandler, M. Global continental and clean basin reconstructions since 200 Ma. *Earth-Science Reviews* **113**, 212–270 (2012). http://dx.doi.org/10.1016/j.earscirev.2012.03.002.

- Shaw, G. *The Porcupine Ant-Eater. The Naturalist's Miscellany, or Coloured Figures of Natural Objects Drawn and Described Immediately from Nature*. Plate 109. London. 7 Vols. (1790–1813, here 1792).

- Shen, S.-z., Crowley, J. L., Wang, Y., Bowring, S. A., Erwin, D. H., Sadler, P. M., Cao, C.-q., Rothman, D. H., Henderson, C. M., Ramezani, J., Zhang, H., Shen, Y., Wang, X.-d., Wang, W., Mu, L., Li, W.-z., Tang, Y.-g., Liu, X.-l., Liu, L.-j., Zeng, Y., Jiang, Y.-f. & Jin, Y.-g. Calibrating the end-Permian mass extinction. *Science* **334**, 1368–1372 (2011). https://doi.org/10.1126/science.1213454.

- Shikama, T. & Kamei, T. Early Triassic ichthyosaurus, *Utatsusaurus hataii* gen. et sp. nov, from the Kitakami Massif, northeast Japan. *Tohoku University Scientific Report, 2nd series* **48**, 77–97 (1978).

- Shikama, T. & Ozaki, H. On a reptilian skeleton from the Palaeozoic formation of San Paulo, Brazil. *Transactions and Proceedings of the Palaeontological Society of Japan* **64**, 351–358 (1966).

- Shirley, M. H., Burtney, B., Oslisly, R., Sebag, D. & Testa, O. Diet and body condition of cave-dwelling dwarf crocodiles (*Osteolaemus tetraspis*, Cope 1861) in Gabon. *African Journal of Ecology* **55**, 411–422 (2017). https://doi.org/10.1111/aje.12365.

- Shockey, B. J. *Xenarthra* E. D. Cope 1889 [B. J. Schoeky], converted clade name. Registration Number 257. In: De Queiroz, K., Cantino, P. D. & Gauthier, J. A. (eds.) *Phylonyms: A Companion to the PhyloCode.* pp. 895–897 (2020).

- Simões, T. R., Vernygora, O., Paparella, I., Jimenez-Huidobro, P. & Caldwell, M. W. Mosasauroid phylogeny under multiple phylogenetic methods provides new insights on the evolution of aquatic adaptations in the group. *PLoS ONE* **12**, e0176773 (2017). https://doi.org/10.1371/journal.pone.0176773.

- Simões, T. R., Caldwell, M. W., Talanda, M., Bernardi, M., Palci, A., Vernygora, O., Bernardini, F., Mancini, L. & Nydam, R. L. The origin of squamates revealed by a Middle Triassic lizard from the Italian Alps. *Nature* **557**, 706–709 (2018). https://doi.org/10.1038/s41586-018-0093-3.

- Simões, T. R., Kammerer, C. F., Caldwell, M. W. & Pierce, S. E. Successive climate crises in the deep past drove the early evolution and radiation of reptiles. *Science Advances* **8**, eabq1898 (2022). https://doi.org/10.1126/sciadv.abq1898.

- Smart, S. & Sakamoto, M. Using linear measurements to diagnose the ecological habitat of *Spinosaurus*. *PeerJ* **12**, e17544 (2024). https://doi.org/10.7717/peerj.17544.

- Soul, L. C. & Benson, R. B. J. Developmental mechanisms of macroevolutionary change in the tetrapod axis: A case study of Sauropterygia. *Evolution* **71**, 1164–1177 (2017). https://doi.org/10.1111/evo.13217.

- Spaulding, M., O'Leary, M. A. & Gatesy, J. Relationships of Cetacea (Artiodactyla) among mammals: Increased taxon sampling alters interpretations of key fossils and character evolution. *PLoS ONE* **4**, e7062 (2009). https://doi.org/10.1371/journal.pone.0007062.

- Spennemann, D. H. R. Cruising the currents: Observations of extra-limital saltwater crocodiles (*Crocodylus porosus* Schneider, 1801) in the Pacific Region. *Pacific Science* **74**, 211–213 (2020). https://doi.org/10.2984/74.3.1.

- Spindler, F., Werneburg, F. & Schneider, J. W. A new mesenosaurine from the lower Permian of Germany and the postcrania of *Mesenosaurus*: implications for early amniote comparative osteology. *Paläontologische Zeitschrift* **93**, 43 pp. (2019). https://doi.org/10.1007/s12542-018-0439-z.

- Spindler, F., Lauer, R., Tischlinger, H. & Mäuser, M. The integumentary of pelagic crocodylomorphs (Thalattosuchia: Metriorhynchidae). *Palaeontologia Electronica* **24**, a25 (2021). http://dx.doi.org/10.26879/1099.

- Stanhope, M. J., Waddell, V. G., Madsen, O., de Jong, W., Hedges, S. B., Cleven, G. C., Kao, D. & Springer, M. S. Molecular evidence for multiple origins of Insectivora and for a new order of endemic African insectivore mammals. *Proceedings of the National Academy of Science* **95**, 9967–9972 (1998). https://doi.org/10.1073/pnas.95.17.9967.

- Staniewicz, A., Behler, N., Dharmasyah, S. & Jones, G. Niche partitioning between juvenile sympatric crocodilians in Mesangat Lake, East Kalimantan, Indonesia. *Raffles Bulletin of Zoology* **66**, 528–537 (2018).

- Stromer, E. Ergebnisse der Forschungsreisen Prof. E. Stromers in den Wüsten Ägyptens. II. Wirbeltier-Reste der Baharîje-Stufe (unterstes Cenoman). 3. Das Original des Theropoden *Spinosaurus aegyptiacus* nov. gen., nov. spec. *Abhandlungen der Königlichen Bayerischen Akademie der Wissenschaften, Mathematisch-physikalische Klasse* **28**, 1–32 (1915).

- Stubbs, T. L. & Benton, M. J. Ecomorphological diversifications of Mesozoic marine reptiles: the roles of ecological opportunity and extinction. *Paleobiology* **42**, 547–573 (2016). https://doi.org/10.1017/pab.2016.15.

- Sues, H.-D. *The Rise of Reptiles: 320 Million Years of Evolution*. Johns Hopkins University Press: Baltimore. 385 pp (2019).

- Thewissen, J. G. M. & Fish, F. E. Locomotor evolution in the earliest cetaceans: functional model, modern analogues, and paleontological evidence. *Paleobiology* 23: 482–490 (1997). https://doi.org/10.1017/S0094837300019850.

- Thewissen, J. G. M. & Nummela, S. "Introduction: On Becoming Aquatic". Chapter 1, in: Thewissen, J. G. M. & Nummela, S. (eds.). *Sensory Evolution on the Threshold: Adaptations in Secondarily Aquatic Vertebrates.* University of California Press: Berkeley, pp. 1–28 (2008).

- Thewissen, J. G. M. & Taylor, M. A. Aquatic adaptations in the limbs of amniotes. Chapter 18 in: Hall, B. K. (ed.) *Fins into Limbs: Evolution, Development, and Transformation*. The University of Chicago Press: Chicago. pp. 310–322 (2007).

- Uwakaneme, C., Rees, M. & Melfi, V. A. Snap happy: crocodile enrichment. In: Macdonald, C. (ed). *Proceedings of the 6th Annual Symposium on Zoo Research*, Edinburgh Zoo, Edinburgh, UK, 8–9th July 2004, pp. 297–301 (2004) [conference paper]. Downloaded from https://cabidigitallibrary.org.

- Verrière, A. & Fröbisch, J. Ontogenetic, dietary, and environmental shifts in Mesosauridae. *PeerJ* **10**, e13866 (2022). https://doi.org/10.7717/peerj.13866.

- Vila, B., Castanera, D., Marmi, J., Canudo, J. I. & Galobart, À. Crocodile swim tracks from the latest Cretaceous of Europe. *Lethaia* **48**, 256–266 (2014). https://doi.org/10.1111/let.12103.

- Vincent, P., Allemand, R., Taylor, P. D., Suan, G. & Maxwell, E. E. New insights on the systematics, palaeoecology and palaeobiology of a plesiosaurian with soft tissue preservation from the Toarcian of Holzmaden, Germany. *The Science of Nature* **104**, 51 (2017). https://doi.org/10.1007/s00114-017-1472-6.

- Vitt, L. J. & Caldwell, J. P. *Herpetology: An Introductory Biology of Amphibians and Reptiles*, Fourth Edition. Elsevier: Amsterdam. 757 pp. (2014).

- Von Fritsch, K. Beitrag zur Kenntnis der Saurier des Halle'schen unteren Muschelkalkes. *Abhandlungen der Jvatulfrschmden Gesellschaft zu Halle* **20**, 273–302 (1894).

- Von Huene, F. *Henodus chelyops*, ein neuer Placodontier. *Palaeontographica* **A84**, 99–148 (1936).

- Von Huene, F. Die großen Stämme der Tetrapoden in den geologischen Zeiten. *Biologisches Zentralblatt* **65**, 266–275 (1946).

- Von Meyer, H. Beiträge zur Petrefactenkunde. In: *Verhandlungen der Kaiserlichen Leopoldinisch-Carolinischen Akademie der Naturforscher [Nova Acta Physico-Medica Académie Caesarea Leopoldino-Carolinae], Fifteenth Volume, Pars Posterior*, Eduard Weber's Bookstore: Bonne, pp. 57–201 (1831).

- Von Meyer, H. *Palaeologica zur Geschichte der Erd und ihrer Geschöpfe.* Siegmund Schmerber: Frankfurt. 560 pp. (1832).

- Von Meyer, H. Mittheilung an Professor Bronn gerichtet. *Neues Jahrbuch für Mineralogie Geogosie und Geol Petrefaktenkunde* **1839**, 559–560 (1839).

- Waddell, P. J., Cao, Y., Hasegawa, M. & Mindell, D. P. Assessing the Cretaceous superordinal divergence times within birds and placental mammals by using whole mitochondrial protein sequences and an extended statistical framework. *Systematic Biology* **48**, 119–137 (1999). https://doi.org/10.1080/106351599260481.

- Wagner, G. P. & Chiu, C.-H. The tetrapod limb: a hypothesis on its origin. *Journal of Experimental Zoology Part B: Molecular and Developmental Evolution* **291**, 226–240 (2001). https://doi.org/10.1002/jez.1100.

- Wang, B. & Secord, R. Paleoecology of Aphelops and Teleoceras (Rhinocerotidae) through an interval of changing climate and vegetation in the Neogene of the Great Plains, central United States. *Palaeogeography, Palaeoclimatology, Palaeoecology* **542**, 109411 (2020). https://doi.org/10.1016/j.palaeo.2019.109411.

- Wang, W., Shang, Q., Cheng, L., Wu, X.-C., Li, C. Ancestral body plan and adaptive radiation of sauropterygian marine reptiles. *iScience* **25**, 105635 (2022). http://dx.doi.org/10.1016/j.isci.2022.105635.

- Watson, D. M. S., 1912, Mesosuchus browni, gen. et spec. nov. a fossil reptile from South Africa. *Records of the Albany Museum* **2**, 296–297. https://hdl.handle.net/2027/uc1.c079647719.

- Whitaker, R. & Basu, D. The gharial (*Gavialis gangeticus*): A review. *Journal, Bombay Natural Hist. Society* **79**, 531–548 (1982).

- Wieland, G. R. *Archelon ischyro*s: a new gigantic cryptodire testudinate from the Fort Pierre Cretaceous of South Dakota. *American Journal of Science*, Fourth Series **2**, 399–412 (1896). https://doi.org/10.2475/ajs.s4-2.12.399.

- Wieland, G. R. Notes on the Cretaceous Turtles, *Toxochelys* and *Archelon*, with a classification of the marine Testudinata. *American Journal of Science*, Fourth Series **80**, 95–108 (1902). https://doi.org/10.2475/ajs.s4-14.80.95.

- Wilberg, E. W., Turner, A. H. & Brochu, C. A. Evolutionary structure and timing of major habitat shifts in Crocodylomorpha. *Scientific Reports* **9**, 514 (2019). https://doi.org/10.1038/s41598-018-36795-1.

- Williston, S. W. New Permian reptiles: Rhachitomous vertebrae. *Journal of Geology* **18**, 585–600 (1910). https://www.jstor.org/stable/30078133.

- Williston, S. W. An ancestral lizard from the Permian of Texas. *Science* **38**, 825–826 (1913). https://doi.org/10.1126/science.38.988.825.

- Williston, S. W. *Water Reptiles of the Past and Present*. University of Chicago Press: Chicago, 251 pp. (1914).

- Wilson, D. E., Bogan, M. A., Brownell, R. L., Burdin, A. M. & Maminov, M. K. Geographic variation in sea otters, *Enhydra lutris*. *Journal of Mammalogy* **72**, 22–36 (1991). https://doi.org/10.2307/1381977.

- Witmer, L. M. The extant phylogenetic bracket and the importance of reconstructing soft tissues in fossils. Chapter 2 in: Thomason, J. (ed.), *Functional Morphology in Vertebrate Paleontology*. Cambridge University Press, pp. 19–33 (1995).

- Wolniewicz, A. S., Shen, Y., Li, Q., Suń, Y., Qiao, Y., Chen, Y., Hu, Y.-W. & Liu, J. Ań armoured marinę reptile from the Early Triassic of South China and its phylogenetic and evolutionary implications. *eLife* **12**, e83163 (2023). https://doi.org/10.7554/eLife.83163.

- Wolpert, L., Tickle, C., Jessell, T., Lawrence, P., Meyerowitz, E., Robertson, E. & Smith, J. *Principles of Development*, Fourth Edition. Oxford University Press: Oxford, 616 pp. (2011).

- Wyneken, J. The anatomy of sea turtles. *Department of Commerce NOAA Technical Memorandum NMFS-SEFSC-470*. Southeast Fisheries Science Center, Seattle, WA, U.S. (2001). Downloaded from https://repository.library.noaa.gov/view/noaa/8502.

- Xie, M., Gol'din, P., Herdina, A. N., Estefa, J., Medvedeva, E. V., Li, L., Newton, P. T., Kotova, S., Shavkuta, B., Saxena, A., Shumate, L. T., Metscher, B. D., Großschmidt, K., Nishimori, S., Akovantseva, A., Usanova, A. P., Kurenkova, A. D., Kumar, A., Arregui, I. L., Tafforeau, P., Fried, K., Carlström, M., Simon, A., Gasser, C., Kronenberg, H. M., Bastepe, M., Cooper, K. L., Timashev, P., Sanchez, S., Adameyko, I., Eriksson, A., Chagin, A. S. Secondary ossification center induces and protects growth plate structure. *eLife* **9**, e55212 (2020). https://doi.org/10.7554/eLife.55212.

- Yin, G. Z., Zhou, X., Cao, Z., Yu, Y. & Luo, Y. A preliminary study on the early Late Triassic marine reptiles from Guanling, Guizhou, China. *Geology, Geochemistry* **28**, 1–22 (2000).

- Youden, W. J. Index for rating diagnostic tests. *Cancer* **3**, 32–35 (1950). https://doi.org/10.1002/1097-0142(1950)3:1%3C32::aid-cncr2820030106%3E3.0.co;2-3.

- Young, C. C. On the new Pachypleurosauroidea from Keichow, south-west China. *Vertebrate PalAsiatica* **9**, 337–356 (1958).

- Young, C. C. A marine lizard from Nanchang, Hupeh Province. *Memoirs of the Institute of Vertebrate Paleontology and Paleoanthropology, Academia Sinica* **9**, 17–28 (1972).

- Young, C. C. & Dong, Z.-M. On the aquatic reptiles of the Triassic in China. *Vertebrate Paleontology Memoirs* **9**, 1–34 (1972).
