## Supplementary R Scripts for "Limb proportions predict aquatic habits and soft-tissue flippers in extinct amniotes": script_S1.R

```
#** First R script (script_S1.R) for Gordon et al., "Limb proportions predict aquatic habits and soft-tissue flippers in extinct amniotes."**

# NOTE ON READABILITY: This script contains many long, multi-line comments. To increase readability within Rstudio, go to Tools > Global Options, select the 'Code' tab, and check 'Soft-wrap R source files'. This will wrap all lines of code to match your personal GUI's margins, so that no lines of code or explanation run offscreen or cutoff at weird places.

# INTRODUCTION: In this R script, we will walk through all the steps involved in analyzing the linear morphometric ("LM") data associated with the Nature Ecology & Evolution manuscript "Limb proportions predict aquatic habits and soft-tissue flippers in extinct amniotes." Our intention is for anyone with experience coding and performing statistical analysis to be able to follow the steps below. If you have any questions about this Rmd file or the analyses we ran, please feel free to email the corresponding author of this manuscript.

# For this project, we generated >11,500 original limb bone measurements from >700 amniote specimens. We scored extant taxa in our dataset for their known aquatic habits (e.g., whether they are fully terrestrial, transiently aquatic, moderately aquatic, highly aquatic, or fully aquatic), their known soft-tissue limb morphologies (e.g., whether their hands are unwebbed, webbed, or flippered), their IUCN Red List status, and a number of other categorical variables. We began our analysis by cleaning the data, performing some measurement-quality-control tests (eg, Bland-Altman analysis), and generating eight alternative time-calibrated supertrees to account for differing hypotheses of phylogenetic relationships among the taxa in our dataset. We then evaluated general patterns and trends in the data——testing for associations among levels of different categorical variables (eg, soft-tissue limb morphology and aquatic habits), pairwise correlations between quantitative (LM) variables, limb proportion patterns within ternary morphospace, and significant differences in the mean and variance of various limb- and limb-bone shape metrics among groups for each categorical variable. Next, we used a phylogenetically informed machine-learning approach to predict the aquatic habits and soft-tissue limb phenotypes of extinct species: This involved training phylogenetic binomial logistic regression models on extant taxa, assessing the relative predictive accuracies of different models using Receiver Operating Characteristic (ROC) curve analysis, using the most accurate models to predict the phenotypes of extinct taxa within our dataset, and using the results of the best models, in tandem with ancestral-state reconstructions, to reconstruct the evolutionary history of flippers and aquatic habits along each major lineage of aquatic amniotes.

# Taken together, this data analysis pipeline requires extensive computational resources to run quickly. In order to maximize parallelization and make it run as efficiently as possible, we have divided the pipeline above into FIVE SCRIPTS, outlined below. These scripts are divided primarily based on their computational requirements. Each of them pursues a different set of tasks with differing parallelization techniques that make it run more efficiently.

#**The five scripts are labeled S1, S2, S3, S4, and S5.**
# You will run script S1 first, and then, after S1 is complete, you will run scripts S2-S4 at the same time. Once those are done running, you will run S5. 

# The first script, S1, is the one you're reading right now. It's the introduction to all the other scripts, it covers the first several steps of the data analysis pipeline for this project, and it should be run before (upstream of) all the others. This script is the least computationally intensive of the five, and should be run interactively. It involves preparing the coding environment, running quality control on measurements, generating time-calibrated trees, defining functions to align trees to datasets, and performing a few simple statistical tests and comparisons that do not require massive computational resources. It ends by saving its R environment to the working directory. This S1 R environment output will serve as a key input file for every subsequent script.

# Scripts S2-S4 can all be run in parallel after S1, as they do not depend on one another. S2 performs all phylogenetic tests comparing quantitative variables among groups, and uses moderate parallelization (in the form of foreach-loops) to increase computation rate. S3 runs pairwise phylogenetic correlation tests for all variable combinations--with raw, log-transformed, and BoxCox-transformed datasets. S4 produces phylogenetic binomial logistic regression (phybLR) models that use raw, log-transformed, and BoxCox-transformed datasets to predict grouping variable classifications. Scripts S2-S4 are very computationally intensive. S2 utilizes moderate parallelization (in the form of foreach-loops) and should be run as a batch job. S3 & S4 are designed to run in massively parallel fashion using Job Arrays. 

# Script S5 is the final script in this series. It must be run after scripts S2-S4. This final script involves applying ROC analysis to select the best phylogenetic binomial logistic regression models for predicting phenotype classifications, using those best models to predict phenotype classifications for extinct tip taxa in the dataset, performing ancestral state reconstructions to reconstruct the evolutionary history of these phenotypes, and then performing a comprehensive set of sensitivity analyses to assess the extent to which supertree topology and transformation method altered the results of our statistical tests from previous scripts (S2-S4). 

# A sixth script, numbered S6, is also associated with this paper, but not detailed here because it does not concern LM data; instead, it concerns geometric morphometric (GM) data, and we provide the required context for it in script_S6.

# A more granular outline of scripts S1-S5 is provided below:

#**[S1] ALL UPSTREAM STUFF**
# --- [S1.1]: PREPARE CODING ENVIRONMENT: Load packages, datasets, etc.
# --- [S1.2]: DO BLAND-ALTMAN ANALYSIS: Compare 2D & 3D measurements.
# --- [S1.3]: MAKE TIME-CALIBRATED TREES: Upload, tip-date, and plot trees.
# --- [S1.4]: EXPLORE RELATIONSHIPS AMONG CATEGORICAL VARIABLES.
# --- [S1.5]: SAVE JOB LISTS & R ENVIRONMENT AS INPUTS FOR DOWNSTREAM SCRIPTS.

#**[S2] COMPARE QUANTITATIVE DATA AMONG GROUPS**
# --- [S2.1]: PREPARE CODING ENVIRONMENT: Load packages, S1 R environment, etc.
# --- [S2.2]: DO PHYLOGENETIC ANOVAs & LEV. TESTs; MAKE ASSOCIATED BOXPLOTs.
# --- -- NOTE: This is repeated for raw, log-trans, and BoxCox-trans datasets.
# --- [S2.3]: MAKE GROUPED BOX PLOTS, TO SHOW HOW TRENDS DIFFER BY SUBCLADE.
# --- [S2.4]: COMPARE LIMB PROPORTIONS IN TERNARY MORPHOSPACE.
# --- [S2.5]: SAVE R ENVIRONMENT AS INPUT FOR SCRIPT_S5.

#**[S3] PERFORM PAIRWISE PHYLOGENETIC CORRELATION TESTS**
# --- [S3.1]: PREPARE CODING ENVIRONMENT: Load pkgs, S1 R environment, etc.
# --- [S3.2]: RUN PAIRWISE CORRELATION TESTS FOR ALL NON-UNITLESS DATA.
# --- -- NOTE: This is repeated for raw, log-trans, and BoxCox-trans datasets.
# --- [S3.3]: SAVE CORRELATION TEST RESULTS & COMPUTATION TIME FOR SCRIPT_S5.

#**[S4] FIT PHYLOGENETIC LOGISTIC REGRESSION MODELS TO DATA**
# --- [S4.1]: PREPARE CODING ENVIRONMENT: Load pkgs, S1 R environment, etc.
# --- [S4.2]: FIT PHYLOGEN. BINOMIAL LOGISTIC REGRESSION MODELS TO DATASETS.
# --- -- NOTE: This is repeated for raw, log-trans, and BoxCox-trans datasets.
# --- [S4.3]: SAVE PHYBLR TEST RESULTS & COMPUTATION TIME FOR SCRIPT_S5.

#**[S5] PERFORM SENSITIVITY TESTS & PREDICT ANCIENT PHENOTYPES**
# --- [S5.1]: PREPARE CODING ENVIRONMENT & STITCH UPSTREAM OUTPUTS.
# --- [S5.2]: RUN SENSITIVITY ANALYSES FOR TRANSFORM METHOD & TREE TOPOLOGY.
# --- [S5.3]: MAKE CORRELATION MATRICES USING PHYCORR RESULTS.
# --- [S5.4]: USE ROC ANALYSIS TO IDENTIFY BEST PHYBLR MODEL(S).
# --- [S5.5]: PREDICT TIP & NODE PROBABILITIES USING BEST PHYBLR MODEL(S).
# --- [S5.6]: SAVE R ENVIRONMENT & COMPUTATION TIME FOR FOR FUTURE REFERENCE.

# In the present script (S1), we will go through each of the S1 steps above in great detail, providing explanations wherever possible. However, before you proceed, please be warned: Taken together, these scripts will save several gigabytes of files to your working directory. Please be prepared to make space as needed. With that warning aside, we can begin!
#------------------------------------------------------------------------------
#**--------------------------------------------------------------------------**

# WHAT FOLLOWS IS THE FULL PIPELINE FOR SCRIPT_S1, SORTED IN SECTIONS LABELED BY THE STEPS ABOVE. EACH MAJOR S1 STEP IS TITLED BELOW, AS ARE INDIVIDUAL "CHUNKS" OF CODE, INTENDED TO COMPLETE A PARTICULAR COMPLEX TASK.

#**"-------------------------------------------------------------------------"*
#**"------------ ### [[S1.1]] PREPARE CODING ENVIRONMENT ### ----------------"*
#**"-------------------------------------------------------------------------"*

#*-----**{ [S1.1.01] See the system we're working with. }*
version # This is the R version and operating system we are using. This is worth noting, as many functions become deprecated or renamed with updates, and may work differently on different operating systems.
environment() # This reports the R environment we are using.
cat("\n","PROGRESS REP: chunk [S1.1.01] complete; starting next chunk...","\n") # Each makeshift 'chunk' of code in this script will end with a "PROGRESS REPORT" output to the console and the slurm .out file. This checkpointing is done to to help localize problems when running the script as a batch job on Yale's HPC cluster.
#*-----**-----*

#*-----**{ [S1.1.01] Load required packages. }*
# NOTE: Optional installation lines are given above each library() command.
# Parallel programming on multiple CPU cores:
 #install.packages('parallel')
library(parallel)
 #install.packages('foreach')
library(foreach)
 #install.packages('doMC')
library(doMC)
 #install.packages('doParallel')
library(doParallel)

# Data object manipulation:
 #library(vctrs) # probably not needed in this version of script; will confirm after troubleshooting on cluster
 #install.packages('forcats')
library(forcats) # for releveling factors with fct_relevel() function
 #install.packages('data.table')
library(data.table) # for converting objects to data tables and matrices
 #install.packages('stringr')
library(stringr) # for parcing through long chr strings (useful to check tree topologies in Nwk format) and using str_detect() function
 #install.packages('plyr')
library(plyr) # useful for in-task parallelization (eg, running function across an object)
 #install.packages('dplyr')
library(dplyr) # useful for in-task parallelization (eg, running function across an object)

# Data visualization and exporting:
 #install.packages('ggplot2')
library(ggplot2) # for effective data visualization with ggplot suite
#install.packages("MESS") 
library(MESS) # for specifying translucent colors as function arguments
 #install.packages('ggrepel')
 #library(ggrepel) # probably not needed in this version of script; will confirm after troubleshooting on cluster
 #install.packages('GGally')
 #library(GGally) # probably not needed in this version of script; will confirm after troubleshooting on cluster
 #install.packages('car')
library(car) # for normal quantile plot construction
 #install.packages('Ternary')
library(Ternary) # for ternary plot construction
 #install.packages('RColorBrewer')
library(RColorBrewer) # for defining fancy color palettes
 #library(grid) # probably not needed in this version of script; will confirm after troubleshooting on cluster
 #install.packages('gridExtra')
library(gridExtra) # useful for generating multi-panel plots and tables
 #install.packages('corrplot')
library(corrplot) # For correlogram construction
 #install.packages('ggimage')
library(ggimage) # For exporting ggplots

# Data analysis:
 #install.packages('devtools')
library(devtools) # required for downloading some packages from github
 #install.packages('BiocManager')
library(BiocManager) # required for downloading some packages from BiocManager
 #install.packages('BlandAltmanLeh')
library(BlandAltmanLeh) # For Bland-Altman analysis
 #install.packages('vcd')
library(vcd) # for Cramer's V and other association stats
 #install.packages('tigerstats') 
library(tigerstats)
 #install.packages('RVAideMemoire')
library(RVAideMemoire) # for Cramer's V, Fisher's Exact Test, PERMANOVAs, and various other statistical tests
 #install.packages('onewaytests')
library(onewaytests) # required in our catPlots function
 #library(rms) # probably not needed in this version of script; will confirm after troubleshooting on cluster
 #library(performance) # contains check_heteroscedasticity(), check_homogeneity(), and check_normality() functions, which might be useful for sensitivity tests; this pkg was previously used to calculate Tjur's Rsq with the r2_tjur() function; since this function is no longer used, the pkg is not currently used in this version of the script.
 #install.packages('MASS')
library(MASS) # used to make Box-Cox transformations with boxcox() function
 #install.packages('pROC')
library(pROC) # contains functions to plot and extract data from ROC curves
 #install.packages('pracma')
library(pracma) # contains diag() and trapz() functions; diag() is useful for making phylogenetic variance-covariance matrices; trapz() is useful for calculating ROC curve AUC

# Phylogenetic comparative methods and tree visualization:
 #install.packages('ape')
library(ape) # for basic tree manipulation and phylogenetic comparative methods
 #devtools::install_github("YuLab-SMU/ggtree") # NOTE: This only works if you update ggplot2 when prompted.
 #BiocManager::install("ggtree")
library(ggtree) # for advanced tree visualization using ggplot2 suite
 #library(paleotree) # contains timePaleoPhy() function which can date trees (but does so unreliably); probably not needed in this version of script; will confirm after troubleshooting on cluster
 #install.packages('strap')
library(strap) # contains DatePhylo() function to time-calibrate trees
 #install.packages('phytools')
library(phytools) # contains phylANOVA() function
 #install.packages('geiger')
library(geiger) # contains aov.phylo() function
#library(phangorn) # probably not needed in this version of script; will confirm after troubleshooting on cluster
#library(phylotools) # probably not needed in this version of script; will confirm after troubleshooting on cluster
 #install.packages('phylolm')
library(phylolm) # contains phyloglm() and phylolm() functions
 #install.packages('vegan')
library(vegan) # contains adonis2(), which should be able to run phyloPERMANOVAs
 #install.packages('castor')
library(castor) # contains asr_independent_contrasts() function
cat("\n","PROGRESS REP: chunk [S1.1.02] complete; starting next chunk...","\n")
#*-----**-----*

#*-----**{ [S1.1.03] Define paths for output files. }*
# Set the current date:
sysdate <- Sys.Date()
current_date <- gsub("-", ".", sysdate)

# Define working directory path:
wd_path <- "/gpfs/gibbs/project/bhullar/cmg89/Flipper_Project/"
setwd(wd_path); getwd()

# Create subdirectory for current run of the script (all output files will be saved to this path):
output_path <- paste(wd_path, current_date, sep="") 

# Create directory for output files:
dir.create(path=output_path, showWarnings=TRUE, recursive=FALSE, mode="0777") 
cat("\n","PROGRESS REP: chunk [S1.1.03] complete; starting next chunk...","\n")
#*-----**-----*

#*-----**{ [S1.1.04] Set up parallel programming environment. }*
# We will use the foreach package to run programs in parallel on multiple CPU cores. The registerDoMC() function below assigns the proper number of cores for this task. Before you begin running the code below, set the CPU_num available to you on your personal computer or HPC cluster.

CPU_num <- 4 # 6 is the total number of available processing cores on my MacBook. 48 is is the highest number that can be requested from Yale's High-Performance Computing cluster. 4 is the total number that can be requested interactively from Yale's HPC via Open On-Demand.

# Set up parallel backend:
cl <- makeCluster(CPU_num, outfile = "") # outfile="" ensures that output prints to the console
registerDoParallel(cl)
cat("\n","PROGRESS REP: chunk [S1.1.04] complete; starting next chunk...","\n")
#*-----**-----*

#*-----**{ [S1.1.05] Import linear morphometric and Bland-Altman data. }*
input_csv_name <- paste0("_LM_all_data_", current_date, ".csv")
all_data_raw <- read.csv(input_csv_name)
rownames(all_data_raw) <- all_data_raw$Tip_label
BlandAltman_data <- read.csv(paste0("_BlandAltman_data_",current_date,".csv"))
cat("\n","PROGRESS REP: chunk [S1.1.05] complete; starting next chunk...","\n")
#*-----**-----*

#*-----**{ [S1.1.06] Clean the data set. }*
# Remove unnecessary columns:
names(all_data_raw)
all_data <- all_data_raw[, c(3:5,8:11,13,15:18,20,22:25,36:54,57:78,80:89)]
  
# Confirm that the correct columns remain:
dim(all_data_raw); dim(all_data); names(all_data)

# Write function to reorder factor levels. Writing this as a function allows us to relevel all required factors within other functions, which avoids certain issues that come up if one wants to convert this file into a knittable Rmd.
relevel_factors <- function(dataset){
  dataset$Flipper_status <- fct_relevel(dataset$Flipper_status, "Flippered", "Non-flippered", "Ambiguous")
  dataset$Ecotype1 <- fct_relevel(dataset$Ecotype1, "Terrestrial", "Transiently Aquatic", "Moderately Aquatic", "Highly Aquatic", "Fully Aquatic", "Data Deficient")
  dataset$Ecotype2 <- fct_relevel(dataset$Ecotype2, "almost always on land", "primarily on land, seldom in water", "primarily on land, often in water", "small or slow-moving water bodies", "large or fast-moving water bodies", "data deficient")
  dataset$Ecotype3 <- fct_relevel(dataset$Ecotype3, "predominantly terrestrial", "terrestrial/freshwater", "terrestrial/euryhaline", "terrestrial/marine", "predominantly freshwater", "freshwater/marine", "predominantly marine", "data deficient")
  dataset$fore_type <- fct_relevel(dataset$fore_type, "Unwebbed", "Webbed", "Flippered", "Unclear, terr.", "Unclear, aquat.")
  dataset$hind_type <- fct_relevel(dataset$hind_type, "Unwebbed", "Webbed", "Flippered", "Unclear, terr.", "Unclear, aquat.")
  dataset$f_subtype <- fct_relevel(dataset$f_subtype, "Unwebbed", "Interdigital Girth", "Stubbed", "Pincer-palmed", "Interdigital Integument", "Partial, slight", "Partial, extensive", "Complete, slight", "Complete, extensive", "Lobate", "Weakly Flippered", "Strongly Flippered", "Unclear, terr.", "Unclear, aquat.")
  dataset$h_subtype <- fct_relevel(dataset$h_subtype, "Unwebbed", "Interdigital Girth", "Stubbed", "Pincer-palmed", "Interdigital Integument", "Partial, slight", "Partial, extensive", "Complete, slight", "Complete, extensive", "Lobate", "Weakly Flippered", "Strongly Flippered", "Unclear, terr.", "Unclear, aquat.")
  dataset$IUCN_cat <- fct_relevel(dataset$IUCN_cat, "DD", "LC", "NT", "VU", "EN", "CR", "EX") 
  dataset$IUCN_cat <- droplevels(dataset$IUCN_cat)
  dataset$MajorClade <- fct_relevel(dataset$MajorClade, "Pan-Reptilia", "Pan-Mammalia", "Outgroups")
  dataset$Subclade <- fct_relevel(dataset$Subclade, "Outgroups", "Stem reptiles", "Euryapsida", "Lepidosauromorpha", "Testudines", "Archosauromorpha", "Non-placentals", "Atlantogenata", "Euarchontoglires", "Eulipotyphla", "Ferae", "Ungulata")
  dataset$Subgroup1 <- fct_relevel(dataset$Subgroup1, "Lissamphibia", "Monotremata", "Marsupialia", "Xenarthra", "Afrotheria", "Primatomorpha", "Rodentia", "Mesonychia", "Perissodactyla", "Artiodactyla", "Permian aquatic stem reptiles", "Thalattosauria", "Ichthyosauromorpha", "Sauropterygiformes", "Thalassochelydia", "Pleurodira", "Cryptodira", "Stem Archosaurs", "Pan-Crocodylia", "Dinosauria", "Rhynchocephalia", "Iguania", "Gekkonomorpha", "Lacertoidea", "Xenosauridae", "Anguimorpha")
  dataset$Subgroup2 <- fct_relevel(dataset$Subgroup2, "Agamidae")
  dataset$Limb_region_measurement_method <- fct_relevel(dataset$Limb_region_measurement_method, "ImageJ", "Slicer", "Calipers")
  return(dataset)
} # closes relevel_factors() function

# Apply relevel_factors function to whole dataset:
all_data <- relevel_factors(dataset=all_data)

# Check that the relevel_factors() function works:
levels(all_data$Ecotype1)
levels(all_data$f_subtype)
  
# Make sure all quantitative variables are numeric: 
names(all_data)
all_data[ , c(21:68)] <- apply(all_data[ , c(21:68)], 2, function(x) as.numeric(as.character(x)))
str(all_data)
head(all_data)

# Optional extra troubleshooting steps: check for missing data:
  # dim(all_data); for(i in 1:length(colnames(all_data))){print(sum(is.na(all_data[, i])))} 
      # No columns should have all (eg, 714) rows of missing data.
cat("\n","PROGRESS REP: chunk [S1.1.06] complete; starting next chunk...","\n")
#*-----**-----*

#*-----**{ [S1.1.07] Define object name vectors to automate downstream work. }*
# In a later section of our script, we will write functions that analyze a given response variable across levels of a given grouping variable within a given dataset for a given tree. To automate this process for each of our functions, we first create data objects (vectors or lists) that will let us run each function on every combination of trees, datasets, response_vars and grouping_vars. We make these objects in the chunks below.

# Vector naming all grouping variables:
grouping_vars <- c("Subclade", "Ecotype1", "Ecotype2", "Ecotype3", "fore_type", "hind_type", "f_subtype", "h_subtype") # Note: "MajorClade", "IUCN_cat", and "Limb_region_measurement_method" were omitted from this section during an earlier draft of the script to save computational time, because they weren't informative.

# Vector naming all quantitative variables:
all_quantitative_vars <- names(all_data[c(21:68)])

# Vector naming all computed ratios found to be potentially informative in previous run:
ratio_vars <- c("f_AZR", "f_SZR", "f_ASR", "f_AnAR", "h_AZR", "h_SZR", "h_ASR", "h_AnAR", "mD_Symmetry_index1", "mD_Symmetry_index2", "pD_Symmetry_index1", "pD_Symmetry_index2", "mD3_up_FI1", "mD3_up_FI2", "mD3_up_FI3", "pD3_up_FI1", "pD3_up_FI2", "pD3_up_FI3", "mD3_proxp_Length.DistWidth", "mD3_proxp_WidthRatios", "pD3_proxp_Length.DistWidth", "pD3_proxp_WidthRatios")

# Vector naming all direct length measurement variables:
direct_measurement_variables <- c("Humerus", "Radius", "Ulna", "Hand", "Femur", "Tibia", "Foot", "mD_Antmost", "mD_Postmost", "mD_Longest", "pD_Antmost", "pD_Postmost", "pD_Longest", "mD3_up_Height", "mD3_up_TopLength", "mD3_up_BottomLength", "pD3_up_Height", "pD3_up_TopLength", "pD3_up_BottomLength", "mD3_proxp_Length", "mD3_proxp_ProxWidth", "mD3_proxp_DistWidth", "pD3_proxp_Length", "pD3_proxp_ProxWidth", "pD3_proxp_DistWidth")
cat("\n","PROGRESS REP: chunk [S1.1.07] complete; starting next chunk...","\n")
#*-----**-----*

#*-----**{ [S1.1.08] Parse cleaned dataset for downstream analysis. }*
# Finalize "base" dataset from which others will be derived:
  rownames(all_data) <- all_data$Tip_label

# Parse data by major clade and measurement unit:
  all_mm <- all_data[all_data$Length_units == "mm", ]
  all_mm <- all_mm[which(is.na(all_mm$Subclade)==F), ] # Note: this second "which" line removes any "NA" rownames introduced artificially by the parsing process. In several of the taxonomic data sets I parse out below, many "NA" rows spontaneously appear unless I include this line. I have added it once for every parsed dataset in this chunk just to be safe and ensure no extra rows are created for no reason.
  reps_all <- all_data[all_data$MajorClade=="Pan-Reptilia", ]
  reps_all <- reps_all[which(is.na(reps_all$Subclade)==F), ]
  mams_all <- all_data[all_data$MajorClade=="Pan-Mammalia", ]
  mams_all <- mams_all[which(is.na(mams_all$Subclade)==F), ]
  reps_mm <- reps_all[reps_all$Length_units == "mm", ]
  reps_mm <- reps_mm[which(is.na(reps_mm$Subclade)==F), ]
  mams_mm <- mams_all[mams_all$Length_units == "mm", ]
  mams_mm <- mams_mm[which(is.na(mams_mm$Subclade)==F), ]
  
# Parse data by smaller taxonomic groupings:
  euryaps_all <- all_data[all_data$Subclade=="Euryapsida", ]
  euryaps_all <- euryaps_all[which(is.na(euryaps_all$Subclade)==F), ]
  ichthyos_all <- all_data[all_data$Subgroup1=="Ichthyosauromorpha", ]
  ichthyos_all <- ichthyos_all[which(is.na(ichthyos_all$Subclade)==F), ]
  sauropter_all <- all_data[all_data$Subgroup1=="Sauropterygiformes", ]
  sauropter_all <- sauropter_all[which(is.na(sauropter_all$Subclade)==F), ]
  turts_all <- all_data[all_data$Subclade=="Testudines", ]
  turts_all <- turts_all[which(is.na(turts_all$Subclade)==F), ]
  archos_all <- all_data[all_data$Subclade=="Archosauromorpha", ]
  archos_all <- archos_all[which(is.na(archos_all$Subclade)==F), ]
  all_BUT_archos <- all_data[!(all_data$Subclade=="Archosauromorpha"), ]
  all_BUT_archos <- all_BUT_archos[which(is.na(all_BUT_archos$Subclade)==F), ]
  dinos_all <- all_data[all_data$Subgroup1=="Dinosauria", ]
  dinos_all <- dinos_all[which(is.na(dinos_all$Subclade)==F), ]
  all_BUT_dinos <- all_data[!(all_data$Subgroup1=="Dinosauria"), ]
  all_BUT_dinos <- all_BUT_dinos[which(is.na(all_BUT_dinos$Subclade)==F), ]
  crocs_all <- all_data[all_data$Subgroup1=="Pan-Crocodylia", ]
  crocs_all <- crocs_all[which(is.na(crocs_all$Subclade)==F), ]
  lepidos_all <- all_data[all_data$Subclade=="Lepidosauromorpha", ]
  lepidos_all <- lepidos_all[which(is.na(lepidos_all$Subclade)==F), ]
  rhyncoceph_all <- all_data[all_data$Subgroup1=="Rhynchocephalia", ]
  rhyncoceph_all <- rhyncoceph_all[which(is.na(rhyncoceph_all$Subclade)==F), ]
  anguimorph_all <- all_data[all_data$Subgroup1=="Anguimorpha", ]
  anguimorph_all <- anguimorph_all[which(is.na(anguimorph_all$Subclade)==F), ]
  iguani_all <- all_data[all_data$Subgroup1=="Iguania", ]
  iguani_all <- iguani_all[which(is.na(iguani_all$Subclade)==F), ]
  agamid_all <- all_data[all_data$Subgroup2=="Agamidae", ]
  agamid_all <- agamid_all[which(is.na(agamid_all$Subclade)==F), ]
  stemreps_all <- all_data[all_data$Subclade=="Stem reptiles", ]
  stemreps_all <- stemreps_all[which(is.na(stemreps_all$Subclade)==F), ]
  ungulates_all <- all_data[all_data$Subclade=="Ungulata", ]
  ungulates_all <- ungulates_all[which(is.na(ungulates_all$Subclade)==F), ]
  nonplacs_all <- all_data[all_data$Subclade=="Non-placentals", ]
  nonplacs_all <- nonplacs_all[which(is.na(nonplacs_all$Subclade)==F), ]
  atlanto_all <- all_data[all_data$Subclade=="Atlantogenata", ]
  atlanto_all <- atlanto_all[which(is.na(atlanto_all$Subclade)==F), ]
  ferae_all <- all_data[all_data$Subclade=="Ferae", ]
  ferae_all <- ferae_all[which(is.na(ferae_all$Subclade)==F), ]
  euarchontagl_all <- all_data[all_data$Subclade=="Euarchontoglires", ]
  euarchontagl_all <- euarchontagl_all[which(is.na(euarchontagl_all$Subclade)==F), ]
    
# Define dataset list and name vector to automate downstream analyses:
  orig_datasets <- list(all_data, turts_all, archos_all, all_BUT_archos, lepidos_all, ungulates_all, atlanto_all, ferae_all) # Note: Additional datasets (e.g., all_mm, reps_all, mams_all, anguimorph_all, dinos_all, all_BUT_dinos, crocs_all, agamid_all, iguani_all, rynchoceph_all, euarchontagl_all) were included in this list in previous versions of the script but omitted because they were less pertinent to our current investigation and added unnecessary extra computational time. Fossil-only datasets (e.g., euryaps_all, ichthyos_all, sauropter_all) aren't informative for some of the downstream functions below, as all or most of their limb type and ecotype values are "data deficient" or "unclear"). Thus, we restricted the pipeline below (and the orig_datasets list above) to include only those datasets with at least some living members. Ideally, given greater computational resources, I would include reps_all, mams_all, anguimorph_all, dinos_all, all_BUT_dinos, crocs_all, agamid_all, and iguani_all in this list as well, but they can be input into orig_datasets separately during a future run.
  orig_dataset_names <- c("all_data", "turts_all", "archos_all", "all_BUT_archos", "lepidos_all", "ungulates_all", "atlanto_all", "ferae_all")
  names(orig_datasets) <- orig_dataset_names
cat("\n","PROGRESS REP: chunk [S1.1.08] complete; starting next chunk...","\n")
#*-----**-----*
  
#*-----**{ [S1.1.09] Write functions to transform datasets. }*
# The statistical tests we will be performing (a simulation-based phylogenetic ANOVA, phylogenetic Levene's test, and phylogenetic correlation tests) should not make strict assumptions about the normality or variance-homogeneity of our data. However, maximizing normality and homoskedasticity can make some statistical tests more reliable, and it is common for paleontologists to log-transform certain kinds of morphometric data prior to phylogenetic statistical analysis (e.g., Soul & Benson 2017 [https://doi.org/10.1111/evo.13217]; Simoes et al. 2022 [https://science.org/doi/10.1126/ sciadv.abq1898]). The Box-Cox transformation method, first developed by Box & Cox in 1964 (https://doi.org/10.1080/01621459.1982.10477788), is a power-transform method that tends to maximize the normality of the dataset in a wider range of cases than logarithmic or other transformations. It also tends to reduce heteroskedasticity. However, the actual impact of log- and BoxCox-transformations on the results of statistical tests remains poorly studied. To address this uncertainty, we sought to perform all of our phylogenetic statistical tests three times--once on our raw data, once following log10-transformation, and once following BoxCox-transformation. In script_S5, we perform a set of sensitivity tests to investigate whether these transformation protocols alter our phylogenetic comparative test results in a significant percentage of cases. In doing so, we hope to shed some light on whether and to what extent the homoskedasticity and normality of the raw data makes a difference for simulation-based nonparametric phylogenetic comparative tests.

# To perform these transformations, we first write functions that allow us to BoxCox-transform and log10-transform a given dataset's numeric variables of interest. We define these functions--BOXCOX.trans() and log10.trans()--below.

# Write function to BoxCox-transform the data:
BOXCOX.trans <- function(dataset, dataset_name, vars_to_transform){
  # Defining directory path for output files within the function: 
  directory_path <- paste0(output_path, "/BoxCox_plots")
  # Creating directory for output files:
  dir.create(path=directory_path, showWarnings = TRUE, recursive = FALSE, mode = "0777")
  ds_trans <- dataset
  
  # Initialize connection to output file:
  png(filename=paste(directory_path, "/", dataset_name, "__", "BoxCox_plots", ".png", sep=""), width=12, height=12, units="in", res=400)
  # Generate Box-Cox power transform plots:
  par(mfrow=c(8,7))
  for(i in 1:length(vars_to_transform)){
    if(sum(is.na(ds_trans[ , vars_to_transform[i]])) < (length(ds_trans[ , vars_to_transform[i]])-2)){
        hist(ds_trans[ , vars_to_transform[i]], main=paste("hist for,",dataset_name,"|",vars_to_transform[i], "untransformed", sep=" "), cex.main=0.3, cex.lab=0.5, xlab = paste(vars_to_transform[i], ", raw values", sep=""))
        y <- ds_trans[ , vars_to_transform[i]]
        #cat("\n", "Transforming", vars_to_transform[i], "...", "\n")
        bc <- boxcox(y ~ 1)
        lambda <- bc$x[which.max(bc$y)]
        y_transformed <- (y^lambda - 1) / lambda
        hist(y_transformed, main=paste("hist for,", dataset_name, "|",vars_to_transform[i],
"BOX-COX-transformed", sep=" "), cex.main=0.3, cex.lab=0.5)
        ds_trans[ , vars_to_transform[i]] <- y_transformed
        cat("\n", dataset_name,"|",vars_to_transform[i], "transformation complete.", "\n")
    } else if(sum(is.na(ds_trans[ , vars_to_transform[i]])) >= (length(ds_trans[ , vars_to_transform[i]])-2)){
        cat("\n", "Transformation not applicable for", dataset_name, "|", vars_to_transform[i], "as it contains only NAs.")}
  }
  dev.off()
  return(ds_trans)
} # closes BOXCOX.trans() function

# We just wrote a function to Box-Cox transform our datasets. We will now write a function to log10-transform our data, for the same internal test to see whether homoskedasticity and normality of the raw data make a difference for our downstream phylogenetic statistical tests.

# Write function to log10-transform the data:
log10.trans <- function(dataset, dataset_name, vars_to_transform){
  # Defining directory path for output files within the function: 
  directory_path <- paste0(output_path, "/log10_plots")
  # Creating directory for output files:
  dir.create(path=directory_path, showWarnings = TRUE, recursive = FALSE, mode = "0777")
  ds_trans <- dataset
  
  # Initialize connection to output file:
  png(filename=paste(directory_path, "/", dataset_name, "__", "log10_plots", ".png", sep=""), width=12, height=12, units="in", res=400)
  # Generate Log10 transformation plots:
  par(mfrow=c(8,7))
  for(i in 1:length(vars_to_transform)){
    if(sum(is.na(ds_trans[ , vars_to_transform[i]])) < (length(ds_trans[ , vars_to_transform[i]])-2)){
      hist(ds_trans[ , vars_to_transform[i]], main=paste("hist for,",dataset_name,"|",vars_to_transform[i], "untransformed", sep=" "), cex.main=0.3, cex.lab=0.5, xlab = paste(vars_to_transform[i], ", raw values", sep=""))
      y <- ds_trans[ , vars_to_transform[i]]
      #cat("\n", "Transforming", vars_to_transform[i], "...", "\n")
      y_transformed <- log10(y)
      hist(y_transformed, main=paste("hist for,", dataset_name, "|",vars_to_transform[i],"log10-transformed", sep=" "), cex.main=0.3, cex.lab=0.5)
      ds_trans[ , vars_to_transform[i]] <- y_transformed
      cat("\n", dataset_name,"|",vars_to_transform[i], "transformation complete.", "\n")
    } else if(sum(is.na(ds_trans[ , vars_to_transform[i]])) >= (length(ds_trans[ , vars_to_transform[i]])-2)){
      cat("\n", "Transformation not applicable for", dataset_name, "|", vars_to_transform[i], "as it contains only NAs.")}
  }
  dev.off()
  return(ds_trans)
} # closes log10.trans() function

# Check that BOXCOX.trans() and log10.trans functions work on full dataset:
  all_trans <- BOXCOX.trans(dataset=all_data, dataset_name="all_data", vars_to_transform=all_quantitative_vars)
  all_trans <- log10.trans(dataset=all_data, dataset_name="all_data", vars_to_transform=all_quantitative_vars)
cat("\n","PROGRESS REP: chunk [S1.1.09] complete; starting next chunk...","\n")
#*-----**-----*
  
#*-----**{ [S1.1.10] Define function to make categorical variables binary. }*
# We will eventually be performing phylogenetic binomial logistic regression (phybLR), which requires a binary response variable. The bLRify() function below rebins all of our existing grouping variables into binary variables. The output for this function is a binarified dataset ready for phybLR.

# Below, we define a function to make "bLR" (binary Logistic Regression-ready) datasets with binary classification schemes for downstream categorical association tests (S1 catPlots, later in this script), phylogenetic comparative tests (S2 script), and binary logistic regression analyses (S4 scripts).

bLRify <- function(dataset, dataset_name, verbose=T){
  ds <- dataset
  Tip_label <- ds$Tip_label
  Taxon <- ds$Taxon
  Subclade <- ds$Subclade
  Subgroup1 <- ds$Subgroup1
  Subgroup2 <- ds$Subgroup2
  Ecotype1 <- ds$Ecotype1
  Ecotype2 <- ds$Ecotype2
  fore_type <- ds$fore_type
  hind_type <- ds$hind_type
  f_subtype <- ds$f_subtype
  h_subtype <- ds$h_subtype
  predictor_vars <- all_quantitative_vars
    ds$Limb_region_measurement_method <- fct_relevel(ds$Limb_region_measurement_method, "ImageJ", "Slicer", "Calipers")
    bin_Method <- gsub("Calipers|Slicer", "3D", ds$Limb_region_measurement_method)
    bin_Method <- gsub("ImageJ", "2D", bin_Method)
    bin_Method <- fct_relevel(bin_Method, "2D", "3D")
    
    if(verbose==T){
      cat("\n", "Rebinned levels(bin_Method):", "\n")
      print(levels(droplevels(bin_Method)))}
    bin_specmType <- gsub("Fossil,.*", "Fossil", ds$Specimen_type)
    bin_specmType <- gsub("Recent.*", "Recent", bin_specmType)
    bin_specmType <- fct_relevel(bin_specmType, "Fossil", "Recent")
    
    if(verbose==T){
      cat("\n", "Rebinned levels(bin_specmType):", "\n")
      print(levels(droplevels(bin_specmType)))}
    bin_eco1_v1 <- gsub("Moderately Aquatic|Highly Aquatic|Fully Aquatic", "Aquatic or Semi-Aquatic", Ecotype1)
    bin_eco1_v1 <- gsub("Transiently Aquatic", "Terrestrial", bin_eco1_v1)
    bin_eco1_v1 <- gsub("Data Deficient", NA, bin_eco1_v1)
    bin_eco1_v1 <- fct_relevel(bin_eco1_v1, "Terrestrial", "Aquatic or Semi-Aquatic")
    
    if(verbose==T){
      cat("\n", "Rebinned levels(bin_eco1_v1):", "\n")
      print(levels(droplevels(bin_eco1_v1)))}
    
    bin_eco1_v2 <- gsub("Terrestrial|Transiently Aquatic|Moderately Aquatic|Highly Aquatic", "NOT Fully Aquatic", Ecotype1)
    bin_eco1_v2 <- gsub("Data Deficient", NA, bin_eco1_v2)
    bin_eco1_v2 <- fct_relevel(bin_eco1_v2, "NOT Fully Aquatic", "Fully Aquatic")
      
    if(verbose==T){
      cat("\n", "Rebinned levels(bin_eco1_v2):", "\n")
      print(levels(droplevels(bin_eco1_v2)))}
    
    bin_eco1_v3 <- gsub("Terrestrial|Transiently Aquatic|Moderately Aquatic", "NOT highly or fully aquatic", Ecotype1)
    bin_eco1_v3 <- gsub("Highly Aquatic|Fully Aquatic", "Highly or fully aquatic", bin_eco1_v3)
    bin_eco1_v3 <- gsub("Data Deficient", NA, bin_eco1_v3)
    bin_eco1_v3 <- fct_relevel(bin_eco1_v3, "NOT highly or fully aquatic", "Highly or fully aquatic")
    
    if(verbose==T){
      cat("\n", "Rebinned levels(bin_eco1_v3):", "\n")
    print(levels(droplevels(bin_eco1_v3)))}
    
    bin_eco1_v4 <- gsub("Transiently Aquatic|Moderately Aquatic", "Transiently or moderately aquatic", Ecotype1)
    bin_eco1_v4 <- gsub("Highly Aquatic|Fully Aquatic|Data Deficient", NA, bin_eco1_v4)
    bin_eco1_v4 <- fct_relevel(bin_eco1_v4, "Terrestrial", "Transiently or moderately aquatic")
    
    if(verbose==T){
      cat("\n", "Rebinned levels(bin_eco1_v4):", "\n")
      print(levels(droplevels(bin_eco1_v4)))}
    bin_eco1_v5 <- gsub("Transiently Aquatic|Highly Aquatic|Fully Aquatic|Data Deficient", NA, Ecotype1)
    bin_eco1_v5 <- fct_relevel(bin_eco1_v5, "Terrestrial","Moderately Aquatic")
    
    if(verbose==T){
      cat("\n", "Rebinned levels(bin_eco1_v5):", "\n")
      print(levels(droplevels(bin_eco1_v5)))}
  
    bin_eco1_v6 <- gsub("Transiently Aquatic|Moderately Aquatic|Fully Aquatic|Data Deficient", NA, Ecotype1)
    bin_eco1_v6 <- fct_relevel(bin_eco1_v6, "Terrestrial", "Highly Aquatic")
    
    if(verbose==T){
      cat("\n", "Rebinned levels(bin_eco1_v6):", "\n")
      print(levels(droplevels(bin_eco1_v6)))}
    bin_eco1_v7 <- gsub("Terrestrial|Transiently Aquatic|Fully Aquatic|Data Deficient", NA, Ecotype1)
    bin_eco1_v7 <- fct_relevel(bin_eco1_v7, "Moderately Aquatic", "Highly Aquatic")
    
    if(verbose==T){
      cat("\n", "Rebinned levels(bin_eco1_v7):", "\n")
      print(levels(droplevels(bin_eco1_v7)))}
    bin_eco1_v8 <- gsub("Terrestrial|Transiently Aquatic|Data Deficient", NA, Ecotype1)
    bin_eco1_v8 <- gsub("Highly Aquatic|Fully Aquatic", "Highly or fully aquatic", bin_eco1_v8)
    bin_eco1_v8 <- fct_relevel(bin_eco1_v8, "Moderately Aquatic", "Highly or fully aquatic")
    
    if(verbose==T){
      cat("\n", "Rebinned levels(bin_eco1_v8):", "\n")
      print(levels(droplevels(bin_eco1_v8)))}
    bin_eco1_v9 <- gsub("Terrestrial|Data Deficient", NA, Ecotype1)
    bin_eco1_v9 <- gsub("Transiently Aquatic|Moderately Aquatic", "Transiently or moderately aquatic", bin_eco1_v9)
    bin_eco1_v9 <- gsub("Highly Aquatic|Fully Aquatic", "Highly or fully aquatic", bin_eco1_v9)
    bin_eco1_v9 <- fct_relevel(bin_eco1_v9, "Transiently or moderately aquatic", "Highly or fully aquatic")
    
    if(verbose==T){
      cat("\n", "Rebinned levels(bin_eco1_v9):", "\n")
      print(levels(droplevels(bin_eco1_v9)))}
    
    for(i in 1:length(Subgroup2)){
      if(is.na(Subgroup2[i])==F & Subgroup2[i]=="Thalattosuchia"){
            bin_eco1_v3[i] <- "Highly or fully aquatic"
            bin_eco1_v8[i] <- "Highly or fully aquatic"
            bin_eco1_v9[i] <- "Highly or fully aquatic"
            }} # This for-loop scores thalattosuchians as being "highly or fully aquatic"; because they were marked as "Unclear, aquat." in the original dataset, they would otherwise be overlooked by the rebinning code below.
  
    bin_eco2 <- gsub("data deficient", NA, ds$Ecotype2)
    bin_eco2 <- gsub("large or fast-moving water bodies", "Lives in deep/fast-moving waters", bin_eco2)
    bin_eco2 <- gsub("almost always on land", "Lives on land or in shallow/slow-moving waters", bin_eco2)
    bin_eco2 <- gsub("primarily on land, seldom in water", "Lives on land or in shallow/slow-moving waters", bin_eco2)
    bin_eco2 <- gsub("primarily on land, often in water", "Lives on land or in shallow/slow-moving waters", bin_eco2)
    bin_eco2 <- gsub("small or slow-moving water bodies", "Lives on land or in shallow/slow-moving waters", bin_eco2)
    bin_eco2 <- fct_relevel(bin_eco2, "Lives on land or in shallow/slow-moving waters", "Lives in deep/fast-moving waters")
    
    if(verbose==T){
      cat("\n", "Rebinned levels(bin_eco2):", "\n")
      print(levels(droplevels(bin_eco2)))}
  bin_fflip <- gsub("Unwebbed|Webbed", "Non-flippered", ds$fore_type)
  bin_fflip <- gsub("Unclear, aquat.|Unclear, terr.", NA, bin_fflip)
  bin_fflip <- fct_relevel(bin_fflip, "Non-flippered", "Flippered")
  
  if(verbose==T){
    cat("\n", "Rebinned levels(bin_fflip):", "\n")
    print(levels(droplevels(bin_fflip)))}
  bin_hflip <- gsub("Unwebbed|Webbed", "Non-flippered", ds$hind_type)
  bin_hflip <- gsub("Unclear, aquat.|Unclear, terr.", NA, bin_hflip)
  bin_hflip <- fct_relevel(bin_hflip, "Non-flippered", "Flippered")
  
  if(verbose==T){
    cat("\n", "Rebinned levels(bin_hflip):", "\n")
    print(levels(droplevels(bin_hflip)))}
    bin_fweb <- gsub("Unclear, aquat.|Unclear, terr.|Flippered", NA, ds$fore_type)
    bin_fweb <- fct_relevel(bin_fweb, "Unwebbed", "Webbed")
    
    if(verbose==T){
      cat("\n", "Rebinned levels(bin_fweb):", "\n")
      print(levels(droplevels(bin_fweb)))}
    bin_hweb <- gsub("Unclear, aquat.|Unclear, terr.|Flippered", NA, ds$hind_type)
    bin_hweb <- fct_relevel(bin_hweb, "Unwebbed", "Webbed")
    
    if(verbose==T){
      cat("\n", "Rebinned levels(bin_hweb):", "\n")
      print(levels(droplevels(bin_hweb)))}
   
  # Combine binarified grouping_var vectors into a single object:  
  bLR_set <- cbind(Tip_label, Taxon, Subclade, Subgroup1, Subgroup2, Ecotype1, Ecotype2, fore_type, hind_type, f_subtype, h_subtype, bin_specmType, bin_Method, bin_fweb, bin_fflip, bin_hweb, bin_hflip, bin_eco1_v1, bin_eco1_v2, bin_eco1_v3, bin_eco1_v4, bin_eco1_v5, bin_eco1_v6, bin_eco1_v7, bin_eco1_v8, bin_eco1_v9, bin_eco2, ds[, c(predictor_vars)])
  
  # Many of the extinct taxa in our dataset (and even some of the extant taxa for which the existing natural history is sparse) have unclear aquatic habits when scored within a five-level scoring system of aquatic affinity, but have reasonably resolved aquatic habits under certain binary scoring systems. In particular, many extinct archosauromorphs in our dataset fall into this category. The code below corrects their scores, so that these taxa get ecotype scores that are correctly binarified.
  for(r in 1:nrow(bLR_set)){
    if(bLR_set[r, "Taxon"]=="Civetticus_civetta" | bLR_set[r, "Taxon"]=="Procyon_lotor" | bLR_set[r, "Taxon"]=="Phataginus_tricuspis" | bLR_set[r, "Taxon"]=="Phataginus_tetradactyla" | bLR_set[r, "Taxon"]=="Tragulus_javanicus" | bLR_set[r, "Taxon"]=="Camelid_sp" | bLR_set[r, "Taxon"]=="Tapirus_indicus" | bLR_set[r, "Taxon"]=="Otolemur_crassicaudatus" | bLR_set[r, "Taxon"]=="Galagoides_demidoff_anomurus" |  bLR_set[r, "Taxon"]=="Pan_paniscus" | bLR_set[r, "Taxon"]=="Cavia_sp" |  bLR_set[r, "Taxon"]=="Echimys_saturnus" | bLR_set[r, "Taxon"]=="Neotoma_floridana" | bLR_set[r, "Taxon"]=="Sciurumimus_albersdoerferi" | bLR_set[r, "Taxon"]=="Heterodontosaurus_sp" | bLR_set[r, "Taxon"]=="Camptosaurus_dispar" | bLR_set[r, "Taxon"]=="Archaeopteryx_lithographica" | bLR_set[r, "Taxon"]=="Chrysolophus_pictus"| bLR_set[r, "Taxon"]== "Bubo_virginianus"| bLR_set[r, "Taxon"]=="Poposaurus_gracilis"| bLR_set[r, "Taxon"]=="Coelophysis_bauri"| bLR_set[r, "Taxon"]=="Popobuddy"| bLR_set[r, "Taxon"]=="Tanycolagreus_topwilsoni"| bLR_set[r, "Taxon"]=="Compsognathus_longipes"| bLR_set[r, "Taxon"]=="Struthiomimus_sedens"| bLR_set[r, "Taxon"]=="Deinonychus_antirrhopus"| bLR_set[r, "Taxon"]=="Mononykus_olecranus"| bLR_set[r, "Taxon"]=="Allosaurus_jimmadseni"| bLR_set[r, "Taxon"]=="Confuciusornis_dui"| bLR_set[r, "Taxon"]=="Eoraptor_lunensis"| bLR_set[r, "Taxon"]=="Mesosuchus_browni"| bLR_set[r, "Taxon"]=="Neoaetosauroides_engaeus"| bLR_set[r, "Taxon"]=="Ornithomimus_edmontonicus"| bLR_set[r, "Taxon"]=="Ornithomimus_sp"| bLR_set[r, "Taxon"]=="Prolacerta_broomi"| bLR_set[r, "Taxon"]=="Protorosaurus_speneri"| bLR_set[r, "Taxon"]=="Sinocalliopteryx_gigas"| bLR_set[r, "Taxon"]=="Sinosauropteryx_prima"| bLR_set[r, "Taxon"]=="Struthiomimus_sp"| bLR_set[r, "Taxon"]=="Oviraptor_philoceratops"){
      bLR_set[r, "bin_fflip"] <- "Non-flippered"
      bLR_set[r, "bin_hflip"] <- "Non-flippered"
      bLR_set[r, "bin_eco1_v1"] <- "Terrestrial"
      bLR_set[r, "bin_eco1_v2"] <- "NOT Fully Aquatic"
      bLR_set[r, "bin_eco1_v3"] <- "NOT highly or fully aquatic"
    }
    if(bLR_set[r, "Taxon"]=="Steneosaurus_bollensis"){
      bLR_set[r, "bin_eco1_v1"] <- "Aquatic or Semi-Aquatic"
    }
    if(bLR_set[r, "Taxon"]=="Dakosaurus_maximus" | bLR_set[r, "Taxon"]=="Geosaurus_sp"){
      bLR_set[r, "bin_eco1_v1"] <- "Aquatic or Semi-Aquatic"
      bLR_set[r, "bin_eco1_v3"] <- "Highly or fully aquatic"
      bLR_set[r, "bin_eco1_v8"] <- "Highly or fully aquatic"
      bLR_set[r, "bin_eco1_v9"] <- "Highly or fully aquatic"
    }
    # The following taxa are mammals whose webbing phenotype we could not verify sufficiently with the zoological specimens, but which we know lack flippers: 
    if(bLR_set[r, "Taxon"]=="Anomalurus_derbianus" | bLR_set[r, "Taxon"]=="Octodon_degus" | bLR_set[r, "Taxon"]=="Dinomys_branickii" | bLR_set[r, "Taxon"]=="Ptilocercus_lowii" | bLR_set[r, "Taxon"]=="Cercopithecus_sp" | bLR_set[r, "Taxon"]=="Loris_tardigradus" | bLR_set[r, "Taxon"]=="Galago_senegalensis" | bLR_set[r, "Taxon"]=="Bison_americanus" | bLR_set[r, "Taxon"]=="Bassaricyon_alleni" | bLR_set[r, "Taxon"]=="Taxidea_taxus" | bLR_set[r, "Taxon"]=="Myrmecophaga_tridactyla" | bLR_set[r, "Taxon"]=="Cyclopes_didactylus" | bLR_set[r, "Taxon"]=="Zaedyus_pichiy" | bLR_set[r, "Taxon"]=="Micoureus_demerarae" | bLR_set[r, "Taxon"]=="Isoodon_obesulus" | bLR_set[r, "Taxon"]=="Antechinus_flavipes" | bLR_set[r, "Taxon"]=="Trichosurus_vulpecula" | bLR_set[r, "Taxon"]=="Equus_caballus" | bLR_set[r, "Taxon"]=="Echinops_telfairi" | bLR_set[r, "Taxon"]=="Orycteropus_afer"){
      bLR_set[r, "bin_fflip"] <- "Non-flippered"
      bLR_set[r, "bin_hflip"] <- "Non-flippered"
    }
  } # closes 'for(r in 1:nrow(bLR_set))' loop
  return(bLR_set)
} # closes bLRify function

# Generate binarified "bLR" dataset from full original dataset:
  all_bLR <- bLRify(dataset=all_data, dataset_name = "all") 
  names(all_bLR) # check that it has all the correct binarified variables
# Check that thalattosuchians were scored correctly:
  all_bLR$bin_eco1_v3[is.na(all_bLR$Subgroup2)==F & all_bLR$Subgroup2=="Thalattosuchia"]

# Define vectors of binary grouping variables for downstream analysis:
bLR_groupers <- names(all_bLR)[grep("bin", names(all_bLR))]
cat("\n","PROGRESS REP: chunk [S1.1.10] complete; starting next chunk...","\n")
#*-----**-----* 

#*-----**{ [S.1.11] Define transformation- and binary-variant datasets. }*
# Define output lists for new datasets:
log10_datasets <- as.list(rep(NA, length(orig_dataset_names)))
log10_dataset_names <- paste(orig_dataset_names, "_log10", sep="")
boxcox_datasets <- as.list(rep(NA, length(orig_dataset_names)))
boxcox_dataset_names <- paste(orig_dataset_names, "_boxcox", sep="")

bLR_raw_datasets <- as.list(rep(NA, length(orig_dataset_names)))
bLR_raw_dataset_names <- paste(orig_dataset_names, "_bLR_raw", sep="")

bLR_boxcox_datasets <- as.list(rep(NA, length(orig_dataset_names)))
bLR_boxcox_dataset_names <- paste(orig_dataset_names, "_bLR_boxcox", sep="")
bLR_log10_datasets <- as.list(rep(NA, length(orig_dataset_names)))
bLR_log10_dataset_names <- paste(orig_dataset_names, "_bLR_log10", sep="")

for(i in 1:length(orig_datasets)){
  # Create list of boxcox-transformed data:
  boxcox_datasets[[i]] <- BOXCOX.trans(dataset=orig_datasets[[i]], dataset_name=boxcox_dataset_names[i], vars_to_transform=all_quantitative_vars)
  
  # Create list of log10-transformed data:
  log10_datasets[[i]] <- log10.trans(dataset=orig_datasets[[i]], dataset_name=log10_dataset_names[i], vars_to_transform=all_quantitative_vars)
  
  # Create list of raw (untransformed) binarified data sets:
  cat("\n", "Running bLRify() function on ", orig_dataset_names[i],"...")
  bLR_raw_datasets[[i]] <- bLRify(dataset=orig_datasets[[i]])
  
  # Create list of boxcox-transformed binarified data sets:
  cat("\n", "Running bLRify() function on ", boxcox_dataset_names[i],"...")
  bLR_boxcox_datasets[[i]] <- bLRify(dataset=boxcox_datasets[[i]])
  
  # Create list of log10-transformed binarified data sets:
  cat("\n", "Running bLRify() function on ", log10_dataset_names[i],"...")
  bLR_log10_datasets[[i]] <- bLRify(dataset=log10_datasets[[i]])
} # closes 'for(i in 1:length(orig_datasets))' loop

# Name dataset list items (so they can be called with text strings):
  # eg, via ' boxcox_datasets[["reps_all_trans"]] '
  names(boxcox_datasets) <- boxcox_dataset_names
  names(log10_datasets) <- log10_dataset_names
  names(bLR_raw_datasets) <- bLR_raw_dataset_names
  names(bLR_boxcox_datasets) <- bLR_boxcox_dataset_names
  names(bLR_log10_datasets) <- bLR_log10_dataset_names
cat("\n","PROGRESS REP: chunk [S1.1.11] complete; starting next chunk...","\n")
#*-----**-----* 
  
#*-----**{ [S.1.12] Report sample sizes by subclade and measurement method. }*
# Define output file:
sink(paste0(output_path, "/*sample_sizes__", current_date, ".txt", sep=""))

# Get total number of specimens:
cat("---------------------------------------------------------------")
cat("\n", "Total number of SPECIMENS in data set:", dim(all_data)[1], "\n")
cat("---------------------------------------------------------------")

# Get total nvals for pan-mammals vs. pan-reptiles vs. outgroup spp:
cat("\n", "Total number of PAN-REPTILES in data set:", length(na.omit(all_data$MajorClade[all_data$MajorClade=="Pan-Reptilia"])), "\n")
cat("\n", "Total number of PAN-MAMMALS in data set:", length(na.omit(all_data$MajorClade[all_data$MajorClade=="Pan-Mammalia"])), "\n")
cat("\n", "Total number of NON-AMNIOTES in data set:", length(na.omit(all_data$MajorClade[all_data$MajorClade=="Outgroups"])), "\n")
cat("---------------------------------------------------------------")

# Get nvals for Recent vs Fossil specimens in full dataset:
cat("\n", "Total number of RECENT specimens in data set:", length(na.omit(all_data$MajorClade[grepl("Recent", all_data$Specimen_type)])), "\n")
cat("\n", "Total number of FOSSIL specimens in data set:", length(na.omit(all_data$MajorClade[grepl("Fossil", all_data$Specimen_type)])), "\n")
cat("---------------------------------------------------------------")

# Get total number of measurements in full dataset:
Measurements_and_Ratios <- sum(is.na(all_data[ , all_quantitative_vars]))
cat("\n", "Total number of MEASUREMENTS AND RATIOS in data set:", Measurements_and_Ratios, "\n")
Just_Measurements <- sum(is.na(all_data[ , direct_measurement_variables]))
cat("\n", "Total number of DIRECT MEASUREMENTS in data set:", Just_Measurements, "\n")
Just_measurements_in_mm <- sum(is.na(all_data[all_data$Length_units == "mm", direct_measurement_variables]))
cat("\n", "Total number of DIRECT MM MEASUREMENTS in data set, excluding pixel measurements:", Just_measurements_in_mm, "\n")
cat("---------------------------------------------------------------")

# Get sample size for each measurement method:
cat("\n", "Total number of specimens measured with CALIPERS (3D):", length(na.omit(all_data$MajorClade[all_data$Limb_region_measurement_method=="Calipers"])), "\n")
cat("\n", "Total number of specimens measured in SLICER (3D):", length(na.omit(all_data$MajorClade[all_data$Limb_region_measurement_method=="Slicer"])),"\n")
cat("\n", "Total number of specimens measured in ImageJ (2D):", length(na.omit(all_data$MajorClade[all_data$Limb_region_measurement_method=="ImageJ"])),"\n")
cat("---------------------------------------------------------------")

# Get sample size for each key subclade/subgroup in dataset:
cat("\n", "Total number of EURYAPSID specimens in data set:", length(rownames(euryaps_all)), "\n")
cat("\n", "Total number of THALATTOSAUR specimens in data set:", length(na.omit(all_data$MajorClade[all_data$Subgroup1=="Thalattosauria"])), "\n")
cat("\n", "Total number of ICHTHYOSAUROMORPH specimens in data set:", length(rownames(ichthyos_all)), "\n")
cat("\n", "Total number of ICHTHYOSAUR specimens in data set:", length(na.omit(all_data$MajorClade[all_data$Subgroup2=="Ichthyosauria"])), "\n")
cat("\n", "Total number of SAUROPTERYGIFORM specimens in data set:", length(rownames(sauropter_all)), "\n")
cat("\n", "Total number of SAUROSPHARGID specimens in data set:", length(na.omit(all_data$MajorClade[all_data$Subgroup2=="Saurosphargidae"])), "\n")
cat("\n", "Total number of PLACODONT specimens in data set:", length(na.omit(all_data$MajorClade[all_data$Subgroup2=="Placodontia"])), "\n")
cat("\n", "Total number of PACHYPLEUROSAUR specimens in data set:", length(na.omit(all_data$MajorClade[all_data$Subgroup2=="Pachypleurosaurs"])), "\n")
cat("\n", "Total number of NOTHOSAURS specimens in data set:", length(na.omit(all_data$MajorClade[all_data$Subgroup2=="Nothosaurs"])), "\n")
cat("\n", "Total number of BASAL (NON-PLESIOSAURIAN) PISTOSAUR specimens in data set:", length(na.omit(all_data$MajorClade[all_data$Subgroup2=="Nothosaurs"])), "\n")
cat("\n", "Total number of PLESIOSAUR specimens in data set:", length(na.omit(all_data$MajorClade[all_data$Subgroup2=="Plesiosauria"])), "\n")
cat("---------------------------------------------------------------")
cat("\n", "Total number of TURTLE specimens in data set:", length(rownames(turts_all)), "\n")
cat("---------------------------------------------------------------")
cat("\n", "Total number of ARCHOSAUROMORPH specimens in data set:", length(rownames(archos_all)), "\n")
cat("\n", "Total number of DINOSAUR specimens in data set (including birds):", length(rownames(dinos_all)), "\n")
cat("\n", "Total number of ORNITHISCHIAN & SAUROPODOMORPH specimens in data set:", length(na.omit(all_data$MajorClade[all_data$Subgroup1=="Dinosauria" &  !(all_data$Subgroup2=="Theropoda")])), "\n")
cat("\n", "Total number of NON-AVIALAN THEROPOD *specimens* in data set:", length(na.omit(all_data$MajorClade[all_data$Subgroup2=="Theropoda" &  is.na(all_data$Radius)==F])), "\n")
cat("\n", "Total number of NON-AVIALAN THEROPOD *species* in data set:", length(unique(na.omit(all_data$Taxon[all_data$Subgroup2=="Theropoda" &  is.na(all_data$Radius)==F]))), "\n")
cat("\n", "Total number of AVIALAN specimens in data set (just hindlimb data):", length(na.omit(all_data$Taxon[all_data$Subgroup2=="Theropoda" &  is.na(all_data$Radius)==T])), "\n")
cat("\n", "Total number of PSEUDOSUCHIAN specimens in data set:", length(rownames(crocs_all)), "\n")
cat("\n", "Total number of STEM-GROUP CROCODILIAN specimens in data set:", length(na.omit(crocs_all$Taxon[is.na(crocs_all$Subgroup2)==T | crocs_all$Subgroup2=="Thalattosuchia"])), "\n")
cat("\n", "Total number of THALATTOSUCHIAN specimens in data set:", length(na.omit(all_data$MajorClade[all_data$Subgroup2=="Thalattosuchia"])), "\n")
cat("\n", "Total number of CROWN-GROUP CROCODILIAN specimens in data set:", length(na.omit(crocs_all$Taxon[crocs_all$Subgroup2=="Alligatoroidea" | crocs_all$Subgroup2=="Crocodyloidea" | crocs_all$Subgroup2=="Gavialoidea"])), "\n")
cat("\n", "Total number of ALLIGATOROID specimens in data set:", length(na.omit(all_data$MajorClade[all_data$Subgroup2=="Alligatoroidea"])), "\n")
cat("\n", "Total number of CROCODYLOID specimens in data set:", length(na.omit(all_data$MajorClade[all_data$Subgroup2=="Crocodyloidea"])), "\n")
cat("\n", "Total number of GAVIALOID specimens in data set:", length(na.omit(all_data$MajorClade[all_data$Subgroup2=="Gavialoidea"])), "\n")
cat("---------------------------------------------------------------")
cat("\n", "Total number of LEPIDOSAUROMORPH specimens in data set:", length(rownames(lepidos_all)), "\n")
cat("\n", "Total number of RHYNCHOCEPHALIAN specimens in data set:", length(rownames(rhyncoceph_all)), "\n")
cat("\n", "Total number of ANGUIMORPH specimens in data set:", length(rownames(anguimorph_all)), "\n")
cat("\n", "Total number of IGUANIAN specimens in data set:", length(rownames(iguani_all)), "\n")
cat("\n", "Total number of AGAMID specimens in data set:", length(rownames(agamid_all)), "\n")
cat("---------------------------------------------------------------")
cat("\n", "Total number of STEM REPTILE specimens in data set:", length(rownames(stemreps_all)), "\n")
cat("---------------------------------------------------------------")
cat("\n", "Total number of UNGULATES specimens in data set:", length(rownames(ungulates_all)), "\n")
cat("\n", "Total number of FERAE specimens in data set:", length(rownames(ferae_all)), "\n")
cat("\n", "Total number of EUARCHONTOGLIRE specimens in data set:", length(rownames(euarchontagl_all)), "\n")
cat("\n", "Total number of ATLANTOGENATAN specimens in data set:", length(rownames(atlanto_all)), "\n")
cat("---------------------------------------------------------------")

# Close the external connection to output file:
sink()
cat("\n","PROGRESS REP: chunk [S1.1.12] complete; starting next chunk...","\n")
#*-----**-----* 

# This concludes the first step in our script_S1 pipeline. The coding environment is set up, the dataset has been cleaned, parsed, and processed for downstream analysis, and we have a record of how many specimens we're working with. In the next section of the script ([S1.2]), we will perform a quality control check on all measurements (i.e., a Bland-Altman check of congruence between measurement methods), and in the following one ([S1.3]) generate a critical input for downstream analyses: our phylogenetic trees.
#------------------------------------------------------------------------------

#**"-------------------------------------------------------------------------"*
#**"-------------- ### [S1.2] DO BLAND-ALTMAN ANALYSES ### ------------------"*
#**"-------------------------------------------------------------------------"*
# Our dataset involved measurements from both 2D digital specimens (high-resolution photographs), 3D digital specimens (meshes), and 3D physical specimens. In the following section, we perform Bland-Altman analyses to test whether these measurement methods produced congruent results. To do this, we measured a subset of samples twice using two separate methods, and then comparing the results below with Bland-Altman analyses to check for any systematic error between measurement methods. More information on the Bland-Altman analysis can be found in the original paper on this method (Altman & Bland 1983 [http://www.jstor.org/stable/2987937]). 

#*-----**{ [S1.2.01] Clean Bland-Altman dataset.  }*
# We begin by renaming and cleaning the Bland-Altman dataset (Table_S2), which contains all paired repeat measurements.
BA <- BlandAltman_data
names(BA)
BA_cleaned <- BA[, c(1,3,7:10, 11:26, 29:50, 52:63)]
names(BA_cleaned)

# Next, we parse the cleaned dataset into smaller objects that contain informative subsets of our measurements:
BAcl_all <- BA_cleaned
BAcl_stylo <- BA_cleaned[, c("Index", "Specimen_ID", "Specimen_type", "Measur_method", "Submethod", "BA_set", "Humerus", "Femur")]
BAcl_zeug <- BA_cleaned[, c("Index", "Specimen_ID", "Specimen_type", "Measur_method", "Submethod", "BA_set", "Radius", "Ulna", "EFZL", "Tibia")]
BAcl_acropod <- BA_cleaned[, c("Index", "Specimen_ID", "Specimen_type", "Measur_method", "Submethod", "BA_set", "Hand", "Foot", "mD_Antmost", "mD_Postmost", "mD_Longest", "pD_Antmost", "pD_Postmost", "pD_Longest")]
BAcl_phalanges <- BA_cleaned[, c("Index", "Specimen_ID", "Specimen_type", "Measur_method", "Submethod", "BA_set", "mD3_up_Height", "mD3_up_TopLength", "mD3_up_BottomLength", "pD3_up_Height", "pD3_up_BottomLength", "mD3_proxp_Length", "mD3_proxp_ProxWidth", "mD3_proxp_DistWidth", "pD3_proxp_Length", "pD3_proxp_ProxWidth", "pD3_proxp_DistWidth")]
BAcl_JUSTmeas <- BA_cleaned[, c("Index", "Specimen_ID", "Specimen_type", "Measur_method", "Submethod", "BA_set", "Humerus", "Radius", "Ulna", "EFZL", "Hand", "Femur", "Tibia", "Foot", "mD_Antmost", "mD_Postmost", "mD_Longest", "pD_Antmost", "pD_Postmost", "pD_Longest", "mD3_up_Height", "mD3_up_TopLength", "mD3_up_BottomLength", "pD3_up_Height", "pD3_up_BottomLength", "mD3_proxp_Length", "mD3_proxp_ProxWidth", "mD3_proxp_DistWidth", "pD3_proxp_Length", "pD3_proxp_ProxWidth", "pD3_proxp_DistWidth")]
BAcl_NOstylo <- BAcl_JUSTmeas[, c(1:6, 8:11, 13:31)]
BAcl_JUSTratios <- BA_cleaned[, c("Index", "Specimen_ID", "Specimen_type", "Measur_method", "Submethod", "BA_set", "AZR_forelimb", "fASR", "fSZR", "fAnAR", "AZR_hindlimb", "hASR", "hSZR", "hAnAR", "mD_Symmetry_index1", "mD_Symmetry_index2","pD_Symmetry_index1", "pD_Symmetry_index2", "mD3_up_FI1", "mD3_up_FI2", "mD3_up_FI3", "pD3_up_FI1", "pD3_up_FI2", "pD3_up_FI3", "mD3_proxp_Length.DistWidth", "mD3_proxp_WidthRatios", "mD3_proxp_WidthRatios.Length", "pD3_proxp_Length.DistWidth", "pD3_proxp_WidthRatios", "pD3_proxp_WidthRatios.Length")]

# To automate downstream analysis, we place these datasets and their names within lists or vectors:
BA_datasets <- list(BAcl_all, BAcl_stylo, BAcl_zeug, BAcl_acropod, BAcl_phalanges, BAcl_JUSTmeas, BAcl_NOstylo, BAcl_JUSTratios)
BA_names <- c("BAcl_all", "BAcl_stylo", "BAcl_zeug", "BAcl_acropod", "BAcl_phalanges", "BAcl_JUSTmeas", "BAcl_NOstylo", "BAcl_JUSTratios")
cat("\n","PROGRESS REP: chunk [S1.2.01] complete; starting next chunk...","\n")
#*-----**-----*

#*-----**{ [S1.2.02] Define BA.plot() function.  }*
# The BA.plot() function creates a Bland-Altman plot boxplot to compare the results of two measurement methods for our cleaned Bland-Altman data. The Bland-Altman plot displays the differences in value between two measurement types against their means. This plot is more effective in gauging agreement between two methods than a correlation metric (Altman and Bland 1983). The associated box plot shows the distribution and mean of the absolute values of these differences. The BA.plot() function also generates a set of diagnostic plots to check for heteroscedasticity among the sampled  differences.

BA.plot <- function(dataset, dataname, units){
  # Create subdirectory for Bland-Altman results:
  directory_path <- paste0(output_path, "/Bland-Altman_results", sep="") # All BA.plot() output files will be saved to this directory path.
  # Create directory for output files:
  dir.create(path=directory_path, showWarnings = TRUE, recursive = FALSE, mode = "0777") 
  
  # Calculating number of data point pairs to compare:
  npairs <- nrow(dataset)/2
  
  # Creating some preliminary objects to make creating the output data set easier:
  num_cols <- unlist(lapply(dataset, is.numeric))
  data_num <- dataset[ , num_cols]
  char_cols <- unlist(lapply(dataset, is.character))
  
  # Creating the output data sets:
  diffs <- data.frame((matrix(ncol = dim(data_num)[2], nrow = npairs)))
  means <- data.frame((matrix(ncol = dim(data_num)[2], nrow = npairs)))
  
  # Separating even and odd rows of the data:
  rows <- nrow(data_num)
  seq_len(rows)%%2
  odd_rows <- seq_len(rows) %% 2
  even_rows <- seq_len(rows) %% 2
  
  # Defining rows that contain Caliper vs. ImageJ data:
  Caliper_rows <- data_num[odd_rows == 1, ]
  ImageJ_rows <- data_num[even_rows == 0, ]
  
  # Computing differences and means between data pairs:
  for(i in 1:npairs){
    for(j in 1:ncol(data_num)){
      diffs[i, ] <- (Caliper_rows[i, ] - ImageJ_rows[i, ])
      means[i,j] <- mean(Caliper_rows[i, j], ImageJ_rows[i, j], trim=0);
      if(is.na(Caliper_rows[i,j])==TRUE){
        means[i,j] <- NA
      }
      if(is.na(ImageJ_rows[i,j])==TRUE){
        means[i,j] <- NA
      }
    } # closes 'for(j in 1:ncol(data_num))' loop
  } # closes 'for(i in 1:npairs)' loop
  
  # Calculating absolute and mean differences:
  absdiffs <- (abs(diffs));
  means_vec <- unlist(means)
  diffs_vec <- unlist(diffs)
  absdiffs_vec <- unlist(absdiffs)
  M_diff <- mean(diffs_vec, na.rm=TRUE)
  M_absdiffs <- mean(absdiffs_vec, na.rm=TRUE)
  ttest_result <- t.test(diffs, mu = 0)
  ttest_pval <- round(ttest_result$p.value, digits=4)
  
  # Generating figures:
  png(filename=paste0(directory_path, "/BAplot_", dataname, "_BlandAltmans.png"), width=4800, height=3000, res=300)
  layout_matrix <- matrix(1:3, ncol = 3)
  par(mfrow=c(3,1), mar=c(6,6,6,6))
  layout(layout_matrix, widths = 2:1, heights = 1:1)
  plot(x = means_vec, y = diffs_vec, type = "p", xlab = paste("Means (", units, ")", sep = ""), ylab = paste("Differences (", units, ")", sep = ""), pch = 19, col = 'red3', cex=2, ylim = c((M_diff + 2*sd(diffs_vec, na.rm=TRUE) + 0.5), (M_diff - 2*sd(diffs_vec, na.rm=TRUE) - 0.5)), cex.lab=2, cex.axis=2); 
  title(main = paste("Bland-Altman plot of ", dataname, sep=""))
  abline(h = M_diff, col = "black", lty = 2, lwd = 2)
  abline(h = (M_diff + 2*sd(diffs_vec, na.rm=TRUE)), col = 'grey50', lty = 2, lwd = 2);
  abline(h = (M_diff - 2*sd(diffs_vec, na.rm=TRUE)), col = 'grey50', lty = 2, lwd = 2);
  abline(h = M_diff + 1, col = 'blue3', lty = 2, lwd = 4);
  abline(h = M_diff - 1, col = 'blue3', lty = 2, lwd = 4);
  #text(x = means + 0.5, y = diffs + 0.5, labels = BA1$Spec..ID, cex = 0.3) # This text() command is optional, for showing the specimen number of each point on the BA plot.
  hist(diffs_vec, col='grey90', border='grey10', xlab="Differences (mm)", main=NA, cex.lab=2, cex.axis=2)
  boxplot(diffs_vec, col = 'red3', lwd = 4, main = NULL, ylab = "Differences (mm)", width = 2, xlab=paste("one-way t-test", "\n", "p = ", ttest_pval, sep=""), cex.lab=2, cex.axis=2);
  points(x = 1, y = round(M_absdiffs, 2), pch = 23, bg = 'gold2', col = 'grey10', cex = 6, lwd=3);
  scaler <- 0.28
  text(x = 1 + scaler, y = M_absdiffs, labels = round(M_diff, 3), col = 'grey10', cex = 1.5, font = 2);
  dev.off()
  
  png(filename=paste0(directory_path, "/BAplot_", dataname, "_heterosced_check.png"), width=4800, height=3000, res=300)
  layout_matrix <- matrix(1:3, ncol = 3)
  par(mfrow=c(3,1), mar=c(6,6,6,6))  
  layout(layout_matrix, widths = 2:1, heights = 1:1)
  
  cortest_result <- cor.test(means_vec, absdiffs_vec, method=c("pearson"))
  Pearson_coef <- round(cortest_result$estimate, digits = 2)
  cortest_pval <- round(cortest_result$p.value, digits = 4)
  
  plot(x = means_vec, y = absdiffs_vec, type = "p", xlab = paste("Means (", units, ")", sep = ""), ylab = paste("Absolute Differences (", units, ")", sep = ""), pch = 19, col = 'red3', cex=2, cex.lab=2, cex.axis=2); 
  abline(lm(absdiffs_vec ~ means_vec), lty = 2, col='grey10', lwd=2);
  mtext(paste("Pearson coef. = ", Pearson_coef, " ; ", "p = ", cortest_pval, sep=""), side=3)
  cormod <- lm(absdiffs_vec ~ means_vec)
  plot(cormod, 2, cex=3, cex.lab=2, cex.axis=2)
  plot(cormod, 4, cex=3, cex.lab=2, cex.axis=2)
  dev.off()
} # closes BA.plot() function

# Check that BA.plot() function works:
BA.plot(dataset = BA_datasets[[1]], dataname = BA_names[7], units = "mm")
cat("\n","PROGRESS REP: chunk [S1.2.02] complete; starting next chunk...","\n")
#*-----**-----*

#*-----**{ [S1.2.03] Apply BA.plot() to all parced Bland-Altman datasets. }*
for(i in 1:length(BA_names)){
  BA.plot(dataset = BA_datasets[[i]], dataname = BA_names[i], units = "mm")}
cat("\n","PROGRESS REP: chunk [S1.2.03] complete; starting next chunk...","\n")
#*-----**-----*

# We have just run Bland-Altman analyses to check whether ImageJ measurements from photographs and caliper measurements from physical specimens provide comparable results. At this point, you should manually review the Bland-Altman output files to assess whether these two measurement methods provide consistent results, and check whether any systematic error appears to result from the use of these two separate measurement methods. In previous runs, we found no evidence of systematic differences between measurement methods, and indeed found that they gave us congruent results. Given the observed variance seen between replicates in unrounded measurements, we rounded all length measurements to either the nearest 0.1 mm (for all lengths ≤ 10 mm) or to the nearest 1.0 mm (for all lengths > 10 mm). Once you have manually reviewed the Bland-Altman files and confirmed that there is no systematic error between measurement methods, you can move onto the third section of this script, where we will generate time-calibrated trees for all our datasets. 
#------------------------------------------------------------------------------

#**"-------------------------------------------------------------------------"*
#**"------------ ### [[S1.3]] MAKE TIME-CALIBRATED TREES ### ----------------"*
#**"-------------------------------------------------------------------------"*

# Time-calibrated trees are required inputs for all of the phylogenetic statistical tests we are conducting. In the section below, we import and check the topologies of previously written tree files, calibrate them with tip-dating using fossil occurrence data from the literature and The Paleobiology Database [https://paleobiodb.org/#/], and write functions to align these trees to our various datasets. Given lingering uncertainty about the interrelations and/or positions of various amniote groups (including turtles, enaliosaurs, and squamates), we produced eight alternative tree topologies to better reflect competing hypotheses, and ultimately run each of our statistical tests on every single tree, to assess how competing phylogenetic assumptions impact our results. These eight trees are labeled "supertree1" through to "supertree8". 

#*-----**{ [S1.3.01] Define tree directory path and diagnostic functions. }*
# Define subdirectory paths:
tree_directory_path <- paste0(wd_path, "*input_trees_", current_date, "_run/")

# Define function to align data to tree:
reorder.data <- function(trimmed_tree, trimmed_dataset){
  cat("\n", "Tip order of maximal tree (first three):", "\n")
  print(head(trimmed_tree$tip.label, 3))
  starting_set <- trimmed_dataset
  cat("\n", "Initial specimen order of dataset (first three):", "\n")
  print(head(rownames(starting_set), 3))
  match_labels <- match(trimmed_tree$tip.label, rownames(starting_set))
  reordered_set <- starting_set[match_labels, ]
  cat("\n", "New specimen order of reordered dataset (first three):", "\n")
  print(head(rownames(reordered_set), 3))
  return(reordered_set) # outputs new reordered dataset
} # closes reorder.data() function

# The code below defines a function to check whether there are an equal number of open & closed parentheses in Nwk file (this is a requirement for downstream analyses using the trees as inputs). Note: The following ReportParentheses function was written with the help of chatGPT v.3.5.
ReportParentheses <- function(newick_string, tree_name) {
  cat("\n", "Checking whether parentheses are balanced for", tree_name, "...", "\n")
  dir.create(path=paste0(tree_directory_path, "parenthesis_reports/"), showWarnings=T)
  full_path <- paste(tree_directory_path, "parenthesis_reports/", tree_name, "__parentheses.txt", sep="")
  sink(full_path)
  cat("PARENTHESIS REPORT FOR NEWICK TREE:", tree_name, "\n", "_____________________________________________", "\n")
  n_open <- 0
  n_close <- 0
  indent_level <- 1
  
  for(i in 1:nchar(newick_string)){
    char_i <- str_sub(string=newick_string, start=i, end=i)
    if(char_i=="("){
      n_open <- n_open + 1
      cat(rep(" ", indent_level), "pos", i, ":", "(", "\n")
      indent_level <- indent_level + 1  
      # Increases indent level for an open parenthesis
    }
    if(char_i==")"){
      n_close <- n_close + 1
      indent_level <- indent_level - 1
      # Decreases indent level for an closed parenthesis
      cat(rep(" ", indent_level), "pos", i, ":", ")", "\n")
    }
  } # closes 'for(i in 1:nchar(newick_string))' loop
  cat("\n","\n","_________________________________________________","\n","\n")
  cat("Total number of open parentheses:", n_open, "\n")
  cat("Total number of closed parentheses:", n_close, "\n")
  cat("\n", "\n", "_________________________________________________")
  sink(file = NULL)
  cat("\n", "Full report of all parentheses for", tree_name, "now available in the following subdirectory:", "\n", tree_directory_path,"\n")
} # closes ReportParentheses() function
cat("\n","PROGRESS REP: chunk [S1.3.01] complete; starting next chunk...","\n")
#*-----**-----*

#*-----**{ [S1.3.02] Run diagnostics on all Nwk files to check tree format. }*
# Write an empty output list to eventually populate with output trees:
tree_variants <- as.list(rep(NA, 8))

# Run for-loop to check parentheses for all trees:
for(i in 1:length(tree_variants)){
  input_path <- paste0(tree_directory_path, "supertree", i, ".txt")
  nwk_string_portions <- read.table(input_path, header=F, sep="")
  length(nwk_string_portions) # then change the number of pasted items below so that it's equal to the length
  maxTree_nwk <- paste(nwk_string_portions[1], nwk_string_portions[2], nwk_string_portions[3], nwk_string_portions[4])
  nchar(maxTree_nwk)
  ReportParentheses(newick_string=maxTree_nwk, tree_name=paste(current_date, "_tree", i, sep=""))
} # closes 'for(i in 1:length(tree_variants))' loop

# NOTE: You should check the newly generated parenthesis_report files before moving on. Once you have confirmed that all Newick files have balanced parentheses, you can proceed. As our next tree-checking step, we confirm that the tip labels match the data. This alignment between tree and dataset is another basic requirement for all of our statistical tests moving forward.

# Run for-loop to match tip_label order for all trees:
for(i in 1:length(tree_variants)){
  input_path <- paste0(tree_directory_path, "/", "supertree", i, ".txt")
  nwk_string_portions <- read.table(input_path, header=F, sep="")
  length(nwk_string_portions) # then change the number of pasted items below so that it's equal to the length
  maxTree_nwk <- paste(nwk_string_portions[1], nwk_string_portions[2], nwk_string_portions[3], nwk_string_portions[4])
  maxTree = read.tree(text=maxTree_nwk)
  tree_moniker <- paste("supertree", i, sep="")
  
  # Align data to tree:
  all_data_ord <- reorder.data(trimmed_tree=maxTree, trimmed_dataset=all_data_raw)
  
  # Define "tips" object for tree calibration:
  Tips <- cbind(all_data_ord$First_occurrence..mya., all_data_ord$last_occurrence..mya.)
  colnames(Tips) <- c("FAD", "LAD")
  rownames(Tips) <- all_data_ord$Tip_label
  
  # Check that the tip labels in the data set and on the tree file match. Each line below gives elements in 1st vec but not 2nd vec.
  setdiff_out1 <- setdiff(maxTree$tip.label, rownames(Tips))
  setdiff_out2 <- setdiff(rownames(Tips), maxTree$tip.label)
  
  # OPTIONAL TROUBLE-SHOOTING STEP:
       #print("DIFFERENCES:"); print(setdiff_out1); print(setdiff_out2)
        # If the output was "character(0)", then they match!
  
  if((length(setdiff_out1)==0) & (length(setdiff_out2)==0)){
    cat("\n", "TIP ORDER REPORT for",  tree_moniker, "|", "The tip and occurrence date orders match.", "\n")
  } else {
    cat("\n", "TIP ORDER REPORT for",  tree_moniker, "|", "The tip and occurrence date orders do NOT match.", "\n")
  } # closes 'if((length(setdiff_out1)==0)&(length(setdiff_out2)==0))' clause
} # closes 'for(i in 1:length(tree_variants))' loop
cat("\n","PROGRESS REP: chunk [S1.3.02] complete; starting next chunk...","\n")
#*-----**-----*

#*-----**{ [S1.3.03] Time-calibrate R 'phylo' objects; save tree tiffs. }*
# Run for-loop to generate chronogram and tiffs for each tree:
for(i in 1:length(tree_variants)) {
  # Report start of current for-loop iteration:
  cat("\n", "-------------------------------------------------------", "\n")
  cat("Running for-loop for tree variant #", i)
  cat("\n", "-------------------------------------------------------", "\n")
  
  # Create input and output paths for files:
  input_path <- paste0(tree_directory_path, "/", "supertree", i, ".txt")
  tree_output_path <- paste0(tree_directory_path, "/tree_tiffs/")
  dir.create(path=tree_output_path, showWarnings = TRUE, recursive = FALSE, mode = "0777")
  
  # Import and process Newick file:
  nwk_string_portions <- read.table(input_path, header=F, sep="")
  length(nwk_string_portions) # change number of pasted items below so it's equal to this length
  maxTree_nwk <- paste0(nwk_string_portions[1], nwk_string_portions[2], nwk_string_portions[3], nwk_string_portions[4])
  maxTree <- read.tree( text = maxTree_nwk )
  
  # Generate ultrametric tree:
  gg_current_tree <- ggtree( maxTree, size=0.1 ) + 
  geom_tiplab( geom="text", size=0.2, fontface = "italic", offset=0.1 ) +
    geom_nodepoint(col="black", alpha=0.6, size=0.04) + 
    xlim(c(-1,150)) + 
    ylim(-2,770) +
    theme(plot.margin = margin(t = 0, r = 0, b = 0, l = 0, unit = "pt"))
  
  # Save ultrametric tree as tiff file in "tree_variants" folder:
  ultrametric_tiff <- paste0(tree_output_path, "supertree", i, "_ultrametric.tif")
  tiff(filename=ultrametric_tiff, width=8, height=16, units='in', res=1200)
  show(gg_current_tree)
  dev.off()
  
  # Align data to tree:
  all_data_ord <- reorder.data(trimmed_tree = maxTree, trimmed_dataset = all_data_raw)
  
  # Define "tips" object for tree calibration:
  Tips <- cbind(all_data_ord$First_occurrence..mya., all_data_ord$last_occurrence..mya.)
  colnames(Tips) <- c("FAD", "LAD")
  rownames(Tips) <- all_data_ord$Tip_label
  
  # Tip-date the supertree:
  time.maxTree <- DatePhylo(tree=maxTree, ages=Tips, rlen=1, method="equal", add.terminal=F)
  tree_variants[[i]] <- time.maxTree
  names(tree_variants)[i] <- paste("supertree", i, sep="")
  
  # Plot tip-dated tree on geological time scale, and save it as high-res tiff:
  tipdated_tiff <- paste0(tree_output_path, "supertree", i, "_tipdated.tif")
  tiff(filename=tipdated_tiff, width=8, height=16, units="in", res=1200)
  geoscalePhylo(tree=time.maxTree, ages=Tips, units=c("Period", "Epoch", "Age"), boxes="Period", cex.tip=0.06, cex.age=0.4, cex.ts=0.5, erotate=0, label.offset=0.5, x.lim=c(390,-12), lwd=0.14, width=0.24)
   dev.off()
  
  # Report completion of for-loop for a given tree:
   cat("\n", "OUTPUT: High-res tiffs of ultrametric and tip-dated trees for tree variant #", i, "\n", "LOCATION:", tree_directory_path, "\n")
} # closes 'for(i in 1:length(tree_variants))' loop

# Check that output list of supertrees is correctly formatted:
(tree_variant_names <- names(tree_variants))
str(tree_variants[["supertree1"]])

# We have just generated eight time-calibrated (tip-dated) trees with alternate topologies, each including all the specimens for which we have LM data. With these supertrees prepared, we will be able to analyze our data within a phylo-genetic comparative framework. However, to do this, we must align our trees to our dataset in a consistent way, which will require original functions.
cat("\n","PROGRESS REP: chunk [S1.3.03] complete; starting next chunk...","\n")
#*-----**-----*

#*-----**{ [S1.3.04] Write functions to trim and align dataset and tree. }*
# As mentioned above, in order perform phylogenetic statistical tests in R, the tip labels of your tree must match the rownames of your dataset exactly. However, different species are missing data for each variable. As a result, different taxa will need to be pruned from the dataset when analyzing different variables, and the tree will need to be pruned accordingly so that the tips match the dataset. We write three functions to do this process below.

rm.NAs <- function(dataset, categorical_var, quantitative_var){
  trimmed_dataset <- dataset[c(!is.na(dataset[ , quantitative_var]) & !is.na(dataset[, categorical_var])), ]
  return(trimmed_dataset)} # closes rm.NAs() function

trim.tree <- function(tree, trimmed_dataset, verbose=T){
  starting_set <- trimmed_dataset
  if(verbose==T){
    cat("\n", "Number of specimens in dataset:", length(rownames(starting_set)), "\n")}
  starting_tree <- tree
  if(verbose==T){
    cat("\n", "Number of tips in starting tree:", length((starting_tree$tip.label)), "\n")}
  match_labels <- match(starting_tree$tip.label, rownames(starting_set))
  missing_tips <- starting_tree$tip.label[is.na(match_labels)==T]
  if(verbose==T){
    cat("\n", "Tree tips to remove:", length(missing_tips), "\n")
    print(head(missing_tips, 3))}
  tree_tr <- drop.tip(phy=starting_tree, tip=missing_tips)
  if(verbose==T){
    cat("\n", "Tree tips remaining:", length(tree_tr$tip.label), "\n")
    print(head(tree_tr$tip.label, 3))}
  return(tree_tr) # outputs new trimmed tree
} # closes trim.tree() function

reorder.data <- function(trimmed_tree, trimmed_dataset, verbose=T){
  if(verbose==T){
    cat("\n", "Tip order of trimmed tree (first three):", "\n")
    print(head(trimmed_tree$tip.label, 3))}
  starting_set <- trimmed_dataset
  if(verbose==T){
    cat("\n", "Initial specimen order of dataset (first three):", "\n")
    print(head(rownames(starting_set), 3))}
  match_labels <- match(trimmed_tree$tip.label, rownames(starting_set))
  reordered_set <- starting_set[match_labels, ]
  if(verbose==T){
    cat("\n", "New specimen order of reordered dataset (first three):", "\n")
  print(head(rownames(reordered_set), 3))}
  return(reordered_set) # outputs new reordered dataset
} # closes reorder.data() function

# Testing new functions to make sure they work:
test_ds <- all_data
test_ds_tr <- rm.NAs(dataset=test_ds, categorical_var="fore_type", quantitative_var = "f_AZR")
phylo_tr <- trim.tree(tree=time.maxTree, trimmed_dataset=test_ds_tr)
test_ds_tr_ord <- reorder.data(trimmed_tree=phylo_tr, trimmed_dataset=test_ds_tr)
  
# Save example pre- and post-pruning trees to check that they look right:
gg_tree_test <- ggtree(time.maxTree, branch.length="none", size=0.1) + geom_text(aes(label=node), hjust=-.1, size=0.3, col='red') + geom_tiplab(size=0.2, hjust=-.2)
tiff(paste0(output_path, "/testTree_beforePruning.tif"), res=1200, height=16, width=8, units="in")
show(gg_tree_test) # pre-pruning tree
dev.off()

gg_tree_test <- ggtree(phylo_tr, branch.length="none", size=0.1) + geom_text(aes(label=node), hjust=-.1, size=0.3, col='red') + geom_tiplab(size=0.2, hjust=-.2, col='grey30')
tiff(paste0(output_path, "/testTree_postPruning.tif"), res=1200, height=16, width=8, units="in")
show(gg_tree_test) # post-pruning tree
dev.off()
cat("\n","PROGRESS REP: chunk [S1.3.04] complete; starting next chunk...","\n")
#*-----**-----*

# Now that our supertrees are prepared and we have all the functions we need to align them to the data, we can begin analyzing the data within a phylogenetic framework. The more intensive phylogenetic statistical tests involved in this data analysis pipeline will be completed in later scripts (S2-S5). However, we have opted to get any less intensive tests out of the way in this script. These 'easier' tests involve assessing relationships among categorical variables. These shorter analyses do not currently include a phylogenetic component, but they could be modified to include one in the future. 
#------------------------------------------------------------------------------

#**"-------------------------------------------------------------------------"*
#**"-- ### [[S1.4]] EXPLORE RELATIONSHIPS AMONG CATEGORICAL VARIABLES ### ---"*
#**"-------------------------------------------------------------------------"*
# In this section, we check for pairwise associations between categorical variables. To do this, we calculate Cramer's V and run Fisher's exact test for each pair of grouping variables, and then visualize the percentage overlap between them with a custom category plot ("catplot") function. We do this to see how different binning systems in our dataset overlap.

#*-----**{ [S1.4.01] Define palettes for all downstream analyses. }*
# To begin data analysis, we first need to define color palettes that will make all our figures look nice and consistent across scripts. We do that with the list below, which gives a separate vector of colors for every grouping variable in the dataset, and which is called in all of our subsequent custom functions that involve plot-building.
grouping_var_palette <- list(
  "MajorClade" = c("Outgroups" = "white", "Pan-Reptilia" =  "white", "Pan-Mammalia" = "white"), 
  "Subclade" = c("Outgroups" = "grey15", "Stem reptiles" = "black", "Euryapsida" = "darkturquoise","Lepidosauromorpha" = "lightgreen","Testudines" = "darkgreen","Archosauromorpha" = "gold2","Non-placentals" = "darkorange", "Atlantogenata" = "orange3", "Euarchontoglires" = "orange4", "Eulipotyphla" = "saddlebrown", "Ferae" = "red2", "Ungulata" = "navy"), 
  
  "Ecotype1" = c("Terrestrial" = "#CA0020", "Transiently Aquatic" = "#F4A582", "Moderately Aquatic" = "lightcyan", "Highly Aquatic" = "#92C5DE", "Fully Aquatic" = "#0571B0", "Data Deficient" = "grey"),
  "Ecotype2" = c("almost always on land" = "#B2182B", "primarily on land, seldom in water" = "#FDDBC7", "primarily on land, often in water" = "#D1E5F0", "small or slow-moving water bodies" = "#67A9CF", "large or fast-moving water bodies" = "#4575B4", "data deficient" = "grey"),
  
  "Ecotype3" = c("predominantly terrestrial" = "#D73027", "terrestrial/freshwater" = "#E0F3F8", "terrestrial/euryhaline" =  "#E0F3F8", "terrestrial/marine" = "#E0F3F8", "predominantly freshwater" = "#91BFDB", "freshwater/marine" = "#91BFDB", "predominantly marine" = "#4575B4", "data deficient" = "grey"),
  
  "IUCN_cat" = c("DD" = "grey", "LC" = "white", "NT" = "orange","VU" = "orange", "EN" = "red", "CR" = "red", "EX" = "grey5"),
  
  "Flipper_status" = c("Non-flippered" = "tomato", "Flippered" = "turquoise", "Ambiguous" =  "grey"), 
  "limb_type" = c("Unwebbed" = "tomato", "Webbed" = "gold", "Flippered" = "turquoise", "Unclear, terr." =  "grey", "Unclear, aquat." = "grey"), 
  
  "fore_type" = c("Unwebbed" = "tomato", "Webbed" = "gold", "Flippered" = "turquoise", "Unclear, terr." =  "grey", "Unclear, aquat." = "grey"),
  
  "hind_type" = c("Unwebbed" = "tomato", "Webbed" = "gold", "Flippered" = "turquoise", "Unclear, terr." =  "grey", "Unclear, aquat." = "grey"),
  
  "limb_subtype" = c("tomato", "tomato", "tomato", "tomato", "tomato", "gold", "gold", "gold", "gold", "gold", "turquoise", "turquoise", "grey", "grey"), 
  
  "f_subtype" = c("Unwebbed" = "tomato", "Interdigital Girth" =  "tomato", "Stubbed" = "tomato", "Pincer-palmed" = "tomato", "Interdigital Integument" = "tomato", "Partial, slight" = "gold", "Partial, extensive" = "gold", "Complete, slight" = "gold","Complete, extensive" = "gold", "Lobate" = "gold", "Weakly Flippered" =  "turquoise", "Strongly Flippered" = "turquoise", "Unclear, terr." = "grey", "Unclear, aquat." = "grey"), 
  
  "h_subtype" = c("Unwebbed" = "tomato", "Interdigital Girth" =  "tomato", "Stubbed" = "tomato", "Pincer-palmed" = "tomato", "Interdigital Integument" = "tomato", "Partial, slight" = "gold", "Partial, extensive" = "gold", "Complete, slight" = "gold","Complete, extensive" = "gold", "Lobate" = "gold", "Weakly Flippered" =  "turquoise", "Strongly Flippered" = "turquoise", "Unclear, terr." = "grey", "Unclear, aquat." = "grey"), 
  
  "bin_eco1_v1" = c("#CA0020", "#0571B0"),
  "bin_eco1_v2" = c("#CA0020", "#0571B0"),
  "bin_eco1_v3" = c("#CA0020", "#0571B0"),
  "bin_eco1_v4" = c("#CA0020", "#0571B0"),
  "bin_eco1_v5" = c("#CA0020", "#0571B0"),
  "bin_eco1_v6" = c("#CA0020", "#0571B0"),
  "bin_eco1_v7" = c("#CA0020", "#0571B0"),
  "bin_eco1_v8" = c("#CA0020", "#0571B0"),
  "bin_eco1_v9" = c("#CA0020", "#0571B0"),
  "bin_eco2" = c("#CA0020", "#0571B0"),
  "bin_fweb" = c("tomato", "gold"),
  "bin_fflip" = c("tomato", "turquoise"), 
  "bin_hweb" = c("tomato", "gold"),
  "bin_hflip" = c("tomato", "turquoise"),
  "bin_Method" = c("white", "white"),
  "bin_specmType" = c("white", "white"),
  "Limb_region_measurement_method" = c("white", "white", "white")
) # closes grouping_var_palette list
cat("\n","PROGRESS REP: chunk [S1.4.01] complete; starting next chunk...","\n")
#*-----**-----*

#*-----**{ [S1.4.02] Define catPlot() function. }*
catPlot <- function(dataset, dataset_name, catVar1, catVar2) {
  # Define directory path for output files: 
  directory_path <- paste0(output_path, "/catPlots")
  # Create directory for output files:
  dir.create(path=directory_path, showWarnings = TRUE, recursive = FALSE, mode = "0777")
  
  # Define variables within function:
  fill_colors <- grouping_var_palette[[catVar2]]
  dataset$catVar1 <- dataset[, catVar1]
  dataset$catVar2 <- dataset[, catVar2]
  
  # Remove missing values from input data:
  ds_tr <- rm.NAs(dataset=dataset, categorical_var=catVar1, quantitative_var=catVar2)
  
  # Create contingency table:
  contingency_table <- table(ds_tr$catVar1, 
                             ds_tr$catVar2)
  # Perform Cramer's V test:
  Vval <- round(assocstats(contingency_table)$cramer, 3)
  
  # Perform Fisher's exact test:
  pval <- chisq.test(contingency_table, simulate.p.value = TRUE)$p.value
  pval <- format(pval, scientific=TRUE, digits=3)
  pval_text <- paste("p = ", pval, sep="")
  
  # Generate output table:
  results_df <- data.table("Cramer's V" = Vval, "Fisher's Exact Test" = pval_text)
  Cramer_table <- tableGrob(results_df, theme = ttheme_default(base_size=20))
  
  # Calculate proportions for stacked barplot: 
    # Prepare data objects:
      cv1_levs <- levels(droplevels(ds_tr$catVar1))
      cv2_levs <- levels(droplevels(ds_tr$catVar2))
      df <- expand.grid(catVar1 = cv1_levs, catVar2 = cv2_levs)
      counts <- table(ds_tr$catVar1, ds_tr$catVar2)
  
  # Calculate the count for each variable pair combination:
    # Create an empty data frame to store the results
      df_count <- as.data.frame(counts)
      colnames(df_count) <- c("catVar1", "catVar2", "freq")
  # Make the stacked barplot:
  catplot <- ggplot(df_count, aes(x = catVar1, y = freq, fill = catVar2)) +
    geom_col(position = "fill", color = 'white', size = 1.4, alpha=0.8) + 
    scale_y_continuous(labels = scales::percent_format()) +
    labs(aes(x = catVar1, y = "Proportion", fill = catVar2)) +
    scale_fill_manual(values = fill_colors) + 
    theme(panel.grid = element_line(color = adjustcolor("white", alpha = 0.1)), panel.background = element_rect(fill = adjustcolor("grey10", alpha = 0.8)), axis.text.x = element_text(angle = 90, vjust = 0.5, hjust=1, size=4))
  
  # Prepare to save high-resolution PND of catplot plot:
  png(filename=paste0(directory_path, "/", "catPlot__", dataset_name, "__", catVar1, "_by_", catVar2, ".png"), width=10, height=10, units="in", res=300)
  
  # Merge plot objects into one final plot:
  (final_catplot <- grid.arrange(catplot, Cramer_table, nrow=2, heights=c(8,2))) # saves output of this command to tiff based on previous tiff() and following dev.off() commands.
  dev.off()
  cat("\n", "A cat plot showing the association between", catVar1, "and", catVar2, "in", dataset_name, "has been saved to the working directory.", "\n")
} # closes catPlot() function

# Test function to make sure it works:
catPlot(dataset=all_data, dataset_name = "***test", catVar1 = "Ecotype2", catVar2 = "Ecotype1")
cat("\n","PROGRESS REP: chunk [S1.4.02] complete; starting next chunk...","\n")
#*-----**-----*

#*-----**{ [S1.4.03] Run catPlot() on NON-binary dataset-variable combos. }*
catPlot_groupers <- c("Subclade", "Ecotype1", "Ecotype2", "Ecotype3", "fore_type", "hind_type")
foreach(i=1:length(orig_datasets), .packages = c("vctrs", "forcats", "data.table", "stringr", "ggplot2", "plyr", "dplyr", "RColorBrewer", "grid", "gridExtra", "vcd", "ggimage", "MASS")) %dopar% {
  # Run rest of nested for-loop within foreach loop:
  for(j in 1:length(catPlot_groupers)){
    for(k in 1:length(catPlot_groupers)){
      if(j==k){
        cat("catPlot skipped for", catPlot_groupers[j], "by", catPlot_groupers[k], "\n", "REASON: They're the same variable, so they overlap completely.")
      } else {
        catPlot(dataset=orig_datasets[[i]], dataset_name = orig_dataset_names[i], catVar1 = catPlot_groupers[j], catVar2 = catPlot_groupers[k])
      }
    } # closes 'for(k in 1:length(catPlot_groupers))' loop
  } # closes 'for(j in 1:length(catPlot_groupers))' loop
} # closes 'foreach(i=1:length(orig_datasets)' loop
cat("\n","PROGRESS REP: chunk [S1.4.03] complete; starting next chunk...","\n")
#*-----**-----*

#*-----**{ [S1.4.04] Run catPlot() on key binary dataset-variable combos. }*
foreach(j=1:length(bLR_groupers), .packages = c("vctrs", "forcats", "data.table", "stringr", "ggplot2", "plyr", "dplyr", "RColorBrewer", "grid", "gridExtra", "vcd", "ggimage", "MASS")) %dopar% {
  for(k in 1:length(bLR_groupers)){
    if(j==k){
      cat("catPlot skipped for", bLR_groupers[j], "by", bLR_groupers[k], "\n", "REASON: They're the same variable, so they overlap completely.")
    } else {
      catPlot(dataset=all_bLR, dataset_name = "all_bLR",
              catVar1 = bLR_groupers[j], 
              catVar2 = bLR_groupers[k])
    }
  } # closes 'for(k in 1:length(bLR_groupers))'
} # closes 'foreach(j=1:length(bLR_groupers)' loop
cat("\n","PROGRESS REP: chunk [S1.4.04] complete; starting next chunk...","\n")
#*-----**-----*

# We have now finished all of the upstream tasks and computationally non-intensive tasks associated with our data analysis pipeline. In what follows, we save all this upstream information to make it quickly accessible for downstream scripts.
#------------------------------------------------------------------------------

#**"-------------------------------------------------------------------------"*
#**"- ### [S1.5] SAVE JOB LISTS & R ENVIRONMENT FOR DOWNSTREAM SCRIPTS  ### -"*
#**"-------------------------------------------------------------------------"*

# In the final section of script_S1, we will write job lists and custom functions for scripts S3-S4, save the final R environment, and report how long the current script took to run. The saved environment file will allow all downstream scripts (S2-S5) to pick up where this one left off, with all the necessary packages, cleaned and parced datasets, tree files, custom functions, grouping variable color schemes, and other information ready from the start.

# The job lists and custom functions will allow us to run script S3-S4 in massively parallel fashion using Job Arrays.

#*-----**{ [S1.5.01] Generate job array lists for downstream scripts. }*
# Generate job list for script_S3 Job Arrays:
  # Define modified data objects for script_S3:
    key_quantitative_vars <- all_quantitative_vars[! all_quantitative_vars %in% c("f_ASR", "f_AnAR", "h_AnAR", "h_ASR", "EFZL", "mD_Antmost", "mD_Postmost", "mD_Longest", "pD_Antmost", "pD_Postmost", "pD_Longest", "mD3_up_BottomLength", "pD3_up_BottomLength")] # This vector shortens the total number of informative quantitative variables used for phylogenetic correlation matrices to 35, in order to reduce the total job array number for phylogenetic correlation tests to below 10,000 (35 corvariables * 35 corvariables * 8 supertrees = 9800 jobs).
    
  log10_mm <- log10.trans(dataset=all_mm, dataset_name="log10_mm", vars_to_transform=all_quantitative_vars)
  rownames(log10_mm) <- log10_mm$Tip_label
  boxcox_mm <- BOXCOX.trans(dataset=all_mm, dataset_name="boxcox_mm", vars_to_transform=all_quantitative_vars)
  rownames(boxcox_mm) <- boxcox_mm$Tip_label
  
  all_cor <- all_mm[, key_quantitative_vars]
  log10_cor <- log10_mm[, key_quantitative_vars]
  boxcox_cor <- boxcox_mm[, key_quantitative_vars]
  corvariables <- names(all_cor)
  sysdate_string <- gsub(pattern="-", replacement=".", x=sysdate)
  script_S3_name <- paste0("script_S3_", sysdate_string, ".R")

  # Specify index lengths: 
  i_max=length(tree_variants)
  j_max=length(corvariables)
  k_max=length(corvariables)
  
  # Specify output file connection:  
  sink(file = paste0(wd_path, "script_S3_joblist.txt"))
  
  # Populate output file:
  for(i in 1:i_max){
    for(j in 1:j_max){
      for(k in 1:k_max){
        cat("module load StdEnv; module load R/4.2.0-foss-2020b; Rscript", script_S3_name, i, j, k, "\n")
      }
    }
  }
  
  # Terminate output file connection:
  sink(file = NULL)

# Generate job list for script_S4 Job Arrays:
  # Define modified data objects for script_S4:
    key_bLR_groupers <- c("bin_fweb", "bin_fflip", "bin_hweb", "bin_hflip", "bin_eco1_v1", "bin_eco1_v3", "bin_eco1_v4", "bin_eco1_v5", "bin_eco2") # This vector shortens the total number of binary grouping variables used for phylogenetic binomial logistic regression (phybLR) analyses, in order to reduce the total job array number to less than 10,000. Ideally, given greater computational resources, I would include bin_eco1_v8 and bin_eco1_v9 in this vector as well, but they can be input separately during a future run if needed.
    baremin_ratio_vars <- c("f_AZR","f_SZR","h_AZR","h_SZR", "mD_Symmetry_index1", "mD_Symmetry_index2", "pD_Symmetry_index1", "pD_Symmetry_index2", "mD3_up_FI1", "mD3_up_FI3", "pD3_up_FI1", "pD3_up_FI3", "mD3_proxp_Length.DistWidth", "mD3_proxp_WidthRatios", "pD3_proxp_Length.DistWidth", "pD3_proxp_WidthRatios") # This vector shortens the total number of informative ratio_vars used for phybLR analyses, in order to reduce the total job array number to less than 10,000. Ideally, given greater computing resources, I would include f_ASR, h_ASR, f_ANAR, and h_AnAR in this vector as well, but they can be input separately during a future run if needed.
    script_S4_name <- paste0("script_S4_", sysdate_string, ".R")
    
  # Specify index lengths: 
  i_max=length(bLR_raw_datasets)
  j_max=length(tree_variants)
  k_max=length(key_bLR_groupers)
  L_max=length(baremin_ratio_vars)
  
  # Specify output file connection:  
  sink(file = paste0(wd_path, "/", "script_S4_joblist.txt"))
  
  # Populate output file:
  for(i in 1:i_max){
    for(j in 1:j_max){
      for(k in 1:k_max){
        for(L in 1:L_max){
          cat("module load StdEnv; module load R/4.2.0-foss-2020b; Rscript", script_S4_name, i, j, k, L, "\n")
        }
      }
    }
  }
  
  # Terminate output file connection:
  sink(file = NULL)
cat("\n","PROGRESS REP: chunk [S1.5.01] complete; starting next chunk...","\n")
#*-----**-----*

# Now that we have written job lists for job arrays, we will define custom functions to apply across those jobs (which will be applied in scripts S3, S4a, S4b, and S4c). We define those functions in the next two chunks of code.

# The original phycorr() function, defined below, will be used to run phylogenetic correlation tests on every pairwise variable combination for every tree using the corphylo() function from the R package ape [https://www.rdocumentation.org/packages/ape/versions/5.4-1].

#*-----**{ [S1.5.02] Define phycorr() wrapper function.}*
# The original function below is a wrapper function designed to run corphylo(), and all required upstream functions, on every pairwise combination of quantitative variables in our dataset. This will make it easier for us to parallelize the function using job arrays in script_S3.

# Specify corphylo parameters for downstream usage:
REML_verdict <- TRUE
optim_method <- "Nelder-Mead"
constraind_verdict <- TRUE
maxitNM_num <- 500 # set to 2 for troubleshooting, and 500 for actual batch job

# Write phycorr() function:
phycorr <- function(input_ds, input_ds_name, corvar1, corvar2, input_tree_name, output_name, method = "Nelder-Mead", REML, constrain.d, maxit.NM, verbose=FALSE){
  # FUNCTION PARAMETERS:
  # input_ds is an input data frame containing corvar1 and corvar2
  # input_ds_name is a chr string naming the input dataset.
  # corvar1 and corvar2 are chr strings specifying the input_ds variables to be correlated with the corphylo function.
  # input_tree_name is a chr string specifying the name of the input tree (must be name of item in tree_variants list).
  # output_name is a chr string specifying the name of the phycorr output object for downstream indexing.
  # method is a chr string specifying whether to use "Nelder-Mead" or "SANN" minimization (specifying corphylo arg of the same name). We default to "Nelder-Mead", and additional modifications would be required to make this wrapper function work with SANN minimization.
  # REML is either TRUE or FALSE, specifies whether REML is used for function instead of ML. We default to TRUE, and additional dditional modifications would be required to make this wrapper function work with ML.
  # constrain.d is either TRUE or FALSE, specifying whether to constrain the d parameter to be btwn 0 and 1 (specifying the corphylo arg of the same name).
  # maxit.NM is a numeric value specifying the maximum number of iterations in the optimization with Nelder-Mead minimization (specifying the corphylo arg of the same name).
  # verbose specifies whether to print intermediate function outputs to the console. 
  
  # Skip variables with perfect correlation 
  # NOTE: This avoids a potential error in chol.default(cov(Xs)).
  if(corvar1==corvar2){
    phycorr_output <- "phycorr skipped because corvar1 = corvar2"
  } else { # continue with function
    
    # Define tree and corvars within function:
    input_tree <- tree_variants[[input_tree_name]]
    input_ds$corvar1 <- input_ds[, corvar1]
    input_ds$corvar2 <- input_ds[, corvar2]
    
    # Trim & align dataset and tree:
    ds_tr <- rm.NAs(dataset=input_ds, categorical_var = corvar1, quantitative_var = corvar2)
    tree_tr <- trim.tree(tree=input_tree, trimmed_dataset = ds_tr, verbose = verbose)
    ds_ord <- reorder.data(trimmed_tree=tree_tr, trimmed_dataset=ds_tr, verbose = verbose)
    ds_ord_cor <- ds_ord[, all_quantitative_vars]
    rownames(ds_ord_cor) <- ds_ord$Tip_label
    
    # Run phylogenetic correlation test:
    phycorr_output <- corphylo(X = ds_ord_cor[, c(corvar1, corvar2)], U = NULL, SeM = NULL, phy=tree_tr, REML = REML, method = method, constrain.d = constrain.d, maxit.NM = maxit.NM, verbose = FALSE)   
    # Record sample size of trimmed and aligned dataset:
    sample_size <- nrow(ds_ord_cor)
    phycorr_output$sample_size <- sample_size
  } # closes 'if(corvar1==corvar2){ } else { }' clause
  
  # Record corvars, output name, & input tree associated w/ phycorr output obj:
  phycorr_output$corvar1 <- corvar1
  phycorr_output$corvar2 <- corvar2
  phycorr_output$input_dataset <- input_ds_name
  output_name <- paste("phycorr output |", corvar1, "x", corvar2, "| DATASET:", input_ds_name, "| TREE:", input_tree_name)
  phycorr_output$output_name <- output_name
  phycorr_output$input_tree <- input_tree_name
  return(phycorr_output)
} # closes phycorr function

# Test phycorr() function:
corvar1_string <- "f_AZR"
corvar2_string <- "Humerus"
test_output <- phycorr(input_ds=all_mm, input_ds_name="all_mm", corvar1=corvar1_string, corvar2=corvar2_string, input_tree_name = "supertree1", method = optim_method,  REML = REML_verdict, constrain.d = constraind_verdict, maxit.NM = maxitNM_num)
cat("\n","PROGRESS REP: chunk [S1.5.02] complete; starting next chunk..","\n")
#*-----**-----*

# We have just defined the phycorr() function. We will run this function on every variable pair & tree combination for raw, log10-transformed, and BoxCox-transformed datasets by running script_S3 in parallel using Job Arrays. 

# The next original function, phybLR(), will be used in our S4 scripts (S4a, S4b, S4c) to fit phylogenetic binomial logistic regression models to the data using the phylolm package [https://cran.r-project.org/web/packages/phylolm/index.html] and assess their accuracy using Receiver Operating Characteristic (ROC) curves. ROC analysis is a method for visualizing and comparing predictive model performance. This method was developed by U.S. military statisticians in World War II. We describe our usage of ROC analysis in more detail within our paper's Methods section, and a great introduction to the approach is given by Fawcett (2006) [doi:10.1016/j.patrec.2005.10.010]. In brief, ROC curves provide a quick way to richly visualize, transparently report, and clearly compare the performance of multiple competing predictive models--in our case, phylogenetic binomial logistic regression (phybLR) models.

#*-----**{ [S1.5.03] Define phybLR() function. }*
# This function fits a phylogenetic binomial logistic regression model with a single predictor variable, saves a predicted probability curve with associated model stats to the working directory, and returns the output sensitivity and specificity data for downstream ROC analysis. The phybLR() function works downstream of the bLRify(), rm.NAs(), trim.tree(), and reorder.data() functions defined in script_S1.

phybLR <- function(dataset_ordered, dataset_name, response, predictor, trimmed_tree, tree_name, boot_num, btol_num, Kfold=3, CV_runs=1000, crossval_reports=T, plot_td_ROCCs=T, plot_Youden=T, plot_McF=T, plot_CIs=T, Firth_method="logistic_MPLE"){
  # FUNCTION ARGUMENTS:
  # dataset_ordered: dataset to start with (data frame)--should be the result of the bLRify(), rm.NAs(), and reorder.data() functions
  # dataset_name: chr string giving name of dataset_ordered (for naming output files)
  # response: name of response variable (chr string)
  # predictor: name of predictor variable (chr string)
  # trimmed_tree: starting tree--should be output from trim.tree() function
  # tree_name: chr string giving name of trimmed_tree (for naming output files)
  # boot_num: assigns the bootstrap replicate number for phyloglm()
  # btol: assigns the linear predictor bound for phyloglm()
  # Kfold: determines the number of bins into which the dataset will be divided for cross-validation-like subsampling. A Kfold=3 means that the dataset will be divided into thirds for cross-validation, such that 2/3 will be used to train the phybLR model and the remaining 1/3 will be used to test it. This cross-validation procedure will be repeated as many times as specified by the CV_runs argument.
  # CV_runs: numeric value specifying the number of cross-validation runs to perform per model.
  # cROCC_CIs: specifies whether function will plot 95% CI bands for cROCC (TRUE or FALSE)
  # plot_td_ROCCs: determines whether function will output a plot of all training-and-testing data ROCs (TRUE or FALSE)
  # plot_McF: determines whether function will output a scatterplot of AUC vs. McFadden's pseudo-R^2 (TRUE or FALSE)
  # plot_Youden: determines whether function will output a plot of predicted probability threshold vs. Youden's J-statistic (TRUE or FALSE)
  # plot_CIs: determines whether function will output dot-and-whisker plots showing 95% AUC, TPR, and FPR confidence intervals (TRUE or FALSE)
  # Firth_method: a chr vector passed onto the phyloglm() function, specifying the method to be used to apply Firth's correction (one of "logistic IG10", "logistic_MPLE", or "poisson_GEE"); see the manpage for phyloglm() for more details.
  
  # IMPORTANT NOTE ON tdROCC averaging method: there are multiple methods by which tdROCC curves can be averaged to generate the consensus ROC curve (cROCC). These are the "aggregative", "vertical", "horizontal", and "threshold" averaging methods, and all but the horizontal method are described in detail by Fawcett 2006 [doi:10.1016/j.patrec.2005.10.010].
  ### The "aggregative" method involves taking all the testing data results generated from K-fold cross-validation sampling runs, and combining them into a single, aggregated testing data set. Then the TPRs and FPRs across thresholds are determined for this aggregated set. As explained by Fawcett 2006 [doi:10.1016/j.patrec.2005.10.010], the tradeoff of this simple method is that it provides no estimate of tdROCC variance, since all testing data sets are aggregated into a single test set. 
  ### The "vertical" averaging method vertically samples all tdROCC TPRs for a given fixed FPR and averages them. As explained by Fawcett 2006 [doi:10.1016/j.patrec.2005.10.010], this method is most appropriate when the researcher can fix the FPR or needs a one-dimensional measure of tdROCC variance. The vertical averaging method is thus discouraged when the researcher lacks direct experimental control of the FPR.
  ### The "horizontal" method should, by analogy with the "vertical" method described by Fawcett 2006 [doi:10.1016/j.patrec.2005.10.010],  sample all tdROCC FPRs for a given fixed TPR and then average them. As with "vertical" averaging, this "horizontal" method is most appropriate when the researcher can fix the TPR or needs a one-dimensional measure of tdROCC variance. The vertical averaging method is thus discouraged when the researcher lacks direct experimental control of the TPR.
  ### The "threshold" averaging method samples all the tdROCC FPRs and TPRs for a given threshold and averages them separately, to produce a single, average (FPR, TPR) coordinate for each threshold. This method is generally preferable because it produces a richer estimate of tdROCC variance across both axes of ROC space and fixes only the classification threshold during averaging calculations, which is typically already under the direct control of the researcher. Given its advantages, we use this "threshold" averaging method in the phybLR() function. To use an alternative tdROCC averaging method to generate a cROCC, one would need to modify the function with a new argument (eg, a 'ROCC_averaging_method' argument with a chr string input of either "threshold", "aggregative", "vertical", or "horizontal"). Although it would be possible to implement these additional averaging methods within the function, it would be difficult, as the current function indexes primarily over thresholds, which is most conducive to the "threshold" averaging method.
  
  # Define dataset and variables within function:
  starting_set <- dataset_ordered
  starting_set$response <- starting_set[, response]
  starting_set$predictor <- starting_set[, predictor]
  
  # Make response variable binary:
  starting_set$response <- as.numeric(factor(starting_set$response)) - 1
  ds_fin <- starting_set
  
  # Skip mismatched forelimb-hindlimb variable combinations:
  proceed_verdict <- "yes" #this is the default start state
  if((str_detect(string=response, pattern="_f")==T) & (str_detect(string=predictor, pattern="h_|pD")==T)){
    warning("\n", "phylobLR() has skipped the following dataset-variable triad:", "\n", "\t", "dataset:", dataset_name, "\n", "\t", "predictor variable:", predictor, "\n", "\t", "response variable:", response, "\n", "REASON: One variable describes forelimbs and the other hindlimbs.", "\n")
    bLR_output <- "phybLR() skipped because one variable describes forelimbs and the other describes hindlimbs."
    return(bLR_output)
    proceed_verdict <- "no"
  }
  if((str_detect(string=response, pattern="_h")==T) & (str_detect(string=predictor, pattern="f_|mD")==T)){
    warning("\n", "phylobLR() has skipped the following dataset-variable triad:", "\n", "\t", "dataset:", dataset_name, "\n", "\t", "predictor:", predictor, "\n", "\t", "response variable:", response, "\n", "REASON: One variable describes forelimbs and the other hindlimbs.)", "\n")
    bLR_output <- "phybLR() skipped because one variable describes forelimbs and the other describes hindlimbs."
    return(bLR_output)
    proceed_verdict <- "no"
  }
  
  # Skip dataset-variable combinations with only one occupied response level:
  checker_ds <- ds_fin$response[is.na(ds_fin$predictor)==F]
  checker_ds <- as.numeric(factor(checker_ds)) - 1 # makes it binary
  obs_w0 <- length(na.omit(checker_ds[checker_ds==0]))
  obs_w1 <- length(na.omit(checker_ds[checker_ds==1]))
  if(obs_w0>0 & obs_w1>0){
    proceed_verdict <- "yes" # both levels of binary variable have data
  } else { # one or both levels of binary variable missing data
    warning("\n", "phylobLR() has skipped the following dataset-variable triad:", "\n", "\t", "dataset:", dataset_name, "\n", "\t", "predictor:", predictor, "\n", "\t", "response variable:", response, "\n", "REASON: At least one level of the binary response variable is missing data.)", "\n")
    bLR_output <- "phybLR() skipped because at least one level of the binary response variable is missing data."
    return(bLR_output)
    proceed_verdict <- "no" 
  } # closes if-else clause
  
  # If proceed_verdict remains "yes", the function will proceed.
  if(proceed_verdict == "yes"){
    # Define directory path for output files: 
    directory_path <- paste0(output_path, "/phybLR_models")
    # Create directory for output files:
    dir.create(path=directory_path, showWarnings = TRUE, recursive = FALSE, mode = "0777")
    
    # Make sure that phyloglm() is only run for response-predictor variable pairs with no overlapping levels, in which predictor values are present for both levels of the given response variable. Otherwise, an error will occur:
    l_dif <- length(setdiff(levels(ds_fin$response), levels(ds_fin$predictor)))
    l_lev <- length(levels(ds_fin$response))
    
    if (l_dif<l_lev) {
      warning("\n", "Phylogenetic binomial logistic regression (phylo-bLR)", "\n", "skipped for", response, "~", predictor, "in", dataset_name, " dataset due to", "\n",  "overlap in levels of response and predictor variables.",  "\n", "\n")
      bLR_output <- "phybLR() skipped due to overlap in levels of response and predictor variables."
      return(bLR_output)
    } else if (l_dif>=l_lev) { # that is, if they don't overlap...
      # We proceed to check if both levels of the response var have data:
      each_level <- tapply(ds_fin$predictor, ds_fin$response, FUN = length)
      if(sum(is.na(each_level))>0){ # If at least one level is missing data...
        warning("\n", "Phylogenetic binomial logistic regression (phylo-bLR)", "\n", "skipped for", response, "~", predictor, "in", dataset_name, "dataset", "\n", "due to absence of predictor variable values for at least one", "\n", "level of the response variable.",  "\n", "\n")
        bLR_output <- "phybLR() skipped due to absence of predictor values for at least one level of the binary response variable."
        return(bLR_output)
      } else if (sum(is.na(each_level))==0) { # If all levels have data...
        # Then we can proceed to run the phylo-bLR!
        
        # The first step is to train and test our phybLR model. We do this with a subsampling approach, by splitting our dataset randomly into thirds, using two thirds to train the model, and one third to test the model in order to assess its performance. We repeat this subsampling process n=CV_runs times, generating one ROC curve for each of the subsampling runs performed. We then average these ROC curves to get a consensus ROC curve. This consensus ROC. curve represents the results of repeated 'cross-validation'-like testing of our model, and we use it to describe model performance.
        
        # Set fold number (K) for cross validation:
        K <- Kfold 
        
        # Report cross-validation process:
        CV_bins <- c(paste("1/", K, sep=""), paste(K-1, "/", K, sep=""))
        cat("\n", "BEGUN: subsampling to train and test dataset with", CV_runs, "100 subsampling iterations...", "\n")
        cat("- for each iteration: using", CV_bins[2], "of data as training data,", CV_bins[1], "of data as testing data", "\n")
        
        # Create sequence of all thresholds to cycle through:
        thresholds <- Thr_seq <- seq(from=0, to=1, by = 0.01) 
        
        # Define n_obs, n_runs, and n_thresholds:
        n_obs <- length(ds_fin$response)
        n_runs <- CV_runs
        n_thresholds <- length(Thr_seq)
        
        # Create output lists to populate with cross-validation results:
        # Initialize output objects:
        td_ROCCs <- as.list(rep(NA, n_runs))
        McF_Rvals <- rep(NA, n_runs)
        McF_pvals <- rep(NA, n_runs)
        Zs <- as.list(rep(NA,length=K))
        # Populate output ROCC list with empty output matrix for each run:
        for(r in 1:length(td_ROCCs)){
          td_ROCCs[[r]] <- as.data.frame(matrix(ncol=2, nrow=n_thresholds))
          colnames(td_ROCCs[[r]]) <- c("TPR","FPR")} 
        
        # Perform subsampling 'cross-validation'-like procedure with 100 runs:
        for(r in 1:n_runs){
          # Initialize vectors to store sensitivity and specificity values
          sensitivity_values <- numeric(length(thresholds))
          specificity_values <- numeric(length(thresholds))
          # Define training and testing data sets:
          # Get rows to sample for subsampling procedure:
          Zs[[r]] <- sample(x=n_obs, size=n_obs/K, replace=FALSE) 
          Z <- Zs[[r]]
          testing_data <- ds_fin[Z, ] 
          training_data <- ds_fin[-Z, ]
          
          testing_data_levels <- length(unique(testing_data$response))
          training_data_levels <- length(unique(training_data$response))
          
          # Avoid situation where testing_data_ord$response has < 2 levels:
          # Give three chances for resampling. If it still has < 2 levels after three tries resampling, it will likely never obtain them. The if-loops below try resampling three times and then (if unsuccessful) report that resampling will not solve the problem and skip the current dataset-variable-tree combination. 
          if( testing_data_levels<2 | training_data_levels<2 ){
            cat("\n", "Warning: sampled testing or training data for run", r, "had <2 levels with data...", "\n", "Resampling for run", r, "now (attempt 1/3)...", "\n")
            # Get rows to sample for subsampling procedure:
            Zs[[r]] <- sample(x=n_obs, size=n_obs/K, replace=FALSE) 
            Z <- Zs[[r]]
            testing_data <- ds_fin[Z, ] 
            training_data <- ds_fin[-Z, ]
          } # closes 'if(testing_data_levels<2|training_data_levels<2)' clause
          testing_data_levels <- length(unique(testing_data$response))
          training_data_levels <- length(unique(training_data$response))
          
          if( testing_data_levels<2 | training_data_levels<2 ){
            cat("\n", "Warning: sampled testing or training data for run", r, "had <2 levels with data...", "\n", "Resampling for run", r, "now (attempt 2/3)...", "\n")
            # Get rows to sample for subsampling procedure:
            Zs[[r]] <- sample(x=n_obs, size=n_obs/K, replace=FALSE) 
            Z <- Zs[[r]]
            testing_data <- ds_fin[Z, ] 
            training_data <- ds_fin[-Z, ]
          } # closes 'if(testing_data_levels<2|training_data_levels<2)' clause
          testing_data_levels <- length(unique(testing_data$response))
          training_data_levels <- length(unique(training_data$response))
          
          if( testing_data_levels<2 | training_data_levels<2 ){
            cat("\n", "Warning: sampled testing or training data for run", r, "had <2 levels with data...", "\n", "Resampling for run", r, "now (attempt 3/3)...", "\n")
            # Get rows to sample for subsampling procedure:
            Zs[[r]] <- sample(x=n_obs, size=n_obs/K, replace=FALSE) 
            Z <- Zs[[r]]
            testing_data <- ds_fin[Z, ] 
            training_data <- ds_fin[-Z, ]
          } # closes 'if(testing_data_levels<2|training_data_levels<2)' clause
          testing_data_levels <- length(unique(testing_data$response))
          training_data_levels <- length(unique(training_data$response))
          
          if( testing_data_levels<2 | training_data_levels<2 ){
            warning("\n", "phylobLR() has skipped the following dataset-variable triad:", "\n", "\t", "dataset:", dataset_name, "\n", "\t", "predictor variable:", predictor, "\n", "\t", "response variable:", response, "\n", "REASON: Cross-validation infeasible (training or testing data missing data for ≥ 1 group levels).", "\n")
            bLR_output <- "phybLR() skipped because subsampling for cross-validation was infeasible (training or testing data missing data for ≥ 1 group levels)."
            return(bLR_output)
            still_proceed_verdict <- "no"
          } # closes 'if( testing_data_levels<2|training_data_levels<2)' clause
          testing_data_levels <- length(unique(testing_data$response))
          training_data_levels <- length(unique(training_data$response))
          
          if( testing_data_levels==2 & training_data_levels==2 ){
            if(crossval_reports==TRUE){
              cat("\n", "Model testing and training data selected for current run of cross-validation.", "\n", "Both datasets have 2 levels of the binary grouping_var. Commencing run", r, "...", "\n")
            }
            still_proceed_verdict <- "yes"
          }
          
          if(still_proceed_verdict=="yes"){
            # Trim and align trees to the new training and testing datasets:
            CV_testing_phylo <- trim.tree(tree=phylo_tr, trimmed_dataset=testing_data, verbose=F)
            testing_data_ord <- reorder.data(trimmed_tree=CV_testing_phylo, trimmed_dataset=testing_data, verbose=F)
            CV_training_phylo <- trim.tree(tree=phylo_tr, trimmed_dataset=training_data, verbose=F)
            
            # Remove NaN & 0 Myr branch lengths from testing & training trees:
            for(l in 1:length(CV_training_phylo$edge.length)){
              if(is.na(CV_training_phylo$edge.length)[l]==T){CV_training_phylo$edge.length[l] <- 0.1}
              if(CV_training_phylo$edge.length[l]==0){CV_training_phylo$edge.length[l] <- 0.1}
            } #closes 'for(l in 1:length(CV_training_phylo$edge.length))' loop
            
            for(l in 1:length(CV_testing_phylo$edge.length)){
              if(is.na(CV_testing_phylo$edge.length)[l]==T){CV_testing_phylo$edge.length[l] <- 0.1}
              if(CV_testing_phylo$edge.length[l]==0){CV_testing_phylo$edge.length[l] <- 0.1}
            } #closes 'for(l in 1:length(CV_testing_phylo$edge.length))' loop
            
            training_data_ord <- reorder.data(trimmed_tree=CV_training_phylo, trimmed_dataset=training_data, verbose=F)
            
            # Train the model with the training dataset:
            trained_mod <- phyloglm(response ~ predictor, data=training_data_ord, phy=CV_training_phylo, boot=boot_num, btol=btol_num, method = Firth_method) 
            
            # Test the model with the testing dataset to get TPR & FPR values:
            # Extract coefficients from the trained model
            trained_mod_coefs <- coef(trained_mod)
            # Compute linear predictors:  
            linear_predictors <- testing_data_ord$predictor * trained_mod_coefs[2] + trained_mod_coefs[1]
            # Use logistic func to calc predicted probabilities for testing_data:
            # logistic function: predProb = 1 / (1 + e ^(−linear predictor))
            e <- exp(1) # outputs the number 'e'
            predProbs_td <- 1 / (1 + e^(-(linear_predictors))) 
            # uses logistic function to get predProbs
            
            # Get TPR and FPR values for each threshold:
            for (t in seq_along(thresholds)) {
              # Set current threshold value:
              threshold <- thresholds[t]
              # Calculate sensitivity and specificity for the current threshold
              sensitivity <- sum(testing_data_ord$response == 1 & predProbs_td >= threshold) / sum(testing_data_ord$response == 1)
              specificity <- sum(testing_data_ord$response == 0 & predProbs_td < threshold) / sum(testing_data_ord$response == 0)
              # Store sensitivity and specificity values
              sensitivity_values[t] <- sensitivity
              specificity_values[t] <- specificity
            } # closes 'for (t in seq_along(thresholds))' loop
            
            # Save TPR and FPR for this run of model testing:
            td_ROCCs[[r]][, "TPR"] <- sensitivity_values
            td_ROCCs[[r]][, "FPR"] <- 1-specificity_values
            
            # Next, we calculate McFadden's pseudo-Rsq for each trained model (as an ROC-independent check of model fit). McFadden's pseudo-Rsq is equal to 1 - logLik(full model)/logLik(intercept-only model). More information on this method can be found in Smith et al. 2021 (https://doi.org/10.1080/02664763.2020.1796940).
            # The full model is the trained one, fit using all model params:
            fullmod <- trained_mod 
            # The null model is fit using only the intercept term:
            nullmod <- phyloglm(response ~ 1, data=training_data_ord, phy=CV_training_phylo) 
            # Calculate R-val for McFadden's pseudo-Rsq:
            logLik_mod <- logLik(fullmod)
            logLik_null <- logLik(nullmod)
            logLik_ratio <- (logLik_mod[[1]] / logLik_null[[1]])
            McF_Rvals[r] <- (1 - logLik_ratio)
            # Calculate p-val for McFadden's pseudo-Rsq:
            test_statistic <- (logLik(fullmod)[[1]] - logLik(nullmod)[[1]]) 
            # This test statistic is the difference in logLik values between the fitted and null models.
            degrees_freedom <- logLik(fullmod)[[2]] - logLik(nullmod)[[2]] 
            # This is the difference in degrees of freedom between the fitted and null models.
            McF_pvals[r] <- (1 - pchisq(test_statistic, degrees_freedom)) 
            # This is the p-val derived from a chi-squared distribution of the test_statistic.
            
            # Optionally report completion of current cross-validation run:
            if(crossval_reports==TRUE){cat("\n", "Run", r, "of cross-validation complete.", "\n")}
          } # closes "if(still_proceed_verdict=='yes')" loop
        } # closes "for(r in 1:n_runs)" loop
        
        # If all runs were completed successfully, tell R to proceed to get avg performance metrics:
        if(r==CV_runs){
          still_still_proceed_verdict <- "yes"
        } else {
          still_still_proceed_verdict <- "no"
        } # closes 'if(r==CV_runs) { } else { }' clause
        
        if(still_still_proceed_verdict=="yes"){    
          # Initialize ROC obj to store avg and 95% confidence interval TPR and FPR values for each threshold across iterations:
          td_ROCC_avg <- as.data.frame(matrix(nrow=n_thresholds, ncol=2))
          td_ROCC_CIs <- as.data.frame(matrix(nrow=n_thresholds, ncol=4))
          colnames(td_ROCC_avg) <- c("TPR", "FPR")
          colnames(td_ROCC_CIs) <- c("TPR_low", "TPR_high", "FPR_low", "FPR_high")
          
          # Perform tdROCC averaging and CI generation to make cROCC:
          vals_across_runs_per_row <- list(rep(NA, length=n_thresholds))
          for(t in 1:n_thresholds){
            vals_across_runs_per_row[[t]] <- as.data.frame(matrix(nrow=n_runs, ncol=2))
            colnames(vals_across_runs_per_row[[t]]) <- c("TPR", "FPR")
          } # closes 'for(t in 1:n_thresholds)' loop
          
          # Get and save avg TPR & FPR values for each threshold across runs:
          for (t in 1:n_thresholds) {
            for (r in 1:n_runs) {
              # Store TPR,FPR pair for each row across runs in their own matrix:
              vals_across_runs_per_row[[t]][r, ] <- td_ROCCs[[r]][t, ] }
            # Average each column of that matrix and save output to td_ROCC_avg
            td_ROCC_avg[t,"TPR"]<-mean(vals_across_runs_per_row[[t]][, "TPR"])
            td_ROCC_avg[t,"FPR"]<-mean(vals_across_runs_per_row[[t]][, "FPR"])
          } # closes 'for (t in 1:n_thresholds)' loop
          
          # Save 95% CIs for TPR & FPR values for each threshold across runs:
          for (t in 1:n_thresholds) {
            for (r in 1:n_runs) {
              # Store TPR,FPR pair for each row across runs in their own matrix:
              vals_across_runs_per_row[[t]][r, ] <- td_ROCCs[[r]][t, ] }
            
            # # Compute the sample size for a given threshold:
            sample_size <- CV_runs # should equal number of runs
            
            # Find the standard deviation of TPR and FPR across thresholds:
            sd_TPR <- sd(vals_across_runs_per_row[[t]]$TPR)
            sd_FPR <- sd(vals_across_runs_per_row[[t]]$FPR)
            
            # Find the standard error for TPR and FPR across thresholds:
            se_TPR <- sd_TPR / sqrt(sample_size)
            se_FPR <- sd_FPR / sqrt(sample_size)
            
            # Calculate the t_score and then margin_error for each parameter:
            alpha <- 0.05
            degrees_of_freedom <- sample_size - 1
            t_score <- qt(p=alpha/2, df=degrees_of_freedom,lower.tail=F)
            margin_error_TPR <- t_score * se_TPR
            margin_error_FPR <- t_score * se_FPR
            
            # Use margin_error to calculate 95% CI bounds:
            lower_bound_TPR <- td_ROCC_avg[t, "TPR"] - margin_error_TPR
            upper_bound_TPR <- td_ROCC_avg[t, "TPR"] + margin_error_TPR
            lower_bound_FPR <- td_ROCC_avg[t, "FPR"] - margin_error_FPR
            upper_bound_FPR <- td_ROCC_avg[t, "FPR"] + margin_error_FPR
            
            # Save 95% CI bounds for downstream plotting:
            td_ROCC_CIs[t, "TPR_low"] <- lower_bound_TPR
            td_ROCC_CIs[t, "TPR_high"] <- upper_bound_TPR
            td_ROCC_CIs[t, "FPR_low"] <- lower_bound_FPR
            td_ROCC_CIs[t, "FPR_high"] <- upper_bound_FPR
          } # closes 'for (t in 1:n_thresholds)' loop
          
          if(plot_td_ROCCs==TRUE){ 
            # Then make a plot of all ROC curves across runs:
            # First, initialize geom plot objects as text, with geom_line text making one curve for each subsampling iteration:
            geom_lines_perRun_text <- ""
            for(r in 1:length(td_ROCCs)){ geom_lines_perRun_text <- paste(geom_lines_perRun_text, " + geom_line(aes(x=td_ROCCs[[", r, "]]$FPR, y=td_ROCCs[[", r, "]]$TPR), color='red', alpha=0.4)", sep="") } # closes geom_lines_perRun_text-populating loop
            
            # Paste the geom_line_set_text into a larger text string that contains all the other commands required to make a nice ggplot:
            ggrocruns_text <- paste("ggplot(td_ROCCs[[1]], aes(x = FPR, y = TPR))", geom_lines_perRun_text, "+ geom_abline(slope = 1, intercept = 0, linetype = 'dashed', color = 'black', alpha=0.5) + geom_line(aes(x=td_ROCC_avg$FPR, y=td_ROCC_avg$TPR), linetype='solid', color='black', size=1.2, alpha=0.8) + labs(title = 'ROCCs from 100 sumsampling iterations with data split into K=", Kfold, "bins per iteration', x = 'False Positive Rate (FPR)', y = 'True Positive Rate (TPR)') + theme(legend.position='none', plot.title=element_text(size=5))", sep="")
            # Convert the ggrocruns_text string into a ggplot object:
            ggrocruns <- eval(parse(text=ggrocruns_text))
            
            # Save the ggrocruns plot as a PNG file:
            png(filename = paste(directory_path, "/", "bLR_", dataset_name, "__", response, "_by_", predictor, "_for_", tree_name, "__testing_ROCs.png", sep=""), width=12.5, height=12.5, units="in", res=300)
            show(ggrocruns)
            dev.off()
            
            cat("\n", "A plot of the ROC curves from all runs of cross-validation has been saved to the working directory.", "\n")
          } # closes 'if(plot_td_ROCCs==TRUE)' clause
          
          cat("\n", "Collecting performance metrics from consensus ROC curve...", "\n")
          # Get values for the diagonal "random guess line" in ROC space:
          ROC_diag_xvals <- seq(0, 1, length.out = 1000)
          ROC_diag_yvals <- ROC_diag_xvals # the x and y vals are equivalent
          
          # Generate consensus ROC output for tested model:
          n_obs <- length(ds_fin$response)
          ROC_yvals <- TPRs <- td_ROCC_avg$TPR
          ROC_xvals <- FPRs <- td_ROCC_avg$FPR
          ROC_95ci_lowbound_yvals <- td_ROCC_CIs$TPR_low
          ROC_95ci_highbound_yvals <- td_ROCC_CIs$TPR_high
          ROC_95ci_lowbound_xvals <- td_ROCC_CIs$FPR_low
          ROC_95ci_highbound_xvals <- td_ROCC_CIs$FPR_high
          
          # Get AUC value for consensus ROC curve:
          AUC <- abs(trapz(x=td_ROCC_avg$FPR, y=td_ROCC_avg$TPR))
          
          # Get the AUC range and a 95% CI across runs:
          AUC_vals <- rep(NA, length(n_runs))
          for(r in 1:n_runs){
            AUC_vals[r]<-abs(trapz(x=td_ROCCs[[r]]$FPR, y=td_ROCCs[[r]]$TPR))}
          AUC_range <- range(AUC_vals, na.rm=T)
          AUC_confint <- quantile(AUC_vals, c(0.025, 0.975), na.rm=T)
          AUC_range_result <- paste(round(AUC_range[1], 3), "-", round(AUC_range[2], 3), sep="")
          AUC_confint_result <- paste(round(AUC_confint[1], 3), "-", round(AUC_confint[2], 3), sep="")
          
          # Get mean & 95% CI values for Rsq and assoc. p-val of trained mods:
          McF_R_mean <- round(mean(McF_Rvals, na.omit=T), 3)
          McF_p_mean <- mean(McF_pvals, na.omit=T)
          McF_R_confint <- quantile(McF_Rvals, c(0.025, 0.975))
          McF_p_confint <- quantile(McF_pvals, c(0.025, 0.975))
          McF_R_confint_result <- paste0(round(McF_R_confint[1], 3), "-", round(McF_R_confint[2]), 3)
          McF_p_confint_result <- paste0(round(McF_p_confint[1], 3), "-", round(McF_p_confint[2]), 3)
          # Get significance-report vector for McF_R values:
          significance_report_vec <- rep(NA, length(McF_Rvals)) # This vector will be used to color-code McFadden's Rsq values by significance of fit (green for sig, red for non-sig) in the next plot.
          for(r in 1:length(McF_pvals)){
            if(McF_pvals[r]<0.05){
              significance_report_vec[r] <- "green2"
            } else {
              significance_report_vec[r] <- "red2"
            }
          } # closes 'for(r in 1:length(McF_pvals))' loop
          
          if(plot_McF==TRUE){ # Make a plot of AUC vs McFadden's pseudo-Rsq across runs to see how they relate.
            # Save McFadden plot as PNG file:
            png(filename = paste0(directory_path, "/", "bLR_", dataset_name, "__", response, "_by_", predictor, "_for_", tree_name, "__AUC_vs_McF_Rsq.png"), width=6, height=6, units="in", res=300)
            plot(x=AUC_vals, y=McF_Rvals, type="p", pch=21, col='grey10', bg=significance_report_vec, xlab="AUC values across runs", ylab="McFadden's pseudo-Rsq values across runs", main = paste("bLR_", dataset_name, "_", response, "_by_", predictor, "_for_", tree_name, " | ", "AUC values vs. McF R-vals |", sep=""), cex.main=0.5)
            dev.off()
            cat("\n","A plot of AUC vs. McFadden R^2 across cross-validation runs has been saved to the working directory.", "\n")
          } # closes 'if(plot_McF==TRUE)' clause
          
          # Get Youden's J stats for consensus ROC curve:
          Jvals <- TPRs - FPRs # J = sensitivity + specificity - 1
          J_thresholds <- Thr_seq # corresponding threshold val for each J val
          J_data <- as.data.frame(cbind(Jvals, J_thresholds))
          Jmax <- Jvals[which.max(Jvals)]
          
          # Use Jmax to get best predicted probability threshold & model stats:
          best_threshold <- Thr_seq[which.max(Jvals)]
          bestmod_TPR_mean <- TPRs[which.max(Jvals)]
          bestmod_FPR_mean <- FPRs[which.max(Jvals)]
          
          if(plot_Youden==TRUE){ # Make a plot of Youden's J statistic values vs. predProb thresholds:
            Youden_plot <- ggplot(data=J_data, aes(x=J_thresholds, y=(Jvals))) + geom_line(size=1.2) + 
              geom_text(aes(x = 0.1, y = Jmax, label = paste("J max = ", round(Jmax, 3), sep="")), vjust = -1, size=3) + 
              geom_text(aes(x = best_threshold-0.025, y = 0.13, label = paste("best threshold = ",round(best_threshold, 3), sep="")), size=3, angle=+90) +
              # Add dotted lines showing max threshold and Youden values:
              geom_segment(aes(x = 0, y = Jmax, xend = best_threshold, yend = Jmax), color = "green2", linetype="dotted", alpha=0.9) + 
              geom_segment(aes(x = best_threshold, y = 0, xend = best_threshold, yend = Jmax), color = "green2", linetype="dotted", alpha=0.9) + 
              geom_point(aes(x = best_threshold, y = Jmax), shape = 21, color = 'black', fill = 'green2', size=4) + 
              ylim(c(0, 1)) +
              labs(x = "Threshold", y = "Youden's Index") +
              ggtitle(paste("bLR_",dataset_name, "_",response, "_by_",predictor, "_for_",tree_name, " | ", "Youden values", sep=""))
            
            # Save Youden plot as PNG file:
            png(filename = paste0(directory_path, "/", "bLR_", dataset_name, "__", response, "_by_", predictor, "_for_", tree_name, "__Youden.png"), width=6, height=6, units="in", res=300)
            show(Youden_plot)
            dev.off()
            cat("\n","A plot of Youden's J-statistic values across thresholds has been saved to the working directory.", "\n")
          } # closes 'if(plot_Youden==TRUE)' clause
          
          # Get range and 95% CIs for the TPR and FPR of this best cROC model:
          best_thresh_index <- which(Thr_seq==best_threshold)
          bestmod_allvals <-  vals_across_runs_per_row[[best_thresh_index]]
          bestmod_FPRs <- bestmod_allvals$FPR
          bestmod_TPRs <- bestmod_allvals$TPR
          bestmod_FPR_range <- range(bestmod_FPRs)
          bestmod_TPR_range <- range(bestmod_TPRs)
          bestmod_FPR_95ci <- quantile(bestmod_FPRs, c(0.025, 0.975))
          bestmod_TPR_95ci <- quantile(bestmod_TPRs, c(0.025, 0.975))
          bestmod_CIs<-as.data.frame(cbind(bestmod_FPR_95ci, bestmod_TPR_95ci))
          colnames(bestmod_CIs) <- c("bestmod_FPRci", "bestmod_TPRci")
          
          if(plot_CIs==TRUE){ 
            # Make plot for 95% CIs for the TPR and FPR of best cROC model:
            bestmod_plot <- ggplot() + geom_errorbar(aes(ymin = bestmod_TPR_95ci[1], ymax = bestmod_TPR_95ci[2], x = mean(bestmod_FPRs)), color = 'grey10', alpha=0.8, linewidth=2, width=0.005) + geom_errorbarh(aes(xmin = bestmod_FPR_95ci[1], xmax = bestmod_FPR_95ci[2], y = mean(bestmod_TPRs)), color = 'grey10', alpha=0.8, linewidth=2, height=0.01) + geom_point(mapping=aes(x=mean(bestmod_FPRs), y=mean(bestmod_TPRs)), pch=19, size=10, color='grey10') + geom_point(mapping=aes(x=mean(bestmod_FPRs), y=mean(bestmod_TPRs)), pch=19, size=7, color='green') + labs(title = "95% CIs of model w/ best threshold ", x = "FPR", y = "TPR") + lims(x=c(min(bestmod_FPR_range), max(bestmod_FPR_range)), y=c(min(bestmod_TPR_range), max(bestmod_TPR_range)))
            
            # Save bestmod CI plot as PNG file:
            png(filename = paste0(directory_path, "/", "bLR_", dataset_name, "__", response, "_by_", predictor, "_for_", tree_name, "__bestmod_CIs.png"), width=6, height=6, units="in", res=300)
            show(bestmod_plot)
            dev.off() 
            cat("\n", "A plot of FPR & TPR CIs for the bestmod (cROCC given best threshold) has saved to the working directory.", "\n")
          } # closes 'if(plot_CIs==TRUE)' clause
          
          cat("\n", "Running phylogenetic binomial logistic regression with full dataset...", "\n") 
          # In machine learning, a model is trained on one subset of known data, and then tested on a separate subset of known data to assess model performance. After model performance is assessed, the assessed model is re-trained on ALL the data (without subsetting) in order to make full use of the known dataset. The model fit with ALL the data is then used to make new predictions. Following this approach, we trained the phybLR model above and tested it with separate subsets of our data to generate a report (in the form of a consensus ROC curve) for the model's performance. However, to make predictions on contentious fossil taxa, we must refit the model based on ALL the data. We do this next step below.
          
          # Fit the true model with the predictor variable and ALL the data:
          bLR_mod <- phyloglm(response ~ predictor, data=ds_fin, phy=trimmed_tree, boot=boot_num, btol=btol_num, method = Firth_method) 
          
          # Fit the null model with only the intercept term and ALL the data:
          nullmod <- phyloglm(response ~ 1, data=ds_fin, phy=trimmed_tree)
          
          # SAVE MODEL METADATA:
          cat("\n", "Saving text file of phybLR model metadata...", "\n")
          assessment_report_filepath_and_filename <- paste0(directory_path, "/", "bLR_", dataset_name, "__", response, "_by_", predictor, "_for_", tree_name, "__model_metadata.txt")
          # Define output text file with all model metadata:
          sink(assessment_report_filepath_and_filename)
          # Write introduction for output file:
          cat("\n", "-------------------------------------------------------------------", "\n", "Phylogenetic Binomial Logistic Regression (phylobLR) Metadata", "\n", "-------------------------------------------------------------------", "\n", "DATASET:", dataset_name, "\n", "QUANTITATIVE PREDICTOR VARIABLE:", predictor, "\n", "BINARY RESPONSE VARIABLE:", response, "\n", "TREE:", tree_name, "\n", "-------------------------------------------------------------------")
          # Report model parameters:
          cat("\n", "MODEL PARAMETERS:", "\n", "• Number of observations:", length(ds_fin$response), "\n", "• Bootstraps =", boot_num, "\n",
              "• btol =", btol_num, "\n", "• phyloglm() 'method':", Firth_method, "\n",  "-------------------------------------------------------------------")
          # Report model coefficients:
          coefficients <- coef(bLR_mod)
          cat("\n", "MODEL COEFFICIENTS:", "\n", coefficients, "\n", "-------------------------------------------------------------------")
          # Report model 'fit' statistics:
          cat("\n", "MODEL 'FIT' AND 'SELECTION' METRICS:", "\n")
          # Calculate McFadden's pseudo-Rsq for the full data set:
          # Recall that McFadden's Rsq = 1 - logLik(model) / logLik(null).
          logLik_mod <- logLik(bLR_mod)
          logLik_null <- logLik(nullmod)
          logLik_ratio <- (logLik_mod[[1]] / logLik_null[[1]])
          McF_Rsq <- round((1 - logLik_ratio), 3)
          # Calculate p-val for McFadden's pseudo-Rsq:
          test_statistic <- (logLik(bLR_mod)[[1]] - logLik(nullmod)[[1]]) 
          # this test statistic is the diff in log likelihoods btwn models
          degrees_freedom <- logLik(bLR_mod)[[2]] - logLik(nullmod)[[2]] 
          # this is the diff in degrees of freedom btwn models
          McF_pval <- (1 - pchisq(test_statistic, degrees_freedom))
          # This is the p-val derived from a chisq distrib of the test stat.
          # Report the McF Rsq results:
          cat("• McFadden's pseudo R-squared: mean Rsq for training models=", McF_R_mean, ";", "95% CI:", McF_R_confint_result, "\n")
          cat("• McFadden's pseudo R-squared: mean p-val for training models=", McF_p_mean, ";", "95% CI:", McF_p_confint_result, "\n")
          cat("• McFadden's pseudo R-squared for model fit with ALL the data=", McF_Rsq, ";", "p =", McF_pval, "\n")
          
          # Report the Akaike Information Criterion:
          cat("• Akaike Information Criterion (AIC):", bLR_mod$aic, "\n", "-------------------------------------------------------------------", "\n")
          
          # Run one-sample binomial test to see if AUC significantly > 0.5:
          binom_output <- binom.test(x = round(AUC * n_obs), n = n_obs, p = 0.5, alternative = "greater", conf.level=0.95)
          AUC_binom_pval <- binom_output$p.value
          AUC_binom_ci <- binom_output$conf.int
          
          # Report cROCC parameters (AUC, Jmax, and best threshold):
          cat("\n", "CONSENSUS RECEIVER OPERATING CHARACTERISTIC CURVE ('cROCC') PARAMETERS:", "\n", "\n", "• DESCRIPTION: To train and test our model, we performed a subsampling procedure with", CV_runs, "runs. For each run, the dataset was randomly split into thirds. First, an initial bLR model was trained on 2/3 of the dataset. Then, the trained model was tested on the remaining 1/3 of the dataset to generate an ROC curve. The resulting", CV_runs, "ROC curves were averaged (individual coordinate pairs averaged across runs) to produce a consensus ROC curve (described below) that summarizes model performance.", "\n", "\n", "• Area Under the Curve (AUC):", AUC, "| 95% CI:", AUC_confint_result, ";", "full range:", AUC_range_result, "\n", "\n", "• ONE-SAMPLE BINOMIAL TEST FOR ROC CURVE:", "\n", "--- Description: Tests whether a given phybLR model makes accurate predictions significantly more than 50% of the time across thresholds. This is equivalent to testing whether the ROC's AUC for this model is significantly greater than that of the null model (0.5).", "\n", "--- Binomial Exact Test p-value: p =", AUC_binom_pval, "\n", "--- Binomial test interpretation: If p<0.05, then this ROC's AUC is significantly greater than 0.5.", "\n", "\n", "• IDEAL PREDICTED PROBABILITY THRESHOLD:", "\n", "--- Selection method: Youden's J-statistic", "\n", "--- Highest Youden's J value (Jmax) =", round(Jmax, 3), "\n", "--- Best predicted probability threshold =", round(best_threshold, 3),  "\n", "--- Conclusion:", round(best_threshold, 3), "is the threshold that most accurately classifies specimens for this dataset-variable triad.", "\n", "\n", "-------------------------------------------------------------------", "\n")
          
          # Compute accuracy metrics for best model:
          P <- sum(ds_fin$response == 1)
          N <- sum(ds_fin$response == 0)
          n_obs <- length(ds_fin$response)
          TPR <- bestmod_TPR_mean
          TP <-  round(TPR*P,0) # Since TPR = TP/P
          FPR <- bestmod_FPR_mean
          FP <- round(FPR*N,0) # Since FPR = FP/N
          TNR <- 1 - FPR
          TN <-  round(TNR*N,0) # Since TNR = TN/N 
          n_successes <- TP + TN
          FNR <- 1-TPR #(wikipedia)
          FN <- round(FNR*P,0) # Since FNR = FN/P
          n_failures <- FP + FN
          Accuracy <- round(((TP+TN)/(P+N)), 2)
          
          # Report accuracy metrics for best model:
          cat("ROC PERFORMANCE METRICS FOR BEST MODEL", "\n", "Description: Performance metrics derived from consensus ROC curve, for model with best predicted probability threshold.", "\n", "• P =", P, "; N =", N, "\n", "• TP =", TP, ";", "TN =", TN, ";", "FP =", FP, ";", "FN =", FN, "\n", "• Number of successes = TP + TN =", n_successes, "\n", "• Number of failures = FP + FN =", n_failures, "\n", "• TPR =", round(TPR, 3), "\n", "• FPR =", round(FPR, 3), "\n", "• TNR =", round(TNR, 3), "\n", "• FNR =", round(FNR, 3), "\n", "• Accuracy = ((TP+TN)/(P+N)) =", Accuracy, "\n", "-------------------------------------------------------------------", "\n")
          
          # Perform Binomial Exact Test to see whether best model (with best threshold) makes more accurate predictions than random guesswork :
          ntrials <- length(ds_fin$response)
          prob_to_beat <- 0.75
          binom_test <- binom.test(x=c(n_successes, n_failures),p=prob_to_beat, alternative = "greater")
          bestmod_binom_pval<-format(binom_test$p.value, scientific=T,digits=5)
          # Report the Binomial Exact Test results:
          cat("ONE-SAMPLE BINOMIAL TEST FOR BEST MODEL:", "\n", "• Description: Tests whether best-performing phybLR model makes accurate predictions significantly more often than 75% of the time.", "\n", "• Binomial test p-value: p =", bestmod_binom_pval, "\n", "• INTERPRETATION: if p<0.05, the model is accurate significantly more than 75% of the time.", "\n", "-------------------------------------------------------------------", "\n")
          
          # Close the external connection to output file:
          sink(); sink(file=NULL) # Finicky behavior seen when running; adding twice prevented glitches.
        } # closes "else if (sum(is.na(each_level))==0)" loop
      } # closes "else if (l_dif>=l_lev)" loop
      # Report completion of function for a given set of inputs:
      cat("\n", "phyblR() function complete!", "\n")
      cat("\n", "-------------------------------------------------------------------", "\n", "DATASET-VARIABLE-TREE COMBO:", dataset_name, "|", response, "by", predictor, "for", tree_name, ":", "\n", "• phybLR object saved to environment as named output", "\n", "• Plot of testing ROCCs for model performance assessment saved to working directory.", "\n", "• phybLR model metadata saved to working directory", "\n", "• Plot of cROCC Youden's index vs. threshold saved to working directory.", "\n", "-------------------------------------------------------------------", "\n") 
      # Save function output to an R data object of class "phylobLR":
      # Initialize object:
      bLR_output <- list()
      # Populate object with model data:
      bLR_output$summary <- cat("\n", "MODEL: phylogenetic binomial logistic regression (phybLR) with", boot_num, "bootstrap reps", "\n", "DATASET-VARIABLE-TREE COMBO:", dataset_name, "|", response, "by", predictor, "for", tree_name, "\n", "METHOD FOR ASSESSING MODEL PERFORMANCE:", "\n", "---Assessed consensus ROC curve (cROCC) generated from 100 subsampling runs.", "\n", "---Each run: phybLR model trained on", CV_bins[2], "of the data, tested on the remaining",  CV_bins[1], "\n", "---All", CV_runs, "ROC output curves were averaged to generate cROCC.", "\n")
      bLR_output$phybLR_mod <- bLR_mod
      bLR_output$input_dataset <- dataset_name
      bLR_output$input_tree <- tree_name
      bLR_output$response_variable <- response
      bLR_output$predictor_variable <- predictor
      bLR_output$cROC_xvals <- ROC_xvals  
      bLR_output$cROC_yvals <- ROC_yvals
      bLR_output$ROC_95ci_lowbound_xvals <- ROC_95ci_lowbound_xvals
      bLR_output$ROC_95ci_lowbound_yvals <- ROC_95ci_lowbound_yvals
      bLR_output$ROC_95ci_highbound_xvals <- ROC_95ci_highbound_xvals
      bLR_output$ROC_95ci_highbound_yvals <- ROC_95ci_highbound_yvals
      bLR_output$AUC_mean <- AUC
      bLR_output$AUC_95ci <- AUC_confint
      bLR_output$AUC_range <- AUC_range_result
      bLR_output$AUC_binomtest_pval <- AUC_binom_pval
      bLR_output$McF_R_mean <- McF_R_mean
      bLR_output$McF_p_mean <- McF_p_mean
      bLR_output$McF_R_confint <- McF_R_confint
      bLR_output$McF_p_confint <- McF_p_confint
      bLR_output$McF_Rsq_for_full_ds <- McF_Rsq
      bLR_output$McF_pval_for_full_ds <- McF_pval
      bLR_output$AIC <- bLR_mod$aic
      bLR_output$Youdens_Jmax <- Jmax
      bLR_output$bestmod_threshold <- best_threshold
      bLR_output$bestmod_binomtest_pval <- bestmod_binom_pval
      bLR_output$n_obs <- n_obs
      bLR_output$bestmod_TPR_mean <- bestmod_TPR_mean
      bLR_output$bestmod_TPR_range <- bestmod_TPR_range
      bLR_output$bestmod_TPR_95ci <- bestmod_TPR_95ci
      bLR_output$bestmod_TP <- TP
      bLR_output$bestmod_FPR_mean <- bestmod_FPR_mean
      bLR_output$bestmod_FPR_range <- bestmod_FPR_range
      bLR_output$bestmod_FPR_95ci <- bestmod_FPR_95ci
      bLR_output$bestmod_FP <- FP
      bLR_output$bestmod_TNR_mean <- TNR
      bLR_output$bestmod_TN <- TN
      bLR_output$bestmod_nSuccesses <- n_successes
      bLR_output$bestmod_FNR <- FNR
      bLR_output$bestmod_FN <- FN
      bLR_output$bestmod_nFailures <- n_failures
      bLR_output$bestmod_Accuracy <- Accuracy
      bLR_output$full_assessment_report <- readLines(assessment_report_filepath_and_filename)
      return(bLR_output)
    } # closes if(still_still_proceed_verdict=="yes") loop
  } # closes if(proceed_verdict == "yes") loop
} # closes main phybLR() function loop
cat("\n","PROGRESS REP: chunk [S1.5.03] complete; starting next chunk..","\n")
#*-----**-----*

#*-----**{ [S1.5.04] Test phybLR() function to make sure it works. }*
# To make sure this function works, we test it below on a range of inputs that frequently produced errors in previous runs. 
# TEST INPUT SET #1:
  # Run upstream functions:
    bLR <- bLRify(all_data)
    bLR_tr <- rm.NAs(dataset=bLR, categorical_var="bin_fflip", quantitative_var="f_AZR")
    phylo_tr <- trim.tree(tree=tree_variants[[1]], trimmed_dataset=bLR_tr)
    ds_ord <- reorder.data(trimmed_tree=phylo_tr, trimmed_dataset=bLR_tr)
  # Test phybLR() function:
    test_bLR_output <- phybLR(dataset_ordered=ds_ord, dataset_name="***test_bLR_", response="bin_fflip", predictor="f_AZR", trimmed_tree=phylo_tr, tree_name = "phylo_tr", boot_num=1, btol_num=10, Kfold=3, CV_runs=5, crossval_reports=T, plot_td_ROCCs=T, plot_Youden=T, plot_McF=T, plot_CIs=T)
  # Verify that phybLR() output object looks right:
    names(test_bLR_output)

# TEST INPUT SET #2:
  # Run upstream functions:
    bLR_tr <- rm.NAs(dataset=bLR_raw_datasets[[1]], categorical_var=key_bLR_groupers[1], quantitative_var=ratio_vars[1])
    phylo_tr <- trim.tree(tree=tree_variants[[1]], trimmed_dataset=bLR_tr)
    ds_ord <- reorder.data(trimmed_tree=phylo_tr, trimmed_dataset=bLR_tr)
  # Test phybLR() function:
    ds_name_input<-"***test_all_bLR_"
    test_bLR_output <- phybLR(dataset_ordered=ds_ord, dataset_name=ds_name_input, response=key_bLR_groupers[1], predictor=ratio_vars[1], trimmed_tree=phylo_tr, tree_name = "phylo_tr", boot_num=1, btol_num=10, CV_runs=5, plot_td_ROCCs=T, plot_Youden=T, plot_McF=T, plot_CIs=T, Kfold = 3)
  
  # Verify that phybLR() output object looks right:
    names(test_bLR_output)

# TEST INPUT SET #3:
  # Run upstream functions:
    bLR_tr <- rm.NAs(dataset=bLR_raw_datasets[[2]], categorical_var=key_bLR_groupers[1], quantitative_var=ratio_vars[1])
    phylo_tr <- trim.tree(tree=tree_variants[[1]], trimmed_dataset=bLR_tr)
    ds_ord <- reorder.data(trimmed_tree=phylo_tr, trimmed_dataset=bLR_tr)
  # Run phybLR() function:
    test_bLR_output <- phybLR(dataset_ordered=ds_ord, dataset_name=bLR_raw_dataset_names[2], response=key_bLR_groupers[1], predictor=ratio_vars[1], trimmed_tree=phylo_tr, tree_name = "supertree1", boot_num=1, btol_num=10, Kfold=3, crossval_reports=F, plot_td_ROCCs=T, plot_McF=F, plot_Youden=F, plot_CIs=T, CV_runs=5,Firth_method="logistic_MPLE")

# Now that we've defined the phybLR() function to train and test a phylogenetic binomial logistic regression on the data, we can apply that function in our S4 scripts to train and test phybLR models for every dataset-variable-tree combination of interest.
cat("\n","PROGRESS REP: chunk [S1.5.04] complete; starting next chunk..","\n")
#*-----**-----*

#*-----**{ [S1.5.05] Save current R environment to working directory. }*
save.image(file=paste0(wd_path, "/R_ENVIR_for_script_S1.RData")) 
cat("\n","PROGRESS REP: chunk [S1.5.05] complete; starting next chunk...","\n")
#*-----**-----*

#*-----**{ [S1.5.06] Record the time that was required for script_S1 to run. }*
# Record running time metrics for entire script:
proctime_output <- proc.time()
elapsed_secs <- proctime_output[3]
elapsed_mins <- elapsed_secs/60
elapsed_hrs <- elapsed_mins/60
elapsed_days <- elapsed_hrs/24
cat("\n","PROGRESS REP: chunk [S1.5.06] complete; starting next chunk...","\n")
#*-----**-----*

#*-----**{ [S1.5.07] Record CPU usage for this script. }*
# The following code calculates CPU usage efficiency based on the following Stack Overflow thread: https://stackoverflow.com/questions/73217110/cpu-usage-measurement-and-calculation-in-r-using-proc-time.
CPU_number <- CPU_num # Recall that this is the number of CPUs used for the current script.
CPU_time_used <- proctime_output[1] + proctime_output[2]
CPU_utilization <- (proctime_output[1] / CPU_time_used) * 100 / CPU_number
cat("\n","PROGRESS REP: chunk [S1.5.07] complete; starting next chunk...","\n")
#*-----**-----*

#*-----**{ [S1.5.08] Save time-elapsed and CPU metrics to a text file. }*
time_report_filepath <- paste0(output_path,"/","TIME_REPORT_for_script_S1.txt")
sink(time_report_filepath)
cat("TIME TAKEN FOR ENTIRE SCRIPT TO RUN", "\n", elapsed_secs, "seconds", "\n", "\t", "=", elapsed_mins, "minutes", "\n", "\t", "=", elapsed_hrs, "hours", "\n", "\t", "=", elapsed_days, "days", "\n")
cat("\n", "CPU USAGE:", "\n", "\t", "Number of CPUs used:",  CPU_number, "\n", "\t", "CPU utilization efficiency:", CPU_utilization, "%", "\n")
sink(file=NULL) # A text file saving the time elapsed for the completion of this script has now been saved to the working directory.
cat("\n","PROGRESS REP: chunk [S1.5.08] complete; starting next chunk...","\n")
#*-----**-----*

#*-----**{ [S1.5.09] Review next steps for script_S2. }*
# TO RUN SCRIPT_S2 AS A BATCH JOB, YOU WILL NEED TO RUN THE FOLLOWING COMMANDS FROM A HIGH-PERFORMANCE COMPUTING CLUSTER TERMINAL WINDOW:
#---# $ cd /path/to/your/working/directory/ # navigates to working directory
#---# $ sbatch _BASH_script_S2.sh # submits BASH script to run script_S2.R
cat("\n","PROGRESS REP: chunk [S1.5.09] complete; starting next chunk...","\n")
#*-----**-----*

#*-----**{ [S1.5.10] Review next steps for scripts S3 & S4. }*
# TO RUN SCRIPTS S3 AND S4 IN MASSIVELY PARALLEL FASHION USING JOB ARRAYS, YOU WILL NEED TO DO THE FOLLOWING FROM WITHIN A HIGH-PERFORMANCE COMPUTING CLUSTER TERMINAL WINDOW:

# (1) CHECK YOUR WORKING DIRECTORY TO MAKE SURE THAT THE JOB LIST OUTPUTS GENERATED ABOVE ARE FORMATTED CORRECTLY AND SPECIFY FEWER THAN 10,000 JOBS. Please note that 10,000 is the hard upper limit for Job Arrays on Yale's HPC.

# (2) RUN THE FOLLOWING COMMANDS TO GENERATE JOB-ARRAY BATCH SCRIPTS:
#...# $ cd /path/to/your/working/directory/ # navigates to working directory
#...# $ module load dSQ # loads dSQ--required for Job Arrays on Yale's HPC
#...# $ dsq --job-file script_S3_joblist.txt --mem-per-cpu 4g -t 12:00:00 --mail-type ALL --batch-file _BASH_script_S3.sh # uses dSQ to write a BASH script to run the S3 job array
#...# $ dsq --job-file script_S4_joblist.txt --mem-per-cpu 4g -p week -t 3- --mail-type ALL --batch-file _BASH_script_S4.sh # uses dSQ to write a BASH script to run the S4 job array

# (3) MAKE NEW SUBDIRECTORIES TO STORE JOB ARRAY OUTPUT REPORTS:
#...# $ mkdir script_S3_JobArray_reportFiles
#...# $ mkdir script_S4_JobArray_reportFiles

# (4) REVISE THE '--output' FLAG IN EACH BASH SCRIPT TO THE FOLLOWING:
#...# $ --output /home/cmg89/palmer_scratch/output_files/script_S3_array_%A/%N/slurm-%A_%a.out # Remember to change the output path to match that of your own scratch folder!
#...# $ --output /home/cmg89/palmer_scratch/output_files/script_S4_array_%A/%N/slurm-%A_%a.out # Remember to change the output path to match that of your own scratch folder!

# (5) SUBMIT THE FOLLOWING COMMANDS TO RUN YOUR JOB-ARRAY BATCH SCRIPTS:
#...# $ sbatch _BASH_script_S3.sh
#...# $ sbatch _BASH_script_S4.sh
cat("\n","PROGRESS REP: chunk [S1.5.10] complete; script_S1 is all done!","\n")
#*-----**-----*

# That concludes the first script in this data analysis pipeline. In this first script, we imported, cleaned, and processed all the LM data for this project; performed a quality-control assessment for LM measurements; read, time-calibrated, and aligned trees to the dataset for downstream phylogenetic statistical tests; began analyzing relationships among categorical variables; and defined job lists and custom functions for downstream scripts to run via batch jobs and Job Arrays. In the following scripts, we will make use of greater parallelization to rigorously assess how all LM data variables differ among groups within a phylogenetic framework (script_S2), systematically assess which quantitative LM variables are correlated (script_S3), and fit phylogenetic binomial logistic regression models to raw, log10-transformed, and BoxCox-transformed datasets (script_S4). Instructions for how to run these scripts from your own local laptop or high-performance computing (HPC) cluster are given in chunks [S1.5.09] and [S1.5.10] above. Once scripts S2, S3, and S4 have finished running, you can open and begin script_S5, in which you will use the combined outputs from all upstream scripts to reconstruct the evolutionary histories of aquatic habits and soft-tissue phenotypes in amniotes, generate some nice plots to visualize pairwise correlations, and run a comprehensive sensitivity analysis to investigate how tree topology and data transformation method alter the results of all previously run phylogenetic statistical tests.

# I hope you have enjoyed this coding journey so far, and that you continue onward to script_S5! In the meantime, if you have any questions about this script or its associated data, or if you would like to discuss a possible collaboration, please feel free to contact me anytime via the address below.

# -- Caleb Gordon
#------------------------------------------------------------------------------
```
