## Supplementary R Scripts for "Limb proportions predict aquatic habits and soft-tissue flippers in extinct amniotes": script_S2.R

```
#** Second R script (script_S2.R) for Gordon et al., "Limb proportions predict aquatic habits and soft-tissue flippers in extinct amniotes."**

# NOTE ON READABILITY: This script contains many long, multi-line comments. To increase readability within Rstudio, go to Tools > Global Options, select the 'Code' tab, and check 'Soft-wrap R source files'. This will wrap all lines of code to match your personal GUI's margins, so that no lines of code run offscreen.

# NOTE ON SCRIPT ORDER: This script is downstream of script_S1.R, upstream of script_S5, and concurrent (can be run in parallel) with scripts S3-S4. If you have not finished running script_S1.R, please close this file and run script_S1.R to completion. Once script_S1.R is done running, you can reopen the current file and proceed. 

# INTRODUCTION: In the previous script (script_S1), we ran all preliminary data- and tree-processing steps required for more intensive downstream statistical analyses. In the current script (script_S2), we perform the first set of those downstream analyses, which involves all phylogenetic tests comparing quantitative variables among groups: phylogenetic ANOVAs with pairwise post-hoc Tukey comparisons, phylogenetic Levene's tests with pairwise post-hoc Tukey comparisons, and boxplots and ternary plots to visualize associated univariate and multivariate differences in variable distributions among groups. This script is computationally intensive, and uses moderate parallelization (in the form of foreach-loops) to increase computation rate by distributing for-loop iterations across multiple CPUs. 

# NOTE ON OUTPUT VOLUME: Please be warned that this script produces thousands of output files (totaling about 2-4 Gigabytes) to your working directory.

# This is the outline of sections for script_S2:
#**[S2] COMPARE QUANTITATIVE DATA AMONG GROUPS**
# --- [S2.1]: PREPARE CODING ENVIRONMENT: Load packages, S1 R environment, etc.
# --- [S2.2]: DO PHYLOGENETIC ANOVAs & LEV. TESTs; MAKE ASSOCIATED BOXPLOTs.
# --- -- NOTE: This is repeated for raw, log-trans, and BoxCox-trans datasets.
# --- [S2.3]: MAKE GROUPED BOX PLOTS, TO SHOW HOW TRENDS DIFFER BY SUBCLADE.
# --- [S2.4]: COMPARE LIMB PROPORTIONS BY GROUP IN TERNARY MORPHOSPACE.
# --- [S2.5]: SAVE R ENVIRONMENT AS INPUT FOR KEY DOWNSTREAM SCRIPTS.

# We go through each of these sections with a series of makeshift code "chunks" below, starting with [S2.1].
#------------------------------------------------------------------------------

#**"-------------------------------------------------------------------------"*
#**"--------------- ### [S2.1] PREPARE CODING ENVIRONMENT ### ---------------"*
#**"-------------------------------------------------------------------------"*

#*-----**{ [S2.1.01] Load required packages. }*
# NOTE: Since all packages were installed for completion of script_S1, we do not repeat the optional installation lines for these packages here; however, lines of code to install all packages are provided in script_S1 [S1.1.01].

# Parallel programming on multiple CPU cores:
library(parallel)
library(foreach)
library(doMC)
library(doParallel)

# Data object manipulation:
library(forcats) # for releveling factors with fct_relevel() function
library(data.table) # for converting objects to data tables and matrices
library(plyr) # useful for in-task parallelization
library(dplyr) # useful for in-task parallelization
library(stringr) # contains str_detect() function

# Data visualization and exporting:
library(ggplot2) # for effective data visualization with ggplot suite
library(MESS) # for specifying translucent colors as function arguments
library(car) # for normal quantile plot construction
library(RColorBrewer) # for defining fancy color palettes
library(gridExtra) # useful for generating multi-panel plots and tables
library(ggimage) # For exporting ggplots
library(Ternary) # for ternary plot construction

# Data analysis:
library(devtools) # required for downloading some other packages from github
library(vcd) # for Cramer's V and other association stats
library(MASS) # used to make Box-Cox transformations with boxcox() function

# Phylogenetic comparative methods and tree visualization:
library(ape) # for basic tree manipulation and phylogenetic comparative methods
library(ggtree) # for advanced tree visualization using ggplot2 suite
library(strap) # contains DatePhylo() function to time-calibrate trees
library(phytools) # contains phylANOVA() function
library(geiger) # contains aov.phylo() function
library(phylolm) # contains phyloglm() and phylolm() functions
library(vegan) # contains adonis2(), which can run phyloPERMANOVAs
cat("\n","PROGRESS REP: chunk [S2.1.01] complete; starting next chunk...","\n")
#*-----**-----*

#*-----**{ [S2.1.02] Set up parallel programming environment. }*
# We will use the foreach package to run programs in parallel on multiple CPU cores. The registerDoMC() function below assigns the proper number of cores for this task. Before you begin running the code below, you should adjust the CPU_num value below to reflect the number of separate CPUs available to you on your personal computer or HPC cluster.

CPU_num <- 48 # 6 is the total number of available processing cores on my MacBook. 48 is the highest number that I can request from Yale's High-Performance Computing cluster.

# Set up parallel backend:
cl <- makeCluster(CPU_num, outfile = "") # outfile="" ensures that the output from parallelized tasks prints to the console
registerDoParallel(cl)
cat("\n","PROGRESS REP: chunk [S2.1.02] complete; starting next chunk...","\n")
#*-----**-----*

#*-----**{ [S2.1.03] Load script_S1 environment and set iter_num. }*
# Define working directory path:
wd_path <- "/gpfs/gibbs/project/bhullar/cmg89/Flipper_Project/"
setwd(wd_path); getwd()
# Load script_S1 environment from working directory:
load(file=paste0(wd_path, "/R_ENVIR_for_script_S1.RData"))

# Set iteration number for all simulation-based comparative phylogenetic tests:
iter_num <- 10000 # This will determine the number of iterations used when performing simulation-based phylogenetic ANOVAs. Increasing the number of iterations will drastically increase the time and computational resources required to run the script below. For troubleshooting this code on a local computer, setting iter_num=3 works well. While running on Yale's HPC and for the paper, we set iter_num=10000. 
cat("\n","PROGRESS REP: chunk [S2.1.03] complete; starting next chunk...","\n")
#*-----**-----*

# We have now set up the coding environment for script_S2. In the next section of the script, we will run the bulk of its data analysis. 
#------------------------------------------------------------------------------

#**"-------------------------------------------------------------------------"*
#**"-- ### [S2.2] DO PHYLOGENETIC ANOVAs, LEVENE'S TESTs, & POST-HOCs ### ---"*
#**"-------------------------------------------------------------------------"*

# We will now generate box plots and run several statistical tests (phylogenetic ANOVAs, phylogenetic Levene's tests, and pairwise post-hoc Tukey comparisons) to compare each key numeric variable among each grouping variable in our data set. Box plots are tree-independent, and one will be generated for each response variable grouping variable pair. Statistical test results depend on a particular tree topology, and will thus be run once for each tree variant.

#*-----**{ [S2.3.02] Write internal phylANOVA and phyloLev functions. }*
# To fully interrogate our linear morphometric data and see how limb proportions and limb bone shapes differ among groups for a given region of tree space, we want to compare the mean and the variance of each quantitative response variable among groups for every combination of grouping_var, response_var, tree, and parced dataset in this script. To do that, we will (for each aforementioned dataset-variable-tree combination) perform one statistical test to compare means among groups (ie, a phylogenetic ANOVA) and one statistical test to compare variances among groups (ie, a phylogenetic Levene's test). In this chunk of code, we write functions to perform these tasks. In a later chunk of code [S3.3.03], we will write a "wrapper" function to at once do both of these tests and make various associated plots to visualize the results.

# To begin, we modify an existing function to perform phylogenetic ANOVAs. We will perform simulation-based phylogenetic ANOVAs following Garland et al. 1993 [https://doi.org/10.1093/sysbio/42.3.265]. To do this, we incorporate the phylANOVA() command from the phytools package (Revell 2012 [https://doi.org/10.1111/j.2041-210X.2011.00169.x]) into an original R function that outputs phylANOVA and post-hoc test results to your working directory as PNGs of easily readable tables. However, the original phylANOVA() function (from phytools v. 1.5-1) has several warnings that aren't helpful in our particular circumstances, so we modify this function below. The resulting phylAOV() function below is nearly identical to the original phylANOVA() function, with a few small modifications: first, it contains a "verbose" argument that lets us selectively reduce function output; second, its original "tree", "x", and "y" arguments have been modified to "trimmed_tree", "reordered_dataset", "grouping_variable", and "response_variable", in order to parallel the inputs of all other custom functions (and these modified inputs are defined accordingly within the function); third, its pairwise post-hoc comparisons are conducted by default with the Benjamini & Hochberg (1995) procedure, which is less prone to false negatives than methods based on the family-wise error rate (Benjamini & Hochberg 1995 [http://www.jstor.org/stable/2346101]).

phylAOV <- function (trimmed_tree, reordered_dataset, grouping_variable, response_variable, nsim = 1000, posthoc = TRUE, p.adj = "BH", verbose=F) {
  # FUNCTION ARGUMENTS:
  # trimmed_tree: an object of class "phylo" that has been trimmed with the trim.tree() function
  # reordered_dataset: a data frame, trimmed and aligned to the trimmed tree--so that it is downstream of the rm.NAs() and reorder.data() functions
  # grouping_variable: a chr string naming the column with grouping variable values (named variable should be a factor)
  # response_variable: a chr string naming the column with response variable values (named variable should be numeric)
  # nsim: a numeric value giving the number of simulations to perform (see Garland et al. 1993 [https://doi.org/10.1093/sysbio/42.3.265])
  # posthoc: chr string specifying which pairwise post-hoc tests to perform--one of "holm", "hochberg", "hommel", "bonferroni", "BH", "BY", "fdr", "none", "tukey", or "games-howell". We set "games-howell" to default because it does not assume equality of sample sizes or homogeneity of variances. Pairwise post-hoc tests are performed using the oneway() function from userfriendlyscience (v.0.7.2).
  # verbose: either TRUE or FALSE; determines whether warnings are output to the console
  
  # Define inputs within function:
  ds <- reordered_dataset
  ds$grouping_var <- ds[, grouping_variable]
  ds$grouping_var <- droplevels(ds$grouping_var)
  ds$response_var <- ds[, response_variable]
  tree <- trimmed_tree
  proceed_verdict <- "yes" # NOTE: The default value for proceed_verdict will change to "no" if feasibility requirements below are not met.
  nvals <- tapply(ds$response_var, INDEX = ds$grouping_var, FUN = function(x) length(na.omit(x)))
  
  # Assign names for variable values:
  names(ds$grouping_var) <- tree$tip.label
  names(ds$response_var) <- tree$tip.label
  
  # Report warnings if verbose=T:
  if (verbose==T) {
    warning("Warning: This function assumes that your dataset is in the order of tree$tip.label.", "\n")}
  
  # First, skip dataset-grouping_var-response_var combinations for which trimming leaves no data:
  if(dim(ds)[1]==0 | dim(ds)[2]==0){ # This checks whether there was any overlap in response_var and grouping_var rows with data. If there was not, then trimming made the dataset empty. In other words, the grouping_var has 0 levels left with data. This happens, for example, with the following combination of inputs: ds = euarchontagl_all_bLR_raw, grouping_var = bin_eco1_v7, response_var = mD_Symmetry_index1, because those specimens with mD_Symmetry_index1 measurements happened not to be scorable under bin_eco1_v7 and vice versa. The code immediately below acts in this case to report the non-feasibility of completing the phylAOV() function and saves the report to the output object:
    obj <- paste("phylAOV() has skipped the following variable pair:", "grouping_var:", grouping_variable, ", response_var:", response_variable, "|", "REASON: None of the rows with grouping_var data and response_var data overlap.")
    return(obj)
    proceed_verdict <- "no"
  } # closes 'if(dim(ds)[1]==0 | dim(ds)[2]==0)' clause
  
  # Next, check that multiple levels of the grouping_var have calculable variance:
  levels_with_calculable_variance <- 0
  for(i in 1:length(nvals)){
    if(nvals[i]>3){levels_with_calculable_variance <- levels_with_calculable_variance + 1}}
  
  if(levels_with_calculable_variance < 2){ # Report non-feasibility and save report to obj:
    warning("\n", "phylAOV() has skipped the following variable combination:", "\n", "\t", "grouping_var:", grouping_variable, "\n", "\t", "response_var:", response_variable, "\n", "REASON: The grouping variable has less than 2 levels with data.", "\n")
    obj <- paste("phylAOV() has skipped the following variable combination:", "grouping_var:", grouping_variable, ", response_var:", response_variable, "|", "REASON: The grouping variable has less than 2 levels with data.")
    return(obj)
  } # closes 'if(levels_with_calculable_variance < 2)' clause
  
  # Next, remove all levels of grouping_var withOUT calculable variance:
  ds_mod <- ds
  for(k in 1:length(nvals)){
    if(as.numeric(nvals[k])<3){
      ds_mod$response_var[ds_mod$grouping_var==levels(ds_mod$grouping_var)[k]] <- NA
    }
  } # closes 'for(k in 1:length(nvals))' loop
  ds_more_trimmed <- ds_mod[is.na(ds_mod$grouping_var)==F, ]
  ds_more_trimmed <- ds_mod[is.na(ds_mod$response_var)==F, ] 
  
  # Remove all empty levels from grouping_var:
  ds_more_trimmed$grouping_var <- droplevels(ds_more_trimmed$grouping_var)
  
  # Trim tree to dataset and order tip labels to tree:
  phylo_tr <- trim.tree(tree=tree,trimmed_dataset=ds_more_trimmed, verbose=F)
  
  # Remove NaN and 0 Myr branch lengths:
  for(i in 1:length(phylo_tr$edge.length)){
    if(is.na(phylo_tr$edge.length[i])==T){
      phylo_tr$edge.length[i] <- 0.1
    }
    if(phylo_tr$edge.length[i]==0)
      phylo_tr$edge.length[i] <- 0.1
  } # closes 'for(i in 1:length(phylo_tr$edge.length))' loop
  
  ds_ordered <- reorder.data(trimmed_tree=phylo_tr,trimmed_dataset=ds_more_trimmed, verbose=F)
  
  # OPTIONAL CONFIRMATION THAT NUMBER OF TREE TIPS AND SPECIMENS MATCH:
    #print("phylo_tr$tip.label:");print(phylo_tr$tip.label)
    #print("ds_ordered$response_var:");print(ds_ordered$response_var)
  
  # If grouping_var has at least 2 levels left, run rest of function:
  if(length(levels(ds_ordered$grouping_var)) > 1 & !(proceed_verdict=="no")){ 
    x <- ds_ordered$grouping_var
    y <- ds_ordered$response_var
    # Perform ANOVA using code from original phyloANOVA() function:
      sig2 <- mean(pic(y, multi2di(phy=phylo_tr, random = FALSE))^2)
      m <- length(levels(x))
      aov <- anova(lm(y ~ x))
      F.obs <- aov[1, 4]
      if (posthoc==TRUE) { 
      # Define tTests() function (source code available at URL=https://github.com/liamrevell/phytools/blob/master/R/phylANOVA.R):
      tTests<-function(x_ob,y_ob){
        if(!is.factor(x_ob)) x_ob<-as.factor(x_ob)
        ybar<-tapply(y_ob,x_ob,mean,na.rm=TRUE)
        s<-tapply(y_ob,x_ob,sd,na.rm=TRUE)
        n<-tapply(!is.na(y_ob),x_ob,sum)
        m<-length(levels(x_ob))
        degf<-n-1
        total.degf<-sum(degf)
        pooled.sd<-sqrt(sum(s^2*degf)/total.degf)
        compare.levels<-function(i_ob,j_ob){
          dif<-ybar[i_ob]-ybar[j_ob]
          se.dif<-pooled.sd*sqrt(1/n[i_ob]+1/n[j_ob])
          t.val<-dif/se.dif
          return(t.val)
        } # closes compare.levels() function definition
        T <- matrix(NA,m,m,dimnames=list(levels(x_ob),levels(x_ob)))
        for(i in 1:m){
          for(j in 1:m){
            T[i,j]<-compare.levels(levels(x_ob)[i],levels(x_ob)[j])
          } # closes for-loop that runs compare.levels function 
        } # closes 'for(i in 1:m)' loop
        return(T)
      } # closes tTests() function definition 
    # Run tTests() function on x and y:
      T.obs <- tTests(x,y)
      sims <- fastBM(tree=phylo_tr, sig2=sig2, nsim = (nsim - 1))
      F.null <- vector()
      T.null <- array(NA, dim = c(m, m, nsim), dimnames = list(levels(x), levels(x), NULL))
      F.null[1] <- F.obs
      T.null[, , 1] <- T.obs
      for (i in 2:nsim) {
        sims_current <- sims[, i-1]
        F.null[i] <- anova(lm(sims_current ~ x))[1, 4]
        T.null[, , i] <- tTests(x, sims[, i - 1])
      } # closes 'for (i in 2:nsim)' loop
      P.F <- sum(F.null >= F.obs)/nsim
      P.T <- matrix(NA, m, m, dimnames = list(levels(x), levels(x)))
      for (i in 1:m){
        for (j in i:m){
          P.T[i, j] <- sum(abs(T.null[i, j, ]) >= abs(T.obs[i, j]))/nsim
          P.T[j, i] <- P.T[i, j]
        }
      } # closes 'for (i in 1:m)' loop
      P.T[lower.tri(P.T)] <- p.adjust(P.T[lower.tri(P.T)], method = p.adj)
      for (i in 1:m){
        for (j in i:m){ P.T[i, j] <- P.T[j, i]
        obj <- list(ds_fin = ds_ordered, F = F.obs, Pf = P.F, T = T.obs, method = p.adj, Pt = P.T, `Sum Sq` = aov$"Sum Sq", `Mean Sq` = aov$"Mean Sq")
        } # closes 'for (j in i:m)' loop
      } # closes 'for (i in 1:m)' loop
    } else { 
      obj <- list( F = F.obs, Pf = P.F, `Sum Sq` = aov$"Sum Sq", 
                   `Mean Sq` = aov$"Mean Sq")
    } # closes big 'if (posthoc==TRUE) { } else { }' clause
    class(obj) <- "phylANOVA"
    return(obj)
  } # closes feasibility-requirements if-then clause: " if(length(levels(ds_ord$grouping_var)) > 1 & !(proceed_verdict=="no")) ".
} # closes phylAOV() function

# We have just defined a phylAOV() function, modified from the earlier phylANOVA() function, to perform a simulation-based phylogenetic ANOVA with post-hoc tests using our preferred inputs. Next, we will write a new function to perform a phylogenetic Levene's test. Levene's test compares the variances of multiple groups to test for the equality of variances. Running a Levene’s Test is equivalent to running an ANOVA on the absolute deviations of all samples from their respective group means. We were unable to find a published phylogenetic Levene's test that accounts for the phylogenetic structure of a dataset before testing for the equality of variances among groups. The following phyloLev() function does just that: first, it calculates the absolute deviation of each response_variable value from its corresponding group mean. It then performs a phylogenetic ANOVA (w/ post-hoc tests) on the resulting absolute deviations, using the phyloAOV() function we just defined above. 

phyloLev <- function(trimmed_tree, reordered_dataset, grouping_variable, response_variable, nsim = 1000, posthoc = TRUE, p.adj = "BH", verbose=F){
  # FUNCTION ARGUMENTS:
  # trimmed_tree: an object of class "phylo" that has been trimmed with the trim.tree() function
  # reordered_dataset: a data frame, trimmed and aligned to the trimmed tree--so that it is downstream of the rm.NAs() and reorder.data() functions
  # grouping_variable: a chr string naming the column with grouping variable values (named variable should be a factor)
  # response_variable: a chr string naming the column with response variable values (named variable should be numeric)
  # nsim: a numeric value giving the number of simulations to perform (see Garland et al. 1993, cited above)
  # posthoc: either TRUE or FALSE; determines whether pairwise posthoc tests will be performed
  # p.adj: chr string--one of "holm", "hochberg", "hommel", "bonferroni", "BH", "BY", "fdr", or "none"; determines whether and how to correct post-hoc test results to avoid false positive test results caused by the family-wise error rate associated with multiple testing; "BH" and "fdr" are preferable for our purposes since they are less prone to false negative test results than some of the more well-known correction methods (eg, holm and bonferroni; see phylAOV() function above).
  # verbose: either TRUE or FALSE; determines whether warnings are output to the console
  
  # Define inputs within function:
  ds <- reordered_dataset
  tree <- trimmed_tree
  x <- ds$grouping_var <- ds[, grouping_variable]
  y <- ds$response_var <- ds[, response_variable]
  nsim_num <- nsim
  p_adjustment_method <- p.adj
  
  if(verbose==T){verbose_verdict <- T
  } else if(verbose==F){verbose_verdict <- F}
  
  # Assign names for variable values:
  names(ds$grouping_var) <- tree$tip.label
  names(ds$response_var) <- tree$tip.label
  
  # Replace each sample’s original value with its absolute deviation from its group mean:
  # Initialize output vector for absolute deviations:
  absolute_deviations <- rep(NA, nrow(ds))
  for(i in 1:nrow(ds)){
    # Get group mean for current specimen:
    current_group <- ds$grouping_var[i]
    mean_of_current_group <- mean(ds$response_var[ds$grouping_var==current_group])
    # Calculate absolute deviation for each sample:
    absolute_deviations[i] <- abs(mean_of_current_group - ds$response_var[i])
  } # closes 'for(i in 1:nrow(ds))' loop
  
  # Add absolute deviations to dataset:
  ds$absolute_deviations <- absolute_deviations
  names(ds$absolute_deviations) <- tree$tip.label
  
  # Run phylogenetic ANOVA on absolute deviations using phylAOV() function:
  phyloLev_obj <- phylAOV(trimmed_tree=tree, reordered_dataset=ds, response_variable="absolute_deviations", grouping_variable=grouping_variable, nsim=nsim_num, p.adj=p_adjustment_method, verbose=verbose_verdict)
  
  # Save output object to R environment:
  class(phyloLev_obj) <- "phyloLev"
  return(phyloLev_obj)
} # closes phyloLev() function
cat("\n","PROGRESS REP: chunk [S2.2.01] complete; starting next chunk...","\n")
#*-----**-----*

#*-----**{ [S2.2.02] Check that phylAOV and phyloLev functions work. }*
# In the following chunk, we will test our new functions using a series of challenging inputs, to make sure they work correctly. This is a useful troubleshooting step for identifying particular inputs that produce errors.

# Run upstream functions:
  test_ds <- all_data
  ds_tr <- rm.NAs(dataset=test_ds, categorical_var = "Ecotype1", quantitative_var = "f_AZR")
  phylo_tr <-trim.tree(tree=tree_variants[[1]],trimmed_dataset=ds_tr,verbose=F)
  ds_ord <-reorder.data(trimmed_dataset=ds_tr, trimmed_tree=phylo_tr,verbose=F)

# Test the phylAOV() function using some 'easier' input combinations:
  test_phylAOV_output <- phylAOV(trimmed_tree=phylo_tr, reordered_dataset=ds_ord, verbose = T, grouping_variable="Ecotype1", response_variable="f_AZR", nsim=1000, posthoc=T, p.adj="BH")
# Test the phyloLev() function:
  test_phyloLev_output <- phyloLev(trimmed_tree=phylo_tr, reordered_dataset=ds_ord, verbose = T, grouping_variable="Ecotype1", response_variable = "f_AZR", nsim = 1000, posthoc = T, p.adj = "BH")

# Next, we will test them using trickier input combinations that previously generated errors during trouble-shooting. 
  
# CHALLENGING INPUT COMBINATION #1:
  # Run upstream functions:
    test_ds <- euarchontagl_all
    ds_tr <- rm.NAs(dataset=test_ds, categorical_var = "fore_type", quantitative_var = "f_AZR")
    phylo_tr <- trim.tree(tree=tree_variants[[1]], trimmed_dataset=ds_tr, verbose=F)
    ds_ord <- reorder.data(trimmed_dataset=ds_tr, trimmed_tree=phylo_tr, verbose = F)
  # Test the phylAOV() and phyloLev() functions:
    test_phylAOV_output <- phylAOV(trimmed_tree=phylo_tr, reordered_dataset=ds_ord, grouping_variable="Ecotype1", response_variable="f_AZR", nsim=1000, posthoc=T, p.adj="BH") 
    test_phyloLev_output <- phyloLev(trimmed_tree=phylo_tr, reordered_dataset=ds_ord, grouping_variable="Ecotype1", response_variable="f_AZR", nsim=1000, posthoc=T, p.adj="BH")
  # Check that post-hoc tests worked:
    test_phylAOV_output$Pt # This matrix should contain actual numeric p-vals, rather than NAs, despite the fact that some grouping_var levels had to be removed because they had less than two rows with data, and subsequently had non-calculable variances).
    test_phyloLev_output$Pt # This matrix should also contain actual numeric p-values, rather than NAs, despite the fact that some grouping_var levels had to be removed because they had less than two rows with data, and subsequently had non-calculable variances).

# CHALLENGING INPUT COMBINATION #2: 
  # Run upstream functions:
    test_ds <- log10_datasets[[7]]
    ds_tr <- rm.NAs(dataset=test_ds, categorical_var = "Subclade", quantitative_var = "mD3_up_FI1")
    phylo_tr <- trim.tree(tree=tree_variants[[1]], trimmed_dataset=ds_tr, verbose=F)
    ds_ord <- reorder.data(trimmed_dataset=ds_tr, trimmed_tree=phylo_tr, verbose = F)
  # Test the phylAOV() and phyloLev() functions:
    test_phylAOV_output <- phylAOV(trimmed_tree=phylo_tr, reordered_dataset=ds_ord, grouping_variable="Subclade", response_variable="mD3_up_FI1", nsim=1000, posthoc=T, p.adj="BH") 
    test_phyloLev_output <- phyloLev(trimmed_tree=phylo_tr, reordered_dataset=ds_ord, grouping_variable="Subclade", response_variable="mD3_up_FI1", nsim=1000, posthoc=T, p.adj="BH") 
  # Check that the outputs are correct:
    test_phylAOV_output # This object should here just be a character string explaining the non-feasibility of running the function on these inputs.
    test_phyloLev_output # This object should here just be a character string explaining the non-feasibility of running the function on these inputs.

# CHALLENGING INPUT COMBINATION #3: 
  # Run upstream functions:
    test_ds <- log10_datasets[[1]]
    ds_tr <- rm.NAs(dataset=test_ds, categorical_var = "Subclade", quantitative_var = "mD3_up_FI1")
    phylo_tr <- trim.tree(tree=tree_variants[[1]], trimmed_dataset=ds_tr, verbose=F)
    ds_ord <- reorder.data(trimmed_dataset=ds_tr, trimmed_tree=phylo_tr, verbose = F)
  # Test the phylAOV() and phyloLev() functions:
    test_phylAOV_output <- phylAOV(trimmed_tree=phylo_tr, reordered_dataset=ds_ord, grouping_variable="Subclade", response_variable="mD3_up_FI1", nsim=1000, posthoc=T, p.adj="BH") 
    test_phyloLev_output <- phyloLev(trimmed_tree=phylo_tr, reordered_dataset=ds_ord, grouping_variable="Subclade", response_variable="mD3_up_FI1", nsim=1000, posthoc=T, p.adj="BH") 
  # Check that post-hoc tests worked:
    test_phylAOV_output$Pt # This matrix should contain actual numeric p-vals, rather than NAs, despite the fact that some grouping_var levels had to be removed because they had less than two rows with data, and subsequently had non-calculable variances).
    test_phyloLev_output$Pt # This matrix should also contain actual numeric p-values, rather than NAs, despite the fact that some grouping_var levels had to be removed because they had less than two rows with data, and subsequently had non-calculable variances).

# CHALLENGING INPUT COMBINATION #4: 
  # Run upstream functions:
    test_ds <- all_data
    ds_tr <- rm.NAs(dataset=test_ds, categorical_var = "f_subtype", quantitative_var = "mD_Symmetry_index1")
    phylo_tr <- trim.tree(tree=tree_variants[[1]], trimmed_dataset=ds_tr, verbose=F)
    ds_ord <- reorder.data(trimmed_dataset=ds_tr, trimmed_tree=phylo_tr, verbose = F)
  # Test the phylAOV() and phyloLev() functions:
    test_phylAOV_output <- phylAOV(trimmed_tree=phylo_tr, reordered_dataset=ds_ord, grouping_variable="f_subtype", response_variable="mD_Symmetry_index1", nsim=1000, posthoc=T, p.adj="BH") 
    test_phyloLev_output <- phyloLev(trimmed_tree=phylo_tr, reordered_dataset=ds_ord, grouping_variable="f_subtype", response_variable="mD_Symmetry_index1", nsim=1000, posthoc=T, p.adj="BH") 
  # Check that post-hoc tests worked:
    test_phylAOV_output$Pt # This matrix should contain actual numeric p-values, rather than NAs, despite the fact that some grouping_var levels had to be removed because they had less than two rows with data, and subsequently had non-calculable variances).
    test_phyloLev_output$Pt # This matrix should also contain actual numeric p-values, rather than NAs, despite the fact that some grouping_var levels had to be removed because they had less than two rows with data, and subsequently had non-calculable variances).

# CHALLENGING INPUT COMBINATION #5: 
  # Run upstream functions:
    test_ds <- ferae_all
    ds_tr <- rm.NAs(dataset=test_ds, categorical_var = "Ecotype1", quantitative_var = "mD3_up_FI1")
    phylo_tr <- trim.tree(tree=tree_variants[[1]], trimmed_dataset=ds_tr, verbose=F)
    ds_ord <- reorder.data(trimmed_dataset=ds_tr, trimmed_tree=phylo_tr, verbose = F)
  # Test the phylAOV() and phyloLev() functions:
    test_phylAOV_output <- phylAOV(trimmed_tree=phylo_tr, reordered_dataset=ds_ord, grouping_variable="Ecotype1", response_variable="mD3_up_FI1", nsim=1000, posthoc=T, p.adj="BH") # Output should report infeasibility of test.
    test_phyloLev_output <- phyloLev(trimmed_tree=phylo_tr, reordered_dataset=ds_ord, grouping_variable="Ecotype1", response_variable="mD3_up_FI1", nsim=1000, posthoc=T, p.adj="BH") # Output should report infeasibility of test.
    
# Now that we have tested the phylAOV() and phyloLev() functions using a combination of difficult inputs, we can proceed to apply them across all possible input combinations. We will automate this process by calling both functions internally within a larger "wrapper" function, which we define next.
cat("\n","PROGRESS REP: chunk [S2.2.02] complete; starting next chunk...","\n")
#*-----**-----*

#*-----**{ [S2.2.03] Define all-in-one across-group-tests wrapper function. }*
# The phyloBoxPlot() function below is a wrapper function that internally calls the previously defined phylAOV() and phyloLev() functions, along with ggplot2 functions, to generate one box plot and eight sets of statistical test outputs for every dataset-variable triad. The phyloBoxPlot() function will output the following items: (1) one custom boxplot for each grouping_var/response_var combination; (2) one high-resolution PNG for each tree/grouping_var/response_var combination, which contains both (i) a data table of phylogenetic ANOVA statistical test results for each tree/grouping_var/response_var combination, and (ii) one data table of phylogenetic Levene's test results for each tree/grouping_var/response_var combination.

phyloBoxPlot <- function(ds, dataset_name, tree_list, response_var, grouping_var, nreps, is.binary){
  # FUNCTION INPUTS:
  # ds: a data frame that has not been processed with rm.NAs(), trim.tree(), or reorder.data(), as these will all be called internally to process the dataset within the function.
  # dataset_name: a chr string naming your dataset (required to name output files)
  # tree_list: a list of tree variants (each of class "phylo") with which to run phylogenetic comparative tests
  # response_var: a chr string naming the numeric response variable within the dataset (should be name of a column within ds)
  # grouping_var: a chr string naming the grouping variable within the dataset (should be name of a column within ds) 
  # nreps: numeric value giving the number of iterations to be run for phylAOV() and phyloLev()
  # is.binary: one of TRUE or FALSE; reports whether dataset has been treated previously with bLRify()
  
  # Define directory path for output files: 
  directory_path <- paste0(output_path, "/phyloBoxPlots")
  # Create directory for output files:
  dir.create(path=directory_path, showWarnings = TRUE, recursive = FALSE, mode = "0777")
  # Define string for dataset-variable combination (for naming output files):
  datasetVariableCombo <- paste0(dataset_name, "__", response_var, "_by_", grouping_var)
  
  # Relevel factors of dataset within function ()
  if(is.binary==F){
    dataset_prepped <- relevel_factors(dataset=ds) 
  } else if (is.binary==T){
    dataset_prepped <- ds
  }
  
  # Trim dataset to remove NAs:
  ds_tr <- rm.NAs(dataset=dataset_prepped, categorical_var=grouping_var, quantitative_var=response_var)
  
  # Define response_var and grouping_var within the function:
  ds_tr$response_var <- ds_tr[ , response_var]
  ds_tr$grouping_var <- ds_tr[ , grouping_var]
  ds_tr$grouping_var <- droplevels(ds_tr$grouping_var)
  
  proceed_verdict <- "yes" # This is the default start state for a dummy variable used to determine whether or not the function can proceed with phylogenetic comparative tests.
  
  # Skip dataset-grouping_var-response_var combinations for which trimming leaves no data:
  if(dim(ds_tr)[1]==0 | dim(ds_tr)[2]==0){ # This checks whether there was any overlap in response_var and grouping_var rows with data. If there was not, then trimming made the dataset empty. In other words, the grouping_var has 0 levels left with data. This happens, for example, with the following combination of inputs: ds = euarchontagl_all_bLR_raw, grouping_var = bin_eco1_v7, response_var = mD_Symmetry_index1, because those specimens with mD_Symmetry_index1 measurements happened not to be scorable under bin_eco1_v7 and vice versa. The code immediately below acts in this case to report the non-feasibility of completing the phyloBoxPlot() function and saves the report to the phyloBoxPlot_output_list:
    warning("\n", "phyloBoxPlot() has skipped the following dataset-variable triad:", "\n", "\t", "dataset:", dataset_name, "\n", "\t", "grouping_var:", grouping_var, "\n", "\t", "response_var:", response_var, "\n", "REASON: None of the rows with grouping_var data and response_var data overlap;", "\n", "thus, trimming with rm.NAs() appropriately left the dataset empty.", "\n")
    phyloBoxPlot_output_list <- paste("phyloBoxPlot() has skipped the following dataset-variable triad:", "dataset:", dataset_name, ",", "grouping_var:", grouping_var, ", response_var:", response_var, "|", "REASON: REASON: None of the rows with grouping_var data and response_var data overlap.")
    return(phyloBoxPlot_output_list)
    proceed_verdict <- "no"
  } # closes 'if(dim(ds_tr)[1]==0 | dim(ds_tr)[2]==0)' clause
  
  # Skip mismatched forelimb-hindlimb variable combinations:
  if((str_detect(string=grouping_var, pattern="fore|f_")==T) & (str_detect(string=response_var, pattern="h_|pD")==T)){ # Report non-feasibility and save report to phyloBoxPlot_output_list.
    warning("\n", "phyloBoxPlot() has skipped the following dataset-variable triad:", "\n", "\t", "dataset:", dataset_name, "\n", "\t", "grouping_var:", grouping_var, "\n", "\t", "response_var:", response_var, "\n", "REASON: One variable describes forelimbs and the other hindlimbs.", "\n")
    phyloBoxPlot_output_list <- paste("phyloBoxPlot() has skipped the following dataset-variable triad:", "dataset:", dataset_name, ",", "grouping_var:", grouping_var, ", response_var:", response_var, "|", "REASON: One variable describes forelimbs and the other hindlimbs.")
    return(phyloBoxPlot_output_list)
    proceed_verdict <- "no"
  } # closes first 'if ( mismatched forelimb-hindlimb variables )' clause
  
  if((str_detect(string=grouping_var, pattern="hind|h_")==T) & (str_detect(string=response_var, pattern="f_|mD")==T)){ # Report non-feasibility and save report to phyloBoxPlot_output_list.
    warning("\n", "phyloBoxPlot() has skipped the following dataset-variable triad:", "\n", "\t", "dataset:", dataset_name, "\n", "\t", "grouping_var:", grouping_var, "\n", "\t", "response_var:", response_var, "\n", "REASON: One variable describes forelimbs and the other hindlimbs.)", "\n")
    phyloBoxPlot_output_list <- paste("phyloBoxPlot() has skipped the following dataset-variable triad:", "dataset:", dataset_name, ",", "grouping_var:", grouping_var, ", response_var:", response_var, "|", "REASON: One variable describes forelimbs and the other hindlimbs.")
    return(phyloBoxPlot_output_list)
    proceed_verdict <- "no"
  } # closes second 'if ( mismatched forelimb-hindlimb variables )' clause
  
  # If proceed_verdict remains "yes", the function will proceed.
  if(proceed_verdict == "yes"){
    # Define tick labels within function:
    group_names <- levels(droplevels(na.omit(ds_tr$grouping_var)))
    nvals <- tapply(ds_tr$response_var, INDEX = droplevels(ds_tr$grouping_var), FUN = function(x) length(na.omit(x)))
    tick_labels <- paste(group_names, "\n n=", nvals, sep="")
    
    # Create output object to record nvals per group:
    output_table_nvals <- paste(group_names, " n=", nvals, ";", sep="")
    output_groups_1string <- output_table_nvals[1]
    for(w in 2:length(output_table_nvals)){
      output_groups_1string <- paste(output_groups_1string, output_table_nvals[w], sep=" ")
    } # closes 'for(w in 2:length(output_table_nvals))' loop
    
    # Next, check that multiple levels of the grouping_var have data:
    levels_with_data <- 0
    for(i in 1:length(nvals)){
      if(nvals[i]>0){levels_with_data <- levels_with_data + 1}}
    if(levels_with_data < 2){ # Report non-feasibility and save report to phyloBoxPlot_output.
      warning("\n", "phyloBoxPlot() has skipped the following dataset-variable triad:", "\n", "\t", "dataset:", dataset_name, "\n", "\t", "grouping_var:", grouping_var, "\n", "\t", "response_var:", response_var, "\n", "REASON: The grouping variable has less than 2 levels with data.", "\n")
      phyloBoxPlot_output_list <- paste("phyloBoxPlot() has skipped the following dataset-variable triad:", "dataset:", dataset_name, ",", "grouping_var:", grouping_var, ", response_var:", response_var, "|", "REASON: The grouping variable has less than 2 levels with data.")
      return(phyloBoxPlot_output_list)
    } # closes 'if(levels_with_data < 2)' clause
    else { #proceeds with rest of function
      # Define palette and box plot aesthetic parameters:
        group_colors <- rep(NA, length(levels(droplevels(na.omit(ds_tr$grouping_var)))))
        for(q in 1:length(group_colors)){
          if(nvals[q]>0){
          group_colors[q] <- grouping_var_palette[[grouping_var]][levels(droplevels(na.omit(ds_tr$grouping_var)))[q]]
          }
        } # closes 'for(q in 1:length(group_colors))' loop
      if(is.binary==TRUE){
        group_colors <- grouping_var_palette[[grouping_var]]
      } # closes 'if(is.binary==TRUE)' clause
      
      subtype_aspect <- 1/3
      subtype_boxsize <- 0.9
      subtype_meanpoint_size <- 1.5
      subtype_meanstroke_size <- 1
      type_aspect <- 3/1
      type_boxsize <- 1.5
      type_meanpoint_size <- 3
      type_meanstroke_size <- 1.5
      
      if((grouping_var=="f_subtype")|(grouping_var=="h_subtype")){
        plot_aspect <- subtype_aspect
        plot_boxsize <- subtype_boxsize
        meanpoint_size <- subtype_meanpoint_size
        meanstroke_size <- subtype_meanstroke_size
      } else {
        plot_aspect <- type_aspect
        plot_boxsize <- type_boxsize
        meanpoint_size <- type_meanpoint_size
        meanstroke_size <- type_meanstroke_size
      } # closes if-else clause about box plot aesthetic parameters
      
    # Generate box plot:
      gg_bp <- ggplot(ds_tr,aes(x=grouping_var, y=response_var)) +
        geom_boxplot( fill=group_colors,
                      position=position_dodge2(preserve="single"),
                      linewidth=plot_boxsize,
                      outlier.size=1,
                      show.legend=TRUE) +
        theme(plot.margin=margin(t=20, r=150, b=40, l=150),
              axis.text=element_text(size=8, angle=90, hjust=1),
              axis.title=element_text(size=12), 
              aspect.ratio=plot_aspect) + 
        xlab(label=NULL) + ylab(label=response_var) +
        stat_summary(fun=mean, geom="point", shape=23, size=meanpoint_size, stroke=meanstroke_size, color="black",fill='white') +
        scale_x_discrete(labels = tick_labels)
      
      # Save high-resolution PNG of box plot:
      png(filename=paste(directory_path, "/", datasetVariableCombo, "__boxPlot", ".png", sep=""), width=10, height=10, units="in", res=200)
      show(gg_bp)
      dev.off()
      
      # Initialize output list of lists (one list item for each tree):
      phyloBoxPlot_output_list <- list(rep(NA, length(tree_list)))
      phyloBoxPlot_output_names <- paste("phyloBoxPlot output:", datasetVariableCombo, "- for", names(tree_list))
      
      # Running per-tree phylAOV() and phyloLev() functions:
      for(j in 1:length(tree_list)){
        tree_name <- names(tree_list)[j]
        # Remove all levels without calculable variance: (where n<2)
        ds_mod <- ds_tr
        for(k in 1:length(nvals)){
          if(nvals[k] < 2){
            ds_mod <- ds_mod[!(ds_mod$grouping_var == levels(ds_mod$grouping_var)[k]), ]
            ds_mod$grouping_var <- droplevels(ds_mod$grouping_var, exclude = levels(ds_tr$grouping_var)[k])
          } # closes 'if(nvals[k] < 2)' clause
        } # closes 'for(k in 1:length(nvals))' loop
        ds_more_trimmed <- ds_mod[is.na(ds_mod$grouping_var)==F, ]
        ds_more_trimmed <- ds_mod[is.na(ds_mod$response_var)==F, ]
        
        # Trim tree to dataset and order tip labels to tree:
        phylo_tr <- trim.tree(tree=tree_list[[j]],trimmed_dataset=ds_more_trimmed, verbose=F)
        
        # Check that the fully trimmed tree contains data; only proceed if yes:
        if(is.null(phylo_tr)==T){ # report non-feasibility and save report to phyloBoxPlot_output object
          cat("\n", "phyloBoxPlot() has skipped the following dataset-variable triad:", "\n", "\t", "dataset:", dataset_name, "\n", "\t", "grouping_var:", grouping_var, "\n", "\t", "response_var:", response_var, "\n", "REASON: The grouping variable has less than 2 levels with data.", "\n")
          phyloBoxPlot_output_list <- paste("phyloBoxPlot() has skipped the following dataset-variable triad:", "dataset:", dataset_name, ",", "grouping_var:", grouping_var, ", response_var:", response_var, "|", "REASON: The grouping variable has less than 2 levels with data.")
          return(phyloBoxPlot_output_list)
        } else { #proceeds with rest of function
          
          # Remove NaN and 0 Myr branch lengths:
          for(i in 1:length(phylo_tr$edge.length)){
            if(is.na(phylo_tr$edge.length[i])==T){
              phylo_tr$edge.length[i] <- 0.1
            }
            if(phylo_tr$edge.length[i]==0)
              phylo_tr$edge.length[i] <- 0.1
          } # closes 'for(i in 1:length(phylo_tr$edge.length))' loop
          ds_ord <- reorder.data(trimmed_tree=phylo_tr,trimmed_dataset=ds_more_trimmed, verbose=F)
          
          # If grouping_var has at least 2 levels left, run phylogenetic ANOVA & Levene's test:
          if(length(levels(ds_ord$grouping_var)) > 1){ 
            # Perform phylogenetic ANOVA & post-hoc tests with phylAOV():
              phylAOV_res <- phylAOV(trimmed_tree=phylo_tr, reordered_dataset=ds_ord, grouping_variable=grouping_var, response_variable=response_var, nsim=nreps, posthoc=T, p.adj="BH")
              if(is.list(phylAOV_res)==T){
              phylANOVA_pval <- paste("overall phylANOVA p-value: p = ", phylAOV_res$Pf, sep="")
                # Make table of p-vals for post-hoc tests:
                  aov_posthocs <- tableGrob(as.table(phylAOV_res$Pt),theme=ttheme_default(base_size=8))
              
                # Define a plot rectangle to output phylANOVA p-value:
                  phylanova_df<-data.frame("x1"=100,"x2"=200,"y1"=0,"y2"=40)
                  x1 <- 100; x2 <- 200; y1 <- 0; y2 <- 40
                  phylanova_rect <- (ggplot(phylanova_df) + geom_rect(mapping=aes(xmin=x1, xmax=x2, ymin=y1, ymax=y2), color="grey", linewidth=1, fill="white") + geom_label(aes(x=150, y=(y2*0.6), label = phylANOVA_pval), size=4, color='black', label.size=1, fontface='bold') + theme_classic() + theme(axis.line  = element_blank(), axis.ticks = element_blank(), axis.text  = element_blank(), axis.title = element_blank())) 
              
                # Save ggplot object of phylAOV stats:
                  phylAOV_grob <- grid.arrange(phylanova_rect, aov_posthocs, nrow=2, heights=c(1,10))
              
                # Perform phylogenetic Levene's test:
                  phyloLev_res <- phyloLev(trimmed_tree=phylo_tr, reordered_dataset=ds_ord, grouping_variable=grouping_var, response_variable=response_var, nsim=nreps, posthoc=T, p.adj="BH")
              
                # Save phylogenetic Levene's test overall p-value:
                  phyloLev_pval <- paste("Overall phylogenetic Levene's test p-val: p = ", phyloLev_res$Pf, sep="")
              
                # Make table of p-vals for post-hoc tests for ggplot output:
                  Lev_posthocs <- tableGrob(as.table(phyloLev_res$Pt),theme=ttheme_default(base_size=8))
              
                # Define a plot rectangle to output phyloLev overall p-value:
                  phyloLev_df <- data.frame("x1"=100,"x2"=200,"y1"=0,"y2"=40)
                  phyloLev_rect <- (ggplot(phyloLev_df) + geom_rect(mapping=aes(xmin=x1, xmax=x2, ymin=y1, ymax=y2), color="grey", linewidth=1, fill="white") + geom_label(aes(x=150, y=(y2*0.6), label = phyloLev_pval), size=4, color='black', label.size=1, fontface='bold') + theme_classic() + theme(axis.line  = element_blank(), axis.ticks = element_blank(), axis.text  = element_blank(), axis.title = element_blank())) 
              
                # Save ggplot object of phyloLev stats:   
                  phyloLev_grob <- grid.arrange(phyloLev_rect, Lev_posthocs, nrow=2, heights=c(1,10))
              
                # Save high-resolution PNG file of phylAOV & phyloLev stats:
                  png(filename=paste(directory_path, "/", datasetVariableCombo, "_for_", tree_name, "__AOV_and_Lev", ".png", sep=""), width=15, height=10, units="in", res=100)
                  grid.arrange(phylAOV_grob, phyloLev_grob)
                  dev.off()
              
            # Save function output to an R object of class "phyloBoxPlot":
              # Initialize output list for present tree:
              pres_phyloBoxPlot_output <- list()
              # Populate pres_phyloBoxPlot_output w/ metadata for this call:
              pres_phyloBoxPlot_output$function_name <- "phyloBoxPlot()"
              pres_phyloBoxPlot_output$bootstrap_number <- nreps
              pres_phyloBoxPlot_output$input_dataset <- dataset_name
              pres_phyloBoxPlot_output$input_tree <- tree_name
              pres_phyloBoxPlot_output$response_variable <- response_var
              pres_phyloBoxPlot_output$grouping_variable <- grouping_var
              pres_phyloBoxPlot_output$full_sample_size <- nrow(ds_ord)
              pres_phyloBoxPlot_output$sample_size_by_group<-output_table_nvals
              pres_phyloBoxPlot_output$sample_size_by_group_singlestring <- output_groups_1string
              pres_phyloBoxPlot_output$phylANOVA_omnibus_Fstat<-phylAOV_res$F
              pres_phyloBoxPlot_output$phylANOVA_omnibus_pval<-phylAOV_res$Pf
              pres_phyloBoxPlot_output$phylANOVA_posthoc_Tstats<-phylAOV_res$T
              pres_phyloBoxPlot_output$phylANOVA_posthoc_correction_method<-"Benjamini-Hochberg (BH)"
              pres_phyloBoxPlot_output$phylANOVA_posthoc_pvals<-phylAOV_res$Pt
              pres_phyloBoxPlot_output$phyloLev_omnibus_Fstat <- phyloLev_res$F
              pres_phyloBoxPlot_output$phyloLev_omnibus_pval <- phyloLev_res$Pf
              pres_phyloBoxPlot_output$phyloLev_posthoc_Tstats<-phyloLev_res$T
              pres_phyloBoxPlot_output$phyloLev_posthoc_correction_method<-"Benjamini-Hochberg (BH)"
              pres_phyloBoxPlot_output$phyloLev_posthoc_pvals<-phyloLev_res$Pt
              pres_phyloBoxPlot_output$phylAOV_full_output <- phylAOV_res
              pres_phyloBoxPlot_output$phyloLev_full_output <- phyloLev_res
              phyloBoxPlot_output_list[[j]] <- pres_phyloBoxPlot_output
            } else if (is.list(phylAOV_res)==F) {
              # Save non-feasibility report to phyloBoxPlot_output object: 
              phyloBoxPlot_output_list <- paste("phyloBoxPlot() has skipped the following dataset-variable triad:", "dataset:", dataset_name, ",", "grouping_var:", grouping_var, ", response_var:", response_var, "|", "REASON: The grouping_var for this dataset has < 2 levels.", sep=" ")
              return(phyloBoxPlot_output_list)
            } # Closes else-if clause for phylAOV not being a list.
            
            # Report case where phylogenetic mean-and-var tests not possible:
          } else if (length(levels((ds_ord$grouping_var))) <= 1){
            warning("\n", dataset_name, "|", grouping_var, ": phylANOVA not possible", "\n", "REASON: The grouping_var for this dataset has < 2 levels.", "\n")
            # Save non-feasibility report to phyloBoxPlot_output object: 
            phyloBoxPlot_output_list <- paste("phyloBoxPlot() has skipped the following dataset-variable triad:", "dataset:", dataset_name, ",", "grouping_var:", grouping_var, ", response_var:", response_var, "|", "REASON: The grouping_var for this dataset has < 2 levels.", sep=" ")
            return(phyloBoxPlot_output_list)
          } # closes else-if clause for when grouping_var has < 2 levels
        } # closes else-if clause for when tree trimming eliminates all tips
      } # closes for-loop iteration through all trees in tree_list
      
      # Report completion of function for a given set of inputs:
      cat("\n", "DATASET-VARIABLE COMBO:", dataset_name, "|", response_var, "by", grouping_var, ":", "\n", "COMPLETE: box plot, phylANOVA & phyloLev test w/ pairwise post-hocs", "\n", "LOCATION:", "Output files saved as PNGs in the working directory.", "\n")
      
      # Name phyloBoxPlot_output_list items:
      names(phyloBoxPlot_output_list) <- phyloBoxPlot_output_names
      # Save phyloBoxPlot_output_list to environment:
      return(phyloBoxPlot_output_list)
    } # closes if-else "levels_with_data < 2" clause
  } # closes "if(proceed_verdict==T)" clause
} # closes phyloBoxPlot() function

# We have just defined a wrapper function that runs phylAOV(), phyloLev(), and ggplot2 commands on a specified set of inputs. In the next chunk of code, we test that function to make sure that it works before proceeding to run it on all possible input combinations. 
cat("\n","PROGRESS REP: chunk [S2.2.03] complete; starting next chunk...","\n")
#*-----**-----*

#*-----**{ [S2.2.04] Test phyloBoxPlot() with difficult input combinations. }*
# In this chunk of code, we test the phyloBoxPlot() function on several different dataset-variable combinations to make sure it works in every case in every way it's supposed to work. Please note: The p-values for phylogenetic ANOVA and phylogenetic Levene's omnibus and posthoc tests are highly variable when nreps<1000, so the "*test_all" results immediately below should not be considered for interpreting the dataset as a whole. They will include many false negatives at low nrep values. As a result, the test outputs below are intended solely for troubleshooting to confirm that the phyloBoxPlot() function and the custom functions called within it are working properly.

test_name <- "*test_all"

test_phyloBoxPlot_output <- phyloBoxPlot(ds=all_data, dataset_name=test_name, tree_list=tree_variants, response_var="f_AZR", grouping_var="Ecotype1",nreps=5, is.binary=F) # should work and save plots to newly created "phyloBoxPlots" folder. 

test_phyloBoxPlot_output <- phyloBoxPlot(ds=euarchontagl_all, dataset_name="*test_euarchontagl", tree_list=tree_variants, response_var="f_AZR", grouping_var="fore_type",nreps=5, is.binary=F) # should work and save plots to newly created "phyloBoxPlots" folder.

test_phyloBoxPlot_output <- phyloBoxPlot(ds=all_data, dataset_name="*test_all", tree_list=tree_variants, response_var="f_ASR", grouping_var="Flipper_status", nreps=5, is.binary=F) # should work and save plots to newly created "phyloBoxPlots" folder.

test_phyloBoxPlot_output <- phyloBoxPlot(ds=reps_all, dataset_name="*test_reps", tree_list=tree_variants, response_var="f_SZR", grouping_var="Flipper_status", nreps=5, is.binary=F) # should work and save plots to newly created "phyloBoxPlots" folder.

test_phyloBoxPlot_output <- phyloBoxPlot(ds=all_data, dataset_name="*test_all", tree_list=tree_variants, response_var="f_AZR", grouping_var="fore_type", nreps=5, is.binary=F) # should work and save plots to newly created "phyloBoxPlots" folder.

test_phyloBoxPlot_output <- phyloBoxPlot(ds=all_trans, dataset_name="*test_trans", tree_list=tree_variants, response_var="f_AZR", grouping_var="Ecotype1", nreps=5, is.binary=F) # should work and save plots to newly created "phyloBoxPlots" folder.

test_phyloBoxPlot_output <- phyloBoxPlot(ds=all_data, dataset_name="*test_all", tree_list=tree_variants, response_var="f_AZR", grouping_var="Subclade", nreps=5, is.binary=F) # should work despite MANY subclade divisions.

test_phyloBoxPlot_output <- phyloBoxPlot(ds=ungulates_all, dataset_name="*test_ungulates_all", tree_list=tree_variants, response_var="f_AZR", grouping_var="Ecotype1", nreps=5, is.binary=F) # should have the correct color for each ecotype bin even though there are no "Highly Aquatic" ungulates in the data set.

test_phyloBoxPlot_output <- phyloBoxPlot(ds=all_data, dataset_name="*test_all", tree_list=tree_variants, response_var="pD3_up_FI2", grouping_var="fore_type", nreps=5, is.binary=F) # should say it skipped because of forelimb-hindlimb variable mismatch.

test_phyloBoxPlot_output <- phyloBoxPlot(ds=all_data, dataset_name="*test_all", tree_list=tree_variants, response_var="mD3_up_FI2", grouping_var="hind_type", nreps=5, is.binary=F) # should say it skipped because of forelimb-hindlimb variable mismatch.

test_phyloBoxPlot_output <- phyloBoxPlot(ds=ungulates_all, dataset_name="*test_ungulates_all", tree_list=tree_variants, response_var="f_AZR", grouping_var="MajorClade", nreps=5, is.binary=F) # should say it skipped because only one level of the grouping variable contained data.

test_phyloBoxPlot_output <- phyloBoxPlot(ds=all_data, dataset_name="all", tree_list=tree_variants, response_var="mD_Symmetry_index1", grouping_var="f_subtype", nreps=5, is.binary=F) # should work without a problem (this dataset-variable triad was particularly challenging for the phyloLev portion of the function while I was troubleshooting it.

test_phyloBoxPlot_output <- phyloBoxPlot(ds=archos_all, dataset_name="*test_archos_all", tree_list=tree_variants, response_var="h_AZR", grouping_var="Ecotype1", nreps=5, is.binary=F) # should work and save plots to newly created "phyloBoxPlots" folder, despite the fact that some levels of the grouping variable are missing data. The boxplot should contain all levels of the grouping variable, whereas the phylANOVA data should only contain that subset of levels containing n>2 samples.

test_phyloBoxPlot_output <- phyloBoxPlot(ds=bLR_raw_datasets[[1]], dataset_name="*test_bLR_ds", tree_list=tree_variants, response_var="f_AZR", grouping_var="Ecotype1", nreps=5, is.binary=T) # should work even though it's a binary dataset.

test_phyloBoxPlot_output <- phyloBoxPlot(ds=log10_datasets[[1]], dataset_name="*test_all_log10", tree_list=tree_variants, response_var="mD3_up_FI1", grouping_var="Subclade", nreps=5, is.binary=F) # should work even though this dataset-variable combination produced frequent errors in previous runs.

test_phyloBoxPlot_output <- phyloBoxPlot(ds=log10_datasets[[1]], dataset_name="*test_dinos_log10", tree_list=tree_variants, response_var="f_AZR", grouping_var="fore_type", nreps=5, is.binary=F) # should work even though this dataset-variable combination produced frequent errors in previous runs.

test_phyloBoxPlot_output <- phyloBoxPlot(ds=ferae_all, dataset_name="ferae_all", tree_list=tree_variants, response_var="mD3_up_FI1", grouping_var="Ecotype1", nreps=5, is.binary=F) # should work even though this dataset-variable combination produced frequent errors in previous runs.

# We have just written and tested an all-in-one function to generate nice box plots and perform phylogenetic ANOVAs and phylogenetic Levene's tests with pairwise post-hoc comparisons for a given set of trees. Next, we will use foreach-loops to run this function on all tree-dataset-variable combinations. 
cat("\n","PROGRESS REP: chunk [S2.2.04] complete; starting next chunk...","\n")
#*-----**-----*

#*-----**{ [S2.2.05] Initialize objects for phyloBoxPlot foreach-loops.}*
# In the next few chunks of code, we will run the phyloBoxPlot() function on every  combination of desired non-binary datasets, grouping variables, and response variables. We will do this using foreach loops, which take advantage of R's ability to run intensive operations on multiple CPUs in parallel. In each case, the total number of iterations (the total number of dataset-variable triads) will equal the product of the number of datasets, the number of response variables, and the number of grouping variables. We have chosen informative subsets of these (specified by the orig_datasets list, and the ratio_vars and grouping_vars vectors) to minimize computation time. We will run one foreach-loop on the un-transformed data sets (orig_datasets), one foreach-loop on the boxcox-transformed data sets (boxcox_datasets), and one foreach-loop on the log10-transformed datasets (log10_datasets). Because these datasets are all the same length (each is a list with 14 items), the lengthof orig_datasets is used to specify the iteration number for all three foreach-loops. However, before running these foreach-loops, we must initialize the objects that will allow us to keep track of foreach-loop progress and outputs. We do this in the present chunk of code.

# Initialize output objects to save phyloBoxPlot test results:
total_iterations <- length(orig_datasets)*length(ratio_vars)*length(grouping_vars)
orig_phyloBoxPlot_object_list <- as.list(rep(NA, total_iterations))
orig_phyloBoxPlot_object_names <- rep(NA, total_iterations)
boxcox_phyloBoxPlot_object_list <- as.list(rep(NA, total_iterations))
boxcox_phyloBoxPlot_object_names <- rep(NA, total_iterations)
log10_phyloBoxPlot_object_list <- as.list(rep(NA, total_iterations))
log10_phyloBoxPlot_object_names <-rep(NA, total_iterations)

# Initialize maximum iteration number for each for-loop below:
imax <- length(orig_datasets)
jmax <- length(ratio_vars)
kmax <- length(grouping_vars)

# Now that we have initialized objects for all foreach loops, we can begin running them. Please be warned that the next few chunks are *very* computationally expensive. They will output thousands of plots, totaling several gigabytes, to your working directory. Depending on your computational resources and the value of iter_num above, they may also take many days to run.
cat("\n","PROGRESS REP: chunk [S2.2.05] complete; starting next chunk...","\n")
#*-----**-----*

#*-----**{ [S2.2.06] Run phyloBoxPlot() on all key dataset-variable combinations for non-binary raw (un-transformed) datasets.}*
# Run phyloBoxPlot() foreach loop on all non-binary (original) datasets:
orig_phyloBoxPlot_object_list <- foreach(i=1:imax, .combine='c', .packages = c("forcats", "data.table", "stringr", "ggplot2", "dplyr", "car", "RColorBrewer", "gridExtra", "ggimage", "vegan", "RVAideMemoire", "MASS", "pracma", "phytools", "ape", "ggtree", "strap", "geiger", "phytools")) %dopar% { 
  # Create an empty list to store results for each iteration of i:
  output_list_per_i <- as.list(rep(NA, jmax*kmax))
  
  # Run nested for-loop within foreach loop:
  for(j in 1:jmax){
    for(k in 1:kmax){
      # Calculate current iteration numbers: 
      current_full_loop_iteration_num <- (i-1)*jmax*kmax + (j-1)*kmax + k
      intra_i_iteration_num <- ((j-1) * kmax) + k
      # Report foreach-loop progress:
      cat("\n", "-----------------------------------------", "\n", "foreach-loop progress:", "\n", "ITERATION:", current_full_loop_iteration_num, "/", total_iterations, "\n", "DATASET:", i, "/", length(orig_datasets), "\n", "ratio_var:",  j, "/", length(ratio_vars), "\n", "grouping_var:", k, "/", length(grouping_vars), "\n", "-----------------------------------------", "\n")
      cat("\n", "~~~", "Current internal (intra-i) iteration number:", intra_i_iteration_num, "~~~", "\n")
      
      # Run phyloBoxPlot() on all original (untransformed) datasets:
      output_list_per_i[[intra_i_iteration_num]] <- phyloBoxPlot(ds=orig_datasets[[i]], dataset_name=names(orig_datasets)[i], tree_list=tree_variants, response_var=ratio_vars[j], grouping_var=grouping_vars[k], nreps=iter_num, is.binary=F)
    } # closes internal 'for(j in 1:jmax)' loop
  } # closes internal 'for(k in 1:kmax)' loop
  output_list_per_i # output from internal j and k loops (for foreach to combine)
} # closes external 'foreach(i=1:imax)' loop

# The output for the foreach-loop above is a fully populated list, called 'orig_phyloBoxPlot_object_list', which contains the test results and associated data and metadata for every run of the phyloBoxPlot() function, on every orig_dataset/ratio_var/grouping_var/tree combination. Next, we will use a regular for-loop to populate a vector of names with which we can name the items in that list.

# Populate orig_phyloBoxPlot_object_names vector:
for(i in 1:imax){
  for(j in 1:jmax){
    for(k in 1:kmax){
      current_full_loop_iteration_num <- (i-1)*jmax*kmax + (j-1)*kmax + k
      orig_phyloBoxPlot_object_names[current_full_loop_iteration_num] <- paste("DATASET:", orig_dataset_names[i], "|", "RESPONSE VARIABLE:", ratio_vars[j], ",", "GROUPING VARIABLE:", grouping_vars[k], sep=" ")
    }
  }
}

# Next, we name the items in the orig_phyloBoxPlot_object_list using our newly populated orig_phyloBoxPlot_object_names vector:
names(orig_phyloBoxPlot_object_list) <- orig_phyloBoxPlot_object_names

# Our R environment now contains a fully populated, named list of phylogenetic ANOVA and Levene's test results for every desired combination of orig_dataset, response variable, grouping variable, and supertree——with corresponding boxplots and test report tables saved as PNGs for every such combination saved to the working directory.
cat("\n","PROGRESS REP: chunk [S2.2.06] complete; starting next chunk...","\n")
#*-----**-----*

#*-----**{ [S2.2.07] Run phyloBoxPlot() function on all key dataset-variable combinations for non-binary BoxCox-transformed datasets.}*
# Run phyloBoxPlot() foreach loop on all non-binary (original) datasets:
boxcox_phyloBoxPlot_object_list <- foreach(i=1:imax, .combine='c', .packages = c("forcats", "data.table", "stringr", "ggplot2", "dplyr", "car", "RColorBrewer", "gridExtra", "ggimage", "vegan", "RVAideMemoire", "MASS", "pracma", "phytools", "ape", "ggtree", "strap", "geiger", "phytools")) %dopar% { 
  
  # Create an empty list to store results for each iteration of i:
  output_list_per_i <- as.list(rep(NA, jmax*kmax))
  
  # Run nested for-loop within foreach loop:
  for(j in 1:jmax){
    for(k in 1:kmax){
      # Calculate current iteration numbers: 
      current_full_loop_iteration_num <- (i-1)*jmax*kmax + (j-1)*kmax + k
      intra_i_iteration_num <- ((j-1) * kmax) + k
      # Report foreach-loop progress:
      cat("\n", "-----------------------------------------", "\n", "foreach-loop progress:", "\n", "ITERATION:", current_full_loop_iteration_num, "/", total_iterations, "\n", "DATASET:", i, "/", length(boxcox_datasets), "\n", "ratio_var:",  j, "/", length(ratio_vars), "\n", "grouping_var:", k, "/", length(grouping_vars), "\n", "-----------------------------------------", "\n")
      cat("\n", "~~~", "Current internal (intra-i) iteration number:", intra_i_iteration_num, "~~~", "\n")
      
      # Run phyloBoxPlot() on all original (untransformed) datasets:
      output_list_per_i[[intra_i_iteration_num]] <- phyloBoxPlot(ds=boxcox_datasets[[i]], dataset_name=boxcox_dataset_names[i], tree_list=tree_variants, response_var=ratio_vars[j], grouping_var=grouping_vars[k], nreps=iter_num, is.binary=F)
    } # closes 'for(k in 1:kmax)' loop
  } # closes 'for(j in 1:jmax)' loop
  output_list_per_i # output from internal j and k loops (for foreach to combine)
} # closes external 'foreach(i=1:imax)' loop

# The output for the foreach-loop above is a fully populated list, called 'boxcox_phyloBoxPlot_object_list', which contains the test results and associated data and metadata for every run of the phyloBoxPlot() function, on every boxcox_dataset/ratio_var/grouping_var/tree combination. Next, repeating the process we did above for the raw datasets, we will use a regular for-loop to populate a vector of names with which we can name the items in that list.

# Populate boxcox_phyloBoxPlot_object_names vector:
for(i in 1:imax){
  for(j in 1:jmax){
    for(k in 1:kmax){
      current_full_loop_iteration_num <- (i-1)*jmax*kmax + (j-1)*kmax + k
      boxcox_phyloBoxPlot_object_names[current_full_loop_iteration_num] <- paste("DATASET:", boxcox_dataset_names[i], "|", "RESPONSE VARIABLE:", ratio_vars[j], ",", "GROUPING VARIABLE:", grouping_vars[k], sep=" ")
    }
  }
}

# Much as we did before, we'll name the items in the boxcox_phyloBoxPlot_object_list using the newly populated boxcox_phyloBoxPlot_object_names vector:
names(boxcox_phyloBoxPlot_object_list) <- boxcox_phyloBoxPlot_object_names

# Our R environment now contains a fully populated, named list of phylogenetic ANOVA and Levene's test results for every desired combination of boxcox_dataset, response variable, grouping variable, and supertree——with corresponding boxplots and test report tables saved as PNGs for every such combination saved to the working directory.
cat("\n","PROGRESS REP: chunk [S2.2.07] complete; starting next chunk...","\n")
#*-----**-----*

#*-----**{ [S2.2.08] Run phyloBoxPlot() function on all key dataset-variable combinations for non-binary log10-transformed datasets.}*
# Run phyloBoxPlot() foreach loop on all non-binary (original) datasets:
log10_phyloBoxPlot_object_list <- foreach(i=1:imax, .combine='c', .packages = c("forcats", "data.table", "stringr", "ggplot2", "dplyr", "car", "RColorBrewer", "gridExtra", "ggimage", "vegan", "RVAideMemoire", "MASS", "pracma", "phytools", "ape", "ggtree", "strap", "geiger", "phytools")) %dopar% { 
  
  # Create an empty list to store results for each iteration of i:
  output_list_per_i <- as.list(rep(NA, jmax*kmax))
  
  # Run nested for-loop within foreach loop:
  for(j in 1:jmax){
    for(k in 1:kmax){
      # Calculate current iteration numbers: 
      current_full_loop_iteration_num <- (i-1)*jmax*kmax + (j-1)*kmax + k
      intra_i_iteration_num <- ((j-1) * kmax) + k
      # Report foreach-loop progress:
      cat("\n", "-----------------------------------------", "\n", "foreach-loop progress:", "\n", "ITERATION:", current_full_loop_iteration_num, "/", total_iterations, "\n", "DATASET:", i, "/", length(log10_datasets), "\n", "ratio_var:",  j, "/", length(ratio_vars), "\n", "grouping_var:", k, "/", length(grouping_vars), "\n", "-----------------------------------------", "\n")
      cat("\n", "~~~", "Current internal (intra-i) iteration number:", intra_i_iteration_num, "~~~", "\n")
      
      # Run phyloBoxPlot() on all log10-transformed datasets:
      output_list_per_i[[intra_i_iteration_num]] <- phyloBoxPlot(ds=log10_datasets[[i]], dataset_name=log10_dataset_names[i], tree_list=tree_variants, response_var=ratio_vars[j], grouping_var=grouping_vars[k], nreps=iter_num, is.binary=F)
    } # closes 'for(k in 1:kmax)' loop
  } # closes 'for(j in 1:jmax)' loop
  output_list_per_i # output from internal j and k loops (for foreach to combine)
} # closes external 'foreach(i=1:imax)' loop

# The output for the foreach-loop above is a fully populated list, called 'log10_phyloBoxPlot_object_list', which contains the test results and associated data and metadata for every run of the phyloBoxPlot() function, on every log10_dataset/ratio_var/grouping_var combination. Next, we will once again use a regular for-loop to populate a vector of names with which we can name the items in that list.

# Populate log10_phyloBoxPlot_object_names vector:
for(i in 1:imax){
  for(j in 1:jmax){
    for(k in 1:kmax){
      current_full_loop_iteration_num <- (i-1)*jmax*kmax + (j-1)*kmax + k
      log10_phyloBoxPlot_object_names[current_full_loop_iteration_num] <- paste("DATASET:", log10_dataset_names[i], "|", "RESPONSE VARIABLE:", ratio_vars[j], ",", "GROUPING VARIABLE:", grouping_vars[k], sep=" ")
    }
  }
}

# Name the items in the log10_phyloBoxPlot_object_list using the new vector:
names(log10_phyloBoxPlot_object_list) <- log10_phyloBoxPlot_object_names

# Our R environment now contains a fully populated, named list of phylogenetic ANOVA and Levene's test results for every desired combination of log10_dataset, response variable, grouping variable, and supertree——with corresponding boxplots and test report tables saved as PNGs for every such combination saved to the working directory.
cat("\n","PROGRESS REP: chunk [S2.2.08] complete; starting next chunk...","\n")
#*-----**-----*

# In the four preceding chunks of code, we ran the phyloBoxPlot() function on all non-binary datasets. In the next four, we will repeat this process on their binary counterparts (those datasets treated with the bLRify() function).

#*-----**{ [S2.2.09] Initialize objects for phyloBoxPlot foreach-loops of all binary datasets.}*
# In the next few chunks of code, we will run the phyloBoxPlot() function on every combination of desired BINARY datasets, grouping variables, and response variables. In each case, the total number of iterations (the total number of dataset-variable triads) will equal the product of the number of datasets, the number of response variables, and the number of grouping variables. We have chosen informative subsets of these (specified by the bLR_raw_datasets list, and the ratio_vars and key_bLR_groupers vectors) to minimize computation time. We will run one foreach loop on the untransformed data sets (bLR_raw_datasets), one foreach loop on the boxcox-transformed data sets (bLR_boxcox_datasets), and one foreach loop on the log10-transformed datasets (bLR_log10_datasets). Because these datasets are all the same length (each is a list with 14 items), length(orig_datasets) is used to specify the iteration number for all three foreach loops. As we did last time, we must (before actually running these foreach loops) must initialize the objects that will allow us to keep track of foreach-loop progress and outputs. We do this in the present chunk of code.

# Initialize output objects to save phyloBoxPlot test results:
key_bLR_groupers <- bLR_groupers[c(3:16)]
total_iterations <- length(bLR_raw_datasets)*length(ratio_vars)*length(key_bLR_groupers)
bin_orig_phyloBoxPlot_object_list <- as.list(rep(NA, total_iterations))
bin_orig_phyloBoxPlot_object_names <- rep(NA, total_iterations)
bin_boxcox_phyloBoxPlot_object_list <- as.list(rep(NA, total_iterations))
bin_boxcox_phyloBoxPlot_object_names <- rep(NA, total_iterations)
bin_log10_phyloBoxPlot_object_list <- as.list(rep(NA, total_iterations))
bin_log10_phyloBoxPlot_object_names <-rep(NA, total_iterations)

# Initialize maximum iteration number for each for-loop below:
imax <- length(bLR_raw_datasets)
jmax <- length(ratio_vars)
kmax <- length(key_bLR_groupers)

# Now that we have initialized objects for all foreach loops, we can begin running them. Please be warned that the next few chunks are *very* computationally expensive. They will output thousands of plots, totaling several gigabytes, to your working directory. Depending on your computational resources and the value of iter_num above, they may also take many days to run.
cat("\n","PROGRESS REP: chunk [S2.2.09] complete; starting next chunk...","\n")
#*-----**-----*

#*-----**{ [S2.2.10] Run phyloBoxPlot() function on all key dataset-variable combinations for binary raw (untransformed) datasets.}*
# Run phyloBoxPlot() foreach loop on all binary (original) datasets:
bin_orig_phyloBoxPlot_object_list <- foreach(i=1:imax, .combine='c', .packages = c("forcats", "data.table", "stringr", "ggplot2", "dplyr", "car", "RColorBrewer", "gridExtra", "ggimage", "vegan", "RVAideMemoire", "MASS", "pracma", "phytools", "ape", "ggtree", "strap", "geiger", "phytools")) %dopar% { 
  
  # Create an empty list to store results for each iteration of i:
  output_list_per_i <- as.list(rep(NA, jmax*kmax))
  
  # Run nested for-loop within foreach loop:
  for(j in 1:jmax){
    for(k in 1:kmax){
      # Calculate current iteration numbers: 
      current_full_loop_iteration_num <- (i-1)*jmax*kmax + (j-1)*kmax + k
      intra_i_iteration_num <- ((j-1) * kmax) + k
      # Report foreach-loop progress:
      cat("\n", "-----------------------------------------", "\n", "foreach-loop progress:", "\n", "ITERATION:", current_full_loop_iteration_num, "/", total_iterations, "\n", "DATASET:", i, "/", length(bLR_raw_datasets), "\n", "ratio_var:",  j, "/", length(ratio_vars), "\n", "grouping_var:", k, "/", length(key_bLR_groupers), "\n", "-----------------------------------------", "\n")
      cat("\n", "~~~", "Current internal (intra-i) iteration number:", intra_i_iteration_num, "~~~", "\n")
      
      # Run phyloBoxPlot() on all original (untransformed) datasets:
      output_list_per_i[[intra_i_iteration_num]] <- phyloBoxPlot(ds=bLR_raw_datasets[[i]], dataset_name=names(bLR_raw_datasets)[i], tree_list=tree_variants, response_var=ratio_vars[j], grouping_var=key_bLR_groupers[k], nreps=iter_num, is.binary=T)
    } # closes internal 'for(k in 1:kmax)' loop
  } # closes internal 'for(j in 1:jmax)' loop
  output_list_per_i # output from internal j and k loops (for foreach to combine)
} # closes external 'foreach(i=1:imax)' loop

# The output for the foreach-loop above is a fully populated list, called 'bin_orig_phyloBoxPlot_object_list', which contains the test results and associated data and metadata for every run of the phyloBoxPlot() function, on every orig_dataset/ratio_var/grouping_var combination. As we did for all non-binary datasets, we will use a regular for-loop to populate a vector of names with which we can name the items in that list.

# Populate bin_orig_phyloBoxPlot_object_names vector:
for(i in 1:imax){
  for(j in 1:jmax){
    for(k in 1:kmax){
      current_full_loop_iteration_num <- (i-1)*jmax*kmax + (j-1)*kmax + k
      bin_orig_phyloBoxPlot_object_names[current_full_loop_iteration_num] <- paste("DATASET:", bLR_raw_dataset_names[i], "|", "RESPONSE VARIABLE:", ratio_vars[j], ",", "GROUPING VARIABLE:", key_bLR_groupers[k], sep=" ")
    }
  }
}

# Name the items in the bin_orig_phyloBoxPlot_object_list using the newly populated names vector:
names(bin_orig_phyloBoxPlot_object_list) <- bin_orig_phyloBoxPlot_object_names

# Our R environment now contains a fully populated, named list of phylogenetic ANOVA and Levene's test results for every desired combination of orig_dataset, response variable, grouping variable, and supertree——with corresponding boxplots and test report tables saved as PNGs for every such combination saved to the working directory.
cat("\n","PROGRESS REP: chunk [S2.2.10] complete; starting next chunk...","\n")
#*-----**-----*

#*-----**{ [S2.2.11] Run phyloBoxPlot() function on all key dataset-variable combinations for binary BoxCox-transformed datasets.}*
# Run phyloBoxPlot() foreach loop on all binary BoxCox-transformed datasets:
bin_boxcox_phyloBoxPlot_object_list <- foreach(i=1:imax, .combine='c', .packages = c("forcats", "data.table", "stringr", "ggplot2", "dplyr", "car", "RColorBrewer", "gridExtra", "ggimage", "vegan", "RVAideMemoire", "MASS", "pracma", "phytools", "ape", "ggtree", "strap", "geiger", "phytools")) %dopar% { 
  
  # Create an empty list to store results for each iteration of i:
  output_list_per_i <- as.list(rep(NA, jmax*kmax))
  
  # Run nested for-loop within foreach loop:
  for(j in 1:jmax){
    for(k in 1:kmax){
      # Calculate current iteration numbers: 
      current_full_loop_iteration_num <- (i-1)*jmax*kmax + (j-1)*kmax + k
      intra_i_iteration_num <- ((j-1) * kmax) + k
      # Report foreach-loop progress:
      cat("\n", "-----------------------------------------", "\n", "foreach-loop progress:", "\n", "ITERATION:", current_full_loop_iteration_num, "/", total_iterations, "\n", "DATASET:", i, "/", length(bLR_boxcox_datasets), "\n", "ratio_var:",  j, "/", length(ratio_vars), "\n", "grouping_var:", k, "/", length(key_bLR_groupers), "\n", "-----------------------------------------", "\n")
      cat("\n", "~~~", "Current internal (intra-i) iteration number:", intra_i_iteration_num, "~~~", "\n")
      
      # Run phyloBoxPlot() on all original (untransformed) datasets:
      output_list_per_i[[intra_i_iteration_num]] <- phyloBoxPlot(ds=bLR_boxcox_datasets[[i]], dataset_name=bLR_boxcox_dataset_names[i], tree_list=tree_variants, response_var=ratio_vars[j], grouping_var=key_bLR_groupers[k], nreps=iter_num, is.binary=T)
    } # closes internal 'for(k in 1:kmax)' loop
  } # closes internal 'for(j in 1:jmax)' loop
  output_list_per_i # output from internal j and k loops (for foreach to combine)
} # closes external 'foreach(i=1:imax)' loop

# The output for the foreach-loop above is the boxcox_phyloBoxPlot_object_list, which contains the test results and associated data and metadata for every run of the phyloBoxPlot() function, on every boxcox_dataset/ratio_var/grouping_var/tree combination. Next, we will use a regular for-loop to populate a vector of names with which we can name the items in that list.

# Populate bin_boxcox_phyloBoxPlot_object_names vector:
for(i in 1:imax){
  for(j in 1:jmax){
    for(k in 1:kmax){
      current_full_loop_iteration_num <- (i-1)*jmax*kmax + (j-1)*kmax + k
      bin_boxcox_phyloBoxPlot_object_names[current_full_loop_iteration_num] <- paste("DATASET:", bLR_boxcox_dataset_names[i], "|", "RESPONSE VARIABLE:", ratio_vars[j], ",", "GROUPING VARIABLE:", key_bLR_groupers[k], sep=" ")
    }
  }
}

# Name the items in the bin_orig_phyloBoxPlot_object_list using the new vector:
names(bin_boxcox_phyloBoxPlot_object_list) <- bin_boxcox_phyloBoxPlot_object_names

# Our R environment now contains a fully populated, named list of phylogenetic ANOVA and Levene's test results for every desired combination of raw binarified dataset, response variable, grouping variable, and supertree——with corresponding boxplots and test report tables saved as PNGs for every such combination saved to the working directory.
cat("\n","PROGRESS REP: chunk [S2.2.11] complete; starting next chunk...","\n")
#*-----**-----*

#*-----**{ [S2.2.12] Run phyloBoxPlot() function on all key dataset-variable combinations for binary log10-transformed datasets.}*
# Run phyloBoxPlot() foreach loop on all binary log10-transformed datasets:
bin_log10_phyloBoxPlot_object_list <- foreach(i=1:imax, .combine='c', .packages = c("forcats", "data.table", "stringr", "ggplot2", "dplyr", "car", "RColorBrewer", "gridExtra", "ggimage", "vegan", "RVAideMemoire", "MASS", "pracma", "phytools", "ape", "ggtree", "strap", "geiger", "phytools")) %dopar% { 
  
  # Create an empty list to store results for each iteration of i:
  output_list_per_i <- as.list(rep(NA, jmax*kmax))
  
  # Run nested for-loop within foreach loop:
  for(j in 1:jmax){
    for(k in 1:kmax){
      # Calculate current iteration numbers: 
      current_full_loop_iteration_num <- (i-1)*jmax*kmax + (j-1)*kmax + k
      intra_i_iteration_num <- ((j-1) * kmax) + k
      # Report foreach-loop progress:
      cat("\n", "-----------------------------------------", "\n", "foreach-loop progress:", "\n", "ITERATION:", current_full_loop_iteration_num, "/", total_iterations, "\n", "DATASET:", i, "/", length(bLR_log10_datasets), "\n", "ratio_var:",  j, "/", length(ratio_vars), "\n", "grouping_var:", k, "/", length(key_bLR_groupers), "\n", "-----------------------------------------", "\n")
      cat("\n", "~~~", "Current internal (intra-i) iteration number:", intra_i_iteration_num, "~~~", "\n")
      
      # Run phyloBoxPlot() on all log10-transformed datasets:
      output_list_per_i[[intra_i_iteration_num]] <- phyloBoxPlot(ds=bLR_log10_datasets[[i]], dataset_name=bLR_log10_dataset_names[i], tree_list=tree_variants, response_var=ratio_vars[j], grouping_var=key_bLR_groupers[k], nreps=iter_num, is.binary=T)
    } # closes internal 'for(k in 1:kmax)' loop
  } # closes internal 'for(j in 1:jmax)' loop
  output_list_per_i # output from internal j and k loops (for foreach to combine)
} # closes external 'foreach(i=1:imax)' loop

# The output for the foreach-loop above is a fully populated list, called 'bin_log10_phyloBoxPlot_object_list', which contains the test results and associated data and metadata for every run of the phyloBoxPlot() function, on every log10_dataset/ratio_var/grouping_var combination. Next, we will use a regular for-loop to populate a vector of names with which we can name the items in that list.

# Populate bin_log10_phyloBoxPlot_object_names vector:
for(i in 1:imax){
  for(j in 1:jmax){
    for(k in 1:kmax){
      current_full_loop_iteration_num <- (i-1)*jmax*kmax + (j-1)*kmax + k
      bin_log10_phyloBoxPlot_object_names[current_full_loop_iteration_num] <- paste("DATASET:", bLR_log10_dataset_names[i], "|", "RESPONSE VARIABLE:", ratio_vars[j], ",", "GROUPING VARIABLE:", key_bLR_groupers[k], sep=" ")
    }
  }
}

# Name the items in the bin_orig_phyloBoxPlot_object_list using the new vector:
names(bin_log10_phyloBoxPlot_object_list) <- bin_log10_phyloBoxPlot_object_names

# Our R environment now contains a fully populated, named list of phylogenetic ANOVA and Levene's test results for every desired combination of log10_dataset, response variable, grouping variable, and supertree——with corresponding boxplots and test report tables saved as PNGs for every such combination saved to the working directory.
cat("\n","PROGRESS REP: chunk [S2.2.12] complete; starting next chunk...","\n")
#*-----**-----*

# This concludes the most statistically intensive portion of script_S2. In what follows, we supplement our results so far with a few more visual comparisons of morphometric differences across groups.
#------------------------------------------------------------------------------

#**"-------------------------------------------------------------------------"*
#**"- ### [S2.3] MAKE BOXPLOTS GROUPED BY SUBCLADE FOR BY-CLADE TRENDS ### -"*
#**"-------------------------------------------------------------------------"*

# In this section,we supplement our previous results with by-clade comparisons using grouped box plots.

#*-----**{ [S2.3.01] Define function to group boxplots by subclade. }*
boxes.byClade <- function(dataset, dataset_name, response_var, grouping_var, subclades_to_omit, outline_size, bp_width, bp_padding){
  # DESCRIBING KEY PARAMETERS:
  # vartype: chr string specifying type of data ("limb", "eco1", or "eco2")
  # this will determine the palette chosen for comparisons 
  
  # Set directory path for output files:
  directory_path <- paste0(output_path, "/boxplots_by_clade")
  dir.create(path=directory_path, showWarnings = TRUE, recursive = FALSE, mode = "0777")
  
  cat("\n", "boxes.byClade() function has started running for", response_var, "by", grouping_var, "\n", "in the", dataset_name, "dataset.", "\n")
  
  # Defining response_var and grouping_var within the function:
  dataset$response_var <- dataset[ , response_var]
  dataset$grouping_var <- dataset[ , grouping_var]
  
  # Removing all rows with missing data:
  ds_tr <- dataset[c(!is.na(dataset$response_var) & !is.na(dataset$grouping_var)), ]
  
  # Define tick labels:
  group_names <- levels(droplevels(na.omit(ds_tr$grouping_var)))
  nvals <- tapply(ds_tr$grouping_var, INDEX = droplevels(ds_tr$grouping_var), FUN = function(x) length(na.omit(x)))
  ticklabs_main <- paste(group_names, "\n n=", nvals, sep="")
  
  # Define palette:
  pal <- grouping_var_palette[[grouping_var]]
  cat("\n", "Generating boxPlot with subclade-partitions...", "\n")
  
  # Get row counts by Subclade and grouping_var
  counts <- table(ds_tr$Subclade, ds_tr$grouping_var)
  
  # Get levels of grouping_var
  levels_grouping <- levels(ds_tr$grouping_var)
  
  # Remove rows with Subclade levels in Subclades_to_cut:
  ds_clade_tr <- ds_tr
  for(i in 1:length(subclades_to_omit)){
    ds_clade_tr<-ds_clade_tr[!ds_clade_tr$Subclade %in% subclades_to_omit[i], ]
  } # closes 'for(i in 1:length(subclades_to_omit))' loop
  clade_names <- levels(droplevels(na.omit(ds_clade_tr$Subclade)))
  clade_nvals <- tapply(ds_clade_tr$grouping_var, INDEX = droplevels(ds_clade_tr$Subclade), FUN = function(x) length(na.omit(x)))
  ticklabs_subclade <- paste(clade_names, "\n n=", clade_nvals, sep="")
  
  # Make grouped box plot:
  bP_byClade <- ggplot(ds_clade_tr, aes(x=Subclade, y=(response_var), fill=grouping_var)) +
    geom_boxplot(size=outline_size, outlier.size=1, show.legend=T, alpha=0.7, position = position_dodge2(width = bp_width, padding = bp_padding, preserve = "single")) +
    theme(plot.margin=margin(t=20, r=30, b=40, l=30), aspect.ratio=3/2,
          axis.text=element_text(size=10, angle=90, hjust=1),
          axis.title=element_text(size=14), legend.position = "none",
          panel.grid.major = element_blank(),
          panel.grid.minor = element_blank(),
          axis.line = element_line(colour = "black"),
          panel.background = element_rect(fill = "transparent"),
          plot.background = element_rect(fill = "transparent")) + #ylim(-0.2,2) + 
    xlab(label=NULL) + ylab(label=response_var) +
    scale_x_discrete(labels = ticklabs_subclade) + 
    scale_fill_manual(values=pal)
  
  # Save high-resolution PNG of grouped boxplot:
  png(filename = paste0(directory_path, "/boxPlot__byClade_", dataset_name, "_", response_var,"_by_", grouping_var, ".png"), width=10, height=12, units="in", res=200)
  show(bP_byClade)
  dev.off()
  
  # Report completion of function for a given set of inputs:
  cat("\n", "boxes.byClade() function complete for", response_var, "by", grouping_var, "in", dataset_name, "dataset.", "\n")
} # closes boxes.byClade() function

# Now that we have defined a function to group morphometric results for a given categorical variable by clade, we will run this function on several key predictor_var-response_var combinations of interest.
cat("\n","PROGRESS REP: chunk [S2.3.01] complete; starting next chunk...","\n")
#*-----**-----*

#*-----**{ [S2.3.02] Run boxes.byClade() function on all_data for each key combination of predictor and non-binary grouping variables. }*

# Run function on key non-binary grouping_variable-dataset combinations:
  # Create a dataset without taxa that have ambiguous phenotypes:
    all_sure <- all_data
    all_sure$fore_type <- gsub("Unclear, terr.|Unclear, aquat.", NA, all_sure$fore_type)
    all_sure$hind_type <- gsub("Unclear, terr.|Unclear, aquat.", NA, all_sure$hind_type)
    all_sure$Ecotype1 <- gsub("Data Deficient", NA, all_sure$Ecotype1)
    all_sure$fore_type <- fct_relevel(all_sure$fore_type, "Unwebbed", "Webbed", "Flippered")
    all_sure$hind_type <- fct_relevel(all_sure$hind_type, "Unwebbed", "Webbed", "Flippered")
    all_sure$Ecotype1 <- fct_relevel(all_sure$Ecotype1, "Terrestrial", "Transiently Aquatic", "Moderately Aquatic", "Highly Aquatic", "Fully Aquatic")

boxes.byClade(dataset=all_sure, dataset_name="*all_sure", response_var="f_AZR", grouping_var="fore_type", subclades_to_omit = c("Outgroups", "Stem reptiles", "Euryapsida"), outline_size=1.2, bp_width=1.35, bp_padding=0.45)

boxes.byClade(dataset=all_sure, dataset_name="*all_sure", response_var="h_AZR", grouping_var="hind_type", subclades_to_omit = c("Outgroups", "Stem reptiles", "Euryapsida"), outline_size=1.2, bp_width=1.35, bp_padding=0.45)

boxes.byClade(dataset=all_sure, dataset_name="*all_sure", response_var="f_AZR", grouping_var="Ecotype1", subclades_to_omit = c("Outgroups", "Stem reptiles", "Euryapsida"), outline_size=1.2, bp_width=1.35, bp_padding=0.45)

boxes.byClade(dataset=all_sure, dataset_name="*all_sure", response_var="h_AZR", grouping_var="Ecotype1", subclades_to_omit = c("Outgroups", "Stem reptiles", "Euryapsida"), outline_size=1.2, bp_width=1.35, bp_padding=0.45)

boxes.byClade(dataset=all_data, dataset_name="all", response_var="f_AZR", grouping_var="fore_type", subclades_to_omit = c("Outgroups"), outline_size=0.8, bp_width=0.6, bp_padding=0.4)

boxes.byClade(dataset=all_data, dataset_name="all", response_var="h_AZR", grouping_var="hind_type", subclades_to_omit = c("Outgroups"), outline_size=0.8, bp_width=0.6, bp_padding=0.4)

boxes.byClade(dataset=all_data, dataset_name="all", response_var="f_AZR", grouping_var="Ecotype1", subclades_to_omit = c("Outgroups"), outline_size=0.8, bp_width=0.6, bp_padding=0.4)

boxes.byClade(dataset=all_data, dataset_name="all", response_var="h_AZR", grouping_var="Ecotype1", subclades_to_omit = c("Outgroups"), outline_size=0.8, bp_width=0.6, bp_padding=0.4)

boxes.byClade(dataset=all_data, dataset_name="all", response_var="f_AZR", grouping_var="Ecotype2", subclades_to_omit = c("Outgroups"), outline_size=0.8, bp_width=0.6, bp_padding=0.4)

boxes.byClade(dataset=all_data, dataset_name="all", response_var="h_AZR", grouping_var="Ecotype2", subclades_to_omit = c("Outgroups"), outline_size=0.8, bp_width=0.6, bp_padding=0.4)

boxes.byClade(dataset=all_data, dataset_name="all", response_var="f_AZR", grouping_var="Ecotype3", subclades_to_omit = c("Outgroups"), outline_size=0.8, bp_width=0.6, bp_padding=0.4)

boxes.byClade(dataset=all_data, dataset_name="all", response_var="h_AZR", grouping_var="Ecotype3", subclades_to_omit = c("Outgroups"), outline_size=0.8, bp_width=0.6, bp_padding=0.4)
cat("\n","PROGRESS REP: chunk [S2.3.02] complete; starting next chunk...","\n")
#*-----**-----*

#*-----**{ [S2.3.03] Run boxes.byClade() function on all_bLR for each key combination of quantitative response and binary grouping variables. }*
# Run function on all binarified datasets:
for(j in 1:length(key_bLR_groupers)){
  for(k in 1:length(ratio_vars)){boxes.byClade(dataset=all_bLR, dataset_name="all_bLR", response_var=ratio_vars[k], grouping_var=key_bLR_groupers[j], subclades_to_omit = c("Outgroups", "Euryapsida", "Stem reptiles"), outline_size=1.4, bp_width=0.8, bp_padding=0.4)
  } # closes 'for(k in 1:length(ratio_vars))' loop
} # closes 'for(j in 1:length(key_bLR_groupers))' loop
cat("\n","PROGRESS REP: chunk [S2.3.03] complete; starting next chunk...","\n")
#*-----**-----*

# This concludes our analysis of how each individual quantitative variable differs among groups in our dataset. In the next section of script_S1, we look at how multiple quantitative variables differ at once among groups. In particular, we look at how limb region (stylopod, zeugopod, and acropod) proportions differ among the levels of each categorical variable within ternary morphospace. This next step is mainly to visualize differences in limb proportions holistically, but does not currently include a phylogenetic component, as this would be computationally expensive and not add much above and beyond what we have just completed in [S2.2].
#------------------------------------------------------------------------------

#**"-------------------------------------------------------------------------"*
#**"-- ### [[S2.4]] COMPARE LIMB PROPs BY GROUP IN TERNARY MORPHOSPACE ### --"*
#**"-------------------------------------------------------------------------"*

#*-----**{ [S2.4.01] Define phyloTernPlot() function to make ternary plots. }*
phyloTernPlot <- function(dataset, dataset_name, input_tree, tree_name, n_iterations, fore_or_hind, ternset_vars, grouping_var, pointsize, alpha_val, showtext=FALSE){
  # FUNCTION ARGUMENTS:
  # dataset: dataset to be fed into the ternary plot function; note that this data frame should NOT have been pre-processed by rm.NAs() or reorder.data(), as all of this processing takes place within the present function.
  # dataset_name: chr string giving name of dataset (to enable output file naming)
  # input_tree: object of class "phylo" used as input for aov.phylo() function to run phyloMANOVA; note that this tree should NOT have been trimmed by trim.tree(), as this takes place within the present function.
  # tree_name: chr string giving name of tree (to enable output file naming)
  # n_iterations: numeric value giving number of iterations for aov.phylo() function to run phyloMANOVA
  # fore_or_hind: chr string equaling either "fore" or "hind", to specify whether this ternary diagram will depict forelimb proportions or hindlimb proportions.
  # ternset_vars: a vector of three chr strings naming the axes of the ternary plot
  # grouping_var: a chr string naming the response variable
  # pointsize: a numeric value specifying the desired size of points on the ternary plot
  # alpha_val: a numeric value specifying the desired opacity of points on the ternary plot
  
  # Report function progress:
  cat("\n","Defining variables within function and checking assumptions...", "\n")
  proceed_verdict <- "yes" # This is the starting state for a dummy variable used to determine whether the function can proceed with or should skip the current dataset-variable combination.
  
  # Set directory path for output files:
  directory_path <- paste0(output_path, "/phyloTernPlots")
  dir.create(path=directory_path, showWarnings = TRUE, recursive = FALSE, mode = "0777")
  
  # Set ternary axis labels:
  if(fore_or_hind == "fore"){tern_labs <- c("Hand (%)", "Zeugopod (%)", "Humerus (%)")}
  if(fore_or_hind == "hind"){tern_labs <- c("Foot (%)", "Tibia (%)", "Femur (%)")}
  
  # Define ternary variables within function:
  dataset$grouping_var <- dataset[ , grouping_var]
  ternvar1 <- ternset_vars[1] 
  ternvar2 <- ternset_vars[2] 
  ternvar3 <- ternset_vars[3] 
  dataset$ternvar1 <- dataset[ , ternvar1]
  dataset$ternvar2 <- dataset[ , ternvar2]
  dataset$ternvar3 <- dataset[ , ternvar3]
  
  # Remove taxa with "unclear" phenotypes:
  for(i in 1:nrow(dataset)){
    if(grepl("eficient", dataset$grouping_var[i])==T | grepl("Unclear", dataset$grouping_var[i])==T | grepl("Ambiguous", dataset$grouping_var[i])==T){
      dataset$grouping_var[i] <- NA # This for-loop converts all specimens groupings marked "Unclear, aquat., "Unclear, terr", "Ambiguous", "Data Deficient", or "data deficient" to NA so they'll be removed and not clutter the ternary plot.
    }
  } # closes 'for(i in 1:nrow(dataset))' loop
  
  # Mark rows with and without quantitative data:
  rows_with_quant_data <- rep(NA, (nrow(dataset)))
  for(i in 1:nrow(dataset)){
    if(is.na(dataset$ternvar1[i])==T | is.na(dataset$ternvar2[i])==T | is.na(dataset$ternvar3[i])==T){
      rows_with_quant_data[i] <- NA
    } else {
      rows_with_quant_data[i] <- "Has data"
    }
  } # closes 'for(i in 1:nrow(dataset))' loop
  
  rows_missing_quant_data <- rep(NA, (nrow(dataset)))
  for(i in 1:nrow(dataset)){
    if(is.na(dataset$ternvar1[i])==T | is.na(dataset$ternvar2[i])==T | is.na(dataset$ternvar3[i])==T){
      rows_missing_quant_data[i] <- 1
    } else {
      rows_missing_quant_data[i] <- 0
    }
  } # closes 'for(i in 1:nrow(dataset))' loop
  
  if(sum(rows_missing_quant_data)==length(rows_missing_quant_data)){ 
    # ie, if all rows of the dataset are missing at least one of their ternary coordinate values
    cat("\n", "phyloTernPlot() has skipped the following dataset-variable combination:", "\n", "\t", "dataset:", dataset_name, "\n", "\t", "grouping_var:", grouping_var, "\n", "REASON: None of the rows with grouping_var data and ternary limb proportion data overlap;", "\n", "thus, trimming with rm.NAs() appropriately left the dataset empty.", "\n")
    proceed_verdict <- "no"
  } # closes first 'if (all rows missing for one ternary coord )' clause
  
  # Trim dataset to remove rows with NAs:
  ds_tr <- dataset[c(!is.na(dataset[ , ternvar1]) & !is.na(dataset[ , ternvar2]) & !is.na(dataset[ , ternvar3]) & !is.na(dataset[, grouping_var])), ]
  
  if(length(ds_tr$grouping_var)<1){
    # ie, if all rows of the dataset are missing at least one of their ternary coordinate values
    cat("\n", "phyloTernPlot() has skipped the following dataset-variable combination:", "\n", "\t", "dataset:", dataset_name, "\n", "\t", "grouping_var:", grouping_var, "\n", "REASON: None of the rows with grouping_var data and ternary limb proportion data overlap;", "\n", "thus, trimming with rm.NAs() appropriately left the dataset empty.", "\n")
    proceed_verdict <- "no"
  } # closes second 'if (all rows missing for one ternary coord )' clause
  
  # Set number of groups remaining:
  ds_tr$grouping_var <- droplevels(ds_tr$grouping_var)
  num_groups <- length(levels(droplevels(ds_tr$grouping_var)))
  
  ds_tern <- ds_tr[, c("grouping_var", "ternvar1", "ternvar2", "ternvar3")]
  ds_tern <- ds_tern[complete.cases(ds_tern), ]
  
  if(num_groups>1){ # ie, if there are fewer than two factors with ternary data remaining
    proceed_verdict2 <- "yes"
  } else if (num_groups<=1){
    proceed_verdict2 <- "no"
    cat("\n", "phyloTernPlot() has skipped the following dataset-variable combination:", "\n", "\t", "dataset:", dataset_name, "\n", "\t", "grouping_var:", grouping_var, "\n", "REASON: Fewer than two group levels with data remained after trimming NAs.", "\n")
  } # closes 'if(num_groups>1) { } else { }' clause
  
  if(proceed_verdict=="yes" & proceed_verdict2=="yes"){ # then proceed with rest of function.
    # Define nvals object:
    group_names <- levels(droplevels(ds_tr$grouping_var))
    nvals <- tapply(ds_tr$grouping_var, INDEX = ds_tr$grouping_var, FUN = function(x) length(na.omit(x)))
    
    # Fix group-color pairings to be robust against missing data: 
    group_colors <- rep(NA, num_groups)
    for(q in 1:length(group_colors)){
      group_q <- group_names[q]
      if(nvals[q]>0){group_colors[q] <- grouping_var_palette[[grouping_var]][group_q]}
    } # closes 'for(q in 1:length(group_colors))' loop
    
    # Below, we define a function (which was written with the help of chatGPT) to assign alpha values using hexadecimal codes, because AddToTernary() doesn't typically recognize alpha arguments.
    color_to_hex_with_alpha <- function(color_names, alpha = 0.5) {
      # Convert color names to RGB values
      rgb_values <- col2rgb(color_names)
      # Convert RGB values to hexadecimal codes
      hex_codes <- rgb(rgb_values[1, ], rgb_values[2, ], rgb_values[3, ], maxColorValue = 255)
      # Add alpha value to the hexadecimal codes
      hex_with_alpha <- paste0(hex_codes, sprintf("%02X", round(alpha * 255)))
      return(hex_with_alpha)
    } # closes color_to_hex_with_alpha() function
    
    color_hexcodes_w_alpha <- color_to_hex_with_alpha(group_colors, alpha = alpha_val)
    fill_colors <- color_hexcodes_w_alpha
    palette(fill_colors)
    
    # Prepare to save png of ternPlot:
    cat("\n","Saving ternary plot and legend as high-resolution PNGs to output directory...", "\n")
    png(filename=paste0(directory_path, "/ternPlot__", dataset_name, "_by_", grouping_var, "__", fore_or_hind, ".png", sep=""), width=12, height=12, units="in", res=600)
    
    # Create and save the ternary plot:
    TernaryPlot(alab=as.character(tern_labs[1]), blab=as.character(tern_labs[2]), clab=as.character(tern_labs[3]), lab.cex=1.2, padding=0.08, grid.lines=10, grid.minor.lines=30, grid.col="lightgray", grid.minor.col="gray95")
    AddToTernary(points, ds_tern[, c("ternvar1", "ternvar2", "ternvar3")], pch=24, cex=pointsize, bg = ds_tern[, "grouping_var"], col = "grey3", lwd = (pointsize/1.5))
    title(main=paste(dataset_name, "_by_", grouping_var, "_ternPlot_", sep=""))
    legend("topright", legend = group_names, pch=24, pt.bg=fill_colors, border="white", pt.cex=1, cex=0.8, bty = "n", title=grouping_var, text.font=2, bg='lightgray', box.lwd=1)
    if(showtext==T){AddToTernary(text, tern_coords, ds_tr$Tip_label, cex=0.2)}
    dev.off()
    cat("\n","phyloTernPlot function successfully run for ", dataset_name, " by ", grouping_var, "\n", "\n")
  } # closes 'if(proceed_verdict=="yes" & proceed_verdict2=="yes")' clause
} # closes phyloTernPlot() function

# Now that we have defined the phyloTernPlot function, we will test it with a series of challenging inputs to make sure that it works. 
cat("\n","PROGRESS REP: chunk [S2.4.01] complete; starting next chunk...","\n")
#*-----**-----*

#*-----**{ [S2.4.02] Test phyloTernPlot() with difficult input combinations. }*
# Define axis label vectors to help automate downstream phyloTernPlot() calls:
ftern_vars <- c("Hand", "EFZL", "Humerus")
htern_vars <- c("Foot", "Tibia", "Femur")

# Here we test the phyloTernPlot() function to make sure it works, using input combinations that previously produced errors while trouble-shooting.

phyloTernPlot(dataset=orig_datasets[[1]], dataset_name="***test_", input_tree=tree_variants[[1]], tree_name="supertree1", n_iterations=10, fore_or_hind="fore", ternset_vars=ftern_vars, grouping_var="Ecotype1", pointsize=2, alpha_val=0.5, showtext=FALSE)

phyloTernPlot(dataset=archos_all, dataset_name="***test_", input_tree=tree_variants[[1]], tree_name="supertree1", n_iterations=10, fore_or_hind="fore", ternset_vars=ftern_vars, grouping_var="fore_type", pointsize=2, alpha_val=0.7, showtext=FALSE)

phyloTernPlot(dataset=orig_datasets[[6]], dataset_name="***test_no_overlap", input_tree=tree_variants[[1]], tree_name="supertree1", n_iterations=10, fore_or_hind="fore", ternset_vars=ftern_vars, grouping_var="IUCN_cat", pointsize=2, alpha_val=0.5, showtext=FALSE) # should skip without error interruption even though there's no overlap between grouping_var rows with data and limb proportion rows with data.

phyloTernPlot(dataset=orig_datasets[[7]], dataset_name="***test_unclear", input_tree=tree_variants[[1]], tree_name="supertree1", n_iterations=10, fore_or_hind="fore", ternset_vars=ftern_vars, grouping_var="IUCN_cat", pointsize=2, alpha_val=0.5, showtext=FALSE) # should skip without error interruption even though there are fewer than two levels with data remaining.

phyloTernPlot(dataset=orig_datasets[[8]], dataset_name="***test_just_one_level", input_tree=tree_variants[[1]], tree_name="supertree1", n_iterations=10, fore_or_hind="fore", ternset_vars=ftern_vars, grouping_var="IUCN_cat", pointsize=2, alpha_val=0.5, showtext=FALSE) # should skip without error interruption even though there are fewer than two levels with data remaining.
cat("\n","PROGRESS REP: chunk [S2.4.02] complete; starting next chunk...","\n")
#*-----**-----*

#*-----**{ [S2.4.03] Run phyloTernPlot() on all fore- and hindlimb datasets. }*
# Define forelimb and hindlimb grouping variables to help automate downstream phyloTernPlot() function calls:
fore_groups <- c("Ecotype1", "Ecotype2", "Ecotype3", "fore_type")
hind_groups <- c("Ecotype1", "Ecotype2", "Ecotype3", "hind_type")

more_orig_datasets <- list(all_data, archos_all, turts_all, lepidos_all, stemreps_all, nonplacs_all, atlanto_all, euarchontagl_all, ferae_all, ungulates_all)
more_orig_dataset_names <- c("all_data", "archos_all", "turts_all", "lepidos_all", "stemreps_all", "nonplacs_all", "atlanto_all", "euarchontagl_all", "ferae_all", "ungulates_all")

# Run phyloTernPlot() on orig_datasets for all forelimb grouping variables:
for(i in 1:length(more_orig_datasets)){
  for(j in 1:length(fore_groups)){
    phyloTernPlot(dataset=more_orig_datasets[[i]], dataset_name=more_orig_dataset_names[i], input_tree=tree_variants[[1]], tree_name="supertree1", fore_or_hind="fore", ternset_vars=ftern_vars, grouping_var=fore_groups[j], pointsize=2, alpha_val=0.7, showtext=FALSE, n_iterations=iter_num)
  } # NB: In order to perform a phyloPERMANOVA, we would need another for-loop layer to iterate through tree_variants as well.
} # closes 'for(i in 1:length(orig_datasets))'

# Run phyloTernPlot() on orig_datasets for all hindlimb grouping variables:
for(i in 1:length(more_orig_datasets)){
  for(j in 1:length(hind_groups)){
    phyloTernPlot(dataset=more_orig_datasets[[i]], dataset_name=more_orig_dataset_names[i], input_tree=tree_variants[[1]], tree_name="supertree1", fore_or_hind="hind", ternset_vars=htern_vars, grouping_var=hind_groups[j], pointsize=2, alpha_val=0.5, showtext=FALSE, n_iterations=iter_num)
  } # NB: In order to perform a phyloPERMANOVA, we would need another for-loop layer to iterate through tree_variants as well.
} # closes 'for(i in 1:length(orig_datasets))'
cat("\n","PROGRESS REP: chunk [S2.4.03] complete; starting next chunk...","\n")
#*-----**-----*

# Please note that the functions above did not involve phylogenetic statistical tests (eg, phylogenetic PERMANOVAs). Again, we did not perform these as they would be computationally expensive and provide little information above and beyond what we obtained from phylogenetic ANOVAs and Levene's tests (see script_S2). However, the interested researcher could implement a phylogenetic PERMANOVA into the phyloTernPlot() function, and we have designed the function keeping in mind the potential to add this extra functionality in the future. 
#------------------------------------------------------------------------------

#**"-------------------------------------------------------------------------"*
#**"------ ### [S2.5] SAVE R ENVIRONMENT FOR DOWNSTREAM SCRIPT S5  ### ------"*
#**"-------------------------------------------------------------------------"*

# In the final section of script_S2, we will save the script's R environment and report how long the script took to run. This saved environment file will enable us to perform critical sensitivity analyses in downstream script_S5, with all the necessary data objects and test results.

#*-----**{ [S2.5.01] Save current R environment to working directory. }*
save.image(file=paste0(wd_path, "/R_ENVIR_for_script_S2.RData"))
cat("\n","PROGRESS REP: chunk [S2.5.01] complete; starting next chunk...","\n")
#*-----**-----*

#*-----**{ [S2.5.02] Record the time that was required for script_S2 to run. }*
# Record running time metrics for entire script:
proctime_output <- proc.time()
elapsed_secs <- proctime_output[3]
elapsed_mins <- elapsed_secs/60
elapsed_hrs <- elapsed_mins/60
elapsed_days <- elapsed_hrs/24
cat("\n","PROGRESS REP: chunk [S2.5.02] complete; starting next chunk...","\n")
#*-----**-----*

#*-----**{ [S2.5.03] Record CPU usage for this script. }*
# The following code calculates CPU usage efficiency based on the following Stack Overflow thread: https://stackoverflow.com/questions/73217110/cpu-usage-measurement-and-calculation-in-r-using-proc-time.
CPU_number <- CPU_num # Recall that this is the number of CPUs used for the current script.
CPU_time_used <- proctime_output[1] + proctime_output[2]
CPU_utilization <- (proctime_output[1] / CPU_time_used) * 100 / CPU_number
cat("\n","PROGRESS REP: chunk [S2.5.03] complete; starting next chunk...","\n")
#*-----**-----*

#*-----**{ [S2.5.04] Save time-elapsed and CPU metrics to a text file. }*
time_report_filepath <- paste0(output_path, "/", "TIME_REPORT_for_script_S2.txt")
sink(time_report_filepath)
cat("TIME TAKEN FOR ENTIRE SCRIPT TO RUN", "\n", elapsed_secs, "seconds", "\n", "\t", "=", elapsed_mins, "minutes", "\n", "\t", "=", elapsed_hrs, "hours", "\n", "\t", "=", elapsed_days, "days", "\n")
cat("\n", "CPU USAGE:", "\n", "\t", "Number of CPUs used:",  CPU_number, "\n", "\t", "CPU utilization efficiency:", CPU_utilization, "%", "\n")
sink(file=NULL) # A text file saving the time elapsed for the completion of this script has now been saved to the working directory.
cat("\n","PROGRESS REP: chunk [S2.5.04] complete; script_S2 has finished running!","\n")
#*-----**-----*

# That concludes script_S2 in our data analysis pipeline. In this script, we ran all phylogenetic comparative tests involved in assessing how single quantitative variables differed among groups. In the downstream script S5, after scripts S2-S4 are all done running, you will see whether these test results differed by data-transformation method and tree topology. I hope you have enjoyed this coding journey so far, and that you continue onward to script_S5! In the meantime, if you have any questions about this script or its associated data, or if you would like to discuss a possible collaboration, please feel free to contact me anytime via the address below.

# -- Caleb Gordon
#------------------------------------------------------------------------------
```
