## Supplementary R Scripts for "Limb proportions predict aquatic habits and soft-tissue flippers in extinct amniotes": script_S3.R

```
#** Third R script (script_S3.R) for Gordon et al., "Limb proportions predict aquatic habits and soft-tissue flippers in extinct amniotes."**

# NOTE ON READABILITY: This script contains many long, multi-line comments. To increase readability within Rstudio, go to Tools > Global Options, select the 'Code' tab, and check 'Soft-wrap R source files'. This will wrap all lines of code to match your personal GUI's margins, so that no lines of code run offscreen.

# NOTE ON SCRIPT ORDER: This script is downstream of script_S1.R, upstream of script_S5, and concurrent (can be run in parallel) with scripts S2-S4. If you have not finished running script_S1.R, please close this file and run script_S1.R to completion. Once script_S1.R is complete, you can reopen the current file and proceed. 

# INTRODUCTION: In the previous R script (script_S1.R), we ran all preliminary data- and tree-processing steps required for more intensive downstream statistical analyses. The current script (script_S3.R) is run many times as part of a job array, in which we perform pairwise phylogenetic correlation tests for all variable pairs in our raw (un-transformed), log10-transformed, and BoxCox-transformed datasets. These tasks are very, very computationally intensive, and moderate parallelization (in the form of foreach-loops) was insufficient to accelerate computation time enough to match our timeframe. Consequently, the S3 scripts are run in a more massively parallel fashion using Job Arrays. 

# This is the outline of the sections for script_S3:
#**[S3] PERFORM PAIRWISE PHYLOGENETIC CORRELATION TESTS**
# --- [S3.1]: PREPARE CODING ENVIRONMENT: Load pkgs, S1 R environment, etc.
# --- [S3.2]: RUN PAIRWISE CORRELATION TESTS FOR ALL NON-UNITLESS DATA.
# --- -- NOTE: This is repeated for raw, log-trans, and BoxCox-trans datasets.
# --- [S3.3]: SAVE CORRELATION TEST RESULTS & COMPUTATION TIME FOR SCRIPT_S5.

# We go through each of these sections with a series of makeshift code "chunks" below, starting with [S3.1].
#------------------------------------------------------------------------------

#**"-------------------------------------------------------------------------"*
#**"-------------- ### [S3.1] PREPARE CODING ENVIRONMENT ### ---------------"*
#**"-------------------------------------------------------------------------"*

#*-----**{ [S3.1.01] Load required packages. }*
# NOTE: Since all packages were installed for completion of script_S1, we do not repeat the optional installation lines for these packages here; however, lines of code to install the packages below are provided in script_S1 [S1.1.01].

# Data object manipulation:
library(forcats) # for releveling factors with fct_relevel() function
library(data.table) # for converting objects to data tables and matrices
library(plyr) # useful for in-task parallelization
library(dplyr) # useful for in-task parallelization

# Data visualization and exporting:
library(ggplot2) # for effective data visualization with ggplot suite
library(MESS) # for specifying translucent colors as function arguments
library(car) # for normal quantile plot construction
library(RColorBrewer) # for defining fancy color palettes
library(gridExtra) # useful for generating multi-panel plots and tables
library(ggimage) # For exporting ggplots
library(corrplot) # For correlogram construction

# Phylogenetic comparative methods and tree visualization:
library(ape) # for basic tree manipulation and phylogenetic comparative methods
library(ggtree) # for advanced tree visualization using ggplot2 suite
library(phylolm) # contains phyloglm() and phylolm() functions
cat("\n","PROGRESS REP: chunk [S3.1.01] complete; starting next chunk..","\n")
#*-----**-----*

#*-----**{ [S3.1.02] Load script_S1 environment. }*
# Define working directory path:
wd_path <- "/gpfs/gibbs/project/bhullar/cmg89/Flipper_Project/"
setwd(wd_path); getwd()
# Load script_S1 environment from working directory:
load(file=paste0(wd_path, "/R_ENVIR_for_script_S1.RData"))
cat("\n","PROGRESS REP: chunk [S3.1.02] complete; starting next chunk..","\n")
#*-----**-----*

#*-----**{ [S3.1.03] Define phycorr() wrapper function.}*
# Below, we define a wrapper function to run corphylo(), and all required upstream functions, on every pairwise combination of quantitative variables in our dataset. This will make it easier for us to parallelize the function using job arrays.

# Specify corphylo parameters for downstream usage:
REML_verdict <- TRUE
optim_method <- "Nelder-Mead"
constraind_verdict <- TRUE
maxitNM_num <- 500 # set to 2 for troubleshooting, and 500 for actual batch job

# Write phycorr() function:
phycorr <- function(input_ds, input_ds_name, corvar1, corvar2, input_tree_name, output_name, method = "Nelder-Mead", REML, constrain.d, maxit.NM, verbose=FALSE){
  # FUNCTION PARAMETERS:
  # input_ds is an input data frame containing corvar1 and corvar2
  # input_ds_name is a chr string naming the input dataset.
  # corvar1 and corvar2 are chr strings specifying the input_ds variables to be correlated with the corphylo function.
  # input_tree_name is a chr string specifying the name of the input tree (must be name of item in tree_variants list).
  # output_name is a chr string specifying the name of the phycorr output object for downstream indexing.
  # method is a chr string specifying whether to use "Nelder-Mead" or "SANN" minimization (specifying corphylo arg of the same name). We default to "Nelder-Mead", and additional modifications would be required to make this wrapper function work with SANN minimization.
  # REML is either TRUE or FALSE, specifies whether REML is used for function instead of ML. We default to TRUE, and additional dditional modifications would be required to make this wrapper function work with ML.
  # constrain.d is either TRUE or FALSE, specifying whether to constrain the d parameter to be btwn 0 and 1 (specifying the corphylo arg of the same name).
  # maxit.NM is a numeric value specifying the maximum number of iterations in the optimization with Nelder-Mead minimization (specifying the corphylo arg of the same name).
  # verbose specifies whether to print intermediate function outputs to the console. 
  
  # Skip variables with perfect correlation 
  # NOTE: This avoids a potential error in chol.default(cov(Xs)).
  if(corvar1==corvar2){
    phycorr_output <- "phycorr skipped because corvar1 = corvar2"
  } else { # continue with function
    
    # Define tree and corvars within function:
    input_tree <- tree_variants[[input_tree_name]]
    input_ds$corvar1 <- input_ds[, corvar1]
    input_ds$corvar2 <- input_ds[, corvar2]
    
    # Trim & align dataset and tree:
    ds_tr <- rm.NAs(dataset=input_ds, categorical_var = corvar1, quantitative_var = corvar2)
    tree_tr <- trim.tree(tree=input_tree, trimmed_dataset = ds_tr, verbose = verbose)
    ds_ord <- reorder.data(trimmed_tree=tree_tr, trimmed_dataset=ds_tr, verbose = verbose)
    ds_ord_cor <- ds_ord[, all_quantitative_vars]
    rownames(ds_ord_cor) <- ds_ord$Tip_label
    
    # Run phylogenetic correlation test:
    phycorr_output <- corphylo(X = ds_ord_cor[, c(corvar1, corvar2)], U = NULL, SeM = NULL, phy=tree_tr, REML = REML, method = method, constrain.d = constrain.d, maxit.NM = maxit.NM, verbose = FALSE)   
    # Record sample size of trimmed and aligned dataset:
    sample_size <- nrow(ds_ord_cor)
    phycorr_output$sample_size <- sample_size
  } # closes 'if(corvar1==corvar2){ } else { }' clause
  
  # Record corvars, output name, & input tree associated w/ phycorr output obj:
  phycorr_output$corvar1 <- corvar1
  phycorr_output$corvar2 <- corvar2
  phycorr_output$input_dataset <- input_ds_name
  output_name <- paste("phycorr output |", corvar1, "x", corvar2, "| DATASET:", input_ds_name, "| TREE:", input_tree_name)
  phycorr_output$output_name <- output_name
  phycorr_output$input_tree <- input_tree_name
  return(phycorr_output)
} # closes phycorr function

# Test phycorr() function:
corvar1_string <- "f_AZR"
corvar2_string <- "Humerus"
test_output1 <- phycorr(input_ds=all_mm, input_ds_name="all_mm", corvar1=corvar1_string, corvar2=corvar2_string, input_tree_name = "supertree1", method = optim_method,  REML=REML_verdict, constrain.d=constraind_verdict, maxit.NM=2)

corvar1_string <- "f_SZR"
corvar2_string <- "Tibia"
test_output2 <- phycorr(input_ds=all_mm, input_ds_name="all_mm", corvar1=corvar1_string, corvar2=corvar2_string, input_tree_name = "supertree3", method = optim_method, REML=REML_verdict, constrain.d=constraind_verdict, maxit.NM=2)

corvar1_string <- "Femur"
corvar2_string <- "Femur"
test_output3 <- phycorr(input_ds=all_mm, input_ds_name="all_mm", corvar1=corvar1_string, corvar2=corvar2_string, input_tree_name = "supertree5", method = optim_method,  REML=REML_verdict, constrain.d=constraind_verdict, maxit.NM=2)
cat("\n","PROGRESS REP: chunk [S3.1.03] complete; starting next chunk..","\n")
#*-----**-----*

#*-----**{ [S3.1.04] Set output directory for correlation test results. }*
cortest_output_path <- paste0(output_path, "/phyloCorrPlots")
dir.create(path=cortest_output_path, showWarnings = TRUE, recursive = FALSE, mode = "0777")
cat("\n","PROGRESS REP: chunk [S3.1.04] complete; starting next chunk..","\n")
#*-----**-----*

# We have now finished setting up the coding environment, and defined the single new custom function, for script_S3. In the next sections of the script, we will use our new phycorr() function to perform pairwise phylogenetic correlation tests on raw, log10-transformed, and BoxCox-transformed datasets.
#------------------------------------------------------------------------------

#**"-------------------------------------------------------------------------"*
#**"-- ### [[S3.2]] PERFORM PAIRWISE CORRELATION TESTS FOR ALL MM DATA ### --"*
#**"-------------------------------------------------------------------------"*

#*-----**{ [S3.2.01] Specify input arguments for this run of Job Arrays. }*
args = commandArgs(trailingOnly=TRUE)
treevar_i = tree_variant_names[as.numeric(args[1])]
corvar_j = corvariables[as.numeric(args[2])]
corvar_k = corvariables[as.numeric(args[3])]
cat("\n","PROGRESS REP: chunk [S3.2.01] complete; starting next chunk..","\n")
#*-----**-----*

#*-----**{ [S3.2.02] Run phycorr() on raw (un-trans) data for this job. }*
phycorr_raw_output <- phycorr(input_ds=all_mm, input_ds_name="all_mm", input_tree_name=treevar_i,corvar1=corvar_j,corvar2=corvar_k,method=optim_method, REML=REML_verdict, constrain.d=constraind_verdict, maxit.NM=maxitNM_num)
cat("\n","PROGRESS REP: chunk [S3.2.02] complete; starting next chunk..","\n")
#*-----**-----*

#*-----**{ [S3.2.03] Run phycorr() on log10-transformed data for this job. }*
phycorr_log10_output <- phycorr(input_ds=log10_mm, input_ds_name="log10_mm", input_tree_name=treevar_i,corvar1=corvar_j,corvar2=corvar_k,method=optim_method, REML=REML_verdict, constrain.d=constraind_verdict, maxit.NM=maxitNM_num)
cat("\n","PROGRESS REP: chunk [S3.2.03] complete; starting next chunk..","\n")
#*-----**-----*

#*-----**{ [S3.2.04] Run phycorr() on BoxCox-transformed data for this job. }*
phycorr_boxcox_output <- phycorr(input_ds=boxcox_mm, input_ds_name="boxcox_mm", input_tree_name=treevar_i,corvar1=corvar_j,corvar2=corvar_k,method=optim_method, REML=REML_verdict, constrain.d=constraind_verdict, maxit.NM=maxitNM_num)
cat("\n","PROGRESS REP: chunk [S3.2.04] complete; starting next chunk..","\n")
#*-----**-----*

#*-----**{ [S3.2.05] Record estimate of computation time for this job. }*
# Record running time metrics for entire script:
proctime_output <- proc.time()
elapsed_secs <- proctime_output[3]
elapsed_mins <- elapsed_secs/60
comp_time_per_dataset <- elapsed_mins / 3 # We divide the total time by three since we ran the computationally intensive phycorr() function three times--once with raw data, once with log10-trans data, and once with BoxCox-trans data.
cat("\n","PROGRESS REP: chunk [S3.2.05] complete; starting next chunk..","\n")
#*-----**-----*

#**"-------------------------------------------------------------------------"*
#**"-- ### [[S3.3]] SAVE CORRELATION TEST RESULTS & COMPUTATION TIME. ### ---"*
#**"-------------------------------------------------------------------------"*

#*-----**{ [S3.3.01] Save phycorr() results for this job. }*
# Initialize output objects:
phycorr_output_list <- list("phycorr_raw_output"=phycorr_raw_output, "phycorr_log10_output"=phycorr_log10_output, "phycorr_boxcox_output"=phycorr_boxcox_output)

phycorr_output_metadata <- data.frame(matrix(ncol=29, nrow=3))
colnames(phycorr_output_metadata) <- c("input_dataset", "input_tree", "corvariable1", "corvariable2", "sample size", "REML", "optim. method", "constrain.d", "maxit.NM", "Rsq", "regression equation", "intercept", "slope", "intercept StdErr", "slope StdErr", "intercept Z-score", "slope Z-score", "intercept p-val", "slope p-val", "logLik", "AIC", "BIC", "param. d1", "param. d2", "computation time", "skipped?", "reason_for_skipping", "JOB_ARRAY_ID", "JOB_TASK_ID")

# Save phylogenetic correlation test results for downstream sensitivity tests:
for(i in 1:length(phycorr_output_list)){
  # Save metadata for current phycorr_output_list element that applies regardless of whether the test was carried out:
  phycorr_output_metadata[i, "input_dataset"] <- phycorr_output_list[[i]]$input_dataset
  phycorr_output_metadata[i, "input_tree"] <- phycorr_output_list[[i]]$input_tree
  phycorr_output_metadata[i,"corvariable1"] <- phycorr_output_list[[i]]$corvar1
  phycorr_output_metadata[i,"corvariable2"] <- phycorr_output_list[[i]]$corvar2
  phycorr_output_metadata[i,"JOB_ARRAY_ID"]<-Sys.getenv(x="SLURM_ARRAY_JOB_ID") # This is the alphanumeric string ID for the Slurm Job Array in which the current row of metadata was produced. It should be the same for all rows in this CSV, and is recorded to support debugging/troubleshooting.
  phycorr_output_metadata[i,"JOB_TASK_ID"]<-Sys.getenv(x="SLURM_ARRAY_TASK_ID") # This is the alphanumeric string ID for the job WITHIN the current Job Array, by which the current row of metadata was produced. Each row in this CSV should correspond to a different job (or "task") within the job array and therefore have a different JOB_TASK_ID value. We record this value here for debugging/troubleshooting purposes.

  if (class(phycorr_output_list[[i]][[1]][1])=="character") { 
    # i.e., if the test was skipped for these two variables...
    # ...then save metadata for any SKIPPED phylogenetic correlation test:
    phycorr_output_metadata[i, c(5:25)] <- NA
    phycorr_output_metadata[i, "skipped?"] <- "Yes"
    phycorr_output_metadata[i, "reason_for_skipping"] <- phycorr_output_list[[i]]
  } else if (!(class(phycorr_output_list[[i]][[1]][1])=="character")){ 
    # i.e., if the test was NOT skipped for these two variables...
    # ...then save metadata for any COMPLETED phylogenetic correlation test:
    phycorr_output_metadata[i, "sample size"] <- phycorr_output_list[[i]]$sample_size
    phycorr_output_metadata[i, "REML"] <- REML_verdict
    phycorr_output_metadata[i, "optim. method"] <- optim_method
    phycorr_output_metadata[i, "constrain.d"] <- constraind_verdict
    phycorr_output_metadata[i, "maxit.NM"] <- maxitNM_num
    phycorr_output_metadata[i, "Rsq"]<-phycorr_output_list[[i]]$cor.matrix[1,2]
    regression_equation <- paste0("y = ", round(phycorr_output_list[[i]]$B[2], 2), "x + ", round(phycorr_output_list[[i]]$B[1], 2)) # for the regression line
    phycorr_output_metadata[i, "regression equation"] <- regression_equation
    phycorr_output_metadata[i, "intercept"] <- phycorr_output_list[[i]]$B[1] 
    phycorr_output_metadata[i, "slope"] <- phycorr_output_list[[i]]$B[2] 
    phycorr_output_metadata[i, "intercept StdErr"] <- phycorr_output_list[[i]]$B.se[1]
    phycorr_output_metadata[i,"slope StdErr"]<-phycorr_output_list[[i]]$B.se[2]
    phycorr_output_metadata[i, "intercept Z-score"] <- phycorr_output_list[[i]]$B.zscore[1]
    phycorr_output_metadata[i, "slope Z-score"] <- phycorr_output_list[[i]]$B.zscore[2]
    phycorr_output_metadata[i, "intercept p-val"] <- phycorr_output_list[[i]]$B.pvalue[1]
    phycorr_output_metadata[i, "slope p-val"] <- phycorr_output_list[[i]]$B.pvalue[2]
    phycorr_output_metadata[i,  "logLik"] <- phycorr_output_list[[i]]$logLik
    phycorr_output_metadata[i, "AIC"] <- phycorr_output_list[[i]]$AIC
    phycorr_output_metadata[i, "BIC"] <- phycorr_output_list[[i]]$BIC
    phycorr_output_metadata[i, "param. d1"] <- phycorr_output_list[[i]]$d[1]
    phycorr_output_metadata[i, "param. d2"] <- phycorr_output_list[[i]]$d[2]
    phycorr_output_metadata[i, "computation time"] <- comp_time_per_dataset
    phycorr_output_metadata[i, "skipped?"] <- "No"
    phycorr_output_metadata[i, "reason_for_skipping"] <- NA
  } # closes big if-else loop for specifying metadata
} # closes 'for(i in 1:length(phycorr_output_list))' loop

# Save phycorr_output_metadata to a CSV file w/in phyloCorrPlots subdirectory:
write.csv(phycorr_output_metadata, paste0(cortest_output_path, "/phycorr_metadata - ", treevar_i, " - ", corvar_j, " - ", corvar_k, ".csv"), row.names=FALSE)
cat("\n","PROGRESS REP: chunk [S3.3.01] complete; script_S3 is finished running!","\n")
#*-----**-----*

# We have just saved a table containing the phycorr() test results from one run of this job array, which includes three rows--one run on raw data, one on log10-transformed data, and one run on BoxCox-transformed data--for the same input tree and variable pair. Running this script in massively parallel fashion with Job Arrays will generate many such output CSV files, which we will stitch together in script_S5. If you have any questions about this script or its associated data, please feel free to contact me anytime via the address below.

# -- Caleb Gordon
#------------------------------------------------------------------------------
```
