## Supplementary R Scripts for "Limb proportions predict aquatic habits and soft-tissue flippers in extinct amniotes": script_S4.R

```
#** Fourth R script (script_S4.R) for Gordon et al., "Limb proportions predict aquatic habits and soft-tissue flippers in extinct amniotes."**

# NOTE ON READABILITY: This script contains many long, multi-line comments. To increase readability within Rstudio, go to Tools > Global Options, select the 'Code' tab, and check 'Soft-wrap R source files'. This will wrap all lines of code to match your personal GUI's margins, so that no lines of code run offscreen.

# NOTE ON SCRIPT ORDER: This script is downstream of script_S1.R, upstream of script_S5, and concurrent (can be run in parallel) with scripts S2-S4. If you have not yet finished running script_S1.R, please close this file and run script_S1.R to completion. Once script_S1.R is complete, you can reopen the current file and proceed. 

# INTRODUCTION: In the upstream R script S1, we ran all preliminary data- and tree-processing steps required for more intensive downstream statistical analyses. In the current script (script_S4.R), we use an original phylogenetically informed machine-learning pipeline to fit phylogenetic logistic regression models to the data for all raw (un-transformed) datasets in order to predict binary phenotype classifications and assess the relative accuracies of different logistic regression models. Scripts S4b and S4c do the same with our log10-transformed and BoxCox-transformed datasets, respectively. These tasks are very, very computationally intensive, and moderate parallelization (in the form of foreach-loops) was insufficient to accelerate computation time to match our timeframe. Consequently, the S4 scripts are run in a massively parallel fashion using Job Arrays. Please note that running this script on job arrays will potentially output thousands of files (totaling several Gigabytes) to your working directory; please proceed with caution and make space accordingly.

# This is the outline of sections for script_S4:
#**[S4] FIT PHYLOGENETIC LOGISTIC REGRESSION MODELS TO DATA**
# --- [S4.1]: PREPARE CODING ENVIRONMENT: Load pkgs, S1 R environment, etc.
# --- [S4.2]: FIT PHYLOGEN. BINOMIAL LOGISTIC REGRESSION MODELS TO DATASETS.
# --- [S4.3]: SAVE PHYBLR TEST RESULTS & COMPUTATION TIME FOR SCRIPT_S5.

# We go through each of these sections with a series of makeshift code "chunks" below, starting with [S4.1].
#------------------------------------------------------------------------------

#**"-------------------------------------------------------------------------"*
#**"-------------- ### [S4.1] PREPARE CODING ENVIRONMENT ### ---------------"*
#**"-------------------------------------------------------------------------"*

#*-----**{ [S4.1.01] Load required packages. }*
# NOTE: Since all packages were installed in script_S1, we do not repeat the optional installation lines for these packages here; however, lines of code to install the following packages are provided in script_S1, in section [S1.1.01].

# Data object manipulation:
library(forcats) # for releveling factors with fct_relevel() function
library(data.table) # for converting objects to data tables and matrices
library(plyr) # useful for in-task parallelization
library(dplyr) # useful for in-task parallelization
library(stringr) # for parcing through long chr strings (useful to check tree topologies in Nwk format) and using str_detect() function

# Data visualization and exporting:
library(ggplot2) # for effective data visualization with ggplot suite
library(MESS) # for specifying translucent colors as function arguments
library(car) # for normal quantile plot construction
library(RColorBrewer) # for defining fancy color palettes
library(gridExtra) # useful for generating multi-panel plots and tables
library(ggimage) # For exporting ggplots
library(corrplot) # For correlogram construction

# Data analysis:
library(devtools) # required for downloading some other packages from github
library(vcd) # for Cramer's V and other association stats
library(MASS) # used to make Box-Cox transformations with boxcox() function
library(pROC) # contains functions to plot and extract data from ROC curves
library(pracma) # contains diag() and trapz() functions; diag() is useful for making phylogenetic variance-covariance matrices; trapz() is useful for calculating ROC curve AUC.

# Phylogenetic comparative methods and tree visualization:
library(ape) # for basic tree manipulation and phylogenetic comparative methods
library(ggtree) # for advanced tree visualization using ggplot2 suite
library(phytools) # contains phylANOVA() function
library(geiger) # contains aov.phylo() function
library(phylolm) # contains phyloglm() and phylolm() functions
library(vegan) # contains adonis2(), which can run phyloPERMANOVAs
cat("\n","PROGRESS REP: chunk [S4.1.01] complete; starting next chunk..","\n")
#*-----**-----*

#*-----**{ [S4.1.02] Load script_S1 environment. }*
# Define working directory path:
wd_path <- "/gpfs/gibbs/project/bhullar/cmg89/Flipper_Project/"
setwd(wd_path); getwd()
# Load script_S1 environment from working directory:
load(file=paste0(wd_path, "/R_ENVIR_for_script_S1.RData"))
cat("\n","PROGRESS REP: chunk [S4.1.02] complete; starting next chunk..","\n")
#*-----**-----*

# We have now set up the coding environment for script_S4. In the next section of the script, we will run the bulk of its data analysis. 
#-----------------------------------------------------------------------------

# Scripts S1-S3 made plots and run tests to describe the relationships between different variables in the dataset. In script_S4, we will use the binarified datasets generated with bLRify() in S1 to fit phylogenetic binomial logistic regression (phybLR) models to the data. These models will give us predicted probabilities based on our limb and limb bone measurements, which we will use to predict the aquatic habits and soft-tissue limb morphologies of contentious fossil taxa in our dataset. In script_S5, we will compare the predictive accuracies of these models and select their best phenotype classifying threshold using Receiver-Operating Characteristic (ROC) curve analysis. 

# As described in script_S1, ROC analysis is a method for visualizing and comparing predictive model performance. This method was developed by U.S. military statisticians in World War II. We describe our usage of ROC analysis in more detail within our paper's Methods section, and a great introduction to the approach is given by Fawcett (2006) [doi:10.1016/j.patrec.2005.10.010]. In brief, ROC curves provide a quick way to richly visualize, transparently report, and clearly compare the performance of multiple competing predictive models--in our case, phylogenetic binomial logistic regression (phybLR) models.

# In what follows, we will fit a phylogenetic binomial logistic regression model to the dataset for each dataset-variable combination [using the phybLR() function]. In script S5, we will select the most accurate of these predictive models for each binary response variable, for each series of metrics (forelimb region proportions, hindlimb region proportions, forelimb acropod dimensions, hindlimb acropod dimensions, mD3 proximal phalanx dimensions, pD3 proximal phalanx dimensions, mD3 ungual dimensions, pD3 ungual dimensions), by using original functions that incorporate ROC analysis.  

#**"-------------------------------------------------------------------------"*
#**"-- ### [[S4.2]] RUN PHYLOGEN. BINOM. LOGISTIC REGRESSION ANALYSES ### --"*
#**"-------------------------------------------------------------------------"*

#*-----**{ [S4.2.01] Initialize data objects for all phybLR() runs. }*
# In the S4 scripts, we will ultimately run the phybLR() function on every combination of desired datasets, supertrees, grouping variables, and response variables. We will do this using Job Arrays. In each case, the total number of iterations (the total number of dataset-tree-variable sets) will equal the product of the number of datasets, the number of tree variants, the number of response variables, and the number of grouping variables. We previously chose informative subsets of these datasets and variables (specified now by the bLR_raw_datasets list, and the baremin_ratio_vars and key_bLR_groupers vectors) to minimize computation time. In script S4 (the one you're working through now), we will run one Job Array on the untransformed data sets (bLR_raw_datasets). In script S4b, we will run one on the log10-transformed data sets (bLR_boxcox_datasets). In script S4c, we will run one on the log10-transformed datasets (bLR_log10_datasets). Because these datasets are all the same length (each is a list with 14 items), length(bLR_raw_datasets) is used to specify the iteration number for all three foreach loops. Before running these foreach loops, we must initialize the objects that will allow us to keep track of foreach-loop progress and outputs. We do this in the present chunk of code.

# Specify parameter values for phybLR() runs:
n_bootstraps <- 10000 # set=3 for troubleshooting, 10000 for final batch job
n_CVruns <- 100 # set=3 for troubleshooting, 100 for final batch job
btol_val <- 35 # set=35 for cluster (values above this lead to this error from phyloglm():"edge.length[i] is NaN. Please reduce btol and/or log.alpha.bound.")
Kfold_val <- 3
cat("\n","PROGRESS REP: chunk [S4.2.01] complete; starting next chunk..","\n")
#*-----**-----*

#*-----**{ [S4.2.02] Specify input arguments for this run of Job Arrays. }*
args = commandArgs(trailingOnly=TRUE)
ds_name_i = bLR_raw_dataset_names[as.numeric(args[1])]
treename_j = tree_variant_names[as.numeric(args[2])]
groupvar_k = key_bLR_groupers[as.numeric(args[3])]
predvar_L = baremin_ratio_vars[as.numeric(args[4])]
cat("\n","PROGRESS REP: chunk [S4.2.02] complete; starting next chunk..","\n")
#*-----**-----*

#*-----**{ [S4.2.03] Run phybLR() on raw (un-trans) data for this job. }*
# Run upstream functions:
  bLR_tr <- rm.NAs(dataset=bLR_raw_datasets[[ds_name_i]], categorical_var=groupvar_k, quantitative_var=predvar_L)
  phylo_tr<-trim.tree(tree=tree_variants[[treename_j]],trimmed_dataset=bLR_tr,verbose=F)
  ds_ord<-reorder.data(trimmed_tree=phylo_tr,trimmed_dataset=bLR_tr,verbose=F)
  
# Run phybLR() function:
  phybLR_current_run_output1 <- phybLR(dataset_ordered=ds_ord, dataset_name=ds_name_i, response=groupvar_k, predictor=predvar_L, trimmed_tree=phylo_tr, tree_name=treename_j, boot_num=n_bootstraps, btol_num=btol_val, Kfold=Kfold_val, CV_runs=n_CVruns, crossval_reports=T, plot_td_ROCCs=T, plot_McF=F, plot_Youden=F, plot_CIs=T)
  
# Specify input dataset name for metadata compilation:
  phybLR_current_run_output1$input_dataset_name <- ds_name_i
cat("\n","PROGRESS REP: chunk [S4.2.03] complete; starting next chunk..","\n")
#*-----**-----*

#*-----**{ [S4.2.04] Run phybLR() on log10-transformed data for this job. }*
# Revise ds_name_i input arg for this run to specify log10 version of dataset:
  ds_name_i = bLR_log10_dataset_names[as.numeric(args[1])]

# Run upstream functions:
  bLR_tr <- rm.NAs(dataset=bLR_log10_datasets[[ds_name_i]], categorical_var=groupvar_k, quantitative_var=predvar_L)
  phylo_tr<-trim.tree(tree=tree_variants[[treename_j]],trimmed_dataset=bLR_tr,verbose=F)
  ds_ord<-reorder.data(trimmed_tree=phylo_tr,trimmed_dataset=bLR_tr,verbose=F)

# Run phybLR() function:
  phybLR_current_run_output2 <- phybLR(dataset_ordered=ds_ord, dataset_name=ds_name_i, response=groupvar_k, predictor=predvar_L, trimmed_tree=phylo_tr, tree_name=treename_j, boot_num=n_bootstraps, btol_num=btol_val, Kfold=Kfold_val, CV_runs=n_CVruns, crossval_reports=T, plot_td_ROCCs=T, plot_McF=F, plot_Youden=F, plot_CIs=T)

# Specify input dataset name for metadata compilation:
  phybLR_current_run_output2$input_dataset_name <- ds_name_i
cat("\n","PROGRESS REP: chunk [S4.2.04] complete; starting next chunk..","\n")
#*-----**-----*

#*-----**{ [S4.2.05] Run phybLR() on BoxCox-transformed data for this job. }*
# Revise ds_name_i input arg for this run to specify log10 version of dataset:
  ds_name_i = bLR_boxcox_dataset_names[as.numeric(args[1])]

# Run upstream functions:
  bLR_tr <- rm.NAs(dataset=bLR_boxcox_datasets[[ds_name_i]], categorical_var=groupvar_k, quantitative_var=predvar_L)
  phylo_tr<-trim.tree(tree=tree_variants[[treename_j]],trimmed_dataset=bLR_tr,verbose=F)
  ds_ord<-reorder.data(trimmed_tree=phylo_tr,trimmed_dataset=bLR_tr,verbose=F)

# Run phybLR() function:
  phybLR_current_run_output3 <- phybLR(dataset_ordered=ds_ord, dataset_name=ds_name_i, response=groupvar_k, predictor=predvar_L, trimmed_tree=phylo_tr, tree_name=treename_j, boot_num=n_bootstraps, btol_num=btol_val, Kfold=Kfold_val, CV_runs=n_CVruns, crossval_reports=T, plot_td_ROCCs=T, plot_McF=F, plot_Youden=F, plot_CIs=T)
  
# Specify input dataset name for metadata compilation:
  phybLR_current_run_output3$input_dataset_name <- ds_name_i
  
cat("\n","PROGRESS REP: chunk [S4.2.05] complete; starting next chunk..","\n")
#*-----**-----*

#*-----**{ [S4.2.06] Record estimate of computation time for this job. }*
# Record running time metrics for entire script:
proctime_output <- proc.time()
elapsed_secs <- proctime_output[3]
elapsed_mins <- elapsed_secs/60
number_phybLRs_successfully_run <- as.numeric((length(phybLR_current_run_output1)>2)==T) + as.numeric((length(phybLR_current_run_output2)>2)==T) + as.numeric((length(phybLR_current_run_output3)>2)==T) # This will give us the number, from 1 to 3, of phybLR runs that were NOT skipped in this script. This number will help us estimate the computation time required for each completed application of phybLR() in this run. 
comp_time_per_dataset <- elapsed_mins / number_phybLRs_successfully_run # We divide the total computation time required in the current script by the number of phybLR() calls resulting in completed phybLR tests (a maximum of 3) to approximate the computation time required for each individual phybLR() call.
cat("\n","PROGRESS REP: chunk [S4.2.06] complete; starting next chunk..","\n")
#*-----**-----*

#**"-------------------------------------------------------------------------"*
#**"--- ### [[S4.3]] SAVE PHYBLR OUTPUT RESULTS & COMPUTATION TIME. ### ----"*
#**"-------------------------------------------------------------------------"*

#*-----**{ [S4.3.01] Stitch together and save phybLR() results for this job. }*
# Set generic dataset name for all datasets in this run:
ds_name_generic <- gsub("boxcox", "", ds_name_i)

# Define directory path for output files: 
phybLR_directory_path <- paste0(output_path, "/phybLR_models")

# Define output list containing all phybLR output objects from this run:
bLR_output_list <- list("phybLR_raw_output"=phybLR_current_run_output1, "phybLR_log10_output"=phybLR_current_run_output2, "phybLR_boxcox_output"=phybLR_current_run_output3)

# Save bLR_output_list for current run to working directory:
save(bLR_output_list, file=paste0(phybLR_directory_path, "/phybLR_list - ", ds_name_generic, " - ", treename_j, " - ", groupvar_k, " - ", predvar_L, ".RData")) 

# Define output table to store the associated metadata for this run:
phybLR_run_metadata <- data.frame(matrix(ncol=49, nrow=3))
colnames(phybLR_run_metadata) <- c("input_dataset", "input_tree", "response_variable", "predictor_variable", "Kfold", "CV_runs", "phyloglm_bootnum", "phyloglm_btol", "AUC_mean", "AUC_95ci_lowbound", "AUC_95ci_highbound", "AUC_range", "AUC_binomt_pval", "McF_Rsq_mean", "McF_pval_mean", "McF_Rsq_95ci_lowb", "McF_Rsq_95ci_highb", "McF_pval_95ci_lowb", "McF_pval_95ci_highb", "McF_Rsq_for_full_ds", "McF_pval_for_full_ds", "AIC", "Youdens_Jmax", "bestmod_threshold", "bestmod_binomt_pval", "n_obs", "bestmod_TPR_mean", "bestmod_TPR_95ci_lowb", "bestmod_TPR_95ci_highb", "bestmod_TPR_range_lowb", "bestmod_TPR_range_highb", "bestmod_FPR_mean", "bestmod_FPR_95ci_lowb", "bestmod_FPR_95ci_highb", "bestmod_FPR_range_lowb", "bestmod_FPR_range_highb", "bestmod_TNR_mean", "bestmod_TP_mean", "bestmod_FP_mean", "bestmod_TN_mean", "bestmod_FN_mean", "bestmod_nSuccesses_mean", "bestmod_nFailures_mean", "bestmod_Accuracy_mean", "comput_time (min)", "skipped?", "reason_for_skipping", "SLURM_JOB_ARRAY_ID", "ARRAY_TASK_NUMBER")

# Save this run's phybLR inputs & results for downstream sensitivity tests:
for(i in 1:length(bLR_output_list)){
  # Report phybLR() function inputs and job metadata:
    phybLR_run_metadata[i, "input_dataset"] <- bLR_output_list[[i]]$input_dataset_name
    phybLR_run_metadata[i, "input_tree"] <- treename_j
    phybLR_run_metadata[i, "response_variable"] <- groupvar_k
    phybLR_run_metadata[i, "predictor_variable"] <- predvar_L
    phybLR_run_metadata[i, "Kfold"] <- Kfold_val
    phybLR_run_metadata[i, "CV_runs"] <- n_CVruns
    phybLR_run_metadata[i, "phyloglm_bootnum"] <- n_bootstraps
    phybLR_run_metadata[i, "phyloglm_btol"] <- btol_val
    phybLR_run_metadata[i, "SLURM_JOB_ARRAY_ID"] <- Sys.getenv(x = "SLURM_ARRAY_JOB_ID") # This is the alphanumeric string ID for the Slurm Job Array in which the current row of metadata was produced. It should be the same for all rows in this CSV, and is recorded to support debugging/troubleshooting.
    phybLR_run_metadata[i, "ARRAY_TASK_NUMBER"] <- Sys.getenv(x = "SLURM_ARRAY_TASK_ID") # This is the alphanumeric string ID for the job WITHIN the current Job Array, by which the current row of metadata was produced. Each row in this CSV should correspond to a different job (or "task") within the job array and therefore have a different JOB_TASK_ID value. We record this value here for debugging/troubleshooting purposes.
    
  # Report phybLR test results (if test was completed):
    if(is.list(bLR_output_list[[i]])==T & length(bLR_output_list[[i]])>2){ # that is, if test could be performed...
      phybLR_run_metadata[i, "AUC_mean"] <- bLR_output_list[[i]]$AUC_mean
      phybLR_run_metadata[i, "AUC_95ci_lowbound"] <- as.numeric(bLR_output_list[[i]]$AUC_95ci[1])
      phybLR_run_metadata[i, "AUC_95ci_highbound"] <- as.numeric(bLR_output_list[[i]]$AUC_95ci[2])
      phybLR_run_metadata[i, "AUC_range"] <- bLR_output_list[[i]]$AUC_range
      phybLR_run_metadata[i, "AUC_binomt_pval"] <- bLR_output_list[[i]]$AUC_binomtest_pval
      phybLR_run_metadata[i, "McF_Rsq_mean"] <- bLR_output_list[[i]]$McF_R_mean
      phybLR_run_metadata[i, "McF_pval_mean"] <- bLR_output_list[[i]]$McF_p_mean
      phybLR_run_metadata[i, "McF_Rsq_95ci_lowb"] <- bLR_output_list[[i]]$McF_R_confint[1]
      phybLR_run_metadata[i, "McF_Rsq_95ci_highb"] <- bLR_output_list[[i]]$McF_R_confint[2]
      phybLR_run_metadata[i, "McF_pval_95ci_lowb"] <- bLR_output_list[[i]]$McF_p_confint[1]
      phybLR_run_metadata[i, "McF_pval_95ci_highb"] <- bLR_output_list[[i]]$McF_p_confint[2]
      phybLR_run_metadata[i, "McF_Rsq_for_full_ds"] <- bLR_output_list[[i]]$McF_Rsq_for_full_ds
      phybLR_run_metadata[i, "McF_pval_for_full_ds"] <- bLR_output_list[[i]]$McF_pval_for_full_ds
      phybLR_run_metadata[i, "AIC"] <- bLR_output_list[[i]]$AIC
      phybLR_run_metadata[i, "Youdens_Jmax"] <- bLR_output_list[[i]]$Youdens_Jmax
      phybLR_run_metadata[i, "bestmod_threshold"] <- bLR_output_list[[i]]$bestmod_threshold
      phybLR_run_metadata[i, "bestmod_binomt_pval"] <- bLR_output_list[[i]]$bestmod_binomtest_pval
      phybLR_run_metadata[i, "n_obs"] <- bLR_output_list[[i]]$n_obs
      phybLR_run_metadata[i, "bestmod_TPR_mean"] <- bLR_output_list[[i]]$bestmod_TPR_mean
      phybLR_run_metadata[i, "bestmod_TPR_95ci_lowb"] <- as.numeric(bLR_output_list[[i]]$bestmod_TPR_95ci[1])
      phybLR_run_metadata[i, "bestmod_TPR_95ci_highb"] <- as.numeric(bLR_output_list[[i]]$bestmod_TPR_95ci[2])
      phybLR_run_metadata[i, "bestmod_TPR_range_lowb"] <- bLR_output_list[[i]]$bestmod_TPR_range[1]
      phybLR_run_metadata[i, "bestmod_TPR_range_highb"] <- bLR_output_list[[i]]$bestmod_TPR_range[2]
      phybLR_run_metadata[i, "bestmod_FPR_mean"] <- bLR_output_list[[i]]$bestmod_FPR_mean
      phybLR_run_metadata[i, "bestmod_FPR_95ci_lowb"] <- as.numeric(bLR_output_list[[i]]$bestmod_FPR_95ci[1])
      phybLR_run_metadata[i, "bestmod_FPR_95ci_highb"] <- as.numeric(bLR_output_list[[i]]$bestmod_FPR_95ci[2])
      phybLR_run_metadata[i, "bestmod_FPR_range_lowb"] <- bLR_output_list[[i]]$bestmod_FPR_range[1]
      phybLR_run_metadata[i, "bestmod_FPR_range_highb"] <- bLR_output_list[[i]]$bestmod_FPR_range[2]
      phybLR_run_metadata[i, "bestmod_TNR_mean"] <- bLR_output_list[[i]]$bestmod_TNR_mean
      phybLR_run_metadata[i, "bestmod_TP_mean"] <- bLR_output_list[[i]]$bestmod_TP
      phybLR_run_metadata[i, "bestmod_FP_mean"] <- bLR_output_list[[i]]$bestmod_FP
      phybLR_run_metadata[i, "bestmod_TN_mean"] <- bLR_output_list[[i]]$bestmod_TN
      phybLR_run_metadata[i, "bestmod_FN_mean"] <- bLR_output_list[[i]]$bestmod_FN
      phybLR_run_metadata[i, "bestmod_nSuccesses_mean"] <- bLR_output_list[[i]]$bestmod_nSuccesses 
      phybLR_run_metadata[i, "bestmod_nFailures_mean"] <- bLR_output_list[[i]]$bestmod_nFailures
      phybLR_run_metadata[i, "bestmod_Accuracy_mean"] <- bLR_output_list[[i]]$bestmod_Accuracy
      phybLR_run_metadata[i, "comput_time (min)"] <- comp_time_per_dataset
      phybLR_run_metadata[i, "skipped?"] <- "No"
      phybLR_run_metadata[i, "reason_for_skipping"] <- NA
      } else if( is.list(bLR_output_list[[i]])==F | length(bLR_output_list[[i]]<=2) ){ #i.e., if test unfeasible...
        phybLR_run_metadata[i, c(9:45)] <- NA
        phybLR_run_metadata[i, "skipped?"] <- "Yes"
        phybLR_run_metadata[i, "reason_for_skipping"] <- bLR_output_list[[i]]
        } # closes 'else if(is.list(bLR_output_list[[i]])==F)' clause
} # closes 'for(i in 1:length(bLR_output_list))' loop

# Save phybLR_run_metadata to a csv file within the phybLR_models subdirectory:
write.csv(phybLR_run_metadata, paste0(phybLR_directory_path, "/phybLR_metadata - ", ds_name_generic, " - ", treename_j, " - ", groupvar_k, " - ", predvar_L, ".csv"), row.names=FALSE)
cat("\n","PROGRESS REP: chunk [S4.3.01] complete; script_S4 has finished running!","\n")
#*-----**-----*

# We have just saved a table containing the phybLR() results from one run of this job array, which includes three rows--one run on raw data, one on log10-transformed data, and one run on BoxCox-transformed data--for the same input tree, input dataset, and variable pair. Running this script in massively parallel fashion with Job Arrays will generate many such output CSV files, which we will stitch together in script_S5. If you have any questions about this script or its associated data, please feel free to contact me anytime via the address below.

# -- Caleb Gordon
#------------------------------------------------------------------------------
```
