## Supplementary R Scripts for "Limb proportions predict aquatic habits and soft-tissue flippers in extinct amniotes": script_S5.R

```
#** Fifth R script (script_S5.R) for Gordon et al., "Limb proportions predict aquatic habits and soft-tissue flippers in extinct amniotes."**

# NOTE ON READABILITY: This script contains many long, multi-line comments. To increase readability within Rstudio, go to Tools > Global Options, select the 'Code' tab, and check 'Soft-wrap R source files'. This will wrap all lines of code to match your personal GUI's margins, so that no lines of code run off-screen.

# NOTE ON SCRIPT ORDER: This script is downstream of all other scripts associated with the paper titled above. If you have not yet finished running scripts S1, S2, and all six S2-S4 scripts, please close this file and run those eight previous scripts to completion. Once those scripts are complete, you can reopen the current file and proceed. 

# NOTE ON REQUIRED STORAGE: For the batch job associated with this script, you will need to request at least 150 gigabytes of storage, as the combined environments from all upstream scripts will be massive (~10 GiB for script_S2 outputs and ~130 GiB for script_S4 outputs). Requesting much less storage then this will cause your R session to crash.

# WHAT WE'VE DONE SO FAR: In script S2, we performed various phylogenetic tests comparing group means and variances for different morphometric variables across phenotypic groups. In script S3, we performed phylogenetic tests checking for pairwise correlations between these variables. In script S4, we fit phylogenetic binomial logistic regression (phybLR) models to the data which used each morphometric variable separately to predict binarified categorical variables. 

# WHAT WE WILL DO HERE--four things: First, we will load the script_S1 environment, and import & stitch all data files generated from upstream script. Second, we will use ROC analysis to determine the best predictor variable for a given tree-response_var combination among each variable series (forelimb proportions, hindlimb proportions, manus planform proportions, pes planform proportions, manual ungual dimensions, pedal ungual dimensions, manual proximal phalanx dimensions, and pedal proximal phalanx dimensions). Third, we will use our previously identified best-performing phybLR models to predict phenotypes for all the extinct tip taxa in our dataset, and use ancestral state reconstructions to reconstruct the evolutionary histories of those phenotypes along the amniote tree. Fourth, we will perform a thorough sensitivity analysis that assesses the impact of tree topology and data-transformation method on the results of all previously performed statistical tests. 

# This script is computationally intensive, but should run within a few days by using moderate parallelization (in the form of foreach-loops) that increases computation rate. Please note that this script may produce thousands of output files (totaling potentially several Gigabytes) to your working directory.

# This is the outline of sections for script_S5:
#**[S5] PERFORM SENSITIVITY TESTS & PREDICT ANCIENT PHENOTYPES**
# --- [S5.1]: PREPARE CODING ENVIRONMENT & STITCH UPSTREAM OUTPUTS.
# --- [S5.2]: RUN SENSITIVITY ANALYSES FOR TRANSFORM METHOD & TREE TOPOLOGY.
# --- [S5.3]: MAKE CORRELATION MATRICES USING PHYCORR RESULTS.
# --- [S5.4]: USE ROC ANALYSIS TO IDENTIFY BEST PHYBLR MODEL(S).
# --- [S5.5]: PREDICT TIP & NODE PROBABILITIES USING BEST PHYBLR MODEL(S).
# --- [S5.6]: SAVE R ENVIRONMENT & COMPUTATION TIME FOR FOR FUTURE REFERENCE.

# We go through each of these sections with a series of makeshift code "chunks" below, starting with [S5.1].
#------------------------------------------------------------------------------

#**"-------------------------------------------------------------------------"*
#**"-------------- ### [S5.1] PREPARE CODING ENVIRONMENT ### ----------------"*
#**"-------------------------------------------------------------------------"*

#*-----**{ [S5.1.01] Load required packages. }*
# NOTE: Since all packages were installed in script_S1, we do not repeat the optional installation lines for these packages here; however, lines of code to install the following packages are provided in script_S1, in section [S1.1.01].

# Parallel programming on multiple CPU cores:
library(parallel)
library(foreach)
library(doMC)
library(doParallel)

# Data object manipulation:
library(forcats) # for releveling factors with fct_relevel() function
library(data.table) # for converting objects to data tables and matrices
library(plyr) # useful for in-task parallelization
library(dplyr) # useful for in-task parallelization
library(stringr) # contains str_detect() function

# Data visualization and exporting:
library(ggplot2) # for effective data visualization with ggplot suite
library(MESS) # for specifying translucent colors as function arguments
library(car) # for normal quantile plot construction
library(RColorBrewer) # for defining fancy color palettes
library(gridExtra) # useful for generating multi-panel plots and tables
library(ggimage) # for exporting ggplots
library(corrplot) # for correlogram construction

# Data analysis:
library(devtools) # required for downloading some other packages from github
library(vcd) # for Cramer's V and other association stats
library(MASS) # used to make Box-Cox transformations with boxcox() function
library(pROC) # contains functions to plot and extract data from ROC curves
library(pracma) # contains diag() and trapz() functions; diag() is useful for making phylogenetic variance-covariance matrices; trapz() is useful for calculating ROC curve AUC.

# Sensitivity analysis:
library(tidyverse)
library(purrr)

# Phylogenetic comparative methods and tree visualization:
library(ape) # for basic tree manipulation and phylogenetic comparative methods
library(ggtree) # for advanced tree visualization using ggplot2 suite
library(strap) # contains DatePhylo() function to time-calibrate trees
library(phytools) # contains phylANOVA() function
library(geiger) # contains aov.phylo() function
library(phylolm) # contains phyloglm() and phylolm() functions
library(castor) # contains asr_independent_contrasts() function
library(vegan) # contains adonis2(), which can run phyloPERMANOVAs
cat("\n","PROGRESS REP: chunk [S5.1.01] complete; starting next chunk..","\n")
#*-----**-----*

#*-----**{ [S5.1.02] Load upstream upstream S1 & S2 script environments. }*
# Define working directory path:
wd_path <- "/gpfs/gibbs/project/bhullar/cmg89/Flipper_Project/"
setwd(wd_path); getwd()
# Load upstream script environments from scripts S1 & S2:
load(file=paste0(wd_path, "/R_ENVIR_for_script_S1.RData")) # S1 output
load(file=paste0(wd_path, "/R_ENVIR_for_script_S2.RData")) # S2 output
cat("\n","PROGRESS REP: chunk [S5.1.02] complete; starting next chunk..","\n")
#*-----**-----*

#*-----**{ [S5.1.03] Set up parallel programming environment. }*
# We will use the foreach package to run programs in parallel on multiple CPU cores. The registerDoMC() function below assigns the proper number of cores for this task. Before you begin running the code below, set the CPU_num available to you on your personal computer or HPC cluster.

CPU_num <- 8 # On Yale's High-Performance Computing cluster, 5 is the total number of processing cores that can be requested for interactive work, whereas 48 is the highest number that can be requested for batch jobs.

# Set up parallel backend:
cl <- makeCluster(CPU_num, outfile = "") # outfile="" ensures that output prints to the console
registerDoParallel(cl)

# Set iteration number for all simulation-based comparative phylogenetic tests:
iter_num=10000 # This will determine the number of iterations through which for-loops will run for performing simulation-based phylogenetic statistical tests. Increasing the number of iterations will drastically increase the time and computational resources required to run the script below. For troubleshooting this code on a local computer, setting iter_num=5 works well. While running on Yale's HPC and for the paper, we set iter_num=10000. 
cat("\n","PROGRESS REP: chunk [S5.1.03] complete; starting next chunk..","\n")
#*-----**-----*

#*-----**{ [S5.1.04] Load & stitch script_S2 outputs for downstream work. }*
# Recall script_S2 output path:
phyloBoxPlot_output_path <- paste0(output_path,"/phyloBoxPlots/") # navigates to the correct directory for phyloBoxPlot outputs

# Initialize script_S2 data objects to store all output metadata:
ALL_phyloBoxPlot_objects <- as.list(rep(NA, length(orig_phyloBoxPlot_object_list)*8*3)) # length = (number of unique dataset-variable triads) * (number of tree variants) * (number of transform methods)

ALL_phyloBoxPlot_metadata <- data.frame(matrix(ncol=18, nrow=length(ALL_phyloBoxPlot_objects))) 

phyBP_iter <- data.frame(matrix(ncol=18,nrow=1)) # An instance of this smaller data frame will store phyloBoxPlot metadata for each list item.

colnames(ALL_phyloBoxPlot_metadata) <- colnames(phyBP_iter) <- c("function_name", "bootstrap_number", "input_dataset", "input_tree", "response_variable", "grouping_variable", "full_sample_size", "sample_size_by_group_singlestring", "phylANOVA_omnibus_Fstat", "phylANOVA_omnibus_pval", "phylANOVA_posthoc_correction_method", "phylANOVA_posthoc_pvals_singlestring", "phyloLev_omnibus_Fstat", "phyloLev_omnibus_pval", "phyloLev_posthoc_correction_method", "phyloLev_posthoc_pvals_singlestring", "skipped?", "reason_for_skipping")

# Populate first third of ALL_phyloBoxPlot_metadata object with raw outputs:
for(i in 1:length(orig_phyloBoxPlot_object_list)){
  for(j in 1:length(tree_variants)){
    FIRST_ITERATION <- ((i-1)*length(tree_variants) + j) # sets iter number
    phyBP_iter_new <- phyBP_iter # initializes phyBP_iter instance
    if(is.list(orig_phyloBoxPlot_object_list[[i]])==T){
      phyBP_iter_new$function_name <-  orig_phyloBoxPlot_object_list[[i]][[j]]$function_name
      phyBP_iter_new$bootstrap_number <- orig_phyloBoxPlot_object_list[[i]][[j]]$bootstrap_number
      phyBP_iter_new$input_dataset <- orig_phyloBoxPlot_object_list[[i]][[j]]$input_dataset
      phyBP_iter_new$input_tree <- orig_phyloBoxPlot_object_list[[i]][[j]]$input_tree
      phyBP_iter_new$response_variable <- orig_phyloBoxPlot_object_list[[i]][[j]]$response_variable
      phyBP_iter_new$grouping_variable <- orig_phyloBoxPlot_object_list[[i]][[j]]$grouping_variable
      phyBP_iter_new$full_sample_size <- orig_phyloBoxPlot_object_list[[i]][[j]]$full_sample_size
      phyBP_iter_new$sample_size_by_group_singlestring <-  orig_phyloBoxPlot_object_list[[i]][[j]]$sample_size_by_group_singlestring
      phyBP_iter_new$phylANOVA_omnibus_Fstat <- orig_phyloBoxPlot_object_list[[i]][[j]]$phylANOVA_omnibus_Fstat
      phyBP_iter_new$phylANOVA_omnibus_pval <- orig_phyloBoxPlot_object_list[[i]][[j]]$phylANOVA_omnibus_pval
      phyBP_iter_new$phylANOVA_posthoc_correction_method <- orig_phyloBoxPlot_object_list[[i]][[j]]$phylANOVA_posthoc_correction_method
      
      phylANOVA_pvals_vec <- as.vector(orig_phyloBoxPlot_object_list[[i]][[j]]$phylANOVA_posthoc_pvals)
      phylANOVA_pvals_string <- paste(phylANOVA_pvals_vec,collapse=", ")
      phyBP_iter_new$phylANOVA_posthoc_pvals_singlestring <- phylANOVA_pvals_string
      
      phyBP_iter_new$phyloLev_omnibus_Fstat <- orig_phyloBoxPlot_object_list[[i]][[j]]$phyloLev_omnibus_Fstat
      phyBP_iter_new$phyloLev_omnibus_pval <- orig_phyloBoxPlot_object_list[[i]][[j]]$phyloLev_omnibus_pval
      phyBP_iter_new$phyloLev_posthoc_correction_method <- orig_phyloBoxPlot_object_list[[i]][[j]]$phyloLev_posthoc_correction_method
        
      phyloLev_pvals_vec <- as.vector(orig_phyloBoxPlot_object_list[[i]][[j]]$phyloLev_posthoc_pvals)
      phyloLev_pvals_string <- paste(phyloLev_pvals_vec,collapse=", ")
      phyBP_iter_new$phyloLev_posthoc_pvals_singlestring <- phyloLev_pvals_string
      
      phyBP_iter_new$'skipped?' <- "No"
      phyBP_iter_new$reason_for_skipping <- "NA"
    } else if (is.list(orig_phyloBoxPlot_object_list[[i]])==F) {
      phyBP_iter_new$function_name <- "phyloBoxPlot()"
      phyBP_iter_new$bootstrap_number <- "NA"
      phyBP_iter_new$input_dataset <- "Detailed in 'reason-for-skipping'"
      phyBP_iter_new$input_tree <- "Detailed in 'reason-for-skipping'"
      phyBP_iter_new$response_variable <- "Detailed in 'reason-for-skipping'"
      phyBP_iter_new$grouping_variable <- "Detailed in 'reason-for-skipping'"
      phyBP_iter_new$full_sample_size <- "NA"
      phyBP_iter_new$sample_size_by_group_singlestring <- "NA"
      phyBP_iter_new$phylANOVA_omnibus_Fstat <- "NA"
      phyBP_iter_new$phylANOVA_omnibus_pval <- "NA"
      phyBP_iter_new$phylANOVA_posthoc_correction_method <- "NA"
      phyBP_iter_new$phylANOVA_posthoc_pvals_singlestring <- "NA"
      phyBP_iter_new$phyloLev_omnibus_Fstat <- "NA"
      phyBP_iter_new$phyloLev_omnibus_pval <- "NA"
      phyBP_iter_new$phyloLev_posthoc_correction_method <- "NA"
      phyBP_iter_new$phyloLev_posthoc_pvals_singlestring <- "NA"
      phyBP_iter_new$'skipped?' <- "Yes"
      phyBP_iter_new$reason_for_skipping <- orig_phyloBoxPlot_object_list[[i]]
    } # closes 'else if(is.list(orig_phyloBoxPlot_object_list[[i]])==F)' clause
    ALL_phyloBoxPlot_objects[[FIRST_ITERATION]] <- phyBP_iter_new
  } # closes 'for(j in 1:length(tree_variants))' loop
} # closes 'for(i in 1:length(orig_phyloBoxPlot_object_list))' loop

# Populate second third of ALL_phyloBoxPlot_metadata object with log10 outputs:
for(i in 1:length(log10_phyloBoxPlot_object_list)){
  for(j in 1:length(tree_variants)){
    SECOND_ITERATION <- FIRST_ITERATION + ((i-1)*length(tree_variants) + j) 
    phyBP_iter_new <- phyBP_iter # initializes phyBP_iter instance
    if(is.list(log10_phyloBoxPlot_object_list[[i]])==T){
      phyBP_iter_new$function_name <-  log10_phyloBoxPlot_object_list[[i]][[j]]$function_name
      phyBP_iter_new$bootstrap_number <- log10_phyloBoxPlot_object_list[[i]][[j]]$bootstrap_number
      phyBP_iter_new$input_dataset <- log10_phyloBoxPlot_object_list[[i]][[j]]$input_dataset
      phyBP_iter_new$input_tree <- log10_phyloBoxPlot_object_list[[i]][[j]]$input_tree
      phyBP_iter_new$response_variable <- log10_phyloBoxPlot_object_list[[i]][[j]]$response_variable
      phyBP_iter_new$grouping_variable <- log10_phyloBoxPlot_object_list[[i]][[j]]$grouping_variable
      phyBP_iter_new$full_sample_size <- log10_phyloBoxPlot_object_list[[i]][[j]]$full_sample_size
      phyBP_iter_new$sample_size_by_group_singlestring <-  log10_phyloBoxPlot_object_list[[i]][[j]]$sample_size_by_group_singlestring
      phyBP_iter_new$phylANOVA_omnibus_Fstat <- log10_phyloBoxPlot_object_list[[i]][[j]]$phylANOVA_omnibus_Fstat
      phyBP_iter_new$phylANOVA_omnibus_pval <- log10_phyloBoxPlot_object_list[[i]][[j]]$phylANOVA_omnibus_pval
      phyBP_iter_new$phylANOVA_posthoc_correction_method <- log10_phyloBoxPlot_object_list[[i]][[j]]$phylANOVA_posthoc_correction_method
      
      phylANOVA_pvals_vec <- as.vector(log10_phyloBoxPlot_object_list[[i]][[j]]$phylANOVA_posthoc_pvals)
      phylANOVA_pvals_string <- paste(phylANOVA_pvals_vec,collapse=", ")
      phyBP_iter_new$phylANOVA_posthoc_pvals_singlestring <- phylANOVA_pvals_string
      
      phyBP_iter_new$phyloLev_omnibus_Fstat <- log10_phyloBoxPlot_object_list[[i]][[j]]$phyloLev_omnibus_Fstat
      phyBP_iter_new$phyloLev_omnibus_pval <- log10_phyloBoxPlot_object_list[[i]][[j]]$phyloLev_omnibus_pval
      phyBP_iter_new$phyloLev_posthoc_correction_method <- log10_phyloBoxPlot_object_list[[i]][[j]]$phyloLev_posthoc_correction_method
      
      phyloLev_pvals_vec <- as.vector(log10_phyloBoxPlot_object_list[[i]][[j]]$phyloLev_posthoc_pvals)
      phyloLev_pvals_string <- paste(phyloLev_pvals_vec,collapse=", ")
      phyBP_iter_new$phyloLev_posthoc_pvals_singlestring <- phyloLev_pvals_string
      
      phyBP_iter_new$'skipped?' <- "No"
      phyBP_iter_new$reason_for_skipping <- "NA"
    } else if (is.list(orig_phyloBoxPlot_object_list[[i]])==F) {
      phyBP_iter_new$function_name <- "phyloBoxPlot()"
      phyBP_iter_new$bootstrap_number <- "NA"
      phyBP_iter_new$input_dataset <- "Detailed in 'reason-for-skipping'"
      phyBP_iter_new$input_tree <- "Detailed in 'reason-for-skipping'"
      phyBP_iter_new$response_variable <- "Detailed in 'reason-for-skipping'"
      phyBP_iter_new$grouping_variable <- "Detailed in 'reason-for-skipping'"
      phyBP_iter_new$full_sample_size <- "NA"
      phyBP_iter_new$sample_size_by_group_singlestring <- "NA"
      phyBP_iter_new$phylANOVA_omnibus_Fstat <- "NA"
      phyBP_iter_new$phylANOVA_omnibus_pval <- "NA"
      phyBP_iter_new$phylANOVA_posthoc_correction_method <- "NA"
      phyBP_iter_new$phylANOVA_posthoc_pvals_singlestring <- "NA"
      phyBP_iter_new$phyloLev_omnibus_Fstat <- "NA"
      phyBP_iter_new$phyloLev_omnibus_pval <- "NA"
      phyBP_iter_new$phyloLev_posthoc_correction_method <- "NA"
      phyBP_iter_new$phyloLev_posthoc_pvals_singlestring <- "NA"
      phyBP_iter_new$'skipped?' <- "Yes"
      phyBP_iter_new$reason_for_skipping <- log10_phyloBoxPlot_object_list[[i]]
    } # closes 'else if(is.list(log10_phyloBoxPlot_object_list[[i]])==F)' 
    ALL_phyloBoxPlot_objects[[SECOND_ITERATION]] <- phyBP_iter_new
  } # closes 'for(j in 1:length(tree_variants))' loop
} # closes 'for(i in 1:length(log10_phyloBoxPlot_object_list))' loop

# Populate third third of ALL_phyloBoxPlot_metadata object with boxcox outputs:
for(i in 1:length(boxcox_phyloBoxPlot_object_list)){
  for(j in 1:length(tree_variants)){
    THIRD_ITERATION <- SECOND_ITERATION + ((i-1)*length(tree_variants) + j) 
    phyBP_iter_new <- phyBP_iter # initializes phyBP_iter instance
    if(is.list(boxcox_phyloBoxPlot_object_list[[i]])==T){
      phyBP_iter_new$function_name <-  boxcox_phyloBoxPlot_object_list[[i]][[j]]$function_name
      phyBP_iter_new$bootstrap_number <- boxcox_phyloBoxPlot_object_list[[i]][[j]]$bootstrap_number
      phyBP_iter_new$input_dataset <- boxcox_phyloBoxPlot_object_list[[i]][[j]]$input_dataset
      phyBP_iter_new$input_tree <- boxcox_phyloBoxPlot_object_list[[i]][[j]]$input_tree
      phyBP_iter_new$response_variable <- boxcox_phyloBoxPlot_object_list[[i]][[j]]$response_variable
      phyBP_iter_new$grouping_variable <- boxcox_phyloBoxPlot_object_list[[i]][[j]]$grouping_variable
      phyBP_iter_new$full_sample_size <- boxcox_phyloBoxPlot_object_list[[i]][[j]]$full_sample_size
      phyBP_iter_new$sample_size_by_group_singlestring <-  boxcox_phyloBoxPlot_object_list[[i]][[j]]$sample_size_by_group_singlestring
      phyBP_iter_new$phylANOVA_omnibus_Fstat <- boxcox_phyloBoxPlot_object_list[[i]][[j]]$phylANOVA_omnibus_Fstat
      phyBP_iter_new$phylANOVA_omnibus_pval <- boxcox_phyloBoxPlot_object_list[[i]][[j]]$phylANOVA_omnibus_pval
      phyBP_iter_new$phylANOVA_posthoc_correction_method <- boxcox_phyloBoxPlot_object_list[[i]][[j]]$phylANOVA_posthoc_correction_method

      phylANOVA_pvals_vec <- as.vector(boxcox_phyloBoxPlot_object_list[[i]][[j]]$phylANOVA_posthoc_pvals)
      phylANOVA_pvals_string <- paste(phylANOVA_pvals_vec,collapse=", ")
      phyBP_iter_new$phylANOVA_posthoc_pvals_singlestring <- phylANOVA_pvals_string
      
      phyBP_iter_new$phyloLev_omnibus_Fstat <- boxcox_phyloBoxPlot_object_list[[i]][[j]]$phyloLev_omnibus_Fstat
      phyBP_iter_new$phyloLev_omnibus_pval <- boxcox_phyloBoxPlot_object_list[[i]][[j]]$phyloLev_omnibus_pval
      phyBP_iter_new$phyloLev_posthoc_correction_method <- boxcox_phyloBoxPlot_object_list[[i]][[j]]$phyloLev_posthoc_correction_method
      
      phyloLev_pvals_vec <- as.vector(boxcox_phyloBoxPlot_object_list[[i]][[j]]$phyloLev_posthoc_pvals)
      phyloLev_pvals_string <- paste(phyloLev_pvals_vec,collapse=", ")
      phyBP_iter_new$phyloLev_posthoc_pvals_singlestring <- phyloLev_pvals_string
      
      phyBP_iter_new$'skipped?' <- "No"
      phyBP_iter_new$reason_for_skipping <- "NA"
    } else if (is.list(orig_phyloBoxPlot_object_list[[i]])==F) {
      phyBP_iter_new$function_name <- "phyloBoxPlot()"
      phyBP_iter_new$bootstrap_number <- "NA"
      phyBP_iter_new$input_dataset <- "Detailed in 'reason-for-skipping'"
      phyBP_iter_new$input_tree <- "Detailed in 'reason-for-skipping'"
      phyBP_iter_new$response_variable <- "Detailed in 'reason-for-skipping'"
      phyBP_iter_new$grouping_variable <- "Detailed in 'reason-for-skipping'"
      phyBP_iter_new$full_sample_size <- "NA"
      phyBP_iter_new$sample_size_by_group_singlestring <- "NA"
      phyBP_iter_new$phylANOVA_omnibus_Fstat <- "NA"
      phyBP_iter_new$phylANOVA_omnibus_pval <- "NA"
      phyBP_iter_new$phylANOVA_posthoc_correction_method <- "NA"
      phyBP_iter_new$phylANOVA_posthoc_pvals_singlestring <- "NA"
      phyBP_iter_new$phyloLev_omnibus_Fstat <- "NA"
      phyBP_iter_new$phyloLev_omnibus_pval <- "NA"
      phyBP_iter_new$phyloLev_posthoc_correction_method <- "NA"
      phyBP_iter_new$phyloLev_posthoc_pvals_singlestring <- "NA"
      phyBP_iter_new$'skipped?' <- "Yes"
      phyBP_iter_new$reason_for_skipping <- boxcox_phyloBoxPlot_object_list[[i]]
    } # closes 'else if(is.list(boxcox_phyloBoxPlot_object_list[[i]])==F)' 
    ALL_phyloBoxPlot_objects[[THIRD_ITERATION]] <- phyBP_iter_new
  } # closes 'for(j in 1:length(tree_variants))' loop
} # closes 'for(i in 1:length(boxcox_phyloBoxPlot_object_list))' loop

# Use ALL_phyloBoxPlot_objects to populate ALL_phyloBoxPlot_metadata table:
for(r in 1:nrow(ALL_phyloBoxPlot_metadata)){
  ALL_phyloBoxPlot_metadata[r, ] <- ALL_phyloBoxPlot_objects[[r]]
} # closes 'for(r in 1:nrow(ALL_phyloBoxPlot_metadata))' loop

# Save ALL_phyloBoxPlot_metadata to CSV file:
write.csv(ALL_phyloBoxPlot_metadata, paste0(phyloBoxPlot_output_path, "*ALL_phyloBoxPlot_metadata.csv"), row.names=FALSE) 
cat("\n","PROGRESS REP: chunk [S5.1.04] complete; starting next chunk..","\n")
#*-----**-----*

#*-----**{ [S5.1.05] Load & stitch script_S3 outputs for downstream work. }*
# Load & stitch script_S3 metadata files:
cortest_output_path <- paste0(output_path,"/phyloCorrPlots/") # navigates to the correct directory for phycorr outputs
phycorr_all_outputs <- list.files(cortest_output_path) # lists all files there
phycorr_key_files <- grep(pattern="phycorr_metadata", phycorr_all_outputs, value=TRUE) # picks out key files representing metadata CSVs to stitch
phycorr_obj_list <- as.list(rep(NA, length=length(phycorr_key_files))) # makes list to be filled with content from all metadata CSVs to be stitched

for(i in 1:length(phycorr_key_files)){
  phycorr_obj_list[[i]] <- read.csv(paste0(cortest_output_path, phycorr_key_files[i]))
} # compiles all CSV outputs into a single list

firstfile <- read.csv(paste0(cortest_output_path,phycorr_key_files[1])) # reads in the first file as a way to initialize the metadata CSV

phycorr_metadata_table <- firstfile # initializes stitched CSV file
for(i in 2:length(phycorr_obj_list)){
  phycorr_metadata_table <- rbind(phycorr_metadata_table,phycorr_obj_list[[i]])
} # cycles through list to create a single stitched CSV file with all metadata

for(i in 1:length(phycorr_key_files)){
  file.remove(paste0(cortest_output_path, phycorr_key_files[i]))
} # This loop deletes all the previous constituent files that we just merged, since we no longer need them and they clutter the output directory.

write.csv(phycorr_metadata_table, paste0(cortest_output_path, "*ALL_phycorr_metadata.csv"), row.names=FALSE) # This line replaces those deleted files with a new stitched file that we created by merging them all.
cat("\n","PROGRESS REP: chunk [S5.1.05] complete; starting next chunk..","\n")
#*-----**-----*

#*-----**{ [S5.1.06] Load & stitch script_S4 outputs for downstream work. }*
phybLR_output_path <- paste0(output_path,"/phybLR_models/") # navigates to the correct directory for phybLR outputs
phybLR_output_path <- "~/project/Flipper_Project/phybLR_models/" ### TO DELETE ###
phybLR_all_outputs <- list.files(phybLR_output_path) # lists all files there
phybLR_CSVs<-grep(pattern="phybLR_metadata",phybLR_all_outputs,value=TRUE) # picks out key files representing metadata CSVs to target for stitching

phybLR_obj_list <- as.list(rep(NA, length=length(phybLR_CSVs))) # makes list to be filled with content from all metadata CSVs to be stitched

for(i in 1:length(phybLR_CSVs)){
  phybLR_obj_list[[i]] <- read.csv(paste0(phybLR_output_path, phybLR_CSVs[i]))
} # compiles all CSV outputs into a single list

firstfile <- read.csv(paste0(phybLR_output_path, phybLR_CSVs[1])) # reads in the first file as a way to initialize the metadata CSV

phybLR_metadata_table <- firstfile # initializes stitched CSV file
for(i in 2:length(phybLR_obj_list)){
  phybLR_metadata_table <- rbind(phybLR_metadata_table, phybLR_obj_list[[i]])
} # cycles through list to create a single stitched CSV file with all metadata

# OPTIONAL DECLUTTERING STEP:
#for(i in 1:length(phybLR_CSVs)){
#  file.remove(paste0(phybLR_output_path, phybLR_CSVs[i]))
#} # This loop deletes all the previous constituent files that we just merged, since we no longer need them and they clutter the output directory.

write.csv(phybLR_metadata_table, paste0(phybLR_output_path, "*ALL_phybLR_metadata.csv"), row.names=FALSE) # This line replaces those (optionally) deleted files with a new stitched file that we created by merging them all.

# Load all R environment files for downstream analysis:
phybLR_env_files <- grep(pattern=".RData",phybLR_all_outputs,value=TRUE) # picks out key files representing R objects saved in script_S4
bLR_mod_list <- as.list(rep(NA, length(phybLR_env_files)))
bLR_mod_names <- rep(NA, length(phybLR_env_files))

load(file=paste0(phybLR_output_path, phybLR_env_files[1]))
test_import <- bLR_output_list
str(test_import)

# WARNING: The next for-loop will take a long time and add ~130-170 GiB to the current R environment.
for(i in 1:length(phybLR_env_files)){
  load(file=paste0(phybLR_output_path, phybLR_env_files[i]))
  bLR_mod_list[[i]]<-bLR_output_list
  rm(bLR_output_list)
  bLR_mod_names[i]<-phybLR_env_files[i]
} # This for-loop should import each of the 9216 bLR_output_list objects generated by the script_S4 job array and save it as a separate element of the bLR_mod_list.

names(bLR_mod_list) <- bLR_mod_names
# Remove all but the raw-data outputs from this list:
bLR_mod_list_copy <- bLR_mod_list
for(i in 1:length(bLR_mod_list_copy)){
  bLR_mod_list[[i]] <- bLR_mod_list_copy[[i]]$phybLR_raw_output
} # This saves just the phybLR output lists for the raw data to the bLR_mod obj.
cat("\n","PROGRESS REP: chunk [S5.1.06] complete; starting next chunk..","\n")
#*-----**-----*  

# We have now set up the coding environment for script_S5, loading required packages and stitching together previous outputs from upstream scripts and job arrays. In the next few sections of the script, we will use these combined outputs to do four things: perform a thorough sensitivity analysis that assesses the impact of transformation method and tree topology on our statistical test results, generate correlation matrices to display pairwise correlations between variables, use ROC analysis to select the best phybLR models, and use those best phybLR models to predict tip and node phenotypes for all contentious taxa in our trees.
#-----------------------------------------------------------------------------

#**"-------------------------------------------------------------------------"*
#**" ### [S5.2] DO SENSITIVITY ANALYSES FOR TRANS. METHOD & TREE SHAPE ### -"*
#**"-------------------------------------------------------------------------"*

# Recall that we ran through our entire data analysis pipeline with eight different supertrees, each assuming a different combination of disputed phylogenetic hypotheses about squamates, enaliosaurs, and stem reptiles. We also ran through our entire data analysis pipeline with three different sets of morphometric data--one raw (un-transformed), one log10-transformed prior to analysis, and one BoxCox-transformed prior to analysis. We saved the results from all of our statistical tests to our working directory, in the form of various "metadata" tables.

# In what follows, we use these tables of test metadata to assess the extent to which these differing tree topologies and data-transformation methods altered our test results. To do this, we use stacked barplots and binomial tests. To assess the impact of tree topology, we visualize with a stacked barplot the percentage of test verdicts (significant vs. not significant) for each test that were the same across all different supertrees and the percentage that were different. We do the same thing to assess the impact of data-transformation method, using one stacked barplot per statistical test to compare the percentage of test verdicts that were the same for raw, log10-trans, and BoxCox-trans datasets, and the percentage of test verdicts that differed among them. We then use binomial exact tests to check whether the percentage of test results that differed was significantly greater than 0.05%.

# The code for the sensitivity analysis in section [S5.2] below was written by Lisa Freisem. 

# NOTE: The code below was written and run on a separate local computer, and the directory paths therefore differ from the others assumed throughout the script. Directory paths may thus need to be changed before successfully running the code below.

#*-----**{ [S5.2.01] Prepare coding environment for sensitivity analysis.}*
# Define working directory path: 
wd_path <- "C:/Users/lisaj/Desktop/Flipper sensitivity script 10_26"
setwd(wd_path); getwd()
input_path  <- "C:/Users/lisaj/Desktop/Flipper sensitivity script 10_26/input"
output_path <- "C:/Users/lisaj/Desktop/Flipper sensitivity script 10_26/output"

# Load csv files:
phybLR_metadata <- read.csv(file = paste0(input_path, "/_ALL_phybLR_metadata.csv"))
phycorr_metadata <- read.csv(file = paste0(input_path, "/_ALL_phycorr_metadata.csv"))
phyloBoxPlot_metadata <- read.csv(paste0(input_path, "/_ALL_phyloBoxPlot_metadata.csv"))

# Create phyloBoxPlot dataset where each phyloLev and phylANOVA posthoc value has its own row, store original metadata as separate variable. Store original format as phyloBoxPlot_metadata_original
phyloBoxPlot_metadata_original <- phyloBoxPlot_metadata

phyloBoxPlot_metadata <- phyloBoxPlot_metadata %>%
  # Separate the strings in both columns into individual rows
  mutate(
    phyloLev_posthoc_pvals = strsplit(as.character(phyloLev_posthoc_pvals_singlestring), ","),
    phylANOVA_posthoc_pvals = strsplit(as.character(phylANOVA_posthoc_pvals_singlestring), ",")
  ) %>%
  unnest(c(phyloLev_posthoc_pvals, phylANOVA_posthoc_pvals)) %>%
  # Delete spaces from both columns to avoid false significance tests 
  mutate(
    phyloLev_posthoc_pvals = gsub(" ", "", phyloLev_posthoc_pvals),
    phylANOVA_posthoc_pvals = gsub(" ", "", phylANOVA_posthoc_pvals)
  ) %>%
  # Drop the original single string columns because they are no longer needed
  select(-phyloLev_posthoc_pvals_singlestring, -phylANOVA_posthoc_pvals_singlestring)
cat("\n","PROGRESS REP.: chunk [S5.2.01] done; starting next chunk...","\n")
#*-----**-----*

#*-----**{ [S5.2.02] Define binomial significance marker for each p-value.}*
# Add new columns to the datasets for each p-value column, labeling the p-values for each test either "significant" or "insignificant" for easier handling later on.

# Define a function to categorize p-values
categorize_significance <- function(pval) {
  ifelse(pval < 0.05, 'significant', 'insignificant')
}

# List of columns for phybLR_metadata
phybLR_pvals <- c("AUC_binomt_pval", "McF_pval_mean", "McF_pval_for_full_ds", "bestmod_binomt_pval")

# Apply the function and create new columns in phybLR_metadata
phybLR_metadata[paste0(phybLR_pvals, "_sign")] <- lapply(phybLR_metadata[phybLR_pvals], categorize_significance)

# List of columns for phycorr_metadata
phycorr_pvals <- c("intercept.p.val", "slope.p.val")

# Apply the function and create new columns in phycorr_metadata
phycorr_metadata[paste0(phycorr_pvals, "_sign")] <- lapply(phycorr_metadata[phycorr_pvals], categorize_significance)

# List of columns for phyloBoxPlot_metadata
phyloBoxPlot_pvals <- c("phylANOVA_omnibus_pval", "phyloLev_omnibus_pval", "phyloLev_posthoc_pvals", "phylANOVA_posthoc_pvals")

# Apply the function and create new columns in phyloBoxPlot_metadata
phyloBoxPlot_metadata[paste0(phyloBoxPlot_pvals, "_sign")] <- lapply(phyloBoxPlot_metadata[phyloBoxPlot_pvals], categorize_significance)
cat("\n","PROGRESS REP.: chunk [S5.2.02] done; starting next chunk...","\n")
#*-----**-----*

#*-----**{ [S5.2.03] Create data subsets filtered by transformation method.}*
# create subsets of each dataset to include only one form of data treatment (raw data, log10-transformed data, or boxcox transformed data). This drops all rows with no specified treatment

# create data subsets for the transformation types "raw", "log10", and "boxcox" for the dataset "phybLR_metadata". WARNING: This will drop all rows with no specified transformation type.
filter_types_phybLR <- c("raw", "log10", "boxcox")
phybLR_filtered <- lapply(filter_types_phybLR, function(type) {
  dplyr::filter(phybLR_metadata, grepl(type, phybLR_metadata$input_dataset))
})

# Name the filtered datasets according to filter type
names(phybLR_filtered) <- paste0("phybLR_", filter_types_phybLR)

# Check whether the number of rows is equal in all type-filtered datasets
if(nrow(phybLR_filtered$phybLR_raw) != nrow(phybLR_filtered$phybLR_log10) | nrow(phybLR_filtered$phybLR_raw) != nrow(phybLR_filtered$phybLR_boxcox))
{print("Warning: Unequal number of rows in raw, log10-transformed and box-cox transformed phybLR_metadata")} 

# Check whether the number of rows in the original dataset is equal to the sum of the filtered datasets. If it is not, some data entries contained no information on transformation method and will be dropped from the sensitivity analysis
if(nrow(phybLR_metadata) != nrow(phybLR_filtered$phybLR_raw)+nrow(phybLR_filtered$phybLR_log10)+nrow(phybLR_filtered$phybLR_boxcox))
{print("Data entries with no information on transformation type (raw, log10, or boxcox) are being dropped from the phybLR_filtered datasets")}

# create data subsets for the transformation types "raw", "log10", and "boxcox" for the dataset "phycorr_metadata"
# 
filter_types_phycorr <- list(
  raw = quote(grepl('all', phycorr_metadata$input_dataset)),
  log10 = quote(grepl('log10', phycorr_metadata$input_dataset)),
  boxcox = quote(grepl('boxcox', phycorr_metadata$input_dataset))
)

phycorr_filtered <- lapply(names(filter_types_phycorr), function(name) {
  dplyr::filter(phycorr_metadata, eval(filter_types_phycorr[[name]]))
})

# Name the filtered datasets according to filter type
names(phycorr_filtered) <- paste0("phycorr_", names(filter_types_phycorr))

# Check whether the number of rows is equal in all type-filtered datasets
if(nrow(phycorr_filtered$phycorr_raw) != nrow(phycorr_filtered$phycorr_log10) | nrow(phycorr_filtered$phycorr_raw) != nrow(phycorr_filtered$phycorr_boxcox))
{print("Warning: Unequal number of rows in raw, log10-transformed and box-cox transformed phycorr_metadata")} 

# Check whether the number of rows in the original dataset is equal to the sum of the filtered datasets. If it is not, some data entries contained no information on transformation method and will be dropped from the sensitivity analysis
if(nrow(phycorr_metadata) != nrow(phycorr_filtered$phycorr_raw)+nrow(phycorr_filtered$phycorr_log10)+nrow(phycorr_filtered$phycorr_boxcox))
{print("Data entries with no information on transformation type (raw, log10, or boxcox) are being dropped from the phycorr_filtered datasets")}

# create data subsets for the transformation types "raw", "log10", and "boxcox" for the dataset "phybLR_metadata"
filter_types_boxplot <- list(
  raw = quote(grepl('all', phyloBoxPlot_metadata$input_dataset) & !grepl('log10|boxcox', phyloBoxPlot_metadata$input_dataset)),
  log10 = quote(grepl('log10', phyloBoxPlot_metadata$input_dataset)),
  boxcox = quote(grepl('boxcox', phyloBoxPlot_metadata$input_dataset))
)

phyloBoxPlot_filtered <- lapply(names(filter_types_boxplot), function(name) {
  dplyr::filter(phyloBoxPlot_metadata, eval(filter_types_boxplot[[name]]))
})

# Name the filtered datasets according to filter type
names(phyloBoxPlot_filtered) <- paste0("phyloBoxPlot_", names(filter_types_boxplot))

# Check whether the number of rows is equal in all type-filtered datasets
if(nrow(phyloBoxPlot_filtered$phyloBoxPlot_raw) != nrow(phyloBoxPlot_filtered$phyloBoxPlot_log10) | nrow(phyloBoxPlot_filtered$phyloBoxPlot_raw) != nrow(phyloBoxPlot_filtered$phyloBoxPlot_boxcox)) 
{print("Warning: Unequal number of rows in raw, log10-transformed and box-cox transformed phyloBoxPlot_metadata")} 

# Check whether the number of rows in the original dataset is equal to the sum of the filtered datasets. If it is not, some data entries contained no information on transformation method and will be dropped from the sensitivity analysis
if(nrow(phyloBoxPlot_metadata) != nrow(phyloBoxPlot_filtered$phyloBoxPlot_raw)+nrow(phyloBoxPlot_filtered$phyloBoxPlot_log10)+nrow(phyloBoxPlot_filtered$phyloBoxPlot_boxcox))
{print("Data entries with no information on transformation type (raw, log10, or boxcox) are being dropped from the phyloBoxPlot_filtered datasets")}
cat("\n","PROGRESS REP.: chunk [S5.2.03] done; starting next chunk...","\n")
#*-----**-----*

#*-----**{ [S5.2.04] Create plots showing influence of transformation method on p-value significance for phybLR tests. }*
# Create function to calculate identical vs. divergent counts
calculate_counts_trans <- function(col_raw, col_log10, col_boxcox) {
  common_values <- intersect(intersect(col_raw, col_log10),col_boxcox)
  
  identical_count <-  sum(col_raw %in% common_values) + 
    sum(col_log10 %in% common_values) + 
    sum(col_boxcox %in% common_values)
  
  total_count <- length(col_raw) + length(col_log10) + length(col_boxcox)
  divergent_count <- total_count - identical_count
  
  return(c(identical_count, divergent_count))
}

results_list_phybLR_trans <- list()

# Prepare results for each column
for (col in phybLR_pvals){
  counts <- calculate_counts_trans(phybLR_filtered$phybLR_raw[[col]],phybLR_filtered$phybLR_log10[[col]],phybLR_filtered$phybLR_boxcox[[col]])
  results_list_phybLR_trans[[col]] <- data.frame(
    Category = c("Identical","Divergent"),
    Count = counts,
    Column = col
  )
}

# Combine results into a single data frame
results_phybLR_trans <- do.call(rbind, results_list_phybLR_trans)

# Convert counts to percentages
results_phybLR_trans <- results_phybLR_trans %>%
  group_by(Column) %>%
  mutate(Percentage = Count / sum(Count) * 100)

# Create the stacked percentage bar plot
ggplot(results_phybLR_trans, aes(x = Column, y = Percentage, fill = Category)) +
  geom_bar(stat = "identity", position = "fill") +
  scale_fill_manual(values=c('brown2', 'springgreen4')) +
  scale_y_continuous(labels = scales::percent) +
  labs(title = "Ratio of identical vs divergent statistical significance between raw, log10-transformed, and boxcox-transformed data in the phybLR dataset",
       x = "value", y = "consensus") + theme(axis.text.x = element_text(angle = 90, vjust = 0.5, hjust=1),plot.title = element_text(hjust = 0.5))
cat("\n","PROGRESS REP.: chunk [S5.2.04] done; starting next chunk...","\n")
#*-----**-----*

#*—————**{ [S5.2.05] Create plots showing influence of transformation method on p-value significance in phycorr dataset. }*
# Create function to calculate identical vs. divergent counts
calculate_counts_trans <- function(col_raw, col_log10, col_boxcox) {
  common_values <- intersect(intersect(col_raw, col_log10),col_boxcox)
  
  identical_count <-  sum(col_raw %in% common_values) + 
    sum(col_log10 %in% common_values) + 
    sum(col_boxcox %in% common_values)
  
  total_count <- length(col_raw) + length(col_log10) + length(col_boxcox)
  divergent_count <- total_count - identical_count
  return(c(identical_count, divergent_count))
}

results_list_phycorr_trans <- list()

# Prepare results for each column
for (col in phycorr_pvals){
  counts <- calculate_counts_trans(phycorr_filtered$phycorr_raw[[col]],phycorr_filtered$phycorr_log10[[col]],phycorr_filtered$phycorr_boxcox[[col]])
  results_list_phycorr_trans[[col]] <- data.frame(
    Category = c("Identical","Divergent"),
    Count = counts,
    Column = col
  )
}

# Combine results into a single data frame
results_phycorr_trans <- do.call(rbind, results_list_phycorr_trans)

# Convert counts to percentages
results_phycorr_trans <- results_phycorr_trans %>%
  group_by(Column) %>%
  mutate(Percentage = Count / sum(Count) * 100)

# Create the stacked percentage bar plot
ggplot(results_phycorr_trans, aes(x = Column, y = Percentage, fill = Category)) +
  geom_bar(stat = "identity", position = "fill") +
  scale_fill_manual(values=c('brown2', 'springgreen4')) +
  scale_y_continuous(labels = scales::percent) +
  labs(title = "Ratio of identical vs divergent statistical significance between raw, log10-transformed, and boxcox-transformed data in the phycorr dataset",
       x = "value", y = "consensus") + theme(axis.text.x = element_text(angle = 90, vjust = 0.5, hjust=1),plot.title = element_text(hjust = 0.5))
cat("\n","PROGRESS REP.: chunk [S5.2.05] done; starting next chunk...","\n")
#*-----**-----*

#*—————**{ [S5.2.06] Create plots showing influence of transformation method on p-value significance in phyloBoxPlot dataset.  }*
# Create function to calculate identical vs. divergent counts
calculate_counts_trans <- function(col_raw, col_log10, col_boxcox) {
  common_values <- intersect(intersect(col_raw, col_log10),col_boxcox)
  
  identical_count <-  sum(col_raw %in% common_values) + 
    sum(col_log10 %in% common_values) + 
    sum(col_boxcox %in% common_values)
  
  total_count <- length(col_raw) + length(col_log10) + length(col_boxcox)
  divergent_count <- total_count - identical_count
  return(c(identical_count, divergent_count))
}

results_list_phyloBoxPlot_trans <- list()

# Prepare results for each column
for (col in phyloBoxPlot_pvals){
  counts <- calculate_counts_trans(phyloBoxPlot_filtered$phyloBoxPlot_raw[[col]],phyloBoxPlot_filtered$phyloBoxPlot_log10[[col]],phyloBoxPlot_filtered$phyloBoxPlot_boxcox[[col]])
  results_list_phyloBoxPlot_trans[[col]] <- data.frame(
    Category = c("Identical","Divergent"),
    Count = counts,
    Column = col
  )
}

# Combine results into a single data frame
results_phyloBoxPlot_trans <- do.call(rbind, results_list_phyloBoxPlot_trans)

# Convert counts to percentages
results_phyloBoxPlot_trans <- results_phyloBoxPlot_trans %>%
  group_by(Column) %>%
  mutate(Percentage = Count / sum(Count) * 100)

# Create the stacked percentage bar plot
ggplot(results_phyloBoxPlot_trans, aes(x = Column, y = Percentage, fill = Category)) +
  geom_bar(stat = "identity", position = "fill") +
  scale_fill_manual(values=c('brown2', 'springgreen4')) +
  scale_y_continuous(labels = scales::percent) +
  labs(title = "Ratio of identical vs divergent statistical significance between raw, log10-transformed, and boxcox-transformed data in the phyloBoxPlot dataset",
       x = "value", y = "consensus") + theme(axis.text.x = element_text(angle = 90, vjust = 0.5, hjust=1),plot.title = element_text(hjust = 0.5))

# Create the stacked percentage bar plot summarizing all transformation method ratios
results_trans_total <- rbind(results_phybLR_trans,results_phycorr_trans,results_phyloBoxPlot_trans) %>%
  arrange(match(Column, c("phylANOVA_omnibus_pval", "phylANOVA_posthoc_pvals", "phyloLev_omnibus_pval", "phyloLev_posthoc_pvals", "intercept.p.val", "slope.p.val", "AUC_binomt_pval", "bestmod_binomt_pval", "McF_pval_mean", "McF_pval_for_full_ds")), desc(Category))

# Mutate the column "Column" into a factor so that the row order in this column is respected by the barplot
results_trans_total$Column <- factor(results_trans_total$Column, levels = unique(results_trans_total$Column))

# Create the stacked percentage bar plot
ggplot(results_trans_total, aes(x = Column, y = Percentage, fill = Category)) +
  geom_bar(stat = "identity", position = "fill") +
  scale_fill_manual(values=c('brown2', 'springgreen4')) +
  scale_y_continuous(labels = scales::percent) +
  labs(title = "Ratio of identical vs divergent statistical significance between raw, log10-transformed, and boxcox-transformed data",
       x = "value", y = "consensus") + theme(axis.text.x = element_text(angle = 90, vjust = 0.5, hjust=1),plot.title = element_text(hjust = 0.5))

# Save the plot as a PDF
ggsave(path = output_path, file = paste0("significance_trans_barplot_", Sys.Date(), ".pdf"), plot = last_plot(), device = "pdf", width = 8, height = 6)
cat("\n","PROGRESS REP.: chunk [S5.2.06] done; starting next chunk...","\n")
#*-----**-----*

#*—————**{ [S5.2.07] Create data subsets filtered by supertree.  }*
# Create subsets of each dataset to include only one supertree. WARNING: This will drop all rows with no specified supertree.

# Create variable to define number of supertrees
supertree_numbers <- 1:8

# Create data subsets for each supertree in the dataset "phybLR_metadata"
phybLR_supertrees <- lapply(supertree_numbers, function(x) {
  dplyr::filter(phybLR_metadata, grepl(paste0('supertree', x), input_tree))
})

# Name the filtered datasets according to numbered supertree
names(phybLR_supertrees) <- paste0("phybLR_supertree_", supertree_numbers)

# Create data subsets for each supertree in the dataset "phycorr_metadata"
phycorr_supertrees <- lapply(supertree_numbers, function(x) {
  dplyr::filter(phycorr_metadata, grepl(paste0('supertree', x), input_tree))
})

# Name the filtered datasets according to numbered supertree
names(phycorr_supertrees) <- paste0("phycorr_supertree_", supertree_numbers)

# Create data subsets for each supertree in the dataset "phyloBoxPlot_metadata"
phyloBoxPlot_supertrees <- lapply(supertree_numbers, function(x) {
  dplyr::filter(phyloBoxPlot_metadata, grepl(paste0('supertree', x), input_tree))
})

# Name the filtered datasets according to numbered supertree
names(phyloBoxPlot_supertrees) <- paste0("phyloBoxPlot_supertree_", supertree_numbers)
cat("\n","PROGRESS REP.: chunk [S5.2.07] done; starting next chunk...","\n")
#*-----**-----*

#*—————**{ [S5.2.08] Create plots showing influence of supertree on p-value significance in phybLR dataset. }*
# Create function to calculate identical vs. divergent counts
calculate_counts_trees <- function(...) {
  # Collect all supertrees into a list
  supertrees <- list(...)
  # Find common values using Reduce to apply intersect across all supertrees
  common_values <- Reduce(intersect, supertrees)
  # Calculate identical counts using sapply
  identical_count <- sum(sapply(supertrees, function(tree) sum(tree %in% common_values)))
  # Calculate total count
  total_count <- sum(sapply(supertrees, length))
  # Calculate divergent count
  divergent_count <- total_count - identical_count
  return(c(identical_count, divergent_count))
}

# Iterate over phybLR_pvals and calculate counts for each column
results_list_phybLR_trees <- lapply(phybLR_pvals, function(col) {
  # Extract supertree values for the current column
  supertree_values <- lapply(phybLR_supertrees, function(tree) tree[[col]])
  # Calculate counts using the previously defined function
  counts <- do.call(calculate_counts_trees, supertree_values)
  # Create a data frame for the results
  data.frame(
    Category = c("Identical", "Divergent"),
    Count = counts,
    Column = col
  )
})

# Convert the list of data frames into a single data frame
results_phybLR_trees <- do.call(rbind, results_list_phybLR_trees)

# Convert counts to percentages
results_phybLR_trees <- results_phybLR_trees %>%
  group_by(Column) %>%
  mutate(Percentage = Count / sum(Count) * 100)

# Create the stacked percentage bar plot
ggplot(results_phybLR_trees, aes(x = Column, y = Percentage, fill = Category)) +
  geom_bar(stat = "identity", position = "fill") +
  scale_fill_manual(values=c('brown2', 'springgreen4')) +
  scale_y_continuous(labels = scales::percent) +
  labs(title = "Ratio of identical vs divergent statistical significance between different supertrees in the phybLR dataset",
       x = "value", y = "consensus") + theme(axis.text.x = element_text(angle = 90, vjust = 0.5, hjust=1),plot.title = element_text(hjust = 0.5))
cat("\n","PROGRESS REP.: chunk [S5.2.08] done; starting next chunk...","\n")
#*-----**-----*

#*—————**{ [S5.2.09] Create plots showing influence of supertree on p-value significance in phycorr dataset. }*
# Iterate over phybLR_pvals and calculate counts for each column
results_list_phycorr_trees <- lapply(phycorr_pvals, function(col) {
  # Extract supertree values for the current column
  supertree_values <- lapply(phycorr_supertrees, function(tree) tree[[col]])
  # Calculate counts using the previously defined function
  counts <- do.call(calculate_counts_trees, supertree_values)
  # Create a data frame for the results
  data.frame(
    Category = c("Identical", "Divergent"),
    Count = counts,
    Column = col
  )
})

# Convert the list of data frames into a single data frame
results_phycorr_trees <- do.call(rbind, results_list_phycorr_trees)

# Convert counts to percentages
results_phycorr_trees <- results_phycorr_trees %>%
  group_by(Column) %>%
  mutate(Percentage = Count / sum(Count) * 100)

# Create the stacked percentage bar plot
ggplot(results_phycorr_trees, aes(x = Column, y = Percentage, fill = Category)) +
  geom_bar(stat = "identity", position = "fill") +
  scale_fill_manual(values=c('brown2', 'springgreen4')) +
  scale_y_continuous(labels = scales::percent) +
  labs(title = "Ratio of identical vs divergent statistical significance between different supertrees in the phycorr dataset",
       x = "value", y = "consensus") + theme(axis.text.x = element_text(angle = 90, vjust = 0.5, hjust=1),plot.title = element_text(hjust = 0.5))
cat("\n","PROGRESS REP.: chunk [S5.2.09] done; starting next chunk...","\n")
#*-----**-----*

#*—————**{ [S5.2.10] Create plots showing influence of supertree on p-value significance in phyloBoxPlot dataset.  }*
# Iterate over phybLR_pvals and calculate counts for each column
results_list_phyloBoxPlot_trees <- lapply(phyloBoxPlot_pvals, function(col) {
  # Extract supertree values for the current column
  supertree_values <- lapply(phyloBoxPlot_supertrees, function(tree) tree[[col]])
  # Calculate counts using the previously defined function
  counts <- do.call(calculate_counts_trees, supertree_values)
  # Create a data frame for the results
  data.frame(
    Category = c("Identical", "Divergent"),
    Count = counts,
    Column = col
  )
})

# Convert the list of data frames into a single data frame
results_phyloBoxPlot_trees <- do.call(rbind, results_list_phyloBoxPlot_trees)

# Convert counts to percentages
results_phyloBoxPlot_trees <- results_phyloBoxPlot_trees %>%
  group_by(Column) %>%
  mutate(Percentage = Count / sum(Count) * 100)

# Create the stacked percentage bar plot
ggplot(results_phyloBoxPlot_trees, aes(x = Column, y = Percentage, fill = Category)) +
  geom_bar(stat = "identity", position = "fill") +
  scale_fill_manual(values=c('brown2', 'springgreen4')) +
  scale_y_continuous(labels = scales::percent) +
  labs(title = "Ratio of identical vs divergent statistical significance between different supertrees in the phyloBoxPlot dataset",
       x = "value", y = "consensus") + theme(axis.text.x = element_text(angle = 90, vjust = 0.5, hjust=1),plot.title = element_text(hjust = 0.5))

# Create the stacked percentage bar plot summarizing all transformation method ratios
results_trees_total <- rbind(results_phybLR_trees,results_phycorr_trees,results_phyloBoxPlot_trees) %>%
  arrange(match(Column, c("phylANOVA_omnibus_pval", "phylANOVA_posthoc_pvals", "phyloLev_omnibus_pval", "phyloLev_posthoc_pvals", "intercept.p.val", "slope.p.val", "AUC_binomt_pval", "bestmod_binomt_pval", "McF_pval_mean", "McF_pval_for_full_ds")), desc(Category))

# Mutate the column "Column" into a factor so that the row order in this column is respected by the barplot
results_trees_total$Column <- factor(results_trees_total$Column, levels = unique(results_trees_total$Column))

# Create the stacked percentage bar plot
ggplot(results_trees_total, aes(x = Column, y = Percentage, fill = Category)) +
  geom_bar(stat = "identity", position = "fill") +
  scale_fill_manual(values=c('brown2', 'springgreen4')) +
  scale_y_continuous(labels = scales::percent) +
  labs(title = "Ratio of identical vs divergent statistical significance between different supertrees",
       x = "value", y = "consensus") + theme(axis.text.x = element_text(angle = 90, vjust = 0.5, hjust=1),plot.title = element_text(hjust = 0.5))

# Save the plot as a PDF
ggsave(path = output_path, file = paste0("significance_trees_barplot_", Sys.Date(), ".pdf"), plot = last_plot(), device = "pdf", width = 8, height = 6)
cat("\n","PROGRESS REP.: chunk [S5.2.10] done; starting next chunk...","\n")
#*-----**-----*

#*—————**{ [S5.2.11] Run binomial exact t-tests to see whether results are significantly different from a random distribution in the transformation datasets}*
# Reorganize results_trans_total dataset to summarize data for each pvalue in one row
results_trans_identical <- subset(results_trans_total, Category == 'Identical') %>%
  rename(Count_identical = Count) %>%
  rename(Percentage_identical = Percentage) %>%
  rename(Pvalue = Column) %>%
  select(-Category)

results_trans_divergent <- subset(results_trans_total, Category == 'Divergent') %>%
  rename(Count_divergent = Count) %>%
  rename(Percentage_divergent = Percentage) %>%
  rename(Pvalue = Column) %>%
  select(-Category)

results_trans_total <- left_join(results_trans_identical, results_trans_divergent, by = 'Pvalue') %>%
  select(Pvalue,Count_identical,everything())

# Perform the binomial test and extract p-value, confidence interval, and estimate for the trans dataset
results_trans_total <- results_trans_total %>%
  rowwise() %>%
  mutate(
    # Perform the binomial test for each row
    binom_test = list(binom.test(
      x = Count_identical,
      n = Count_identical + Count_divergent,
      p = 0.25,
      alternative = "two.sided",
      conf.level = 0.95
    )),
    
    # Extract the p-value from the test
    Binom_test_pval = binom_test$p.value,
    # Extract the confidence interval from the test
    Binom_test_conf_low = binom_test$conf.int[1],
    Binom_test_conf_high = binom_test$conf.int[2],
    # Extract the estimate from the test
    Binom_test_estimate = binom_test$estimate
  ) %>%
  ungroup() %>%
  select(-binom_test)  # Remove the temporary 'binom_test' column
cat("\n","PROGRESS REP.: chunk [S5.2.11] done; starting next chunk...","\n")
#*-----**-----*

#*—————**{ [S5.2.12] Run binomial exact t-tests to see whether results are significantly different from a random distribution in the tree datasets}*
# Reorganize results_trees_total dataset to summarize data for each pvalue in one row
results_trees_identical <- subset(results_trees_total, Category == 'Identical') %>%
  rename(Count_identical = Count) %>%
  rename(Percentage_identical = Percentage) %>%
  rename(Pvalue = Column) %>%
  select(-Category)

results_trees_divergent <- subset(results_trees_total, Category == 'Divergent') %>%
  rename(Count_divergent = Count) %>%
  rename(Percentage_divergent = Percentage) %>%
  rename(Pvalue = Column) %>%
  select(-Category)

results_trees_total <- left_join(results_trees_identical, results_trees_divergent, by = 'Pvalue')%>%
  select(Pvalue,Count_identical,everything())

# Perform the binomial test and extract p-value, confidence interval, and estimate for the trees dataset
results_trees_total <- results_trees_total %>%
  rowwise() %>%
  mutate(
    # Perform the binomial test for each row
    binom_test = list(binom.test(
      x = Count_identical,
      n = Count_identical + Count_divergent,
      p = 0.0078,
      alternative = "two.sided",
      conf.level = 0.95
    )),
    
    # Extract the p-value from the test
    Binom_test_pval = binom_test$p.value,
    # Extract the confidence interval from the test
    Binom_test_conf_low = binom_test$conf.int[1],
    Binom_test_conf_high = binom_test$conf.int[2],
    # Extract the estimate from the test
    Binom_test_estimate = binom_test$estimate
  ) %>%
  ungroup() %>%
  select(-binom_test)  # Remove the temporary 'binom_test' column
cat("\n","PROGRESS REP.: chunk [S5.2.12] done; starting next chunk...","\n")
#*-----**-----*

#*—————**{ [S5.2.13] Save the results_trans_total and results_trees_total files to the working directory. }*
# Save the dataframe "results_trees_total" as a CSV file
write.csv(results_trans_total, file = paste0(output_path, "/results_trans_total_", Sys.Date(), ".csv"), row.names = FALSE)

# Save the dataframe "results_trees_total" as a CSV file
write.csv(results_trees_total, file = paste0(output_path, "/results_trees_total_", Sys.Date(), ".csv"), row.names = FALSE)
cat("\n","PROGRESS REP.: chunk [S5.2.13] done; starting next chunk...","\n")
#*-----**-----*

# The results of this sensitivity analysis are presented in supplementary Table S6 and Extended Data Fig. 8, and discussed in detail in the main-text, Online Methods, and Supplementary Text S4. In brief, we recover very strong consensus across data transformation methods and supertrees for all phybLR-, phylANOVA-, and phyloLev-associated statistical tests, but poor consensus across both treatment types for phylogenetic correlation tests. As a result, we ultimately present a separate phylogenetic correlation matrix for each tree (see Supplementary Figures), and generate these separate correlation matrices for each tree below.
#------------------------------------------------------------------------------

#**"-------------------------------------------------------------------------"*
#**"---- ### [S5.3] MAKE CORRELATION MATRICES USING PHYCORR RESULTS. ### ----"*
#**"-------------------------------------------------------------------------"*

#*-----**{ [S5.3.01] Make correlogram for raw (un-transformed), phylogenetically UN-corrected data). }*
# We will begin this section by generating a correlogram for raw, phylogenetically un-corrected data. These correlation tests are not valid (since they fail to account for phylogenetic covariance), but they provide a visual reference to the researcher who wants a sense for how the uncorrected data are correlated prior to being corrected for phylogenetic autocorrelation.

# Calculate raw correlation coefficients and p-vals (w/out phylo correction):
sigcorr <- cor.mtest(na.omit(all_mm[,all_quantitative_vars]), conf.level = .95)
par(mar = c(2,2,2,2), oma = c(0,1,0,1))

# Generate correlogram:
tiff(filename=paste0(cortest_output_path, "/NOphy__all_mm__corrplot.tif"), width=8000, height=8000, units="px", res=250)
options(repr.plot.width=20, repr.plot.height=20)
corrplot.mixed(cor(na.omit(all_mm[,all_quantitative_vars])), lower.col = "black", upper = "square", lower="number", tl.col = "black", number.cex =0.8, tl.pos = "lt", tl.cex=1.5, p.mat = sigcorr$p, sig.level = .05, diag = "n", main = "Correlogram of quantitative LM data, NOT phylo-corrected", cex.main = 1.5, mar = c(1,1,1,1))
dev.off()
cat("\n","PROGRESS REP.: chunk [S5.3.01] done; starting next chunk...","\n")
#*-----**-----*

#*-----**{ [S5.3.02] Make correlograms for raw (un-transformed), phylogenetically CORRECTED data). }*
# We will next generate eight high-resolution correlograms for raw (un-transformed) but phylogenetically corrected data--one for each of our supertrees. 

phycorr_raw_table <- phycorr_metadata_table[phycorr_metadata_table$input_dataset=="all_mm", ]

# Make separate correlogram showing phycorr() results for each tree:
for(t in 1:length(tree_variants)){
  # Pick out all phycorr output rows associated with raw data for current tree:
  phycorr_for_this_tree <- phycorr_raw_table[phycorr_raw_table$input_tree == tree_variant_names[t], ]
  
  # Initialize output matrices for Pearson correlation coefficients and p-vals:
  cor_mat <- matrix(nrow=length(corvariables), ncol=length(corvariables))
  rownames(cor_mat) <- corvariables
  colnames(cor_mat) <- corvariables
  p_mat <- matrix(nrow=length(corvariables), ncol=length(corvariables))
  rownames(p_mat) <- corvariables
  colnames(p_mat) <- corvariables
  
  # Populate cor_mat and p_mat for current tree:
  for(r in 1:nrow(phycorr_for_this_tree)){
    corvar1 <- phycorr_for_this_tree$corvariable1[r]
    corvar2 <- phycorr_for_this_tree$corvariable2[r]
    if (corvar1==corvar2) {
      cor_mat[corvar1,corvar2] <- 1
      p_mat[corvar1,corvar2] <- 0 
    } else if (!(corvar1==corvar2)) {
      cor_mat[corvar1,corvar2] <- phycorr_for_this_tree$'Rsq'[r] # misnamed Rsq
      cor_mat[corvar2,corvar1] <- phycorr_for_this_tree$'Rsq'[r] # misnamed Rsq
      p_mat[corvar1,corvar2] <- phycorr_for_this_tree$'slope.p.val'[r]
      p_mat[corvar2,corvar1] <- phycorr_for_this_tree$'slope.p.val'[r]
      # Note that the p-val cannot be NA for corrplot.mixed() function.
    } # closes 'if (corvar1==corvar2) { } else { }' clause
  } # closes 'for(r in 1:nrow(phycorr_for_this_tree))' loop
  
  # The loop below ensures that there will not be a mismatch between matrices:
  for(r in 1:nrow(p_mat)){
    for(c in 1:ncol(p_mat)){
      if(is.na(p_mat[r,c])==T){
        cor_mat[r,c] <- NA
        cor_mat[c,r] <- NA
        p_mat[c,r] <- NA
        p_mat[r,c] <- NA
      }
      if(is.na(cor_mat[r,c])==T){
        cor_mat[r,c] <- NA
        cor_mat[c,r] <- NA
        p_mat[c,r] <- NA
        p_mat[r,c] <- NA
    }
    }
  }

  # Generate high-resolution tiff of correlogram for current tree:
  tiff(filename=paste0(cortest_output_path, "/YESphy__all_mm__RAW_for_supertree", t, "_corrplot.tif"), width=8000, height=8000, units="px", res=300)
  corrplot.mixed(corr=cor_mat, is.corr=T, upper='square', lower='number', tl.pos='lt', tl.cex=1.5, tl.col='black', lower.col='black', number.cex=0.7, p.mat=p_mat, sig.level=.05, diag="n", main="Correlogram of phylogenetically corrected raw (untransformed) data", mar=c(1,1,1,1))
  dev.off() 
} # closes 'for(t in 1:length(tree_variants))' loop

# Once this chunk is done running, a separate correlogram for each tree, with all pairwise phylogenetically corrected correlations among quantitative variables, will have been saved to the working directory, along with a CSV file containing all test results.
cat("\n","PROGRESS REP.: chunk [S5.3.02] done; starting next chunk...","\n")
#*-----**-----*

#**"-------------------------------------------------------------------------"*
#**"- ### [[S5.4]] USE ROC ANALYSIS TO PICK BEST MODEL-THRESHOLD PAIRS ### -"*
#**"-------------------------------------------------------------------------"*
# In the following section, we will define a pair of original functions that let us select, for a given tree and binary response variable, the single best-performing model (ie, the best predictor variable) for each of "forelimb region proportions" (f_AZR, f_ASR, f_SZR), "forelimb acropod planform dimensions" (mD_Symmetry_index1, mD_Symmetry_index2), "mD3 ungual proportions" (mD3_up_FI1, mD3_up_FI2, mD3_up_FI3), "mD3 proximal phalanx proportions" (mD3_proxp_Length.DistWidth, mD3_proxp_WidthRatios), "hindlimb region proportions" (h_AZR, h_SZR, h_ASR), "hindlimb acropod planform dimensions" (pD_Symmetry_index1, pD_Symmetry_index2), "pD3 ungual proportions"(pD3_up_FI1, pD3_up_FI2, pD3_up_FI3), and "pD3 proximal phalanx proportions" (pD3_proxp_ Length.DistWidth, pD3_proxp_WidthRatios). We will ultimately use the previously generated phybLR output list items for these best models to make ROC plots that let us visualize relative model performance for different sets of predictors. 

# To start, we will define the phycROCCs() function below, which makes an ROC plot from a given set of phybLR object inputs.

#*-----**{ [S5.4.01] Define phycROCCs() function. }*
phycROCCs <- function(phybLR_mod_list, ROC_curve_names, ROC_colors, plot_cROCC_CIs=FALSE, predictor_set_names, output_name){
  # DESCRIBING FUNCTION PARAMETERS:
  # phybLR_mod_list: A list of ROC curves (output by phybLR() or similar)
  # ROC_curve_names: A chr vector with names for all listed cROCCs (must be in same order as ROC_object_list)
  # ROC_colors: A chr vector with colors for all listed cROCCs (must be in same order as ROC_object_list)
  # plot_cROCC_CIs: TRUE or FALSE, specifies whether 95% CI bands will be plotted for each cROCC. 
  # predictor_set_names: A chr vector with name for each ROC's predictor set (must be in same order as ROC_object_list)
  # output_name: A character string specifying the title of the ROC plot and associated output file (note: this should NOT include '.png')
  
  # Define directory path for output files: 
  directory_path <- paste0(output_path, "/phybLR_cROCCs")
  # Create directory for output files:
  dir.create(path=directory_path, showWarnings = TRUE, recursive = FALSE, mode = "0777")
  
  # Check that phybLR_mod_list and its constituent items are formatted right:
  if(is.list(phybLR_mod_list)==F){
    stop("Error: The phybLR_mod_list input is not a list; it must be a list of phybLR() outputs.")
    } # closes 'if(is.list(phybLR_mod_list)==F)' clause
  for(i in 1:length(phybLR_mod_list)){
    if(is.list(phybLR_mod_list[[i]])==F){
      stop("Error: One of the items in phybLR_mod_list is not an object of class 'phybLR'.")
    } # closes 'if(is.list(phybLR_mod_list[[i]])==F)' clause
  } # closes 'for(i in 1:length(phybLR_mod_list))' loop
  
  # Shorten ROC_colors vector to length of phybLR_mod_list:
    # NOTE: The following single line was written with the help of chatGPT:
  ROC_colors <- ROC_colors[1:length(phybLR_mod_list)]
  
  # Create ROC_dfs object to contain coordinates for each ROC curve:
  ROC_dfs <- as.list(rep(NA, length(phybLR_mod_list)))
  
  for(i in 1:length(phybLR_mod_list)){
    ROC_dfs[[i]] <- data.table("FPRs" = phybLR_mod_list[[i]]$cROC_xvals, "TPRs" = phybLR_mod_list[[i]]$cROC_yvals)
    } # closes 'for(i in 1:length(phybLR_mod_list))' loop
  
  # Create ROC_CI_dfs object to contain coords for each set of ROC_CI bounds:
  ROC_CI_dfs <- as.list(rep(NA, length(phybLR_mod_list)))
  
  for(i in 1:length(phybLR_mod_list)){
    ROC_CI_dfs[[i]] <- data.table("FPRs" = phybLR_mod_list[[i]]$cROC_xvals, "TPRs" = phybLR_mod_list[[i]]$cROC_yvals, "ROC_95ci_lowbound_xvals" = phybLR_mod_list[[i]]$ROC_95ci_lowbound_xvals, "ROC_95ci_lowbound_yvals" = phybLR_mod_list[[i]]$ROC_95ci_lowbound_yvals, "ROC_95ci_highbound_xvals" = phybLR_mod_list[[i]]$ROC_95ci_highbound_xvals, "ROC_95ci_highbound_yvals" = phybLR_mod_list[[i]]$ROC_95ci_highbound_yvals)
    } # closes 'for(i in 1:length(phybLR_mod_list))' loop
  
  # Create vectors to store model performance metrics:
  AUCs <- rep(NA, length(phybLR_mod_list))
  AUC_pvals <- rep(NA, length(phybLR_mod_list))
  best_thresholds <- rep(NA, length(phybLR_mod_list))
  bestmod_pvals <- rep(NA, length(phybLR_mod_list))
  for(i in 1:length(phybLR_mod_list)){
    AUCs[i] <- phybLR_mod_list[[i]]$AUC_mean
    AUC_pvals[i] <- phybLR_mod_list[[i]]$AUC_binomtest_pval
    best_thresholds[i] <- phybLR_mod_list[[i]]$bestmod_threshold
    bestmod_pvals[i] <- phybLR_mod_list[[i]]$bestmod_binomtest_pval
  } # closes 'for(i in 1:length(phybLR_mod_list))' loop
  
  # Name roctab columns: 
  rocvecs <- as.list(rep(NA, length(ROC_dfs)))
  ROCols <- ROC_colors
  
  # Populate rocvec output vectors:
  for(i in 1:length(rocvecs)){
    AUC_val <- round(phybLR_mod_list[[i]]$AUC_mean, 2)
    AUC_plab<-paste("p = ",format(AUC_pvals[i],scientific=T,digits=4),sep="")
    rocvecs[[i]] <- c("Predictors"=predictor_set_names[i], "AUC"=AUC_val, "Binomial Test"=AUC_plab)
  } # closes 'for(i in 1:length(rocvecs))' loop
  
  # Populate roctab output table:
  roctab <- rocvecs[[1]] # jump-start roctab with rocvec[[1]] as first row
  for(i in 2:length(rocvecs)){
    roctab <- rbind(roctab, rocvecs[[i]])}
  
  # Set table dimensions:
  # most aesthetic total height and width were estimated by trial & error
  total_table_height <- 5.6
  first_column_width <- 9.5
  second_col_width <- 3
  third_col_width <- 6
  cell_heights <- unit(rep(total_table_height/nrow(roctab),nrow(roctab)),"cm")
  cell_widths_unscaled <- c(first_column_width, second_col_width, third_col_width)
  cell_widths <- unit(cell_widths_unscaled, "cm") 
  
  # Set table column and row labels:
  Predictors_lab <- "Predictors"
  AUC_lab <- "AUC"
  colnames(roctab) <- c(Predictors_lab, AUC_lab, "Binomial Test")
  rownames(roctab) <- NULL
  
  # Setting tableGrob theme:
  font_matrix <- matrix(data=2, nrow(roctab), ncol(roctab))
  font_matrix[,1] <- rep(2, nrow(roctab))
  fill_matrix <- matrix("black", nrow(roctab), ncol(roctab))
  border_matrix <- matrix("whitesmoke", nrow(roctab), ncol(roctab))
  border_matrix[,1] <- rep("whitesmoke", nrow(roctab))
  col_matrix <- matrix("white", nrow(roctab), ncol(roctab))
  
  for(i in 1:nrow(col_matrix)){
    col_matrix[i,1] <- ROCols[i]}
  custom_theme <- ttheme_default(base_size=20, border = TRUE, core=list(fg_params = list(col = col_matrix, fontface = font_matrix, hjust=0.5), bg_params = list(fill = fill_matrix, col = border_matrix, lwd=2)), rowhead=list(bg_params = list(col="whitesmoke", fill = "whitesmoke", lwd=8), fg_params = list(col="black", fontsize=0)), colhead=list(bg_params = list(col="whitesmoke", fill="white", lwd=8)))
  
  roctab_grob <- tableGrob(roctab, theme=custom_theme, heights=cell_heights, widths=cell_widths)
  
  # Define a set of geom_line commands--one for each ROC curve in the ROC_list:
  geom_line_set_text <- ""
  reverse_sequence <- seq(from=length(ROC_dfs), to=1) # Making this a reverse sequence ensures that earlier ROC curves will plot on top of later ones, which is what we want, since the first entry in each ROC_list (limb region proportions) was found to be the best model. We want that curve to be the most visible, so it should not be covered by any of the others.
  for(i in 1:length(ROC_dfs)){
    if(i == length(ROC_dfs)){lwd_val <- 6} 
    # this makes the first ROC_list item the thickest curve
    else {lwd_val <- 3}
    i_mod <- reverse_sequence[i]
    if(!(i == length(ROC_dfs))){
      geom_line_set_text <- paste(geom_line_set_text, "geom_line(data = ROC_dfs[[", i_mod, "]], show.legend = TRUE, aes(x = FPRs, y = TPRs), linetype='solid', linewidth=", lwd_val, ", color = ROCols[", i_mod, "], alpha=0.9) + ", sep="")
    } else {
      geom_line_set_text <- paste(geom_line_set_text, "geom_line(data = ROC_dfs[[", i_mod, "]], show.legend = TRUE, aes(x = FPRs, y = TPRs), linetype='solid', linewidth=", lwd_val, ", color = ROCols[", i_mod, "], alpha=0.9)", sep="")
    }
  } # closes 'for(i in 1:length(ROC_dfs))' loop
  
  # Define a set of geom_errorbar commands—one for each cROCC in the ROC_list:
  geom_CI_bands_set_text <- ""
  if(plot_cROCC_CIs==TRUE){
    reverse_sequence <- seq(from=length(ROC_CI_dfs), to=1) # Making this a reverse sequence ensures that earlier ROC curves will plot on top of later ones, which is what we want, since the first entry in each ROC_list (limb region proportions) was found to be the best model. We want that curve to be the most visible, so it should not be covered by any of the others.
    for(i in 1:length(ROC_CI_dfs)){
      i_mod <- reverse_sequence[i]
      geom_CI_bands_set_text <- paste(geom_CI_bands_set_text, "geom_point(data = ROC_CI_dfs[[", i_mod, "]], inherit.aes=F, aes(x = FPRs, y = TPRs)) + geom_ribbon(data = ROC_CI_dfs[[", i_mod, "]], aes(ymin = ROC_95ci_lowbound_yvals, ymax = ROC_95ci_highbound_yvals, x = FPRs), fill = ROCols[", i_mod, "], outline.type='full', alpha=0.2) + ", sep="")
    } # closes 'for(i in 1:length(ROC_CI_dfs))' loop
  } # closes 'if(plot_cROCC_CIs==TRUE)' clause
  
  # Define one set of geom_point commands for each cROCC bestmod in ROC_list:
  geom_point_set_text <- ""
  reverse_sequence <- seq(from=length(ROC_dfs), to=1) # Making this a reverse sequence ensures that earlier ROC curves will plot on top of later ones, which is what we want, since the first entry in each ROC_list (limb region proportions) was found to be the best model. We want that curve to be the most visible, so it should not be covered by any of the others.
  for(i in 1:length(ROC_dfs)){
    i_mod <- reverse_sequence[i]
    if(i == length(ROC_dfs)){size_val <- 8} 
    # this makes the first ROC_list point size match curve thickness
    else {size_val <- 4}
    point_coords <- c(phybLR_mod_list[[i_mod]]$bestmod_FPR_mean, phybLR_mod_list[[i_mod]]$bestmod_TPR_mean)
    if(!(i == length(ROC_dfs))){
      geom_point_set_text <- paste(geom_point_set_text, "geom_point(aes(x =", point_coords[1], ", y =", point_coords[2], "), shape = 21, color = 'white', fill = ROCols[", i_mod, "], alpha=0.9, size=", size_val, ", stroke=2) +", sep="")
    } else {
      geom_point_set_text <- paste(geom_point_set_text, "geom_point(aes(x =", point_coords[1], ", y =", point_coords[2], "), shape = 21, color = 'white', fill = ROCols[", i_mod, "], alpha=0.9, size=", size_val, ", stroke=2.5)", sep="")
    }
  } # closes 'for(i in 1:length(ROC_dfs))' loop
  
  # Define first ROC curve to initialize plot:
  ROC_base_data <- as.data.frame(cbind(phybLR_mod_list[[1]]$cROC_xvals, phybLR_mod_list[[1]]$cROC_yvals))
  colnames(ROC_base_data) <- c("cROC_xvals", "cROC_yvals")
  # Paste the geom_line_set_text into a larger text string that contains all the other commands required to make a nice ggplot:
  ggrocplot_text <- paste("ggplot(data=ROC_base_data, linewidth=1.3, alpha=0) + geom_segment(x=0, xend=1, y=0, yend=1, linetype='longdash', linewidth=2.2, color='white', alpha=0.05) + ylim(0, 1) + xlim(0,1) + xlab('False Positive Rate') + ylab('True Positive Rate') + ", geom_CI_bands_set_text, geom_line_set_text, "+", geom_point_set_text, "+ theme(legend.position = 'bottom', axis.text = element_text(size=20), axis.title = element_text(size=20), panel.background=element_rect(fill='black'), axis.line = element_line(color = 'grey20'), panel.grid = element_line(color = 'grey20'), aspect.ratio=1) + ggtitle(paste('ROC plot for', output_name, sep=' ')) + annotation_custom(roctab_grob, xmin = 0.49, xmax = 0.82, ymin = -0.05, ymax = 0.25)", sep="")
  
  # Convert the ggrocplot_text string into a ggplot object:
  ggrocplot <- eval(parse(text=ggrocplot_text))
  
  # Save the plot as a png file:
  png(filename = paste0(directory_path, "/", output_name, "__cROCCs.png"), width=12.5, height=12.5, units="in", res=300)
  show(ggrocplot)
  dev.off()
  
  # Save a text file with the associated dataset-variable-tree combination info for each ROC curve:
  sink(file = paste0(directory_path, "/", output_name, "_ds-var-tree_combos.txt"))
  cat("\n", "Dataset-variable-tree combinations for each ROC curve in order from top to bottom:", "\n")
  for(i in 1:length(ROC_curve_names)){
    cat("---------------------------------------------------------------", "\n", "CURVE NUMBER", i, ":", "COLOR:", ROC_colors[i], "|", ROC_curve_names[[i]], "\n", "---------------------------------------------------------------", "\n")
  } # closes 'for(i in 1:length(ROC_curve_names))' loop
  sink(file = NULL); sink(file = NULL)
  
  # Save associated 95% CI plot for AUC:
  # Define a set of geom commands--one for each ROC curve in the ROC_list:
  reverse_sequence <- seq(from=length(ROC_curve_names), to=1)
  geom_set_text <- c("")
  for(i in 1:length(ROC_curve_names)){
    i_mod <- reverse_sequence[i]
    AUC_ci_min <- round(phybLR_mod_list[[i]]$AUC_95ci[1], 3)
    AUC_ci_max <- round(phybLR_mod_list[[i]]$AUC_95ci[2], 3)
    AUC_mean <- round(phybLR_mod_list[[i]]$AUC_mean, 3)
    geom_set_text <- paste(geom_set_text, "+ geom_segment(aes(x=", i_mod, ", xend=", i_mod, ", y=", AUC_ci_min, ", yend=", AUC_ci_max, "), color = 'grey30', alpha=0.6, size=4) + geom_segment(aes(x=", i_mod, ", xend=", i_mod, ", y=", AUC_ci_min, ", yend=", AUC_ci_max, "), color = ROC_colors[", i, "], alpha=0.6, size=2) + geom_point(aes(x=", i_mod, ", y=", AUC_mean, "), shape=21, color='grey10', fill=ROCols[", i, "], size=6, stroke=4, alpha=0.8)", sep="")
  } # closes 'for(i in 1:length(ROC_curve_names))' loop
  
  # Paste the geom_set_text into a larger text string that contains all the other commands required to make a nice ggplot:
  ggauc_text <- paste("ggplot() + geom_hline(aes(yintercept=0.5), linetype='dashed', color='grey15', alpha=0.5, size=1)", geom_set_text, "+ lims(y=c(0,1), x=c(0.5,length(ROC_curve_names)+0.5)) + ggtitle('95% CIs for AUC from 100 phybLR subsampling iterations') + ylab('ROC curves (key in .txt file)') + xlab('AUC') + coord_flip()", sep=" ")
  
  # Convert the ggauc_text string into a ggplot object:
  ggauc_plot <- eval(parse(text=ggauc_text))
  
  # Save the plot as a png file:
  png(filename = paste(directory_path, "/", output_name, "__AUC_confints.png", sep=""), width=4, height=12, units="in", res=300)
  show(ggauc_plot)
  dev.off()  
  
  # Save associated 95% CI plot for TPR:
  # Define a set of geom commands--one for each ROC curve in the ROC_list:
  reverse_sequence <- seq(from=length(ROC_colors), to=1)
  TPR_set_text <- c("")
  for(i in 1:length(ROC_colors)){
    i_mod <- reverse_sequence[i]
    ci_min <- round(phybLR_mod_list[[i]]$bestmod_TPR_95ci[1], 3)
    ci_max <- round(phybLR_mod_list[[i]]$bestmod_TPR_95ci[2], 3)
    TPR_mean <- round(phybLR_mod_list[[i]]$bestmod_TPR_mean, 3)
    TPR_set_text <- paste(TPR_set_text, "+ geom_segment(aes(x=", i_mod, ", xend=", i_mod, ", y=", ci_min, ", yend=", ci_max, "), color = 'grey30', alpha=0.6, size=4) + geom_segment(aes(x=", i_mod, ", xend=", i_mod, ", y=", ci_min, ", yend=", ci_max, "), color = ROC_colors[", i, "], alpha=0.6, size=2) + geom_point(aes(x=", i_mod, ", y=", TPR_mean, "), shape=21, color='grey10', fill=ROCols[", i, "], size=6, stroke=4, alpha=0.8)", sep="")
  } # closes 'for(i in 1:length(ROC_colors))' loop
  
  # Paste the TPR_set_text into a larger text string that contains all the other commands required to make a nice ggplot:
  ggTPR_text <- paste("ggplot() + geom_hline(aes(yintercept=0.5), linetype='dashed', color='grey15', alpha=0.5, size=1)", TPR_set_text, "+ lims(y=c(0,1), x=c(0.5,length(ROC_colors)+0.5)) + ggtitle('95% CIs for AUC from 100 phybLR subsampling iterations') + ylab('ROC curves (key in .txt file)') + xlab('TPR') + coord_flip()", sep=" ")
  
  # Convert the ggTPR_text string into a ggplot object:
  ggTPR_plot <- eval(parse(text=ggTPR_text))
  
  # Save the plot as a png file:
  png(filename = paste0(directory_path, "/", output_name, "__TPR_confints.png"), width=4, height=12, units="in", res=300)
  show(ggTPR_plot)
  dev.off()  
  
  # Save associated 95% CI plot for FPR:
  # Define a set of geom commands--one for each ROC curve in the ROC_list:
  reverse_sequence <- seq(from=length(ROC_curve_names), to=1)
  FPR_set_text <- c("")
  for(i in 1:length(ROC_curve_names)){
    i_mod <- reverse_sequence[i]
    ci_min <- round(phybLR_mod_list[[i]]$bestmod_FPR_95ci[1], 3)
    ci_max <- round(phybLR_mod_list[[i]]$bestmod_FPR_95ci[2], 3)
    FPR_mean <- round(phybLR_mod_list[[i]]$bestmod_FPR_mean, 3)
    FPR_set_text <- paste(FPR_set_text, "+ geom_segment(aes(x=", i_mod, ", xend=", i_mod, ", y=", ci_min, ", yend=", ci_max, "), color = 'grey30', alpha=0.6, size=4) + geom_segment(aes(x=", i_mod, ", xend=", i_mod, ", y=", ci_min, ", yend=", ci_max, "), color = ROC_colors[", i, "], alpha=0.6, size=2) + geom_point(aes(x=", i_mod, ", y=", FPR_mean, "), shape=21, color='grey10', fill=ROCols[", i, "], size=6, stroke=4, alpha=0.8)", sep="")
  } # closes 'for(i in 1:length(ROC_curve_names))' loop
  
  # Paste the FPR_set_text into a larger text string that contains all the other commands required to make a nice ggplot:
  ggFPR_text <- paste("ggplot() + geom_hline(aes(yintercept=0.5), linetype='dashed', color='grey15', alpha=0.5, size=1)", FPR_set_text, "+ lims(y=c(0,1), x=c(0.5,length(ROC_curve_names)+0.5)) + ggtitle('95% CIs for AUC from 100 phybLR subsampling iterations') + ylab('ROC curves (key in .txt file)') + xlab('FPR') + coord_flip()", sep=" ")
  
  # Convert the ggFPR_text string into a ggplot object:
  ggFPR_plot <- eval(parse(text=ggFPR_text))
  
  # Save the plot as a png file:
  png(filename = paste0(directory_path, "/", output_name, "__FPR_confints.png"), width=4, height=12, units="in", res=300)
  show(ggFPR_plot)
  dev.off()  
} # closes phycROCCs() function
cat("\n","PROGRESS REP: chunk [S5.4.01] complete; starting next chunk..","\n")
#*-----**-----*

#*-----**{ [S5.4.02] Test phycROCCs() function on sample cROCC curve list. }*
# Define test_ROClist object and associated vectors:
test_ROClist <- bLR_mod_list[c(1,10,14,999)]
test_ROCnames <- bLR_mod_names[c(1,10,14,999)]
roc_colors_vec <- c("green", "hotpink", "purple", "grey60")
predictor_names_vec <- c("f_AnAR", "mD_Symmetry_index2", "mD3_up_FI2", "mD3_proxp_WidthRatios")

# Check test_ROClist object to confirm that it looks right:
test_ROClist[[1]]$AUC_95ci[1]
test_ROClist[[1]]$AUC_mean

# Test phycROCCs() function:
phycROCCs(phybLR_mod_list = test_ROClist, ROC_curve_names = test_ROCnames, predictor_set_names = predictor_names_vec, ROC_colors = roc_colors_vec, plot_cROCC_CIs=TRUE, output_name = "*test_ROC_plot")
cat("\n","PROGRESS REP: chunk [S5.4.02] complete; starting next chunk..","\n")
#*-----**-----*

#*-----**{ [S5.4.03] Define phybLR-model filtering and selector functions. }*
# The phycROCCs() function above lets us plot any set of phybLR cROCCs together in a single plot. We will now use this function and its outputs (i.e., use ROC analysis) to select the best-performing models for each series of morphometric variables in our dataset: forelimb proportions, hindlimb proportions, manus planform dimensions, pes planform dimensions, mD3 ungual dimensions, pD3 ungual dimensions, mD3 proximal phalanx dimensions, and pD3 proximal phalanx dimensions. To do this, we will initialize several objects to automate down-stream ROC plotting, define new model-filtering and model-selector functions to identify the best predictor out of each variable series, and use phycROCCs() to plot the four series winners for forelimbs and the four series winners for hindlimbs.

# Initialize objects to automate downstream ROC plotting:
f_props_predictors <- c("f_AZR", "f_ASR", "f_SZR")
h_props_predictors <- c("h_AZR", "h_ASR", "h_SZR")
f_acr_predictors <- c("mD_Symmetry_index1", "mD_Symmetry_index2")
h_acr_predictors <- c("pD_Symmetry_index1", "pD_Symmetry_index2")
f_mD3_ungp_predictors <- c("mD3_up_FI1", "mD3_up_FI2", "mD3_up_FI3")
h_pD3_ungp_predictors <- c("pD3_up_FI1", "pD3_up_FI3")
f_mD3_proxp_predictors <- c("mD3_proxp_Length.DistWidth", "mD3_proxp_WidthRatios")
h_pD3_proxp_predictors <- c("pD3_proxp_Length.DistWidth", "pD3_proxp_WidthRatios")

# Create separate list for forelimb and hindlimb variable series:
f_predictor_sets <- list(f_props_predictors, f_acr_predictors, f_mD3_ungp_predictors, f_mD3_proxp_predictors)
f_predictor_set_names <- c("forelimb_proportions", "f_acropod_dimensions", "mD3_ungual_proportions", "mD3_proxp_proportions")

h_predictor_sets <- list(h_props_predictors, h_acr_predictors, h_pD3_ungp_predictors, h_pD3_proxp_predictors)
h_predictor_set_names <- c("hindlimb_proportions", "h_acropod_dimensions", "pD3_ungual_proportions", "pD3_proxp_proportions")

# Define grouping variable & tree name vectors to automate downstream analyses:
bin_ecotype_groupers <- key_bLR_groupers[grep("eco", key_bLR_groupers)]
bin_forelimb_groupers <- key_bLR_groupers[grep("eco|fw|ff", key_bLR_groupers)]
bin_hindlimb_groupers <- key_bLR_groupers[grep("eco|hw|hf", key_bLR_groupers)]
tree_names <- c("supertree1", "supertree2", "supertree3", "supertree4", "supertree5", "supertree6", "supertree7", "supertree8")

# Write a function to subset the bLR_mod_list to only include those for a certain tree-dataset-response triad for a given set of predictor variables:
filter_bLR_mods <- function(full_bLR_mod_list, target_tree, target_dataset, target_response_variable, target_predictor_variables) {
  # Initialize filtered list:
  filtered_list <- list() 
  # Once populated, this list will be output to the working environment.
  for (i in 1:length(full_bLR_mod_list)) {
    if (length(names(bLR_mod_list[[i]]))>2) { # Since a skipped combination of phybLR inputs will output a list of length 2, this clause only proceeds with those phybLR input combinations that were not skipped. 
      # Extract the predictor variable for the current element
      current_predictor <- full_bLR_mod_list[[i]]$predictor_variable
      # OPTIONAL TROUBLESHOOTING LINE:
        # cat("\n", i, ":", "proceeded", "-", "current predictor :", current_predictor) 
      # Check if the predictor variable matches any of the target predictors
      if ((current_predictor %in% target_predictor_variables)==T) {
        # Extract tree, dataset, and response variable for the current element
        current_tree <- full_bLR_mod_list[[i]]$input_tree
        current_dataset <- full_bLR_mod_list[[i]]$input_dataset
        current_response_variable <- full_bLR_mod_list[[i]]$response_variable
        # Check if current element matches target tree, ds, & response_var:
        if (current_tree == target_tree && 
            current_dataset == target_dataset && 
            current_response_variable == target_response_variable) {
          # Add the current element to the filtered list
          filtered_list[[length(filtered_list) + 1]] <- full_bLR_mod_list[[i]]
        } # closes if-clause for matching current list element
      } # closes 'if(current_predictor %in% target_predictor_variables)'
    } # closes 'if (is.list(full_bLR_mod_list[[i]])==T)' clause
  } # closes 'for (i in 1:length(full_bLR_mod_list))' loop
  # Save filtered_list object to console:
  return(filtered_list)
} # closes filter_bLR_mods() function

# Test new filter_bLR_mods() function on a single tree-dataset-response-predictor_set combination:
test_filtered_list <- filter_bLR_mods(full_bLR_mod_list=bLR_mod_list, target_tree="supertree1", target_dataset="all_data_bLR_raw", target_response_variable = "bin_fweb", target_predictor_variables = f_props_predictors)

# Run diagnostics to check that the function worked:
length(test_filtered_list)
for(i in 1:length(test_filtered_list)){
  print(test_filtered_list[[i]]$input_tree)
  print(test_filtered_list[[i]]$input_dataset)
  print(test_filtered_list[[i]]$response_variable)
  print(test_filtered_list[[i]]$predictor_variable)
  print("-----------------------------------")
} # closes 'for(i in 1:length(test_filtered_list))' loop

# Test new filter_bLR_mods() function on another, more challenging tree-dataset-response-predictor_set combination:
test_filtered_list <- filter_bLR_mods(full_bLR_mod_list=bLR_mod_list, target_tree=tree_names[1], target_dataset=bLR_raw_dataset_names[1], target_response_variable = bin_forelimb_groupers[1], target_predictor_variables = f_mD3_proxp_predictors) 

# Run diagnostics to check that the function worked:
length(test_filtered_list)
for(i in 1:length(test_filtered_list)){
  print(test_filtered_list[[i]]$input_tree)
  print(test_filtered_list[[i]]$input_dataset)
  print(test_filtered_list[[i]]$response_variable)
  print(test_filtered_list[[i]]$predictor_variable)
  print("-----------------------------------")
} # closes 'for(i in 1:length(test_filtered_list))' loop

# Test new filter_bLR_mods() function on another challenging tree-dataset-response-predictor_set combination:
test_filtered_list <- filter_bLR_mods(full_bLR_mod_list=bLR_mod_list, target_tree=tree_names[1], target_dataset="archos_all_bLR_raw", target_response_variable = "bin_fweb", target_predictor_variables = f_mD3_proxp_predictors) 

# Run diagnostics to check that the function worked:
length(test_filtered_list) # This output should be empty since there are no non-flippered archosaur forelimbs in the dataset.

# This diagnostic test output should show that the test_filtered_list object contains two phybLR objects, each with the same tree, dataset, and response variable, and each with a different predictor variable in the designated target_predictor_variables set.

# Define a new bLR_mod_selector function to pick the best among multiple models for the same dataset-response_var-tree triad:
bLR_mod_selector <- function(filtered_props_predictor_mods, filtered_acr_predictor_mods, filtered_ungual_predictor_mods, filtered_proxp_predictor_mods){
  # Each of the arguments above is a list of models generated by the filter_bLR_mods function. The bLR_mod_selector function picks the best single function from each of these lists to output a new single list of four best-performing models--one from each of the four filtered lists provided as inputs. This is done by picking that phybLR model from each filtered list whose corresponding cROCC has the highest AUC.
  
  # Rename objects within function to save space:
  props_mods <- filtered_props_predictor_mods
  acr_mods <- filtered_acr_predictor_mods
  ungp_mods <- filtered_ungual_predictor_mods
  proxp_mods <- filtered_proxp_predictor_mods
  
  # Initialize output object:
  selected_bLR_mods <- as.list(rep(NA, 4))
  
  # Initialize index-keeper to remove defunct model sets:
  defunct_index <- rep(NA, 4)
  
  # Pick the most accurate phybLR model from filtered_props_predictor_mods:
    # NOTE: The next 15 lines of code were written with the help of chatGPT.
      # Initialize object to track highest AUC among models:
      max_auc_value <- 0  # initialize with a very low value
      if( (length(props_mods)>0) & !(is.null(props_mods)) ){ 
        # If there is a props model for this combo...
        defunct_index[1] <- 0 # ...then this is NOT a defunct model series.
      # Run a for-loop to find the element with the highest AUC:
      for(i in 1:length(props_mods)) {
        # Check if the current element has a valid AUC value (it should):
        if(!is.null(props_mods[[i]]$AUC_mean)) {
          # Compare current element's AUC to the max_auc_value
          if(props_mods[[i]]$AUC_mean > max_auc_value) {
            max_auc_value <- props_mods[[i]]$AUC_mean  # updates max AUC value
            selected_bLR_mods[[1]] <- props_mods[[i]] # stores selected model
          } # closes 'if(!is.null(props_mods[[i]]$AUC))' clause
        } # closes 'if(props_mods[[i]]$AUC_mean > max_auc_value)' clause
      } # This for-loop should populate selected_bLR_mods[[1]] with the most accurate phybLR model from the props_mods list.
      } else {
        defunct_index[1] <- 1 # ...then this IS a defunct model series.
      } # closes if / else if '(length(props_mods)>0)' clause pair
      
  # Pick the most accurate phybLR model from filtered_acr_predictor_mods:
      # Re-initialize object to track highest AUC among models:
      max_auc_value <- 0  # initialize with a very low value
      if( (length(acr_mods)>0) & !(is.null(acr_mods)) ){ 
        # If there even is an acr model for this combo...
        defunct_index[2] <- 0 # ...then this is NOT a defunct model series.
        # Run a for-loop to find the element with the highest AUC:
      # Run a for-loop to find the element with the highest AUC:
      for(i in 1:length(acr_mods)) {
        # Check if the current element has a valid AUC value (it should):
        if(!is.null(acr_mods[[i]]$AUC_mean)) {
          # Compare current element's AUC to the max_auc_value
          if(acr_mods[[i]]$AUC_mean > max_auc_value) {
            max_auc_value <- acr_mods[[i]]$AUC_mean # updates max AUC value
            selected_bLR_mods[[2]] <- acr_mods[[i]] # stores selected model
          } # closes 'if(!is.null(props_mods[[i]]$AUC))' clause
        } # closes 'if(props_mods[[i]]$AUC_mean > max_auc_value)' clause
      } # This for-loop should populate selected_bLR_mods[[2]] with the most accurate phybLR model from the acr_mods list.
      } else {
        defunct_index[2] <- 1 # ...then this IS a defunct model series.
      } # closes if / else if (length(mods_obj)>0) clause pair

  # Pick the most accurate phybLR model from filtered_ungp_predictor_mods:
      # Re-initialize object to track highest AUC among models:
      max_auc_value <- 0  # initialize with a very low value
      if( (length(ungp_mods)>0) & !(is.null(ungp_mods)) ){ # If there is an ungp model for this combo...
        defunct_index[3] <- 0 # ...then this is NOT a defunct model series.
      # Run a for-loop to find the element with the highest AUC:
      for(i in 1:length(ungp_mods)) {
        # Check if the current element has a valid AUC value (it should):
        if(!is.null(ungp_mods[[i]]$AUC_mean)) {
          # Compare current element's AUC to the max_auc_value
          if(ungp_mods[[i]]$AUC_mean > max_auc_value) {
            max_auc_value <- ungp_mods[[i]]$AUC_mean # updates max AUC value
            selected_bLR_mods[[3]] <- ungp_mods[[i]] # stores selected model
          } # closes 'if(!is.null(props_mods[[i]]$AUC))' clause
        } # closes 'if(props_mods[[i]]$AUC_mean > max_auc_value)' clause
      } # This for-loop should populate selected_bLR_mods[[3] with the most accurate phybLR model from the ungp_mods list.
      } else {
        defunct_index[3] <- 1 # ...then this IS a defunct model series.
      } # closes if / else if (length(mods_obj)>0) clause pair

  # Pick the most accurate phybLR model from filtered_proxp_predictor_mods:
      # Re-initialize object to track highest AUC among models:
      max_auc_value <- 0  # initialize with a very low value
      # Run a for-loop to find the element with the highest AUC
      if( (length(proxp_mods)>0) & !(is.null(proxp_mods)) ){ 
        # If there is a proxp model for this combo...
        defunct_index[4] <- 0 # ...then this is NOT a defunct model series.
      for(i in 1:length(proxp_mods)) {
        # Check if the current element has a valid AUC value (it should):
        if(!is.null(proxp_mods[[i]]$AUC_mean)) {
          # Compare current element's AUC to the max_auc_value
          if(proxp_mods[[i]]$AUC_mean > max_auc_value) {
            max_auc_value <- proxp_mods[[i]]$AUC_mean # updates max AUC value
            selected_bLR_mods[[4]] <- proxp_mods[[i]] # stores selected model
          } # closes 'if(!is.null(props_mods[[i]]$AUC))' clause
        } # closes 'if(props_mods[[i]]$AUC_mean > max_auc_value)' clause
      } # This for-loop should populate selected_bLR_mods[[4]] with the most accurate phybLR model from the proxp_mods list. 
      } else {
        defunct_index[4] <- 1 # ...then this IS a defunct model series.
      } # closes if / else if (length(mods_obj)>0) clause pair
      
  # Remove any defunct models from the series:
      selected_bLR_mods_screened <- selected_bLR_mods[defunct_index != 1]
  # Output the screened selected_bLR_mods list to the working environment:
  return(selected_bLR_mods_screened)
} # closes bLR_mod_selector function
cat("\n","PROGRESS REP: chunk [S5.4.03] complete; starting next chunk..","\n")
#*-----**-----*

#*-----**{ [S5.4.04] Use new functions to plot best phybLR cROCCs for each series of morphometric variables. }*
# Now we can apply the new filter_bLR_mods, bLR_mod_selector, and phycROCCs functions to generate a single ROC plot (showing the four highest-AUC models across data series) for each tree-dataset-response-predictor_set combination. 

# Make one ROC plot for each *forelimb* tree-dataset-response-predictors combo:
for(t in 1:length(tree_names)){
  for(d in 1:length(bLR_raw_dataset_names)){
    for(b in 1:length(bin_forelimb_groupers)){
      cat("\n", "CURRENT TREE:", t, "\n", "CURRENT DATASET NUMBER:", d, "\n", "CURRENT DATASET NAME:", bLR_raw_dataset_names[d], "\n", "CURRENT RESPONSE_VAR NUMBER:", b, "\n", "CURRENT RESPONSE_VAR NAME:", bin_forelimb_groupers[b], "\n")
      cat("\n","Filtering models for current tree-dataset-response_var combination...", "\n")
      
      # FIRST, for forelimb region proportions: Run filter_bLR_mods function on current tree-dataset-variable combination to get a list of bLR_mods specific to one set of predictor vars:
      f_props_mods <- filter_bLR_mods(full_bLR_mod_list=bLR_mod_list, target_tree=tree_names[t], target_dataset=bLR_raw_dataset_names[d], target_response_variable = bin_forelimb_groupers[b], target_predictor_variables = f_props_predictors)
      
      # SECOND, for forelimb acropod dimensions: Run filter_bLR_mods function on current tree-dataset-variable combination to get a list of bLR_mods specific to one set of predictor vars:
      f_acr_mods <- filter_bLR_mods(full_bLR_mod_list=bLR_mod_list, target_tree=tree_names[t], target_dataset=bLR_raw_dataset_names[d], target_response_variable = bin_forelimb_groupers[b], target_predictor_variables = f_acr_predictors)
      
      # THIRD, for mD3 ungual dimensions: # Run filter_bLR_mods function on current tree-dataset-variable combination to get a list of bLR_mods specific to one set of predictor vars:
      f_ungp_mods <- filter_bLR_mods(full_bLR_mod_list=bLR_mod_list, target_tree=tree_names[t], target_dataset=bLR_raw_dataset_names[d], target_response_variable = bin_forelimb_groupers[b], target_predictor_variables = f_mD3_ungp_predictors)
      
      # FOURTH, for mD3 proximal phalanx dimensions: # Run filter_bLR_mods function on current tree-dataset-variable combination to get a list of bLR_mods specific to one set of predictor vars:
      f_proxp_mods <- filter_bLR_mods(full_bLR_mod_list=bLR_mod_list, target_tree=tree_names[t], target_dataset=bLR_raw_dataset_names[d], target_response_variable = bin_forelimb_groupers[b], target_predictor_variables = f_mD3_proxp_predictors)
      
      # Report completion of filter_bLR_mods calls for this iteration:
      cat("\n", "Filtering complete for this iteration. Now selecting best models...", "\n")
      
      #### PORTION TO DELETE ####
      cat("\n", "length(f_props_mods):", length(f_props_mods), ", length(f_acr_mods):", length(f_acr_mods), ", length(f_ungp_mods):", length(f_ungp_mods), ", length(f_proxp_mods):", length(f_proxp_mods), "\n")
      #### PORTION TO DELETE ####
      
      # NEXT: Check to make sure there are models with the selected criteria:
      if( length(f_props_mods)==0 | length(f_acr_mods)==0 | length(f_ungp_mods)==0 | length(f_proxp_mods)==0 ) {
        cat("\n", "WARNING: phycROCCs has skipped this iteration because at least one set of models failed to meet the selection criteria.", "\n", "It may be that there are no data for at least one of the levels of the current binary response variable in the selected dataset.", "\n")
      } else {
        # IF IT DOES, THEN: Run bLR_mod_selector on the list of filtered models for the current tree-dataset-response_variable combination to get a single list of the selected phybLR model for each data series:
        selected_forelimb_mods <- bLR_mod_selector(filtered_props_predictor_mods = f_props_mods, filtered_acr_predictor_mods = f_acr_mods, filtered_ungual_predictor_mods = f_ungp_mods, filtered_proxp_predictor_mods = f_proxp_mods)
        
        # Report completion of model selection calls for this iteration:
        cat("\n", "Model selection complete for this iteration. Now plotting cROCCs...", "\n")
        
        # THEN: Save an ROC plot of the four selected phybLR models for this tree-dataset-response_variable combination:
        # Define current list of predictors in correct order:
        selected_mod_predictors <- rep(NA, length(selected_forelimb_mods))
        for(f in 1:length(selected_mod_predictors)){
          selected_mod_predictors[f]<-selected_forelimb_mods[[f]]$predictor_variable}
        # Define output_name string for phycROCCs() function:
        current_output_name <- paste("FORELIMBS_", bLR_raw_dataset_names[d], bin_forelimb_groupers[b], "for", tree_names[t], sep="_")
        # Run phycROCCs() function on current list of selected bLR models:
        phycROCCs(phybLR_mod_list=selected_forelimb_mods, ROC_curve_names=selected_mod_predictors, predictor_set_names=selected_mod_predictors, output_name = current_output_name, ROC_colors = roc_colors_vec, plot_cROCC_CIs = T)
        
        # FINALLY: Report for-loop completion for this iteration:
        cat("\n", "Selected ROC curves saved for", current_output_name, "\n", "--------------------------------------------------------------------------")
      } # closes 'if-else' clause
    } # closes 'for(b in 1:length(bin_forelimb_groupers))' loop
  } # closes 'for(d in 1:length(bLR_raw_dataset_names))' loop
} # closes 'for(t in 1:length(tree_names))' loop

# We have just generated one ROC plot for every combination of tree, dataset, forelimb response variable, and forelimb predictor variable series. Next, we will do the same for every combination of tree, dataset, hindlimb response variable, and hindlimb predictor variable series.

# Make one ROC plot for each *hindlimb* tree-dataset-response-predictors combo:
for(t in 1:length(tree_names)){
  for(d in 1:length(bLR_raw_dataset_names)){
    for(b in 1:length(bin_hindlimb_groupers)){
      cat("\n", "CURRENT TREE:", t, "\n", "CURRENT DATASET NUMBER:", d, "\n", "CURRENT DATASET NAME:", bLR_raw_dataset_names[d], "\n", "CURRENT RESPONSE_VAR NUMBER:", b, "\n", "CURRENT RESPONSE_VAR NAME:", bin_hindlimb_groupers[b], "\n")
      # FIRST, for hindlimb region proportions: Run filter_bLR_mods function on current tree-dataset-variable combination to get a list of bLR_mods specific to one set of predictor vars:
      h_props_mods <- filter_bLR_mods(full_bLR_mod_list=bLR_mod_list, target_tree=tree_names[t], target_dataset=bLR_raw_dataset_names[d], target_response_variable = bin_hindlimb_groupers[b], target_predictor_variables = h_props_predictors)
      
      # SECOND, for hindlimb acropod dimensions: Run filter_bLR_mods function on current tree-dataset-variable combination to get a list of bLR_mods specific to one set of predictor vars:
      h_acr_mods <- filter_bLR_mods(full_bLR_mod_list=bLR_mod_list, target_tree=tree_names[t], target_dataset=bLR_raw_dataset_names[d], target_response_variable = bin_hindlimb_groupers[b], target_predictor_variables = h_acr_predictors)
      
      # THIRD, for pD3 ungual dimensions: # Run filter_bLR_mods function on current tree-dataset-variable combination to get a list of bLR_mods specific to one set of predictor vars:
      h_ungp_mods <- filter_bLR_mods(full_bLR_mod_list=bLR_mod_list, target_tree=tree_names[t], target_dataset=bLR_raw_dataset_names[d], target_response_variable = bin_hindlimb_groupers[b], target_predictor_variables = h_pD3_ungp_predictors)
      
      # FOURTH, for pD3 proximal phalanx dimensions: # Run filter_bLR_mods function on current tree-dataset-variable combination to get a list of bLR_mods specific to one set of predictor vars:
      h_proxp_mods <- filter_bLR_mods(full_bLR_mod_list=bLR_mod_list, target_tree=tree_names[t], target_dataset=bLR_raw_dataset_names[d], target_response_variable = bin_hindlimb_groupers[b], target_predictor_variables = h_pD3_proxp_predictors)
      
      # NEXT: Check to make sure there are models with the selected criteria:
      if( length(h_props_mods)==0 | length(h_acr_mods)==0 | length(h_ungp_mods)==0 | length(h_proxp_mods)==0 ) {
        cat("\n", "WARNING: phycROCCs has skipped this iteration because at least one set of models failed to meet the selection criteria.", "\n", "It may be that there are no data for at least one of the levels of the current binary response variable in the selected dataset.", "\n")
      } else {
        # IF IT DOES, THEN: Run bLR_mod_selector on the list of filtered models for the current tree-dataset-response_variable combination to get a single list of the selected phybLR model for each data series:
        selected_hindlimb_mods <- bLR_mod_selector(filtered_props_predictor_mods = h_props_mods, filtered_acr_predictor_mods = h_acr_mods, filtered_ungual_predictor_mods = h_ungp_mods, filtered_proxp_predictor_mods = h_proxp_mods)
        
        # THEN: Save an ROC plot of the four selected phybLR models for this tree-dataset-response_variable combination:
        # Define current list of predictors in correct order:
        selected_mod_predictors <- rep(NA, length(selected_hindlimb_mods))
        for(f in 1:length(selected_mod_predictors)){
          selected_mod_predictors[f]<-selected_hindlimb_mods[[f]]$predictor_variable}
        # Define output_name string for phycROCCs() function:
        current_output_name <- paste("HINDLIMBS_", bLR_raw_dataset_names[d], bin_hindlimb_groupers[b], "for", tree_names[t], sep="_")
        
        # Run phycROCCs() function on current list of selected bLR models:
        phycROCCs(phybLR_mod_list=selected_hindlimb_mods, ROC_curve_names=selected_mod_predictors, predictor_set_names=selected_mod_predictors, output_name = current_output_name, ROC_colors = roc_colors_vec, plot_cROCC_CIs = T)
        
        # FINALLY: Report for-loop progress:
        cat("\n", "Selected ROC curves saved for", current_output_name, "\n", "--------------------------------------------------------------------------")
      } # closes 'if else' clause
    } # closes 'for(b in 1:length(bin_hindlimb_groupers))' loop
  } # closes 'for(d in 1:length(bLR_raw_dataset_names))' loop
} # closes 'for(t in 1:length(tree_names))' loop

cat("\n","PROGRESS REP: chunk [S5.4.04] complete; starting next chunk..","\n")
#*-----**-----*

# In the chunk above, we just created one set of ROC plots for every series of forelimb and hindlimb measurements for every tree-dataset-variable combination in our pipeline. Once this script has run, you will need to manually go through these and determine which phybLR models are worth using to reconstruct the evolutionary histories of certain phenotypes in script_S5.

# NOTE ON PROCEEDING: Before you proceed, please go through the phybLR subdirectory, review the output ROC plots, and manually select the most informative phybLR models to use for reconstructing ancient phenotypes. In our study, we found that relative hand length (f_AZR) and relative foot length (h_AZR) made by far the most accurate predictions for the following binary response variables: eco1_v1, eco1_v3, bin_fflip, and bin_hflip. We therefore opted to use those predictor variables to reconstruct the tip and node states for those response variables on each of our trees. However, depending on your own research interests, you may want to reconstruct a different binary phenotype from our dataset using different morphometric predictive variables. If so, you will need to modify the code below accordingly.
#------------------------------------------------------------------------------

#**"-------------------------------------------------------------------------"*
#**"- ### [S5.5] PREDICT TIP & NODE PHENOTYPES W/ BEST PHYBLR MODEL(s). ### -"*
#**"-------------------------------------------------------------------------"*

#*-----**{ [S5.5.01] Write tipPreds() function to predict tip phenotypes.}*
# Now that we have tested and compared the predictive performance of all our phybLR models in ROC space, we will use the best-performing models to predict the probabilities for predictable binary phenotypes in contentious extinct tip taxa and all the internal nodes of our trees. We will do this with two original functions. The first function, tipPreds(), will use a given phybLR model and its associated tree and dataset to predict the phenotype for the associated binary response variable of every tree tip, by testing whether its predicted probability of having the phenotype is significantly higher than the phybLR bestmod classification threshold. The second function, nodePreds(), will be a wrapper function that uses the asr_independent_contrasts() function in the castor package to perform ancestral state reconstructions for every internal node in the given tree. In the chunk below, we define the tipPreds() function.

tipPreds <- function(dataset, dataset_name, response, predictor, phybLR_obj, tree, tree_name){
  # FUNCTION ARGUMENTS:
  # dataset: dataset to start with (data frame)--should be downstream of the bLRify() function but not the rm.NAs() or reorder.data() functions.
  # dataset_name: chr string giving name of dataset for output files 
  # response: name of response variable (chr string)
  # predictor: name of predictor variable (chr string)
  # phybLR_obj: output object from phybLR() function--gives the best-performing phybLR model used to get predProbs for contentious taxa
  # tree: object of class "phylo" for performing ancestral state reconstructions using the asr_independent_contrasts() function
  # tree_name: chr string giving name of tree for output files
  
  cat("\n", "Initializing data environment and calling phybLR model...", "\n")
  # Define predictor and response variables within function:
  dataset$predictor <- dataset[, predictor]
  dataset$response <- dataset[, response]
  
  # Define palette for output plots within function:
  fill_colors <- grouping_var_palette[[response]]
  
  # Define base character string for output files:
  output_name <- paste0(dataset_name, "__", response, "__by__", predictor, "__for__", tree_name)
  
  # Create directory path for output files: 
  transitional_path <- paste0(output_path, "/BESTMOD_PREDICTIONS/")
  dir.create(path=transitional_path, showWarnings = TRUE, recursive = FALSE, mode = "0777")
  directory_path <- paste0(transitional_path, output_name, "/")
  dir.create(path=directory_path, showWarnings = TRUE, recursive = FALSE, mode = "0777")
  tipPreds_subdir <- paste0(directory_path, "tipPreds/")
  dir.create(path=tipPreds_subdir, showWarnings = TRUE, recursive = FALSE, mode = "0777")
  
  # Check that phybLR object is properly formatted:
  if(is.list(phybLR_obj)==F){stop("phybLR_obj input must be a list of class 'phybLR'.")}
  
  # Trim dataset to remove all rows with missing predictor variable data:
  ds_tr <- dataset[is.na(dataset$predictor)==F,]
  
  # Align data and tree:
  tree_tr <- trim.tree(tree=tree, trimmed_dataset=ds_tr, verbose=F)
  ds_ord <- reorder.data(trimmed_tree=tree_tr,trimmed_dataset=ds_tr,verbose=F)
  
  # Assign known probabilities (0 and 1) for unambiguous taxa:
  ds_ord$known_probs<-as.integer(ds_ord$response == levels(ds_tr$response)[2])
  
  # Call phybLR model within function:
  bLRmod <- phybLR_obj$phybLR_mod
  bestmod_threshold <- phybLR_obj$bestmod_threshold
  
  # Internally define shade_density() function for plotting annotated kernel density curves:
  shade_density <- function(data, value, histogram_title, ...) {
    # data: vector of values with which to generate density curve
    # value: limit below which to shade in
    # histogram_title: chr string passed on to 'main' arg of plot() function
    # Calculate the kernel density estimate:
    density_curve <- density(data, ...)
    # Plot the density curve:
    plot(density_curve, main=histogram_title, xlab='Predicted Probability', ylab='Kernel Density', lwd=3, xlim=c(0,1), cex.lab=1.5, cex.axis=2, cex.main=0.4)
    # Fill the entire area under the curve with yellow
    polygon(c(density_curve$x, rev(density_curve$x)), c(density_curve$y, rep(0, length(density_curve$y))), col = col.alpha("green", 0.2), border = NA)
    # Identify the x and y coordinates where x is less than or equal to the specified value
    x_coords <- density_curve$x[density_curve$x <= value]
    y_coords <- density_curve$y[density_curve$x <= value]
    # Add a polygon to shade the area under the curve
    polygon(c(x_coords, rev(x_coords)), c(y_coords, rep(0, length(y_coords))), col = col.alpha("red", 0.2), border = NA)
    # Add a vertical line at the specified value for reference
    abline(v = value, col = "red", lwd=1, lty='dashed')
    # Add another vertical line for the mean of the distribution:
    abline(v=mean(data), col='black', lwd=5, lty='dotted')
  } # closes shade_density() function
  
  cat("\n", "Predicting probabilities for", response, "phenotype in all contentious tip taxa...", "\n") 
  
  # Get phybLR model parameters:
  phybLR_mod_coef_means <- coef(bLRmod)
  phybLR_mod_coef_SDs <- bLRmod$sd
  
  # Initialize dataframe containing all predicted probability results:
  tipProbs_data <- as.data.frame(matrix(nrow=nrow(ds_ord), ncol=24))
  colnames(tipProbs_data) <- c("input_dataset", "input_tree", "response_variable", "predictor_variable", "bestmod_binomt_pval", "bestmod_TPR_mean", "bestmod_FPR_mean", "bestmod_Accuracy_mean", "|", "tip #", "tip_label", "prob_type", "knownProb", "tx_score_mean", "tx_score_sd", "predProb_mean", "predProb_sd", "bestmod_threshold", "ttest_tval", "ttest_df", "ttest_pval", "ttest_verdict", "has_phenotype", "final_verdict") # NOTE: The final_verdict is either the known prob (if there is one) or the verdict classification (0 or 1) based on the results of the t-test.
  
  # Populate this dataframe with predicted probability results for each tip:
  for(i in 1:nrow(tipProbs_data)){ # For each row in the output table...
    # Save the phybLR model inputs and metadata:
    tipProbs_data[i, "input_dataset"] <- phybLR_obj$input_dataset
    tipProbs_data[i, "input_tree"] <- phybLR_obj$input_tree
    tipProbs_data[i, "response_variable"] <- phybLR_obj$response_variable
    tipProbs_data[i, "predictor_variable"] <- phybLR_obj$predictor_variable
    tipProbs_data[i, "bestmod_binomt_pval"] <- phybLR_obj$bestmod_binomtest_pval
    tipProbs_data[i, "bestmod_TPR_mean"] <- phybLR_obj$bestmod_TPR_mean
    tipProbs_data[i, "bestmod_FPR_mean"] <- phybLR_obj$bestmod_FPR_mean
    tipProbs_data[i, "bestmod_Accuracy_mean"] <- phybLR_obj$bestmod_Accuracy
    tipProbs_data[i, "bestmod_threshold"] <- phybLR_obj$bestmod_threshold
    tipProbs_data[i, "|"] <- "|" # This column is purely aesthetic, to separate the previously tabulated phybLR metadata put here for quick reference (and available in the phybLR_metadata CSV saved above) from the tipPreds metadata that's currently being tabulated for the first time by this function.
    
    # Set tip number string to ensure output files are saved alphabetically:
    ival <- i
    if(i<10){tipnum <- paste("00", ival, sep="")}
    if(i>=10 & i<100){tipnum <- paste("0", ival, sep="")}
    if(i>100){tipnum <- as.character(ival)}
    
    tipProbs_data[i, "tip #"] <- tipnum
    tipProbs_data[i, "tip_label"] <- ds_ord$Tip_label[i]
    
    # Next, determine whether the probability of this tip is known or unknown.
    if(is.na(ds_ord$known_probs[i])==F){ # If it's known, set its probability type to "tip_known" and other predProb results to NA. 
      tipProbs_data$knownProb[i] <- ds_ord$known_probs[i]
      tipProbs_data$prob_type[i] <- "tip_known"
      tipProbs_data$tx_score_mean[i] <- NA
      tipProbs_data$tx_score_sd[i] <- NA
      tipProbs_data$predProb_mean[i] <- NA
      tipProbs_data$predProb_sd[i] <- NA
      tipProbs_data$ttest_tval[i] <- NA
      tipProbs_data$ttest_df[i] <- NA
      tipProbs_data$ttest_pval[i] <- NA
      tipProbs_data$ttest_verdict[i] <- NA
      if(ds_ord$known_probs[i]==1){
        tipProbs_data$has_phenotype[i] <- "Yes"} 
      if(ds_ord$known_probs[i]==0){
        tipProbs_data$has_phenotype[i] <- "No" }
      tipProbs_data$final_verdict[i] <- ds_ord$known_probs[i]
    } # closes 'if(is.na(ds_ord$known_probs[i])==F)' clause
    
    if(is.na(ds_ord$known_probs[i])==T){ # If it's unknown, then predict it using the best given phybLR model, and set its knownProb and probability type accordingly:
      cat("\n", "Predicting probability for tip", tipnum, ":", ds_ord$Tip_label[i], "...")
      tipProbs_data$knownProb[i] <- NA
      tipProbs_data$prob_type[i] <- "tip_predicted"
      
      # Compute and save the associated fixed linear predictor value for this:
      tx_score_mean <- ds_ord$predictor[i]*phybLR_mod_coef_means[2] + phybLR_mod_coef_means[1]
      tipProbs_data$tx_score_mean[i] <- tx_score_mean
      
      tx_score_sd <- ds_ord$predictor[i]*phybLR_mod_coef_SDs[2] + phybLR_mod_coef_SDs[1]
      tipProbs_data$tx_score_sd[i] <- tx_score_sd
      
      # Use logistic function to calculate associated mean predicted probability:
      # logistic function: predProb = 1 / (1 + e ^(−linear predictor))
      e <- exp(1) # outputs the number 'e'
      predProb_mean <- 1 / (1 + e^(-(tx_score_mean))) 
      tipProbs_data$predProb_mean[i] <- predProb_mean
      
      # Use logistic function to calculate associated standard deviation:
      # logistic function: predProb = 1 / (1 + e ^(−linear predictor))
      e <- exp(1) # outputs the number 'e'
      predProb_sd <- 1 / (1 + e^(-(tx_score_sd))) 
      tipProbs_data$predProb_sd[i] <- predProb_sd
      
      # Sample 10,000x from predicted probability distribution for this tip:
        # First, make a normal distrib. defined by the mean and sd given above:
      tip_prob_distribution <- normal_sample <- rnorm(1000, mean = tipProbs_data$predProb_mean[i], sd = tipProbs_data$predProb_sd[i]) 
        sample_probs_df <- data.frame("sample number"=seq(from=1, to=1000), "sample_values"=tip_prob_distribution) # Note: We assume above that the linear predictors and associated predicted probabilities are normally distributed about their mean values.
      
      # Test whether the mean predicted probability for this tip is significantly greater than the classifying threshold for the best given phybLR model:
      ttest_output <- t.test(x=tip_prob_distribution, mu=bestmod_threshold, alternative="greater", conf.level=0.99)
      tipProbs_data$ttest_tval[i] <- ttest_output$statistic[[1]]
      tipProbs_data$ttest_df[i] <- ttest_output$parameter[[1]]
      tipProbs_data$ttest_pval[i] <- ttest_output$p.value[[1]]
      if(tipProbs_data$ttest_pval[i]<=0.05){
        tipProbs_data$ttest_verdict[i] <- "Mean predicted probability for this tip is significantly higher than the predicted probability threshold given by the associated phybLR model for classifying phenotypes."
        tipProbs_data$has_phenotype[i] <- "Yes!"
        tipProbs_data$final_verdict[i] <- 1
      } else if (tipProbs_data$ttest_pval[i]>0.05){
        tipProbs_data$ttest_verdict[i] <- "Mean predicted probability for this tip is NOT significantly higher than the predicted probability threshold given by the associated phybLR model for classifying phenotypes."
        tipProbs_data$has_phenotype[i] <- "No"
        tipProbs_data$final_verdict[i] <- 0
      } # closes 'if(tipProbs_data$ttest_pval[i]<=0.05) { } else { }' clause
      
      # Define other geom object strings for plot calling:
      histogram_title_text <- paste0('Density curve of phybLR predProbs |', output_name, ' | for tip ', tipnum, ' : ', ds_ord$Tip_label[i])
      
      # Save the kernel density plot of predicted probability as a PNG file:
      png_file_path <- paste0(tipPreds_subdir, "/tip_", tipnum, "__", ds_ord$Tip_label[i], "__predProb_hist.png")
      png(filename = png_file_path, width=12, height=4, units="in", res=300)
      shade_density(data=sample_probs_df$sample_values, value=bestmod_threshold, histogram_title=histogram_title_text)
      #show(gg_hist_plot)
      dev.off()
    } # closes 'if(is.na(ds_ord$known_probs[i])==T)' clause
  } # closes 'for(i in 1:nrow(ds_ord))' loop
  
  # Report function progress:
  cat("\n", "The tipPreds function has finished making its predictions for this set of inputs.", "\n")
  
  # Save CSV file containing predicted probability results:
  write.csv(tipProbs_data, paste0(tipPreds_subdir, "/", "*tipProbs_table.csv"), row.names=FALSE)
  
  # Save tipPreds output object for the downstream nodePreds function:
  tipPreds_output <- list("dataset"=ds_ord, "dataset_name"=dataset_name, "predictor_variable"=predictor, "response_variable"=response, "tree"=tree_tr, "tree_name"=tree_name, "tipPreds"=tipProbs_data, "directory_path"=directory_path, "output_string"=output_name)
  class(tipPreds_output) <- "tipPreds"
  return(tipPreds_output)
} # closes tipPreds() function

# Test tipPreds() function on fflip predictor for all_data:
  # Run upstream functions:
    bLR <- bLRify(all_data)
    bLR_tr <- rm.NAs(dataset=bLR, categorical_var="bin_fflip", quantitative_var="f_AZR")
    phylo_tr <- trim.tree(tree=tree_variants[[1]], trimmed_dataset=bLR_tr)
    ds_ord <- reorder.data(trimmed_tree=phylo_tr, trimmed_dataset=bLR_tr)
    flip_bLR_output <- phybLR(dataset_ordered=ds_ord, dataset_name="***test_fflip_bLR_", response="bin_fflip", predictor="f_AZR", trimmed_tree=phylo_tr, tree_name = "phylo_tr", boot_num=100, CV_runs=3, btol_num=35, Kfold=3, crossval_reports=T, plot_td_ROCCs=T, plot_Youden=T, plot_McF=T, plot_CIs=T)
  # Test tipPreds() function:
    predictor_string <- "f_AZR"
    ds_string <- "test_ds"
    test_tipPreds_output <- tipPreds(dataset=bLR, dataset_name=ds_string, predictor=predictor_string, response="bin_fflip", tree=tree_variants[[1]], tree_name="test_tree", phybLR_obj=flip_bLR_output)
  # Test tipPreds() function again with DATA FRAMES BLR_MOD_LIST....:
    predictor_string <- "f_AZR"
    ds_string <- "test_ds"
    test_tipPreds_output <- tipPreds(dataset=bLR, dataset_name="all_BUT_archos_bLR_raw", predictor="f_AnAR", response="bin_eco1_v1", tree=tree_variants[[1]], tree_name="test_tree", phybLR_obj=bLR_mod_list[[1]])
  # We have now finished defining and testing the tipPreds() function.
cat("\n","PROGRESS REP.: chunk [S5.5.01] done; starting next chunk...","\n")
#*-----**-----*

#*-----**{ [S5.5.02] Define function to predict internal node phenotypes.}*
# In the previous chunk, we defined the tipPreds() function to predict binary phenotypes for all tips in a given tree, effectively setting all tips to either 0 or 1. In the present chunk, we define a nodePreds() function that will use the tipPreds() output, in tandem with the asr_independent_contrasts() function from the R package castor, to predict the phenotypes for all internal nodes in the given tree.

nodePreds <- function(tipPreds_object){
  # FUNCTION ARGUMENTS: tipPreds_object: an object of class "tipPreds", produced by the tipPreds() function
  
  # To begin, make sure that tipPreds_object is in the correct format:
  if(!(class(tipPreds_object)=="tipPreds")){stop("tipPreds_object input must be a list of class 'tipPreds'. You must generate this object with the tipPreds() function.")}
  
  # Define all key inputs within function:
  dataset <- tipPreds_object$dataset
  dataset_name <- tipPreds_object$dataset_name
  tree <- tipPreds_object$tree
  tree_name <- tipPreds_object$tree_name
  predictor <- tipPreds_object$predictor
  response <- tipPreds_object$response
  output_string <- tipPreds_object$output_string
  base_directory_path <- tipPreds_object$directory_path
  tipPreds_data <- tipPreds_object$tipPreds
  
  # Define palette for output plots within function:
  fill_colors <- grouping_var_palette[[response]]
  
  # Define directory path for output files: 
  nodePreds_subdir <- paste0(base_directory_path, "nodePreds/")
  dir.create(path=nodePreds_subdir, showWarnings = TRUE, recursive = FALSE, mode = "0777")
  
  cat("\n", "Reconstructing predicted probs for", response, "at each internal node...", "\n")
  
  # Remove singleton nodes from the tree:
    tree_collapsed <- collapse.singles(tree=tree, root.edge=T)
    #tree_collapsed <- tree # This is an optional alternative line for defining the tree that avoids collapsing singleton nodes.
  
  # Assign final probabilities to dataset:
    ds_fin <- dataset
    ds_fin$tip_phenotypes <- tipPreds_data$final_verdict
  
  # OPTIONAL TROUBLESHOOTING STEP: Confirm alignment of tree and dataset:
    # print(setdiff(rownames(dataset), tree$tip.label)); print(setdiff(tree$tip.label, rownames(dataset)))
  
  # Calculate ancestral predProb for each internal node:
    asr_output <- asr_independent_contrasts(tree=tree_collapsed, tip_states=ds_fin$tip_phenotypes, weighted=T, include_CI=F, check_input=F) # NOTE: Unfortunately, my R session always crashes when I set include_CI=T. I have no idea why. To avoid this, I have set include_CI=F. All tip values have confidence intervals of either 100% (for knownProbs) or 95% (for predProbs).
    ASRs <- asr_output$ancestral_states 
  
  # We now have ancestral states for all the internal nodes of the tree. Next, we will tabulate this data for every node and iterate over every node in the tree to create a pie chart of predicted probabilities for each one.
    cat("\n", "Saving predicted probability data for all nodes to CSV file...", "\n")
  
  # Get set of all nodes:
    all_nodes<-sort(unique(c(tree_collapsed$edge[,1],tree_collapsed$edge[,2])))
    cat("\n", "Total number of nodes in tree:", length(all_nodes)) 
  
  # Get all internal node numbers:
    internal_nodes <- sort(unique(tree_collapsed$edge[,1]))
    cat("\n","Total number of internal nodes in tree:",length(internal_nodes)) 
  
  # Filter out internal nodes to get tip numbers:
  tip_nodes <- all_nodes[!(all_nodes %in% internal_nodes)]
  cat("\n","Total number of external nodes (tips) in tree:", length(tip_nodes), "\n")
  
  # Initialize data frame containing predProb data for all nodes:
  prob1_lab <- paste("asr P(", levels(ds_fin$response)[2], ")", sep="")
  prob2_lab <- paste("asr P(", levels(ds_fin$response)[1], ")", sep="")
  ASR_table <- as.data.frame(matrix(nrow=length(internal_nodes), ncol=9))
  colnames(ASR_table) <- c("input_dataset", "input_tree", "response_variable", "predictor_variable", "node #", "tips defining this internal node (tip numbers)", "tips defining this internal node (tip labels)", prob1_lab, prob2_lab)
  
  # Internally define function get minimum set of tips uniquely specifying a given internal node:
  # NOTE: This internal function was generated with the help of chatGPT v. 3.5.
  get_min_tips_to_specify_node <- function(tree, internal_node) {
    # Check if the node is an internal node
    if(internal_node<=length(tree$tip.label)){
      stop("The provided node is not an internal node.")}
    
    # Get all descendants of the internal node
    all_descendants <- phangorn::Descendants(tree, internal_node, "tips")[[1]]
      # NOTE: The minimum set of tips to uniquely specify the node is just the descendant tips because all these tips are required to uniquely specify the internal node. To proceed, we can return the smallest set of tips from each child of the internal node.
    
    # Find the children of the internal node
    children <- phangorn::Children(tree, internal_node)
    
    min_tips <- c()
    for (child in children) {
      min_tips <- c(min_tips, phangorn::Descendants(tree, child, "tips")[[1]][1]) } # closes 'for (child in children)' loop
    
    # Compile the minimum tip numbers and associated tip labels:
    min_tip_labels <- tree$tip.label[min_tips]
    min_tip_labels_string <- paste0(min_tip_labels[[1]], ", ", min_tip_labels[[2]])
    min_tips_numbers_string <- paste0(min_tips[[1]], ", ", min_tips[[2]])
    
    # Save these minimum tip strings as a single output list:
    min_tips_output<-as.list(c(min_tips_numbers_string, min_tip_labels_string))
    return(min_tips_output)
  } # closes get_min_tips_to_specify_node() function
  
  # Populate ASR_table:
  for(n in 1:length(internal_nodes)){
    if(internal_nodes[n]<1000){nodenum<-paste("0", internal_nodes[n], sep="")}
    ASR_table[n, "input_dataset"] <- dataset_name
    ASR_table[n, "input_tree"] <- tree_name
    ASR_table[n, "response_variable"] <- response
    ASR_table[n, "predictor_variable"] <- predictor
    ASR_table[n, "node #"] <- nodenum
    min_tips_for_current_node <- get_min_tips_to_specify_node(tree=tree_collapsed, internal_node=internal_nodes[n])
    ASR_table[n, "tips defining this internal node (tip numbers)"] <- min_tips_for_current_node[[1]]
    ASR_table[n, "tips defining this internal node (tip labels)"] <- min_tips_for_current_node[[2]]
    ASR_table[n, prob1_lab] <- ASRs[n]
    ASR_table[n, prob2_lab] <- 1-ASRs[n]
  } # closes 'for(n in 1:length(internal_nodes))' loop
  
  # Save ASR_table to CSV file in the nodePreds subdirectory:
  write.csv(ASR_table, paste0(nodePreds_subdir, "/*ASR_predProbs_table.csv"), row.names=FALSE)
  
  # Make pie chart with mean predProb for each tip and node in tree:
  cat("\n", "Saving pie chart of the predicted probability for each node...", "\n")
  # Make input dataframe to populate for pie chart:
  pie_df <- matrix(data=NA, nrow=length(levels(ds_fin$response)), ncol=length(levels(ds_fin$response))) # For binary variables, the pie_df data frame should be a 2x2 matrix.
  # Set group labels (levels of response variable) within pie_df: 
  for(i in 1:nrow(pie_df)){pie_df[i, 1] <- levels(ds_fin$response)[i]}
  
  # Make lists to populate with pie_dfs (one per internal node):
  pie_node_list <- as.list(rep(NA, length(internal_nodes)))
  pie_node_geoms <- as.list(rep(NA, length(internal_nodes)))
  pie_node_names <- internal_nodes
  
  # Initialize output objects for internal nodes:
  for(i in 1:length(internal_nodes)){pie_node_list[[i]] <- pie_df}
  # Iterate over the internal nodes:
  for(i in 1:length(internal_nodes)){
    # Set node number string for output file:
    if(internal_nodes[i]<1000){nodenum<-paste("0", internal_nodes[i], sep="")}
    node_i <- paste("int_node", nodenum, sep="")
    
    # Make input dataframe for pie chart:
    pie_node_list[[i]] <- matrix(data=NA, nrow=length(levels(ds_fin$response)), ncol=length(levels(ds_fin$response))) 
    pie_node_list[[i]][, 1] <- levels(ds_fin$response)
    pie_node_list[[i]] <- as.data.frame(pie_node_list[[i]])
    pie_node_list[[i]][2,2] <- ASRs[i]
    pie_node_list[[i]][1,2] <- 1-ASRs[i]
    colnames(pie_node_list[[i]]) <- c("Group", "ASR_predProb")
    pie_node_list[[i]][, 2] <- as.numeric(pie_node_list[[i]][, 2])
    pie_node_list[[i]][, 1] <- fct_relevel(pie_node_list[[i]][, 1], levels(ds_fin$response))
    node_output_df <- pie_node_list[[i]]
    
    # Make ggplot pie chart for current tree node:
    gg_pie_full <- ggplot(data=pie_node_list[[i]], aes(x="", y=ASR_predProb, fill=Group)) + geom_bar(stat="identity", size=2, width=2, color="grey10", fill=fill_colors) + coord_polar("y", start=0) + theme_void() + ggtitle(paste( output_string, "|", "phybLR bestmod predProb for node", internal_nodes[i], sep=" ")) + theme(title = element_text(size=3)) # + scale_color_manual(values = fill_colors)
    
    pie_node_geoms[[i]] <- ggplot(data=pie_node_list[[i]], aes(x='', y=ASR_predProb, fill=Group)) + geom_bar(stat='identity', size=2, fill = fill_colors) + coord_polar('y', start=0) + theme_void() + theme(legend.position = "none") #+ scale_color_manual(values = fill_colors)
    
    # Save PNG of gg_pie plot to working directory:
    png(filename = paste0(nodePreds_subdir, "/", node_i, "__ASR_piechart.png"), width=4, height=4, units="in", res=300)
    show(gg_pie_full)
    dev.off()  
  } # closes 'for(i in 1:length(internal_nodes))' loop
  
  # Name pie_node_geoms:
    names(pie_node_geoms) <- pie_node_names
    
  cat("\n", "Saving high-resolution reference trees with nodes labeled...", "\n")
  
  # Save reference trees with labeled nodes to help place pie charts manually:
    # Pre-processed (original) tree:
      ggtree(tree, branch.length="none", size=0.1) + geom_text(aes(label=node), hjust=-.1, size=0.3, col='red') + geom_tiplab(size=0.2,hjust=-.2,col='grey30')
      ggsave(paste0(nodePreds_subdir, "/", "*", output_string, "__phylo_beforePruning.png"), dpi=1200, height=12, units="in")
      
    # Post-processed (trimmed, aligned, & collapsed) tree:
      ggtree(tree_collapsed, branch.length="none", size=0.1) + geom_text(aes(label=node), hjust=-.1, size=0.3, col='red') + geom_tiplab(size=0.2, hjust=-.2, col='grey30')
      ggsave(paste0(nodePreds_subdir, "/", "*", output_string, "__phylo_postPruning.png"), dpi=1200, height=12, units="in")
  
  cat("\n", "Saving high-resolution tree with overlaid pie charts...", "\n")
  # Now create post-processed tree with pie charts on each tip and node:
    # Convert node and tip lists to data frames:
    node_pies <- as.data.frame(matrix(nrow=length(pie_node_list), ncol=3))
    colnames(node_pies) <- c("Node #", levels(ds_fin$response))
    for(i in 1:length(pie_node_list)){
      node_pies[i, 1] <- pie_node_names[i] # node number
      node_pies[i, 2] <- pie_node_list[[i]][2,2] # ASR for P(yes)
      node_pies[i, 3] <- pie_node_list[[i]][1,2] # ASR for P(no)
    } # closes 'for(i in 1:length(pie_node_list))' loop
  
  gg_all_pies_on_tree<-ggtree(tree_collapsed,branch.length="none",size=0.1) + 
    geom_tiplab(geom="text",size=0.2,fontface="italic",hjust=-.1,col='grey30') + geom_inset(insets=pie_node_geoms, width=0.015, height=0.015, x="node")
  ggsave(paste0(nodePreds_subdir, "/", "*", output_string, "__phylo__postPruning_w_ASRs.png"), dpi=1200, height=12, units="in")
  
  # Report completion of function for a given set of inputs:
  cat("\n", "nodePreds() function complete!", "\n")
  cat("\n", "-------------------------------------------------------------------", "\n", "DATASET-VARIABLE-TREE COMBO:", dataset_name, "|", response, "by", predictor, "for", tree_name, ":", "\n", "• ds_fin object saved to environment as named output", "\n", "• Pie chart of predicted probs for each node and tip saved to working directory.", "\n", "• Reference tree with all node and tip numbers labeled also saved.", "\n", "• Maximal tree with overlaid ASR pie charts saved to working directory.",  "\n",  "-------------------------------------------------------------------", "\n")
  # Save ASR_table object as output:
  return(ASR_table)
} # closes nodePreds() function

# Test nodePreds() function on fflip predictor for all_data:
  test_nodePreds_output <- nodePreds(tipPreds_object = test_tipPreds_output)

# Test tipPreds() and nodePreds() functions on eco1v1 predictor for all_BUT_archos:
  # Run upstream functions:
    bLR <- bLRify(all_BUT_archos)
    bLR_tr <- rm.NAs(dataset=bLR, categorical_var="bin_eco1_v1", quantitative_var="f_AZR")
    phylo_tr <- trim.tree(tree=tree_variants[[1]], trimmed_dataset=bLR_tr)
    ds_ord <- reorder.data(trimmed_tree=phylo_tr, trimmed_dataset=bLR_tr)
    test_bLR_output <- phybLR(dataset_ordered=ds_ord, dataset_name="*test_ds", response="bin_eco1_v1", predictor="f_AZR", trimmed_tree=phylo_tr, tree_name = "phylo_tr", boot_num=2, btol_num=35, Kfold=3, crossval_reports=T, plot_td_ROCCs=T, plot_Youden=T, plot_McF=T, plot_CIs=T)
  # Test tipPreds() and nodePreds() functions:
    test_tipPreds_output <- tipPreds(dataset=bLR, dataset_name="*test_ds", predictor="f_AZR", response="bin_eco1_v1",tree=tree_variants[[1]], tree_name="test_tree", phybLR_obj=test_bLR_output)
    test_nodePreds_output <- nodePreds(tipPreds_object=test_tipPreds_output)
cat("\n","PROGRESS REP.: chunk [S5.5.02] done; starting next chunk...","\n")
#*-----**-----*

# In the previous few chunks of code, we defined two functions that could use a given phybLR() output object, in tandem with the correct associated tree, dataset, and predictor and response variables to predict the response variable phenotypes for all of a tree's tips and internal nodes. 

#*-----**{ [S5.5.03] Run nodePreds() and phyPreds() on key input combos.}*
# In the last few chunks of code, we defined two functions that could use a given phybLR() output object, in tandem with the correct associated tree, dataset, and predictor variables to predict the correct associated response variable phenotypes for all of a tree's tips and internal nodes. In addition, in script S4a, you generated a list of phybLR output objects from raw (un-transformed) datasets: the phybLR_output_list. Before starting the present script, you manually sorted through this list to select those phybLR models that were sufficiently accurate and interesting to use for making phenotype predictions. Assuming you have already chosen those phybLR models and adjusted the following code accordingly, you are ready to proceed. 

# In this chunk, I manually ran the nodePreds() and phyPreds() functions on all those models that I identified as being sufficiently accurate and interesting to use for making phenotype predictions. However, depending on the focus of your own study, you may want to revise this code to focus on other models or predictors. I found in my manual investigation of phybLR_output_list items that f_AZR and h_AZR were generally the best predictors of aquatic habits and flipper phenotypes, and that bin_fflip, bin_hflip, bin_eco1_v1, and bin_eco1_v3 were the most reliable and useful binary response variables for which to make phybLR predictions. I also found a clear difference in model accuracy with and without archosaurs included in the dataset, so I opted to run through the entire set of commands once with archosaurs and once without them. I also repeat the set of commands once for each supertree. I do this because, even though their differing topologies seem to have little effect on phybLR tip predictions (see outputs of section [S5.2] above), they do greatly affect ancestral state reconstructions. 

# Each set of commands below includes optional upstream function calls (including a phybLR() call), which we keep here for reference and reproducibility, though we would recommend skipping each phybLR() call to save computational time, and instead just isolating the appropriate phybLR_output_list item. Note that 'CV_runs' cannot be set to a smaller number (e.g., CV_runs=3) in the phybLR() calls below, because phybLR cross-validation runs not only to generate confidence intervals for cROCCs, but also set the optimal bestmod threshold probability for this curve, which is required to predict tip phenotypes and (by extension) ancestral state reconstructions.

ALL_BESTMOD_PREDICTIONS_list <- foreach(t=1:length(tree_variants), .combine='c', .packages = c("ggplot2", "forcats", "data.table", "stringr", "plyr", "dplyr", "car", "gridExtra", "ggimage", "vegan", "MASS", "pracma", "phytools", "ape", "ggtree", "pROC", "phylolm", "geiger", "phytools", "phangorn", "MESS", "castor")) %do% {
  # Create an empty list to store results for each iteration of i:
  output_list_per_t <- as.list(rep(NA, 8)) # one per prediction_set-dataset pair--so eight total per supertree.
  for(i in 1:length(output_list_per_t)){
    output_list_per_t[[i]] <- as.list(rep(NA, 5)) # Each list element will contain five objects--the name of the input tree, the name of the prediction set, the name of the input dataset, the tipPreds output, and the nodePreds output.
  } # closes 'for(i in 1:length(output_list_per_t))' loop
  
  # Define moniker string for current tree:
  tree_moniker <- names(tree_variants)[t]
  
  if(tree_variant_names[t]==tree_moniker){
  #-------------------------------------------------------------------------#
  ############ * FIRST SET OF PREDICTIONS: f_AZR -> bin_fflip * #############
  #-------------------------------------------------------------------------#
  cat("\n", "tipPreds and nodePreds commenced for the following combination:", "\n", "TREE:", tree_variant_names[t], "-", "PREDICTION SET: f_AZR -> bin_fflip")
  
  # Use f_AZR to predict bin_fflip phenotype for all_data:
    # Select best phybLR model (code written with help of chatGPT v. 3.5):
      bLR_output_element <- Filter(function(x) {
        x$input_dataset == "all_data_bLR_raw" && 
          identical(x$input_tree, tree_variant_names[t]) && 
          x$predictor_variable == "f_AZR" && 
          x$response_variable == "bin_fflip"
        }, bLR_mod_list)
      bLR_output <- bLR_output_element[[1]]
      
    # Define input dataset (required for tipPreds() function):
      bLR <- bLRify(all_data, verbose = F)
      
    # Optional code to generate bLR_output from scratch:
      # bLR_tr <- rm.NAs(dataset=bLR, categorical_var="bin_fflip", quantitative_var="f_AZR")
      # phylo_tr<-trim.tree(tree=tree_variants[[t]],trimmed_dataset=bLR_tr)
      # ds_ord <- reorder.data(trimmed_tree=phylo_tr, trimmed_dataset=bLR_tr)
      # bLR_output <- phybLR(dataset_ordered=ds_ord, dataset_name="all_data", response="bin_fflip", predictor="f_AZR", trimmed_tree=phylo_tr, tree_name=tree_moniker, boot_num=10000, btol_num=35, Kfold=3, CV_runs=1000, crossval_reports=F, plot_td_ROCCs=T, plot_Youden=F, plot_McF=F, plot_CIs=T, Firth_method="logistic_MPLE") # WARNING: Running this last line of code would take a very long time. I would not recommend running it interactively unless you reduce the boot_num and CV_runs values.
      
    # Run tipPreds() and nodePreds() functions:
      tipPreds_output <- tipPreds(dataset=bLR, dataset_name="all_data", predictor="f_AZR", response="bin_fflip",tree=tree_variants[[t]], tree_name=tree_moniker, phybLR_obj=bLR_output)
      nodePreds_output <- nodePreds(tipPreds_object=tipPreds_output)

    # Specify outputs and record inputs for this prediction set:
      output_list_per_t[[1]][[1]] <- bLR_output$input_tree
      output_list_per_t[[1]][[2]] <- "PREDICTION SET: f_AZR -> bin_fflip"
      output_list_per_t[[1]][[3]] <- bLR_output$input_dataset
      output_list_per_t[[1]][[4]] <- tipPreds_output
      output_list_per_t[[1]][[5]] <- nodePreds_output
      
  # Use f_AZR to predict bin_fflip phenotype for all_BUT_archos:
    # Select best phybLR model (code written with help of chatGPT v. 3.5):
      bLR_output_element <- Filter(function(x) {
        x$input_dataset == "all_BUT_archos_bLR_raw" && 
          identical(x$input_tree, tree_variant_names[t]) && 
          x$predictor_variable == "f_AZR" && 
          x$response_variable == "bin_fflip"
      }, bLR_mod_list)
      bLR_output <- bLR_output_element[[1]]
    # Define input dataset (required for tipPreds() function):
      bLR <- bLRify(all_BUT_archos, verbose = F)
      
    # Optional code to generate bLR_output from scratch:
      # bLR_tr <- rm.NAs(dataset=bLR, categorical_var="bin_fflip", quantitative_var="f_AZR")
      # phylo_tr <- trim.tree(tree=tree_variants[[t]], trimmed_dataset=bLR_tr)
      # ds_ord <- reorder.data(trimmed_tree=phylo_tr, trimmed_dataset=bLR_tr)
      # bLR_output <- phybLR(dataset_ordered=ds_ord, dataset_name="all_BUT_archos", response="bin_fflip", predictor="f_AZR", trimmed_tree=phylo_tr, tree_name=tree_moniker, boot_num=10000, btol_num=35, Kfold=3, CV_runs=1000, crossval_reports=F, plot_td_ROCCs=T, plot_Youden=F, plot_McF=F, plot_CIs=T, Firth_method="logistic_MPLE") # WARNING: Running this last line of code would take a very long time. I would not recommend running it interactively unless you reduce the boot_num and CV_runs values.
      
    # Run tipPreds() and nodePreds() functions:
      tipPreds_output <- tipPreds(dataset=bLR, dataset_name="all_BUT_archos", predictor="f_AZR", response="bin_fflip",tree=tree_variants[[t]], tree_name=tree_moniker, phybLR_obj=bLR_output)
      nodePreds_output <- nodePreds(tipPreds_object=tipPreds_output)
      
    # Specify outputs and record inputs for this prediction set:
      output_list_per_t[[2]][[1]] <- bLR_output$input_tree
      output_list_per_t[[2]][[2]] <- "PREDICTION SET: f_AZR -> bin_fflip"
      output_list_per_t[[2]][[3]] <- bLR_output$input_dataset
      output_list_per_t[[2]][[4]] <- tipPreds_output
      output_list_per_t[[2]][[5]] <- nodePreds_output
      
  #-------------------------------------------------------------------------#
  ############ * SECOND SET OF PREDICTIONS: h_AZR -> bin_hflip * ############
  #-------------------------------------------------------------------------#
  
  cat("\n", "tipPreds and nodePreds commenced for the following combination:", "\n", "TREE:", tree_variant_names[t], "-", "PREDICTION SET: h_AZR -> bin_hflip")
      
  # Use h_AZR to predict bin_hflip phenotype for all_data:
      # Select best phybLR model (code written with help of chatGPT v. 3.5):
      bLR_output_element <- Filter(function(x) {
          x$input_dataset == "all_data_bLR_raw" && 
            identical(x$input_tree, tree_variant_names[t]) && 
            x$predictor_variable == "h_AZR" && 
            x$response_variable == "bin_hflip"
        }, bLR_mod_list)
        bLR_output <- bLR_output_element[[1]]
      # Define input dataset (required for tipPreds() function):
        bLR <- bLRify(all_data, verbose = F)
        
      # Optional code to generate bLR_output from scratch:
        # bLR_tr <- rm.NAs(dataset=bLR, categorical_var="bin_hflip", quantitative_var="h_AZR")
        # phylo_tr <- trim.tree(tree=tree_variants[[t]],trimmed_dataset=bLR_tr)
        # ds_ord <- reorder.data(trimmed_tree=phylo_tr, trimmed_dataset=bLR_tr)
        # bLR_output <- phybLR(dataset_ordered=ds_ord, dataset_name="all_data", response="bin_hflip", predictor="h_AZR", trimmed_tree=phylo_tr, tree_name=tree_moniker, boot_num=10000, btol_num=35, Kfold=3, CV_runs=1000, crossval_reports=F, plot_td_ROCCs=T, plot_Youden=F, plot_McF=F, plot_CIs=T, Firth_method="logistic_MPLE") # WARNING: Running this last line of code would take a very long time. I would not recommend running it interactively unless you reduce the boot_num and CV_runs values.
        
      # Test tipPreds() and nodePreds() functions:
        tipPreds_output <- tipPreds(dataset=bLR, dataset_name="all_data", predictor="h_AZR", response="bin_hflip",tree=tree_variants[[t]], tree_name=tree_moniker, phybLR_obj=bLR_output)
        nodePreds_output <- nodePreds(tipPreds_object=tipPreds_output)
      
      # Specify outputs and record inputs for this prediction set:
        output_list_per_t[[3]][[1]] <- bLR_output$input_tree
        output_list_per_t[[3]][[2]] <- "PREDICTION SET: h_AZR -> bin_hflip"
        output_list_per_t[[3]][[3]] <- bLR_output$input_dataset
        output_list_per_t[[3]][[4]] <- tipPreds_output
        output_list_per_t[[3]][[5]] <- nodePreds_output
        
  # Use h_AZR to predict bin_hflip phenotype for all_BUT_archos:
      # Select best phybLR model (code written with help of chatGPT):
        bLR_output_element <- Filter(function(x) {
          x$input_dataset == "all_BUT_archos_bLR_raw" && 
            identical(x$input_tree, tree_variant_names[t]) && 
            x$predictor_variable == "h_AZR" && 
            x$response_variable == "bin_hflip"
        }, bLR_mod_list)
        bLR_output <- bLR_output_element[[1]]
      # Define input dataset (required for tipPreds() function):
        bLR <- bLRify(all_data, verbose = F)
        
      # Optional code to generate bLR_output from scratch:
        # bLR_tr <- rm.NAs(dataset=bLR, categorical_var="bin_hflip", quantitative_var="h_AZR")
        # phylo_tr <- trim.tree(tree=tree_variants[[t]],trimmed_dataset=bLR_tr)
        # ds_ord <- reorder.data(trimmed_tree=phylo_tr, trimmed_dataset=bLR_tr)
        # bLR_output <- phybLR(dataset_ordered=ds_ord, dataset_name="all_BUT_archos", response="bin_hflip", predictor="h_AZR", trimmed_tree=phylo_tr, tree_name=tree_moniker, boot_num=10000, btol_num=35, Kfold=3, CV_runs=1000, crossval_reports=F, plot_td_ROCCs=T, plot_Youden=F, plot_McF=F, plot_CIs=T, Firth_method="logistic_MPLE") # WARNING: Running this last line of code would take a very long time. I would not recommend running it interactively unless you reduce the boot_num and CV_runs values.
        
    # Run tipPreds() and nodePreds() functions:
      tipPreds_output <- tipPreds(dataset=bLR, dataset_name="all_BUT_archos", predictor="h_AZR", response="bin_hflip",tree=tree_variants[[t]], tree_name=tree_moniker, phybLR_obj=bLR_output)
      nodePreds_output <- nodePreds(tipPreds_object=tipPreds_output)
    
    # Specify outputs and record inputs for this prediction set:
      output_list_per_t[[4]][[1]] <- bLR_output$input_tree
      output_list_per_t[[4]][[2]] <- "PREDICTION SET: h_AZR -> bin_hflip"
      output_list_per_t[[4]][[3]] <- bLR_output$input_dataset
      output_list_per_t[[4]][[4]] <- tipPreds_output
      output_list_per_t[[4]][[5]] <- nodePreds_output
      
  #-------------------------------------------------------------------------#
  ########### * THIRD SET OF PREDICTIONS: f_AZR -> bin_eco1_v1 * ############
  #-------------------------------------------------------------------------#
  
  cat("\n", "tipPreds and nodePreds commenced for the following combination:", "\n", "TREE:", tree_variant_names[t], "-", "PREDICTION SET: f_AZR -> bin_eco1_v1")
      
  # Use f_AZR to predict eco1_v1 phenotype for all_data:
    # Select best phybLR model (code written with help of chatGPT v. 3.5):
      bLR_output_element <- Filter(function(x) {
        x$input_dataset == "all_data_bLR_raw" && 
          identical(x$input_tree, tree_variant_names[t]) && 
          x$predictor_variable == "f_AZR" && 
          x$response_variable == "bin_eco1_v1"
      }, bLR_mod_list)
      bLR_output <- bLR_output_element[[1]]
    # Define input dataset (required for tipPreds() function):
      bLR <- bLRify(all_data, verbose = F)
      
    # Optional code to generate bLR_output from scratch:
      # bLR_tr <- rm.NAs(dataset=bLR, categorical_var="bin_eco1_v1", quantitative_var="f_AZR")
      # phylo_tr <- trim.tree(tree=tree_variants[[t]], trimmed_dataset=bLR_tr)
      # ds_ord <- reorder.data(trimmed_tree=phylo_tr, trimmed_dataset=bLR_tr)
      # bLR_output <- phybLR(dataset_ordered=ds_ord, dataset_name="all_data", response="bin_eco1_v1", predictor="f_AZR", trimmed_tree=phylo_tr, tree_name=tree_moniker, boot_num=10000, btol_num=35, Kfold=3, CV_runs=1000, crossval_reports=F, plot_td_ROCCs=T, plot_Youden=F, plot_McF=F, plot_CIs=T, Firth_method="logistic_MPLE") # WARNING: Running this last line of code would take a very long time. I would not recommend running it interactively unless you reduce the boot_num and CV_runs values.
      
    # Run tipPreds() and nodePreds() functions:
      tipPreds_output <- tipPreds(dataset=bLR, dataset_name="all_data", predictor="f_AZR", response="bin_eco1_v1",tree=tree_variants[[t]], tree_name=tree_moniker, phybLR_obj=bLR_output)
      nodePreds_output <- nodePreds(tipPreds_object=tipPreds_output)
    
    # Specify outputs and record inputs for this prediction set:
      output_list_per_t[[5]][[1]] <- bLR_output$input_tree
      output_list_per_t[[5]][[2]] <- "PREDICTION SET: f_AZR -> bin_eco1_v1"
      output_list_per_t[[5]][[3]] <- bLR_output$input_dataset
      output_list_per_t[[5]][[4]] <- tipPreds_output
      output_list_per_t[[5]][[5]] <- nodePreds_output
      
  # Use f_AZR to predict eco1_v1 phenotype for all_BUT_archos:
    # Select best phybLR model (code written with help of chatGPT v. 3.5):
      bLR_output_element <- Filter(function(x) {
        x$input_dataset == "all_BUT_archos_bLR_raw" && 
          identical(x$input_tree, tree_variant_names[t]) && 
          x$predictor_variable == "f_AZR" && 
          x$response_variable == "bin_eco1_v1"
      }, bLR_mod_list)
      bLR_output <- bLR_output_element[[1]]
    # Define input dataset (required for tipPreds() function):
      bLR <- bLRify(all_data, verbose = F)
      
    # Optional code to generate bLR_output from scratch:
      # bLR_tr <- rm.NAs(dataset=bLR, categorical_var="bin_eco1_v1", quantitative_var="f_AZR")
      # phylo_tr <- trim.tree(tree=tree_variants[[t]], trimmed_dataset=bLR_tr)
      # ds_ord <- reorder.data(trimmed_tree=phylo_tr, trimmed_dataset=bLR_tr)
      # bLR_output <- phybLR(dataset_ordered=ds_ord, dataset_name="all_BUT_archos", response="bin_eco1_v1", predictor="f_AZR", trimmed_tree=phylo_tr, tree_name=tree_moniker, boot_num=10000, btol_num=35, Kfold=3, CV_runs=1000, crossval_reports=F, plot_td_ROCCs=T, plot_Youden=F, plot_McF=F, plot_CIs=T, Firth_method="logistic_MPLE") # WARNING: Running this last line of code would take a very long time. I would not recommend running it interactively unless you reduce the boot_num and CV_runs values.
      
    # Run tipPreds() and nodePreds() functions:
      tipPreds_output <- tipPreds(dataset=bLR, dataset_name="all_BUT_archos", predictor="f_AZR", response="bin_eco1_v1",tree=tree_variants[[t]], tree_name=tree_moniker, phybLR_obj=bLR_output)
      nodePreds_output <- nodePreds(tipPreds_object=tipPreds_output)
    # Specify outputs and record inputs for this prediction set:
      output_list_per_t[[6]][[1]] <- bLR_output$input_tree
      output_list_per_t[[6]][[2]] <- "PREDICTION SET: f_AZR -> bin_eco1_v1"
      output_list_per_t[[6]][[3]] <- bLR_output$input_dataset
      output_list_per_t[[6]][[4]] <- tipPreds_output
      output_list_per_t[[6]][[5]] <- nodePreds_output
      
  #-------------------------------------------------------------------------#
  ########### * FOURTH SET OF PREDICTIONS: f_AZR -> bin_eco1_v3 * ###########
  #-------------------------------------------------------------------------#
  
  cat("\n", "tipPreds and nodePreds commenced for the following combination:", "\n", "TREE:", tree_variant_names[t], "-", "PREDICTION SET: f_AZR -> bin_eco1_v3")
      
  # Use f_AZR to predict eco1_v3 phenotype for all_data:
    # Select best phybLR model (code written with help of chatGPT v. 3.5):
      bLR_output_element <- Filter(function(x) {
        x$input_dataset == "all_data_bLR_raw" && 
          identical(x$input_tree, tree_variant_names[t]) && 
          x$predictor_variable == "f_AZR" && 
          x$response_variable == "bin_eco1_v3"
      }, bLR_mod_list)
      bLR_output <- bLR_output_element[[1]]
    # Define input dataset (required for tipPreds() function):
      bLR <- bLRify(all_data, verbose = F)
      
    # Optional code to generate bLR_output from scratch:
      # bLR_tr <- rm.NAs(dataset=bLR, categorical_var="bin_eco1_v3", quantitative_var="f_AZR")
      # phylo_tr <- trim.tree(tree=tree_variants[[t]], trimmed_dataset=bLR_tr)
      # ds_ord <- reorder.data(trimmed_tree=phylo_tr, trimmed_dataset=bLR_tr)
      # bLR_output <- phybLR(dataset_ordered=ds_ord, dataset_name="all_data", response="bin_eco1_v3", predictor="f_AZR", trimmed_tree=phylo_tr, tree_name=tree_moniker, boot_num=10000, btol_num=35, Kfold=3, CV_runs=1000, crossval_reports=F, plot_td_ROCCs=T, plot_Youden=F, plot_McF=F, plot_CIs=T, Firth_method="logistic_MPLE") # WARNING: Running this last line of code would take a very long time. I would not recommend running it interactively unless you reduce the boot_num and CV_runs values.
      
    # Run tipPreds() and nodePreds() functions:
      tipPreds_output <- tipPreds(dataset=bLR, dataset_name="all_data", predictor="f_AZR", response="bin_eco1_v3",tree=tree_variants[[t]], tree_name=tree_moniker, phybLR_obj=bLR_output)
      nodePreds_output <- nodePreds(tipPreds_object=tipPreds_output)
  
    # Specify outputs and record inputs for this prediction set:
      output_list_per_t[[7]][[1]] <- bLR_output$input_tree
      output_list_per_t[[7]][[2]] <- "PREDICTION SET: f_AZR -> bin_eco1_v3"
      output_list_per_t[[7]][[3]] <- bLR_output$input_dataset
      output_list_per_t[[7]][[4]] <- tipPreds_output
      output_list_per_t[[7]][[5]] <- nodePreds_output
      
  # Use f_AZR to predict eco1_v3 phenotype for all_BUT_archos:
    # Select best phybLR model (code written with help of chatGPT v. 3.5):
      bLR_output_element <- Filter(function(x) {
        x$input_dataset == "all_BUT_archos_bLR_raw" && 
          identical(x$input_tree, tree_variant_names[t]) && 
          x$predictor_variable == "f_AZR" && 
          x$response_variable == "bin_eco1_v3"
      }, bLR_mod_list)
      bLR_output <- bLR_output_element[[1]]
    # Define input dataset (required for tipPreds() function):
      bLR <- bLRify(all_data, verbose = F)
      
    # Optional code to generate bLR_output from scratch:
      # bLR_tr <- rm.NAs(dataset=bLR, categorical_var="bin_eco1_v3", quantitative_var="f_AZR")
      # phylo_tr <- trim.tree(tree=tree_variants[[t]], trimmed_dataset=bLR_tr)
      # ds_ord <- reorder.data(trimmed_tree=phylo_tr, trimmed_dataset=bLR_tr)
      # bLR_output <- phybLR(dataset_ordered=ds_ord, dataset_name="all_BUT_archos", response="bin_eco1_v3", predictor="f_AZR", trimmed_tree=phylo_tr, tree_name=tree_moniker, boot_num=10000, btol_num=35, Kfold=3, CV_runs=1000, crossval_reports=F, plot_td_ROCCs=T, plot_Youden=F, plot_McF=F, plot_CIs=T, Firth_method="logistic_MPLE") # WARNING: Running this last line of code would take a very long time. I would not recommend running it interactively unless you reduce the boot_num and CV_runs values.
      
    # Run tipPreds() and nodePreds() functions:
      tipPreds_output <- tipPreds(dataset=bLR, dataset_name="all_BUT_archos", predictor="f_AZR", response="bin_eco1_v3",tree=tree_variants[[t]], tree_name=tree_moniker, phybLR_obj=bLR_output)
      nodePreds_output <- nodePreds(tipPreds_object=tipPreds_output)
      
    # Specify outputs and record inputs for this prediction set:
      output_list_per_t[[8]][[1]] <- bLR_output$input_tree
      output_list_per_t[[8]][[2]] <- "PREDICTION SET: f_AZR -> bin_eco1_v3"
      output_list_per_t[[8]][[3]] <- bLR_output$input_dataset
      output_list_per_t[[8]][[4]] <- tipPreds_output
      output_list_per_t[[8]][[5]] <- nodePreds_output
  } # closes 'if(tree_variant_names[t]==tree_moniker)' clause
  output_list_per_t # output from single foreach run and thus a single supertree (for foreach to combine for all eight trees)
  # When this foreach-loop is done, each of the prediction sets above will generate one set of predictions for the tips and nodes of each supertree considered in our analysis.
} # closes 'foreach(t=1:length(tree_variants)' loop
cat("\n","PROGRESS REP.: chunk [S5.5.03] done; starting next chunk...","\n")
#*-----**-----*

#*-----**{ [S5.5.04] Save output predictions to RData object and CSV. }*
# Save output_list_per_t file from previous chunk to its own output object:
save(ALL_BESTMOD_PREDICTIONS_list, file=paste0(output_path, "/BESTMOD_PREDICTIONS/ALL_BESTMOD_PREDICTIONS_OUTPUT_OBJ.RData")) 
# OPTIONAL LINE TO LOAD PREVIOUSLY SAVED ALL_BESTMOD_PREDICTIONS_list OBJECT: load(file=paste0(output_path, "/BESTMOD_PREDICTIONS/ALL_BESTMOD_PREDICTIONS_OUTPUT_OBJ.RData"))

# Save ALL_BESTMOD_PREDICTIONS_list content to separate tip & node data frames:
  # Initialize data frames:
ALL_TIP_PREDICTIONS_table <- data.frame() 
ALL_NODE_PREDICTIONS_table <- data.frame() 

for(i in 1:length(ALL_BESTMOD_PREDICTIONS_list)){
  # Combine all tipPreds output objects:
  ALL_TIP_PREDICTIONS_table <- rbind(ALL_TIP_PREDICTIONS_table, ALL_BESTMOD_PREDICTIONS_list[[i]][[4]][[7]])
  # Rename last two columns in each nodePreds dataframe so that they're all the same (this is required to make rbind work):
  names(ALL_BESTMOD_PREDICTIONS_list[[i]][[5]])[8] <- "asr P(HAS phenotype)"
  names(ALL_BESTMOD_PREDICTIONS_list[[i]][[5]])[9] <- "asr P(LACKS phenotype)"
  # Combine all nodePreds output objects:
  ALL_NODE_PREDICTIONS_table <- rbind(ALL_NODE_PREDICTIONS_table, ALL_BESTMOD_PREDICTIONS_list[[i]][[5]])
} # closes 'for(i in 1:length(ALL_BESTMOD_PREDICTIONS_list))' loop

# Add extra response_var column to node predictions table for ease of reference:
ALL_NODE_PREDICTIONS_table$'response variable' <-ALL_NODE_PREDICTIONS_table$response_variable

# Save the new data frames to CSV files within the BESTMOD_PREDICTIONS subdir:
write.csv(x = ALL_TIP_PREDICTIONS_table, file = paste0(output_path, "/BESTMOD_PREDICTIONS/*ALL_TIP_PREDICTIONS_table.csv"))

write.csv(x = ALL_NODE_PREDICTIONS_table, file = paste0(output_path, "/BESTMOD_PREDICTIONS/*ALL_NODE_PREDICTIONS_table.csv"))

cat("\n","PROGRESS REP.: chunk [S5.5.04] done; starting next chunk...","\n")
#*-----**-----*
# We have now completed the bulk of data analysis for this project, and that concludes our machine learning approach to reconstruct the evolutionary history of aquatic habits and flipper form in amniotes. Additional functions to perform multivariate machine learning methods (eg, phylogenetic discriminant analysis or make partial dependence plots) may be appended here in the future as needed. At this stage, you should comb through the output plots from the tipPreds() and nodePreds() loops above, and use them to investigate evolutionary trends in particular subclades of interest (e.g., Enaliosauria, Mosasauria, etc.). 

#**"-------------------------------------------------------------------------"*
#**"------ ### [S5.6] SAVE R ENVIRONMENT FOR DOWNSTREAM SCRIPT S5  ### ------"*
#**"-------------------------------------------------------------------------"*

# In the final section of script_S5, we will save the script's R environment and report how long the script took to run.

#*-----**{ [S5.6.01] Save current R environment to working directory. }*
save.image(file=paste0(wd_path, "/R_ENVIR_for_script_S5.RData"))
cat("\n","PROGRESS REP: chunk [S5.6.01] complete; starting next chunk..","\n")
#*-----**-----*

#*-----**{ [S5.6.02] Record the time required for script_S5 to run. }*
# Record running time metrics for entire script:
proctime_output <- proc.time()
elapsed_secs <- proctime_output[3]
elapsed_mins <- elapsed_secs/60
elapsed_hrs <- elapsed_mins/60
elapsed_days <- elapsed_hrs/24
cat("\n","PROGRESS REP: chunk [S5.6.02] complete; starting next chunk..","\n")
#*-----**-----*

#*-----**{ [S5.6.03] Record CPU usage for this script. }*
# The following code calculates CPU usage efficiency based on the following Stack Overflow thread: https://stackoverflow.com/questions/73217110/cpu-usage-measurement-and-calculation-in-r-using-proc-time.
CPU_number <- CPU_num # the number of CPUs used for the current script
CPU_time_used <- proctime_output[1] + proctime_output[2]
CPU_utilization <- (proctime_output[1] / CPU_time_used) * 100 / CPU_number
cat("\n","PROGRESS REP: chunk [S5.6.03] complete; starting next chunk..","\n")
#*-----**-----*

#*-----**{ [S5.6.04] Save time-elapsed and CPU metrics to a text file. }*
time_report_filepath <- paste0(wd_path, "/", "TIME_REPORT_for_script_S4c.txt")
sink(time_report_filepath)
cat("TIME TAKEN FOR ENTIRE SCRIPT TO RUN", "\n", elapsed_secs, "seconds", "\n", "\t", "=", elapsed_mins, "minutes", "\n", "\t", "=", elapsed_hrs, "hours", "\n", "\t", "=", elapsed_days, "days", "\n")
cat("\n", "CPU USAGE:", "\n", "\t", "Number of CPUs used:",  CPU_number, "\n", "\t", "CPU utilization efficiency:", CPU_utilization, "%", "\n")
sink(file=NULL) # A text file saving the time elapsed for the completion of this script has now been saved to the working directory.
cat("\n","PROGRESS REP: chunk [S5.6.04] is complete.", "\n", "script_S5 has finished running!..","\n")
#*-----**-----*

#------------------------------------------------------------------------------
#**"-------------------------------------------------------------------------"*
#**"--------------------- ### [[!]] CONCLUSION ### --------------------------"*
#**"-------------------------------------------------------------------------"*

cat("\n","PROGRESS REP.: All chunks done! Data analysis complete!","\n")

# The steps above conclude our data analysis pipeline for linear morphometric data. Taken as a whole, this pipeline interrogated and described the largest dataset of amniote-wide limb measurements to date within a phylogenetic framework, and applied a rigorous machine-learning approach to predict the aquatic affinities and soft-tissue limb phenotypes of many extinct amniote fossils and their common ancestors. Thank you for taking this coding journey and congrats on making it to the end! If you have any questions about this data analysis pipeline, please do not hesitate to email with any questions.

# Sincerely,
# Caleb Gordon
```
