## Supplementary R Scripts for "Limb proportions predict aquatic habits and soft-tissue flippers in extinct amniotes": script_S6.Rmd

````
---
title: "Geometric morphometric analyses for Gordon et al."
author: "Caleb Gordon"
date: "9/27/2020"
output: html_document
---

```{r setup, include=FALSE}
knitr::opts_chunk$set(echo = TRUE)
```

## R Markdown

Sixth R script (script_S6.Rmd) for Gordon et al., "Limb proportions predict aquatic habits and soft-tissue flippers in extinct amniotes."

NOTE ON READABILITY: This script contains many long, multi-line comments. To increase readability within Rstudio, go to Tools > Global Options, select the 'Code' tab, and check 'Soft-wrap R source files'. This will wrap all lines of code to match your personal GUI's margins, so that no lines of code run offscreen.

NOTE ON SCRIPT ORIGIN: Please note that this single Rmd file was compiled on October 3rd, 2024, from earlier R scripts originally written in 2020 and 2022, and some R functions may have since become deprecated. Please proceed with caution and, whenever possible, use R v. 4.2.0-foss-2020b to minimize unnecessary error messages.

NOTE ON SCRIPT TASKS: Herein, we will run three GM analyses: one for the forelimb zeugoautopod, one for the forelimb acropodium, and one for the ungual phalanx. In each, we will begin by converting image files to be landmarked so that they are in the correct format, define required grouping variables and data objects, optionally proceed with landmark using R geomorph's digitize2d function (alternatively, you can place landmarks in ImageJ), define sliders, perform a generalized procrustes superimposition of all raw landmark coordinates, perform a Principal Coordinates Analysis (PCA) and PERMANOVAs in PAST, and then compare individual PC scores among groups in R.

NOTE ON SCRIPT ORDER: The current script can be run independently of all other scripts (S1-S5) associated with this project, as they concern linear morphometric (LM) data, whereas the current script concerns geometric morphometric (GM) data. This script assumes that you have already defined a landmark scheme and compiled all the 2D images for each of your 3 GM analyses within a separate subfolder in your working directory. Please note that these subdirectories are not specified below (as each of the following GM analyses was previously run in separate scripts), so the code below will have to be revised to work on your three separate (zeugoautopod, acropod, & ungual) subdirectories.

### ----------------------- SET UP CODING ENVIRONMENT --------------------- ###

```{r Load relevant packages and set working directory.}
# For file conversions:
library(png)
library(jpeg)
library(tiff)

# For placing landmarks and performing geometric morphometric analyses:
library(geomorph)

# For downstream data analysis and visualization:
library(car)
library(ggplot2)

# Write path of your working directory to an object, to make later steps easier:
base_dir <- "/Users/caleb/Desktop/Flippers_KEY/Flipper GM files PRESSING/CURRENT SPECIMENS"
```

### ---------------- [GM ANALYSIS 1] ZEUGOAUTOPOD PLANFORM ---------------- ###

```{r Convert all image files to the correct format.}
# To work in geomorph, all image files must be in JPEG format. The following code tells you which files in your working directory are in which format, converts PNG and TIFF files to JPEGs.

list.files(base_dir, pattern='*jpg')
list.files(base_dir, pattern='*png')
list.files(base_dir, pattern='*tif')

png_set <- list.files(base_dir, pattern='*png')
tif_set <- list.files(base_dir, pattern = '*tif')

for (png_i in png_set){
  print(png_i)
  jpg_i <- gsub(".png", ".jpg", x = png_i)
  jpg_out_path <- paste0(base_dir, '/', jpg_i)
  png_in_path = paste0(base_dir, '/', png_i)
  img <- readPNG(png_in_path)
  writeJPEG(img, target = jpg_out_path, quality = 1)
}

for (tif_i in tif_set){
  print(tif_i)
  jpg_i <- gsub(".tif", ".jpg", x = tif_i)
  jpg_out_path <- paste0(base_dir, '/', jpg_i)
  tif_in_path = paste0(base_dir, '/', tif_i)
  img <- readTIFF(tif_in_path)
  writeJPEG(img, target = jpg_out_path, quality = 1)
}

# Note: If the tif code doesn't work (due to "incompatible type for 'RichTIFFIPTC'), just open the tif and export it manually to the working directory as a jpg.

# Now that you've made jpg versions of all your specimen image files, you can remove all former PNG and TIFF files in your working directory:
unlink("/Users/calebgordon/Desktop/220802_GM/*png")
unlink("/Users/calebgordon/Desktop/220802_GM/*tif")

# The code below verifies the absence of PNGs & TIFFs in the working directory:
list.files(base_dir, pattern='*png')
list.files(base_dir, pattern='*tif')
# It should say there are none ("character(0)").
```

```{r Define data objects and grouping variables.}
# Now that all of your image files are in the correct format, there's one more thing you should do before you place landmarks: define data objects that will make it much easier to automate landmark placement and downstream analysis for a large number of specimens.

# First, define a path designating a plain text file to which landmark coordinate data will be read. The following code defines this path and removes this file if it already exists (to avoid the formation of duplicates during multiple runs).
tps_path <- paste0(base_dir, '/raw_coordinates.txt')
#if (file.exists(tps_path)){file.remove(tps_path)}

# Second, define a shorthand object listing all of your input files:
specimens <- list.files(base_dir, pattern='*jpg')

# Thirdly, define grouping variables. In your geometric morphometric analyses, you'll probably want to bin specimens by group. To do this, it's best to define grouping variables that you can input into geomorph commands later.

# First, manually create a file grouping specimens in excel, and save it as a .csv file in the working directory (e.g., "GM_groups.csv"). You may need to open this CSV file in Numbers, manually delete blank rows and columns, and then overwrite the original CSV file. Then, once you have a correctly formatted CSV file, run the following code:
GM_groups <- read.csv("GM_groups.csv", header=T, row.names=NULL)

# Check out your data frame of grouping variables, to make sure it looks right:
dim(GM_groups)
head(GM_groups)

# Set your working directory one more time in preparation for the next step:
setwd(base_dir)
```

```{r Collect landmarks.}
# Now that we have defined useful data objects and grouping variables, we are ready to place landmarks.

# The code below 'uploads' the specimen image files in your working directory to the RStudio GUI, so that you can manually place landmarks on each file in R. In this line of code, you should also set the number of landmarks you plan to place, and the scale, if necessary. Once you run this line, you'll be prompted to place landmarks on each image sequentially. Don't worry if some of these landmarks are sliders. You'll define sliders retroactively in a bit.

# Note: You must copy this text into the console and run it there. Do not run this code from within the Rmd chunk or it will probably not work.
digitize2d(filelist=specimens, nlandmarks=31, scale=NULL, MultScale=FALSE, tpsfile=tps_path, verbose=FALSE)
```

Troubleshooting with the digitize2d() function above:
The digitize2d() function can be finicky sometimes. If it is not working, try these five things:
1. Do not run the code from within the Rmd chunk above. Instead, copy and paste it into the R console window and run it there.
2. Make sure your input filelist (eg, specimens) is a character vector. You can do this by typing "class(specimens)" in the console. A character vector is expected as input.
3. Set your working directory and make sure your files are correctly named, your filepath is specified correctly, and your files are in the working directory.
4. Make sure that none of your specimen image files begin with a numerical character (1234567890) or contain any spaces. Whenever I've had inexplicable issues with digitize2d() in the past, taking this step has fixed it.
5. If you've done (1) to (4) and it's still not working, it might be a readJPEG() issue. The readJPEG() command is a dependency of digitize2d() and apparently it can run into iOS permissions issues. Run this code to make sure the readJPEG() function isn't the issue:
jpgtest <- readJPEG("/Users/calebgordon/Desktop/Morphometric analyses 9.27.20/GM analysis - 9.27.20/Working_directory/Cryptocleidus_Sauropterygia.jpg", native = TRUE)

Once you place landmarks on all of your specimens, you can move on to the next step.

```{r Define sliders.}
# Read your raw_coordinates.txt file to an object to make downstream work easier:
raw_coordinates <- readland.tps("/Users/caleb/Desktop/Flippers_KEY/Flipper GM files PRESSING/CURRENT SPECIMENS/raw_coordinates.txt", specID = "ID") # This object contains the raw pixel coordinates of all of your landmarks for each specimen.

# The following code lets you manually (and retroactively) define sliders for all of your specimens in the RStudio GUI.
define.sliders(landmarks = raw_coordinates, nsliders = 20, surfsliders = NULL, write.file = TRUE)
#—————————Note: Do not run this command from within the chunk. Copy and paste this code into the R console window and run it there.
#—————————This command will have you define sliders for your first specimen, and then write them to the csv file "curveslide.csv". But the sliders are the same for all specimens, so this is ultimately all you need.

# Now write the slider data to an object for downstream use:
sliders <- as.matrix(read.csv("/Users/caleb/Desktop/Flippers_KEY/Flipper GM files PRESSING/CURRENT SPECIMENS/curveslide.csv", header=T)) 
#This file will be one of the direct inputs for Generalized Procrustes Analysis.
```

```{r Perform a Generalized Procrustes Superimposition}
# Perform generalized procrustes superimposition and write the output to a new object containing the Procrustes coordinates and metadata:
GM.gpa <- gpagen(raw_coordinates, curves = sliders, surfaces = NULL, PrinAxes = TRUE, max.iter = NULL, ProcD = FALSE, Proj = TRUE, print.progress = TRUE)

# GM.gpa$coords contains the Procrustes coordinates of your landmarks and sliding semi-landmarks. Check out the Procrustes coordinates of your specimens:
head(GM.gpa$coords)

# Now write the output of GM.gpa$coords to a tps file that PAST can read:
writeland.tps(GM.gpa$coords, "Procrustes_coordinates.txt", scale = NULL, specID = "ID")
```

Now, at this point, PAST and RStudio diverge. I prefer the PCA figure output of PAST to that of geomorph, so I would recommend doing PCA and PERMANOVAs in PAST. However, geomorph also has some useful tools that PAST doesn't (for example, warp grid comparisons). As a result, below are instructions for all of the downstream analyses I've found useful in PAST *and* RStudio.

```{r Perform PCA and PERMANOVAs in PAST}
# In the plain text file you just created ("Procrustes_coordinates.txt"), manually type in specimen names as "ID=specimen_name" (eg, "ID=Adelosaurus_sp.") below each stanza of Procrustes coordinates.This is necessary to put the file in tps format so that PAST can read it. Next, open this file in PAST3.

# Manually define your grouping variables, perhaps using the Groups csv you defined earlier.

# Run a PCA. Follow in-app instructions to make and modify convex hulls. Save elbow-plots and all other metadata for reference. Also save spline plots of low and high extreme scores for each relevant PC.
```

```{r Compare individual PC scores between groups in R.}
# The following code looks at PC score distributions for key PC scores from the PCA you ran above, and compares them between limb types and clades. 

# Set working directory:
setwd("/Users/calebgordon/Desktop/GM analysis - 9.29.20")
# Read data into an object:
PC_data <- read.csv("GM_forelimb_data.csv")
# Check out the data:
dim(PC_data); head(PC_data)

# Define new data frames and means for use in box plots:
  #For Clade comparisons:
  PC_data_Clade <- PC_data[, c(3,2,6,7,8,9)]
  dim(PC_data_Clade); head(PC_data_Clade)

  #For Limb Type comparisons:
  PC_data_LimbType <- PC_data[, c(3,4,6,7,8,9)]
  dim(PC_data_LimbType); head(PC_data_LimbType)

  #Defining a mean PC1/2 score metric:
  PC_1_2_mean <- rep(NA, 38)
  for (i in 1:38){PC_1_2_mean[i] <- mean(c(PC_data_LimbType$PC1[i], PC_data_LimbType$PC2[i]))}
  PC_data_LimbType_exp <- cbind(PC_data_LimbType[, c(1:6)], PC_1_2_mean)

  #Defining means for quantitative variables:
  PC1_mean_F <- mean(PC_data_LimbType$PC1[PC_data_LimbType$Forelimb.Type == "Flippered"])
  PC1_mean_Nf <- mean(PC_data_LimbType$PC1[PC_data_LimbType$Forelimb.Type == "Non-flippered"])
  PC1_mean_Amb <- mean(PC_data_LimbType$PC1[PC_data_LimbType$Forelimb.Type == "Ambiguous"])
  PC1_groupmeans <- c(PC1_mean_F, PC1_mean_Nf, PC1_mean_Amb)
  PC2_mean_F <- mean(PC_data_LimbType$PC2[PC_data_LimbType$Forelimb.Type == "Flippered"])
  PC2_mean_Nf <- mean(PC_data_LimbType$PC2[PC_data_LimbType$Forelimb.Type == "Non-flippered"])
  PC2_mean_Amb <- mean(PC_data_LimbType$PC2[PC_data_LimbType$Forelimb.Type == "Ambiguous"])
  PC1_2_mean__mean_F <- mean(PC_data_LimbType_exp$PC_1_2_mean[PC_data_LimbType_exp$Forelimb.Type == "Flippered"])
  PC1_2_mean__mean_Nf <- mean(PC_data_LimbType_exp$PC_1_2_mean[PC_data_LimbType_exp$Forelimb.Type == "Non-flippered"])
  PC1_2_mean__mean_Amb <- mean(PC_data_LimbType_exp$PC_1_2_mean[PC_data_LimbType_exp$Forelimb.Type == "Ambiguous"])

# Define a function to provide distributions of all quantitative variables with normal quantile plots to gage normality of distribution:
distrib.plot <- function(data, title = "title") {
  par(mfrow = c(2, length(names(data))), mar = c(2,0.1,2,1.5), oma = c(0,3,0, 0), cex.axis = 0.6, pch = 19, cex.main = 0.8, mgp=c(2, 0.5, 0));
  for (i in 1:length(names(data))){hist(data[, i], col = 'seagreen3', breaks = 5, main = NULL, xlab = NULL)}
  mtext("Frequency", side = 2, line = 47.5, cex = 0.7)
  mtext(title, side = 3, adj = 1.4, font = 2, cex = 0.8, line = 0.8); 
  for (i in 1:length(names(data))){
  qqPlot(data[, i], col = 'seagreen3', col.lines = 'grey40', lwd = 1, pch = 19, main = NULL, ylab = NULL, xlab = "", id = FALSE)
  title(names(data)[i], line = 1)}
  mtext("Normal quantiles", side = 2, line = 47.5, cex = 0.7)}

# Run function on untransformed variables:
distrib.plot(PC_data_LimbType_exp[, 3:7], title = "Distributions of PC scores for different reptile forelimbs")

# Run function on transformed variables:
distrib.plot(log(abs(PC_data_LimbType_exp[, 3:7])), title = "Distributions of log-transformed PC scores for different reptile forelimbs")

# Make box plots:
  #Comparing PC1 scores:
(bp_PC1_LimbType_untrans <- ggplot(PC_data_LimbType, aes(x=Forelimb.Type, y=PC1, fill=Forelimb.Type)) + geom_boxplot(width=0.8) + theme(legend.position="none") + scale_x_discrete(limits=c("Flippered", "Non-flippered", "Ambiguous"), name = "Limb Type", labels = c("Flippered", "Non-flippered", "Ambiguous")) + scale_fill_manual(values=c("grey", "skyblue", "salmon1")) + geom_point(x=1, y=PC1_mean_F, col = "yellow", size = 2) + geom_point(x=2, y=PC1_mean_Nf, col = "yellow", size = 2) + geom_point(x=3, y=PC1_mean_Amb, col = "yellow", size = 2) + theme(aspect.ratio = 4))

  #Comparing absolute-value transformed PC1 scores:
(bp_PC1_LimbType_trans <- ggplot(PC_data_LimbType, aes(x=Forelimb.Type, y=abs(PC1), fill=Forelimb.Type)) + geom_boxplot(width=0.8) + theme(legend.position="none") + scale_x_discrete(limits=c("Flippered", "Non-flippered", "Ambiguous"), name = "Limb Type", labels = c("Flippered", "Non-flippered", "Ambiguous")) + scale_fill_manual(values=c("grey", "skyblue", "salmon1")) + geom_point(x=1, y=abs(PC1_mean_F), col = "yellow", size = 2) + geom_point(x=2, y=abs(PC1_mean_Nf), col = "yellow", size = 2) + geom_point(x=3, y=abs(PC1_mean_Amb), col = "yellow", size = 2) + theme(aspect.ratio = 4))

  #Comparing PC2 scores:
(bp_PC2_LimbType_untrans <- ggplot(PC_data_LimbType, aes(x=Forelimb.Type, y=PC2, fill=Forelimb.Type)) + geom_boxplot(width=0.8) + theme(legend.position="none") + scale_x_discrete(limits=c("Flippered", "Non-flippered", "Ambiguous"), name = "Limb Type", labels = c("Flippered", "Non-flippered", "Ambiguous")) + scale_fill_manual(values=c("grey", "skyblue", "salmon1")) + geom_point(x=1, y=PC2_mean_F, col = "yellow", size = 2) + geom_point(x=2, y=PC2_mean_Nf, col = "yellow", size = 2) + geom_point(x=3, y=PC2_mean_Amb, col = "yellow", size = 2) + theme(aspect.ratio = 4))

  #Comparing PC3 scores:
(bp_PC3_LimbType_untrans <- ggplot(PC_data_LimbType, aes(x=Forelimb.Type, y=PC3, fill=Forelimb.Type)) + geom_boxplot(width=0.8) + theme(legend.position="none") + scale_x_discrete(limits=c("Flippered", "Non-flippered", "Ambiguous"), name = "Limb Type", labels = c("Flippered", "Non-flippered", "Ambiguous")) + scale_fill_manual(values=c("grey", "skyblue", "salmon1")) + theme(aspect.ratio = 4))

  #Comparing mean PC1&2 scores (a PC1/2 score metric):
(bp_PC3_LimbType_untrans <- ggplot(PC_data_LimbType_exp, aes(x=Forelimb.Type, y=PC_1_2_mean, fill=Forelimb.Type)) + geom_boxplot(width=0.8) + theme(legend.position="none") + scale_x_discrete(limits=c("Flippered", "Non-flippered", "Ambiguous"), name = "Limb Type", labels = c("Flippered", "Non-flippered", "Ambiguous")) + scale_fill_manual(values=c("grey", "skyblue", "salmon1")) + geom_point(x=1, y=PC1_2_mean__mean_F, col = "yellow", size = 2) + geom_point(x=2, y=PC1_2_mean__mean_Nf, col = "yellow", size = 2) + geom_point(x=3, y=PC1_2_mean__mean_Amb, col = "yellow", size = 2) + theme(aspect.ratio = 4))
```

```{r Perform t-tests to compare flippered and non-flippered groups}
# NOTE: The code below performs regular t-tests on the PC scores for all specimens in the dataset. These test results are not technically valid, as they do not correct for phylogenetic autocorrelation among specimens. In order to perform the statistical analyses below more rigorously, one would need to define a time-calibrated tree including all the landmarked specimens in the dataset. You would then need to use the tree to either (i) perform phylogenetic ANOVAs comparing each PC score among groups, or (ii) compute phylogenetic independent contrasts (PICs) for all PC score values in the data set and then compare those PIC values among groups.

#First, create a new data frame that excludes the "Ambiguous" taxa:
dim(PC_data_LimbType_exp)
PC_data_LimbType_exp2 <- PC_data_LimbType_exp[c(1:14, 16:29, 31:38), ]
dim(PC_data_LimbType_exp2)

PC_data_LimbType_exp2$Forelimb.Type <- droplevels(PC_data_LimbType_exp2$Forelimb.Type)
levels(PC_data_LimbType_exp2$Forelimb.Type)

# Now you can perform t-tests comparing normally distributed variables between flippered and non-flippered taxa:
t.test(PC_data_LimbType_exp2$PC1 ~ PC_data_LimbType_exp2$Forelimb.Type)
t.test(PC_data_LimbType_exp2$PC2 ~ PC_data_LimbType_exp2$Forelimb.Type)
t.test(PC_data_LimbType_exp2$PC3 ~ PC_data_LimbType_exp2$Forelimb.Type)
t.test(PC_data_LimbType_exp2$PC_1_2_mean ~ PC_data_LimbType_exp2$Forelimb.Type)
```

This concludes our geometric morphometric analysis of the forelimb zeugoautopod. In the next portion of the script, we repeat the steps above to perform a similar 2D GM analysis of the forelimb acropodial planform.

### ---------------- [GM ANALYSIS 2] ACROPODIAL PLANFORM ------------------ ###

```{r Convert all image files to the correct format.}
# To work in geomorph, all image files must be in JPEG  format. The following code tells you which files in your working directory are in which format, converts PNG and TIFF files to JPEGs.
list.files(base_dir, pattern='*jpg')
list.files(base_dir, pattern='*png')
list.files(base_dir, pattern='*tif')
png_set <- list.files(base_dir, pattern='*png')
tif_set <- list.files(base_dir, pattern = '*tif')

for (png_i in png_set){
  print(png_i)
  jpg_i <- gsub(".png", ".jpg", x = png_i)
  jpg_out_path <- paste0(base_dir, '/', jpg_i)
  png_in_path = paste0(base_dir, '/', png_i)
  img <- readPNG(png_in_path)
  writeJPEG(img, target = jpg_out_path, quality = 1)
}

for (tif_i in tif_set){
  print(tif_i)
  jpg_i <- gsub(".tif", ".jpg", x = tif_i)
  jpg_out_path <- paste0(base_dir, '/', jpg_i)
  tif_in_path = paste0(base_dir, '/', tif_i)
  img <- readTIFF(tif_in_path)
  writeJPEG(img, target = jpg_out_path, quality = 1)
}

# Note: If the tiff code doesn't work (due to "incompatible type for 'RichTIFFIPTC'), just open the tif and export it manually to the working directory as a jpg.

# Now that you've made jpg versions of all your specimen image files, you can remove all former PNG and TIFF files in your working directory:
unlink("/Users/calebgordon/Desktop/220802_GM/*png")
unlink("/Users/calebgordon/Desktop/220802_GM/*tif")

# The code below verifies the absence of PNGs & TIFFs in the working directory:
list.files(base_dir, pattern='*png')
list.files(base_dir, pattern='*tif')
# It should say there are none ("character(0)").
```

```{r Define data objects and grouping variables.}
# Now that all of your image files are in the correct format, there's one more thing you should do before you place landmarks: define data objects that will make it much easier to automate landmark placement and downstream analysis for a large number of specimens.

# First, define a path designating a plain text file to which landmark coordinate data will be read. The following code defines this path and removes this file if it already exists (to avoid the formation of duplicates during multiple runs).
tps_path <- paste0(base_dir, '/raw_coordinates.txt')
#if (file.exists(tps_path)){file.remove(tps_path)}

# Second, define a shorthand object listing all of your input files:
specimens <- list.files(base_dir, pattern='*jpg')

# Thirdly, define grouping variables. In your geometric morphometric analyses, you'll probably want to bin specimens by group. To do this, it's best to define grouping variables that you can input into geomorph commands later.

# First, manually create a file grouping specimens in excel, and save it as a .csv file in the working directory (e.g., "GM_groups.csv"). You may need to open this CSV file in Numbers, manually delete blank rows and columns, and then overwrite the original CSV file. Then, once you have a correctly formatted CSV file, run the following code:
GM_groups <- read.csv("02.08.23_acropod_GM.csv", header=T, row.names=NULL)

# Check out your data frame of grouping variables, to make sure it looks right:
dim(GM_groups); head(GM_groups)

# Set your working directory one more time in preparation for the next step:
setwd(base_dir)
```

```{r Collect landmarks.}
# Now that we have defined useful data objects and grouping variables, we are ready to place landmarks.

# The code below 'uploads' the specimen image files in your working directory to the RStudio GUI, so that you can manually place landmarks on each file in R. In this line of code, you should also set the number of landmarks you plan to place, and the scale, if necessary. Once you run this line, you'll be prompted to place landmarks on each image sequentially. Don't worry if some of these landmarks are sliders. You'll define sliders retroactively in a bit.

# Note: You must copy this text into the console and run it there. Do not run this code from within the Rmd chunk or it won't work.

digitize2d(filelist=specimens, nlandmarks=31, scale=NULL, MultScale=FALSE, tpsfile=tps_path, verbose=FALSE)
```

Troubleshooting with the digitize2d() function above:
The digitize2d() function can be finicky sometimes. If it's not working, try these five things:

1. Do not run the code from within the Rmd chunk above. Instead, copy and paste it into the R console window and run it there.

2. Make sure your input filelist (eg, specimens) is a character vector. You can do this by typing "class(specimens)" in the console. A character vector is expected as input.

3. Set your working directory and make sure your files are correctly named, your filepath is specified correctly, and your files are in the working directory.

4. Make sure that none of your specimen image files begin with a numerical character (1234567890) or contain any spaces. Whenever I've had inexplicable issues with digitize2d() in the past, taking this step has fixed it.

5. If you've done (1) to (4) and it's still not working, it might be a readJPEG() issue. The readJPEG() command is a dependency of digitize2d() and apparently it can run into iOS permissions issues. Run this code to make sure the readJPEG() function isn't the issue:
jpgtest <- readJPEG("/Users/calebgordon/Desktop/Morphometric analyses 9.27.20/GM analysis - 9.27.20/Working_directory/Cryptocleidus_Sauropterygia.jpg", native = TRUE)

Once you place landmarks on all specimens, you can move on to the next step.

```{r Define sliders.}
# Read your raw_coordinates.txt file to an object to make downstream work easier:
raw_coordinates <- readland.tps("/Users/caleb/Desktop/raw_coordinates.txt", specID = "ID") # This object contains the raw pixel coordinates of all of your landmarks for each specimen.

# The following code lets you manually (and retroactively) define sliders for all of your specimens in the RStudio GUI.
define.sliders(landmarks = raw_coordinates, nsliders = 20, surfsliders = NULL, write.file = TRUE)
#—————————Note: Do not run this command from within the chunk. Copy and paste this code into the R console window and run it there.
#—————————This command will have you define sliders for your first specimen, and then write them to the csv file "curveslide.csv". But the sliders are the same for all specimens, so this is ultimately all you need.

# Now write the slider data to an object for downstream use:
sliders <- as.matrix(read.csv("/Users/caleb/Desktop/Flippers_KEY/Flipper GM files PRESSING/CURRENT SPECIMENS/curveslide.csv", header=T)) 
#This file will be one of the direct inputs for Generalized Procrustes Analysis.
```

```{r Perform Generalized Procrustes Superimposition}
# Read your raw_coordinates.txt file to an object to make downstream work easier:
raw_coordinates <- readland.tps("/Users/caleb/Desktop/raw_coordinates.txt", specID = "ID") # This object contains the raw pixel coordinates of all of your landmarks for each specimen. If you defined sliders, you've already done this. But it won't overwrite anything important, so you can run this line again without worry.

# Perform generalized procrustes superimposition and write the output to a new object containing the Procrustes coordinates and metadata:

# WITH SLIDERS:
#GM.gpa <- gpagen(raw_coordinates, curves = sliders, surfaces = NULL, PrinAxes = TRUE, max.iter = NULL, ProcD = FALSE, Proj = TRUE, print.progress = TRUE)

# WITHOUT SLIDERS:
GM.gpa <- gpagen(raw_coordinates, curves = NULL, surfaces = NULL, PrinAxes = TRUE, max.iter = NULL, ProcD = FALSE, Proj = TRUE, print.progress = TRUE)

# GM.gpa$coords contains the Procrustes coordinates of your landmarks and sliding semi-landmarks. Check out the Procrustes coordinates of your specimens:
head(GM.gpa$coords)

# Now write the output of GM.gpa$coords to a tps file that PAST can read:
writeland.tps(GM.gpa$coords, "Procrustes_coordinates_unprocessed.txt", scale = NULL, specID = "ID")
```

```{r Process your Procrustes coordinate data.}
#In the plain text file you just created ("Procrustes_coordinates.txt"), either manually type in specimen names as "ID=specimen_name" (eg, "ID=Adelosaurus_sp.") below each stanza of Procrustes coordinates or run the following lines of code (which do the same thing).This is necessary to put the file in tps format so that PAST can read it.

  # This line reads your unprocessed Procrustes coordinates from the previous step into an object we can modify.
  Procrustes_coords <- readLines(con <- file("Procrustes_coordinates_unprocessed.txt"))

  # This line defines a new vector containing all the names of the files in your dataset.
  specimen_filenames <- list.files("/Users/caleb/Desktop/02.07.23_Acropod_GM/specimen_files")
  length(specimen_filenames)

  
  # This line replaces the blank lines with "ID=File_name".
  for(i in 1:length(specimen_filenames)){Procrustes_coords[c(i*17)] <- paste("ID=", specimen_filenames[i], sep="")}
  Procrustes_coords # This line lets you check that the replacement worked.
  write.table(Procrustes_coords, file = "Procrustes_coordinates_FIN.txt", sep = "\n", row.names = FALSE, col.names = FALSE) # This line writes the resulting modified data object (Procrustes_coords) to a new txt file.

# Now, open the new file you've just created ("Procrustes_coordinates_FIN.txt"), manually remove the quotation marks, and manually add a line break between every stanza. After this, you will be ready to perform the rest of the analysis.
```

Now, at this point, PAST and RStudio diverge. I prefer the PCA figure output of PAST to that of geomorph, so I would recommend doing PCA and PERMANOVAs in PAST. However, geomorph also has some useful tools that PAST doesn't (for example, warp grid comparisons). As a result, below are instructions for all of the downstream analyses I've found useful in PAST *and* RStudio.

```{r Perform PCA and run PERMANOVAs in PAST.}
# Open "Procrustes_coordinates_FIN.txt" in PAST4
# Manually enter in your grouping variables, using the Groups csv file you defined earlier.

# Run a PCA. Follow in-app instructions to make and modify convex hulls. Save elbow-plots and all other metadata for reference. Also save spline plots of low and high extreme scores for each relevant PC.
```

```{r Compare individual PC scores among groups in R.}
# The following code looks at PC score distributions for key PC scores from the PCA you ran above, and compares them between limb types and clades. It's basically the same as the first few chunks of code in my Ternary script ("Ternary script.Rmd").

# Set working directory:
setwd("/Users/calebgordon/Desktop/GM analysis - 9.29.20")
# Read data into an object:
PC_data <- read.csv("GM_forelimb_data.csv")
# Check out the data:
dim(PC_data); head(PC_data)

# Define new data frames and means for use in boxplots:
  #For Clade comparisons:
  PC_data_Clade <- PC_data[, c(3,2,6,7,8,9)]
  dim(PC_data_Clade); head(PC_data_Clade)
  #For Limb Type comparisons:
  PC_data_LimbType <- PC_data[, c(3,4,6,7,8,9)]
  dim(PC_data_LimbType); head(PC_data_LimbType)
  #Defining a mean PC1/2 score metric:
  PC_1_2_mean <- rep(NA, 38)
  for (i in 1:38){PC_1_2_mean[i]<-mean(c(PC_data_LimbType$PC1[i], PC_data_LimbType$PC2[i]))}
  PC_data_LimbType_exp <- cbind(PC_data_LimbType[, c(1:6)], PC_1_2_mean)
  #Defining means for quantitative variables:
  PC1_mean_F <- mean(PC_data_LimbType$PC1[PC_data_LimbType$Forelimb.Type == "Flippered"])
  PC1_mean_Nf <- mean(PC_data_LimbType$PC1[PC_data_LimbType$Forelimb.Type == "Non-flippered"])
  PC1_mean_Amb <- mean(PC_data_LimbType$PC1[PC_data_LimbType$Forelimb.Type == "Ambiguous"])
  PC1_groupmeans <- c(PC1_mean_F, PC1_mean_Nf, PC1_mean_Amb)
  PC2_mean_F <- mean(PC_data_LimbType$PC2[PC_data_LimbType$Forelimb.Type == "Flippered"])
  PC2_mean_Nf <- mean(PC_data_LimbType$PC2[PC_data_LimbType$Forelimb.Type == "Non-flippered"])
  PC2_mean_Amb <- mean(PC_data_LimbType$PC2[PC_data_LimbType$Forelimb.Type == "Ambiguous"])
  PC1_2_mean__mean_F <- mean(PC_data_LimbType_exp$PC_1_2_mean[PC_data_LimbType_exp$Forelimb.Type == "Flippered"])
  PC1_2_mean__mean_Nf <- mean(PC_data_LimbType_exp$PC_1_2_mean[PC_data_LimbType_exp$Forelimb.Type == "Non-flippered"])
  PC1_2_mean__mean_Amb <- mean(PC_data_LimbType_exp$PC_1_2_mean[PC_data_LimbType_exp$Forelimb.Type == "Ambiguous"])

# Run distrib.plot function on untransformed variables:
distrib.plot(PC_data_LimbType_exp[, 3:7], title = "Distributions of PC scores for different reptile forelimbs")

# Run distrib.plot function on transformed variables:
distrib.plot(log(abs(PC_data_LimbType_exp[, 3:7])), title = "Distributions of log-transformed PC scores for different reptile forelimbs")

# Make box plots:
  #Comparing PC1 scores:
(bp_PC1_LimbType_untrans <- ggplot(PC_data_LimbType, aes(x=Forelimb.Type, y=PC1, fill=Forelimb.Type)) + geom_boxplot(width=0.8) + theme(legend.position="none") + scale_x_discrete(limits=c("Flippered", "Non-flippered", "Ambiguous"), name = "Limb Type", labels = c("Flippered", "Non-flippered", "Ambiguous")) + scale_fill_manual(values=c("grey", "skyblue", "salmon1")) + geom_point(x=1, y=PC1_mean_F, col = "yellow", size = 2) + geom_point(x=2, y=PC1_mean_Nf, col = "yellow", size = 2) + geom_point(x=3, y=PC1_mean_Amb, col = "yellow", size = 2) + theme(aspect.ratio = 4))

  #Comparing absolute-value transformed PC1 scores:
(bp_PC1_LimbType_trans <- ggplot(PC_data_LimbType, aes(x=Forelimb.Type, y=abs(PC1), fill=Forelimb.Type)) + geom_boxplot(width=0.8) + theme(legend.position="none") + scale_x_discrete(limits=c("Flippered", "Non-flippered", "Ambiguous"), name = "Limb Type", labels = c("Flippered", "Non-flippered", "Ambiguous")) + scale_fill_manual(values=c("grey", "skyblue", "salmon1")) + geom_point(x=1, y=abs(PC1_mean_F), col = "yellow", size = 2) + geom_point(x=2, y=abs(PC1_mean_Nf), col = "yellow", size = 2) + geom_point(x=3, y=abs(PC1_mean_Amb), col = "yellow", size = 2) + theme(aspect.ratio = 4))

  #Comparing PC2 scores:
(bp_PC2_LimbType_untrans <- ggplot(PC_data_LimbType, aes(x=Forelimb.Type, y=PC2, fill=Forelimb.Type)) + geom_boxplot(width=0.8) + theme(legend.position="none") + scale_x_discrete(limits=c("Flippered", "Non-flippered", "Ambiguous"), name = "Limb Type", labels = c("Flippered", "Non-flippered", "Ambiguous")) + scale_fill_manual(values=c("grey", "skyblue", "salmon1")) + geom_point(x=1, y=PC2_mean_F, col = "yellow", size = 2) + geom_point(x=2, y=PC2_mean_Nf, col = "yellow", size = 2) + geom_point(x=3, y=PC2_mean_Amb, col = "yellow", size = 2) + theme(aspect.ratio = 4))

  #Comparing PC3 scores:
(bp_PC3_LimbType_untrans <- ggplot(PC_data_LimbType, aes(x=Forelimb.Type, y=PC3, fill=Forelimb.Type)) + geom_boxplot(width=0.8) + theme(legend.position="none") + scale_x_discrete(limits=c("Flippered", "Non-flippered", "Ambiguous"), name = "Limb Type", labels = c("Flippered", "Non-flippered", "Ambiguous")) + scale_fill_manual(values=c("grey", "skyblue", "salmon1")) + theme(aspect.ratio = 4))

  #Comparing mean PC1&2 scores (a PC1/2 score metric):
(bp_PC3_LimbType_untrans <- ggplot(PC_data_LimbType_exp, aes(x=Forelimb.Type, y=PC_1_2_mean, fill=Forelimb.Type)) + geom_boxplot(width=0.8) + theme(legend.position="none") + scale_x_discrete(limits=c("Flippered", "Non-flippered", "Ambiguous"), name = "Limb Type", labels = c("Flippered", "Non-flippered", "Ambiguous")) + scale_fill_manual(values=c("grey", "skyblue", "salmon1")) + geom_point(x=1, y=PC1_2_mean__mean_F, col = "yellow", size = 2) + geom_point(x=2, y=PC1_2_mean__mean_Nf, col = "yellow", size = 2) + geom_point(x=3, y=PC1_2_mean__mean_Amb, col = "yellow", size = 2) + theme(aspect.ratio = 4))
```

```{r Perform t-tests to compare flippered and non-flippered groups}
# NOTE: The code below performs regular t-tests on the PC scores for all specimens in the dataset. These test results are not technically valid, as they do not correct for phylogenetic autocorrelation among specimens. In order to perform the statistical analyses below more rigorously, one would need to define a time-calibrated tree including all the landmarked specimens in the dataset. You would then need to use the tree to either (i) perform phylogenetic ANOVAs comparing each PC score among groups, or (ii) compute phylogenetic independent contrasts (PICs) for all PC score values in the data set and then compare those PIC values among groups.

#First, create a new data frame that excludes the "Ambiguous" taxa:
dim(PC_data_LimbType_exp)
PC_data_LimbType_exp2 <- PC_data_LimbType_exp[c(1:14, 16:29, 31:38), ]
dim(PC_data_LimbType_exp2)
PC_data_LimbType_exp2$Forelimb.Type <- droplevels(PC_data_LimbType_exp2$Forelimb.Type)
levels(PC_data_LimbType_exp2$Forelimb.Type)

# Now you can perform t-tests comparing normally distributed variables between flippered and non-flippered taxa:
t.test(PC_data_LimbType_exp2$PC1 ~ PC_data_LimbType_exp2$Forelimb.Type)
t.test(PC_data_LimbType_exp2$PC2 ~ PC_data_LimbType_exp2$Forelimb.Type)
t.test(PC_data_LimbType_exp2$PC3 ~ PC_data_LimbType_exp2$Forelimb.Type)
t.test(PC_data_LimbType_exp2$PC_1_2_mean ~ PC_data_LimbType_exp2$Forelimb.Type)
```

### ---------------- [GM ANALYSIS 3] UNGUAL PHALANX SHAPE ----------------- ###

```{r Convert all image files so that they are in the correct format.}
# To work in geomorph, all image files must be in JPEG  format. The following code tells you which files in your working directory are in which format, converts PNG and TIFF files to JPEGs.

list.files(base_dir, pattern='*jpg')
list.files(base_dir, pattern='*png')
list.files(base_dir, pattern='*tif')
png_set <- list.files(base_dir, pattern='*png')
tif_set <- list.files(base_dir, pattern = '*tif')

for (png_i in png_set){
  print(png_i)
  jpg_i <- gsub(".png", ".jpg", x = png_i)
  jpg_out_path <- paste0(base_dir, '/', jpg_i)
  png_in_path = paste0(base_dir, '/', png_i)
  img <- readPNG(png_in_path)
  writeJPEG(img, target = jpg_out_path, quality = 1)
}

for (tif_i in tif_set){
  print(tif_i)
  jpg_i <- gsub(".tif", ".jpg", x = tif_i)
  jpg_out_path <- paste0(base_dir, '/', jpg_i)
  tif_in_path = paste0(base_dir, '/', tif_i)
  img <- readTIFF(tif_in_path)
  writeJPEG(img, target = jpg_out_path, quality = 1)
}

# Note: If the tiff code doesn't work (due to "incompatible type for 'RichTIFFIPTC'), just open the tif and export it manually to the working directory as a jpg.

# Now that you've made jpg versions of all your specimen image files, you can remove all former PNG and TIFF files in your working directory:
unlink("/Users/calebgordon/Desktop/220802_GM/*png")
unlink("/Users/calebgordon/Desktop/220802_GM/*tif")

# The code below verifies the absence of PNGs and TIFFs in your working directory:
list.files(base_dir, pattern='*png')
list.files(base_dir, pattern='*tif')
# It should say there are none ("character(0)").
```

```{r Define data objects and grouping variables.}
# Now that all of your image files are in the correct format, there's one more thing you should do before you place landmarks: define data objects that will make it much easier to automate landmark placement and downstream analysis for a large number of specimens.

# First, define a path designating a plain text file to which landmark coordinate data will be read. The following code defines this path and removes this file if it already exists (to avoid the formation of duplicates during multiple runs).
tps_path <- paste0(base_dir, '/raw_coordinates.txt')
#if (file.exists(tps_path)){file.remove(tps_path)}

# Second, define a shorthand object listing all of your input files:
specimens <- list.files(base_dir, pattern='*jpg')

# Thirdly, define grouping variables. In your geometric morphometric analyses, you'll probably want to bin specimens by group. To do this, it's best to define grouping variables that you can input into geomorph commands later.

# First, manually create a file grouping specimens in excel, and save it as a .csv file in the working directory (e.g., "GM_groups.csv"). You may need to open this CSV file in Numbers, manually delete blank rows and columns, and then overwrite the original CSV file. Then, once you have a correctly formatted CSV file, run the following code:
GM_groups <- read.csv("GM_groups.csv", header=T, row.names=NULL)

# Check out your data frame of grouping variables, to make sure it looks right:
dim(GM_groups)
head(GM_groups)

# Set your working directory one more time in preparation for the next step:
setwd(base_dir)
```

```{r Collect landmarks.}
# Now that we have defined useful data objects and grouping variables, we are ready to place landmarks.

# The code below 'uploads' the specimen image files in your working directory to the RStudio GUI, so that you can manually place landmarks on each file in R. In this line of code, you should also set the number of landmarks you plan to place, and the scale, if necessary. Once you run this line, you'll be prompted to place landmarks on each image sequentially. Don't worry if some of these landmarks are sliders. You'll define sliders retroactively in a bit.

# Note: You must copy this text into the console and run it there. Do not run this code from within the Rmd chunk or it won't work.

digitize2d(filelist=specimens, nlandmarks=31, scale=NULL, MultScale=FALSE, tpsfile=tps_path, verbose=FALSE)
```

Troubleshooting with the digitize2d() function above:
The digitize2d() function can be finicky sometimes. If it's not working, try these five things:
---1. Do not run the code from within the Rmd chunk above. Instead, copy and paste it into the R console window and run it there.

---2. Make sure your input filelist (eg, specimens) is a character vector. You can do this by typing "class(specimens)" in the console. A character vector is expected as input.

---3. Set your working directory and make sure your files are correctly named, your filepath is specified correctly, and your files are in the working directory.

---4. Make sure that none of your specimen image files begin with a numerical character (1234567890) or contain any spaces. Whenever I've had inexplicable issues with digitize2d() in the past, taking this step has fixed it.

---5. If you've done (1) to (4) and it's still not working, it might be a readJPEG() issue. The readJPEG() command is a dependency of digitize2d() and apparently it can run into iOS permissions issues. Run this code to make sure the readJPEG() function isn't the issue:
jpgtest <- readJPEG("/Users/calebgordon/Desktop/Morphometric analyses 9.27.20/GM analysis - 9.27.20/Working_directory/Cryptocleidus_Sauropterygia.jpg", native = TRUE)

Once you place landmarks on all specimens, you can move on to the next step.

```{r Define sliders.}
# Read raw_coordinates.txt file to an object to make downstream work easier:
raw_coordinates <- readland.tps("/Users/caleb/Desktop/raw_coordinates.txt", specID = "ID") # This object contains the raw pixel coordinates of all of your landmarks for each specimen.

# The following code lets you manually (and retroactively) define sliders for all of your specimens in the RStudio GUI.
define.sliders(landmarks = raw_coordinates, nsliders = 14, surfsliders = NULL, write.file = TRUE)
#—————————Note: Do not run this command from within the chunk. Copy and paste this code into the R console window and run it there.
#—————————This command will have you define sliders for your first specimen, and then write them to the csv file "curveslide.csv". But the sliders are the same for all specimens, so this is ultimately all you need.

# Now write the slider data to an object for downstream use:
sliders <- as.matrix(read.csv("/Users/caleb/Desktop/curveslide.csv", header=T)) 
#This file will be one of the direct inputs for Generalized Procrustes Analysis.
```

```{r Perform Generalized Procrustes Superimposition}
# Perform Generalized Procrustes Superimposition and write the output to a new object containing the Procrustes coordinates and metadata:
GM.gpa <- gpagen(raw_coordinates, curves = sliders, surfaces = NULL, PrinAxes = TRUE, max.iter = NULL, ProcD = FALSE, Proj = TRUE, print.progress = TRUE)

# GM.gpa$coords contains the Procrustes coordinates of your landmarks and sliding semi-landmarks. Check out the Procrustes coordinates of your specimens:
head(GM.gpa$coords)

# Now write the output of GM.gpa$coords to a tps file that PAST can read:
writeland.tps(GM.gpa$coords, "Procrustes_coordinates.txt", scale = NULL, specID = "ID")
```

Now, at this point, PAST and RStudio diverge. I prefer the PCA figure output of PAST to that of geomorph, so I would recommend doing PCA and PERMANOVAs in PAST. However, geomorph also has some useful tools that PAST doesn't (for example, warp grid comparisons). As a result, below are instructions for all of the downstream analyses I've found useful in PAST *and* RStudio.

```{r Perform PCA and phyloPERMANOVAs in PAST.}
#In the plain text file you just created ("Procrustes_coordinates.txt"), manually type in specimen names as "ID=specimen_name" (eg, "ID=Adelosaurus_sp.") below each stanza of Procrustes coordinates.This is necessary to put the file in tps format so that PAST can read it.

# Then open this file in PAST4
# Manually define grouping variables, perhaps using the Groups csv you defined earlier.

# Run a PCA. Follow in-app instructions to make and modify convex hulls. Save elbow-plots and all other metadata for reference. Also save spline plots of low and high extreme scores for each relevant PC.
```

```{r Compare individual PC scores among groups in R.}
# The following code looks at PC score distributions for key PC scores from the PCA you ran in (8), and compares them between limb types and clades. It's basically the same as the first few chunks of code in my Ternary script ("Ternary script.Rmd").

# Read data into an object:
PC_data <- read.csv("Ungual_2D_GM_10.25.22.csv")

# Check out the data:
dim(PC_data)
head(PC_data)
names(PC_data)
PC_data <- PC_data[ , c(1:11)]

# Defining group means for quantitative variables:
PC1_means <- tapply(PC_data$PC.1, INDEX=PC_data$LT2, FUN=mean)
PC2_means <- tapply(PC_data$PC.2, INDEX=PC_data$LT2, FUN=mean)
PC3_means <- tapply(PC_data$PC.3, INDEX=PC_data$LT2, FUN=mean)
PC4_means <- tapply(PC_data$PC.4, INDEX=PC_data$LT2, FUN=mean)

# Run distrib.plot function on untransformed variables:
distrib.plot(PC_data[, c(8:11)], title = "Distributions of PC scores for amniote unguals") # VERDICT: ALL NORMAL!

# Make box plots:
  #Comparing PC1 scores:
(bp_PC1 <- ggplot(PC_data, aes(x=LT2, y=PC.1, fill=LT2)) + geom_boxplot(lwd=1.5) + theme(legend.position="none") + scale_x_discrete(limits=c("Flippered", "Unwebbed", "Webbed"), name = "Limb Type", labels = c("Flippered", "Unwebbed", "Webbed")) + scale_fill_manual(values=c("skyblue", "salmon1", "gold")) +       theme(plot.margin=margin(t=20, r=100, b=40, l=100), axis.text=element_text(size=12), axis.title=element_text(size=14)) + xlab(label=NULL) + ylab(label="PC1 score") + stat_summary(fun=mean, geom="point", shape=23, size=5, stroke=1.5, color="black",fill='azure') + scale_x_discrete(labels = c(paste("Flippered \n n=",length(PC_data$Taxon[PC_data$LT2 == "Flippered"]), sep=""), paste("Unwebbed \n n=",length(PC_data$Taxon[PC_data$LT2 == "Unwebbed"]), sep=""), paste("Webbed \n n=", length(PC_data$Taxon[PC_data$LT2 == "Webbed"]), sep=""))) + ggtitle(paste("Raw PC1 score", " by limb type", sep="")))
    
  #Comparing PC2 scores:
(bp_PC2 <- ggplot(PC_data, aes(x=LT2, y=PC.2, fill=LT2)) + geom_boxplot(lwd=1.5) + theme(legend.position="none") + scale_x_discrete(limits=c("Flippered", "Unwebbed", "Webbed"), name = "Limb Type", labels = c("Flippered", "Unwebbed", "Webbed")) + scale_fill_manual(values=c("skyblue", "salmon1", "gold")) +       theme(plot.margin=margin(t=20, r=100, b=40, l=100), axis.text=element_text(size=12), axis.title=element_text(size=14)) + xlab(label=NULL) + ylab(label="PC2 score") + stat_summary(fun=mean, geom="point", shape=23, size=5, stroke=1.5, color="black",fill='azure') + scale_x_discrete(labels = c(paste("Flippered \n n=",length(PC_data$Taxon[PC_data$LT2 == "Flippered"]), sep=""), paste("Unwebbed \n n=",length(PC_data$Taxon[PC_data$LT2 == "Unwebbed"]), sep=""), paste("Webbed \n n=", length(PC_data$Taxon[PC_data$LT2 == "Webbed"]), sep=""))) + ggtitle(paste("Raw PC2 score", " by limb type", sep="")))    

  #Comparing PC3 scores:
(bp_PC3 <- ggplot(PC_data, aes(x=LT2, y=PC.3, fill=LT2)) + geom_boxplot(lwd=1.5) + theme(legend.position="none") + scale_x_discrete(limits=c("Flippered", "Unwebbed", "Webbed"), name = "Limb Type", labels = c("Flippered", "Unwebbed", "Webbed")) + scale_fill_manual(values=c("skyblue", "salmon1", "gold")) +       theme(plot.margin=margin(t=20, r=100, b=40, l=100), axis.text=element_text(size=12), axis.title=element_text(size=14)) + xlab(label=NULL) + ylab(label="PC3 score") + stat_summary(fun=mean, geom="point", shape=23, size=5, stroke=1.5, color="black",fill='azure') + scale_x_discrete(labels = c(paste("Flippered \n n=",length(PC_data$Taxon[PC_data$LT2 == "Flippered"]), sep=""), paste("Unwebbed \n n=",length(PC_data$Taxon[PC_data$LT2 == "Unwebbed"]), sep=""), paste("Webbed \n n=", length(PC_data$Taxon[PC_data$LT2 == "Webbed"]), sep=""))) + ggtitle(paste("Raw PC3 score", " by limb type", sep="")))   

  #Comparing PC4 scores:
(bp_PC4 <- ggplot(PC_data, aes(x=LT2, y=PC.4, fill=LT2)) + geom_boxplot(lwd=1.5) + theme(legend.position="none") + scale_x_discrete(limits=c("Flippered", "Unwebbed", "Webbed"), name = "Limb Type", labels = c("Flippered", "Unwebbed", "Webbed")) + scale_fill_manual(values=c("skyblue", "salmon1", "gold")) +       theme(plot.margin=margin(t=20, r=100, b=40, l=100), axis.text=element_text(size=12), axis.title=element_text(size=14)) + xlab(label=NULL) + ylab(label="PC4 score") + stat_summary(fun=mean, geom="point", shape=23, size=5, stroke=1.5, color="black",fill='azure') + scale_x_discrete(labels = c(paste("Flippered \n n=",length(PC_data$Taxon[PC_data$LT2 == "Flippered"]), sep=""), paste("Unwebbed \n n=",length(PC_data$Taxon[PC_data$LT2 == "Unwebbed"]), sep=""), paste("Webbed \n n=", length(PC_data$Taxon[PC_data$LT2 == "Webbed"]), sep=""))) + ggtitle(paste("Raw PC4 score", " by limb type", sep=""))) 
```

```{r Perform PIC on GM data.}
# Read in the time-calibrated phylogeny generated in TimeTree:
Ungual_2D_GM_tree <-read.newick("Ungual_tree.nwk")

# Check specimen order:
Ungual_2D_GM_tree$tip.label
PC_data$Taxon

# Import CSV:
PC_data_forPIC <- read.csv("Ungual_2D_GM_10.25.22_forPIC.csv")

# Create numeric vectors for the traits you want to perform corrections on:
PC1 <- PC_data_forPIC$PC.1
PC2 <- PC_data_forPIC$PC.2
PC3 <- PC_data_forPIC$PC.3
PC4 <- PC_data_forPIC$PC.4

# Compute PICs:
PC1_contrasts <- pic(PC1, Ungual_2D_GM_tree)
PC2_contrasts <- pic(PC2, Ungual_2D_GM_tree)
PC3_contrasts <- pic(PC3, Ungual_2D_GM_tree)
PC4_contrasts <- pic(PC4, Ungual_2D_GM_tree)
all_PC_contrasts <- cbind(PC1_contrasts, PC2_contrasts, PC3_contrasts, PC4_contrasts)

# Compute contrasts and associated variances
PC1_contrasts.var <- pic(PC1, Ungual_2D_GM_tree, var.contrasts = TRUE)
PC2_contrasts.var <- pic(PC2, Ungual_2D_GM_tree, var.contrasts = TRUE)
PC3_contrasts.var <- pic(PC3, Ungual_2D_GM_tree, var.contrasts = TRUE)
PC4_contrasts.var <- pic(PC4, Ungual_2D_GM_tree, var.contrasts = TRUE)

# Plot PICs onto specimen tree:
plot (Ungual_2D_GM_tree)
  nodelabels(round(PC1_contrasts.var [,1], 3), adj = c(0, -0.5), frame="n")
  #nodelabels(round(PC2_contrasts.var [,1], 3), adj = c(0, 1), frame="rect")
```

```{r Perform phylogenetic ANOVA with post-hoc Tukey tests.}
# Confirming order is correct:
PC_data_forPIC$Taxon
Ungual_2D_GM_tree$tip.label

# phylANOVA for PC1:
phylANOVA(Ungual_2D_GM_tree, x=PC_data_forPIC$LT2, y=PC_data_forPIC$PC.1, nsim=1000, posthoc=TRUE, p.adj="bonferroni")

# phylANOVA for PC2:
phylANOVA(Ungual_2D_GM_tree, x=PC_data_forPIC$LT2, y=PC_data_forPIC$PC.2, nsim=1000, posthoc=TRUE, p.adj="bonferroni")

# phylANOVA for PC3:
phylANOVA(Ungual_2D_GM_tree, x=PC_data_forPIC$LT2, y=PC_data_forPIC$PC.3, nsim=1000, posthoc=TRUE, p.adj="bonferroni")

# phylANOVA for PC4:
phylANOVA(Ungual_2D_GM_tree, x=PC_data_forPIC$LT2, y=PC_data_forPIC$PC.4, nsim=1000, posthoc=TRUE, p.adj="bonferroni")
```

This concludes all geometric morphometric analyses for the above paper. If you have any questions about these analyses, please feel free to contact me at.

Sincerely,
Caleb Gordon
````
